# SFRP1 upregulation causes hippocampal synaptic dysfunction and memory impairment

**DOI:** 10.1101/2024.04.04.588100

**Authors:** Guadalupe Pereyra, María Inés Mateo, María Jesús Martin-Bermejo, Pablo Miaja, Remco Klaassen, Agnès Gruart, Javier Rueda-Carrasco, Alba Fernández-Rodrigo, Esperanza López-Merino, Pilar Esteve, José A. Esteban, August B. Smit, José M. Delgado-García, Paola Bovolenta

**Affiliations:** Centro de Biología Molecular Severo Ochoa, CSIC-UAM, Campus de la Universidad Autónoma de Madrid, Madrid, Spain; CIBER de Enfermedades Raras (CIBERER) Madrid, Spain; Center for Neurogenomics and Cognitive Research, VU University Amsterdam, Amsterdam, The Netherlands; División de Neurociencias, Universidad Pablo de Olavide, Seville, Spain

**Keywords:** Alzheimer’s disease, dendritic spines, long-term potentiation, neurodegeneration, proteomics

## Abstract

Decreased dendritic complexity and impaired synaptic function are strongly linked to cognitive decline in Alzheimer’s disease (AD), and precede the emergence of other neuropathological traits that establish a harmful cycle exacerbating synaptic dysfunction. SFRP1, a glial-derived protein regulating cell-cell communication, is abnormally elevated in the brain of AD patients and related mouse models already at early disease stages. Neutralization of SFRP1 activity in mice reduces the occurrence of protein aggregates, neuroinflammation and prevents the loss of synaptic long-term potentiation (LTP). In this study, we generated transgenic mice that overexpress *Sfrp1* in astrocytes to investigate whether LTP loss is due to an early influence of SFRP1 on synaptic function or results from other alterations driving disease progression. We report that SFRP1-overexpressing mice show reduced dendritic complexity and spine density in dentate gyrus granule cells during early adulthood, prior to a significant deficit in LTP response and late onset cognitive impairment. Ultrastructural analysis revealed the loss of small-sized synapses and presynaptic alterations in transgenic mice. Analysis of proteomic changes points to a general decrease in protein synthesis and modifications in the synaptic proteome, particularly of proteins related to synaptic vesicle cycle and synaptic organizers, like neurexin and neuroligin. We propose a model wherein SFRP1 directly impacts on synaptic function, by increasing the availability of synaptic organizing molecules at the synapse. These observations, combined with documented SFRP1 effects on APP processing and microglial activation, imply that SFRP1 contributes to multiple pathological effects in AD, emerging as a promising therapeutic target for this devastating disease.

## Introduction

Synaptic plasticity is a fundamental cellular mechanism that allows neurons to regulate their functional connectivity in response to different stimuli, thereby modulating neuronal and circuitry function. This plasticity may involve structural changes of the dendritic arbor and remodeling or pruning of the associated spines in postsynaptic neurons^1–3^, as well as structural and/or functional modifications of the presynaptic component. The latter includes, for example, changes in the number or mobilization of presynaptic vesicles and alterations in the expression and/or function of different transsynaptic organizing molecules, such as Neurexins, Eph or Cadherins^4–6^. Synaptic plasticity is under continuous homeostatic remodeling, which, when dysregulated, leads to synaptic dysfunction. This dysfunction is a prevalent feature of neurodegenerative diseases, particularly of those involving cognitive decline, such as Alzheimer’s disease (AD). Indeed, reduced complexity of the neuronal dendritic arbor and synaptic dysfunction strongly correlate with cognitive decline in AD, and are among the earliest changes observed in the brains of both AD patients^7–10^ and related animal models^11–13^. These modifications are thereafter fostered by additional neuropathological events such as the characteristic accumulation of unfolded proteins and aggregated peptides (e.g. Aβ, hyper-phosphorylated tau, α-synuclein) that, together with chronic neuroinflammation, establish a feed-forward mechanism that drives disease progression^11,14–16^. However, the onset and temporal sequence of neuronal alterations in AD patients and mouse models is still poorly understood.

Recent studies have demonstrated that the brain levels of the glial derived SFRP1^17,18^ significantly increase with aging^19^, becoming a cause of non-pathological cognitive decline and a primary risk factor for AD and related dementias^20^. Notably, SFRP1 is significantly enriched in brain samples from different cohorts of AD patients, as compared to cognitively normal, age-matched individuals from preclinical stages of the disease^17,21–24^. A similar increase is also present in different AD mouse models, such as APP^695swe^;PS1^dE9^ and 5xFAD, well before the appearance of amyloid plaques and neuroinflammatory signs^17,22,25^. Increased SFRP1 levels positively correlate with AD severity^17,22^ and with the concentration of soluble Aβ peptides^17^. Furthermore, SFRP1 promotes the formation and aggregation of Aβ peptides^17^ and sustains microglial activation via the upregulation of the HIF-dependent inflammatory pathway, thereby acting as an astrocyte-to-microglia amplifier of neuroinflammation^18^. Consistent with a key role of SFRP1 in AD, antibody-mediated neutralization of SFRP1 in APP^695swe^;PS1^dE9^ mice is sufficient to significantly reduce inflammation, amyloid plaque burden and to prevent the decrease in the LTP response normally observed in these mice^17^.

The latter observation has two possible and not mutually exclusive interpretations. In the absence of SFRP1, synaptic function in APP^695swe^;PS1^dE9^ mice is preserved as a result of reduced neuroinflammation and decreased generation of toxic Aβ peptides, which have been shown to alter synaptic activity^14,15^. Alternatively, there is a direct association between synaptic function and SFRP1 activity, which, when abnormally elevated, could contribute to the initial synaptic alterations observed in AD. The latter possibility is supported by the notion that SFRP1 is implicated in the regulation of Wnt signaling and ADAM10 shedding activity, which are both established molecular drivers of synaptic function. Wnt signaling is involved in synapse formation and plasticity and its dysregulation associates with aging and AD^26,27^. ADAM10 localizes pre- and post-synaptically in rodent and human neurons and its conditional genetic inactivation in mice impairs synaptic plasticity^28–30^ in concomitance with an abnormal processing of a number of structural synaptic proteins, including APP, N-cadherin, neurexin and Eph receptors^29,31,32^.

To dissect SFRP1 functions and disease related mechanisms, we generated a transgenic mouse model that overexpresses *Sfrp1* in astrocytes, thereby mimicking what happens in APP^695swe^;PS1^dE9^ mice but in the absence of the formation of Aβ aggregates. Using morphological, ultrastructural, electrophysiological, behavioral and proteomic approaches, we determined that high SFRP1 levels directly impact on dendritic complexity and spine density of dentate gyrus (DG) granule cells (GCs). Elevated SFRP1 levels lead to the upregulation of proteins involved in establishing synaptic structure and vesicle cycle, impair hippocampal activity and lead to late onset cognitive decline. We thus propose that SFRP1 plays an important role in early changes occurring at the synapse in AD.

## Results

### Generation of a transgenic mouse line with elevated Sfrp1 levels in the brain

To recreate an environment in which adult neurons are exposed to increased levels of astrocyte-derived SFRP1 without the presence of amyloid plaque or toxic Aβ peptides, we crossed two available mouse lines. The first line carries a *LacZ*;TRE;*Sfrp1* construct, containing the bovine *Sfrp1* ortholog^33^. *Sfrp1* is highly expressed in the embryonic forebrain^34,35^ (**Fig S1A**) and its loss or gain of function interfere with patterning and neurogenesis^34–36^. The promoter of the *GFAP* gene, encoding an astrocyte specific intermediate filament, is poorly active during embryogenesis^37^. We thus chose the *hGFAP*;tTA,tetO line^38^ as a transgene driver, to avoid any possible interference with embryonic and postnatal brain development. The resulting *hGFAP*;tTA/*LacZ*;TRE;*Sfrp1* mice (*Sfrp1^TG^*, hereafter; **Fig. 1A**) were bred in the absence or presence of doxycycline in the drinking water up to one month of age to determine possible developmental alterations induced by transgene expression. SFRP1 levels in brain homogenates from untreated *Sfrp1^TG^* mice were significantly increased already at late embryonic stages (**Fig. S1A**) as compared to *LacZ*;TRE;*Sfrp1* animals that were routinely used as controls. Nevertheless, no differences were observed in the overall pup growth. Furthermore, no histological abnormalities were detected in the different brain structures of one-month old mice with or without doxycycline treatment, upon immunohistochemical staining of brain sections with cell-specific markers. We thus performed all subsequent experiments in untreated mice.

**Figure 1.**
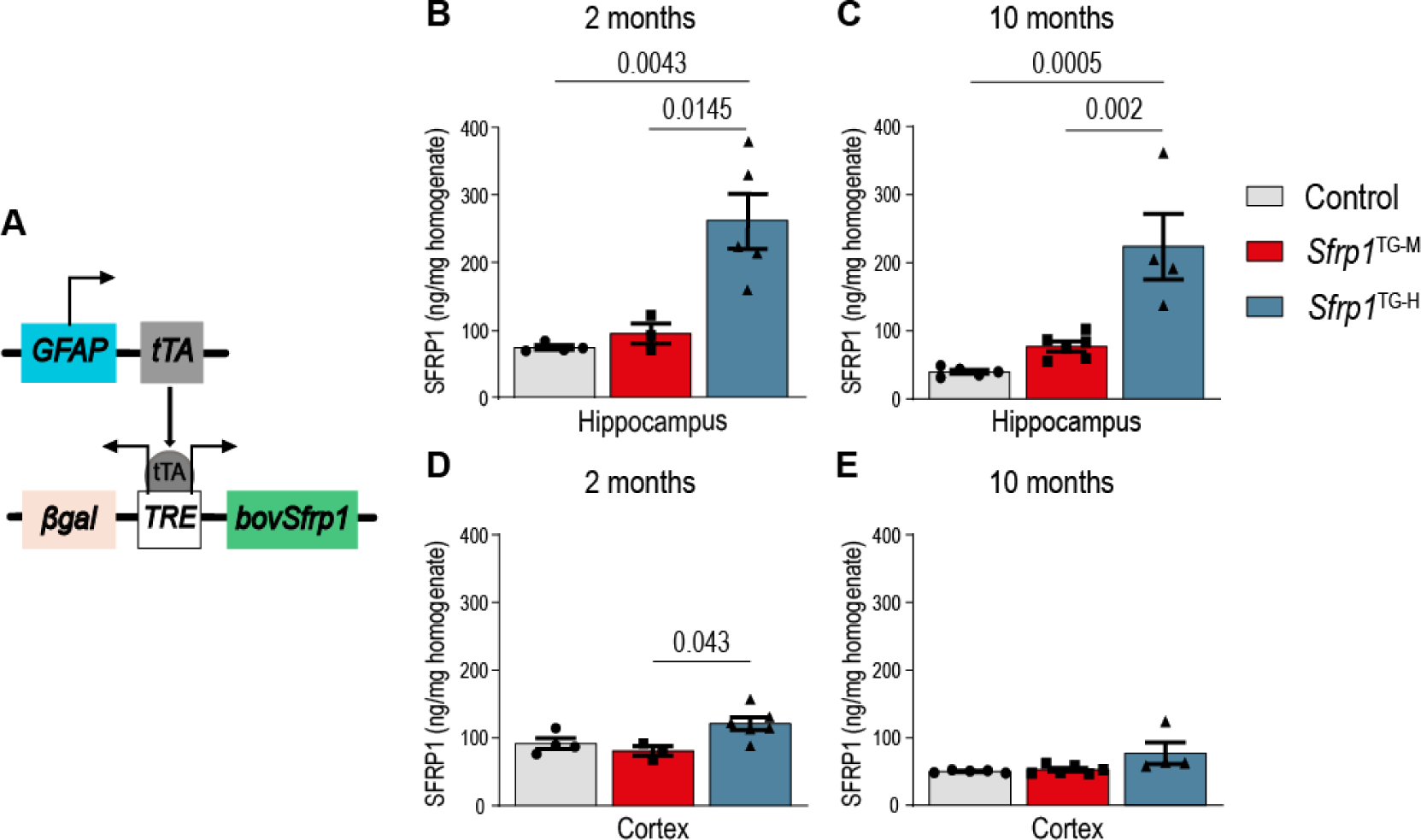
*Sfrp1^TG^* mice express high SFRP1 levels in the hippocampus. **A)** Diagram of the strategy used to generate *Sfrp1^TG^* mice. **B-E**) ELISA determination of SFRP1 protein levels in hippocampal (B, C) or cortical (D, E) homogenates from 2 and 10 months-old control and transgenic mice. n = 4 control, 3 *Sfrp1^TG-M^*, 5 *Sfrp1^TG-H^* mice in (B); 5 control, 6 *Sfrp1^TG-M^* and 4 *Sfrp1^TG-H^* mice in (C, E); 4 control, 3 *Sfrp1^TG-M^*, 6 *Sfrp1^TG-H^* mice in (D). Graph bars represent mean ± SEM. Data were analyzed with one-way ANOVA followed by Bonferroni multiple comparisons test.

The *GFAP* promoter is particularly active in the hippocampus but much less so in the cortex^37^. In alignment with this pattern and the anticipated difference between mice carrying homozygous or heterozygous copies of the *LacZ*;TRE;*Sfrp1* transgene, staining for the β-galactosidase (βgal) reporter and the GFAP astrocytic marker revealed a differential distribution of double-positive cells in the hippocampus of *Sfrp1^TG^* mice bred with homozygous and heterozygous copies of the transgene. Double-positive cells in the cortex were instead less frequent (**Fig. S1B**) and no βgal expression was observed in controls. Consistent with this distribution, SFRP1 protein levels — determined using a highly specific ELISA^17^ — in hippocampal homogenates of two and ten months old homozygous *Sfrp1^TG^* mice were about 3.5-fold higher than those of age-matched controls (**Fig. 1B, C**). In contrast, only a slight up-regulation was observed in cortical homogenates (**Fig. 1D, E**). Hippocampal homogenates from heterozygous *Sfrp1^TG^* mice showed lower SFRP1 accumulation (about 1.5-fold of controls), although this did not reach statistical significance when evaluated with one-way ANOVA (**Fig. 1B, C**). For most experiments, we bred *Sfrp1^TG^* mice at homozygosis for the *LacZ*;TRE;*Sfrp1* transgene. But, acknowledging the variable expression of transgenic animals independently of the genotype, we grouped the analyzed mice based on SFRP1 hippocampal levels determined with ELISA in *Sfrp1^TG-M^* and *Sfrp1^TG-H^*, according to their medium (40 to 80 ng/mg) or high (>100 ng/mg) SFRP1 levels as compared to the indistinguishable low levels of wildtype (wt) and control *LacZ*;TRE;*Sfrp1* mice (20-30 ng/mg; **Fig. S1C**). Male and female mice did not show significant differences in their SFRP1 content in total brain homogenates (**Fig. S1D**).

### SFRP1 upregulation alters dendritic complexity, spine numbers and presynaptic terminals

SFRP1 is a secreted and highly dispersible protein^39,40^ that localizes to the brain matrisome, influencing several cells types^18,24^. Several studies have indirectly implicated SFRP1 in modulating synaptogenesis and synaptic vulnerability in the hippocampus^27,41,42^. However, there is no information on the consequences of its chronically increased level on neuronal structure and function, a condition observed in the brain of AD patients^17,22,24^. To address this question, we focused on granule cells (GCs) of the dentate gyrus (DG), the first relay of the tri-synaptic pathway originating from the entorhinal cortex, which sustains crucial cognitive functions^43,44^. GCs have a compact dendritic arbor, the modification of which has been well characterized in AD patients and related mouse models^45–47^. We analyzed ten-months old mice as a paradigm for an aging brain, and young two-months old animals because we previously showed that, at this age, SFRP1 levels are already increased in the brain of APP^695swe^;PS1^dE9^ mice, thus preceding amyloid plaque deposition^17^.

To visualize the morphology of GCs from control and *Sfrp1^TG^* mice, we transduced GCs with Sindbis viral particles carrying the green fluorescent protein (GFP) and subsequently analyzed the transduced cells in hippocampal sections with confocal microscopy (**Fig. 2A**). In control mice, GCs exhibited a well-developed dendritic arbor with a Y shaped morphology composed of a short primary dendrite that ramifies in branches of different order invading the molecular layer and decorated by numerous spines. This morphology, comparable to previous descriptions^45–47^, was observed in both two- and ten-months old control mice as well as in two-months-old *Sfrp1^TG-M^* animals (**Fig. 2A**). The dendritic arbor of GCs from *Sfrp1^TG-H^* mice at both analyzed stages was instead simpler, predominantly with a V shape, reduced branching and fewer spines, resembling that described for AD patients^45^ (**Fig. 2A**). Reduced dendritic branches were also observed in ten-months-old *Sfrp1^TG-M^*animals (**Fig. 2A**). Quantitative analysis of these differences using Sholl analysis supported a simplified and less branched dendritic morphology in two-months-old *Sfrp1^TG-H^* GCs (**Fig. 2B**), which only slightly worsened with age compared to controls (**Fig. 2C**). In contrast, the dendritic arbor of GCs from *Sfrp1^TG-M^* mice lost its complexity only in older mice (**Fig. 2B, C**). Quantification of spine density in secondary dendrites from GCs demonstrated a significant 20% reduction in two-months-old *Sfrp1^TG-H^* mice compared to age-matched controls and *Sfrp1^TG-M^* mice, which presented similar densities (**Fig. 2A, D**). At older ages both *Sfrp1^TG-M^* and *Sfrp1^TG-H^* GCs presented a similar decrease in spine density (**Fig. 2A, E**), strongly suggesting that chronic exposure to increased SFRP1 levels leads to synaptic loss.

**Figure 2.**
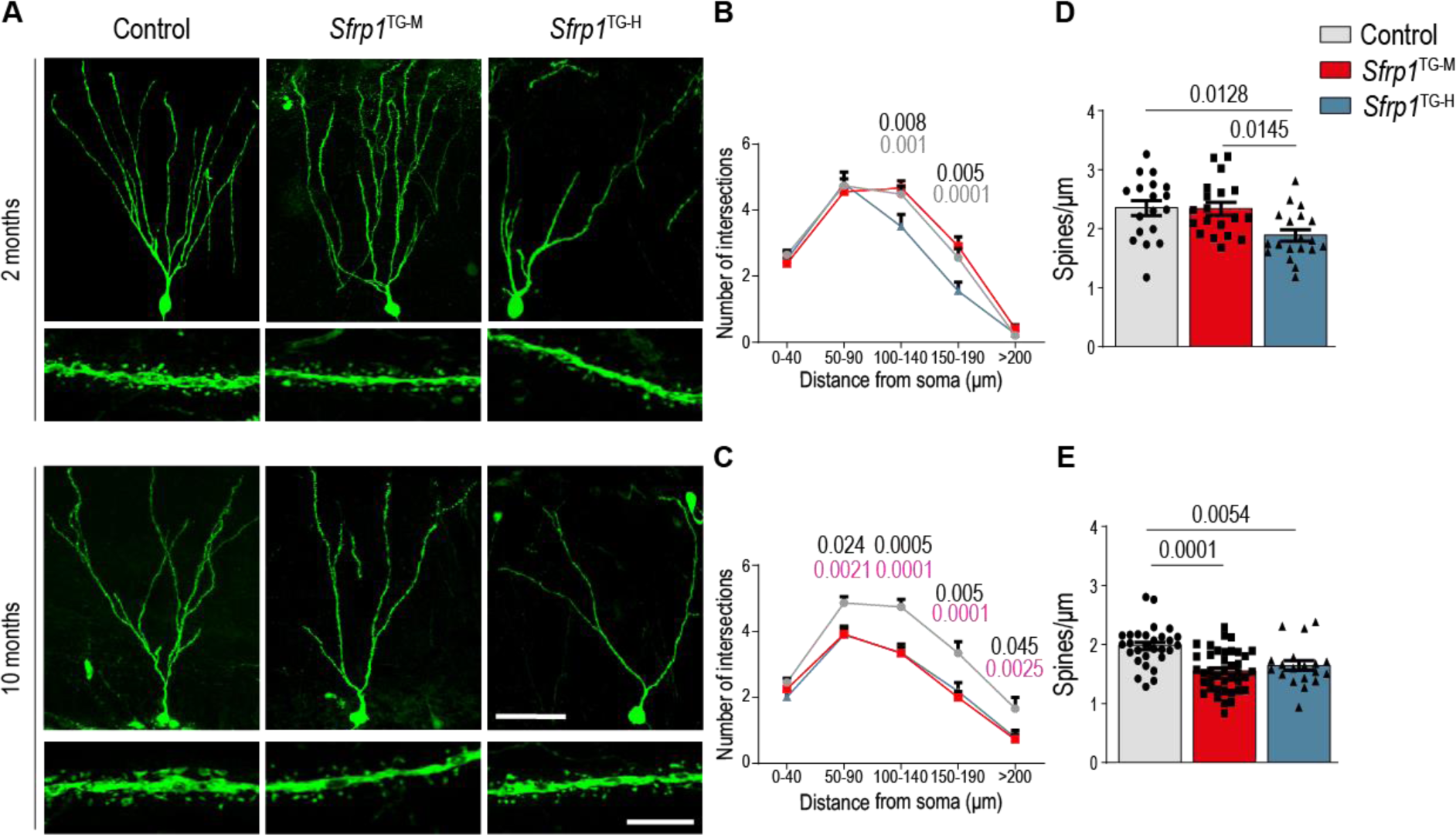
GCs of *Sfrp1^TG^* are poorly ramified and present low spine density. **A**) Representative confocal images of GFP^+^ GCs and dendritic segments observed in the DG of 2 (top) and 10 (bottom) months-old control and transgenic mice after viral transduction. Scale bars: 50 µm (dendritic trees), 5 µm (dendritic segments). **B, C**) Sholl profiles showing the number of intersections of the dendritic trees of GCs with imaginary circles drawn at increasing distances from the soma in two (B) and ten (C) months-old control (n= 33 neurons, 4 mice; n=41 neurons, 4 mice), *Sfrp1^TG-M^* (n=38, 4 mice; n=67, 7 mice) and *Sfrp1^TG-H^* (n=24, 3 mice; n=24, 3 mice). The dots represent mean + SEM. Statistical significance was calculated with two-way ANOVA followed by Bonferroni multiple comparisons test. P values indicated in black indicate control vs. *Sfrp1^TG-H^* comparisons; in grey, *Sfrp1^TG-M^* vs. *Sfrp1^TG-H^*, and in pink, control vs. *Sfrp1^TG-M^*. **D, E**) Quantifications of spine density (calculated as spines/μm) in secondary dendrites from two (D) and ten (E) months old control (n=18 dendritic segments, 3 mice; n=30, 4 mice), *Sfrp1^TG-M^* (n=19, 3 mice; n=40, 7 mice) and *Sfrp1^TG-H^* (n=19, 3 mice; n=19, 3 mice). Graphs represent mean ± SEM. Statistical significance was evaluated by one-way ANOVA followed by Bonferroni multiple comparisons.

To further investigate the effect of SFRP1 on synapses, we used transmission electron microscopy (TEM). We analyzed the structure and distribution of the asymmetric excitatory synapses that entorhinal cortex afferents form with the dendritic spines of the GCs in the molecular layer of the DG^48^. In agreement with the above observations, synaptic density in the molecular layer of both two- and ten-months old *Sfrp1^TG-H^* mice was reduced as compared to age matched controls (**Fig. 3A, B; Fig. S2A, B**). Furthermore, cumulative frequency analysis of the length of the post-synaptic densities (PSD) revealed that, at ten months of age, transgenic mice mostly lose synapses with narrow PSDs but proportionally increase the number of those with larger PSDs (**Fig. 3C**). A similar increase of synapses with larger PSD was also observed at 2 months of age, but, notably, the loss of narrow synapses was much less evident (**Fig. S2C)**. This suggests that *Sfrp1^TG-H^* mice preferentially lose thinner spines with age but stabilize larger ones, given that there seems to be a correlation between PSD and spine’s size^49^. TEM analysis revealed other notable synaptic changes: compared to controls, the presynaptic terminals of *Sfrp1^TG-H^* mice presented a significantly reduced area and evidenced fewer synaptic vesicles at both two and ten months of age (**Fig. 3D, E, F**; **Fig. S2D, E, F**). Notably, similar characteristics have been observed in mice deficient in APP and APP-like proteins^50^ and in the DG molecular layer from postmortem AD brains^51^. Additional but less quantifiable alterations included the increased presence of multi-vesicular bodies, isolated postsynaptic compartments and increased glial cells profiles (**Fig. S2G-I**).

**Figure 3.**
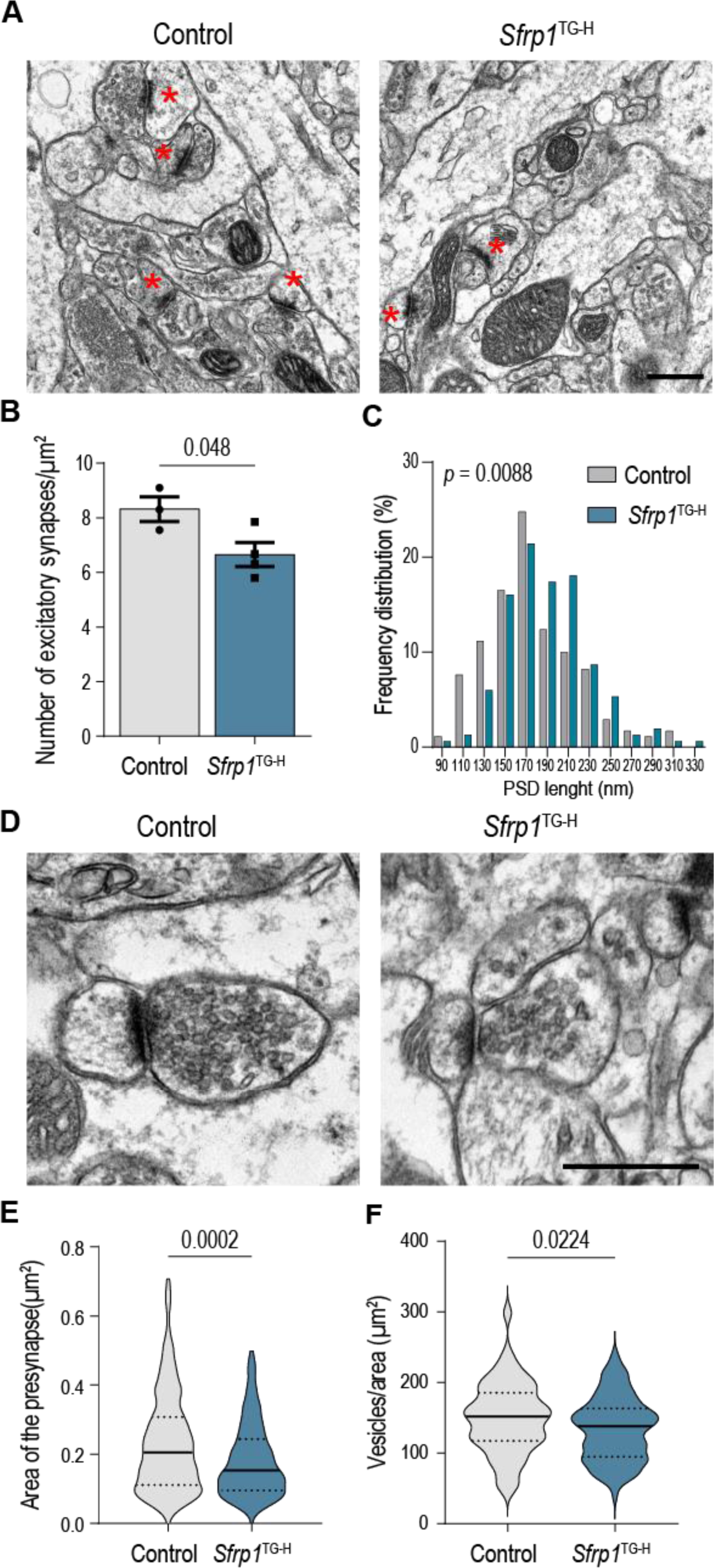
Chronic exposure to high SFRP1 levels correlates with ultrastructural synaptic alterations. **A, D**) Representative low (A) and high power (D) electron micrographs showing asymmetric synapses (red asterisks in A) in the molecular layer of the DG in 10-months-old control and *Sfrp1^TG-H^* mice. Scale bars 0.5 µm. Note the paucity of synaptic vesicles in the presynaptic terminals of transgenic mice. **B**) Quantification of the density of asymmetric synapses in control (n=3) vs transgenic (n=4) mice. The graph represents mean ± SEM. Statistical significance was determined with two-tailed Student’s t-test. **C**) Histogram of control and transgenic synapses according to their PSD length. N = 171 synapses from 3 control, 149 synapses from 4 *Sfrp1^TG-H^* mice. Statistical significance was calculated with Kolmogorov-Smirnov test. **E, F**) Quantification of the presynaptic area (E) and vesicle content (F) in asymmetric synapses. Violin plots represent data distribution, median (solid line) and 25% and 75% quartiles (dotted lines). N = 268 synapses from 3 control and 217 synapses from 4 *Sfrp1^TG-H^* mice in (E), 102 control and 105 transgenic synapses in (F). Statistical significance was calculated with two-tailed Mann-Whitney test.

Taken altogether these observations indicate that chronic exposure to abnormally elevated SFRP1 levels negatively impacts on GCs morphology and structural aspects of synaptic connectivity, with alterations mimicking those described in the DG of AD patients.

### Sfrp1 directly impacts on neuronal structure

The morphological changes observed in transgenic GCs could result from direct SFRP1 activity on the neurons or a from a glia-mediated effect. To distinguish between these possibilities, we treated primary cultures of hippocampal neurons from embryonic wt mice with human recombinant SFRP1 (hrSFRP1, 400 ng/ml) conjugated to the Alexa-488 fluorophore from 10 to 180 min. The cultures were thereafter fixed and immunostained with anti-MAP2 to visualize the neurons. The fluorescent signal decorated the surface of MAP2-positive neurons after 30 min of incubation (**Fig. 4A**) and seemed to localize with increasing frequency on intracellular vesicles at 30 and 180 min (**Fig. 4A, B**), indicating that the protein binds to the neuronal surface and is likely internalized. Furthermore, the hrSFRP-Alexa-488 signal colocalized with an anti-SFRP1 staining (**Fig. S3A, B)**, ruling out artifacts due to unconjugated Alexa-488 binding. The association of SFRP1 with the neuronal membrane was confirmed using hippocampal synaptosomal preparations from ten-months old control and *Sfrp1^TG-H^* animals (see last section of the results). Western blot and ELISA analysis of the synaptosomes’ soluble and membrane fractions, obtained by ultracentrifugation, revealed that SFRP1 highly associates to the membranes in both control and transgenic samples, with a significant enrichment in the latter (**Fig. 4C, D**). Notably, prolonged (from 5 to 14 DIV) exposure of neurons to hrSFRP1 resulted in a simplified morphology with fewer and shorter processes than saline treated control neurons (**Fig. S3C, D**).

**Figure 4.**
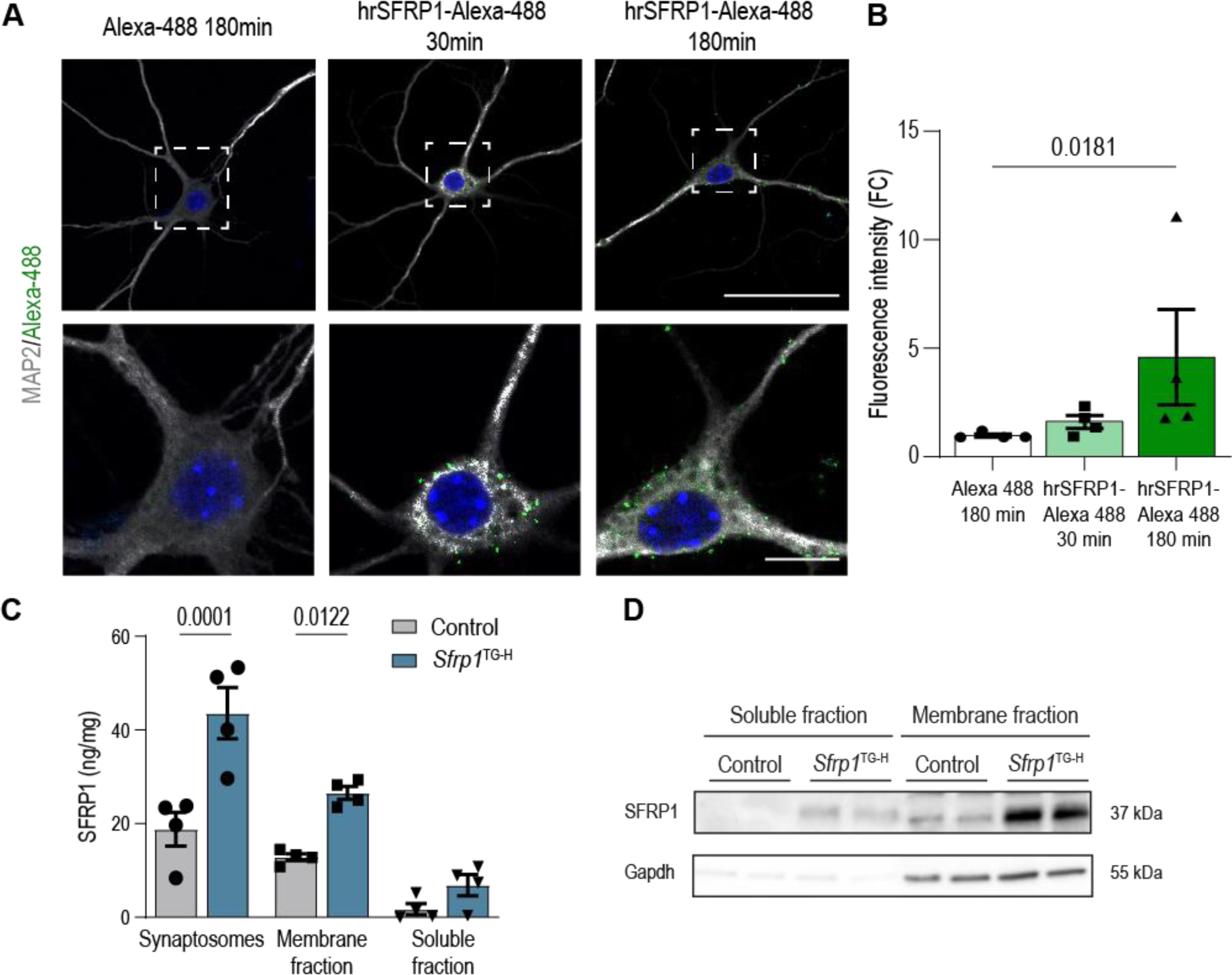
Hippocampal neurons bind and internalize SFRP1. **A)** Representative confocal images of wt hippocampal neurons treated with 400 ng/ml hrSFRP1-Alexa488 for 30 and 180 min and immunostained with anti-MAP2. **B)** The graph represents the mean fluorescence intensity of Alexa488 alone or conjugated with hrSFRP1 colocalizing with MAP2 in wt hippocampal neurons. Data (mean ± SEM) are presented as fold change (FC) over the control condition. N = 4 independent cultures. Statistical significance was calculated with Krustal-Wallis followed by Dunn’s multiple comparisons post-hoc analysis. **C**) ELISA determination of SFRP1 concentration in hippocampal synaptosomes from 10 months-old control and *Sfrp^TG-H^* mice and in the corresponding membrane and soluble fractions obtained by ultracentrifugation. Data are shown as mean ± SEM, n = 4 samples for each genotype and fraction. Statistical significance was evaluated with two-way ANOVA followed by Bonferroni post-hoc analysis. **D)** Western blot analysis of SFRP1 in membrane and soluble fractions obtained from control and transgenic hippocampal synaptosomes.

Taken together these data indicate that SFRP1 directly binds to neuronal membranes and negatively influences neurite branching.

### Alert-behaving Sfrp1^TG^ mice present an altered synaptic function

To determine whether the morphological alteration observed in *Sfrp1^TG^* mice had functional consequences, we compared the electrophysiological properties of the perforant pathway (PP) to DG synapses in ten-months-old alert-behaving control and *Sfrp1^TG^* mice (**Fig. 5A**), using input/output (I/O), paired-pulse facilitation (PPF) and LTP protocols^52^.

**Figure 5.**
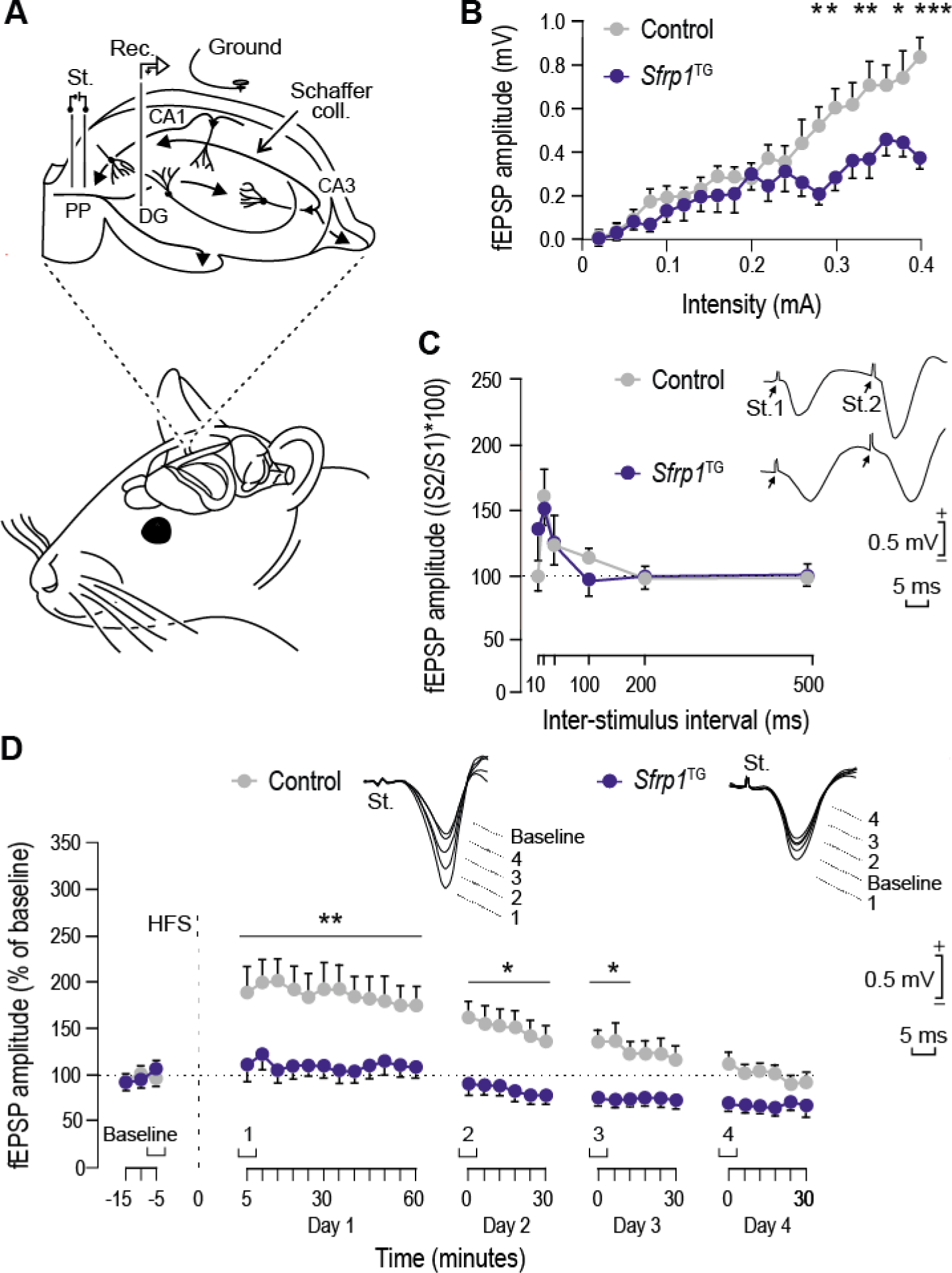
*Sfrp1^TG^* mice present reduced basal synaptic transmission and LTP response of the PP-DG pathway. **A)** A bipolar stimulating plus two recording electrodes were implanted in the performant pathway (PP) and ipsilateral dentate gyrus (DG), respectively. Abbreviations: Coll, collaterals; Rec, recording; St, stimulation**. B)** To perform input/output curves, single pulses were presented to the PP at increasing intensities while recording the evoked fEPSP at the DG area for controls (n=4) and *Sfrp1^TG^* (n=6) mice. Data are presented as mean ± SEM and analyzed with two-way ANOVA followed by the Holm-Sidak method. **C)** Double pulse facilitation evoked in the two groups of mice (n=5 mice/group). Representative examples (averaged five times) of fEPSPs evoked by paired pulses at 40 ms of inter-pulse interval in the two groups of mice are illustrated in the right side of the graph. Data shown are mean ± SEM amplitudes of the second fEPSP expressed as the percentage of the first for six inter-pulse intervals. Data were analyzed with two-way ANOVA followed by the Holm-Sidak method. **D)** LTP evoked in controls (n=12) and *Sfrp1^TG^* (n=13) mice. Following 15 min of baseline recordings, mice were presented with a high frequency stimulation (HFS) protocol indicated by the dashed line. LTP evolution was followed for a total of four days. Illustrative examples (averaged five times) of fEPSPs recorded at the times indicated in the bottom graph and collected from representative control and transgenic mice are shown at the top. Data are mean ± SEM amplitudes. Two-way ANOVA followed by the Holm-Sidak method was used to evaluate statistical significance. *p < 0.05, **p<0.01, ***p<0.001.

When the PP from transgenic mice was stimulated with single pulses of increasing intensities (I/O protocol; 0.02–0.4 mA in 0.02 mA steps), the field excitatory postsynaptic potentials (fEPSPs) evoked at higher intensities were significantly smaller [F(_1,19_) = 5.709; p < 0.001] than that of control mice (**Fig. 5B, Table S1**), suggesting that SFRP1 overexpression impairs basal synaptic transmission at the PP-DG synapses. In contrast, no significant difference was observed between control and transgenic mice when the PP was stimulated using a PPF protocol consisting of paired pulses with different inter-stimulus intervals (10-500 ms). Both genotypes presented an enhanced amplitude of the evoked fEPSPs after the second pulse at short inter-stimulus intervals (**Fig. 5C, Table S1**). High-frequency stimulation (HFS) of the PP-DG synapses evoked LTP in controls [F(_1,32_) = 16.176; p < 0.001], but not in *Sfrp1^TG^* mice (**Fig. 5D**). PP stimulation evoked fEPSPs that were consistently smaller [F(_1,32_) = 4.229; p < 0.001] in transgenic mice as compared to controls across four sessions, recorded right after the HFS presentation and at 24, 48 and 72 hrs post-stimulation (**Fig. 5D, Table S1**). In other words, whereas controls presented I/O, PPF and LTP values comparable to those previously described for wt using similar approaches^52^, *Sfrp1^TG^* mice presented significant deficits in I/O curves and LTP.

We next asked whether these alterations were specific of the PP-DG connections or were common to additional downstream hippocampal synapses. To test this possibility, we analyzed the functionality of the CA3 Shaffer projections onto CA1 pyramidal cells (CA3-CA1 synapses; **Fig. S4A**), which lie downstream of the PP-DG input. No significant differences were observed in I/O curves or in PPF between controls and *Sfrp1^TG^* mice (**Fig. S4B, C, Table S1**). However, the amplitude of the LTP response following HFS was smaller in transgenic than in control mice, [two-way ANOVA, F_(1,38)_ = 1.155; *P* = 0.242, followed by the Holm-Sidak method, t = 2.073; *P* < 0.05] in the first recording session, up to 60 minutes after the HFS (**Fig. S4D, Table S4**).

Two months old *Sfrp1^TG-H^* mice already show structural alterations in their synapses as compared to age-matched controls (**Fig. S2**). Therefore, we asked whether these mice already have an impaired electrophysiological response. To this end, we evaluated CA3-CA1 connectivity using a more amenable experimental set up, using acute hippocampal slices form 2 months old control and *Sfrp1^TG-H^* mice. Electrophysiological recordings of *Sfrp1^TG-H^* slices showed no differences in PPF or LTP compared to control littermates (**Fig. S5B-D**). However, the transgenic hippocampi presented a slight decrease in basal synaptic transmission, evidenced by the lower response to the I/O protocol (**Fig. S5A**).

Taken together, these data show that, with age, high SFRP1 levels impair basal synaptic transmission and synaptic plasticity in the intrinsic hippocampal circuit.

### SFRP1 overexpression impairs hippocampal-dependent memory with age

The observed synaptic defects in transgenic mice suggest potential alterations in hippocampal-dependent cognition. To test this, we compared the performance of control and *Sfrp1^TG^* mice at two and ten months of age using the Y maze and novel object recognition tests.

The Y maze exploits rodents’ innate exploratory behavior and assesses short-term memory^53^ (**Fig. 6A**). At two months, all transgenic mice, regardless of brain SFRP1 content, matched the controls’ exploration time of the novel arm (**Fig. 6B**). However, at ten months, *Sfrp1^TG-H^* mice spent significantly less time in the new arm than controls (28% vs. 40% of total exploration time; **Fig. 6C**). *Sfrp1^TG-M^* mice also performed worse than controls (35%; **Fig. 6C**), though not reaching statistical significance in one-way ANOVA testing.

**Figure 6.**
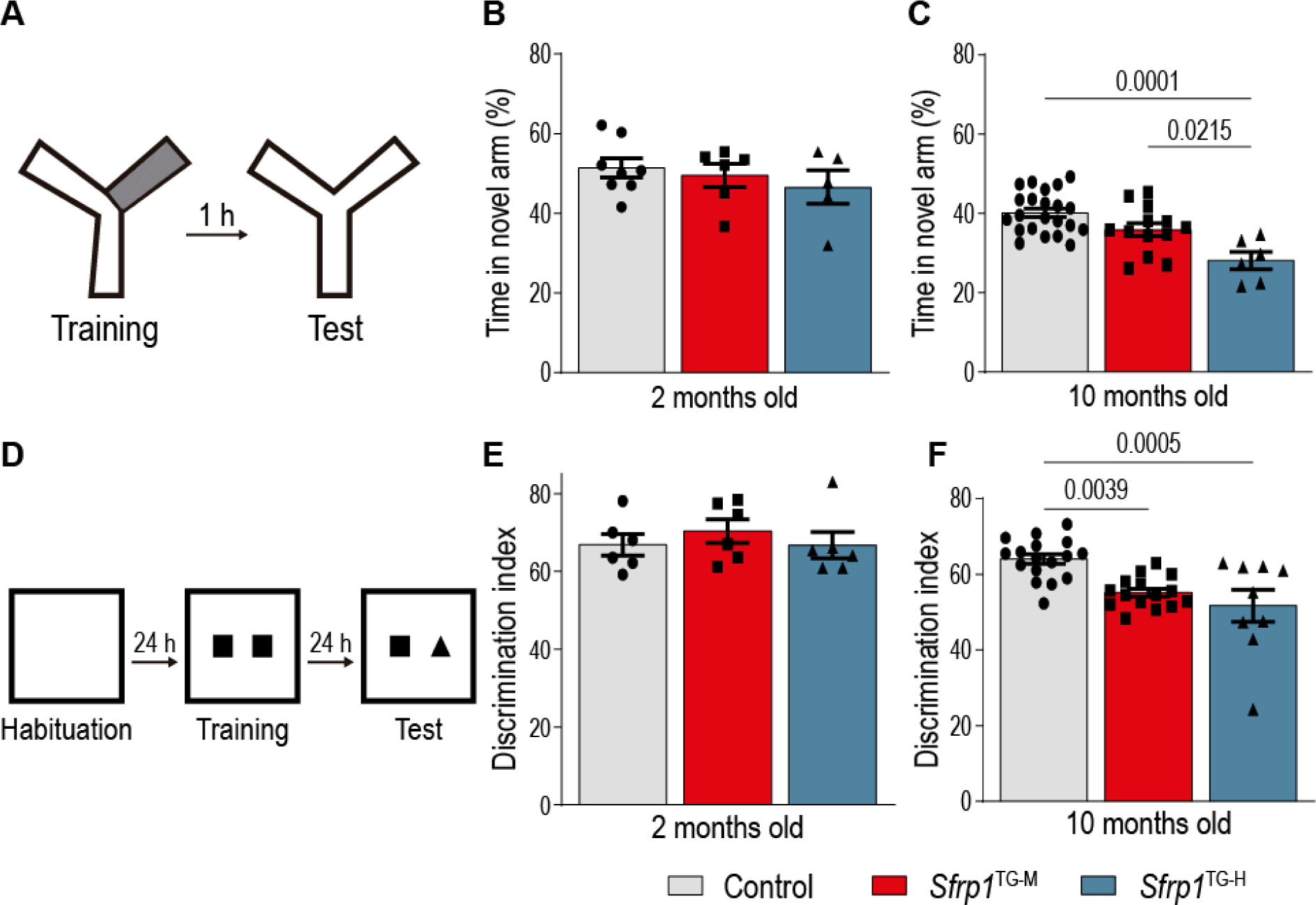
*Sfrp1^TG^* mice present cognitive impairment as they age. **A**) Schematic representation of the Y maze set up. The grey area represents the arm that was kept closed during the training session. **B, C)** The graphs show the performance of two (B) and ten (C) months-old control and transgenic mice during the testing phase of the Y maze. Data represent the percentage of time spent in the new arm over the total exploration time. Control, n=8; *Sfrp1^TG-M^*, n=6 and *Sfrp1^TG-H^* n=5 in (B); control n=22, *Sfrp1*^TG-M^, n=13; *Sfrp1^TG-H^* n=6 mice in (C). **D)** Schematic representation of the NOR test. **E, F)** The graphs represent the discrimination index, calculated as the percentage of time exploring the new object vs. total object exploration time in two (D) and ten (E) months old mice. Each experimental group included n=6 in (E) and control n=17, *Sfrp1^TG-M^* n=15 and *Sfrp1^TG-H^* n=9 mice in (F). All data are represented as mean ± SEM, statistical significance was evaluated with one-way ANOVA followed by Bonferroni multiple comparisons test.

The novel object recognition test (NORT) exploits the mouse innate preference for novelty and evaluates different aspects of their learning and memory ability^54^ (**Fig. 6D**). In this test, performed 24 hrs after the training phase, 2-months-old *Sfrp1^TG^* behaved like controls (**Fig. 6E**), indicating that spatial memory is not impaired at a younger age, well in line with the electrophysiological recordings at this time point. However, at ten months of age, both *Sfrp1^TG-M^* and *Sfrp1^TG-H^* mice failed to discriminate the novel versus the familiar object, as indicated by the discrimination index (time spent exploring the new object vs. total exploration time) (**Fig. 6F**). Notably, when tested in an open field and rotarod apparatus, all transgenic mice showed locomotor activity, exploratory behavior and motor coordination indistinguishable from those of controls (**Fig. S6**).

All in all, these observations indicate that despite chronic exposure to high SFRP1 levels, transgenic mice show resilience to behavioral changes, given that most morphological alterations well precede cognitive deterioration.

*Sfrp1^TG-H^* mice present a 3.5-fold increase in SRFP1 level in the hippocampus but the increase is much more modest in the cortex (**Fig. 1**). To test whether this difference has a behavioral correlate, we compared the associative learning capabilities of 10 months old control and *Sfrp1^TG^* mice, given that the acquisition of operant conditioning tasks implies the proper functioning of motor^55^ and prefrontal circuits^56^. Mice were individually trained in Skinner boxes to press a lever for collecting a food pellet with a fixed schedule for a total of ten sessions of 20 min (**Fig. S7A**). Both control and *Sfrp1^TG^* mice reached the selected criterion (to press the lever ≥ 20 times for two successive sessions) within four-six training sessions with no significant difference (*P* = 0.227) and reached asymptotic values by the fifth training session (**Fig. S7B**). The learning curves of the two genotypes were also similar with no significant difference (F_(1,9)_ = 0.768, *P* = 0.646), indicating that *Sfrp1^TG^* and control mice have similar associative learning ability (**Fig. S7C**). These data indicate a strong correlation between SFRP1 accumulation, synaptic defects and behavioral alterations: hippocampal physiology and related behavioral functions are altered whereas those supported by the neocortex, where SFRP1 accumulates poorly, are instead intact.

### Sfrp1 overexpression modifies the synaptic proteome

To determine what are the maladaptive molecular pathways underlying the phenotype observed in the hippocampus of *Sfrp1^TG-H^* mice, we opted for unbiased approach by comparing the hippocampal proteome of 10-months-old control and *Sfrp1^TG-H^* mice.

We prepared hippocampal homogenates and the corresponding synaptosomal fractions^57^ (**Fig. S8**) to enable the identification of both broader and synaptic-specific changes. Both sets of samples were subjected to analysis using data-independent analysis liquid chromatography with tandem mass spectrometry (DIA LC-MS/MS)^58^. This analysis identified, on average, 51.000 unique peptides that mapped to 6.700 unique proteins. The MS-DAP platform^59^ (mass spectrometry downstream analysis pipeline) was used for normalization, quality control and differential abundance analysis of the proteomic data. Analysis of peptide detection frequency and coefficient of variation (CoV) within each experimental group indicated a high reproducibility among samples (**Fig. S8B**), with the exception of a control homogenate that was thus omitted from further analysis. For differential abundance analysis, we only considered peptides detected in at least 6 out of the 8 samples within each group. An adjusted p-value threshold of q ≤ 0.05 was used to discriminate significantly altered proteins. SFRP1 did not pass the first filtering criterion, as it was not consistently detected in all control samples likely due to its low molecular weight, glycosylation, content of disulfide bridges^60^ and relatively low concentration under homeostatic conditions. However, SFRP1 emerged as a differentially abundant protein in transgenic vs. control homogenates when we consider every protein found in at least one sample from each group for differential abundance analysis (**Table S2**). A significant enrichment was also found when the same samples (homogenates and synaptosomes) were tested for SFRP1 content with ELISA (**Fig. S8C**).

Out of the approximately 6.700 proteins identified across all the controls and *Sfrp1^TG-H^* samples, we identified 30 (homogenates) and 29 (synaptosomes) differentially abundant proteins (DAPs; **Fig. 7A, 8A**; **Table S3**) with only one protein common to both fractions, Wdr7, involved in vesicle acidification. Gene ontology (GO) analysis of the homogenate DAPs using the g:Profiler webtool^61^ highlighted “protein translation initiation / regulation” as categories downregulated in the transgenic samples (**Fig. 7B**). These include proteins mostly expressed in neurons^62^, such as the translation preinitiation or initiation factors eIF3b, eIF4g3, eIF3e, eIF3l or the regulators Mtrex, Ewsr1 and Sf1. Seventeen additional proteins (e.g. eIF3g; eIF3m; eIF3f; eIF3b) fell in this category when p-value ≤0.05 was considered (**Table S3**). A negative effect of SFRP1 on global protein synthesis was supported by the use of the surface sensing of translation (SUnSET) assay, based on the detection of the incorporation of puromycin, a protein synthesis inhibitor, in cell cultures^63^. Indeed, SFRP1 addition to hippocampal neuronal cultures reduced puromycin incorporation compared to saline treated cultures (**Fig. 7C**). Notably, the astrocyte-derived extracellular proteins thrombospondin 4 (Thbs4) and SPARC-like protein 1 (Sparcl1), which have been shown to increase dendritic complexity and spine density in cultured neurons^64^, were also significantly decreased in the *Sfrp1^TG-H^* homogenates (**Table S3**). Analysis of the proteins identified in the synaptosomal samples using the Synaptic Gene Ontologies (SynGO) database^59^ (https://www.syngoportal.org/index.html) confirmed an enrichment in synaptic proteins in this fraction (**Fig. 8A, B**). GO and SynGO analysis of the DAPs identified the terms “presynapse”, “synaptic vesicle”, “postsynapse” and “postsynaptic density” as significantly enriched in the transgenics (**Fig. 8B, C**). Elevated proteins associated with synaptic vesicle function and endosomal compartments comprised Wdr7, Syt12, Atp6v0a1, Ap2a2, and Rogdi, whereas the presynaptic Nrxn3, Epha4, Cacna1a, Pip5k1c and the post-synaptic Nrp1 emerged among the upregulated synaptic organizing molecules (**Fig. 8A**, **Table S3).** Additional upregulated synaptic proteins included, among others, Ephb2, Dnm1, Cadm3, Nptn; Psen1 Camk2b, Homer3, L1-CAM (**Table S3**), when p-value ≤0.05 was considered.

**Figure 7.**
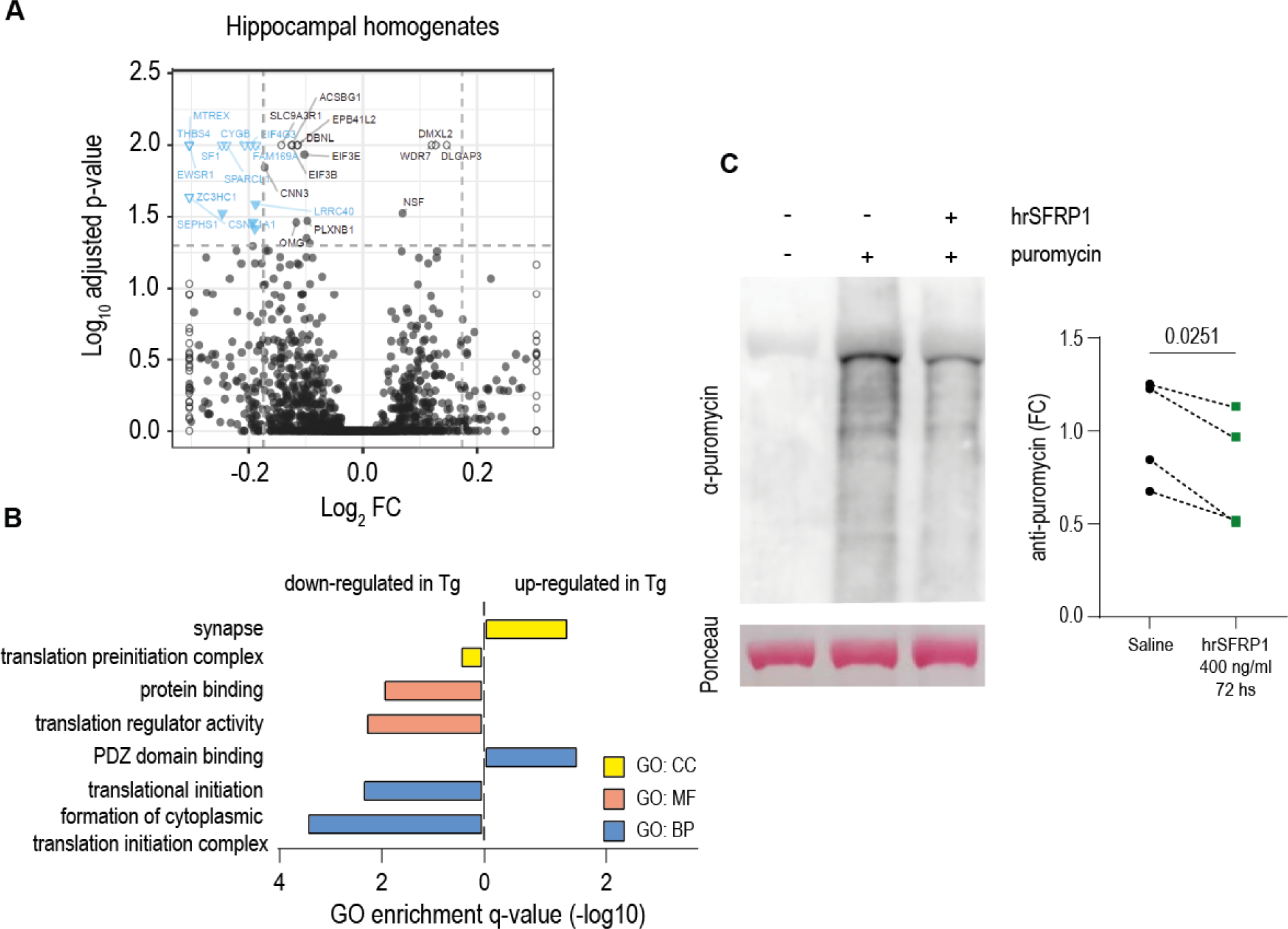
*Sfrp1* overexpression dampens hippocampal protein synthesis. **A)** Volcano plot showing the log10 adjusted p-value (y-axis) and the log2 fold change (x-axis) of all quantified proteins in hippocampal homogenates from 10 months old control and *Sfrp1*^TG-H^ mice. **B**) GO term enrichment analysis of the differentially abundant proteins found in *Sfrp1^TG^*^-*H*^ hippocampal homogenates as compared to controls. The three classical GO categories were assessed for upregulated and downregulated proteins separately. CC: Cellular components, MF: molecular function, BP: biological process. **C**) Representative image (top) and quantification (bottom) of a SUnSET Western blot used to compare puromycin-labelled proteins in hippocampal neuronal cultures (DIV 17) grown in the presence or absence of SFRP1 for 72hrs. The Ponceau staining is shown as a loading control. Cultures were incubated with or without (as control, saline) puromycin (1 µg/ml) for 1 hr. The data in the graph correspond to each individual culture (n=4) presented as fold change (FC) over the control condition. Statistical significance was evaluated with two-tailed paired-t-test.

**Figure 8.**
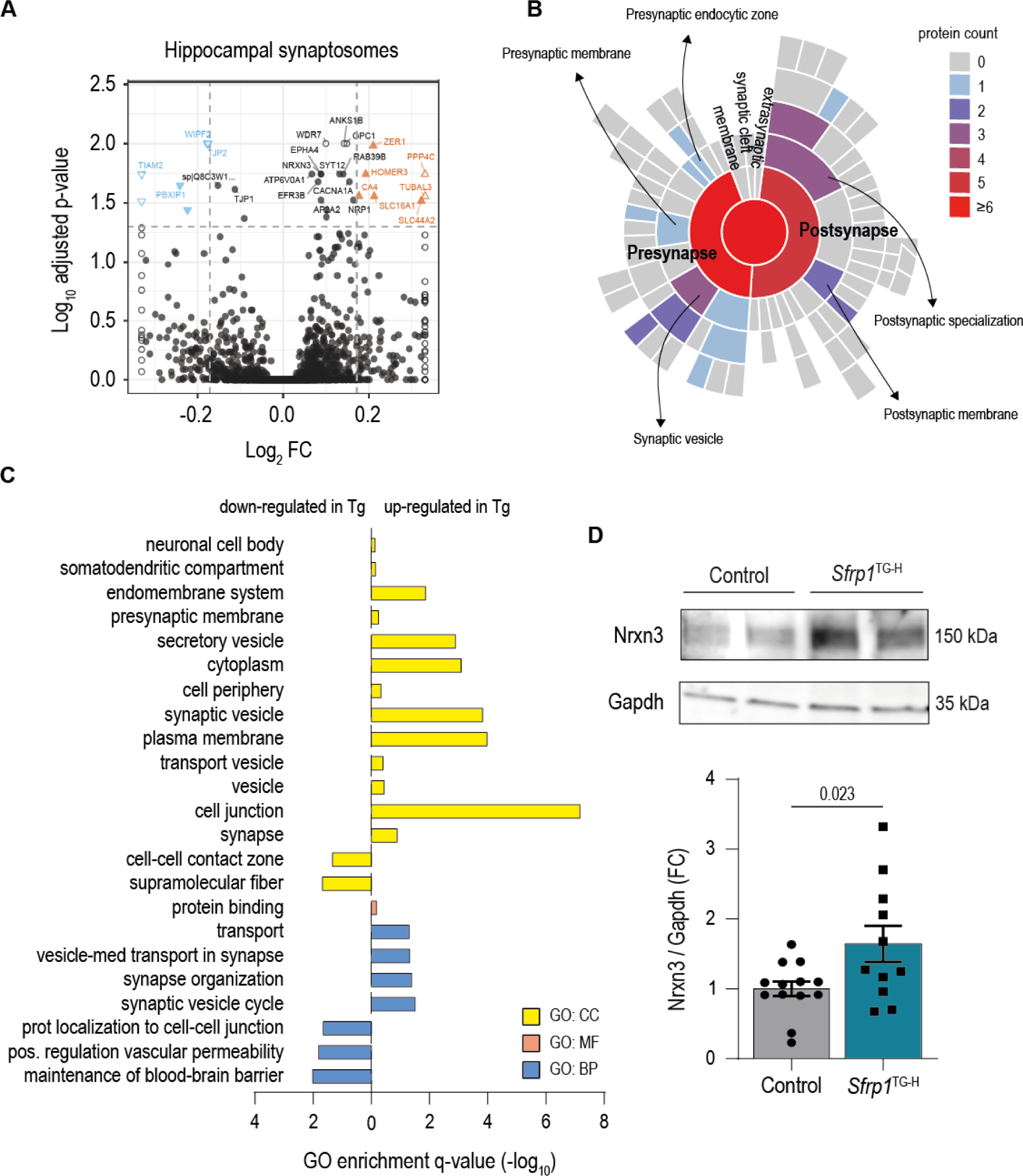
*Sfrp1* overexpression modifies the hippocampal synaptic proteome. **A)** Volcano plot showing the log10 FDR adjusted p-value (y-axis) and the log2 fold change (x-axis) of all quantified proteins in hippocampal synaptosomes from 10 months old control and *Sfrp1*^TG-H^ mice. **B**) Sunburst blot showing the SynGO terms that were found enriched in the transgenic synaptosomes. **C**) GO term enrichment analysis of the differentially abundant proteins found in *Sfrp1^TG^*^-*H*^ hippocampal homogenates as compared to controls. The three classical GO categories were assessed for upregulated and downregulated proteins separately. CC: Cellular components, MF: molecular function, BP: biological process. **D)** Representative image (top) and quantification (bottom) of a Western blot analysis of Neurexin 3 in the membrane fraction from control (n=13) and *Sfrp1*^TG-H^ (n=11) hippocampal synaptosomes, presented as fold change (FC) over control values. Data are presented as mean ± SEM and were analyzed with two-tailed Student-t test.

The upregulated levels of synaptic organizing molecules appeared particularly interesting given their critical role in synaptic plasticity and synapse elimination^6^. We hypothesized that at least part of the synaptic changes we observed could be linked to the persistency of molecules such as neurexins, L1-CAM, Eph4 and Eph2 at the synapses, thereby interfering with synaptic plasticity. To test this possibility, we focused on neurexins. Neurexins constitute a protein family with a large number of isoforms involved in synaptic assembly and remodeling^65^ and their expression is consistently upregulated in the brain of AD patients^21^. Furthermore, their assembly in synaptic nanoclusters seems to be regulated by ADAM10^32^, the activity of which is down modulated by SFRP1^66^. Western blot analysis of synaptic membrane preparations supported this possibility showing significantly increased levels of Neurexin3 in synaptic membranes from *Sfrp1^TG-H^* compared to those of controls (**Fig. 8D**).

A recent study has compiled a large number of data sets reporting proteomic changes occurring in the brain of AD patients at different stages of the disease (preclinical, mild cognitive and advanced) and generated a searchable database (https://neuropro.biomedical.hosting) that can be interrogated for meta-analysis^21^. This compendium highlights that proteins belonging to the synapse or the matrisome (including SFRP1) are among those that show consistent changes starting from preclinical stages of the disease^21^. We thus used the Neuropro database to identify possible DEG similarities between AD and *Sfrp1^TG-H^*. Out of the 29 *Sfrp1^TG-H^* synaptosome associated DAPs (**Table S3**), 26 (89%) were also listed in Neuropro (23/30; 76% in homogenates), although the differential expression values were not always in the same direction, as also observed among different original sources included in the study^21^. On the other hand, and consistent with our proteomic data, Neuropro does not identify the main components of the Wnt signaling pathways or its common down-stream targets (e.g. Axin2, CyclinD1 and c-Myc) among the AD associated DEGs, as possibly expected if SFRP1 would act as a Wnt signaling modulator^60^. An exception is the Wnt signaling inhibitor Dkk3, which is consistently upregulated in AD brains and not differentially detected in our proteomic data. The lack of Wnt signaling components among the identified DAPs may simply reflect that cascade activation relies mostly on post-translational modifications and thus proteins levels may remain unchanged. We therefore tested whether the transcription of Wnt target genes is affected in *Sfrp1^TG^* mice. We assessed *Axin2*, *CyclinD1* and *c-Myc* mRNA levels in control and *Sfrp1^TG-H^*hippocampi using qPCR, but found no expression differences (**Fig. S9**), indicating that Wnt signaling alterations are an unlikely explanation for the phenotype observed in *Sfrp1^TG^* mice.

Taking all these data together we propose that chronically elevated levels of SFRP1 negatively influence the turnover of synaptic organizing molecules, thereby leading to additional molecular and cellular alterations that, with age, result in loss of cognitive functions.

## Discussion

A growing number of genetic and "omic" studies on human brain employing large patient cohorts are yielding insights into the intricate molecular pathways disrupted in AD^21,67^. These investigations have underscored an important role of glial cells not only in disease’s progression but also at its onset, exerting a consequential influence on synaptic function^14^. In particular, proteomic analyses of human brains have unveiled that preclinical AD stages are characterized by significant molecular alterations in both the matrisome and synapse components^21–23,66,68^. In line with this finding, our study unveils a crucial relationship between the sustained overexpression of the glial-derived matrisome protein SFRP1 and the emergence of early dendritic and synaptic dysfunction in the mouse hippocampus, ultimately culminating in a poor LTP response and cognitive impairment as the animals age. These structural and functional SFRP1-induced maladjustments are linked to an array of molecular changes, including the increased expression of structural synaptic proteins. These observations, coupled with the notion that SFRP1 fosters Aβ generation and aggregation as well as neuroinflammation^17,18^, position SFRP1 among the factors that contribute to multiple aspects of AD pathogenesis, and present it as a potentially important therapeutic target.

The present study originated from the recognition that SFRP1 functions as a negative modulator of ADAM10 shedding activity^66^ and thereby has the potential to interfere with the processing of the numerous ADAM10 substrates, encompassing APP, proteins involved in the control of neuroinflammation and synaptic adhesion molecules^69^ — all of which play roles in AD pathology. Consequently, we postulated that the consistently elevated SFRP1 levels observed in the brains of AD patients^21–23,66,68^ might influence synaptic function directly and independently of amyloid plaque accumulation. The results of this study substantiate this hypothesis and further suggest resemblances between the phenotype of *Sfrp1^TG^* mice and specific features observed in AD patients.

GCs of *Sfrp1^TG-H^* mice exhibit a distinctive V shaped dendritic arbour with the preferential loss of distal dendrites and small spines mirroring the observations in the GCs from the brain of AD patients^45–47^. These structural abnormalities are linked to the hippocampal levels of SFRP1, given that *Sfrp1^TG-H^* mice manifest an earlier onset of dendritic and spine loss than *Sfrp1^TG-M^* mice. These defects are likely attributable to the direct binding and internalization of SFRP1 in neurons. Indeed, we observed that fluorescently labelled SFRP1 protein decorates the plasma-membrane and endocytic vesicles of cultured neurons, and the membrane fraction of synaptosomal preparations is enriched in SFRP1. Furthermore, neurons cultured in virtual absence of glial cells and exposed for several days to SFRP1 have a poorly developed neuritic tree. This does not rule out the possibility that, in vivo, additional mechanisms, potentially involving ECM and glial cells, may contribute to the observed *Sfrp1^TG^*phenotype. In fact, SFRP1 mediates astrocytes-microglia cross talk^18^ and both glial cell types contribute to synaptic elimination in homeostatic and pathological conditions^70,71^.

A second notable similarity between our *Sfrp1^TG^* model and AD is the resilience to functional changes (loss of cognitive capability and prominent electrophysiological alterations) despite early morphological and structural alterations in neurons. Indeed, despite the presence of structural alterations, *Sfrp1*^TG-H^ mice present a normal synaptic function at 2 months of age, but both synaptic structure and function are impaired by 10 months of age. The most notable change between the two analyzed time points is the late selective loss of thin spines, which are thought to be important for maintaining the levels of synaptic plasticity required for cognitive function^72^. Thus, it is tempting to speculate that, in *Sfrp1^TG-H^* mice, both functional and behavioural changes become apparent when the number of small spines goes below a critical threshold. In addition, compensatory mechanisms may further counteract the harmful effect of high SFRP1 levels, as proposed to explain the resilience of the human/mouse brain to the accumulation of toxic Aβ peptides^73^. These harmful effects include an impaired basal synaptic transmission and long term potentiation of the PP-DG and, to a lesser extent, CA3-CA1 synapses, well in line with the notion that antibody-mediated neutralization of SFRP1 in adult APP^695swe^;PS1^dE9^ mice rescues the LTP defects that are typically observed in this AD model. In contrast, *Sfrp1* genetic inactivation improves cognitive performance in APP^695swe^;PS1^dE9^ mice^17^, supporting the idea that excessive SFRP1 inhibits hippocampal synaptic plasticity. Why CA3-CA1 synapses are less affected by SFRP1 accumulation, is just a matter of speculation, but the simpler dendritic structure of GCs might make these more susceptible to dendritic and synaptic elimination compared to CA3 and CA1 pyramidal neurons.

A third commonality between the brains of *Sfrp1^TG^* and AD patients is the occurrence of a number of proteomic changes, predominantly affecting proteins involved in vesicle cycle and synaptic adhesion. The presynaptic terminals of *Sfrp1^TG^* mice are characterised by a depleted reservoir of synaptic vesicles, suggesting that the concomitant higher abundance of proteins involved in vesicle recycling may reflect a compensatory mechanism to preserve presynaptic function. This compensation may be associated to a more rapid turnover of the exocytic vesicles or an enhanced synaptic vesicle release, as demonstrated in iPSC-derived neurons from Down syndrome patients^74^, who suffer from early-onset AD. The latter possibility would also align with the idea that synapses in *Sfrp1^TG^* hippocampi are less plastic due to higher levels of synaptic adhesion proteins, such as L1CAM, Eph4, neurexin and neuroligin, all increased in *Sfrp1^TG^* synaptosomes. Among them, neurexins, a protein family with a large number of isoforms^65^ might be particularly relevant for the *Sfrp1^TG^* phenotype.

Neurexins isoforms are cell type specific and localize at the presynaptic terminals, where they can interact with a number of postsynaptic proteins such as neuroligin, as well as extracellular proteins, including the astrocyte derived Thbs4 and SPARC^64^, that we found downregulated in *Sfrp1^TG^* mice. Apart from their heterogeneity, neurexins and the molecular interactor neuroligin are well-characterized substrates of ADAM10^32,75,76^. Thus, neurexin shedding, mediated by ADAM10, is an additional mechanism that enhances the turnability and plasticity of synaptic contacts. ADAM10 localises at both the pre- and post-synaptic compartments of excitatory synapses^28,29^. Its conditional inactivation in the brain impairs synaptic plasticity^30^. Therefore, it is plausible that SFRP1 accumulation in the synaptic membrane fraction may reflect its interaction with ADAM10, as we have demonstrated previously in the synaptosomes of APP^695swe^;PS1^dE9^ mice^17^. This interaction is expected to limit ADAM10 activity, consequently affecting the turnover of neurexin, neuroligin and other ADAM10 substrates (e.g. L1CAM or Eph4)^76^ at the synaptic plasma membrane. This reduced turnover could explain the observed increase of spines with larger PSD in *Sfrp1^TG^*. This feature resembles the morphological alterations of dendritic spines in the hippocampus of *Adam10* conditional knock-out mice, which are associated with abnormal processing of APP, N-Cadherin and Nectin-1^30^. In our study we could not prove abnormal Neurexin3 processing due to the lack of appropriate tools to detect the proteolytic fragments. Nevertheless, inhibition of ADAM10-mediated shedding of neurexin has been shown to substantially increase its synaptic content^32^, mirroring our observations in *Sfrp1^TG^* synaptosomes.

In conclusion, we propose that chronic exposure to SFRP1 reduces ADAM10 shedding of synaptic organizing molecules, limiting their turnover and, consequently, reducing synaptic plasticity and LTP response, thereby causing memory impairment. These alterations are likely the cause of the additional molecular changes observed in the *Sfrp1^TG^* hippocampus. The decreased levels of proteins involved in translational control pathways found in the homogenates from *Sfrp1^TG^* hippocampi are particularly relevant because protein translation is critical for synaptic plasticity and memory consolidation^77,78^. Furthermore, addition of SFRP1 to hippocampal cultures lead to reduced puromycin incorporation, indicating that SFRP1 directly influences neuronal translation mechanisms. In patients suffering from AD, elevated SFRP1 levels may lead to similar synaptic alterations, as well as increased plaques accumulation and enhanced neuroinflammation, making of SFRP1 an interesting therapeutic target.

While we propose that the phenotype of *Sfrp1^TG^* is largely attributable to the regulation of synaptic remodeling, we acknowledge alternative interpretations. For example, impairment in protein translation might not be a secondary effect of synaptic loss but rather the cause of such a loss, although we have no explanation for how SFRP1 initially impacts translation. An additional interpretation may include increased neuroinflammation, which we have only observed in older mice. Additional studies are thus necessary to comprehensively understand the mechanisms underlying the loss of cognitive ability in aging *Sfrp1^TG^* mice. Our discussion on the role of neurexins touches on a complex area, raising numerous questions, particularly regarding the identification of the involved isoforms. While we have suggested that the observed phenotypes result from the SFRP1-mediated down-regulation of ADAM10 proteolytic activity in synaptosomes^17^, we cannot rule out the possibility that SFRP1 may influence Wnt signaling, despite the absence of evidence for the modulation of this pathway. Additionally, the punctate staining along the neuronal cell body and processes observed *in vitro* suggests potential interactions of SFRP1 with other neuronal proteins. Further studies are therefore needed to elucidate these aspects.

An early decline in dendritic complexity and spine density is commonly observed in many neurodegenerative diseases^79^, though the specific neuronal population affected varies according to the type of neurodegenerative disease. This raises the question of whether increased SFRP1 levels are AD specific and whether its activity is neuron type-selective. We currently lack a clear answer to these questions. However, SFRP1 has not emerged as a differentially expressed protein in five proteomic studies of the substantia nigra from Parkinson’s disease patients^80,81^, nor in the limited studies analysing the brain of individuals with frontotemporal dementia^82–84^. Resolving these issues might contribute to a better understanding of the potential for selecting SFRP1 as an AD therapeutic target and/or additional diagnostic marker.

## Material and methods

### Animals

*Sfrp1^TG^* mice were generated by crossing *hGFAP;tTA* transgenic mice^85^ and *LacZ*;TRE;Sfrp1 mice^33^. Mice carrying only the *LacZ*;TRE;Sfrp1 transgene were used as controls. Males and females were used indistinctly for all experiments, as no sex related differences were observed, with the exception of electrophysiological recordings, for which only males were used. Animals were maintained in the Animal Facility of the Centro de Biología Molecular Severo Ochoa, in a temperature-controlled, pathogen-free environment under 12-12 light-dark cycles. Food and water were available *ad libitum*. All animal procedures were approved by the CBMSO and Comunidad Autónoma de Madrid ethical committees under the following protocol approval number (PROEX 092.6/21).

### Antibodies

The following primary antibodies were used: chicken anti-βgalactosidase (1:2000, abcam, #ab9361), rabbit anti-GFAP (1:1000, DakoCytomation, #z0334), rabbit anti-MAP2 (1:1000, in-house generation; (Sánchez et al., 1998), rabbit anti-Sfrp1 (1:1000, abcam, #ab4193), rabbit anti-Nrnx1 (1:1000, Millipore #abn161), mouse anti-puromycin (1:500, DHSB #PMY-2A4), mouse anti-GluN2b (1:1000, BD #610416), mouse anti-PSD95 (1;1000, Millipore #mab1596), mouse anti-synaptophysin (1:1000, Sigma #SAB4200544), mouse anti-Tubulin (1:1000, Sigma #T5168), mouse anti-GAPDH (1:5000, Santa Cruz Biotechnology #sc-32233), sheep anti-Nrxn3 (1:1000, RnD Systems #AF5269). The corresponding secondary antibodies conjugated with Alexa fluorophores (Thermo Fisher Scientific) were used at a 1:1000 concentration, while secondary antibodies conjugated with peroxidase (Jackson Immune Research) were used at a 1:25.000 dilution.

### Brain extract preparation

Whole brains or dissected hippocampi were homogenized in RIPA buffer (20 mM Tris-HCl pH 7.4, 150 mM NaCl, 1 mM EDTA, 1 mM EGTA, 1% Triton X100, 0.5% deoxycholate, 0.1% SDS) supplemented with protease inhibitors (cOmplete™ Protease Inhibitor Cocktail, Roche) and centrifuged at 21000 g for 30 min at 4°C. The resulting supernatants were used as brain or hippocampal homogenates. Protein concentration was determined using the BCA Protein Assay Kit (Thermo Fisher Scientific) following the manufacturer’s indications.

### Hippocampal synaptosome preparation

Synaptosomes were obtained using the Percoll-based protocol described in^57^. Briefly, mice hippocampi were homogenized in 1 ml sucrose buffer (0.32 M sucrose, 1 mM EDTA, 1 mg/ml bovine serum albumin (BSA), 5 mM HEPES pH 7.4) supplemented with protease inhibitors. Homogenates were centrifuged at 1000 g for 10 minutes at 4°C. The supernatant was further centrifuged for 12 minutes at 14000 g at 4°C. The resulting pellet was resuspended in 300 μl Krebbs-Ringer buffer (140 mM NaCl, 5 mM KCl, 5 mM glucose, 1 mM EDTA, 10 mM HEPES, pH 7.4) and mixed with Percoll (45% v/v). The Percoll mixture was centrifuged for 2 min at 18500 g at 4°C. The synaptosomes were collected from the surface with a syringe, resuspended in Krebs-Ringer buffer and centrifuged at 18500 g for 30 seconds at 4°C. The resulting pellet was resuspended in 100 μl sucrose buffer containing protease inhibitors. Protein concentration was determined with the BCA Protein Assay Kit.

### ELISA

SFRP1 was quantified using a capture ELISA according to a previously described protocol^17^. Briefly, 96-well plates were coated overnight at 4°C with 50µl of anti-SFRP1 (IgG1, clone 10.5.6) diluted in PBS (1.5 µg/ml). Plates were washed 3x with 0.05% Tween20 in PBS (PBST) and blocked with 100 µl of blocking buffer (2% BSA/0.05/PBST) for 3hs at RT. Samples were prepared at 0.1 µg/µl in blocking buffer and incubated for 2hs at 37°C. After washing, plates were incubated with 50 µl of biotin-labelled anti-SFRP1 (IgG2b, clone 17.8.13) diluted in PBS (0.1 µg/ml) for 1h at 37°C. Plates were washed again and incubated with 50 µl Streptavidin-POD at a dilution of 1:25000 in PBS for one hour at 25°C. After extensive washing, the bound protein was visualized using 100 µl of Tetra-methyl-benzidine liquid substrate (TMB slow kinetic form, Sigma-Aldrich) at RT for 20 min. The reaction was terminated by the addition of 100 µl of HCl 2N and the results were measured at 450 nm in a microtiter plate ELISA reader (FLUOstar OPTIMA, BMG LABTECH).

### Immunofluorescence

Mice were perfused transcardially with saline solution (0.9% NaCl) followed by 4% PFA in 0.1M phosphate buffer (PB). Brains were collected and post-fixed by immersion in 4% PFA PB O/N at 4°C. After extensive washing in PBS, brains were incubated in 30% sucrose (weight/volume) diluted in 0.1M PB for 48hr and embedded in a 7.5% gelatin/15% sucrose solution in PB. The embedded brains were frozen in a bath of isopentane at −30°C and stored at −80°C until sectioning using a cryostat (Leica). Immunostainings were performed following standard protocols. Sections were thawed, blocked with 5% BSA/1% FBS/0.1% TritonX-100/PBS and incubated with primary antibodies diluted in blocking buffer O/N at 4°C. Secondary antibodies were incubated for 1h at RT at a dilution of 1:1000 in blocking buffer. Sections were stained with Hoechst (1:2000, Invitrogen, #33342) and mounted on microscope slides. Confocal images were acquired using a Laser Scanning Confocal LSM800 apparatus equipped with an Axio Observer inverted microscope or with a LSM900 apparatus coupled to an upright Axio Imager 2 Microscope (Zeiss). For neuronal cultures immunostaining, primary antibodies were incubated for 2 hrs at RT followed by procedures similar to those used for tissue sections.

### Sindbis virus stereotactic injection

In order to visualize single neurons, sindbis virus carrying GFP were injected into the dentate gyrus (DG) of 2- and 10-month-old control and transgenic mice. Mice were anesthetised with 4% inhaled isoflurane (Forane, Abbvie), which was maintained at 2.5% during the procedure, and placed in the stereotactic apparatus. After its exposure, the scull was drill to allow the injection of the virus using the following coordinates from Bregma: −1.8 mm antero-posterior (A-P), 1.5 mm lateral, −2.2 mm dorso-ventral (D-V)^86^. The viral solution (1 µl) was infused (0.2 µl/min) with a stereotactic injector (Stoelting) connected to a Hamilton syringe (34G needle). After the infusion, the syringe was left in position for 5 min before its retraction. Mice were sacrificed 15 hrs after the injection, perfused and fixed as described above. The brains were collected and processed to obtain 50 µm thick cryostat coronal sections.

### Analysis of dendritic complexity and spine density

Sections containing the hippocampus were stained with Hoechst and visualized in a LSM800 confocal microscope (Zeiss). Z-stacks images of single GFP positive GCs, identified according to their anatomical localization, were captured using a 25X objective to visualize the entire dendritic tree. Images were processed with the Simple Neurite Tracer plugging of the FIJI Software, and morphological complexity was determined by Sholl analysis, in which imaginary circles of increasing radius are drawn from the soma at 10 µm intervals. To visualize dendritic segments, Z-stack images were obtained with a 63X objective with a 3X zoom in a LSM800 confocal microscope, and deconvoluted using the Huygens software (Scientific Volume Imaging). NeuronStudio^87^ was used to measure the number of dendritic spines. Spine density was calculated by dividing the number of spines by the dendritic length.

### Transmission electron microscopy

Control and *Sfrp1^TG-H^* mice were anaesthetized and perfused with saline solution followed by a mixture of PFA (4%) and glutaraldehyde (2%) in 0.1M PB. Brains were post-fixed in the same solution O/N at 4°C and extensively washed with PBS before embedding in 4% agarose, 2% sucrose in 0.1% PB. 200 µm coronal sections were obtained using a vibratome (Leica) and treated with 2% osmium tetroxide in 0.1M PB for 1.5 hrs at RT. After washing with distilled water, the sections were incubated with 2% uranyl acetate in water for 1 h at RT, washed again and dehydrated in grades of ethanol (EtOH 30% 5 min, EtOH 50% 5 min, EtOH 70% 2×10min, EtOH 96% 2×10min, EtOH 100% 3×10min) at RT under agitation. Dehydration was completed with a mixture of ethanol: propylene oxide (PO) 1:1 for 5 min and pure PO 3×10 min. Infiltration of the resin was accomplished with PO:Epon 1:1 for 45 min followed by incubation in pure Epon resin (TAAB 812 Resin, TAAB Laboratories) O/N at RT. Then, the sections were flat embedded between two acetate sheets. Serial ultrathin sections of the DG were obtained using an ultramicrotome (Leica Ultracut UCT) with a diamond blade and collected on slot grids, stained with 2% uranyl acetate and lead citrate and examined with a JEM1400 Flash transmission electron microscope at different magnifications.

### Primary hippocampal cultures

Hippocampal cultures were prepared as described^88^, with minor modifications. Hippocampi were dissected from E18.5 mouse embryos in ice-cold Ca^2+^, Mg^2+^ free Hank’s buffer salt solution (HBSS, Gibco, Life Technologies Co.) and digested with 0.005% trypsin (Trypsin-EDTA 0.05%, Gibco, Life Technologies Co., #25300054), 50 µg/ml DNAseI (Merck, #DN25) in HBSS at 37°C for 15 minutes. The digestion was ended by the addition of plating medium (Minimum Essential Medium supplemented with 10% horse serum and 20% glucose (Thermo Fisher Scientific). The cell suspension was centrifuged at 150 g for 10 min at 37°C. Cells were resuspended from the pellet in plating medium and dissociated by pipetting. Cells were counted using a Neubauer Chamber and seeded on coverslips into multi-well culture dishes (Falcon) pre-treated with 0.1% poly-D-lysine (Sigma-Aldrich, #P7280) at a density of 12.500 cells/cm^2^ in plating medium containing antibiotics (streptomycin/penicillin 100 µg/ml, PanReac). After 3 hrs, the medium was replaced with Neurobasal (NB, Gibco, Thermo Fisher Scientific) supplemented with B27, GlutaMAX (Thermo Fisher Scientific) and antibiotics. Hippocampal cultures were treated with 5µM cytosine arabino-furanoside (Merck, #C1768) 24 hrs after plating to prevent the growth of glial, endothelial and other possible dividing cells. The culture medium was refreshed after 7 days *in vitro* (DIV) and thereafter half of its volume was replaced by fresh medium every 2-3 days.

### Culture treatments

Hippocampal neurons were cultured in the presence or absence of human recombinant SFRP1 (hrSFRP1, 400 ng/ml; RD #5396-SF-025). Coverslips were then fixed with 2% PFA in 0.1 M PB previously equilibrated to 37°C, followed by a 10 min incubation with warm 4% PFA. After fixation, the coverslips were extensively washed with PBS. In the case of short-term incubations, hrSFRP1 was first conjugated with Alexa-488 following the manufacturer’s protocol (DyLight 488 Conjugation Kit (Fast), abcam), and then added to the culture medium (400 ng/ml) for periods variable from 10 to 180 min, when the coverslips were washed and fixed as described above.

### Surface Sensing of Translation assay

To measure protein synthesis in response to SFRP1, hippocampal neuronal cultures were prepared and treated with the recombinant protein for 72 hrs (from DIV14 to DIV17). Then, the cultures were treated with 1 µg/ml puromycin, a structural analogue of transfer RNAs, for 1 hr at 37°C. This compound is incorporated into elongating peptide chains, therefore its detection using specific anti-puromycin antibodies allows for the detection of changes in protein synthesis^63^. After this treatment, neurons were washed with fresh culture medium for 10 minutes and processed for western blot analysis.

### Electrophoresis and Western blot

Protein extracts (20 μg) were diluted in Laemmli Buffer (62,5 mM Tris-HCl pH 6.8, 2% SDS, 10 % glycerol, 10 % β-Mercaptoethanol and 0.005 % bromophenol blue) and boiled for 5 min at 100°C. The samples were loaded into acrylamide/bis-acrylamide gels, electrophoretically resolved within reduced and denaturing conditions and transferred to 0.45 μm pore nitrocellulose membranes by using Mini-Protean system (Bio-Rad), for 90 min at 380 mA. Membranes were washed with Tris buffer saline (TBS, NaCl 150 mM, Tris-HCl 10 mM pH 8) containing 0.1% Tween20 (TBST) and then blocked with 10% non-fat milk in TBST for 1 h at RT. After 3x 5 min washes with TBST, the membranes were incubated O/N at 4°C with the primary antibodies diluted in TBST. After 3x 5 min washing steps, membranes were incubated with peroxidase-conjugated secondary antibodies prepared in TBST for 1 h at RT. Finally, the membranes were further washed 3x with TBST and once with TBS before visualization of the resulting bands with ECL Advanced Western Blotting Detection Kit (GE Healthcare, #RPN2135) in an Amersham Imager 600 (GE Healthcare). Bands were quantified with FIJI.

### In vivo electrophysiological recordings

In vivo electrophysiological recordings were performed in the laboratory of J.M. Delgado-García at the Universidad Pablo de Olavide (Seville). In accordance with habitual procedures from this laboratory^52^, mice were anesthetized with 0.8% isoflurane delivered from a calibrated Fluotec 5 vaporizer (Fluotec-Ohmeda, Tewksbury, MA) at a flow rate of 1-1.2 L/min oxygen and placed in a stereotaxic frame (David Kopf Instruments, Tujunga, CA, USA). Anesthesia was maintained during surgery through an adaptable mouse mask (David Kopf Instruments). A first group of animals was implanted with a) a bipolar stimulating electrode in the right perforant pathway using the following stereotaxic coordinates from Bregma: −3.8 mm antero-posterior (A-P); 2 mm lateral and −1 mm dorso-ventral (D-V); b) two recording electrodes in the molecular layer of the ipsilateral dentate gyrus using the following coordinates: −2.3 mm A-P, 1.5 mm lateral and - 1.75 mm D-V^86^. A second group of animals was implanted with a bipolar stimulating electrode in the right Schaffer collaterals of CA3 pyramidal neurons (−1.5 mm A-P, 2 mm lateral, −1 mm D-V from Bregma) and with two recording electrodes in the ipsilateral CA1 stratum radiatum (−2.2 mm A-P, 1.2 mm lateral, −1 mm D-V). All electrodes were made of Teflon-coated tungsten wire (50 mm, Advent Research Materials Ltd, Eynsham, UK). The final position of the recording electrodes was determined using the field potential depth profile evoked by paired pulses (40 ms intervals) presented at the stimulating projecting pathways. Two bare silver wires (0.1 mm) were affixed to the skull as ground. Wires were connected to a six-pin socket, which was fixed to the skull through two small screws and dental cement as described^52^.

#### Input/output curves (I/O), paired-pulse facilitation and LTP induction

I/O curves were generated with stimulus intensities ranging from 0.02 to 0.4 mA, in steps of 0.02 mA. The stimulus intensity for paired-pulse facilitation was set well below the threshold for evoking a population spike, usually 30–40% of the intensity (mA) necessary for evoking a maximal fEPSP response (Madroñal et al., 2009). Paired pulses were presented at six (10, 20, 40, 100, 200 and 500 ms) inter-pulse intervals. The stimulus intensity for LTP induction was also set at 30–40% of peak fEPSP values. After 15 min of baseline recording (1 stimulus/20 s), each mouse was stimulated with a high-frequency protocol (HFS) consisting of 5 trains of pulses (200 Hz, 100 ms) at a rate of 1/s. This protocol was repeated six times, at intervals of 1 min. Evolution of fEPSPs after the HFS was followed for 60 min at the same stimulation rate. Additional recording sessions (30 min each) were performed for 3/4 days.

#### Data collection, analysis, representations and statistical tests

Rectangular pulses (1V) corresponding to stimulus presentations and recorded fEPSPs were stored digitally on a computer via an analogue/digital converter (CED 1401 Plus, Cambridge, UK) at a sampling frequency of 11-22 kHz and with an amplitude resolution of 12 bits. Electrophysiological data were analysed off-line for quantification of fEPSP amplitudes using commercial programs (Spike 2 and SIGAVG from CED). Up to 5 successive fEPSPs were averaged, determining the mean value of the amplitude. Computed results were processed for statistical analysis using the Signal program (Systat Software, San Jose, CA, USA).

### Acute brain slices electrophysiological recordings

Two months old mice from both sexes were anesthetized under isoflurane (Forane, Abbvie) and quickly decapitated. The brains were removed and immersed in ice-cold Ca^2+^-free dissection solution (10 mM _D_-glucose, 4 mM KCl, 26 mM NaHCO_3_, 233.7 mM sucrose, 5 mM MgCl_2_, and 0.001% (w/v) phenol red as a pH indicator) infused with carbogen (5% CO_2_ and 95% O_2_). Coronal slices (300 µm) were then prepared in the same carbogenated solution by cutting the brain with a vibrotome (Leica, VT1200S) and left in carbon-gassed artificial cerebrospinal fluid (aCSF; 119 mM NaCl, 2.5 mM KCl, 1 mM NaH_2_PO_4_, 26 mM NaHCO_3_, 11 mM glucose, 1.2 mM MgCl_2_, 2.5 mM CaCl_2_, and osmolarity adjusted to 290 mOsm) at 37°C to recover. After 1 hour, they were kept at 25°C until used for recordings. Electrophysiological recordings were performed at 25°C in an immersion chamber under a constant flow of carbogenated aCSF supplemented with picrotoxin (100 µM, Sigma-Aldrich) to block γ-amino-butyric acid type-A receptors. fEPSPs were recorded from the stratum radiatum of CA1 pyramidal neurons after Schaffer collateral fiber stimulation; using glass recording electrodes (0.5 - 1 mOhm) filled with the same aCSF as the extracellular medium. I/O curves were generated with stimulus intensities ranging from 0.002 to 0.25 mA. The stimulus intensity for PPF and LTP was set well below the threshold for evoking a population spike, usually 30–40% of the intensity (mA) necessary for evoking a maximal fEPSP response (Madroñal et al., 2009). Paired pulses were presented at 4 (50, 100, 200 and 400 ms) inter-pulse intervals. Later, after a minimum of 20 min of baseline recording (1 stimulus/15 s), each slice was stimulated with a theta-burst protocol (TBS) consisting of 10 trains of bursts (4 pulses at 100 Hz with a 200 ms interval) that were repeated for 4 cycles with a 20-second intercycle interval. fEPSPs were recorded after the TBS for 60 min at the same stimulation rate. Data acquisition was carried out with MultiClamp 700 A/B amplifiers and pClamp software (Molecular Devices). Data analysis was performed using custom-made Excel (Microsoft) macros.

### Behavioural tests

All behavioural tests were performed in a separate room to reduce noise. Mice were habituated to the room at least one week before the experiments. To avoid the influence of circadian alterations, all tests were performed during light hours (3.00 to 7.00 PM). All training sessions and tests were recorded for subsequent analysis with the AnyMaze software (Stoelting) and revised manually.

#### Open field

Exploratory behaviour and spontaneous activity were assessed by allowing the animals to explore a 40 × 40 × 35 cm open arena for 10 min. The distance travelled, mean speed, mobile time and time spent in the inner region of the arena were determined.

#### *Rotarod* (Ugo Basile)

On the first training session, mice were placed on the apparatus for 1 min at a constant speed (4 rpm). This was repeated 4X with 1h intervals. After 24 hrs, the mice were placed on the rotarod with accelerated speed (4 to 8 rpm in 2 min), 4X at 1h intervals. The test was conducted on the third day allowing the mice to remain on the rotarod with an acceleration of 4 to 40 rpm for 5 min. Each mouse was tested 4X at 1h intervals, automatically recording the latency to fall. The mean value of the 4 tests was used for analysis.

#### Y maze

A “Y” shaped maze was positioned in the testing room, and 3 different visual cues were placed at equal distances from each arm, one per arm. Mice were allowed to explore 2 of the 3 arms of the maze for 7 min during the training session. After 1h, the mice were placed in the maze again with all the arms accessible and allowed to explore for 5 min. The time spent in each arm was measured, and the exploration index was calculated as the percentage of time in the new arm vs the total exploring time.

#### Novel Object Recognition test

In the first training session, each mouse was placed in an open field cage of 40 × 40 × 35 cm for 10 minutes. On day 2, two identical objects were placed in the centre of the arena and mice were allowed to explore for 10 min. After 24 hrs, one object was replaced by a new one and the mice were tested for 10 min. The time exploring each of the objects was quantified to calculate the discrimination index (time interacting with the novel object vs. time exploring both objects).

#### Operant conditioning

Following previous descriptions^55^, mice training took place in 5 Skinner modules (MED Associates, St Albans, Vermont, USA), each one provided with a lever and a food dispenser. Modules were housed within separated sound attenuating boxes, provided with a 45 dB white noise and dimly (19 W lamp) illuminated (Cibertec SA, Madrid, Spain). Training took place for 20 min during 10 successive days, in which mice were allowed to press the lever to receive pellets (Noyes formula P; 45 mg; Sandown Scientific, Hampton, UK) from the feeder using a fixed-ratio (1:1) schedule.

### Quantitative real time PCR

RNA was isolated from hippocampal homogenates from 10 months old control and transgenic mice by using the TRI Reagent protocol (Sigma) following the manufacturer’s instructions. The RNA concentration was determined in a Nanodrop spectrophotometer by 260 nm absorbance. RNA (5 μg) was used to synthesize the first-strand complementary DNA (cDNA) using the First-Strand cDNA synthesis kit (GE Healthcare) with a pd(N)6 primer following the manufacturer’s indications. The quantitative real-time PCR was performed using the GoTaq qPCR Master Mix kit (Promega) following the manufacturer’s guidelines. Each reaction was performed in triplicates using 10 ng of the cDNA, 0.25 μM of the primer mix and 4 μl of the GoTaq master mix. The primers used for cDNA detection are as follows: *Axin2* Fw 5’-GGATTCAGGTCCTTCAAGAGA-3’, Rv 5’-GTGCGTCGCTGGATAACT-3’; *CyclinD1* Fw 5’-TTCCTCTCCAAAATGCCAGA-3’, Rv 5’-TACCATGGAGGGTGGGTTGG-3’; *Myc* Fw 5’-AGACACCGCCCACCACCAGCA-3’, Rv 5’-CGGGATGGAGATGAGCCC-3’; *Gapdh* Fw 5’-AAAATGGTGAAGGTCGGTGTGA-3’, Rv 5’-ATGGGCTTCCCGTTGATGAC-3’; *Hprt* Fw 5’-TCCTCCTCAGACCGCTTTT-3’, Rv 5’-CTAAAGGTGGCCAAGCCCAGCAA-3’. The qPCR was performed in an ABI Prism 7900 Sequence Detection System (Applied Biosystems) using standard parameters. *Gapdh* and *Hprt* were used as housekeeping genes. Data analysis was performed using the ΔΔCt method.

### Proteomic analysis

Hippocampal homogenates and synaptosomes were obtained from control and *Sfrp1^TG^* mice. 1 µg aliquots of each sample were processed for mass spectrometry.

#### MS sample preparation using in-gel Tryptic digestion

Homogenate and synaptosome samples were extracted and reduced in Laemmli sample buffer, containing 10 mM dithiothreitol (DTT), by incubation at 37 °C for 15 minutes in a thermomixer set to 1400 RPM, followed by a second incubation at 98 °C for 5 minutes. Free sulfhydryl groups were alkylated by incubation with 20 mM iodoacetamide (IAA) for 30 minutes at RT. Next, 8 µg of protein was loaded onto a 10% SurePAGE polyacrylamide gel (Genscript) and resolved for 1 cm. The gels were fixed overnight and stained with colloidal Coomassie Blue G-250. Sample lanes were cut into 1 mm^3^ squares, transferred to a MultiScreen HV 96-well filter plate (Millipore) and unstained until clear using repeated applications of 50 mm NH_3_HCO_3_ in 50% acetonitrile. After dehydration with 100% acetonitrile, each well was supplemented with 0.32µg MS grade Trypsin/Lys-C (Promega) in 50 mM NH_4_HCO_3_ and incubated overnight at 37 °C within a humidified incubator. Tryptic peptides were extracted and pooled by two incubations with 0.1% trifluoroacetic acid in 50% acetonitrile, dried by SpeedVac and stored at −80 °C.

#### LC-MS analysis

Each sample of tryptic digest was dissolved in 0.1% formic acid and the peptide concentration was determined by tryptophan-fluorescence assay^91^; 75 ng of peptide was loaded onto an Evotip Pure (Evosep). Peptide samples were separated by standardized 30 samples per day method on the Evosep One liquid chromatography system, using a 15 cm × 150 μm reverse-phase column packed with 1.9 µm C18-beads (EV1106 from Evosep) connected to a 20 µm ID ZDV emitter (Bruker Daltonics). Peptides were electro-sprayed into the timsTOF Pro 2 mass spectrometer (Bruker Daltonics) equipped with CaptiveSpray source and measured with the following settings: Scan range 100-1700 m/z, ion mobility 0.6 to 1.6 Vs/cm2, ramp time 100 ms, accumulation time 100 ms, and collision energy decreasing linearly with inverse ion mobility from 59 eV at 1.6 Vs/cm2 to 20 eV at 0.6 Vs/cm2. Operating in dia-PASEF mode, each cycle took 1.8 s and consisted of 1 MS1 full scan and 16 dia-PASEF scans. Each dia-PASEF scan contained two isolation windows, in total covering 400-1201 m/z (1 Th window overlap) and ion mobility 0.6 to 1.43 Vs/cm2. Ion mobility was auto calibrated at the start of each sample (calibrant m/z, 1/K0: 622.029, 0.992 Vs/cm2; 922.010, 1.199 Vs/cm2; 1221.991, 1.393 Vs/cm2).

#### MS data analysis

DIA-PASEF raw data were processed with DIA-NN 1.8^92^. An in-silico spectral library was generated from the uniprot mouse proteome (SwissProt and TrEMBL, canonical and additional isoforms, release 2022-03) using Trypsin/P digestion and at most 1 missed cleavage. Fixed modification was set to carbamidomethylation (C) and variable modifications were oxidation(M) and N-term M excision (at most 1 per peptide). Peptide length was set to 7-30, precursor charge range was set to 2-4 and precursor m/z was limited to 380– 1220. Both MS1 and MS2 mass accuracy were set to 10 ppm, scan window was set to 12, double-pass-mode and match-between-runs were enabled. Protein identifiers (isoforms) were used for protein inference. All other settings were left as default. MS-DAP 1.0.6^59^ was used for downstream analyses of the DIA-NN results. Filtering and normalization were applied to respective samples per statistical contrast. Peptide-level filtering was configured to retain only peptides that were confidently identified in at least 75% of samples per sample group. Peptide abundance values were normalized using the VSN algorithm, followed by protein-level mode-between normalization. One homogenate control sample was identified as outlier in the quality control analyses presented in the MS-DAP report, which we subsequently excluded from statistical analyses. Differential expression analysis was performed by the MSqRob algorithm and resulting p-values were adjusted for multiple testing using the Benjamini-Hochberg False Discovery Rate (FDR) procedure.

### Statistical analysis

Statistical analysis was performed using Prism v9 software (GraphPad), except for electrophysiological recordings, which were analysed using the Signal program (Systat Software, San Jose, CA, USA). Data distribution was evaluated with Shapiro-Wilk normality test. The statistical tests used to analyse each dataset are indicated in the legend of the figures. Statistical significance was set at p<0.05, significant differences are indicated in the figures.

## Acknowledgements

We thank José M. González Martín and María Sánchez Enciso for their help with electrophysiological recording, and the CBM Advanced Light Microscopy and Electron Microscopy facilities for their support and advice during the course of this study.This work was supported by grants from the Spanish AEI (BFU2013-43213-P; BFU2016-75412-R with FEDER support and PID2019-104186RB-I00), Cure Alzheimer Fund, Fundación Tatiana Perez de Guzman el Bueno and CIBERER to PB and by grants PID2021-122446NB-I00 funded by MCIN/AEI/ 10.13039/501100011033 and “ERDF A way of making Europe” and by Spanish Junta de Andalucía Bio122 to A.G. and J.M.D.-G. FPI fellowships from the AEI supported the work of GP (BES-2017-080318), MIM (BES-2014-068797) and JRC (BES-2011-047189). We also acknowledge a CBM Institutional grant from the Fundación Ramon Areces. The CBM is a Center of Excellence Severo Ochoa CEX2021-001154-S funded by MCIN/AEI/10.13039/501100011033.

## Author contributions

GP, MIM, JRC, PE and PB set up the research project and designed the experiments. GP, MIM and MJMB conducted the experiments and acquired the data, with the following exceptions: AG carried out the experiments and analyzed the data in Fig. 5, Fig. S4 and Fig. S7; RK carried out the LC-MS analysis and quality control assessment of the data in Fig. 7 and Fig 8 under the supervision of ABS; and PM performed the experiments and analyzed the data in Fig. S5 with the help of AFR and ELM under the supervision of JAE. GP, MIM and PB interpreted the data. GP assembled the figures. GP and PB wrote the manuscript. ABS, JMDG and JAE provided invaluable conceptual advice. PB obtained financial support and provided supervision.

## Data availability statement

The data supporting the findings of this study are available within the paper and its Supplementary Information. The data related to proteomic analysis are being deposited in PIRIDE (https://www.ebi.ac.uk/pride/) and will be available with the following accession number: XXX

