## Supplementary figures for "SFRP1 upregulation causes hippocampal synaptic dysfunction and memory impairment"

**Figure S1**

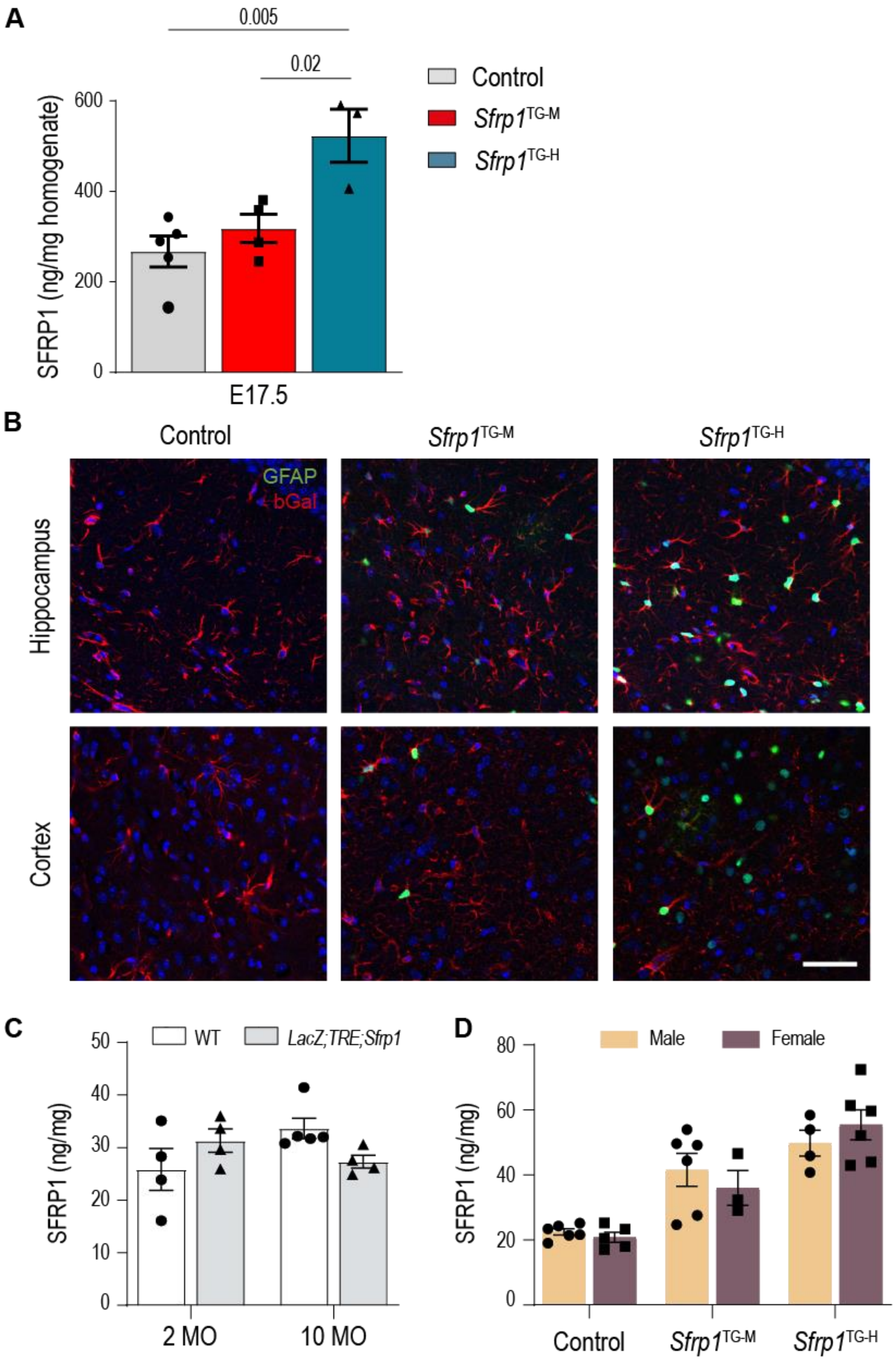


**Figure S1. Characterization of SFRP1 expression in *Sfrp1*$^{TG}$ mice**. **A)** ELISA determination of SFRP1 protein levels in whole brain homogenates from E17.5 control (n=5), *Sfrp1*$^{TG\text{-}M}$ (n=4) and *Sfrp1*$^{TG\text{-}H}$ (n=3) brains. The bars represent mean ± SEM. Statistical significance was evaluated with one-way ANOVA followed by Bonferroni post-hoc analysis. **B)** Representative confocal images of hippocampal (top) and cortical (bottom) coronal sections from control and *Sfrp1*$^{TG}$ mouse brains stained for GFAP (red) and β-galactosidase (green). Scale bar 50 µm. **C**) ELISA determination of SFRP1 protein levels in whole brain homogenates from two and ten months old wt (n=4) and *LacZ*;TRE;*Sfrp1* (n=4) mice. The bars represent mean ± SEM. Statistical significance was evaluated with two-way ANOVA followed by Bonferroni post-hoc analysis. **D)** ELISA determination of SFRP1 protein levels in whole brain homogenates from male and female control (n=6; n=5) *Sfrp*$^{TG\text{-}M}$ (n=6; n=3) and *Sfrp*$^{TG\text{-}H}$ (n=4; n=6) mice. Graph bars represent mean ± SEM. Statistical significance was evaluated by two-way ANOVA followed by Bonferroni multiple comparisons test.

### Figure S2

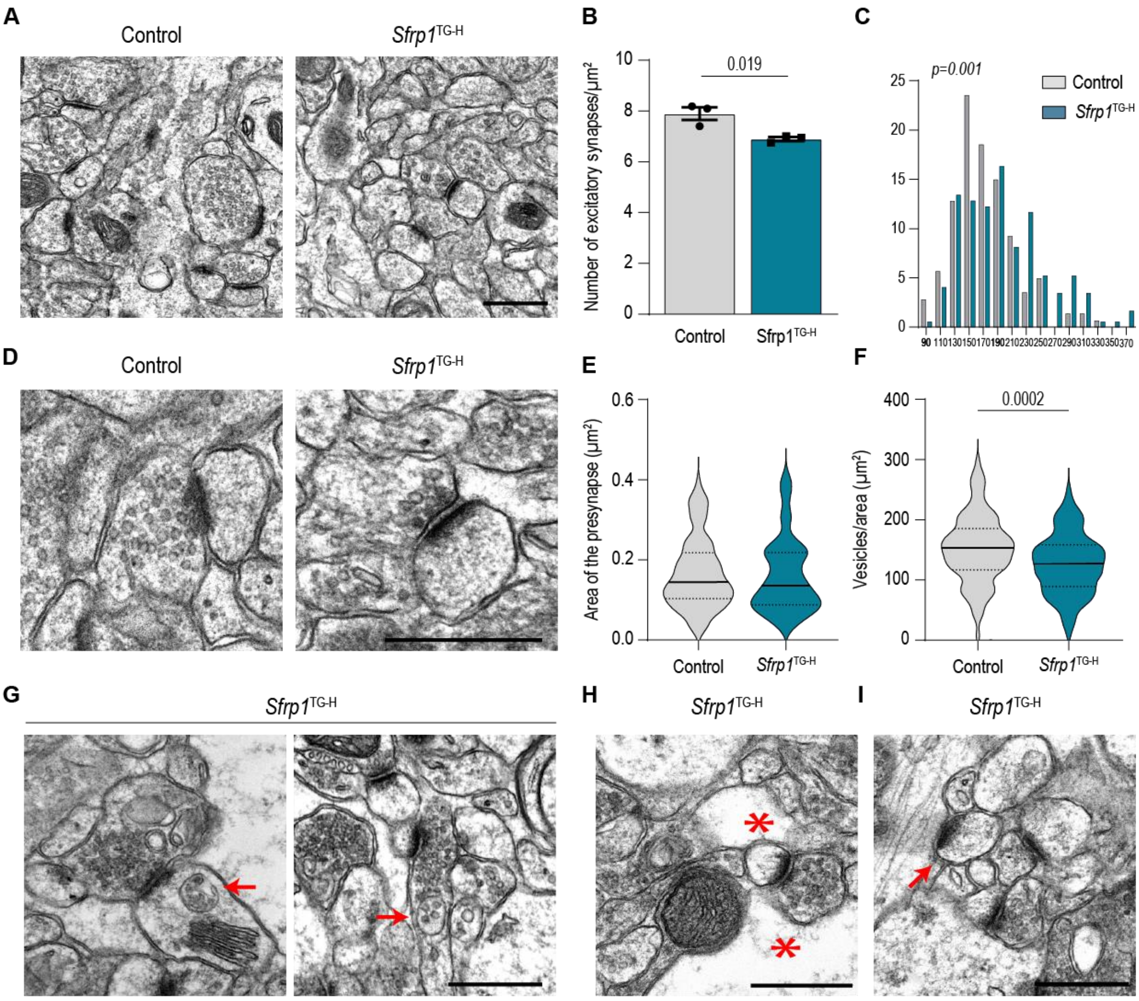


**Figure S2. Ultrastructural synaptic alterations in GCs of young *Sfrp1*$^{TG-H}$ mice. A, D)** Representative low (A) and high power (D) electron micrographs showing asymmetric synapses in the molecular layer of the DG in two months old control and *Sfrp1*$^{TG-H}$ mice. Scale bars 0.5 µm. **B**) Quantification of the density of asymmetric synapses in control (n=3) and *Sfrp1*$^{TG-H}$ (n=3) mice. The graph represents mean ± SEM. Statistical significance was determined with two-tailed Student's t-test. **C**) Frequency distribution of the PSD length in control (n=140 synapses; 3 mice) and *Sfrp1*$^{TG-H}$ (n=172 synapses; 3 mice). Statistical significance was calculated with Kolmogorov-Smirnov test. **E, F**) Quantification of the presynaptic area (E) and synaptic vesicle content (F) in asymmetric synapses from control (n=110) and *Sfrp1*$^{TG-H}$ (n=117) mice. The violin plots represent data distribution, median (solid line) and 25% and 75% quartiles (dotted lines). Statistical significance was calculated with two-tailed Mann-Whitney test in (E) and two-tailed Student's t-test in (F). **G-I**) Representative examples of other structural alterations observed in two- and ten-months old *Sfrp1*$^{TG-H}$ GCs: accumulation of multi-vesicular bodies in pre- and postsynaptic compartments (G), enlarged glial cells (H) and isolated postsynaptic compartments (I).

**Figure S3**

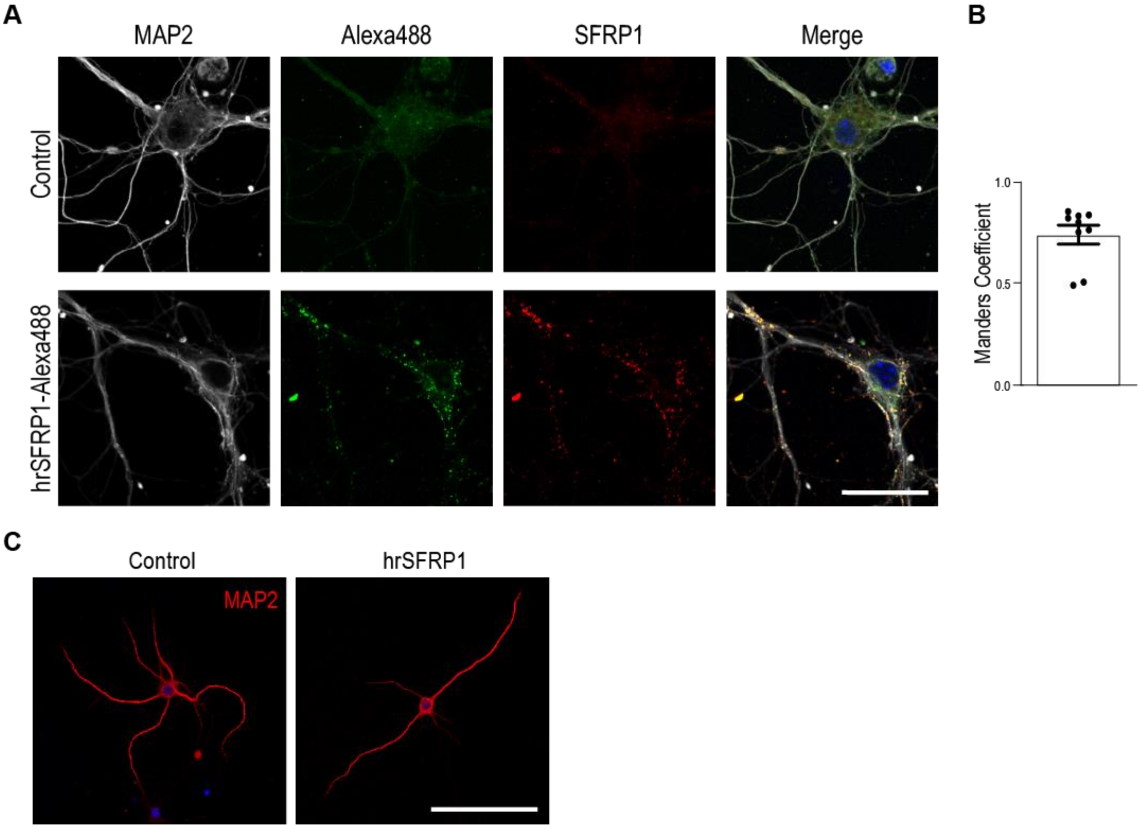


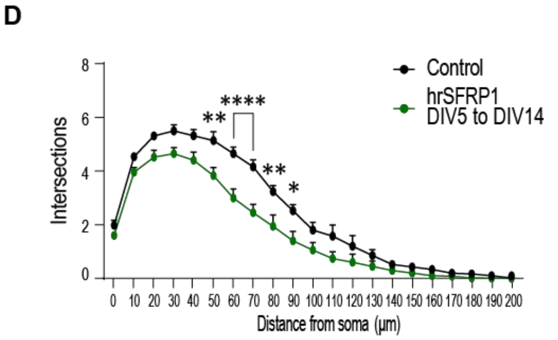


| Exact p-values from the statistical analysis of the experiment in FigS3D | | | | | |
|---|---|---|---|---|---|
| Test: Two-way ANOVA | | | | | |
| Multiple comparisons test: Bonferroni's | | | | | |
| P values: | | | | | |
| Distance from soma (μm) | p-value | Distance from soma (μm) | p-value | Distance from soma (μm) | p-value |
| 0 | >0,9999 | 70 | <0,0001 | 140 | >0,9999 |
| 10 | >0,9999 | 80 | 0,0017 | 150 | >0,9999 |
| 20 | 0,3037 | 90 | 0,0133 | 160 | >0,9999 |
| 30 | 0,1973 | 100 | 0,4601 | 170 | >0,9999 |
| 40 | 0,1085 | 110 | 0,2156 | 180 | >0,9999 |
| 50 | 0,0016 | 120 | >0,9999 | 190 | >0,9999 |
| 60 | <0,0001 | 130 | >0,9999 | 200 | >0,9999 |

**Figure S3. hrSFRP1 colocalizes with MAP2 and Sfrp1. A**) Representative confocal images of wt hippocampal neurons treated with 400 ng/ml hrSFRP1-Alexa488 conjugate (bottom line) or with Alexa488 (top line) for 3 hrs and immunostained for MAP2 (grey) and SFRP1 (red). Scale bar 25 μm. **B)** Quantification of the colocalization between hrSFRP1-Alexa488 and anti-SFRP1 signals. **C**) Representative confocal images of wt hippocampal neurons cultured in the presence (right) or absence (left) of hrSFRP1 (400 ng/ml, DIV5-DIV14) and immunostained for MAP2. Scale bar 100 μm. **D**) Sholl analysis of neuronal complexity in the presence or absence of hrSFRP1. The dots represent mean + SEM. N = 5 independent cultures. Statistical significance was evaluated by one-way ANOVA followed by Bonferroni multiple comparisons test. The exact p-values are indicated in the table in the right. *p < 0.05, **p<0.01, ****p<0.0001.

**Figure S4**

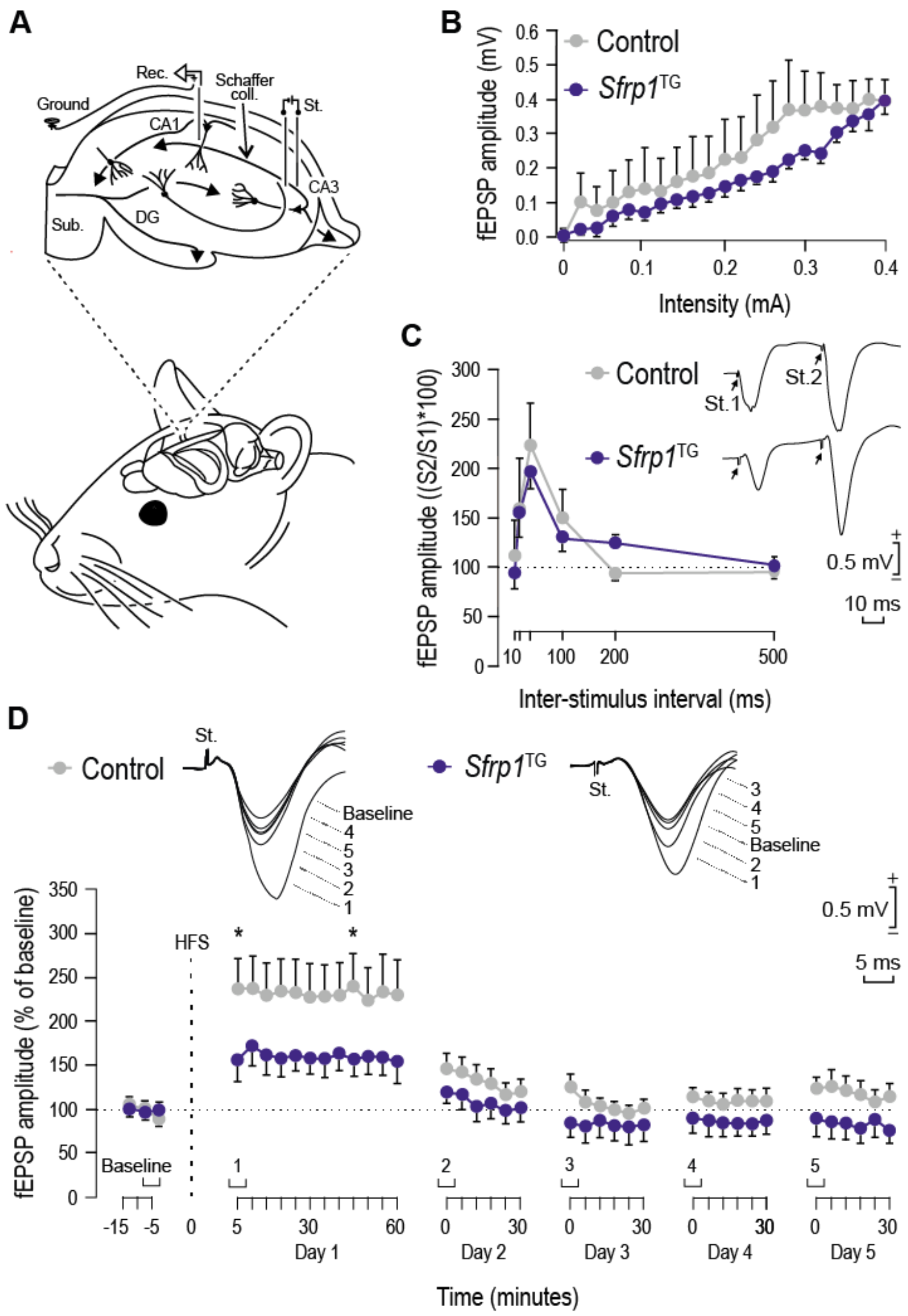


**Figure S4**. **The LTP response of CA3-CA1 synapses is decreased in *Sfrp1$^{TG}$* mice**. **A)** Mice were implanted with a bipolar stimulating and a recording electrode in the hippocampal CA3 Schaffer collaterals and ipsilateral CA1 area, respectively. **B)** Input/output curves recorded in CA1 for control (n=3) and *Sfrp1$^{TG}$* (n=6) mice. Data are presented as mean ± SEM and analyzed with two-way ANOVA followed by the Holm-Sidak method. **C)** Double pulse facilitation evoked in control and *Sfrp1$^{TG}$* mice. Representative examples (averaged five times) of fEPSPs evoked by paired pulses at 40 ms of inter-pulse interval in control (n=3) and *Sfrp1*$^{TG}$ (n=6) mice are illustrated on the right side of the graph. Data are mean ± SEM amplitudes of the second fEPSP expressed as the percentage of the first for six inter-pulse intervals. Data were analyzed with two-way ANOVA followed by the Holm-Sidak method. **D)** LTP evoked by HFS in CA3-CA1 synapses in ten-months-old control (n=12) and *Sfrp1*$^{TG}$ (n=10) mice. LTP evolution was followed for a total of 5 days. Illustrative examples (averaged five times) of fEPSPs recorded each day are presented in the right. Data are mean ± SEM amplitudes. Two-way ANOVA followed by the Holm-Sidak method was used to evaluate statistical significance. *$p < 0.05$.

**Figure S5**

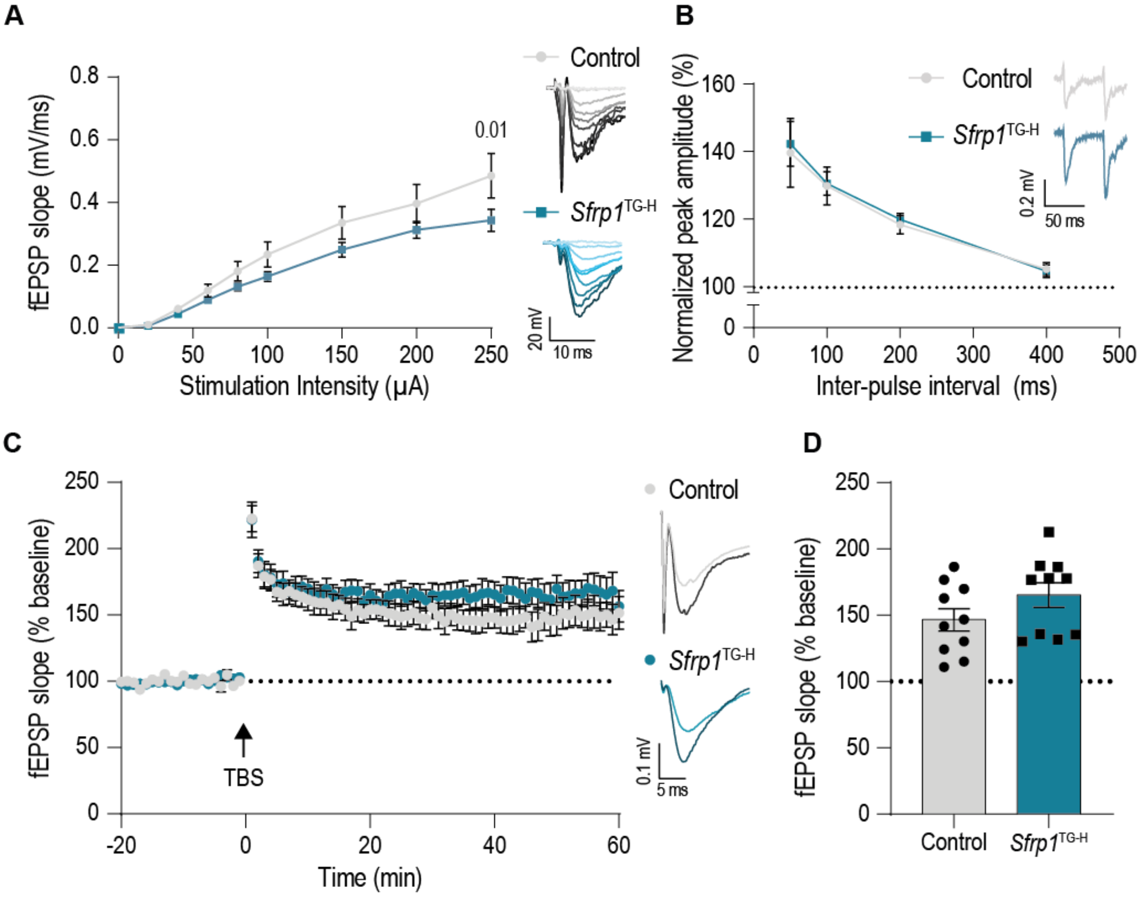


**Figure S5. Sfrp1$^{TG\text{-}H}$ mice show reduced basal transmission and unaltered synaptic plasticity at 2 months of age in the CA3-CA1 pathway of hippocampal acute slices. A)** Input/output curves of the extracellular fEPSP recorded at different intensities (0 – 250 µA) in control (n = 12 slices from 7 mice) and *Sfrp1$^{TG\text{-}H}$* (n = 13 slices from 7 mice) mice. Data are shown as mean ± SEM of the fEPSP slope and analyzed using two-way ANOVA followed by Hold-Sidak correction for multiple comparisons. Representative images from each group at each stimulation intensity are depicted on the right. **B)** Average values recorded from a double pulse facilitation protocol at intervals from 50 – 400 ms in control (n = 9 slices from 6 mice) and Sfrp1$^{TG\text{-}H}$ (n = 9 slices from 6 mice) mice. Representative traces of the 50 ms paired pulse interval are depicted on the top right. Data are shown as the mean amplitude percentage ± SEM of the second evoked response normalized to the first. Statistical significance was evaluated with two-way ANOVA. **C)** LTP time course induced by TBS in control (n = 8 slices from 5 mice) and Sfrp1$^{TG\text{-}H}$ (n = 9 slices from 5 mice) mice represented as mean percentage over baseline ± SEM. Representative traces are depicted on the right of the graph, showing the basal and potentiated responses of both groups. **D)** Average fEPSP slopes from the last 5 minutes of the LTP recordings shown in C). Two-tailed unpaired Student's t-test was used for statistical analysis. *p<0.05.

**Figure S6**

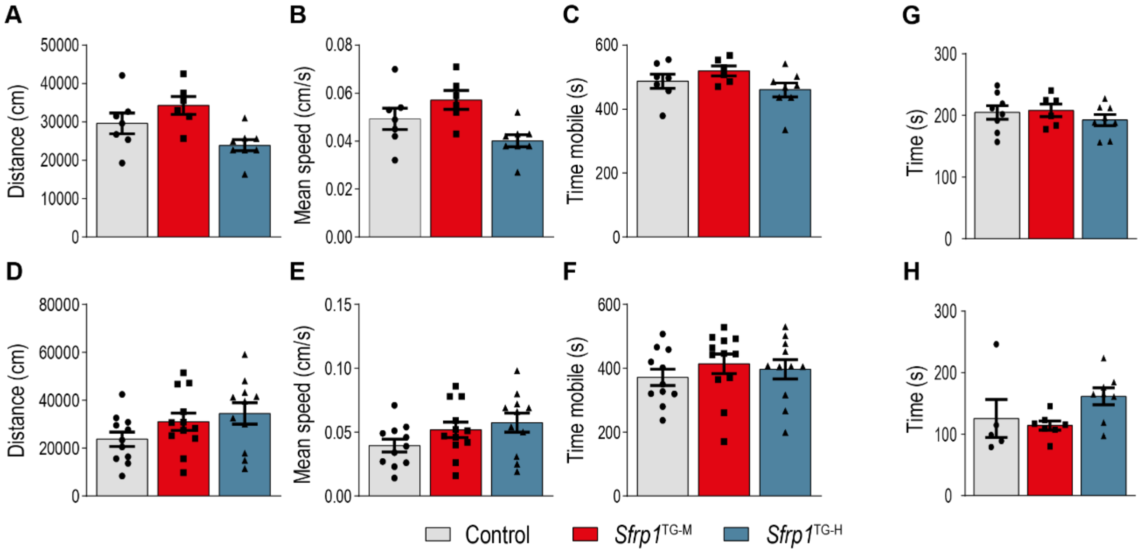


**Figure S6. *Sfrp1*$^{TG}$ mice present a normal locomotor activity**. **A-C; D-F)** The graphs show the total distance travelled (A, D), the mean speed (B, E) and the total time mobile (C, F) in two (A-C) and ten (D-F) months old control (n=7; n=11), *Sfrp1*$^{TG-M}$ (n=6; n=12) and *Sfrp1*$^{TG-H}$ (n=8; n=11) mice evaluated in the open field. No difference was observed between genotypes. **G, H)** The graphs show the time spent by two (G) and ten (H) months old control (n=8, n=5), *Sfrp1*$^{TG-M}$ (n=6, n=7) and *Sfrp1*$^{TG-H}$ (n=8, n=8) mice on the rotarod. Motor coordination was comparable between genotypes. The bars represent mean ± SEM. Data were analyzed by one-way ANOVA followed by Bonferroni post-hoc analysis.

**Figure S7**

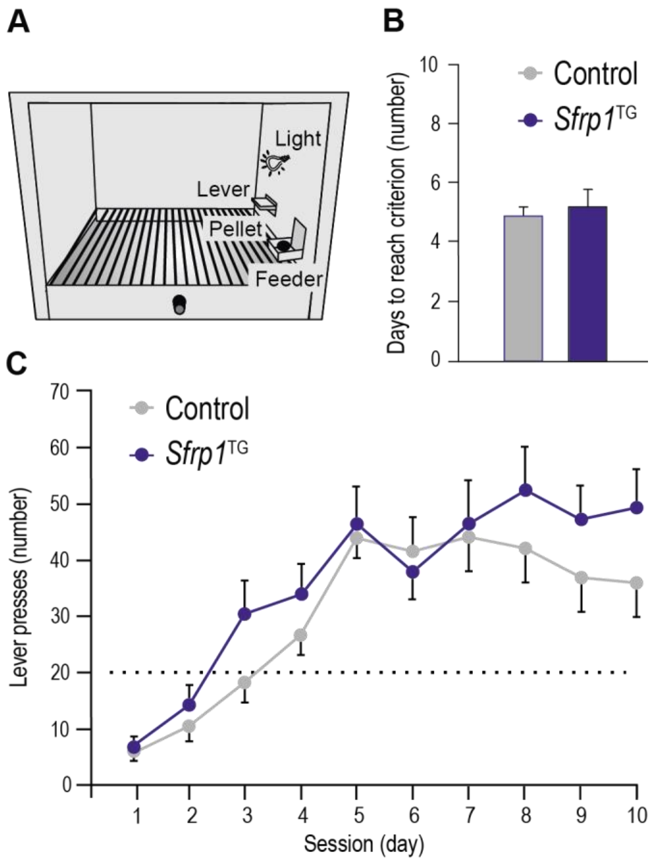


**Figure S7. Associative learning performance is not altered in *Sfrp1*$^{TG}$ mice**. **A)** Schematic representation of the Skinner box. **B)** The graph represents the number of days needed for mice to reach the selected criterion: to press the lever ≥ 20 times for two successive sessions. No difference between control (n=13) and *Sfrp1*$^{TG}$ (n=15) mice was observed. The bars represent mean + SEM. Statistical significance was calculated by two-tailed Student's t-test. **C)** Evolution of the number of lever presses during the successive training sessions. The dots represent mean ± SEM and were analyzed by two-way ANOVA followed by the Holm-Sidak method.

**Figure S8**

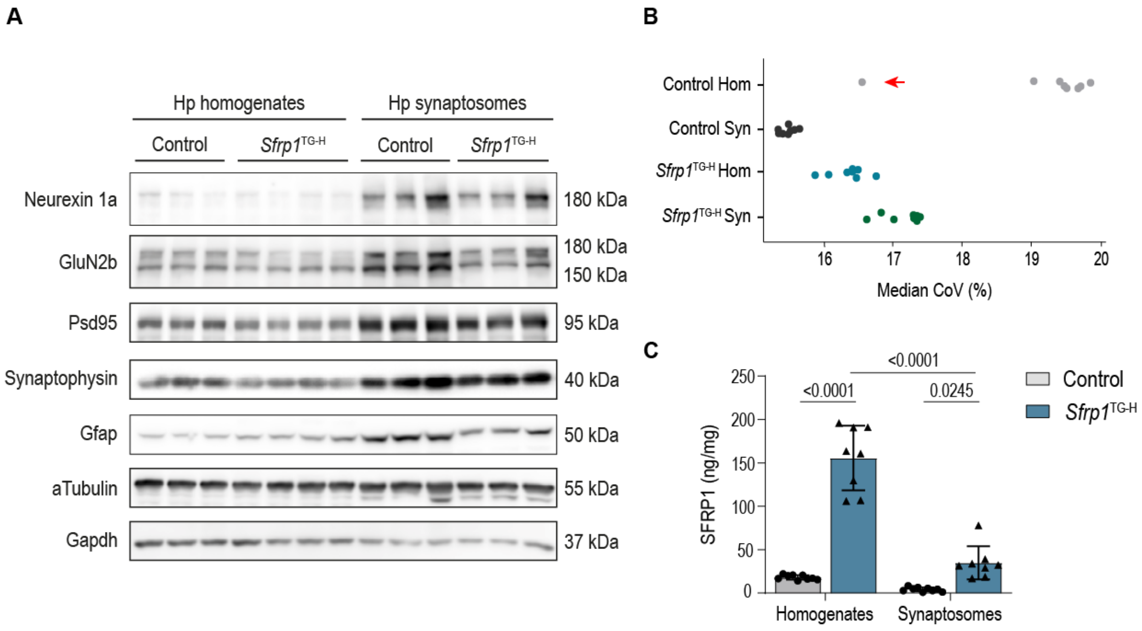


**Figure S8**. **Quality control assessment of hippocampal homogenates and synaptosomes used in proteomic experiments**. **A**) Hippocampal homogenates and synaptosomes obtained from 10-month-old control and *Sfrp1*$^{TG\text{-}H}$ mice were subjected to Western blot and probed with antibodies against synaptic proteins to corroborate their enrichment in the synaptosomal fractions. Alpha-tubulin and Gapdh were used as loading controls. **B**) Coefficient of variation (CoV) analysis determines how much the CoV of an experimental group improves by removing a single sample. This analysis confirmed the presence of an outlier among the control hippocampal homogenates, which was therefore eliminated from further analysis. **C**) ELISA determination of SFRP1 protein levels in control and *Sfrp1*$^{TG\text{-}H}$ hippocampal homogenates and synaptosomes (n=8). Note that SFRP1 is almost absent in control synaptosomes. Data are mean ± SEM. Statistical significance was calculated with two-way ANOVA and Bonferroni multiple comparisons test.

**Figure S9**

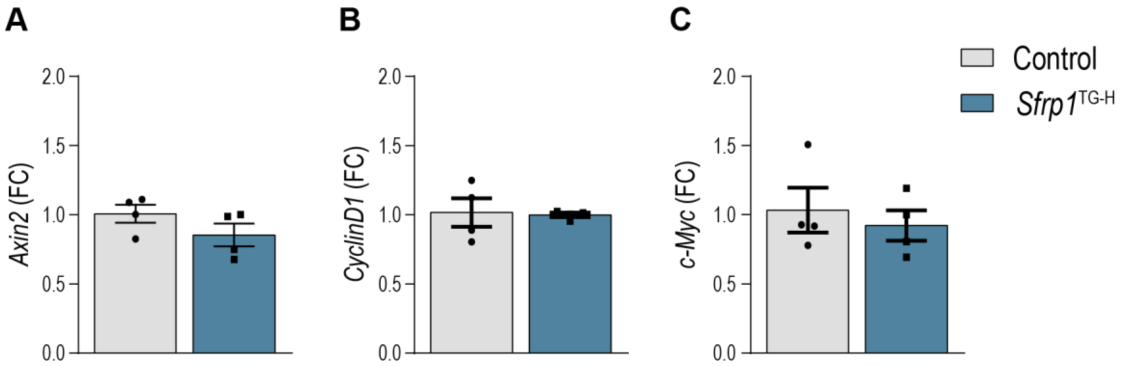


**Figure S9. SFRP1 upregulation does not modify the expression of canonical Wnt target genes.** qPCR determination of the relative expression of *Axin2* (A), *CyclinD1* (B) and *c-Myc* (C) in hippocampal homogenates from ten months old control (n=4) and *Sfrp1*$^{TG\text{-}H}$ (n=4) mice. Data are represented as fold change (FC) over the control group, mean ± SEM are shown in the graphs. Statistical significance was evaluated with two-tailed Student's t-test.

**Figure S10**

**Uncroped Western blots - Figure 4**

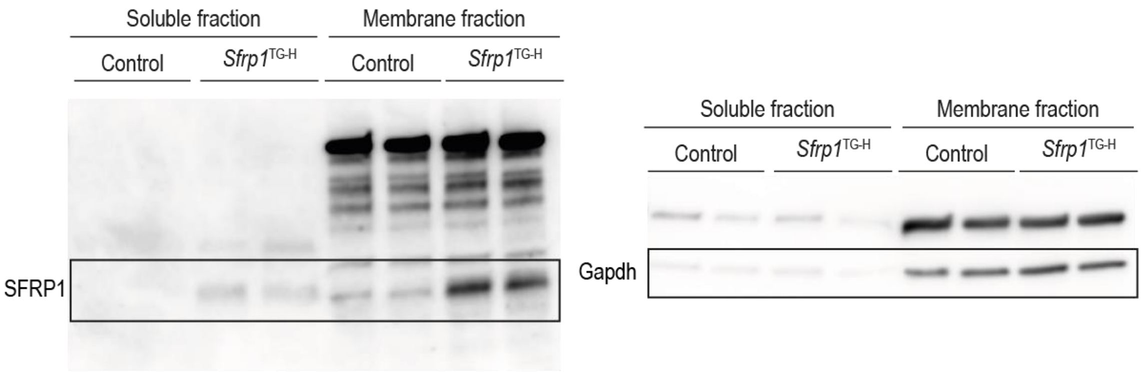


**Uncroped Western blots - Figure 8**

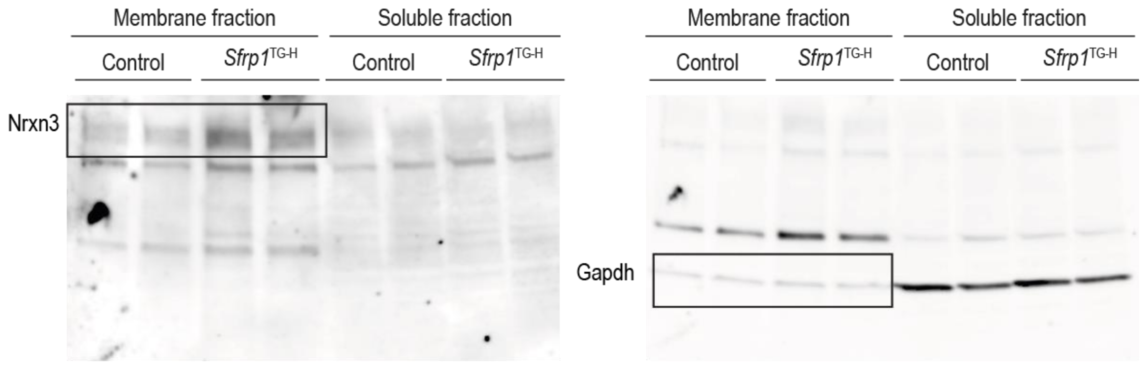


**Figure S10. Uncropped Western blots membranes from figures 4D and 8D**

**Table S1. Statistical comparisons from the experiments in Figure 5.**

**Table S2. List of all proteins detected in at least one sample from each experimental group.**

**Table S3. List of the differentially abundant proteins** (adjusted p-value < 0.05) and non-adjusted differentially abundant proteins (p-value < 0.05) found in hippocampal homogenates and synaptosomes.
