## Supplementary material for "SFRP1 upregulation causes hippocampal synaptic dysfunction and memory impairment": Suppl Table 1a

### Supplementary Figure 4

| | Normal Distribution | Test | Multiple comparisons | | p-val |
|---|---|---|---|---|---|
| S4B | | Two-way ANOVA + Holm-Sidak | Control vs. Sfrp1$^{TG}$ after each input (mA) | 0,02 | 0,9971 |
| | | | | 0,04 | 0,9972 |
| | | | | 0,06 | 0,9972 |
| | | | | 0,08 | 0,9972 |
| | | | | 0,1 | 0,9972 |
| | | | | 0,12 | 0,9972 |
| | | | | 0,14 | 0,9972 |
| | | | | 0,16 | 0,9972 |
| | | | | 0,18 | 0,9972 |
| | | | | 0,2 | 0,9971 |
| | | | | 0,22 | 0,9972 |
| | | | | 0,24 | 0,9919 |
| | | | | 0,26 | 0,8656 |
| | | | | 0,28 | 0,9219 |
| | | | | 0,3 | 0,9919 |
| | | | | 0,32 | 0,989 |
| | | | | 0,34 | 0,9972 |
| | | | | 0,36 | 0,9972 |
| | | | | 0,38 | 0,9972 |
| | | | | 0,4 | 0,9972 |
| S4C | Yes | Two-way ANOVA + Holm-Sidak | Control vs. Sfrp1$^{TG}$ at each time interval | 10 ms | 0,9919 |
| | | | | 20 ms | 0,9919 |
| | | | | 40 ms | 0,9919 |
| | | | | 100 ms | 0,9919 |
| | | | | 200 ms | 0,9695 |
| | | | | 500 ms | 0,9919 |
| S4D | Yes | Two-way ANOVA + Holm-Sidak | Control vs. Sfrp1$^{TG}$ baseline | -15 | 0,995 |
| | | | | -10 | 0,995 |
| | | | | -5 | 0,995 |
| | | | Control vs. Sfrp1$^{TG}$ Day 1 (min) | 5 | 0,0482 |
| | | | | 10 | 0,1748 |
| | | | | 15 | 0,1433 |
| | | | | 20 | 0,0751 |
| | | | | 25 | 0,1426 |
| | | | | 30 | 0,1433 |
| | | | | 35 | 0,1433 |
| | | | | 40 | 0,182 |
| | | | | 45 | 0,0375 |
| | | | | 50 | 0,2409 |
| | | | | 55 | 0,087 |
| | | | | 60 | 0,0973 |
| | | | Control vs. Sfrp1$^{TG}$ Day 2 (min) | 5 | 0,9892 |
| | | | | 10 | 0,9825 |
| | | | | 15 | 0,9675 |
| | | | | 20 | 0,9888 |
| | | | | 25 | 0,9892 |

| | | | | | |
|---|---|---|---|---|---|
| | | | | 30 | 0,9892 |
| | | | Control vs. Sfrp1$^{TG}$ Day 3 (min) | 5 | 0,9234 |
| | | | | 10 | 0,9825 |
| | | | | 15 | 0,9892 |
| | | | | 20 | 0,9892 |
| | | | | 25 | 0,9892 |
| | | | | 30 | 0,9892 |
| | | | Control vs. Sfrp1$^{TG}$ Day 4 (min) | 5 | 0,9892 |
| | | | | 10 | 0,9892 |
| | | | | 15 | 0,9892 |
| | | | | 20 | 0,9892 |
| | | | | 25 | 0,9892 |
| | | | | 30 | 0,9892 |
