## Supplementary material for "SFRP1 upregulation causes hippocampal synaptic dysfunction and memory impairment": Suppl Table 1b

| gene_symbols_or_id | unique_peptides | peptides_used_for_dea_contrast: Control_Hom vs SFRP1-TG_Hom | foldchange.log2_msqrob_contrast: Control_Hom vs SFRP1-TG_Hom | pvalue_msqrob_contrast: Control_Hom vs SFRP1-TG_Hom | qvalue_msqrob_contrast: Control_Hom vs SFRP1-TG_Hom |
|---|---|---|---|---|---|
| DMXL2 | 123 | 122 | 0,129985557 | 1,88E-07 | 0,000694635 |
| SLC9A3R1 | 19 | 18 | -0,135057196 | 1,69999E-07 | 0,000694635 |
| EIF4G3 | 24 | 23 | -0,181734975 | 4,40698E-07 | 0,001084998 |
| ACSBG1 | 34 | 32 | -0,132802494 | 1,12114E-06 | 0,001192664 |
| WDR7 | 66 | 65 | 0,112129979 | 1,13033E-06 | 0,001192664 |
| DBNL | 23 | 23 | -0,107386728 | 9,20926E-07 | 0,001192664 |
| THBS4 | 6 | 6 | -0,444103189 | 7,12346E-07 | 0,001192664 |
| FAM169A | 12 | 12 | -0,189816188 | 2,07576E-06 | 0,001916448 |
| CNN3 | 10 | 10 | -0,164339765 | 7,2005E-06 | 0,005909213 |
| SFRP1 | 2 | 1 | 1,858842808 | 1,30824E-05 | 0,009662644 |
| EIF3B | 22 | 21 | -0,110162557 | 1,64564E-05 | 0,01104972 |
| SPARCL1 | 15 | 15 | -0,205997224 | 2,14718E-05 | 0,012855979 |
| TBC1D15 | 8 | 8 | -0,199057316 | 2,5467E-05 | 0,012855979 |
| ZC3HC1 | 2 | 2 | -0,475438574 | 2,61088E-05 | 0,012855979 |
| LZIC | 7 | 7 | -0,203377085 | 2,39805E-05 | 0,012855979 |
| NSF | 72 | 71 | 0,074253555 | 3,94603E-05 | 0,01619188 |
| EPB41L2 | 37 | 37 | -0,110891757 | 3,86683E-05 | 0,01619188 |
| SEC31A | 30 | 30 | -0,141027258 | 3,84678E-05 | 0,01619188 |
| OSCP1 | 9 | 9 | -0,181384841 | 5,01913E-05 | 0,019511218 |
| HDGFL2 | 8 | 8 | -0,174903698 | 5,56828E-05 | 0,020563671 |
| EIF3E | 26 | 25 | -0,111069293 | 6,7411E-05 | 0,023709428 |
| SF1 | 6 | 6 | -0,232840555 | 7,11414E-05 | 0,023884113 |
| UFL1 | 5 | 4 | -0,187064539 | 8,41431E-05 | 0,027020908 |
| CPNE4 | 18 | 18 | 0,148309706 | 8,85625E-05 | 0,027255103 |
| DLGAP3 | 21 | 21 | 0,13076178 | 0,000109795 | 0,032437689 |
| ERC2 | 45 | 44 | 0,093989389 | 0,000137828 | 0,039153864 |
| SEPHS1 | 4 | 4 | -0,238767248 | 0,00015579 | 0,042617174 |

| | | | | | |
|---|---|---|---|---|---|
| CDC37L1 | 3 | 3 | -0,230455695 | 0,000184994 | 0,045545438 |
| HOOK3 | 21 | 21 | -0,150951961 | 0,000180444 | 0,045545438 |
| COQ7 | 6 | 5 | 0,244015724 | 0,000183724 | 0,045545438 |
| OMG | 10 | 10 | -0,145886449 | 0,000193232 | 0,046038985 |
| RPLP0 | 13 | 13 | -0,135486411 | 0,000209149 | 0,048274134 |
| PLXNB1 | 28 | 27 | -0,093349281 | 0,000264694 | 0,059243226 |
| DOCK9 | 2 | 1 | -1,770078003 | 0,000276304 | 0,060023004 |
| DOCK10 | 11 | 11 | -0,130906229 | 0,000295107 | 0,062276031 |
| GJA1 | 19 | 18 | -0,173925508 | 0,000310246 | 0,063652133 |
| EWSR1 | 5 | 5 | -0,269993087 | 0,00036771 | 0,067128592 |
| STRN4 | 16 | 16 | -0,101025874 | 0,000348823 | 0,067128592 |
| DTNA | 9 | 9 | -0,159251518 | 0,000372634 | 0,067128592 |
| CLU | 16 | 16 | -0,141652818 | 0,000343929 | 0,067128592 |
| SRSF1 | 3 | 3 | -0,262721113 | 0,00036389 | 0,067128592 |
| LYZ1 | 1 | 1 | -0,566879178 | 0,000382988 | 0,067351128 |
| MCUB | 3 | 2 | 0,405428866 | 0,000443718 | 0,076216256 |
| MARCKSL1 | 2 | 2 | -0,403105318 | 0,00051374 | 0,085191133 |
| PCK2 | 23 | 21 | 0,177910779 | 0,000519036 | 0,085191133 |
| TOMM70 | 28 | 28 | 0,097138131 | 0,000544009 | 0,087348973 |
| SYT12 | 13 | 13 | 0,104187608 | 0,000591418 | 0,092940773 |
| CYGB | 9 | 9 | -0,168341878 | 0,000620052 | 0,095410573 |
| CACNB2 | 8 | 8 | 0,190293933 | 0,000640052 | 0,096478022 |
| GNL1 | 13 | 13 | 0,215293194 | 0,000661049 | 0,097650205 |
| PCCB | 28 | 27 | 0,12823769 | 0,000711826 | 0,103089104 |
| KCMF1 | 3 | 3 | -0,195993396 | 0,000742619 | 0,104420097 |
| HNRNPC | 6 | 6 | -0,320238402 | 0,000763429 | 0,104420097 |
| UBL4A | 7 | 7 | -0,133619368 | 0,000758338 | 0,104420097 |
| SFXN3 | 15 | 14 | 0,085364386 | 0,000855533 | 0,110092678 |
| SEC23IP | 18 | 18 | -0,10574871 | 0,000856785 | 0,110092678 |
| HAL | 3 | 1 | -1,484948352 | 0,000864524 | 0,110092678 |
| GNG4 | 1 | 1 | -0,388063124 | 0,000861971 | 0,110092678 |
| EPHA4 | 26 | 26 | 0,107080738 | 0,00090734 | 0,111693544 |
| SUPT6H | 4 | 4 | -0,21720837 | 0,000894964 | 0,111693544 |

| | | | | | |
|---|---|---|---|---|---|
| NRP1 | 16 | 16 | 0,096436913 | 0,000987756 | 0,117258039 |
| HNRNPK | 22 | 22 | -0,177238182 | 0,001031922 | 0,117258039 |
| PHB2 | 20 | 19 | 0,090809398 | 0,001029922 | 0,117258039 |
| MTREX | 1 | 1 | -0,497227599 | 0,00097448 | 0,117258039 |
| MON1A | 3 | 3 | -0,2947106 | 0,001028512 | 0,117258039 |
| VPS37A | 1 | 1 | -0,550209378 | 0,001050407 | 0,117550042 |
| NDUFS2 | 27 | 26 | 0,104256565 | 0,001092835 | 0,120472778 |
| ROBO1 | 14 | 14 | 0,133664576 | 0,001227345 | 0,131379252 |
| AP2M1 | 35 | 35 | 0,066843367 | 0,001226344 | 0,131379252 |
| SEC16A | 11 | 11 | -0,138464271 | 0,001272358 | 0,133036393 |
| USP14 | 27 | 27 | -0,079051471 | 0,00127885 | 0,133036393 |
| NT5DC3 | 30 | 30 | 0,122210931 | 0,001301688 | 0,133531453 |
| SEC13 | 3 | 3 | -0,167871813 | 0,001339081 | 0,13548568 |
| PITPNM2 | 21 | 21 | -0,100429674 | 0,001362497 | 0,135991881 |
| FUBP1 | 3 | 3 | -0,213725455 | 0,001398481 | 0,137722447 |
| CSE1L | 36 | 35 | -0,104830949 | 0,001458255 | 0,140470797 |
| EIF4G1 | 22 | 22 | -0,106887103 | 0,001464426 | 0,140470797 |
| CSTF3 | 1 | 1 | -0,905864278 | 0,001600319 | 0,151537858 |
| CSNK1A1 | 1 | 1 | -0,435848061 | 0,001658723 | 0,153141564 |
| DLGAP1 | 16 | 16 | 0,083520601 | 0,001650861 | 0,153141564 |
| EIF3L | 23 | 23 | -0,081760774 | 0,0016931 | 0,154385653 |
| TBC1D17 | 11 | 11 | -0,090470228 | 0,001830253 | 0,164856671 |
| TECPR1 | 20 | 20 | 0,087030497 | 0,001858993 | 0,165134389 |
| CFAP36 | 11 | 11 | -0,1563494 | 0,001878052 | 0,165134389 |
| STXBP6 | 4 | 4 | -0,239896519 | 0,002061542 | 0,179135861 |
| UQCR10 | 3 | 3 | 0,135478828 | 0,0021206 | 0,180031614 |
| WDR91 | 7 | 7 | -0,126777849 | 0,002097531 | 0,180031614 |
| FAM120A | 23 | 23 | -0,072421606 | 0,002145217 | 0,18005198 |
| KATNB1 | 10 | 10 | -0,143038661 | 0,00222062 | 0,184230377 |
| PPP3CA | 1 | 1 | -0,275395849 | 0,002244887 | 0,184230377 |
| EFHD2 | 20 | 20 | -0,125474693 | 0,002337742 | 0,185661989 |
| MADD | 1 | 1 | 0,641542969 | 0,002301364 | 0,185661989 |
| PSMD4 | 8 | 8 | -0,092539144 | 0,00231625 | 0,185661989 |

| | | | | | |
|---|---|---|---|---|---|
| CAMK2D | 14 | 13 | -0,137020068 | 0,002420576 | 0,190195465 |
| BZW2 | 3 | 3 | -0,21187782 | 0,002520443 | 0,195957809 |
| ATP6V0A1 | 43 | 43 | 0,06527346 | 0,002580774 | 0,196511271 |
| KCNF1 | 1 | 1 | 1,002772567 | 0,002574191 | 0,196511271 |
| SRPK2 | 11 | 11 | -0,106798874 | 0,002799453 | 0,196796425 |
| SBF1 | 58 | 58 | 0,061899039 | 0,002903624 | 0,196796425 |
| SHANK1 | 52 | 51 | 0,100911911 | 0,002966625 | 0,196796425 |
| GPAM | 1 | 1 | -1,150552732 | 0,002887858 | 0,196796425 |
| GORASP2 | 9 | 9 | -0,192515661 | 0,002940437 | 0,196796425 |
| GRIA1 | 37 | 37 | 0,071596061 | 0,003037475 | 0,196796425 |
| SYNGAP1 | 74 | 72 | 0,112162973 | 0,002989258 | 0,196796425 |
| TCEA1 | 13 | 12 | -0,097059783 | 0,002716652 | 0,196796425 |
| ERC1 | 17 | 17 | 0,12225764 | 0,002723184 | 0,196796425 |
| CA4 | 7 | 7 | 0,171232848 | 0,002815554 | 0,196796425 |
| CRKL | 14 | 14 | -0,089964119 | 0,002624594 | 0,196796425 |
| ALDH1B1 | 21 | 20 | 0,078956769 | 0,002987724 | 0,196796425 |
| NGEF | 14 | 14 | -0,147819472 | 0,002876116 | 0,196796425 |
| GNAS | 1 | 1 | -0,546130617 | 0,002742795 | 0,196796425 |
| GET1 | 1 | 1 | -0,503985429 | 0,003012529 | 0,196796425 |
| ARF4 | 6 | 6 | -0,209875365 | 0,002776717 | 0,196796425 |
| NASP | 2 | 2 | -0,217170417 | 0,002978175 | 0,196796425 |
| MADD | 47 | 47 | 0,052782502 | 0,003195246 | 0,205218162 |
| GK | 19 | 19 | 0,075566712 | 0,003229944 | 0,205658332 |
| RASGRF1 | 18 | 18 | -0,081361067 | 0,003345848 | 0,208371589 |
| ASS1 | 15 | 15 | -0,115859586 | 0,003349039 | 0,208371589 |
| MTCO2 | 12 | 12 | 0,134148225 | 0,003357192 | 0,208371589 |
| PDCD4 | 3 | 3 | -0,17127176 | 0,00345889 | 0,21231245 |
| EIF3F | 15 | 14 | -0,097174143 | 0,003478176 | 0,21231245 |
| AP1S2 | 2 | 2 | -0,177977181 | 0,003541352 | 0,212653857 |
| RPL18 | 6 | 6 | -0,195421279 | 0,003535415 | 0,212653857 |
| ANK2 | 3 | 3 | 0,186942335 | 0,003571192 | 0,212716333 |
| RPS6KA1 | 5 | 5 | 0,18184087 | 0,003603928 | 0,212948888 |
| SERPINA3K | 23 | 23 | -0,610287488 | 0,003684072 | 0,214794181 |

| | | | | | |
|---|---|---|---|---|---|
| RGS6 | 5 | 5 | -0,147588423 | 0,003722401 | 0,214794181 |
| ANKRD44 | 1 | 1 | -0,477225338 | 0,003707248 | 0,214794181 |
| BABAM2 | 6 | 5 | -0,131120127 | 0,003839648 | 0,219842182 |
| OGDH | 50 | 50 | 0,093921857 | 0,003898064 | 0,220505522 |
| ADCY9 | 31 | 31 | 0,114978937 | 0,003910943 | 0,220505522 |
| FUS | 9 | 9 | -0,13918993 | 0,004007069 | 0,221973305 |
| ZCCHC8 | 1 | 1 | -0,360756988 | 0,00399216 | 0,221973305 |
| GAS7 | 1 | 1 | 0,385803596 | 0,004027136 | 0,221973305 |
| CDKN1B | 2 | 2 | -0,305747581 | 0,004076885 | 0,223050904 |
| IGFBP5 | 1 | 1 | -1,079561786 | 0,004139563 | 0,224814821 |
| sp\|Q8C3W1\|CA198 | 9 | 9 | -0,121811752 | 0,004230887 | 0,228097317 |
| COX6A1 | 3 | 3 | 0,254325775 | 0,00435517 | 0,233096286 |
| PRUNE2 | 4 | 4 | 0,14186144 | 0,004417855 | 0,23475017 |
| SH3GL3 | 12 | 12 | 0,101220204 | 0,004558311 | 0,240483457 |
| EIF2S2 | 9 | 9 | -0,091731502 | 0,004686721 | 0,243775492 |
| ADAM11 | 10 | 10 | 0,096891655 | 0,004662005 | 0,243775492 |
| KBTBD11 | 18 | 18 | -0,099074694 | 0,004756142 | 0,243950458 |
| ARHGEF6 | 6 | 6 | 0,163533917 | 0,004747832 | 0,243950458 |
| IQSEC3 | 16 | 16 | 0,102188462 | 0,004869515 | 0,245438624 |
| GUCY1B1 | 18 | 18 | -0,091830608 | 0,004911399 | 0,245438624 |
| PTPRZ1 | 34 | 34 | -0,058598442 | 0,004935613 | 0,245438624 |
| GLIPR2 | 2 | 2 | 0,199221177 | 0,004930137 | 0,245438624 |
| USP10 | 9 | 9 | -0,105264556 | 0,004951307 | 0,245438624 |
| PA2G4 | 20 | 20 | -0,092090055 | 0,005016906 | 0,247032448 |
| OGFR | 9 | 9 | -0,146425531 | 0,005126847 | 0,249124285 |
| SERBP1 | 15 | 15 | -0,131964257 | 0,005094512 | 0,249124285 |
| SNRPA1 | 1 | 1 | -0,281944874 | 0,005209438 | 0,251483046 |
| RYR2 | 19 | 19 | 0,078790532 | 0,00529832 | 0,254112943 |
| SLC25A12;SLC25A1 | 6 | 5 | 0,127458904 | 0,005385286 | 0,256617585 |
| NIPSNAP2 | 15 | 14 | 0,102223245 | 0,00556888 | 0,260327499 |
| DPYSL4 | 28 | 28 | -0,101786225 | 0,005508573 | 0,260327499 |
| PDXDC1 | 5 | 5 | -0,144748415 | 0,005548043 | 0,260327499 |
| PCCA | 41 | 40 | 0,089157315 | 0,005713079 | 0,263729997 |

| | | | | | |
|---|---|---|---|---|---|
| MUC19 | 1 | 1 | -1,002082545 | 0,005688528 | 0,263729997 |
| PITPNC1 | 9 | 9 | -0,176541455 | 0,005847473 | 0,268257385 |
| MPP2 | 22 | 21 | -0,072700924 | 0,005961427 | 0,268482304 |
| EGLN3 | 1 | 1 | 0,30488974 | 0,005933067 | 0,268482304 |
| CAMK2A | 16 | 16 | 0,097485163 | 0,005899535 | 0,268482304 |
| MFF | 11 | 11 | 0,143019338 | 0,005999122 | 0,26854251 |
| ZCCHC7 | 1 | 1 | 0,298788645 | 0,00606376 | 0,269800801 |
| RIMS1 | 12 | 11 | 0,119207269 | 0,006337488 | 0,276237207 |
| DDX19A | 8 | 8 | -0,104442057 | 0,006352942 | 0,276237207 |
| DYNLT1 | 1 | 1 | -0,286668862 | 0,006358019 | 0,276237207 |
| SMTNL2 | 1 | 1 | -0,471103964 | 0,00631151 | 0,276237207 |
| LPP | 1 | 1 | -0,550029925 | 0,006439019 | 0,27812045 |
| COX20 | 2 | 2 | 0,300956728 | 0,006588276 | 0,282912821 |
| CNKSR2 | 35 | 34 | 0,060945617 | 0,006770289 | 0,283662321 |
| ANKS1B | 15 | 15 | 0,098583364 | 0,006912973 | 0,283662321 |
| MYO18A | 70 | 69 | 0,037511043 | 0,0067391 | 0,283662321 |
| KRT17 | 9 | 5 | 0,515054784 | 0,006688671 | 0,283662321 |
| TSPYL4 | 4 | 4 | -0,200695698 | 0,006912149 | 0,283662321 |
| FAM177A1 | 4 | 3 | 0,292771839 | 0,006863131 | 0,283662321 |
| TIMM10 | 4 | 4 | 0,124132639 | 0,00687398 | 0,283662321 |
| PTPRD | 1 | 1 | -0,987513604 | 0,006781871 | 0,283662321 |
| USO1 | 36 | 36 | -0,09534593 | 0,007526151 | 0,307116838 |
| APLP1 | 12 | 11 | -0,113693393 | 0,007704513 | 0,312269043 |
| NRBP1 | 7 | 7 | -0,105842949 | 0,007736967 | 0,312269043 |
| SEPTIN8 | 30 | 30 | 0,056604 | 0,008029834 | 0,318862108 |
| SORBS1 | 8 | 8 | -0,156766824 | 0,008003614 | 0,318862108 |
| ATP6V1B2 | 26 | 26 | 0,062707638 | 0,00795387 | 0,318862108 |
| NAPB | 26 | 26 | 0,050103594 | 0,008389186 | 0,321438651 |
| WDR41 | 6 | 5 | 0,205138968 | 0,008403805 | 0,321438651 |
| ME1 | 29 | 29 | 0,153990509 | 0,008194883 | 0,321438651 |
| CUL1 | 27 | 27 | 0,055872115 | 0,008397364 | 0,321438651 |
| PDK1 | 10 | 9 | 0,120539943 | 0,008580389 | 0,321438651 |
| MAPT | 6 | 6 | -0,23390008 | 0,008552148 | 0,321438651 |

| | | | | | |
|---|---|---|---|---|---|
| CSRP1 | 10 | 10 | -0,106791503 | 0,008314489 | 0,321438651 |
| RANGAP1 | 11 | 11 | -0,119509861 | 0,00854571 | 0,321438651 |
| GPC4 | 12 | 11 | -0,097364423 | 0,008336389 | 0,321438651 |
| ALYREF;ALYREF2 | 1 | 1 | -0,434416247 | 0,008537253 | 0,321438651 |
| EIF4E2 | 1 | 1 | -0,313062707 | 0,008616958 | 0,321438651 |
| KCNAB3 | 1 | 1 | -0,286949408 | 0,008327216 | 0,321438651 |
| SGTA | 15 | 15 | -0,135104936 | 0,008675012 | 0,321978085 |
| H2-Q10;H2-D1 | 1 | 1 | 1,071102664 | 0,008852762 | 0,326932485 |
| DYNC1I2 | 9 | 9 | -0,105358133 | 0,008932108 | 0,328221639 |
| GOLGA2 | 5 | 5 | -0,191976333 | 0,008997713 | 0,328995583 |
| NCDN | 35 | 35 | -0,16568926 | 0,009104722 | 0,329644499 |
| MFSD6 | 3 | 2 | 0,317327463 | 0,009095735 | 0,329644499 |
| PLCH2 | 27 | 26 | 0,129888376 | 0,009184371 | 0,329780113 |
| SNX5 | 14 | 13 | 0,081680396 | 0,009240875 | 0,329780113 |
| NOVA1 | 4 | 4 | -0,197851783 | 0,009245466 | 0,329780113 |
| HNRNPF | 2 | 2 | -0,174673344 | 0,009300581 | 0,329780113 |
| VSTM2B | 1 | 1 | -0,900508502 | 0,009331715 | 0,329780113 |
| IPO5 | 35 | 35 | -0,073234078 | 0,009687278 | 0,333674446 |
| BZW1 | 12 | 12 | -0,186093166 | 0,009720759 | 0,333674446 |
| G3BP2 | 16 | 16 | -0,148723565 | 0,009727663 | 0,333674446 |
| AGO2 | 12 | 12 | -0,094049329 | 0,00973973 | 0,333674446 |
| APPL2 | 15 | 15 | -0,08253219 | 0,009758148 | 0,333674446 |
| SYNGAP1;DAB2IP | 2 | 2 | 0,205714081 | 0,00948734 | 0,333674446 |
| POU3F2 | 1 | 1 | 0,312923961 | 0,009710243 | 0,333674446 |
| DCXR | 3 | 3 | -0,208733385 | 0,009952618 | 0,336728038 |
| MFN2 | 31 | 31 | 0,064629237 | 0,009979683 | 0,336728038 |
| SCN9A | 1 | 1 | 0,350146824 | 0,009984219 | 0,336728038 |
| PPP1R3G | 4 | 4 | -0,154126085 | 0,010121302 | 0,33979972 |
| ACACA | 27 | 26 | 0,055319359 | 0,010182305 | 0,340297012 |
| MTSS2 | 6 | 6 | -0,173876653 | 0,010274334 | 0,340297012 |
| OLFR1234 | 1 | 1 | 0,437294837 | 0,010245707 | 0,340297012 |
| LGALS8 | 1 | 1 | -0,467833922 | 0,010388308 | 0,341809411 |
| COL24A1 | 1 | 1 | -0,266286192 | 0,010412553 | 0,341809411 |

| | | | | | |
|---|---|---|---|---|---|
| EIF4G1 | 1 | 1 | -0,254788813 | 0,010513032 | 0,343580775 |
| EDC3 | 2 | 2 | -0,411463866 | 0,010647229 | 0,346433621 |
| PHB1 | 22 | 21 | 0,070582367 | 0,010710613 | 0,346967491 |
| SETD3 | 10 | 10 | -0,094433271 | 0,011095055 | 0,347942105 |
| GFM1 | 21 | 19 | 0,081585144 | 0,011117565 | 0,347942105 |
| ELMOD1 | 3 | 3 | 0,188160944 | 0,010903512 | 0,347942105 |
| LRRC40 | 9 | 9 | -0,139671624 | 0,011068713 | 0,347942105 |
| TMEM263 | 2 | 2 | 0,247935777 | 0,010944524 | 0,347942105 |
| CBLN4 | 1 | 1 | -0,259940285 | 0,011094801 | 0,347942105 |
| PUF60 | 8 | 8 | -0,125997214 | 0,010789777 | 0,347942105 |
| MS4A6D | 1 | 1 | 0,453467863 | 0,011077878 | 0,347942105 |
| GNAI1 | 9 | 9 | 0,086810343 | 0,011266432 | 0,349004933 |
| SIDT2 | 1 | 1 | 0,290497668 | 0,011201657 | 0,349004933 |
| LRRTM4 | 4 | 4 | -0,213744314 | 0,011293282 | 0,349004933 |
| IMMT | 55 | 54 | 0,074498644 | 0,011496162 | 0,353422628 |
| CIAO2B | 3 | 3 | -0,132372259 | 0,011531932 | 0,353422628 |
| RPN2 | 16 | 15 | -0,071577111 | 0,011645754 | 0,355436105 |
| ANKS1B | 4 | 4 | 0,137463981 | 0,011804523 | 0,357328717 |
| SLC17A7 | 14 | 13 | 0,093561634 | 0,011764892 | 0,357328717 |
| AACS | 11 | 11 | -0,128186509 | 0,012002011 | 0,361823889 |
| ALDH3A2 | 1 | 1 | 0,259810721 | 0,012163292 | 0,361864813 |
| SRI | 6 | 4 | -0,110843706 | 0,012199342 | 0,361864813 |
| KALRN;TRIO | 8 | 8 | 0,083849627 | 0,012191519 | 0,361864813 |
| RAB26 | 4 | 3 | 0,271675926 | 0,012061097 | 0,361864813 |
| UPF1 | 26 | 26 | -0,053949025 | 0,012355917 | 0,365043199 |
| PCBP1 | 12 | 12 | -0,094824541 | 0,012703181 | 0,373807535 |
| ATXN10 | 23 | 22 | -0,069593893 | 0,01280353 | 0,375265376 |
| TSC22D1 | 4 | 4 | -0,208642819 | 0,012971327 | 0,37868073 |
| GRM2 | 15 | 14 | -0,101872185 | 0,013084789 | 0,380489176 |
| MINAR1 | 1 | 1 | -0,779769552 | 0,013487129 | 0,390650729 |
| PSMC2 | 27 | 27 | -0,08015628 | 0,013639135 | 0,39097468 |
| ALDH1A1 | 19 | 19 | 0,16710906 | 0,013717819 | 0,39097468 |
| ARIH1 | 5 | 5 | -0,096491775 | 0,013748245 | 0,39097468 |

| | | | | | |
|---|---|---|---|---|---|
| UIMC1 | 1 | 1 | 0,300113455 | 0,013617093 | 0,39097468 |
| GABPA | 1 | 1 | -0,453678889 | 0,013767109 | 0,39097468 |
| FAM184B | 1 | 1 | 0,313771822 | 0,013815921 | 0,39097468 |
| VDAC3 | 12 | 12 | 0,087414629 | 0,013984799 | 0,391548703 |
| HNRNPUL2 | 29 | 29 | -0,122423637 | 0,01389882 | 0,391548703 |
| ZNF148 | 1 | 1 | 0,40686566 | 0,013995242 | 0,391548703 |
| KLC2 | 15 | 15 | -0,093696429 | 0,014306512 | 0,398746793 |
| SDHB | 18 | 17 | 0,101735042 | 0,014422663 | 0,400472887 |
| KIF3A | 17 | 17 | -0,07310992 | 0,014586278 | 0,403499067 |
| WDFY3 | 27 | 26 | 0,120405604 | 0,014650589 | 0,403765853 |
| MADD | 3 | 2 | -0,201982761 | 0,01495127 | 0,410520732 |
| BMP2K | 2 | 2 | 0,212850103 | 0,015251842 | 0,4172226 |
| CHPF | 1 | 1 | 0,363505522 | 0,01536415 | 0,418743948 |
| MRTFB;MRTFA;MY | 1 | 1 | -0,182952008 | 0,015433558 | 0,419089178 |
| CYCS | 14 | 14 | 0,091059738 | 0,015671421 | 0,420954568 |
| KHSRP | 9 | 9 | -0,135232378 | 0,015730228 | 0,420954568 |
| SAT2 | 1 | 1 | 0,248524532 | 0,015566877 | 0,420954568 |
| ANKRD27 | 1 | 1 | 0,255381961 | 0,015690281 | 0,420954568 |
| ACBD3 | 6 | 6 | -0,110180736 | 0,015804397 | 0,421412544 |
| RPL11 | 6 | 6 | -0,089579692 | 0,015909128 | 0,422679204 |
| IGLON5 | 4 | 4 | -0,103725533 | 0,016134177 | 0,42508188 |
| DOC2A | 3 | 3 | -0,187378479 | 0,016172219 | 0,42508188 |
| RYR1;RYR2 | 1 | 1 | 0,220033113 | 0,016064344 | 0,42508188 |
| NPM1 | 10 | 8 | -0,238126778 | 0,016284972 | 0,426527658 |
| ACTN1;ACTN2 | 13 | 13 | -0,077555474 | 0,016901572 | 0,43168962 |
| ANKRD34A | 6 | 6 | -0,132232725 | 0,016809357 | 0,43168962 |
| WIPF2 | 7 | 7 | -0,141629223 | 0,016914792 | 0,43168962 |
| VBP1 | 12 | 12 | -0,089293245 | 0,016949633 | 0,43168962 |
| NDUFA10 | 19 | 18 | 0,108262625 | 0,01668241 | 0,43168962 |
| PHKB | 7 | 7 | -0,097299442 | 0,016664825 | 0,43168962 |
| CAPNS1 | 9 | 9 | -0,139747045 | 0,01669992 | 0,43168962 |
| PLA2G6 | 2 | 2 | 0,226756833 | 0,016876643 | 0,43168962 |
| LRRC7 | 46 | 44 | 0,081049746 | 0,0172632 | 0,434417198 |

| | | | | | |
|---|---|---|---|---|---|
| GPD2 | 56 | 52 | 0,080237937 | 0,017198216 | 0,434417198 |
| ITIH2 | 2 | 2 | 0,29059576 | 0,017409625 | 0,434417198 |
| NFYB | 2 | 2 | -0,23930488 | 0,017337056 | 0,434417198 |
| LMCD1 | 3 | 3 | -0,163624609 | 0,017367255 | 0,434417198 |
| SERPINA1B;SERPIN. | 6 | 6 | -0,438487795 | 0,017237484 | 0,434417198 |
| DNAJB4 | 8 | 8 | -0,117690398 | 0,017569 | 0,434781359 |
| ELMO3 | 1 | 1 | 0,857495371 | 0,017600816 | 0,434781359 |
| S1PR1 | 4 | 4 | -0,203453852 | 0,017507132 | 0,434781359 |
| CES1C;CES1D | 1 | 1 | 0,303689013 | 0,017797103 | 0,43526292 |
| SLITRK4 | 1 | 1 | -0,562510287 | 0,017700641 | 0,43526292 |
| TEKT3 | 1 | 1 | 0,313298478 | 0,017758142 | 0,43526292 |
| VARS1 | 31 | 31 | -0,040022493 | 0,017859608 | 0,435350061 |
| SLC12A6 | 7 | 7 | -0,096444513 | 0,018254354 | 0,44205463 |
| SLC35A4 | 1 | 1 | 0,265680538 | 0,018247947 | 0,44205463 |
| DOCK4 | 35 | 35 | 0,04482228 | 0,018363137 | 0,443235718 |
| CISD1 | 9 | 9 | 0,090367333 | 0,018444389 | 0,443746757 |
| MICU3 | 21 | 20 | 0,084978469 | 0,018580885 | 0,445579267 |
| NDUFS7 | 9 | 9 | 0,095604003 | 0,0189337 | 0,451110681 |
| GRIN1 | 1 | 1 | 0,357576384 | 0,01891933 | 0,451110681 |
| GRIN2B | 39 | 33 | 0,078473341 | 0,019190145 | 0,45428978 |
| ME3 | 28 | 26 | 0,129744422 | 0,019174041 | 0,45428978 |
| NDUFA5 | 8 | 6 | 0,118426482 | 0,019781476 | 0,466792283 |
| CHAT | 6 | 6 | -0,086900779 | 0,019971614 | 0,468286804 |
| OPHN1 | 2 | 2 | 0,259711583 | 0,019966486 | 0,468286804 |
| SNX3 | 6 | 6 | -0,094859757 | 0,02009936 | 0,469790741 |
| ANKRD28 | 4 | 3 | 0,106903043 | 0,020251035 | 0,471842734 |
| ABHD12 | 17 | 17 | -0,059367733 | 0,020447284 | 0,473428346 |
| UNC13A;UNC13C;U | 5 | 5 | 0,154722317 | 0,020415792 | 0,473428346 |
| IQSEC2 | 32 | 30 | 0,052678354 | 0,020650894 | 0,474111094 |
| NACA | 8 | 7 | -0,078062738 | 0,020657209 | 0,474111094 |
| LIN7B;LIN7C;LIN7A | 3 | 3 | -0,217177941 | 0,020669344 | 0,474111094 |
| ELAVL1 | 7 | 7 | -0,104342122 | 0,020887874 | 0,476166174 |
| APOH | 1 | 1 | -0,470300346 | 0,020874712 | 0,476166174 |

| | | | | | |
|---|---|---|---|---|---|
| ACO2 | 53 | 52 | 0,086131042 | 0,021066667 | 0,47876432 |
| GPR162 | 3 | 3 | -0,1425902 | 0,021133561 | 0,478811282 |
| VDAC2 | 14 | 14 | 0,069935157 | 0,021207215 | 0,479010677 |
| MYO9A | 1 | 1 | -0,200386452 | 0,021289129 | 0,479394841 |
| GABRG2 | 9 | 9 | -0,088751625 | 0,021738675 | 0,48508112 |
| COQ8A | 7 | 5 | 0,141133868 | 0,021727992 | 0,48508112 |
| RPAP1 | 1 | 1 | 0,607347506 | 0,021622683 | 0,48508112 |
| ACAP3 | 2 | 2 | 0,226326397 | 0,021902799 | 0,487271308 |
| GRM3 | 22 | 22 | -0,079480617 | 0,021991172 | 0,487768166 |
| AP2A2 | 54 | 54 | 0,067577395 | 0,022135514 | 0,488634704 |
| ADCY5;ADCY6 | 2 | 2 | 0,145152066 | 0,022228711 | 0,488634704 |
| SERINC5 | 1 | 1 | -0,412173245 | 0,022164779 | 0,488634704 |
| SARNP | 2 | 2 | -0,230790598 | 0,022312207 | 0,48901472 |
| AGK | 26 | 25 | 0,113089733 | 0,022666727 | 0,495314933 |
| UBXN2A | 1 | 1 | -0,397039947 | 0,022796791 | 0,496687612 |
| ARHGEF2 | 17 | 17 | 0,067147123 | 0,022940774 | 0,496893133 |
| UTP20 | 1 | 1 | -0,338808864 | 0,022885613 | 0,496893133 |
| SLC30A1 | 8 | 8 | 0,086330795 | 0,023207676 | 0,500743631 |
| ASAP1 | 20 | 20 | -0,066891759 | 0,023305033 | 0,500743631 |
| SHTN1 | 2 | 2 | -0,179804579 | 0,023341573 | 0,500743631 |
| ABI3 | 3 | 3 | -0,216470378 | 0,023389731 | 0,500743631 |
| PHACTR1;RAI14 | 1 | 1 | -0,216031797 | 0,023744146 | 0,506862027 |
| PDE1B | 18 | 18 | -0,133266178 | 0,024078549 | 0,512519208 |
| PHGDH | 18 | 18 | -0,096686184 | 0,024208211 | 0,513798409 |
| FTO | 6 | 6 | -0,123697213 | 0,024343859 | 0,515196971 |
| MIGA2 | 7 | 6 | 0,113387765 | 0,024427088 | 0,515481348 |
| PHYHIPL | 16 | 16 | -0,088110017 | 0,024674186 | 0,517778278 |
| KIF2A;KIF2B | 1 | 1 | 0,318007533 | 0,024676138 | 0,517778278 |
| BNIP3L | 2 | 2 | 0,25276644 | 0,024763301 | 0,518135237 |
| NPY2R | 2 | 1 | -0,529017554 | 0,024870664 | 0,518911647 |
| DNAJC6 | 24 | 24 | 0,055123577 | 0,024982848 | 0,519783986 |
| HAPLN1 | 21 | 20 | -0,170691531 | 0,025100759 | 0,520770243 |
| FSD1 | 21 | 21 | -0,056314509 | 0,025450465 | 0,526546592 |

| | | | | | |
|---|---|---|---|---|---|
| DLD | 22 | 22 | 0,083767172 | 0,025579531 | 0,527738598 |
| CLTC | 130 | 129 | 0,066661808 | 0,026115569 | 0,531011028 |
| ATP6V0D1 | 19 | 19 | 0,054478777 | 0,026515152 | 0,531011028 |
| AP3D1 | 31 | 31 | 0,213409461 | 0,026525069 | 0,531011028 |
| SPAG9 | 29 | 29 | -0,078390935 | 0,026600877 | 0,531011028 |
| NUMA1 | 1 | 1 | -0,402051997 | 0,026553462 | 0,531011028 |
| NDUFS1 | 55 | 55 | 0,069449807 | 0,026067762 | 0,531011028 |
| DGKB | 16 | 16 | 0,075747936 | 0,026127415 | 0,531011028 |
| CYB5R1 | 15 | 14 | 0,108228235 | 0,026339892 | 0,531011028 |
| SERPINA1C | 6 | 6 | -0,39770215 | 0,025912298 | 0,531011028 |
| ISCA2 | 6 | 5 | 0,110186592 | 0,026489654 | 0,531011028 |
| CAMLG | 2 | 2 | -0,27694913 | 0,026334274 | 0,531011028 |
| FLNB | 1 | 1 | 0,429046212 | 0,026011143 | 0,531011028 |
| LPCAT4 | 5 | 5 | 0,123788727 | 0,027001187 | 0,533676529 |
| CYRIA | 15 | 14 | -0,068411791 | 0,027095681 | 0,533676529 |
| CELSR2 | 3 | 3 | 0,165742929 | 0,027054914 | 0,533676529 |
| GALNT16 | 3 | 3 | 0,307215812 | 0,027081609 | 0,533676529 |
| RAB3D;RAB3B | 1 | 1 | 0,466090519 | 0,026977648 | 0,533676529 |
| HNRNPA3 | 10 | 10 | -0,114225604 | 0,027381159 | 0,537865002 |
| NECTIN1 | 8 | 8 | 0,122458376 | 0,027660796 | 0,541916822 |
| EPB41L1 | 62 | 62 | -0,035530396 | 0,027846304 | 0,544107945 |
| G3BP1 | 8 | 8 | -0,083671 | 0,028203276 | 0,548417978 |
| NT5C1A | 1 | 1 | -0,278666885 | 0,028215385 | 0,548417978 |
| DECR1 | 11 | 10 | 0,076971788 | 0,028417772 | 0,550902003 |
| DLG4 | 33 | 33 | 0,036071981 | 0,028602091 | 0,551089641 |
| ATP2B3 | 33 | 33 | 0,044088616 | 0,028626114 | 0,551089641 |
| GM26992 | 1 | 1 | 0,2279629 | 0,028651289 | 0,551089641 |
| PPP1R9A | 18 | 18 | -0,071521076 | 0,028936558 | 0,553222112 |
| UBQLN4;UBQLN2 | 1 | 1 | -0,201886916 | 0,028983939 | 0,553222112 |
| SLC20A1 | 2 | 2 | -0,160423495 | 0,028986861 | 0,553222112 |
| RELN | 4 | 4 | -0,114579806 | 0,029097462 | 0,553901685 |
| TNPO2 | 9 | 9 | -0,095699543 | 0,029174329 | 0,553937259 |
| EPN2 | 9 | 9 | -0,105375574 | 0,02951239 | 0,558596009 |

| | | | | | |
|---|---|---|---|---|---|
| KRT6A | 19 | 6 | 0,576070605 | 0,029582514 | 0,558596009 |
| VPS13B | 2 | 1 | -0,479478453 | 0,029646579 | 0,558596009 |
| NUDT16 | 4 | 4 | 0,119592202 | 0,029962661 | 0,563115041 |
| PRKACA | 21 | 19 | -0,065308857 | 0,030512205 | 0,571579006 |
| PALMD | 2 | 2 | -0,114700165 | 0,030567791 | 0,571579006 |
| BAG6 | 22 | 21 | -0,101886625 | 0,031396299 | 0,577438661 |
| PDK2 | 12 | 12 | 0,087456482 | 0,031390745 | 0,577438661 |
| ATP5A1 | 50 | 50 | 0,074224648 | 0,031340181 | 0,577438661 |
| KIF21B | 1 | 1 | -0,317549685 | 0,031382545 | 0,577438661 |
| CEBPZOS | 1 | 1 | 0,678123656 | 0,031221309 | 0,577438661 |
| STRN3 | 2 | 1 | -0,283933171 | 0,03111059 | 0,577438661 |
| KLHDC7A | 1 | 1 | 0,221154881 | 0,031428424 | 0,577438661 |
| SLC24A2 | 7 | 7 | 0,091456068 | 0,031791394 | 0,581215926 |
| RNF126 | 1 | 1 | -0,235898592 | 0,031749275 | 0,581215926 |
| CNOT3 | 3 | 3 | -0,147695875 | 0,032250577 | 0,585092637 |
| HSPD1 | 52 | 50 | 0,081250579 | 0,032320308 | 0,585092637 |
| PNPT1 | 17 | 15 | 0,070736298 | 0,032097915 | 0,585092637 |
| EIF4G2 | 22 | 22 | -0,089055629 | 0,032235884 | 0,585092637 |
| USP9X | 63 | 63 | 0,027764608 | 0,032428591 | 0,585204077 |
| COPB1 | 18 | 17 | -0,060963668 | 0,032605804 | 0,585204077 |
| HSD11B1 | 3 | 3 | -0,22761881 | 0,032494165 | 0,585204077 |
| GPR155 | 2 | 2 | 0,242805745 | 0,032722622 | 0,585204077 |
| ENTPD2 | 9 | 9 | -0,123691673 | 0,032671062 | 0,585204077 |
| RAB5A | 14 | 14 | -0,061136643 | 0,033203883 | 0,592376518 |
| YBX1;YBX3 | 3 | 3 | -0,228370423 | 0,033418691 | 0,594772168 |
| LAMP5 | 2 | 2 | -0,12397539 | 0,033656432 | 0,597563469 |
| SF3B1 | 2 | 2 | -0,207503616 | 0,033826575 | 0,599144089 |
| PRKAR1A | 17 | 17 | -0,064117422 | 0,034281837 | 0,605755142 |
| TOM1L2 | 23 | 23 | -0,099067787 | 0,035054433 | 0,611651632 |
| WFS1 | 19 | 19 | -0,084293556 | 0,034907386 | 0,611651632 |
| STK24 | 8 | 8 | -0,130606399 | 0,034781679 | 0,611651632 |
| BSN | 157 | 156 | 0,089546685 | 0,035006595 | 0,611651632 |
| TRIM32 | 6 | 6 | -0,11606531 | 0,034888156 | 0,611651632 |

| | | | | | |
|---|---|---|---|---|---|
| ELAVL4 | 4 | 4 | -0,157082132 | 0,035153884 | 0,611651632 |
| HSPA12A | 1 | 1 | 0,664088116 | 0,035195227 | 0,611651632 |
| ITCH;NEDD4L | 1 | 1 | -0,314545027 | 0,035476767 | 0,612222892 |
| AMIGO1 | 3 | 1 | -0,459214318 | 0,035406666 | 0,612222892 |
| SEC24A | 1 | 1 | 0,209515018 | 0,035325829 | 0,612222892 |
| VAMP2 | 4 | 4 | 0,082541794 | 0,036422237 | 0,627073765 |
| GIT1 | 35 | 34 | 0,040281298 | 0,036537699 | 0,627598711 |
| RAPH1 | 1 | 1 | -0,446290801 | 0,036651454 | 0,628091972 |
| ALDH18A1 | 23 | 21 | 0,068638011 | 0,036790516 | 0,629015622 |
| TAB3 | 1 | 1 | 0,236947932 | 0,03711712 | 0,63313406 |
| OOSP2 | 1 | 1 | -0,150322557 | 0,037614993 | 0,640148241 |
| IQSEC1 | 43 | 43 | 0,039620747 | 0,037989104 | 0,644205258 |
| EIF4G2 | 1 | 1 | -0,207761739 | 0,038027822 | 0,644205258 |
| CNPY2 | 5 | 5 | 0,105054418 | 0,038383116 | 0,646589276 |
| CIAPIN1 | 5 | 5 | -0,101212276 | 0,03843118 | 0,646589276 |
| GGA3 | 8 | 7 | -0,1218216 | 0,038307135 | 0,646589276 |
| GSTK1 | 15 | 15 | 0,068869966 | 0,039151203 | 0,657206336 |
| PLXNA1 | 40 | 40 | 0,076670533 | 0,039494098 | 0,661458969 |
| HNRNPA2B1 | 13 | 13 | -0,117047091 | 0,039686129 | 0,663171385 |
| NOMO1 | 27 | 26 | -0,0638799 | 0,040008692 | 0,664054377 |
| RBM12 | 3 | 3 | -0,188304527 | 0,039951587 | 0,664054377 |
| SERPINA1B;SERPIN | 1 | 1 | -0,458985047 | 0,039844148 | 0,664054377 |
| C3 | 18 | 17 | 0,280729854 | 0,040276711 | 0,664403167 |
| HOMER3 | 17 | 17 | 0,134144512 | 0,040322102 | 0,664403167 |
| PRKACB | 8 | 7 | -0,094745999 | 0,040538803 | 0,664403167 |
| DMTN | 24 | 23 | -0,083700887 | 0,040242339 | 0,664403167 |
| DOCK1 | 7 | 6 | -0,109337988 | 0,040559809 | 0,664403167 |
| C4BPA | 1 | 1 | -0,30624135 | 0,040569432 | 0,664403167 |
| SELENOO | 4 | 4 | -0,151483702 | 0,040793761 | 0,664985988 |
| UCK1 | 5 | 5 | 0,096973372 | 0,04087512 | 0,664985988 |
| SLC39A11 | 1 | 1 | -0,356282472 | 0,040728397 | 0,664985988 |
| TAGLN2 | 12 | 10 | -0,109797129 | 0,041281159 | 0,670115686 |
| DLG2 | 3 | 3 | 0,129969917 | 0,041446352 | 0,671321836 |

| | | | | | |
|---|---|---|---|---|---|
| IREB2 | 2 | 2 | -0,141726553 | 0,041659768 | 0,672467921 |
| RHOF | 2 | 2 | 0,185532838 | 0,041699202 | 0,672467921 |
| KYAT3 | 12 | 9 | 0,114471629 | 0,04262147 | 0,679390682 |
| SLC25A5 | 12 | 11 | 0,065914973 | 0,042230163 | 0,679390682 |
| ATXN2L | 18 | 18 | -0,099551194 | 0,042465798 | 0,679390682 |
| MUG1 | 22 | 22 | -0,457845858 | 0,042600064 | 0,679390682 |
| API5 | 8 | 8 | -0,152884759 | 0,04276054 | 0,679390682 |
| CRACDL | 2 | 2 | -0,232077498 | 0,042748976 | 0,679390682 |
| ADCY8 | 5 | 5 | -0,152267888 | 0,042772362 | 0,679390682 |
| PREPL | 18 | 18 | -0,065970166 | 0,042912826 | 0,680159087 |
| RNF214 | 14 | 14 | -0,112180473 | 0,043325851 | 0,680278881 |
| PRKAG2 | 16 | 16 | 0,075633656 | 0,043195891 | 0,680278881 |
| NIPSNAP1 | 16 | 15 | 0,093658082 | 0,043360331 | 0,680278881 |
| AHCYL1 | 39 | 39 | -0,071976057 | 0,043458051 | 0,680278881 |
| PCBP3 | 6 | 6 | -0,114369181 | 0,043473007 | 0,680278881 |
| NOVA2 | 4 | 4 | -0,133324292 | 0,043167257 | 0,680278881 |
| SLC7A11 | 4 | 4 | -0,220804825 | 0,043955701 | 0,685731048 |
| 0610012G03RIK | 1 | 1 | 0,323855963 | 0,04400711 | 0,685731048 |
| PTPRS | 36 | 36 | 0,058225183 | 0,044150198 | 0,68651234 |
| TUB | 2 | 2 | 0,157641895 | 0,044356849 | 0,686833717 |
| SHROOM2 | 13 | 13 | -0,106961263 | 0,044314315 | 0,686833717 |
| THNSL1 | 21 | 17 | 0,070888181 | 0,045428289 | 0,69339959 |
| TMEM126A | 5 | 5 | 0,088682221 | 0,045022333 | 0,69339959 |
| ANKRD29 | 4 | 4 | -0,155728721 | 0,045164988 | 0,69339959 |
| IDH3G | 13 | 12 | 0,073108254 | 0,045305105 | 0,69339959 |
| TPT1 | 7 | 7 | -0,160054833 | 0,04511896 | 0,69339959 |
| CHMP7 | 7 | 7 | -0,140058484 | 0,045438045 | 0,69339959 |
| SNX9 | 1 | 1 | 0,236186379 | 0,045129062 | 0,69339959 |
| CPE | 22 | 22 | 0,073019082 | 0,045589816 | 0,694281197 |
| SCO1 | 2 | 2 | 0,160947839 | 0,045788594 | 0,69444468 |
| ARNT | 1 | 1 | -0,214089739 | 0,045770556 | 0,69444468 |
| COPZ1 | 5 | 5 | -0,099976619 | 0,046106159 | 0,695212531 |
| TMEM100 | 1 | 1 | -0,362136506 | 0,046197226 | 0,695212531 |

| | | | | | |
|---|---|---|---|---|---|
| EPB41L2 | 3 | 3 | -0,134214357 | 0,046215726 | 0,695212531 |
| PKIA | 1 | 1 | -0,401843916 | 0,046084068 | 0,695212531 |
| PLIN3 | 8 | 7 | -0,17394221 | 0,046334495 | 0,695582485 |
| VPS13C | 63 | 63 | 0,046771158 | 0,046935765 | 0,696119603 |
| USP47 | 26 | 25 | -0,041062862 | 0,046497361 | 0,696119603 |
| DGKE | 9 | 9 | -0,085206427 | 0,046563346 | 0,696119603 |
| SNX18 | 5 | 5 | -0,142076073 | 0,046852991 | 0,696119603 |
| PDCD2 | 1 | 1 | 0,192157352 | 0,046842271 | 0,696119603 |
| CDK5RAP3 | 1 | 1 | -0,27885387 | 0,046856775 | 0,696119603 |
| AFDN | 28 | 27 | 0,050347869 | 0,047101487 | 0,697177517 |
| MICAL1 | 4 | 4 | -0,095976993 | 0,047397 | 0,69735905 |
| CES1C | 13 | 13 | 0,191830633 | 0,047298917 | 0,69735905 |
| MMP15 | 1 | 1 | -0,189903209 | 0,047320697 | 0,69735905 |
| MPP3 | 8 | 8 | 0,070345709 | 0,047657544 | 0,698559654 |
| PCDHGA9 | 1 | 1 | 0,517220732 | 0,047762338 | 0,698559654 |
| TIMMDC1 | 6 | 5 | -0,123691642 | 0,047722056 | 0,698559654 |
| SLC1A7 | 1 | 1 | -0,652393907 | 0,047926584 | 0,699576583 |
| CYP46A1 | 21 | 21 | -0,054624447 | 0,048593181 | 0,70439925 |
| RTRAF | 10 | 10 | -0,147421897 | 0,048575376 | 0,70439925 |
| DLG5 | 1 | 1 | 0,300365182 | 0,048534948 | 0,70439925 |
| MAP3K15 | 1 | 1 | 0,189135541 | 0,048638454 | 0,70439925 |
| PRUNE1 | 18 | 18 | -0,06887593 | 0,048830483 | 0,705796375 |
| COG2 | 2 | 2 | -0,161876436 | 0,049318267 | 0,710067679 |
| SYVN1 | 2 | 2 | -0,220345063 | 0,049267133 | 0,710067679 |
| GABBR1 | 15 | 14 | 0,073916856 | 0,050084237 | 0,719692947 |
| WASL | 10 | 10 | -0,095578106 | 0,051059654 | 0,728775831 |
| SORBS1 | 2 | 2 | -0,24569885 | 0,051335347 | 0,728775831 |
| CDC42EP4 | 10 | 10 | -0,109816542 | 0,051604354 | 0,728775831 |
| SQSTM1 | 6 | 6 | -0,086308627 | 0,051520059 | 0,728775831 |
| CACNA2D3 | 31 | 31 | 0,04250782 | 0,051558179 | 0,728775831 |
| ACTN4 | 23 | 22 | -0,080901279 | 0,050839922 | 0,728775831 |
| SARM1 | 11 | 11 | 0,062577656 | 0,051357061 | 0,728775831 |
| NDUFA4 | 6 | 6 | 0,101762039 | 0,051542223 | 0,728775831 |

| | | | | | |
|---|---|---|---|---|---|
| CUSTOS | 2 | 2 | -0,19020524 | 0,050943909 | 0,728775831 |
| CKB | 30 | 29 | -0,055700832 | 0,051926578 | 0,731926921 |
| HK1 | 61 | 61 | 0,061590821 | 0,052154891 | 0,732349853 |
| COX7A2L | 5 | 5 | 0,121757844 | 0,052063479 | 0,732349853 |
| SLC6A9 | 4 | 3 | -0,168392055 | 0,052273444 | 0,732621737 |
| SH3KBP1 | 15 | 14 | -0,082406449 | 0,052644486 | 0,734695728 |
| EIF4H | 11 | 11 | -0,058123961 | 0,052561468 | 0,734695728 |
| MRPS10 | 2 | 2 | -0,140958747 | 0,052910774 | 0,734695728 |
| RAB27A | 2 | 2 | 0,123779045 | 0,052856219 | 0,734695728 |
| NTRK2 | 5 | 5 | -0,094256107 | 0,052946703 | 0,734695728 |
| WDR35 | 1 | 1 | 0,277111696 | 0,053018254 | 0,734695728 |
| RIMS1 | 29 | 29 | 0,085485247 | 0,05340684 | 0,737317151 |
| SNX27 | 18 | 17 | -0,042100475 | 0,053407078 | 0,737317151 |
| EPB41L3 | 17 | 17 | -0,087437735 | 0,053661024 | 0,739153747 |
| ALDH16A1 | 1 | 1 | -0,266264455 | 0,05374026 | 0,739153747 |
| SUCLG1 | 14 | 13 | 0,068261542 | 0,053948399 | 0,740313899 |
| SYT1 | 32 | 32 | 0,060783301 | 0,054025073 | 0,740313899 |
| IDH3B | 36 | 35 | 0,064496748 | 0,054282116 | 0,741086336 |
| DGKH | 18 | 18 | -0,134583078 | 0,054188038 | 0,741086336 |
| RAPGEF4 | 24 | 23 | 0,051222954 | 0,054552796 | 0,743407654 |
| NRXN1 | 26 | 26 | -0,04096526 | 0,056012973 | 0,757785409 |
| ATG5 | 4 | 4 | 0,097850268 | 0,056009748 | 0,757785409 |
| SERPINB12 | 1 | 1 | -0,279314977 | 0,055751845 | 0,757785409 |
| CYBC1 | 1 | 1 | 0,235589833 | 0,056018255 | 0,757785409 |
| OLFR1047 | 1 | 1 | 0,185511396 | 0,056218983 | 0,759110435 |
| PLXNA3 | 1 | 1 | 0,418129167 | 0,056346757 | 0,75944735 |
| VDAC1 | 23 | 23 | 0,089916951 | 0,05709125 | 0,766683584 |
| CEP120 | 1 | 1 | -0,22599683 | 0,057081782 | 0,766683584 |
| PACSIN1 | 43 | 42 | 0,035541097 | 0,058034487 | 0,768489221 |
| SRP72 | 7 | 7 | -0,071490211 | 0,057807223 | 0,768489221 |
| RAD23B | 18 | 17 | -0,067321591 | 0,057986008 | 0,768489221 |
| EPM2AIP1 | 22 | 22 | -0,077074147 | 0,057765429 | 0,768489221 |
| GLMN | 3 | 3 | -0,130118989 | 0,057591896 | 0,768489221 |

| | | | | | |
|---|---|---|---|---|---|
| PLXNA1;PLXNA2;PL | 7 | 7 | 0,068056014 | 0,05745123 | 0,768489221 |
| COPS5 | 21 | 21 | -0,057339879 | 0,057914408 | 0,768489221 |
| LAMTOR1 | 5 | 5 | 0,102512884 | 0,058058081 | 0,768489221 |
| MEGF9 | 1 | 1 | 0,38022607 | 0,058312635 | 0,77047786 |
| KRT18;KRT17 | 1 | 1 | -0,67733299 | 0,058521959 | 0,771862842 |
| VSNL1 | 23 | 23 | -0,110034731 | 0,058653197 | 0,772214818 |
| SUPT16H | 1 | 1 | 0,315202565 | 0,058818057 | 0,773007417 |
| SLC8A1 | 30 | 29 | 0,05312841 | 0,059636672 | 0,779604351 |
| CCAR1 | 1 | 1 | -0,397275238 | 0,059572648 | 0,779604351 |
| PTPN6 | 2 | 2 | 0,214394597 | 0,059466707 | 0,779604351 |
| HNRNPH2 | 6 | 6 | -0,069025414 | 0,059842616 | 0,780090745 |
| PGS1 | 8 | 7 | 0,084603301 | 0,059885114 | 0,780090745 |
| MYO15A | 2 | 2 | 0,144868188 | 0,060546583 | 0,785722567 |
| PAIP1 | 6 | 6 | -0,078214704 | 0,060636591 | 0,785722567 |
| UQCRQ | 9 | 9 | 0,106163086 | 0,060528071 | 0,785722567 |
| SF3B3 | 11 | 11 | -0,066248447 | 0,061002221 | 0,787696512 |
| NECAB1 | 10 | 9 | -0,142916936 | 0,060932437 | 0,787696512 |
| ELFN2 | 18 | 18 | 0,054668613 | 0,061518167 | 0,789636368 |
| IKZF3 | 1 | 1 | 0,145336242 | 0,061475496 | 0,789636368 |
| TIAL1 | 2 | 2 | -0,101860769 | 0,061376467 | 0,789636368 |
| ANKS1A | 2 | 1 | -0,366456436 | 0,06158009 | 0,789636368 |
| SYT7 | 3 | 2 | -0,099245269 | 0,061731567 | 0,79020685 |
| EEF1G | 20 | 20 | -0,074265098 | 0,062147891 | 0,79415973 |
| DLGAP1 | 9 | 9 | 0,080778533 | 0,062425543 | 0,794957002 |
| GRM8 | 3 | 2 | -0,262544931 | 0,06232797 | 0,794957002 |
| DPY19L4 | 1 | 1 | -0,198356877 | 0,06254105 | 0,794992109 |
| PHF24 | 15 | 15 | -0,051170591 | 0,06264357 | 0,794992109 |
| TMOD2 | 20 | 20 | -0,099445035 | 0,063264216 | 0,800119006 |
| ADGRB2 | 8 | 8 | 0,077773266 | 0,063212692 | 0,800119006 |
| GAN | 1 | 1 | -0,303824309 | 0,063480289 | 0,801479337 |
| UQCRC1 | 24 | 24 | 0,077272229 | 0,063819211 | 0,804383432 |
| KIAA1549 | 17 | 17 | 0,058797748 | 0,064334242 | 0,809493543 |
| DDX17;DDX5 | 1 | 1 | -0,145526148 | 0,064772415 | 0,813620846 |

| | | | | | |
|---|---|---|---|---|---|
| ERC2;ERC1 | 14 | 14 | 0,053147313 | 0,064960991 | 0,814604206 |
| PPP6R2 | 13 | 13 | -0,055155004 | 0,065150002 | 0,81558969 |
| UNC13A | 35 | 33 | 0,052012913 | 0,066077522 | 0,821630608 |
| SLC25A12 | 38 | 37 | 0,054648231 | 0,065871016 | 0,821630608 |
| STK32C | 5 | 5 | -0,089373016 | 0,065988161 | 0,821630608 |
| R3HDM2 | 5 | 5 | -0,08731773 | 0,065752128 | 0,821630608 |
| NDUFA12 | 15 | 15 | 0,067452514 | 0,066335547 | 0,823452695 |
| HSPA1L | 11 | 10 | 0,067197036 | 0,06648122 | 0,823876332 |
| CP | 9 | 9 | 0,176781585 | 0,067011359 | 0,827668731 |
| CCDC91 | 3 | 3 | -0,140650508 | 0,066911841 | 0,827668731 |
| DFFA | 1 | 1 | -0,17523893 | 0,06742738 | 0,831416738 |
| RASGRF2 | 15 | 14 | -0,098656189 | 0,067896222 | 0,832272162 |
| PLXND1 | 9 | 9 | -0,100898579 | 0,06782679 | 0,832272162 |
| PCBP2 | 5 | 5 | -0,084493904 | 0,067801607 | 0,832272162 |
| IGDCC4 | 1 | 1 | -0,295961376 | 0,067947484 | 0,832272162 |
| SPHK2 | 6 | 6 | 0,099289525 | 0,068365092 | 0,836000943 |
| PYCR3 | 6 | 6 | 0,08756902 | 0,069165012 | 0,841602605 |
| TNPO1;TNPO2 | 5 | 5 | -0,132025701 | 0,068990704 | 0,841602605 |
| ATP2B1 | 6 | 6 | 0,114199034 | 0,069063724 | 0,841602605 |
| EXOC5 | 13 | 13 | 0,052411639 | 0,069344551 | 0,842399436 |
| PTPRS | 1 | 1 | -0,270806049 | 0,069688836 | 0,845191691 |
| CSNK1D | 2 | 2 | -0,163504859 | 0,07002595 | 0,84703219 |
| EGFLAM | 1 | 1 | 0,190331671 | 0,070069952 | 0,84703219 |
| TPM1 | 8 | 8 | -0,112781845 | 0,07032502 | 0,848726467 |
| EIF3G | 6 | 6 | -0,10125729 | 0,070671459 | 0,848746984 |
| CNTN6 | 2 | 2 | 0,197410855 | 0,07045714 | 0,848746984 |
| TXNDC12 | 2 | 2 | 0,131344465 | 0,070665715 | 0,848746984 |
| ACSL1 | 22 | 21 | 0,04837207 | 0,071217426 | 0,851151955 |
| ZSCAN22 | 1 | 1 | -0,184379308 | 0,07118436 | 0,851151955 |
| NRXN2;NRXN1 | 3 | 3 | -0,146251935 | 0,07105993 | 0,851151955 |
| SART3 | 17 | 17 | -0,089335282 | 0,071795915 | 0,854976981 |
| TIMM29 | 11 | 10 | 0,079318992 | 0,071884742 | 0,854976981 |
| EVI2A | 1 | 1 | -0,201701964 | 0,071769722 | 0,854976981 |

| | | | | | |
|---|---|---|---|---|---|
| TARS3 | 24 | 24 | -0,040797724 | 0,073083708 | 0,858034204 |
| ISCA1 | 4 | 4 | 0,089396337 | 0,072287492 | 0,858034204 |
| SRRT | 5 | 5 | -0,089651956 | 0,073255983 | 0,858034204 |
| PREB | 6 | 6 | 0,090781568 | 0,073120823 | 0,858034204 |
| APOA1 | 18 | 18 | -0,291953783 | 0,072663875 | 0,858034204 |
| ME3;ME2 | 1 | 1 | 0,188266201 | 0,072979027 | 0,858034204 |
| SLC1A2 | 16 | 16 | -0,077897854 | 0,073419661 | 0,858034204 |
| SGSM1 | 11 | 11 | 0,069043037 | 0,072792456 | 0,858034204 |
| URI1 | 1 | 1 | 0,358673757 | 0,073346347 | 0,858034204 |
| CACNA1C;CACNA1I | 1 | 1 | 0,355327514 | 0,072941156 | 0,858034204 |
| PSME3 | 6 | 6 | -0,079703198 | 0,072864148 | 0,858034204 |
| OPA1 | 62 | 59 | 0,051732067 | 0,075125199 | 0,859644523 |
| VCL | 42 | 41 | -0,079390694 | 0,075398662 | 0,859644523 |
| GAMT | 5 | 5 | 0,092795575 | 0,075050087 | 0,859644523 |
| MDH2 | 24 | 24 | 0,062322768 | 0,075216091 | 0,859644523 |
| AIP | 9 | 9 | -0,063581187 | 0,075027272 | 0,859644523 |
| PPM1G | 15 | 15 | -0,08918057 | 0,075053391 | 0,859644523 |
| EPDR1 | 8 | 8 | 0,048572508 | 0,074874775 | 0,859644523 |
| EPN1 | 15 | 15 | -0,054867683 | 0,075418339 | 0,859644523 |
| EIF3H | 12 | 11 | -0,061239192 | 0,075419666 | 0,859644523 |
| CDK18 | 4 | 4 | -0,087596862 | 0,074485162 | 0,859644523 |
| PSMD13 | 20 | 20 | -0,060686825 | 0,074609039 | 0,859644523 |
| NDUFB7 | 8 | 8 | 0,067121163 | 0,074077039 | 0,859644523 |
| IMMT | 16 | 15 | 0,062142197 | 0,074312754 | 0,859644523 |
| RBFOX3;RBFOX1 | 5 | 4 | -0,077026081 | 0,074670023 | 0,859644523 |
| PSMD10 | 1 | 1 | 0,191223475 | 0,075103389 | 0,859644523 |
| SUMO2 | 1 | 1 | -0,183268405 | 0,07429199 | 0,859644523 |
| SV2B | 20 | 20 | 0,059259275 | 0,075758256 | 0,862173312 |
| NDUFA2 | 5 | 5 | 0,096522475 | 0,076056804 | 0,862911762 |
| CERS6 | 2 | 2 | 0,170967495 | 0,07601075 | 0,862911762 |
| VCAN | 19 | 18 | -0,110486152 | 0,07632488 | 0,863335324 |
| OSGEP | 6 | 6 | -0,076534196 | 0,076327913 | 0,863335324 |
| DDX6 | 14 | 14 | -0,044522876 | 0,076703095 | 0,866252384 |

| | | | | | |
|---|---|---|---|---|---|
| ARL3 | 12 | 12 | 0,063663842 | 0,077239921 | 0,870983297 |
| RAB11A | 1 | 1 | -0,193085051 | 0,077754466 | 0,875448917 |
| KIF5C;KIF5A;KIF5B | 11 | 11 | 0,055144546 | 0,077942867 | 0,876234422 |
| RPL22 | 4 | 4 | -0,07383525 | 0,078562824 | 0,881861725 |
| UCHL5 | 5 | 5 | -0,089810945 | 0,07870713 | 0,882140912 |
| MTMR12 | 6 | 6 | -0,081428086 | 0,079024454 | 0,884355478 |
| GRK6 | 4 | 4 | -0,078285892 | 0,079410326 | 0,887329295 |
| RPL7 | 16 | 15 | -0,064273897 | 0,079857217 | 0,889645209 |
| MINDY2 | 3 | 3 | -0,126637623 | 0,079858485 | 0,889645209 |
| EEFSEC | 3 | 3 | 0,100969187 | 0,08028539 | 0,891711113 |
| LPCAT1 | 1 | 1 | -0,345327964 | 0,0802197 | 0,891711113 |
| PJA1 | 1 | 1 | 0,179005158 | 0,080424532 | 0,891915307 |
| KIF2A | 2 | 2 | 0,133646222 | 0,08070984 | 0,893737453 |
| CCDC177 | 19 | 18 | -0,054299596 | 0,081728199 | 0,89898014 |
| EHD4 | 11 | 11 | -0,056948392 | 0,082130755 | 0,89898014 |
| NDUFC2 | 8 | 8 | 0,062042822 | 0,081990278 | 0,89898014 |
| PHKA1 | 9 | 9 | -0,072454594 | 0,081574976 | 0,89898014 |
| CTTN | 27 | 27 | -0,07345718 | 0,082142856 | 0,89898014 |
| DDX5 | 6 | 6 | -0,083281405 | 0,081310568 | 0,89898014 |
| PIP4P1 | 2 | 2 | 0,125403186 | 0,082156999 | 0,89898014 |
| ERICH5 | 1 | 1 | -0,213419765 | 0,082117778 | 0,89898014 |
| PLXNA4 | 41 | 41 | 0,03071944 | 0,082603279 | 0,900624597 |
| ELP1 | 15 | 15 | -0,042815929 | 0,083010547 | 0,900624597 |
| DLG2 | 28 | 28 | 0,029130948 | 0,083038905 | 0,900624597 |
| CAMK2A;CAMK2B | 1 | 1 | 0,49460649 | 0,083015951 | 0,900624597 |
| CHMP1B1;CHMP1E | 2 | 2 | -0,15014213 | 0,082982707 | 0,900624597 |
| DRP2 | 1 | 1 | -0,473046337 | 0,082616013 | 0,900624597 |
| CHORDC1 | 13 | 13 | 0,06405948 | 0,083281012 | 0,901926032 |
| DMXL1 | 15 | 14 | 0,080814812 | 0,083791137 | 0,906122016 |
| GUCY1A2 | 9 | 9 | -0,064078411 | 0,085265644 | 0,906146834 |
| GPHN | 25 | 25 | -0,037461027 | 0,084110667 | 0,906146834 |
| SLC8A2 | 31 | 31 | 0,030020886 | 0,084734361 | 0,906146834 |
| CACNA1B | 14 | 14 | 0,056726644 | 0,084269479 | 0,906146834 |

| | | | | | |
|---|---|---|---|---|---|
| ATP5F1E | 3 | 3 | 0,09805771 | 0,084618275 | 0,906146834 |
| PEF1 | 6 | 6 | -0,084490297 | 0,085015725 | 0,906146834 |
| CERT1 | 7 | 7 | -0,082086302 | 0,08506928 | 0,906146834 |
| KPNA1 | 8 | 8 | -0,058762688 | 0,084533572 | 0,906146834 |
| MOB1B;MOB1A | 2 | 2 | -0,11965355 | 0,084959146 | 0,906146834 |
| MYD88 | 1 | 1 | 0,125366186 | 0,084032803 | 0,906146834 |
| CHRM2 | 1 | 1 | -0,145965078 | 0,085060902 | 0,906146834 |
| KCTD10 | 1 | 1 | -0,283887589 | 0,085252976 | 0,906146834 |
| RBM3 | 6 | 6 | -0,221402959 | 0,085804002 | 0,909251586 |
| JCAD | 1 | 1 | 0,180977516 | 0,08577722 | 0,909251586 |
| HNRNPAB | 8 | 8 | -0,108541435 | 0,086093492 | 0,909625198 |
| ATXN7L2 | 1 | 1 | -0,259671237 | 0,086085904 | 0,909625198 |
| NUDT16L1 | 1 | 1 | -0,136479625 | 0,086208724 | 0,909625198 |
| SERPINI1 | 6 | 6 | -0,115556647 | 0,086365084 | 0,909975054 |
| AP3B2 | 50 | 49 | 0,034754618 | 0,08831366 | 0,912759526 |
| NIF3L1 | 11 | 11 | 0,070252074 | 0,086971885 | 0,912759526 |
| APRT | 7 | 7 | 0,062624052 | 0,088870932 | 0,912759526 |
| SRSF7 | 3 | 3 | -0,21275774 | 0,088443819 | 0,912759526 |
| EEF2 | 46 | 45 | -0,07178514 | 0,08896229 | 0,912759526 |
| HECTD1 | 21 | 21 | -0,040489168 | 0,087680415 | 0,912759526 |
| TPM3 | 11 | 11 | -0,092897868 | 0,08826854 | 0,912759526 |
| ISOC1 | 9 | 9 | -0,127017736 | 0,088717748 | 0,912759526 |
| DBN1 | 34 | 34 | -0,070698183 | 0,088415171 | 0,912759526 |
| COG3 | 2 | 2 | -0,11996877 | 0,088679469 | 0,912759526 |
| RHOB | 9 | 9 | 0,068797018 | 0,087611668 | 0,912759526 |
| PFDN2 | 8 | 8 | -0,057268384 | 0,088465993 | 0,912759526 |
| RWDD1 | 4 | 4 | -0,072713301 | 0,088040759 | 0,912759526 |
| NDUFS5 | 8 | 8 | 0,071961115 | 0,088977371 | 0,912759526 |
| UBA5 | 15 | 14 | -0,039585934 | 0,0870733 | 0,912759526 |
| CLDN1 | 1 | 1 | -0,221535706 | 0,088276236 | 0,912759526 |
| CLIC6;CLIC4 | 1 | 1 | -0,661858257 | 0,088849782 | 0,912759526 |
| FAM136A | 2 | 2 | 0,116655009 | 0,087362243 | 0,912759526 |
| PPP1CC | 2 | 2 | -0,067616651 | 0,088245903 | 0,912759526 |

| | | | | | |
|---|---|---|---|---|---|
| GNA13 | 19 | 19 | -0,040173512 | 0,089615744 | 0,918033131 |
| NF1 | 13 | 13 | -0,05884215 | 0,090004644 | 0,918196546 |
| SCN1A;SCN3A;SCN2 | 4 | 4 | 0,094733573 | 0,08977546 | 0,918196546 |
| UBL7 | 2 | 2 | -0,113565853 | 0,089937552 | 0,918196546 |
| REPS2 | 15 | 15 | -0,074926759 | 0,090402993 | 0,920988282 |
| CAMK2A | 9 | 9 | 0,085976405 | 0,090634301 | 0,921884631 |
| SPRYD4 | 7 | 7 | -0,064037589 | 0,090740607 | 0,921884631 |
| RAP2B | 7 | 7 | 0,067498699 | 0,091449208 | 0,925265553 |
| PALM | 1 | 1 | -0,145924719 | 0,091418255 | 0,925265553 |
| LIN9 | 1 | 1 | 0,34639678 | 0,091395483 | 0,925265553 |
| GOT2 | 39 | 38 | 0,06310949 | 0,091591014 | 0,925432602 |
| NCKIPSD | 34 | 32 | -0,039043745 | 0,091904972 | 0,926071107 |
| PPP4R3A | 8 | 8 | -0,065300386 | 0,091808541 | 0,926071107 |
| NAMPT | 18 | 18 | -0,06689554 | 0,092072237 | 0,926492568 |
| DHRS7B | 11 | 9 | 0,076989761 | 0,092402092 | 0,927285122 |
| ARL6 | 7 | 7 | 0,084986221 | 0,092359099 | 0,927285122 |
| PRPSAP2 | 18 | 18 | -0,060395465 | 0,092610075 | 0,928111279 |
| UQCRC2 | 25 | 25 | 0,064009938 | 0,093138314 | 0,929849514 |
| PTGR3 | 11 | 11 | 0,071338777 | 0,093161202 | 0,929849514 |
| PSMF1 | 4 | 4 | -0,096135678 | 0,093156461 | 0,929849514 |
| RYR2 | 77 | 76 | 0,050422167 | 0,093345721 | 0,930203585 |
| AS3MT | 7 | 7 | -0,076888027 | 0,093450984 | 0,930203585 |
| sp\|P01680\|KV4A1_ | 1 | 1 | 0,331420045 | 0,093574501 | 0,930203585 |
| COPE | 4 | 4 | -0,111391312 | 0,09408298 | 0,934001196 |
| EIF3D | 14 | 14 | -0,062587002 | 0,094328283 | 0,935179461 |
| BCAT1 | 11 | 10 | 0,0707995 | 0,094502533 | 0,935651086 |
| FHIP1B | 8 | 8 | -0,06409564 | 0,094937882 | 0,938153013 |
| SERPINA1B;SERPIN | 4 | 4 | -0,213146152 | 0,095009268 | 0,938153013 |
| SCN8A | 5 | 5 | 0,108797605 | 0,095248081 | 0,939255442 |
| EIF3C | 28 | 28 | -0,038641503 | 0,095442338 | 0,939916144 |
| PSD3 | 34 | 34 | 0,031464323 | 0,096209421 | 0,941355705 |
| GCAT | 6 | 6 | -0,152661133 | 0,095717084 | 0,941355705 |
| PSMD5 | 15 | 15 | -0,048279101 | 0,096225773 | 0,941355705 |

| | | | | | |
|---|---|---|---|---|---|
| NECAB2 | 15 | 15 | 0,067465679 | 0,095881967 | 0,941355705 |
| PPP1R14C | 1 | 1 | -0,274689414 | 0,096040612 | 0,941355705 |
| SGTB | 6 | 5 | -0,103054061 | 0,09656697 | 0,943443964 |
| TMEM165 | 3 | 3 | 0,188357613 | 0,097131863 | 0,945212039 |
| NHSL2 | 10 | 10 | -0,078534646 | 0,096987412 | 0,945212039 |
| NDUFA6 | 9 | 8 | 0,059661187 | 0,097093064 | 0,945212039 |
| CLPTM1 | 12 | 12 | -0,057823985 | 0,097561379 | 0,947497387 |
| ARHGEF12 | 19 | 19 | -0,049634624 | 0,097623275 | 0,947497387 |
| ITPKA | 27 | 25 | -0,067551129 | 0,098160613 | 0,951462317 |
| AK5 | 19 | 19 | 0,068830596 | 0,098596903 | 0,953937917 |
| RANBP3 | 15 | 15 | -0,045965741 | 0,09880348 | 0,953937917 |
| TDP2 | 1 | 1 | -0,278182924 | 0,098778387 | 0,953937917 |
| PPFIA4 | 25 | 25 | 0,051472549 | 0,0991954 | 0,956471573 |
| SRCIN1 | 68 | 65 | 0,038414754 | 0,101621289 | 0,956573486 |
| TUFM | 34 | 32 | 0,055806602 | 0,101666692 | 0,956573486 |
| CRACDL | 20 | 20 | -0,080911858 | 0,099839508 | 0,956573486 |
| ACTN4 | 16 | 16 | -0,057492983 | 0,099761837 | 0,956573486 |
| MGLL | 20 | 20 | -0,055205691 | 0,101576531 | 0,956573486 |
| MAST4 | 2 | 2 | 0,161404236 | 0,101173899 | 0,956573486 |
| PPP1R1B | 5 | 5 | -0,146982972 | 0,101194701 | 0,956573486 |
| ISLR2 | 7 | 7 | 0,071931165 | 0,100239685 | 0,956573486 |
| CFAP418 | 3 | 3 | -0,101483149 | 0,099474127 | 0,956573486 |
| CCDC85A | 6 | 6 | -0,114150472 | 0,100299065 | 0,956573486 |
| RTCB | 21 | 20 | -0,053885961 | 0,100838445 | 0,956573486 |
| CORO2A | 8 | 8 | 0,058660659 | 0,101573377 | 0,956573486 |
| CIAO3 | 4 | 4 | -0,094867353 | 0,100873259 | 0,956573486 |
| GGA1 | 11 | 11 | -0,111088095 | 0,10055899 | 0,956573486 |
| ADCYAP1R1 | 3 | 3 | -0,121560503 | 0,101396698 | 0,956573486 |
| MYH10;MYH14 | 2 | 2 | 0,105153056 | 0,100574205 | 0,956573486 |
| UBAP1 | 1 | 1 | 0,458431796 | 0,099593542 | 0,956573486 |
| NWD1 | 1 | 1 | -0,202549548 | 0,100737636 | 0,956573486 |
| MYO1A;MYO1B | 1 | 1 | -0,179246642 | 0,100726106 | 0,956573486 |
| LETM1 | 31 | 29 | 0,050697749 | 0,10190864 | 0,957630043 |

| | | | | | |
|---|---|---|---|---|---|
| EIF3I | 11 | 11 | -0,063937559 | 0,102381716 | 0,960524427 |
| SBF2 | 2 | 2 | 0,16095299 | 0,102476746 | 0,960524427 |
| PRKCG | 50 | 49 | 0,052105524 | 0,103453761 | 0,968453082 |
| MCAT | 7 | 7 | 0,077556358 | 0,103854521 | 0,970974045 |
| TUBA8 | 10 | 10 | 0,086156586 | 0,104268418 | 0,972556869 |
| sp\|Q9CWU4\|CA05 | 1 | 1 | -0,172587566 | 0,10428717 | 0,972556869 |
| PPFIA2 | 1 | 1 | 0,153394423 | 0,104569678 | 0,973961716 |
| CDS2 | 7 | 7 | -0,064840646 | 0,104964446 | 0,97513101 |
| HNRNPH3 | 1 | 1 | 0,198320895 | 0,105136314 | 0,97513101 |
| SLC25A42 | 8 | 8 | 0,064043124 | 0,105223316 | 0,97513101 |
| CDK17 | 5 | 5 | -0,073502057 | 0,104968588 | 0,97513101 |
| RYR2 | 5 | 5 | 0,101143098 | 0,105669706 | 0,978040665 |
| ALDOART2 | 14 | 14 | 0,06368464 | 0,106867292 | 0,98461906 |
| SLC4A3 | 9 | 9 | 0,072496743 | 0,106540026 | 0,98461906 |
| LRRTM1 | 9 | 9 | 0,073423478 | 0,106758885 | 0,98461906 |
| POLR2M | 2 | 2 | -0,128001955 | 0,106989068 | 0,98461906 |
| MOB2 | 1 | 1 | -0,131506054 | 0,107046995 | 0,98461906 |
| TAGLN3 | 18 | 18 | -0,041292291 | 0,107314886 | 0,984630744 |
| ERMP1 | 8 | 8 | -0,058563699 | 0,107271369 | 0,984630744 |
| ATP5F1C | 20 | 19 | 0,054742088 | 0,107702567 | 0,98696174 |
| SYN3 | 14 | 14 | -0,046846254 | 0,107859584 | 0,987175817 |
| IGBP1B | 1 | 1 | 0,165650813 | 0,108255355 | 0,987811117 |
| TMED7 | 3 | 3 | -0,088695848 | 0,108281094 | 0,987811117 |
| KCTD8;KCTD16 | 1 | 1 | 0,335017891 | 0,108463961 | 0,987811117 |
| SMPDL3B | 3 | 3 | -0,123656627 | 0,108390455 | 0,987811117 |
| DNM1 | 4 | 4 | 0,099222837 | 0,108608053 | 0,987905271 |
| CIAO2A | 1 | 1 | 0,178334465 | 0,109122048 | 0,991359713 |
| SLC9A7 | 7 | 7 | 0,061595066 | 0,109366412 | 0,992359116 |
| TACO1 | 5 | 5 | 0,135361193 | 0,109672036 | 0,993911238 |
| PSMC1 | 29 | 29 | -0,059655517 | 0,110127948 | 0,99416081 |
| RAB1B | 9 | 9 | -0,080393591 | 0,110346979 | 0,99416081 |
| GOLM2 | 3 | 3 | 0,197704809 | 0,110146982 | 0,99416081 |
| MAP4K4 | 1 | 1 | 0,282419934 | 0,110372578 | 0,99416081 |

| | | | | | |
|---|---|---|---|---|---|
| LRBA | 1 | 1 | -0,138282912 | 0,11009167 | 0,99416081 |
| ICAM5 | 24 | 24 | -0,041042074 | 0,111080081 | 0,995381596 |
| CDK12;CDK13 | 2 | 2 | -0,256526941 | 0,111043566 | 0,995381596 |
| NLRX1 | 8 | 5 | 0,11587384 | 0,110979269 | 0,995381596 |
| KIF20B | 1 | 1 | 0,347528671 | 0,111181941 | 0,995381596 |
| WDSUB1 | 1 | 1 | 0,137519266 | 0,110890033 | 0,995381596 |
| CCDC9 | 2 | 2 | -0,13933595 | 0,111561241 | 0,997568188 |
| DDRGK1 | 3 | 3 | -2,45118E-17 | 0,999999993 | 1 |
| UAP1L1 | 18 | 18 | -5,92428E-16 | 0,999999948 | 1 |
| PSMD4 | 3 | 2 | -7,10047E-16 | 0,999999927 | 1 |
| PSMD4 | 1 | 1 | 0 | 1 | 1 |
| MRPS7 | 5 | 5 | 0 | 1 | 1 |
| SRGAP3 | 43 | 43 | 4,20824E-18 | 0,999999995 | 1 |
| ZFPL1 | 1 | 1 | -0,172360298 | 0,310592201 | 1 |
| EDC4 | 9 | 9 | -0,078450588 | 0,136211413 | 1 |
| PFDN4 | 4 | 4 | 0 | 1 | 1 |
| SEPTIN6 | 10 | 10 | 0 | 1 | 1 |
| FH | 29 | 28 | 0,028673304 | 0,35542988 | 1 |
| PIGS | 7 | 6 | 0 | 1 | 1 |
| RUFY1 | 7 | 7 | 0 | 1 | 1 |
| PCLO | 170 | 160 | 0,029355166 | 0,403954013 | 1 |
| ACBD6 | 6 | 6 | 3,34993E-17 | 0,999999991 | 1 |
| SEMA7A | 8 | 8 | 0 | 1 | 1 |
| STIP1 | 55 | 55 | 0 | 1 | 1 |
| TSC1 | 17 | 17 | 0 | 1 | 1 |
| ANK3 | 38 | 38 | 0 | 1 | 1 |
| CTNND1 | 18 | 18 | 0 | 1 | 1 |
| CHCHD3 | 14 | 13 | 2,83301E-16 | 0,99999995 | 1 |
| GOLGB1 | 4 | 3 | -0,08741343 | 0,318979235 | 1 |
| SEPTIN11 | 11 | 11 | -0,004683509 | 0,680279384 | 1 |
| PNP | 22 | 21 | -1,86576E-17 | 0,999999992 | 1 |
| PKP4 | 34 | 31 | 0,002282692 | 0,744519074 | 1 |
| TTC4 | 2 | 2 | 0 | 1 | 1 |

| | | | | | |
|---|---|---|---|---|---|
| RAB3GAP2 | 32 | 32 | 0 | 1 | 1 |
| ITM2C | 5 | 5 | 0,057781181 | 0,123685165 | 1 |
| CAMSAP3 | 14 | 14 | 2,49253E-17 | 0,999999983 | 1 |
| PDHA1 | 37 | 36 | 0,047561436 | 0,129912228 | 1 |
| NDUFA6 | 1 | 1 | -0,075218865 | 0,620876182 | 1 |
| CASKIN1 | 52 | 51 | 0 | 1 | 1 |
| CEP170B | 35 | 35 | -7,37286E-20 | 1 | 1 |
| ACOX1 | 15 | 14 | -1,09617E-18 | 0,999999998 | 1 |
| CAMKV | 25 | 25 | -6,17281E-16 | 0,999999931 | 1 |
| SND1 | 37 | 37 | -0,020162373 | 0,42751023 | 1 |
| PANK2 | 5 | 5 | 1,83628E-15 | 0,999999852 | 1 |
| BCR | 22 | 22 | 0 | 1 | 1 |
| RAB3GAP1 | 23 | 22 | 7,72343E-18 | 0,999999994 | 1 |
| EXOC8 | 20 | 20 | -2,3411E-18 | 0,999999998 | 1 |
| PSMD9 | 9 | 9 | -0,011120215 | 0,545052545 | 1 |
| FMNL2;FMNL3 | 4 | 4 | 0 | 1 | 1 |
| HDLBP | 21 | 21 | -1,70651E-17 | 0,999999987 | 1 |
| UBE3A | 26 | 25 | -1,55901E-16 | 0,999999953 | 1 |
| MICU1 | 15 | 13 | 1,41349E-17 | 0,99999999 | 1 |
| TPM3 | 3 | 3 | -0,042330105 | 0,28645319 | 1 |
| ADK | 17 | 17 | -0,093783495 | 0,147235911 | 1 |
| PACSIN2 | 15 | 15 | -2,27883E-16 | 0,999999952 | 1 |
| VPS52 | 24 | 24 | -0,00972284 | 0,552004831 | 1 |
| RABL6 | 17 | 17 | 3,54112E-17 | 0,999999983 | 1 |
| LMTK3 | 9 | 9 | -5,06927E-17 | 0,999999992 | 1 |
| DGLUCY | 8 | 8 | 2,07317E-17 | 0,999999991 | 1 |
| CAND1 | 44 | 43 | -0,031583849 | 0,232999178 | 1 |
| ATP2C1 | 8 | 8 | 0 | 1 | 1 |
| ALB | 40 | 40 | 0 | 1 | 1 |
| SORL1 | 23 | 23 | 3,81775E-16 | 0,999999936 | 1 |
| AP2A1 | 45 | 45 | 5,22978E-16 | 0,999999914 | 1 |
| ADD2 | 36 | 36 | -0,025698267 | 0,454728358 | 1 |
| ALDH4A1 | 13 | 12 | 1,08234E-15 | 0,999999905 | 1 |

| | | | | | |
|---|---|---|---|---|---|
| PPP1R12C | 10 | 10 | -2,19884E-16 | 0,999999958 | 1 |
| YBX1 | 3 | 3 | -0,090731795 | 0,159706135 | 1 |
| PAG1 | 2 | 2 | 0,09031723 | 0,128144549 | 1 |
| PKM | 52 | 52 | 0 | 1 | 1 |
| PRKRA | 6 | 6 | -1,9527E-18 | 0,999999997 | 1 |
| MAP1B | 109 | 105 | -4,81086E-16 | 0,999999925 | 1 |
| ANKFY1 | 13 | 13 | 0,004059488 | 0,733820955 | 1 |
| SPG7 | 12 | 9 | -6,49471E-18 | 0,999999994 | 1 |
| NCKAP1 | 69 | 68 | 1,34655E-16 | 0,999999943 | 1 |
| AAK1 | 37 | 37 | 0 | 1 | 1 |
| TAGLN | 10 | 9 | -9,88037E-18 | 0,999999998 | 1 |
| PLEC | 194 | 188 | 1,22222E-15 | 0,999999937 | 1 |
| CLTA | 1 | 1 | 0 | 1 | 1 |
| MTHFD1 | 34 | 34 | 0 | 1 | 1 |
| MAP7D1 | 15 | 15 | -0,025827274 | 0,394707132 | 1 |
| CENPE | 1 | 1 | 0 | 1 | 1 |
| MARCHF6 | 1 | 1 | 0 | 1 | 1 |
| MAP4 | 24 | 24 | -0,033358091 | 0,475154231 | 1 |
| CNNM1 | 16 | 16 | -2,29207E-15 | 0,99999983 | 1 |
| RELCH | 2 | 2 | 0 | 1 | 1 |
| SPTB | 51 | 48 | -1,38229E-16 | 0,999999968 | 1 |
| PIK3CB | 1 | 1 | 0 | 1 | 1 |
| PPP1R16B | 1 | 1 | 0 | 1 | 1 |
| 2210016F16RIK | 8 | 8 | -9,08399E-17 | 0,999999979 | 1 |
| SPTBN2 | 114 | 111 | -2,40461E-15 | 0,999999869 | 1 |
| PDE2A | 39 | 39 | 0,011827108 | 0,507436563 | 1 |
| ECI3;ECI2 | 4 | 4 | -0,013375117 | 0,618696502 | 1 |
| FKBP4 | 32 | 32 | -0,00543152 | 0,719652446 | 1 |
| PCDHB14;PCDHB6 | 1 | 1 | 0 | 1 | 1 |
| KSR1 | 20 | 20 | 7,91058E-13 | 0,999996253 | 1 |
| NEMF | 4 | 4 | 0 | 1 | 1 |
| TXNDC15 | 2 | 2 | 4,64588E-17 | 0,999999991 | 1 |
| MYH14 | 42 | 41 | -0,028891893 | 0,125554281 | 1 |

| | | | | | |
|---|---|---|---|---|---|
| COPB2 | 16 | 16 | -6,10153E-17 | 0,999999972 | 1 |
| NDUFA7 | 16 | 16 | 5,27342E-17 | 0,999999984 | 1 |
| MRPS35 | 6 | 4 | 3,33361E-16 | 0,999999966 | 1 |
| RFTN1 | 5 | 4 | -5,70106E-18 | 0,999999994 | 1 |
| ABCB10 | 10 | 9 | 0,056336405 | 0,194190099 | 1 |
| MAP4 | 21 | 21 | -0,069077186 | 0,153448883 | 1 |
| CLIP2 | 42 | 42 | -0,00748849 | 0,675151847 | 1 |
| YWHAE | 35 | 35 | 1,35125E-16 | 0,999999984 | 1 |
| SDHA | 29 | 26 | 6,35849E-17 | 0,999999971 | 1 |
| ITGA3 | 1 | 1 | 0 | 1 | 1 |
| VPS51 | 32 | 32 | -1,98443E-17 | 0,999999996 | 1 |
| METTL13 | 3 | 3 | 2,8086E-16 | 0,999999962 | 1 |
| ALDH2 | 27 | 26 | 1,37925E-15 | 0,999999924 | 1 |
| COMMD3 | 3 | 3 | 0,077676217 | 0,180802631 | 1 |
| NWD2 | 23 | 22 | 0,004911465 | 0,753476216 | 1 |
| CCDC124 | 3 | 3 | 0 | 1 | 1 |
| CTSA | 6 | 6 | 0 | 1 | 1 |
| SRM | 12 | 12 | 0 | 1 | 1 |
| IQGAP2 | 45 | 45 | 0 | 1 | 1 |
| CNP | 34 | 34 | -8,90762E-16 | 0,999999958 | 1 |
| FRMD4A | 6 | 6 | 4,27997E-17 | 0,999999989 | 1 |
| PPP1R9B | 35 | 34 | -0,007901516 | 0,671528932 | 1 |
| NNT | 30 | 26 | 0,058336775 | 0,429077929 | 1 |
| BCL2L2 | 3 | 3 | -8,32986E-18 | 0,999999996 | 1 |
| HMGN5 | 5 | 5 | -0,077801593 | 0,280156893 | 1 |
| ATP6V1D | 21 | 21 | 4,85938E-16 | 0,999999921 | 1 |
| GART | 26 | 26 | 0 | 1 | 1 |
| OTUD7A | 4 | 4 | -2,30492E-10 | 0,999957005 | 1 |
| FBXO41 | 18 | 17 | 1,30275E-16 | 0,999999962 | 1 |
| RAP1GAP | 7 | 6 | -0,001504055 | 0,857926681 | 1 |
| RAP1GAP | 16 | 16 | -0,049017259 | 0,2576218 | 1 |
| PICALM | 6 | 6 | -0,038582949 | 0,353312629 | 1 |
| IGBP1 | 4 | 4 | -2,83622E-18 | 0,999999997 | 1 |

| | | | | | |
|---|---|---|---|---|---|
| AJM1 | 20 | 16 | 0,03353734 | 0,389817925 | 1 |
| RRBP1 | 20 | 20 | 1,07898E-17 | 0,999999993 | 1 |
| CTNND2 | 40 | 38 | 0 | 1 | 1 |
| MACROD1 | 4 | 4 | 0 | 1 | 1 |
| JAGN1 | 2 | 2 | -3,85354E-18 | 0,999999997 | 1 |
| OXCT1 | 31 | 31 | 5,57972E-17 | 0,999999985 | 1 |
| RPLP1 | 2 | 2 | 0 | 1 | 1 |
| RUVBL2 | 21 | 19 | 0 | 1 | 1 |
| PCBD2 | 2 | 2 | -2,87572E-19 | 0,999999999 | 1 |
| DUT | 2 | 2 | -7,71906E-16 | 0,99999995 | 1 |
| RAB11B | 3 | 3 | -9,22356E-13 | 0,999996904 | 1 |
| NCAM1 | 35 | 35 | -3,97625E-18 | 0,999999999 | 1 |
| CCT2 | 47 | 47 | -1,48703E-16 | 0,999999948 | 1 |
| STXBP1 | 60 | 59 | 0 | 1 | 1 |
| ANKIB1 | 2 | 2 | 0 | 1 | 1 |
| MAT2A | 20 | 20 | 0 | 1 | 1 |
| TNR | 42 | 41 | -0,003769543 | 0,762922306 | 1 |
| TDRKH | 15 | 14 | 3,11784E-17 | 0,999999984 | 1 |
| NPRL3 | 2 | 2 | -7,81466E-16 | 0,999999936 | 1 |
| PRMT5 | 11 | 11 | 0 | 1 | 1 |
| TTC39B | 3 | 3 | -0,102306762 | 0,119219772 | 1 |
| MIEF1 | 1 | 1 | 0 | 1 | 1 |
| EMC1 | 22 | 21 | -2,70239E-15 | 0,999999819 | 1 |
| EXOC3 | 13 | 13 | 0,0019546 | 0,815371587 | 1 |
| RYR2 | 6 | 6 | 0 | 1 | 1 |
| ACP6 | 6 | 6 | 4,60446E-17 | 0,999999982 | 1 |
| GPR158 | 36 | 35 | 3,3623E-16 | 0,999999949 | 1 |
| SLC27A1;SLC27A4 | 2 | 2 | 0 | 1 | 1 |
| ITPR1 | 39 | 38 | 0,007668629 | 0,621434112 | 1 |
| NCAN | 34 | 34 | -0,043793779 | 0,245245654 | 1 |
| DPYSL3 | 31 | 29 | 1,35307E-15 | 0,99999987 | 1 |
| CRMP1 | 31 | 31 | 0 | 1 | 1 |
| DPYSL2 | 44 | 44 | 0 | 1 | 1 |

| | | | | | |
|---|---|---|---|---|---|
| CARS2 | 6 | 6 | 0 | 1 | 1 |
| SHANK3 | 46 | 42 | 0,038129353 | 0,166011922 | 1 |
| KIF1A | 49 | 48 | 0 | 1 | 1 |
| QDPR | 12 | 12 | 0 | 1 | 1 |
| CELF2 | 10 | 9 | -0,057029042 | 0,336210166 | 1 |
| CELF1 | 5 | 5 | 0 | 1 | 1 |
| HARS1 | 16 | 16 | 0 | 1 | 1 |
| ACAP2 | 13 | 13 | -5,32226E-17 | 0,999999981 | 1 |
| EEA1 | 52 | 51 | -0,03863815 | 0,452474158 | 1 |
| CPLX2 | 6 | 6 | -9,77371E-16 | 0,999999944 | 1 |
| MTOR | 45 | 44 | 3,56585E-16 | 0,999999918 | 1 |
| PRXL2A | 9 | 9 | -4,52707E-16 | 0,999999945 | 1 |
| PTPRF;PTPRD | 5 | 5 | 5,26114E-17 | 0,999999979 | 1 |
| TRHDE | 9 | 9 | 0,02880892 | 0,375920699 | 1 |
| CCDC43 | 5 | 5 | -3,65252E-16 | 0,99999996 | 1 |
| UBE2O | 36 | 36 | 1,97594E-18 | 1 | 1 |
| CALCOCO1 | 14 | 14 | 0 | 1 | 1 |
| EPRS1 | 53 | 52 | 1,7674E-16 | 0,999999974 | 1 |
| ELAC2 | 5 | 5 | 0,04768825 | 0,263016316 | 1 |
| VMA21 | 1 | 1 | 0,073877682 | 0,239739479 | 1 |
| AMPH | 21 | 21 | -1,78679E-16 | 0,999999976 | 1 |
| HUWE1 | 54 | 54 | -2,18588E-16 | 0,99999993 | 1 |
| PRR36 | 14 | 14 | -0,02376558 | 0,558954617 | 1 |
| HTT | 51 | 51 | 0 | 1 | 1 |
| PTPN23 | 20 | 20 | -1,24238E-16 | 0,999999969 | 1 |
| FOXO3 | 1 | 1 | 0 | 1 | 1 |
| CHMP4B | 9 | 9 | -0,044691408 | 0,261132457 | 1 |
| APOB | 1 | 1 | 0 | 1 | 1 |
| KIF2A | 29 | 29 | 0,02688208 | 0,216941109 | 1 |
| RPS25 | 5 | 5 | -0,071166429 | 0,126389958 | 1 |
| ASPSCR1 | 9 | 9 | 0 | 1 | 1 |
| PITPNM1 | 25 | 25 | -1,9486E-17 | 0,999999983 | 1 |
| PPP5C | 25 | 24 | 3,90808E-16 | 0,999999921 | 1 |

| | | | | | |
|---|---|---|---|---|---|
| LRP1 | 52 | 51 | 9,73953E-18 | 0,999999991 | 1 |
| HADHB | 26 | 24 | 1,14338E-17 | 0,999999996 | 1 |
| XPO7 | 34 | 34 | -0,02714182 | 0,315485496 | 1 |
| MAST1 | 13 | 13 | 0 | 1 | 1 |
| COPS2 | 31 | 30 | -2,81304E-16 | 0,999999944 | 1 |
| TUBB6 | 7 | 7 | 0 | 1 | 1 |
| SLC25A27 | 10 | 10 | 7,27904E-19 | 0,999999999 | 1 |
| PRMT8 | 10 | 9 | 0 | 1 | 1 |
| TLN1 | 40 | 39 | 4,9241E-17 | 0,999999987 | 1 |
| ILVBL | 5 | 5 | 0,051543379 | 0,336532632 | 1 |
| JUP | 16 | 14 | 0,019484618 | 0,721647182 | 1 |
| AP1B1;AP2B1 | 33 | 33 | 0 | 1 | 1 |
| FASN | 83 | 82 | 0 | 1 | 1 |
| SPTBN1 | 208 | 204 | -4,70769E-16 | 0,999999967 | 1 |
| CPT1C | 11 | 11 | 0 | 1 | 1 |
| WARS1 | 33 | 33 | 8,11569E-17 | 0,999999978 | 1 |
| ABCF1 | 8 | 8 | 2,07771E-17 | 0,999999984 | 1 |
| CDV3 | 6 | 6 | -0,102013882 | 0,127525657 | 1 |
| ACAA2 | 27 | 25 | 0 | 1 | 1 |
| LSAMP | 14 | 14 | 7,65956E-18 | 0,999999996 | 1 |
| CAMK4 | 13 | 13 | -1,3445E-17 | 0,999999991 | 1 |
| PSMA2 | 12 | 12 | 3,38069E-17 | 0,999999989 | 1 |
| ITGB8 | 8 | 8 | -0,046950736 | 0,174608132 | 1 |
| WIPI2 | 6 | 6 | 0,041799794 | 0,191158485 | 1 |
| CYC1 | 17 | 16 | 0,044359733 | 0,16513946 | 1 |
| THUMPD3 | 3 | 3 | -0,035660943 | 0,441854987 | 1 |
| HYOU1 | 36 | 36 | -1,98057E-17 | 0,999999993 | 1 |
| STK25 | 4 | 4 | 8,69561E-14 | 0,99999927 | 1 |
| DLAT | 32 | 32 | 2,39881E-15 | 0,999999854 | 1 |
| MTUS2 | 5 | 5 | 0,086249672 | 0,224729432 | 1 |
| NPTN | 10 | 10 | 1,94645E-13 | 0,999998787 | 1 |
| PPA1 | 19 | 19 | 4,50945E-17 | 0,999999987 | 1 |
| HINT2 | 5 | 5 | 0 | 1 | 1 |

| | | | | | |
|---|---|---|---|---|---|
| PHOSPHO1 | 4 | 4 | 2,50917E-17 | 0,999999993 | 1 |
| PRRT3 | 14 | 14 | 0 | 1 | 1 |
| PYGB;PYGL | 11 | 11 | -1,93775E-17 | 0,999999998 | 1 |
| MRPL37 | 9 | 7 | 3,73206E-18 | 0,999999999 | 1 |
| SV2A | 22 | 22 | 2,74294E-16 | 0,999999941 | 1 |
| PDK3 | 17 | 16 | 0,023734318 | 0,450710046 | 1 |
| UBR4 | 61 | 61 | 2,30264E-17 | 0,99999999 | 1 |
| PPP3CB | 17 | 17 | -1,2318E-14 | 0,999999528 | 1 |
| SPON1 | 7 | 7 | -0,024158797 | 0,465995662 | 1 |
| BAG3 | 3 | 3 | -0,0416649 | 0,402925993 | 1 |
| TKFC | 7 | 7 | -1,9702E-16 | 0,999999966 | 1 |
| NFASC | 45 | 45 | -8,05063E-17 | 0,999999969 | 1 |
| CRYZ | 13 | 13 | 0 | 1 | 1 |
| ZNRD2 | 2 | 2 | 0 | 1 | 1 |
| ALDOA | 21 | 21 | 4,01934E-17 | 0,99999998 | 1 |
| TTC9B | 5 | 5 | 5,05375E-18 | 0,999999997 | 1 |
| ACAD8 | 16 | 14 | 5,84069E-17 | 0,999999987 | 1 |
| GAP43 | 18 | 18 | -0,007372351 | 0,743592554 | 1 |
| CSNK1A1 | 1 | 1 | 0 | 1 | 1 |
| SLC16A1 | 5 | 5 | 1,66855E-17 | 0,999999992 | 1 |
| VPS53 | 17 | 17 | 0,015539834 | 0,416765225 | 1 |
| AGT | 4 | 4 | -0,079454387 | 0,177653885 | 1 |
| CCDC50 | 1 | 1 | 0 | 1 | 1 |
| TBCD | 16 | 16 | 0 | 1 | 1 |
| AGPS | 7 | 6 | -2,17572E-16 | 0,999999957 | 1 |
| ACOT7 | 26 | 26 | 0,040188291 | 0,3368125 | 1 |
| PRKAR2B | 15 | 15 | -0,031091042 | 0,267332274 | 1 |
| TUBGCP2 | 9 | 9 | -0,044075203 | 0,302815348 | 1 |
| MAP7 | 1 | 1 | -0,023134098 | 0,665043814 | 1 |
| PRKAR2A | 1 | 1 | 0 | 1 | 1 |
| CLSTN1 | 12 | 12 | 2,67178E-17 | 0,999999985 | 1 |
| ITSN1 | 34 | 33 | -4,48852E-17 | 0,999999981 | 1 |
| DMAC2L | 7 | 6 | -0,060855797 | 0,166346592 | 1 |

| | | | | | |
|---|---|---|---|---|---|
| ANXA4 | 6 | 6 | 1,91703E-16 | 0,999999975 | 1 |
| TAOK2 | 11 | 11 | 4,98708E-16 | 0,999999923 | 1 |
| ACSF2 | 20 | 19 | 1,66865E-17 | 0,999999997 | 1 |
| IAH1 | 2 | 2 | 0 | 1 | 1 |
| COX5B | 8 | 8 | 0,048869376 | 0,329359077 | 1 |
| DLG1 | 29 | 28 | 0 | 1 | 1 |
| ATXN2 | 8 | 8 | -4,01048E-16 | 0,999999944 | 1 |
| EIF4B | 14 | 14 | -1,67513E-16 | 0,99999996 | 1 |
| SDHD | 1 | 1 | 0 | 1 | 1 |
| BEGAIN | 18 | 17 | 1,26589E-15 | 0,999999867 | 1 |
| STRIP1 | 6 | 6 | -0,034262955 | 0,3052146 | 1 |
| SH3PXD2A | 6 | 6 | 4,54234E-17 | 0,999999993 | 1 |
| SEL1L | 6 | 6 | -1,79606E-17 | 0,999999988 | 1 |
| ARHGAP21 | 25 | 22 | 0 | 1 | 1 |
| CAPN5 | 22 | 22 | 2,30762E-17 | 0,999999982 | 1 |
| LXN | 6 | 6 | -5,87117E-17 | 0,999999984 | 1 |
| SHANK2 | 47 | 45 | 2,42721E-16 | 0,999999941 | 1 |
| GEMIN5 | 7 | 6 | -2,26851E-18 | 0,999999999 | 1 |
| STRAP | 19 | 19 | 0 | 1 | 1 |
| PDIA6 | 9 | 9 | 8,11886E-18 | 0,999999994 | 1 |
| LTBP2 | 1 | 1 | 0 | 1 | 1 |
| OGT | 37 | 37 | 3,53147E-13 | 0,999998228 | 1 |
| MCCC2 | 21 | 20 | 1,63681E-17 | 0,999999985 | 1 |
| RUSF1 | 3 | 3 | 1,24199E-18 | 1 | 1 |
| PRKAR2A | 16 | 15 | 0 | 1 | 1 |
| ADGRB3 | 16 | 15 | 0,013748116 | 0,571260148 | 1 |
| MYO5A | 90 | 89 | 0 | 1 | 1 |
| ADGRL3 | 25 | 25 | 0 | 1 | 1 |
| DCUN1D2;DCUN1D | 2 | 2 | -5,79498E-21 | 1 | 1 |
| GALK2 | 8 | 8 | 4,44748E-17 | 0,999999991 | 1 |
| DYNC1H1 | 275 | 270 | 0,005672368 | 0,658160333 | 1 |
| RASAL1 | 31 | 31 | 2,49469E-17 | 0,999999983 | 1 |
| NCAM1 | 15 | 15 | -0,04414952 | 0,345907889 | 1 |

| | | | | | |
|---|---|---|---|---|---|
| BCAR1 | 5 | 5 | 0,049320651 | 0,36930725 | 1 |
| ATAT1 | 8 | 8 | 0 | 1 | 1 |
| UBA1 | 60 | 59 | 1,78985E-17 | 0,999999991 | 1 |
| ARHGEF2 | 12 | 12 | 0,020588495 | 0,381313438 | 1 |
| HSD17B4 | 17 | 17 | -0,004729984 | 0,65930445 | 1 |
| ADH5 | 12 | 12 | -3,08859E-15 | 0,999999872 | 1 |
| ETFA | 18 | 18 | 1,25454E-17 | 0,999999989 | 1 |
| MTHFD1L | 39 | 36 | 0 | 1 | 1 |
| KPNB1 | 33 | 33 | -3,5694E-17 | 0,999999984 | 1 |
| PEX5L | 7 | 7 | 1,34742E-18 | 1 | 1 |
| PSMD1 | 42 | 42 | -0,019820076 | 0,293103525 | 1 |
| TLCD4 | 2 | 2 | 3,8515E-15 | 0,999999852 | 1 |
| MAP4 | 16 | 15 | -0,038058142 | 0,425735139 | 1 |
| APIP | 5 | 5 | 0 | 1 | 1 |
| CTNNB1 | 27 | 27 | -1,51012E-16 | 0,999999961 | 1 |
| ENAH | 13 | 12 | -0,027734675 | 0,490880631 | 1 |
| ATP1A2;ATP1A1 | 19 | 19 | 5,02113E-17 | 0,999999989 | 1 |
| ENO2;ENO3 | 2 | 2 | 0 | 1 | 1 |
| ENO1 | 33 | 33 | 8,60687E-16 | 0,999999914 | 1 |
| ENO1;ENO2;ENO3 | 3 | 3 | 2,58854E-15 | 0,999999868 | 1 |
| PSEN1 | 1 | 1 | 0,118036158 | 0,350474007 | 1 |
| CBR3 | 15 | 15 | -7,34663E-17 | 0,999999981 | 1 |
| MMUT | 21 | 18 | 1,00427E-17 | 0,999999991 | 1 |
| RAP1GAP2 | 6 | 6 | -0,010129283 | 0,675616393 | 1 |
| ADD1 | 34 | 34 | -0,042386071 | 0,239328847 | 1 |
| ATRX | 1 | 1 | 0 | 1 | 1 |
| OPA3 | 3 | 3 | 1,01228E-14 | 0,999999764 | 1 |
| WDR47 | 31 | 31 | 3,70955E-18 | 0,999999998 | 1 |
| IBA57 | 6 | 6 | 1,7533E-15 | 0,999999886 | 1 |
| ALDH5A1 | 28 | 27 | 0,021148882 | 0,361359149 | 1 |
| RALBP1 | 6 | 6 | 7,50747E-17 | 0,999999993 | 1 |
| YKT6 | 10 | 10 | -0,005266092 | 0,778159348 | 1 |
| LMNA | 31 | 1 | 0 | 1 | 1 |

| | | | | | |
|---|---|---|---|---|---|
| SLC25A4 | 18 | 16 | 1,47066E-16 | 0,999999965 | 1 |
| NDUFV1 | 26 | 26 | 0,068788255 | 0,148369171 | 1 |
| RABGGTA | 20 | 20 | -1,4107E-16 | 0,999999947 | 1 |
| COTL1 | 16 | 16 | -2,52952E-16 | 0,999999973 | 1 |
| CCAR2 | 30 | 30 | -0,062644665 | 0,133157317 | 1 |
| RARS1 | 33 | 33 | 0,012870471 | 0,342326478 | 1 |
| JAKMIP1 | 2 | 2 | 0 | 1 | 1 |
| RIDA | 9 | 9 | 1,54255E-16 | 0,99999999 | 1 |
| USP15;USP11 | 1 | 1 | 2,89687E-21 | 1 | 1 |
| GNB1 | 15 | 14 | 0,002316583 | 0,807117094 | 1 |
| IVD | 17 | 16 | 7,24853E-19 | 0,999999999 | 1 |
| NUB1 | 3 | 3 | -0,093239 | 0,196059639 | 1 |
| L2HGDH | 18 | 16 | -1,18907E-17 | 0,999999996 | 1 |
| GNB2 | 9 | 8 | 1,16559E-18 | 0,999999999 | 1 |
| ARRB1 | 16 | 16 | -0,052136138 | 0,150115341 | 1 |
| SLC39A6 | 5 | 4 | -0,074467324 | 0,319562666 | 1 |
| GLUL | 25 | 23 | -0,033367002 | 0,283830387 | 1 |
| RPS3A | 19 | 19 | 0,013977025 | 0,698572728 | 1 |
| NISCH | 15 | 15 | 0 | 1 | 1 |
| TPP2 | 46 | 45 | -3,94573E-18 | 0,999999998 | 1 |
| ATP1A3 | 49 | 48 | 0,037132727 | 0,145362264 | 1 |
| ATP1A1 | 43 | 43 | -6,91344E-16 | 0,999999914 | 1 |
| ATP1A2 | 56 | 55 | 0 | 1 | 1 |
| ACTR1A;ACTR1B | 12 | 12 | 4,05893E-17 | 0,999999981 | 1 |
| JPH1 | 2 | 2 | -0,059493446 | 0,380498699 | 1 |
| SACS | 12 | 12 | 0 | 1 | 1 |
| ERLIN2 | 15 | 15 | -3,751E-17 | 0,999999983 | 1 |
| LMBRD2 | 7 | 7 | 0 | 1 | 1 |
| CACNA2D2 | 11 | 11 | -0,029542718 | 0,390379402 | 1 |
| ANXA5 | 25 | 23 | 8,22488E-17 | 0,999999985 | 1 |
| ERLIN1 | 2 | 2 | -0,135929436 | 0,149460983 | 1 |
| CASK | 21 | 21 | 0 | 1 | 1 |
| ATG3 | 6 | 6 | -0,035182287 | 0,3330728 | 1 |

| | | | | | |
|---|---|---|---|---|---|
| ACOT2 | 8 | 8 | 0 | 1 | 1 |
| SNTA1 | 8 | 8 | -0,011788568 | 0,587958736 | 1 |
| CNTN3 | 7 | 7 | -2,08324E-19 | 1 | 1 |
| VAMP1 | 6 | 6 | 0,023372215 | 0,623361013 | 1 |
| VAMP3 | 6 | 6 | 0,09289507 | 0,247869546 | 1 |
| ARHGAP44 | 15 | 15 | -0,014702953 | 0,419002173 | 1 |
| CCDC115 | 1 | 1 | 0 | 1 | 1 |
| AHSA1 | 25 | 25 | 0 | 1 | 1 |
| ADAM22 | 25 | 23 | -1,32965E-16 | 0,999999965 | 1 |
| PPA2 | 20 | 20 | 0 | 1 | 1 |
| SEPTIN5 | 29 | 27 | 1,38558E-18 | 0,999999997 | 1 |
| TLN2 | 7 | 7 | -8,21468E-18 | 0,999999996 | 1 |
| MAP2K6 | 5 | 5 | -6,87321E-17 | 0,999999981 | 1 |
| CLVS1 | 12 | 12 | 2,23262E-18 | 0,999999999 | 1 |
| ELOB | 11 | 11 | 6,50187E-17 | 0,999999978 | 1 |
| DGKG | 16 | 16 | 2,42192E-16 | 0,999999948 | 1 |
| PC | 57 | 56 | 0,012825753 | 0,572457852 | 1 |
| COX10 | 1 | 1 | 0,213491202 | 0,251922439 | 1 |
| PITPNB | 12 | 12 | -4,02712E-17 | 0,99999998 | 1 |
| NUTF2 | 8 | 8 | 1,89991E-16 | 0,999999975 | 1 |
| KCNC1 | 6 | 6 | -1,12618E-17 | 0,999999993 | 1 |
| PRDX1 | 20 | 20 | -1,55357E-16 | 0,999999984 | 1 |
| BIN1 | 33 | 33 | -3,84342E-16 | 0,999999947 | 1 |
| PALM | 15 | 15 | -0,054356273 | 0,317971694 | 1 |
| MYH9 | 99 | 98 | 1,32534E-15 | 0,999999866 | 1 |
| GSR | 10 | 10 | 0,018999097 | 0,507828037 | 1 |
| DAD1 | 2 | 2 | 3,03767E-18 | 0,999999997 | 1 |
| TTYH1 | 6 | 4 | 0 | 1 | 1 |
| OCIAD1 | 10 | 10 | 0 | 1 | 1 |
| SERPINB6 | 25 | 25 | 0,000134344 | 0,961832308 | 1 |
| FKBP8 | 10 | 10 | 6,20267E-17 | 0,999999988 | 1 |
| ICAM1 | 1 | 1 | 0 | 1 | 1 |
| LINGO1 | 16 | 15 | 0,024207566 | 0,312009399 | 1 |

| | | | | | |
|---|---|---|---|---|---|
| SMS | 16 | 16 | -3,86281E-17 | 0,999999996 | 1 |
| IPCEF1 | 2 | 2 | -0,073011804 | 0,246528416 | 1 |
| FUOM | 3 | 3 | -4,78984E-17 | 0,999999994 | 1 |
| MBLAC1 | 3 | 3 | 0 | 1 | 1 |
| CAMK2B | 3 | 3 | 1,01061E-15 | 0,999999925 | 1 |
| RPS8 | 5 | 5 | -6,84409E-19 | 0,999999999 | 1 |
| ANK3 | 42 | 36 | 4,17827E-17 | 0,999999983 | 1 |
| PSMC6 | 23 | 21 | -0,011118813 | 0,516731219 | 1 |
| HSP90AB1 | 36 | 35 | 1,09016E-16 | 0,999999975 | 1 |
| RPSA | 11 | 11 | -0,025919472 | 0,316460999 | 1 |
| GRIA2 | 36 | 36 | 0 | 1 | 1 |
| RAB11FIP5 | 16 | 15 | -2,76346E-11 | 0,99997755 | 1 |
| GRIA3 | 23 | 23 | 0 | 1 | 1 |
| MAP6 | 68 | 67 | -0,140365246 | 0,139167482 | 1 |
| OTUD4 | 2 | 2 | 7,21844E-20 | 1 | 1 |
| TM9SF2 | 7 | 7 | 0 | 1 | 1 |
| ABCE1 | 14 | 14 | -2,23747E-16 | 0,999999949 | 1 |
| ASPA | 9 | 9 | -3,13616E-17 | 0,999999997 | 1 |
| HNRNPM | 4 | 4 | 0 | 1 | 1 |
| MRPS23 | 7 | 7 | 1,42253E-12 | 0,999995724 | 1 |
| GLTP | 4 | 4 | -1,79589E-17 | 0,999999995 | 1 |
| GSTM1 | 28 | 27 | -1,51459E-16 | 0,999999973 | 1 |
| BRSK2 | 14 | 14 | 0 | 1 | 1 |
| RAB28 | 4 | 4 | -3,50061E-19 | 1 | 1 |
| MARS1 | 32 | 32 | 0 | 1 | 1 |
| SLC30A9 | 13 | 13 | 0 | 1 | 1 |
| ENPP6 | 11 | 11 | -6,9986E-16 | 0,999999937 | 1 |
| UBE2L3 | 8 | 8 | 0 | 1 | 1 |
| MPDZ | 10 | 10 | -0,048031924 | 0,268929608 | 1 |
| SPATA2L | 3 | 3 | -6,33342E-17 | 0,999999988 | 1 |
| GRIK2 | 3 | 1 | 3,45756E-18 | 0,999999997 | 1 |
| RAB14 | 20 | 20 | 1,26157E-14 | 0,99999947 | 1 |
| KRT14 | 18 | 4 | 0 | 1 | 1 |

| | | | | | |
|---|---|---|---|---|---|
| OXR1 | 24 | 23 | 0 | 1 | 1 |
| SORT1 | 9 | 9 | 2,27648E-11 | 0,999980264 | 1 |
| HECTD4 | 28 | 28 | 0 | 1 | 1 |
| HSP90AA1;HSP90A | 16 | 16 | 0,001385228 | 0,849275799 | 1 |
| MLYCD | 8 | 8 | 1,14888E-17 | 0,999999995 | 1 |
| RAB27B | 8 | 8 | 0,022893353 | 0,427533815 | 1 |
| SORBS2 | 11 | 10 | 0 | 1 | 1 |
| ADSL | 13 | 13 | -3,5108E-17 | 0,999999991 | 1 |
| PACS1 | 31 | 31 | -1,64031E-16 | 0,999999968 | 1 |
| SERPINB1A | 21 | 21 | 1,12674E-15 | 0,99999995 | 1 |
| FLII | 8 | 8 | 0,042725271 | 0,165406372 | 1 |
| EIF2B1 | 5 | 5 | -2,94983E-19 | 1 | 1 |
| NIT2 | 14 | 14 | 0,038491666 | 0,325301272 | 1 |
| APLP2 | 9 | 9 | -1,92186E-17 | 0,999999992 | 1 |
| TARS1 | 35 | 34 | 1,65051E-17 | 0,999999984 | 1 |
| HADHA | 36 | 34 | 7,51854E-17 | 0,999999974 | 1 |
| MYH10 | 106 | 106 | 0 | 1 | 1 |
| ABCB7 | 18 | 15 | 0,000712909 | 0,901372993 | 1 |
| PTEN | 9 | 9 | 0 | 1 | 1 |
| PGM1 | 43 | 43 | 0 | 1 | 1 |
| ISYNA1 | 12 | 12 | 2,06164E-17 | 0,999999995 | 1 |
| ANPEP | 3 | 3 | 1,01595E-15 | 0,999999963 | 1 |
| GATB | 4 | 4 | -1,19786E-18 | 1 | 1 |
| UBA2 | 23 | 23 | -0,063432195 | 0,181079801 | 1 |
| RUFY3 | 1 | 1 | 0 | 1 | 1 |
| ALDH1L1 | 44 | 41 | -0,047291072 | 0,221257983 | 1 |
| NCS1 | 11 | 10 | -2,6928E-16 | 0,999999947 | 1 |
| RPS6KA2 | 10 | 10 | -0,027546179 | 0,470057837 | 1 |
| SAMHD1 | 2 | 2 | 2,70789E-18 | 0,999999999 | 1 |
| CORO1A | 22 | 22 | -4,15657E-17 | 0,999999978 | 1 |
| POLR2H | 7 | 7 | -0,05500534 | 0,157741162 | 1 |
| MAP2 | 103 | 101 | -0,009157368 | 0,710485533 | 1 |
| EHD3;EHD1 | 11 | 11 | 1,63109E-16 | 0,999999979 | 1 |

| | | | | | |
|---|---|---|---|---|---|
| SNAP25 | 26 | 26 | 0,020737789 | 0,433955519 | 1 |
| SPTBN4 | 44 | 44 | -0,023353255 | 0,358682929 | 1 |
| RALA | 7 | 7 | 0,051084642 | 0,17855375 | 1 |
| RPS14 | 8 | 8 | 0,104692689 | 0,168564668 | 1 |
| TF | 38 | 36 | 3,41053E-17 | 0,999999997 | 1 |
| ATAD3 | 27 | 25 | 1,95698E-16 | 0,999999952 | 1 |
| GNB5 | 10 | 10 | 0 | 1 | 1 |
| ACE | 12 | 11 | -2,91615E-17 | 0,999999996 | 1 |
| SRGAP1 | 4 | 4 | -6,94718E-18 | 0,999999994 | 1 |
| DLGAP2 | 20 | 19 | 3,50406E-17 | 0,999999978 | 1 |
| ZNF365 | 2 | 2 | 1,71281E-16 | 0,999999988 | 1 |
| CMTR1 | 2 | 2 | -5,97512E-19 | 0,999999999 | 1 |
| NEO1 | 21 | 21 | 0 | 1 | 1 |
| PACSIN3 | 7 | 7 | 0 | 1 | 1 |
| TBC1D9B | 10 | 10 | -5,47783E-18 | 0,999999998 | 1 |
| PTPRN2 | 24 | 23 | -2,30566E-17 | 0,999999992 | 1 |
| BCAS1 | 16 | 16 | -2,13857E-13 | 0,999998835 | 1 |
| LNPK | 12 | 12 | 2,14741E-18 | 0,999999998 | 1 |
| GALE | 8 | 7 | 6,55911E-18 | 0,999999997 | 1 |
| NUCB1 | 11 | 11 | -1,31992E-16 | 0,999999972 | 1 |
| MAGED1 | 4 | 4 | -1,09799E-16 | 0,999999979 | 1 |
| PSMC4 | 18 | 18 | -0,008930385 | 0,629584909 | 1 |
| SEPTIN9 | 30 | 28 | 0 | 1 | 1 |
| SEPTIN7 | 38 | 38 | 0,026098137 | 0,208273246 | 1 |
| SEPTIN4 | 13 | 12 | -9,62552E-13 | 0,999995988 | 1 |
| SEPTIN3 | 26 | 26 | 0 | 1 | 1 |
| ERH | 1 | 1 | -0,045944677 | 0,592630296 | 1 |
| NUFIP2 | 2 | 2 | -1,28917E-18 | 0,999999999 | 1 |
| AIFM3 | 10 | 9 | 1,85432E-17 | 0,999999991 | 1 |
| SCLY | 3 | 3 | 0 | 1 | 1 |
| ATP2B1 | 62 | 61 | 0,009858814 | 0,537477525 | 1 |
| BTBD17 | 11 | 11 | 5,15728E-16 | 0,999999931 | 1 |
| AMACR | 8 | 8 | 8,60973E-17 | 0,999999983 | 1 |

| | | | | | |
|---|---|---|---|---|---|
| TRIM28 | 15 | 14 | -2,60671E-15 | 0,999999851 | 1 |
| SPTAN1 | 235 | 231 | -5,24618E-14 | 0,999999395 | 1 |
| SPTA1 | 2 | 2 | 0 | 1 | 1 |
| GBP2 | 3 | 3 | -5,29397E-16 | 0,999999954 | 1 |
| COX4I1 | 17 | 17 | -3,50758E-17 | 0,999999989 | 1 |
| ATP6V1A | 58 | 58 | 0,030556285 | 0,334173405 | 1 |
| RCN2 | 14 | 14 | -0,039987965 | 0,357294879 | 1 |
| MTMR7 | 6 | 6 | -2,12821E-17 | 0,999999989 | 1 |
| FNBP1 | 9 | 9 | 1,03897E-17 | 0,999999996 | 1 |
| ADCY2 | 8 | 8 | 0,03418357 | 0,264189149 | 1 |
| AGAP2 | 31 | 31 | 0 | 1 | 1 |
| JPH3 | 4 | 4 | -7,2528E-16 | 0,999999926 | 1 |
| DCUN1D3 | 3 | 3 | 0 | 1 | 1 |
| DCPS | 15 | 15 | -0,08969354 | 0,113153335 | 1 |
| KRT76 | 4 | 4 | 3,26052E-15 | 0,999999933 | 1 |
| TOMM22 | 6 | 6 | 0,072238223 | 0,186575304 | 1 |
| ASL | 16 | 16 | -0,024140501 | 0,358411163 | 1 |
| PAPSS2 | 2 | 2 | -1,11508E-14 | 0,999999758 | 1 |
| RPS10 | 9 | 9 | -1,54516E-17 | 0,999999992 | 1 |
| PDXK | 16 | 16 | 0 | 1 | 1 |
| WBP2 | 9 | 9 | 0,004625919 | 0,753798267 | 1 |
| ARMC1 | 8 | 8 | -4,54023E-17 | 0,999999983 | 1 |
| ANAPC5 | 3 | 3 | 1,54478E-17 | 0,999999994 | 1 |
| WASHC2 | 5 | 5 | 0 | 1 | 1 |
| IGF2R | 4 | 4 | -1,80001E-17 | 0,999999993 | 1 |
| NUDCD3 | 18 | 18 | 5,7997E-17 | 0,999999987 | 1 |
| COMMD9 | 3 | 3 | -8,33529E-20 | 1 | 1 |
| PCSK1N | 6 | 6 | -3,5315E-18 | 0,999999997 | 1 |
| TARS2 | 12 | 11 | 0 | 1 | 1 |
| USP8 | 9 | 9 | 0 | 1 | 1 |
| CHMP2A | 5 | 5 | -5,98792E-17 | 0,99999998 | 1 |
| SLC12A5 | 32 | 32 | -2,07277E-17 | 0,999999991 | 1 |
| UHRF1BP1L | 26 | 25 | 1,84036E-14 | 0,999999385 | 1 |

| | | | | | |
|---|---|---|---|---|---|
| MARCKS | 11 | 11 | -2,22949E-16 | 0,999999968 | 1 |
| TBCA | 6 | 6 | -1,08585E-16 | 0,99999997 | 1 |
| ANXA2 | 16 | 13 | 3,34135E-16 | 0,999999961 | 1 |
| RPS2 | 9 | 9 | 0 | 1 | 1 |
| ADO | 9 | 9 | -0,042970952 | 0,269011356 | 1 |
| PCNA | 3 | 3 | -0,095828285 | 0,171504641 | 1 |
| USP15 | 25 | 25 | 3,91022E-17 | 0,999999978 | 1 |
| HSPH1 | 51 | 50 | 3,54403E-17 | 0,999999996 | 1 |
| CHID1 | 10 | 9 | -1,20945E-15 | 0,999999896 | 1 |
| ROCK2 | 52 | 51 | 5,48183E-17 | 0,999999972 | 1 |
| RDH14 | 8 | 8 | 0 | 1 | 1 |
| SCN3B | 4 | 4 | 8,29349E-16 | 0,999999985 | 1 |
| STRN3 | 14 | 14 | 4,18118E-17 | 0,999999987 | 1 |
| MAPT | 1 | 1 | -0,060283289 | 0,323996016 | 1 |
| EPB41L3 | 31 | 31 | -0,043457733 | 0,325064189 | 1 |
| CANX | 25 | 24 | 1,48021E-17 | 0,999999994 | 1 |
| PITPNA | 27 | 27 | 3,39348E-17 | 0,999999986 | 1 |
| ATPIF1 | 6 | 6 | 0 | 1 | 1 |
| CDC37 | 20 | 20 | -1,78386E-17 | 0,999999994 | 1 |
| ATP6V1E1 | 28 | 26 | 4,53868E-15 | 0,999999755 | 1 |
| BIN1 | 1 | 1 | 0 | 1 | 1 |
| RAB30 | 8 | 8 | -2,28652E-15 | 0,999999862 | 1 |
| LRPAP1 | 17 | 17 | -0,008663106 | 0,62570067 | 1 |
| RHOT2 | 8 | 8 | 0 | 1 | 1 |
| RHOT1 | 15 | 14 | 0 | 1 | 1 |
| ACAA1A | 18 | 17 | -3,45355E-17 | 0,99999998 | 1 |
| RABEP2 | 2 | 2 | 0 | 1 | 1 |
| DNM1L | 40 | 40 | -0,01332028 | 0,464666427 | 1 |
| RMDN3 | 14 | 13 | 0,046600094 | 0,132453345 | 1 |
| TMEM132A | 4 | 4 | 2,73575E-18 | 0,999999997 | 1 |
| C1QBP | 7 | 7 | -1,8652E-14 | 0,999999623 | 1 |
| VAT1L | 18 | 18 | -2,76618E-16 | 0,999999968 | 1 |
| RMDN1 | 8 | 6 | 0,044171406 | 0,307274254 | 1 |

| | | | | | |
|---|---|---|---|---|---|
| CACNA1A | 13 | 12 | 0,027093127 | 0,39058445 | 1 |
| FUNDC2 | 5 | 5 | 0 | 1 | 1 |
| RALB | 4 | 4 | 3,28277E-14 | 0,999999622 | 1 |
| ARHGDIA | 15 | 15 | 0 | 1 | 1 |
| ENOPH1 | 8 | 8 | 2,53946E-17 | 0,999999988 | 1 |
| DHDH | 8 | 8 | 0 | 1 | 1 |
| RNPEP | 21 | 21 | 6,29849E-17 | 0,999999968 | 1 |
| MAST3 | 9 | 9 | -1,10195E-16 | 0,999999983 | 1 |
| BCL2L13 | 6 | 6 | -1,98899E-17 | 0,999999989 | 1 |
| BASP1 | 16 | 16 | 4,77794E-16 | 0,999999962 | 1 |
| TPM4 | 8 | 8 | -0,004559229 | 0,765600635 | 1 |
| ALS2 | 8 | 8 | -3,419E-14 | 0,999999824 | 1 |
| ARHGAP23 | 21 | 20 | -0,016950562 | 0,457210061 | 1 |
| MAP1A | 126 | 125 | -0,056295613 | 0,240892947 | 1 |
| ANK2 | 150 | 149 | 0 | 1 | 1 |
| SLC25A18 | 11 | 9 | -5,10943E-18 | 0,999999995 | 1 |
| VCPIP1 | 29 | 29 | 0 | 1 | 1 |
| PTPRA | 21 | 21 | 1,22461E-17 | 0,999999987 | 1 |
| GPRIN1 | 45 | 45 | -0,097665113 | 0,137729443 | 1 |
| PTPRE | 12 | 12 | 5,03696E-17 | 0,999999984 | 1 |
| AHNAK | 4 | 4 | -3,13976E-17 | 0,99999999 | 1 |
| PTPRN | 11 | 11 | -7,67986E-16 | 0,999999947 | 1 |
| LRPPRC | 62 | 59 | 3,08379E-16 | 0,999999944 | 1 |
| ABCF3 | 7 | 7 | -1,26253E-05 | 0,989043464 | 1 |
| SYNE1 | 43 | 38 | -0,033095378 | 0,166148992 | 1 |
| RTN1 | 27 | 27 | -0,037315432 | 0,464024307 | 1 |
| ATG7 | 13 | 13 | 1,42278E-06 | 0,994529424 | 1 |
| PPP2R2D | 2 | 2 | -3,43745E-16 | 0,999999988 | 1 |
| RELCH | 32 | 32 | -0,063423172 | 0,128183299 | 1 |
| EXTL2 | 4 | 4 | -0,034483657 | 0,500305986 | 1 |
| TLN2 | 75 | 74 | 7,18137E-17 | 0,999999972 | 1 |
| RTN3 | 40 | 40 | -0,054251757 | 0,329426667 | 1 |
| MAP7D2 | 13 | 13 | -0,067123748 | 0,230395926 | 1 |

| | | | | | |
|---|---|---|---|---|---|
| BNIP3 | 4 | 4 | 0,017479405 | 0,60314376 | 1 |
| SRGAP2 | 25 | 25 | 0,005001933 | 0,704857706 | 1 |
| KIF3B | 15 | 15 | 0 | 1 | 1 |
| VPS25 | 5 | 5 | -6,77707E-17 | 0,999999989 | 1 |
| CIT | 16 | 13 | 0 | 1 | 1 |
| RABEP1 | 32 | 32 | -1,31118E-15 | 0,999999862 | 1 |
| TRAPPC10 | 13 | 13 | 1,84745E-17 | 0,999999994 | 1 |
| EML1 | 7 | 7 | -7,49538E-18 | 0,999999998 | 1 |
| DCTN1 | 57 | 57 | 0 | 1 | 1 |
| GHITM | 2 | 2 | 1,17458E-15 | 0,999999932 | 1 |
| EPS15 | 14 | 14 | 0 | 1 | 1 |
| GANAB | 39 | 37 | 0,020934221 | 0,419540542 | 1 |
| PDCD10 | 7 | 7 | 0 | 1 | 1 |
| VPS35 | 33 | 33 | -0,01446847 | 0,530138283 | 1 |
| TRAPPC14 | 6 | 6 | 3,69447E-17 | 0,99999999 | 1 |
| SOGA3 | 34 | 33 | 0 | 1 | 1 |
| RB1CC1 | 7 | 7 | 1,01631E-18 | 0,999999999 | 1 |
| ABCG2 | 6 | 6 | 0,043568827 | 0,261284509 | 1 |
| CTTNBP2 | 25 | 25 | -0,053969194 | 0,190472733 | 1 |
| CCDC6 | 12 | 12 | -5,33929E-16 | 0,999999912 | 1 |
| DENND4B | 4 | 3 | 0,10922754 | 0,252515714 | 1 |
| NLN | 13 | 13 | 2,25531E-18 | 0,999999997 | 1 |
| IDI1 | 7 | 7 | -2,00707E-18 | 0,999999997 | 1 |
| STRIP2 | 1 | 1 | 0 | 1 | 1 |
| ACSS2 | 11 | 11 | -8,38417E-18 | 0,999999995 | 1 |
| DSP | 53 | 22 | 6,10188E-16 | 0,999999955 | 1 |
| KCNH3 | 1 | 1 | 4,86401E-20 | 1 | 1 |
| TPD52L1 | 3 | 3 | -0,060623904 | 0,156375076 | 1 |
| CCDC93 | 10 | 10 | 0 | 1 | 1 |
| NPTXR | 13 | 13 | 0,058023564 | 0,130368615 | 1 |
| BNIP1 | 2 | 2 | -2,69996E-18 | 0,999999996 | 1 |
| RPS3 | 24 | 24 | -0,054956248 | 0,124565541 | 1 |
| UMPS | 7 | 6 | -1,56598E-15 | 0,99999988 | 1 |

| | | | | | |
|---|---|---|---|---|---|
| STRN | 27 | 27 | 0 | 1 | 1 |
| GOLGA3 | 13 | 13 | -0,021929193 | 0,472006715 | 1 |
| HAPLN4 | 10 | 10 | 1,13511E-15 | 0,999999899 | 1 |
| PSMD11 | 28 | 27 | -0,036215637 | 0,147262018 | 1 |
| WDR11 | 9 | 9 | 0 | 1 | 1 |
| MPRIP | 20 | 17 | 2,81419E-17 | 0,999999985 | 1 |
| APOL9B;APOL9A | 1 | 1 | 0 | 1 | 1 |
| PALM2 | 13 | 13 | -6,6146E-17 | 0,999999984 | 1 |
| DNAJC13 | 29 | 28 | 2,69346E-17 | 0,999999985 | 1 |
| GPM6B | 9 | 9 | -2,41188E-15 | 0,99999986 | 1 |
| USP20 | 1 | 1 | 0 | 1 | 1 |
| DPYSL5 | 30 | 29 | -5,40737E-17 | 0,999999981 | 1 |
| NCALD | 13 | 13 | -0,049007858 | 0,308658897 | 1 |
| GFAP | 29 | 26 | -7,11455E-18 | 0,999999998 | 1 |
| ALDOC | 37 | 37 | -1,14572E-15 | 0,9999999 | 1 |
| ATP13A1 | 12 | 12 | -5,66365E-17 | 0,999999984 | 1 |
| ADCK2 | 1 | 1 | 0 | 1 | 1 |
| BCAP31 | 8 | 8 | -0,000128678 | 0,964057703 | 1 |
| ARPC4 | 7 | 7 | 3,14241E-17 | 0,999999981 | 1 |
| MARK1 | 18 | 17 | -1,12189E-16 | 0,999999971 | 1 |
| SIK3 | 16 | 16 | -1,30029E-16 | 0,999999952 | 1 |
| HEXIM1 | 3 | 3 | -1,04709E-13 | 0,999999204 | 1 |
| MACF1 | 41 | 40 | 0 | 1 | 1 |
| HMOX2 | 16 | 15 | -2,61305E-19 | 1 | 1 |
| PRRC2A | 11 | 11 | -0,056655342 | 0,33569641 | 1 |
| UBE2N | 11 | 11 | -7,78632E-17 | 0,999999972 | 1 |
| ACTR3 | 19 | 19 | 0 | 1 | 1 |
| ACTR3B | 15 | 15 | 0 | 1 | 1 |
| AKAP12 | 16 | 16 | -2,60286E-17 | 0,999999987 | 1 |
| PTPRD | 33 | 31 | -8,95863E-17 | 0,999999956 | 1 |
| DGKZ | 8 | 8 | 0,009073318 | 0,590756293 | 1 |
| BMERB1 | 3 | 3 | -3,91137E-17 | 0,999999983 | 1 |
| RACK1 | 20 | 20 | -0,00749308 | 0,71482559 | 1 |

| | | | | | |
|---|---|---|---|---|---|
| CADPS | 81 | 80 | 2,52165E-15 | 0,999999797 | 1 |
| HOMER1 | 23 | 23 | -0,021502289 | 0,495468462 | 1 |
| DOCK9 | 31 | 31 | 0,036849704 | 0,242535707 | 1 |
| LRRFIP2 | 2 | 2 | 0 | 1 | 1 |
| STOML2 | 12 | 11 | 0 | 1 | 1 |
| TJP1 | 30 | 28 | -4,14449E-16 | 0,999999932 | 1 |
| PDXP | 16 | 16 | 4,87844E-17 | 0,999999978 | 1 |
| LUC7L | 1 | 1 | 0 | 1 | 1 |
| DNAH7C;DNAH7B;I | 1 | 1 | 1,22835E-17 | 0,999999991 | 1 |
| SPECC1 | 15 | 14 | -2,87438E-17 | 0,999999986 | 1 |
| TJP2 | 18 | 18 | -9,11949E-18 | 0,999999995 | 1 |
| IDE | 7 | 7 | 6,55981E-14 | 0,999999335 | 1 |
| LRP1B | 10 | 9 | -1,65986E-15 | 0,999999916 | 1 |
| CXXC5 | 1 | 1 | -0,105960683 | 0,245733404 | 1 |
| TACC1 | 8 | 8 | -0,059764995 | 0,32343734 | 1 |
| SLC9A6 | 8 | 8 | 0 | 1 | 1 |
| KRT2 | 43 | 11 | 7,16228E-17 | 0,999999996 | 1 |
| HSPA4 | 74 | 73 | 0 | 1 | 1 |
| RPGR | 5 | 5 | 0 | 1 | 1 |
| FARP1 | 21 | 21 | -0,03778559 | 0,114962798 | 1 |
| FECH | 14 | 12 | 3,26314E-17 | 0,999999985 | 1 |
| PLS3 | 29 | 29 | -0,045132643 | 0,150006031 | 1 |
| WDR44 | 27 | 27 | -1,67435E-16 | 0,999999961 | 1 |
| GPS1 | 9 | 9 | -1,27789E-15 | 0,999999902 | 1 |
| SGIP1 | 2 | 2 | 0 | 1 | 1 |
| RNF14 | 1 | 1 | 0 | 1 | 1 |
| OPTC | 1 | 1 | 0 | 1 | 1 |
| PLPPR3 | 8 | 8 | 0 | 1 | 1 |
| KRT10 | 26 | 8 | -2,11707E-17 | 0,999999995 | 1 |
| SEC23A | 14 | 14 | 0 | 1 | 1 |
| SHPK | 3 | 3 | 0 | 1 | 1 |
| SEH1L | 4 | 4 | -1,68412E-17 | 0,999999992 | 1 |
| PPFIA3 | 52 | 48 | 0 | 1 | 1 |

| | | | | | |
|---|---|---|---|---|---|
| PPFIA2 | 25 | 25 | 9,71251E-32 | 1 | 1 |
| LLGL1 | 14 | 14 | -2,77021E-17 | 0,999999989 | 1 |
| NUDC | 26 | 26 | -6,71804E-15 | 0,999999695 | 1 |
| FAM114A2 | 11 | 10 | -0,088302352 | 0,180913473 | 1 |
| CLMP | 1 | 1 | 0,011184063 | 0,784683874 | 1 |
| NRCAM | 32 | 31 | -1,52661E-16 | 0,999999948 | 1 |
| GET4 | 7 | 7 | -1,3723E-17 | 0,999999991 | 1 |
| HNRNPU | 16 | 15 | -0,052553274 | 0,264740356 | 1 |
| MOG | 13 | 13 | -1,15687E-15 | 0,999999941 | 1 |
| CLIP3 | 6 | 6 | -2,01694E-17 | 0,999999991 | 1 |
| KCNAB1 | 13 | 12 | 4,75839E-16 | 0,999999937 | 1 |
| UQCRFS1 | 16 | 16 | 0,029990695 | 0,359655338 | 1 |
| THTPA | 7 | 7 | -6,63994E-17 | 0,999999978 | 1 |
| MICAL3 | 27 | 27 | -0,029179776 | 0,367417873 | 1 |
| NTAN1 | 1 | 1 | 0 | 1 | 1 |
| GARS1 | 30 | 30 | -6,3996E-19 | 1 | 1 |
| GPR37L1 | 3 | 3 | 0,027589448 | 0,422724074 | 1 |
| LHPP | 4 | 4 | 7,61908E-16 | 0,999999951 | 1 |
| KCNAB2 | 21 | 21 | 0 | 1 | 1 |
| HABP4 | 6 | 6 | -0,04876987 | 0,278905259 | 1 |
| GABRA1 | 10 | 10 | -1,21886E-17 | 0,999999992 | 1 |
| DNM3 | 36 | 36 | 0 | 1 | 1 |
| CAMK1D;CAMK1 | 5 | 5 | 9,84018E-17 | 0,999999983 | 1 |
| PAICS | 18 | 18 | -2,34128E-16 | 0,999999949 | 1 |
| PEA15 | 11 | 11 | -3,26033E-17 | 0,999999987 | 1 |
| NSMF | 8 | 8 | 0,024905075 | 0,346232057 | 1 |
| HACE1 | 10 | 10 | -8,76214E-18 | 0,999999993 | 1 |
| ECHS1 | 17 | 15 | 2,52784E-17 | 0,999999991 | 1 |
| RDX | 16 | 16 | -0,020060862 | 0,382603001 | 1 |
| SPHKAP | 15 | 15 | -0,049366391 | 0,269569944 | 1 |
| CCT5 | 36 | 35 | 6,183E-18 | 0,999999996 | 1 |
| CCT4 | 38 | 37 | -0,02183537 | 0,190991481 | 1 |
| OCRL | 22 | 21 | 1,24838E-15 | 0,999999896 | 1 |

| | | | | | |
|---|---|---|---|---|---|
| PLCD1 | 15 | 15 | -2,178E-17 | 0,999999994 | 1 |
| TARDBP | 6 | 6 | -0,061424779 | 0,133598334 | 1 |
| MTMR11 | 1 | 1 | 0 | 1 | 1 |
| PZP | 33 | 33 | -1,25777E-16 | 0,999999992 | 1 |
| SAG | 2 | 2 | 6,30437E-17 | 1 | 1 |
| IPO7 | 22 | 22 | -3,52633E-17 | 0,999999982 | 1 |
| HIBCH | 24 | 24 | 0,014298884 | 0,458139258 | 1 |
| QRSL1 | 9 | 9 | 0 | 1 | 1 |
| NRDC | 24 | 24 | 6,4499E-17 | 0,99999999 | 1 |
| CLASP1 | 27 | 27 | -7,76545E-17 | 0,999999978 | 1 |
| PGM2 | 7 | 6 | 0 | 1 | 1 |
| GDPD1 | 14 | 14 | 0 | 1 | 1 |
| ACLY | 56 | 54 | 0,015353028 | 0,250278177 | 1 |
| OLA1 | 25 | 23 | 0 | 1 | 1 |
| ARFGAP1 | 17 | 16 | -1,13062E-16 | 0,999999967 | 1 |
| VCP | 73 | 73 | 0 | 1 | 1 |
| CPNE7 | 22 | 20 | 0,103696106 | 0,142263124 | 1 |
| SLC2A3 | 14 | 14 | -1,1028E-13 | 0,999998751 | 1 |
| CSMD2 | 7 | 7 | -6,1667E-19 | 0,999999999 | 1 |
| PSMA3 | 15 | 15 | 0,039454999 | 0,327224483 | 1 |
| ALDH9A1 | 19 | 19 | 1,9833E-17 | 0,99999999 | 1 |
| HCN2;HCN1 | 4 | 4 | 0,059987896 | 0,344559555 | 1 |
| PRKAG1 | 6 | 6 | -0,015831208 | 0,572208963 | 1 |
| PGAP1 | 8 | 6 | -0,032831686 | 0,445486784 | 1 |
| RYR3 | 4 | 4 | 0 | 1 | 1 |
| NPEPPS | 49 | 48 | 0,040600313 | 0,202278192 | 1 |
| SLC32A1 | 13 | 13 | -0,004716156 | 0,720666382 | 1 |
| NPLOC4 | 7 | 7 | -0,045409199 | 0,315696016 | 1 |
| PICK1 | 2 | 2 | 0 | 1 | 1 |
| GM49486 | 6 | 6 | -0,045336168 | 0,28130048 | 1 |
| VWA5A | 18 | 18 | 0,032866806 | 0,154747944 | 1 |
| PPP1R11 | 8 | 8 | 0,012555907 | 0,627383925 | 1 |
| OTUB1 | 18 | 17 | 6,91444E-17 | 0,999999977 | 1 |

| | | | | | |
|---|---|---|---|---|---|
| MPST | 15 | 14 | 0 | 1 | 1 |
| FLNA | 20 | 16 | -1,73893E-16 | 0,999999974 | 1 |
| PRRC2C | 12 | 12 | -0,06481665 | 0,24316566 | 1 |
| SRSF3 | 2 | 2 | -3,75745E-16 | 0,999999972 | 1 |
| PFAS | 27 | 27 | 0 | 1 | 1 |
| TCP1 | 34 | 34 | 5,28923E-17 | 0,99999999 | 1 |
| IGSF11 | 3 | 3 | 0,097656906 | 0,191786769 | 1 |
| FOLH1 | 13 | 12 | -0,098378167 | 0,161275476 | 1 |
| IDE | 18 | 18 | 9,12144E-17 | 0,99999998 | 1 |
| EIF3A | 45 | 44 | -0,004978227 | 0,658292304 | 1 |
| B3GALT6 | 1 | 1 | 0 | 1 | 1 |
| PTGES2 | 13 | 13 | 1,27859E-18 | 0,999999998 | 1 |
| C9ORF72 | 3 | 3 | 0 | 1 | 1 |
| TRAP1 | 22 | 22 | 2,76841E-16 | 0,999999954 | 1 |
| THUMPD1 | 13 | 13 | -0,026072091 | 0,435944784 | 1 |
| VPS11 | 18 | 18 | -1,07471E-13 | 0,999998952 | 1 |
| RRAGC | 16 | 15 | -0,004740093 | 0,744884968 | 1 |
| NELL2 | 3 | 3 | 0 | 1 | 1 |
| RPL34 | 5 | 5 | -2,66055E-16 | 0,999999981 | 1 |
| CAP2 | 31 | 31 | -3,78574E-17 | 0,999999978 | 1 |
| ZFP933 | 1 | 1 | 0 | 1 | 1 |
| MIA2 | 6 | 6 | -0,019726647 | 0,592677521 | 1 |
| GSTP1 | 14 | 14 | 1,71708E-16 | 0,999999987 | 1 |
| RAB10 | 11 | 11 | 0 | 1 | 1 |
| ACSL6 | 40 | 39 | 0,01385874 | 0,339288147 | 1 |
| FRY | 12 | 12 | 0 | 1 | 1 |
| TRAPPC12 | 12 | 12 | -4,29851E-17 | 0,999999979 | 1 |
| GALK1 | 9 | 9 | -1,16658E-16 | 0,999999966 | 1 |
| NBEA | 65 | 64 | -8,75655E-16 | 0,999999923 | 1 |
| LYST | 10 | 10 | -1,76249E-16 | 0,999999984 | 1 |
| PPP2R5E | 14 | 14 | -1,26722E-17 | 0,999999988 | 1 |
| PNCK | 5 | 5 | -1,93814E-15 | 0,999999879 | 1 |
| DCC | 11 | 11 | 0 | 1 | 1 |

| | | | | | |
|---|---|---|---|---|---|
| ARFGEF2 | 16 | 16 | 2,49522E-17 | 0,999999984 | 1 |
| DNAJC2 | 2 | 2 | -2,84493E-19 | 0,999999999 | 1 |
| ALDH6A1 | 31 | 29 | -4,59646E-17 | 0,999999984 | 1 |
| YJEFN3 | 2 | 2 | -0,149426958 | 0,361747208 | 1 |
| LANCL1 | 10 | 10 | 0,008285261 | 0,56166569 | 1 |
| BAIAP2 | 34 | 34 | 0 | 1 | 1 |
| SLC9A1 | 11 | 10 | 0 | 1 | 1 |
| NAPRT | 8 | 8 | -0,020746603 | 0,61671752 | 1 |
| LAMP2 | 5 | 5 | 3,70848E-17 | 0,999999993 | 1 |
| AKR1B7 | 2 | 2 | 0,015273645 | 0,757118621 | 1 |
| GNS | 5 | 5 | 0 | 1 | 1 |
| OPLAH | 30 | 29 | -0,005076798 | 0,728108256 | 1 |
| SYNJ1 | 51 | 51 | 0,031115919 | 0,162025589 | 1 |
| GFOD1 | 9 | 9 | -3,55104E-18 | 0,999999998 | 1 |
| PROM1 | 4 | 4 | -0,070962252 | 0,264730014 | 1 |
| UBXN1 | 5 | 5 | -0,046704833 | 0,196257932 | 1 |
| ARHGAP39 | 27 | 27 | 0,026340894 | 0,323544156 | 1 |
| VAMP7 | 9 | 9 | 0,051080487 | 0,205784932 | 1 |
| ESD | 13 | 13 | 0,01511867 | 0,597157368 | 1 |
| CLMN | 14 | 14 | -0,033563367 | 0,293482369 | 1 |
| TOLLIP | 12 | 12 | -0,042466322 | 0,14289232 | 1 |
| GPRC5B | 3 | 3 | -5,19784E-18 | 0,999999998 | 1 |
| SEC61A2 | 1 | 1 | 0 | 1 | 1 |
| SEC61A1 | 2 | 2 | -1,21899E-17 | 0,999999996 | 1 |
| FARSA | 19 | 19 | -0,006787177 | 0,747680445 | 1 |
| CLASP2 | 6 | 6 | -9,12545E-18 | 0,999999995 | 1 |
| ATG2B | 12 | 12 | 5,76419E-18 | 0,999999995 | 1 |
| VIPAS39 | 11 | 11 | 0 | 1 | 1 |
| ACADM | 16 | 14 | 1,1457E-16 | 0,999999971 | 1 |
| ATP2A2 | 56 | 56 | 2,92351E-17 | 0,999999973 | 1 |
| ADD3 | 25 | 25 | -0,01985168 | 0,364413019 | 1 |
| ATIC | 29 | 27 | -2,73523E-16 | 0,999999953 | 1 |
| TGFBRAP1 | 5 | 5 | 0 | 1 | 1 |

| | | | | | |
|---|---|---|---|---|---|
| RPP14 | 1 | 1 | 0 | 1 | 1 |
| FAH | 9 | 8 | -4,15523E-17 | 0,999999988 | 1 |
| TFRC | 21 | 21 | 1,7365E-16 | 0,999999955 | 1 |
| RTCA | 7 | 7 | -8,79899E-17 | 0,999999968 | 1 |
| SLC12A9 | 7 | 6 | -0,02708951 | 0,53624103 | 1 |
| PGM3 | 18 | 18 | 0 | 1 | 1 |
| ABR | 23 | 23 | -0,013602029 | 0,481158211 | 1 |
| PREP | 44 | 43 | 1,95263E-17 | 0,999999992 | 1 |
| VPS45 | 20 | 20 | 0,025052605 | 0,2871104 | 1 |
| EEF1D | 6 | 6 | -0,010942757 | 0,601566753 | 1 |
| TIAM1 | 1 | 1 | 0 | 1 | 1 |
| RGS1 | 1 | 1 | 0 | 1 | 1 |
| TUBA1C;TUBA8 | 17 | 17 | 0 | 1 | 1 |
| TUBA1C | 15 | 15 | -2,70185E-17 | 0,99999999 | 1 |
| UPF2 | 7 | 7 | -4,2846E-18 | 0,999999995 | 1 |
| XPNPEP1 | 17 | 17 | 1,24086E-18 | 0,999999999 | 1 |
| CYP2D22 | 6 | 5 | -0,013792113 | 0,721233154 | 1 |
| PSME2 | 8 | 8 | 0,011954506 | 0,680479257 | 1 |
| SLC12A2 | 24 | 22 | -5,79641E-17 | 0,999999991 | 1 |
| ARAP2 | 1 | 1 | 0 | 1 | 1 |
| OAT | 17 | 17 | 0,006699765 | 0,753736461 | 1 |
| CLTB | 2 | 2 | 0,149574056 | 0,202678504 | 1 |
| CMPK2 | 14 | 14 | 6,97543E-17 | 0,999999976 | 1 |
| ARHGAP12 | 4 | 4 | -0,067857446 | 0,204885425 | 1 |
| ENO2 | 29 | 29 | 0 | 1 | 1 |
| SYNGR3 | 4 | 4 | 6,89177E-18 | 0,999999999 | 1 |
| PCNP | 4 | 4 | -0,035981043 | 0,596067185 | 1 |
| BCAM | 1 | 1 | 2,81095E-18 | 0,999999997 | 1 |
| GAD1 | 13 | 13 | -0,031860236 | 0,319283298 | 1 |
| PLCL1 | 14 | 14 | -9,55985E-17 | 0,999999978 | 1 |
| ALAD | 12 | 12 | 0 | 1 | 1 |
| UBR3 | 3 | 3 | 0 | 1 | 1 |
| ATP5MK | 4 | 4 | 1,21516E-15 | 0,999999928 | 1 |

| | | | | | |
|---|---|---|---|---|---|
| NUMBL | 11 | 11 | -4,0879E-17 | 0,999999987 | 1 |
| MRI1 | 5 | 5 | -3,4577E-18 | 0,999999997 | 1 |
| SRCIN1 | 10 | 10 | 1,17687E-12 | 0,999996343 | 1 |
| EPHB2 | 10 | 10 | -2,09658E-17 | 0,999999991 | 1 |
| ANKRD63 | 3 | 3 | 0 | 1 | 1 |
| DCTN4 | 13 | 13 | 0 | 1 | 1 |
| TMEM160 | 4 | 4 | 0 | 1 | 1 |
| DYNC1LI1 | 26 | 26 | -2,31462E-16 | 0,99999997 | 1 |
| LGI3 | 11 | 10 | 0,01251308 | 0,592147447 | 1 |
| DHRS1 | 10 | 10 | 0 | 1 | 1 |
| TNKS1BP1 | 36 | 36 | -0,016646685 | 0,566428085 | 1 |
| SEC11C | 4 | 4 | 0 | 1 | 1 |
| DIAPH1 | 12 | 12 | -0,045451775 | 0,151593026 | 1 |
| HADH | 7 | 7 | 1,16161E-17 | 0,999999993 | 1 |
| TTC7B | 29 | 26 | 0,009767371 | 0,58858706 | 1 |
| ETFB | 6 | 6 | 1,78105E-17 | 0,999999992 | 1 |
| SLC6A1 | 11 | 11 | 9,15087E-18 | 0,999999996 | 1 |
| MRPL2 | 2 | 2 | 0 | 1 | 1 |
| LYSMD2 | 3 | 3 | -4,42156E-17 | 0,999999992 | 1 |
| HDGF | 12 | 12 | -3,6071E-17 | 0,999999989 | 1 |
| LMO7 | 4 | 1 | 0 | 1 | 1 |
| DDC | 14 | 14 | -4,1382E-17 | 0,999999987 | 1 |
| STAM | 14 | 14 | -0,055229185 | 0,244734757 | 1 |
| PDCL | 3 | 3 | 0 | 1 | 1 |
| NRAP | 1 | 1 | 0 | 1 | 1 |
| RPS18 | 12 | 12 | 3,33324E-17 | 0,999999989 | 1 |
| MRPL58 | 2 | 2 | 0 | 1 | 1 |
| SEC14L2 | 18 | 18 | 1,27002E-16 | 0,999999962 | 1 |
| PALS2 | 22 | 22 | -0,058025409 | 0,181739805 | 1 |
| SERPINA6 | 3 | 3 | 0,51093955 | 0,132444163 | 1 |
| ECPAS | 20 | 20 | 0,027688929 | 0,194111356 | 1 |
| USP11 | 16 | 16 | -4,1749E-16 | 0,999999945 | 1 |
| DMWD | 7 | 7 | 1,0604E-16 | 0,99999997 | 1 |

| | | | | | |
|---|---|---|---|---|---|
| MTFP1 | 6 | 6 | 0,040156696 | 0,290029666 | 1 |
| ACTA1 | 23 | 22 | 0 | 1 | 1 |
| DNAJC11 | 22 | 22 | 2,84248E-15 | 0,999999764 | 1 |
| USP7 | 29 | 29 | 2,90133E-17 | 0,999999989 | 1 |
| IPO4 | 9 | 9 | -0,028163095 | 0,274171125 | 1 |
| PCYOX1 | 14 | 13 | 4,16979E-18 | 0,999999994 | 1 |
| LGI1 | 24 | 24 | 9,14034E-18 | 0,999999998 | 1 |
| FBLN1 | 1 | 1 | 0 | 1 | 1 |
| SYNGR1 | 3 | 3 | 9,87265E-17 | 0,999999974 | 1 |
| CLASP2 | 42 | 41 | -7,85973E-18 | 0,999999994 | 1 |
| MIB2 | 2 | 2 | 2,26758E-18 | 0,999999998 | 1 |
| EIF2S3X | 16 | 15 | -4,63079E-16 | 0,999999916 | 1 |
| PGK1 | 33 | 32 | 0 | 1 | 1 |
| XPO5 | 14 | 14 | 0 | 1 | 1 |
| CACNG8 | 11 | 11 | -5,78271E-17 | 0,99999998 | 1 |
| USP31 | 8 | 7 | -5,86208E-15 | 0,999999829 | 1 |
| CACNA1C | 7 | 7 | -3,51482E-19 | 1 | 1 |
| TRAPPC4 | 4 | 4 | 0 | 1 | 1 |
| PSMB1 | 12 | 12 | -2,68033E-16 | 0,999999962 | 1 |
| ARMC6 | 7 | 7 | -1,52201E-16 | 0,999999978 | 1 |
| TANGO2 | 2 | 2 | 0,045224175 | 0,326633037 | 1 |
| ACAT3 | 11 | 11 | 0 | 1 | 1 |
| PADI2 | 10 | 10 | -0,045359611 | 0,234140702 | 1 |
| YARS2 | 9 | 7 | -7,28152E-16 | 0,999999932 | 1 |
| PAM16 | 7 | 6 | 0 | 1 | 1 |
| GAPDHS | 1 | 1 | 0 | 1 | 1 |
| COX6C | 11 | 11 | 0 | 1 | 1 |
| IRGQ | 13 | 13 | 0 | 1 | 1 |
| TRAPPC11 | 12 | 11 | -0,003447105 | 0,773215863 | 1 |
| ACAT1 | 27 | 27 | 0,056628509 | 0,145225019 | 1 |
| AHCY | 28 | 27 | 3,79015E-17 | 0,999999994 | 1 |
| TMCC2 | 6 | 6 | 0 | 1 | 1 |
| SMIM20 | 1 | 1 | 0 | 1 | 1 |

| | | | | | |
|---|---|---|---|---|---|
| GSN | 20 | 20 | 0 | 1 | 1 |
| RPL10A | 14 | 14 | -0,021658534 | 0,404620836 | 1 |
| DPP8 | 10 | 10 | -0,001593098 | 0,860447943 | 1 |
| DPP9 | 13 | 13 | 4,04478E-16 | 0,999999946 | 1 |
| PIP5K1C | 25 | 25 | 0 | 1 | 1 |
| UROD | 9 | 9 | 0 | 1 | 1 |
| CALB1 | 26 | 26 | -0,073484678 | 0,14318933 | 1 |
| FAAH | 14 | 13 | 5,79413E-17 | 0,999999981 | 1 |
| DPP3 | 31 | 31 | 6,04442E-18 | 0,999999996 | 1 |
| CPT2 | 11 | 9 | 0 | 1 | 1 |
| TIMM9 | 4 | 4 | 0 | 1 | 1 |
| ARL5A | 2 | 2 | -1,28057E-06 | 0,996584416 | 1 |
| FDXR | 11 | 11 | 2,08573E-17 | 0,999999987 | 1 |
| VPS33B | 10 | 10 | -6,00603E-17 | 0,999999976 | 1 |
| EIF2S1 | 19 | 19 | -0,015242564 | 0,415209016 | 1 |
| PPP2R5D | 24 | 24 | 5,1878E-16 | 0,999999922 | 1 |
| MACF1 | 72 | 65 | 1,03387E-18 | 0,999999999 | 1 |
| ADGRL1 | 25 | 25 | 7,73392E-17 | 0,999999968 | 1 |
| HECTD3 | 11 | 11 | 6,12797E-18 | 0,999999994 | 1 |
| MTCH1 | 13 | 13 | 0,020518929 | 0,397793162 | 1 |
| AMPD2 | 22 | 22 | -5,89088E-18 | 0,999999999 | 1 |
| H2AC20;HIST2H2A/ | 4 | 3 | -1,6752E-18 | 0,999999998 | 1 |
| TOM1 | 12 | 12 | -0,041285003 | 0,385138925 | 1 |
| HTRA2 | 9 | 8 | -5,25304E-17 | 0,999999978 | 1 |
| MON2 | 12 | 12 | -3,05487E-16 | 0,999999947 | 1 |
| PRPSAP1 | 11 | 11 | -2,26264E-16 | 0,999999949 | 1 |
| SNTB2 | 4 | 3 | 0,083218839 | 0,160467511 | 1 |
| MRPL45 | 3 | 2 | 0 | 1 | 1 |
| LRRC8C | 5 | 5 | 0 | 1 | 1 |
| SLC27A1 | 8 | 8 | -0,074846731 | 0,145262618 | 1 |
| DGKD | 4 | 4 | 0 | 1 | 1 |
| PDPK1 | 17 | 17 | -7,76461E-19 | 0,999999998 | 1 |
| SLK | 20 | 19 | -2,23191E-16 | 0,999999942 | 1 |

| | | | | | |
|---|---|---|---|---|---|
| STK4 | 4 | 4 | -4,4766E-18 | 0,999999994 | 1 |
| TAOK1 | 15 | 13 | 0,002512882 | 0,76667021 | 1 |
| CUL4A | 15 | 15 | -5,10112E-17 | 0,999999976 | 1 |
| CBS | 8 | 8 | -0,149670225 | 0,13930411 | 1 |
| CUL9 | 2 | 2 | 1,55823E-18 | 0,999999999 | 1 |
| NF1 | 10 | 10 | 0 | 1 | 1 |
| CPT1A | 9 | 8 | -3,63608E-18 | 0,999999997 | 1 |
| CNTNAP4 | 11 | 10 | 0 | 1 | 1 |
| CAMKK2 | 14 | 14 | 0 | 1 | 1 |
| STAU2 | 4 | 4 | -2,80984E-18 | 0,999999997 | 1 |
| COG7 | 6 | 6 | -6,64181E-18 | 0,999999997 | 1 |
| ANKRD34B | 3 | 3 | -0,054437905 | 0,25534811 | 1 |
| SYNCRIP | 21 | 21 | -5,94172E-17 | 0,999999993 | 1 |
| SRC | 18 | 17 | 1,24712E-16 | 0,999999961 | 1 |
| PHACTR1 | 15 | 15 | 5,09163E-17 | 0,999999971 | 1 |
| MAP2K5 | 2 | 2 | -3,93541E-15 | 0,999999867 | 1 |
| ATL1 | 26 | 25 | -3,75653E-17 | 0,999999988 | 1 |
| MRPL11 | 6 | 5 | 0,153257191 | 0,171445847 | 1 |
| PGD | 27 | 26 | 1,35517E-16 | 0,99999997 | 1 |
| LMAN2L | 5 | 5 | 0 | 1 | 1 |
| SMPD3 | 11 | 11 | -0,030334781 | 0,299841158 | 1 |
| TBC1D24 | 21 | 21 | 2,76596E-19 | 1 | 1 |
| ITSN2 | 14 | 13 | 0 | 1 | 1 |
| MIA3 | 3 | 3 | -5,66749E-15 | 0,999999857 | 1 |
| DLG3 | 44 | 43 | 1,10634E-17 | 0,999999992 | 1 |
| MYO5B;MYO5A | 6 | 6 | -9,71123E-18 | 0,999999992 | 1 |
| ATP2B2 | 37 | 37 | -0,030828609 | 0,192369 | 1 |
| MAP2K1 | 20 | 19 | -0,043301692 | 0,277812736 | 1 |
| REM2 | 7 | 7 | 0,036728551 | 0,401056339 | 1 |
| TPPP | 14 | 14 | -0,05562186 | 0,385857026 | 1 |
| ASTN1 | 25 | 25 | 0,047285404 | 0,132726718 | 1 |
| CARMIL2 | 15 | 15 | 8,99959E-17 | 0,999999993 | 1 |
| CLIC4 | 9 | 9 | 2,46583E-15 | 0,999999891 | 1 |

| | | | | | |
|---|---|---|---|---|---|
| TMEM229A | 1 | 1 | -0,026646163 | 0,704573311 | 1 |
| WNK2 | 20 | 20 | -0,033623368 | 0,209089813 | 1 |
| MYO6 | 11 | 11 | 0 | 1 | 1 |
| PPIP5K1 | 7 | 7 | 0 | 1 | 1 |
| SLC39A12 | 6 | 6 | -0,069186051 | 0,276276393 | 1 |
| FMNL1 | 17 | 17 | -7,63797E-17 | 0,999999985 | 1 |
| DPY19L1 | 1 | 1 | 0 | 1 | 1 |
| TTBK1 | 4 | 4 | 4,70779E-15 | 0,999999911 | 1 |
| PSMG1 | 3 | 3 | 6,70236E-16 | 0,999999962 | 1 |
| PPM1H | 20 | 19 | 0 | 1 | 1 |
| FAM13C | 7 | 7 | 1,4302E-18 | 0,999999999 | 1 |
| CAPN1 | 15 | 15 | -4,4798E-16 | 0,99999994 | 1 |
| ACTN1 | 48 | 47 | -0,011877214 | 0,533152325 | 1 |
| ACTN2 | 31 | 29 | 0,021278864 | 0,341595457 | 1 |
| ITGAV | 17 | 17 | -3,72332E-17 | 0,999999995 | 1 |
| PCDH19 | 6 | 6 | 1,40987E-20 | 1 | 1 |
| TTC9C | 4 | 4 | 1,2633E-17 | 0,999999993 | 1 |
| PRKCB | 38 | 38 | 1,16247E-15 | 0,999999902 | 1 |
| NAGK | 10 | 10 | 0,012951885 | 0,617402682 | 1 |
| CKMT1 | 28 | 26 | 1,49321E-15 | 0,999999862 | 1 |
| HSD17B8 | 10 | 10 | 0 | 1 | 1 |
| UNC45B | 3 | 3 | 0 | 1 | 1 |
| BIRC6 | 27 | 27 | -0,007820471 | 0,574150988 | 1 |
| PDCD6 | 9 | 9 | -1,45426E-16 | 0,999999963 | 1 |
| RPL7A | 9 | 9 | 5,58315E-16 | 0,999999949 | 1 |
| CMBL | 5 | 4 | 9,16922E-20 | 1 | 1 |
| CLIC6 | 16 | 15 | -0,28818495 | 0,450796632 | 1 |
| PAK5 | 7 | 6 | -1,2391E-18 | 0,999999998 | 1 |
| FXR2 | 10 | 10 | -0,012241741 | 0,507830742 | 1 |
| WDR77 | 8 | 8 | -1,49021E-17 | 0,999999991 | 1 |
| WDR1 | 33 | 33 | 0 | 1 | 1 |
| GJC2 | 2 | 2 | 6,77532E-19 | 0,999999999 | 1 |
| PPP3CA | 29 | 29 | -1,48706E-17 | 0,999999989 | 1 |

| | | | | | |
|---|---|---|---|---|---|
| TRAF3 | 14 | 13 | 9,2004E-16 | 0,999999927 | 1 |
| ARPC1A | 18 | 18 | 0 | 1 | 1 |
| ARSA | 4 | 4 | 0,08411512 | 0,151163556 | 1 |
| AP1M1 | 17 | 17 | -0,02866709 | 0,237596632 | 1 |
| ILDR2 | 7 | 7 | 0,01310427 | 0,573447942 | 1 |
| ATP5F1B | 43 | 43 | 0,071466528 | 0,11447989 | 1 |
| NTRK3 | 5 | 5 | 1,0533E-16 | 0,999999967 | 1 |
| HDHD3 | 7 | 7 | 0 | 1 | 1 |
| PCYT2 | 14 | 14 | 0 | 1 | 1 |
| GGACT | 3 | 3 | 1,53679E-16 | 0,999999979 | 1 |
| sp\|Q91V76\|CK054 | 9 | 9 | 1,84824E-16 | 0,99999998 | 1 |
| SRR | 12 | 12 | -0,038166053 | 0,302077871 | 1 |
| ADSS2 | 17 | 17 | 0 | 1 | 1 |
| ATP9B | 3 | 3 | 0 | 1 | 1 |
| VPS39 | 6 | 6 | 0,016259438 | 0,607910383 | 1 |
| RPS23 | 5 | 5 | -9,69308E-18 | 0,999999994 | 1 |
| ARMT1 | 17 | 17 | 4,08311E-17 | 0,999999983 | 1 |
| VTI1A | 6 | 6 | 8,96303E-18 | 0,999999993 | 1 |
| FBXO7 | 5 | 5 | -2,46612E-18 | 0,999999997 | 1 |
| PABPC4 | 12 | 12 | -0,033175639 | 0,316342201 | 1 |
| ANXA11 | 18 | 18 | 0 | 1 | 1 |
| ADSS1 | 17 | 17 | 0 | 1 | 1 |
| PRCP | 2 | 2 | -2,32872E-18 | 0,999999998 | 1 |
| KIF21A | 37 | 35 | -2,24357E-16 | 0,999999936 | 1 |
| STXBP5 | 23 | 22 | 0,025008345 | 0,362498794 | 1 |
| GPX1 | 13 | 13 | 0,036874575 | 0,310575559 | 1 |
| TPD52L2 | 3 | 3 | -6,29743E-18 | 0,999999996 | 1 |
| PLD2 | 4 | 4 | 1,83221E-18 | 0,999999998 | 1 |
| FAF2 | 10 | 10 | 0 | 1 | 1 |
| CRAT | 17 | 16 | 0 | 1 | 1 |
| PPP6C | 8 | 8 | 4,20631E-18 | 0,999999998 | 1 |
| PPP2CB;PPP2CA | 14 | 14 | -1,53622E-17 | 0,999999993 | 1 |
| MGST3 | 5 | 5 | 0,049265429 | 0,408621025 | 1 |

| | | | | | |
|---|---|---|---|---|---|
| SNAP23 | 3 | 3 | 2,75706E-19 | 1 | 1 |
| PPP1CA;PPP1CB | 9 | 9 | -9,02882E-18 | 0,999999994 | 1 |
| CNTN1 | 68 | 67 | 0 | 1 | 1 |
| CADM1 | 11 | 11 | 0 | 1 | 1 |
| MINK1 | 22 | 22 | 4,97936E-17 | 0,999999967 | 1 |
| RPL18A | 6 | 6 | 0 | 1 | 1 |
| CSDE1 | 32 | 32 | -0,012354987 | 0,411344239 | 1 |
| DIP2A | 14 | 14 | 0 | 1 | 1 |
| UGP2 | 24 | 24 | 0 | 1 | 1 |
| MTND4 | 10 | 10 | 0 | 1 | 1 |
| HOMER2 | 14 | 13 | 0 | 1 | 1 |
| CTNNA1 | 21 | 21 | -1,04884E-16 | 0,999999974 | 1 |
| CTNNA2 | 47 | 44 | 3,06649E-16 | 0,99999993 | 1 |
| DAAM1 | 18 | 17 | 9,79756E-15 | 0,999999687 | 1 |
| DAAM2 | 9 | 8 | 0 | 1 | 1 |
| FGB | 12 | 4 | -2,67628E-19 | 1 | 1 |
| LRRC4B | 12 | 12 | 0,029441324 | 0,272888753 | 1 |
| ZC2HC1A | 13 | 13 | 0 | 1 | 1 |
| ADCY5 | 14 | 13 | -3,52093E-17 | 0,999999985 | 1 |
| PAFAH1B1 | 30 | 30 | 4,5054E-17 | 0,999999985 | 1 |
| BPNT1 | 1 | 1 | 3,08531E-20 | 1 | 1 |
| BPNT1 | 17 | 16 | -4,44224E-17 | 0,999999992 | 1 |
| CCT8 | 50 | 49 | -0,010626786 | 0,507382669 | 1 |
| NAPB;NAPA | 4 | 4 | 0,003921396 | 0,79592125 | 1 |
| NLGN2 | 15 | 15 | 7,27092E-16 | 0,999999911 | 1 |
| AIFM1 | 28 | 28 | 2,86116E-17 | 0,999999993 | 1 |
| SLC4A4 | 23 | 23 | 0,076340513 | 0,133325801 | 1 |
| PFKM | 40 | 37 | 1,92555E-16 | 0,999999957 | 1 |
| LRP5 | 1 | 1 | 0 | 1 | 1 |
| MRPS9 | 5 | 5 | -4,53329E-18 | 0,999999996 | 1 |
| GATD1 | 5 | 5 | 0 | 1 | 1 |
| NCAM2 | 27 | 27 | -0,02655511 | 0,374796516 | 1 |
| NAPA | 20 | 20 | 3,63069E-16 | 0,999999913 | 1 |

| | | | | | |
|---|---|---|---|---|---|
| ST13 | 16 | 14 | 0 | 1 | 1 |
| ADAM10 | 6 | 6 | 0 | 1 | 1 |
| DLGAP4 | 17 | 17 | 7,05356E-17 | 0,999999964 | 1 |
| DHCR7 | 4 | 4 | 1,2361E-19 | 1 | 1 |
| GNPDA1 | 12 | 12 | 0 | 1 | 1 |
| EPHA4;EPHA7 | 1 | 1 | 0 | 1 | 1 |
| STMN1 | 7 | 7 | -0,053051163 | 0,150562261 | 1 |
| TRMT5 | 3 | 3 | -4,65152E-16 | 0,999999965 | 1 |
| MTMR6 | 6 | 6 | 0 | 1 | 1 |
| TTC22 | 1 | 1 | 0 | 1 | 1 |
| NMT1 | 17 | 16 | -9,21065E-17 | 0,999999964 | 1 |
| OTULIN | 3 | 3 | 0 | 1 | 1 |
| ACOT11 | 6 | 6 | 0 | 1 | 1 |
| PPP1R7 | 32 | 32 | -2,75491E-12 | 0,999993606 | 1 |
| GGCT | 10 | 10 | 5,6635E-18 | 0,999999993 | 1 |
| STX4 | 7 | 7 | -0,01431731 | 0,662938586 | 1 |
| BAG5 | 12 | 12 | -4,97763E-17 | 0,999999987 | 1 |
| AP1S1 | 7 | 7 | 1,63507E-17 | 0,999999992 | 1 |
| EIF2A | 9 | 9 | -6,11278E-16 | 0,999999922 | 1 |
| STX1B | 24 | 24 | 0,017667573 | 0,288368342 | 1 |
| ASPH | 3 | 3 | 0 | 1 | 1 |
| RPL35A | 9 | 9 | -0,011551101 | 0,626928272 | 1 |
| PNPLA8 | 17 | 14 | 0,04650869 | 0,291721193 | 1 |
| CORO1C | 23 | 22 | 7,08772E-18 | 0,999999993 | 1 |
| GABARAPL2 | 6 | 6 | -1,27469E-17 | 0,99999999 | 1 |
| DHX30 | 11 | 11 | -3,58668E-17 | 0,999999987 | 1 |
| TCP11L1 | 6 | 6 | 2,38966E-17 | 0,999999987 | 1 |
| PVALB | 10 | 9 | 0 | 1 | 1 |
| CTSD | 19 | 19 | 0 | 1 | 1 |
| PPM1F | 9 | 9 | 0 | 1 | 1 |
| EXOG | 16 | 14 | 4,86001E-17 | 0,999999983 | 1 |
| AUH | 17 | 16 | 0,019411132 | 0,39431439 | 1 |
| DGKQ | 14 | 14 | 0,008185118 | 0,58616495 | 1 |

| | | | | | |
|---|---|---|---|---|---|
| PSMA5 | 12 | 12 | -7,03925E-16 | 0,999999919 | 1 |
| NFS1 | 20 | 20 | -7,4932E-18 | 0,999999992 | 1 |
| PFKL | 37 | 37 | 0,056527777 | 0,123407558 | 1 |
| APPL2;APPL1 | 1 | 1 | -5,32744E-20 | 1 | 1 |
| PSMB4 | 13 | 13 | 0 | 1 | 1 |
| FMNL2 | 9 | 9 | 0 | 1 | 1 |
| XPO1 | 35 | 35 | -0,062972601 | 0,160186896 | 1 |
| RPS7 | 11 | 11 | -0,049142577 | 0,28694679 | 1 |
| ABCA2 | 3 | 3 | 0 | 1 | 1 |
| NLGN3 | 13 | 13 | 0 | 1 | 1 |
| CLTA | 10 | 9 | -0,056701355 | 0,319272129 | 1 |
| TBC1D10A | 3 | 3 | -6,30824E-17 | 0,999999979 | 1 |
| PTGR2 | 10 | 10 | -2,59436E-15 | 0,999999857 | 1 |
| PTPN9 | 16 | 16 | 2,13947E-16 | 0,999999955 | 1 |
| AP3S1 | 7 | 7 | 0,018972586 | 0,513879474 | 1 |
| BDH1 | 23 | 22 | 0,002932696 | 0,783111826 | 1 |
| GSTA2;GSTA1 | 1 | 1 | 0 | 1 | 1 |
| LTA4H | 41 | 41 | -1,32832E-16 | 0,999999975 | 1 |
| AIMP1 | 13 | 13 | -0,026155586 | 0,310258852 | 1 |
| ESRRB | 1 | 1 | 0 | 1 | 1 |
| CARM1 | 7 | 7 | 7,54943E-19 | 0,999999999 | 1 |
| TBC1D1 | 2 | 2 | -1,76492E-19 | 1 | 1 |
| KALRN | 35 | 34 | 0,040092368 | 0,165443787 | 1 |
| DMXL1;DMXL2 | 2 | 2 | -1,64932E-17 | 0,999999996 | 1 |
| STAT3 | 4 | 4 | -0,015659977 | 0,628813522 | 1 |
| GSTA4 | 13 | 13 | 9,5325E-18 | 0,999999999 | 1 |
| NAXD | 15 | 15 | 3,41142E-16 | 0,999999932 | 1 |
| TUBB5;TUBB2A;TU | 5 | 5 | -6,68138E-17 | 0,999999989 | 1 |
| ARL1 | 4 | 4 | -4,64302E-19 | 0,999999999 | 1 |
| SDR39U1 | 10 | 9 | 9,09961E-17 | 0,999999978 | 1 |
| SYT7 | 23 | 22 | 5,24441E-16 | 0,999999936 | 1 |
| ANXA6 | 52 | 52 | -1,09165E-17 | 0,999999994 | 1 |
| QARS1 | 25 | 25 | 0 | 1 | 1 |

| | | | | | |
|---|---|---|---|---|---|
| ASAP2 | 14 | 14 | -5,0762E-18 | 0,999999995 | 1 |
| ZER1 | 10 | 9 | 0 | 1 | 1 |
| ARHGAP1 | 18 | 18 | 7,33394E-18 | 0,999999993 | 1 |
| ROBO2 | 21 | 19 | 7,7578E-17 | 0,999999965 | 1 |
| PSMA6 | 13 | 12 | 6,25145E-18 | 0,999999998 | 1 |
| GTF2A2 | 1 | 1 | -0,074942064 | 0,316154271 | 1 |
| HIP1R | 24 | 24 | 0 | 1 | 1 |
| IGSF8 | 18 | 17 | 5,33887E-17 | 0,999999983 | 1 |
| EFL1 | 6 | 6 | 0 | 1 | 1 |
| EDF1 | 4 | 3 | 5,60315E-18 | 0,999999997 | 1 |
| DNM2 | 13 | 12 | 0 | 1 | 1 |
| KIT | 17 | 17 | 0,062441326 | 0,154618981 | 1 |
| PPID | 21 | 21 | 0,032980473 | 0,21310026 | 1 |
| ABCB6 | 3 | 3 | -1,68121E-19 | 1 | 1 |
| sp\|Q9CYI0\|NJMU_ | 1 | 1 | 0,113584902 | 0,384798565 | 1 |
| THEM4 | 9 | 9 | 0,018272702 | 0,499642956 | 1 |
| MIOS | 9 | 9 | 0,034244398 | 0,35008427 | 1 |
| PSMD3 | 34 | 34 | -3,06016E-15 | 0,999999777 | 1 |
| TTC1 | 8 | 8 | -1,83136E-17 | 0,999999991 | 1 |
| ANAPC7 | 2 | 2 | 0 | 1 | 1 |
| COPS4 | 30 | 30 | -0,028401262 | 0,28365102 | 1 |
| ATXN3 | 3 | 3 | -6,8346E-16 | 0,999999966 | 1 |
| AKT2 | 3 | 3 | -9,62854E-20 | 1 | 1 |
| DNAJB12 | 4 | 4 | 1,80301E-14 | 0,9999996 | 1 |
| SNPH | 22 | 21 | 2,80358E-16 | 0,999999945 | 1 |
| MRPL46 | 5 | 5 | 2,37749E-18 | 0,999999998 | 1 |
| FGG | 10 | 9 | 1,43426E-17 | 0,999999993 | 1 |
| NDUFAF1 | 9 | 8 | 0 | 1 | 1 |
| D430041D05RIK | 15 | 15 | -2,25097E-18 | 0,999999999 | 1 |
| ICA1L | 3 | 3 | 1,74037E-18 | 0,999999999 | 1 |
| ICA1 | 11 | 11 | 0 | 1 | 1 |
| TMEM121B | 4 | 4 | -6,92402E-17 | 0,999999981 | 1 |
| ZPR1 | 5 | 5 | -0,011341806 | 0,664038093 | 1 |

| | | | | | |
|---|---|---|---|---|---|
| WIPF3 | 13 | 13 | -0,15626028 | 0,115922802 | 1 |
| PPP2R1A | 35 | 35 | -5,7087E-17 | 0,999999992 | 1 |
| SHISA6 | 21 | 19 | 6,94817E-06 | 0,988605803 | 1 |
| OLFR1135 | 1 | 1 | 0 | 1 | 1 |
| TIMM44 | 25 | 25 | 2,47139E-18 | 0,999999999 | 1 |
| LRSAM1 | 7 | 7 | 0 | 1 | 1 |
| ARFGAP2 | 8 | 8 | -7,86402E-13 | 0,999996751 | 1 |
| TSC2 | 20 | 20 | 0,043181762 | 0,139705913 | 1 |
| ACAN | 22 | 22 | 0 | 1 | 1 |
| TBRG4 | 12 | 12 | 4,18762E-17 | 0,999999993 | 1 |
| YTHDF3 | 5 | 5 | 0 | 1 | 1 |
| RABGEF1 | 6 | 6 | 0 | 1 | 1 |
| PLXNB2 | 16 | 16 | -8,46085E-18 | 0,999999992 | 1 |
| RPL24 | 5 | 5 | -4,78069E-17 | 0,99999999 | 1 |
| GCDH | 10 | 9 | 0,024091386 | 0,435856049 | 1 |
| HNRNPL | 3 | 2 | 1,11292E-16 | 0,999999978 | 1 |
| RAB11B;RAB11A | 14 | 14 | -1,19649E-18 | 0,999999998 | 1 |
| GRIN1 | 29 | 28 | 0,020415663 | 0,389049118 | 1 |
| UGGT1 | 21 | 20 | 1,92476E-16 | 0,999999971 | 1 |
| PSMA7 | 13 | 13 | 0 | 1 | 1 |
| NRXN2 | 19 | 19 | -5,07562E-17 | 0,999999971 | 1 |
| FHL1 | 3 | 3 | 0 | 1 | 1 |
| CHM | 10 | 10 | 1,66083E-17 | 0,99999999 | 1 |
| EFR3B | 24 | 24 | 0,046464282 | 0,171632678 | 1 |
| ADGRL2 | 7 | 6 | 0,032726671 | 0,466667268 | 1 |
| WNK3 | 2 | 2 | 0 | 1 | 1 |
| HNRNPUL1 | 9 | 9 | -2,36505E-05 | 0,982947893 | 1 |
| PAFAH1B3 | 9 | 8 | -0,011293467 | 0,649714722 | 1 |
| NCKAP1L | 8 | 7 | -4,59564E-22 | 1 | 1 |
| SOD2 | 13 | 12 | 0,043049505 | 0,221559426 | 1 |
| ABLIM1 | 17 | 15 | 0 | 1 | 1 |
| PEPD | 15 | 15 | 1,28448E-17 | 0,999999994 | 1 |
| ABLIM2 | 14 | 14 | 0 | 1 | 1 |

| | | | | | |
|---|---|---|---|---|---|
| PSMB5 | 14 | 14 | 8,99953E-17 | 0,999999968 | 1 |
| ABRAXAS2 | 8 | 8 | -4,81428E-16 | 0,999999934 | 1 |
| SCRN1 | 19 | 19 | -3,22346E-17 | 0,999999988 | 1 |
| FERMT2 | 15 | 14 | -0,062872923 | 0,16850487 | 1 |
| EZR | 25 | 25 | -3,33364E-17 | 0,999999986 | 1 |
| ATP1B1 | 19 | 19 | 0,028215243 | 0,152326873 | 1 |
| TMEM38A | 1 | 1 | 0 | 1 | 1 |
| GPM6B | 5 | 5 | 0 | 1 | 1 |
| 4932415D10RIK | 3 | 3 | -0,086160949 | 0,179242685 | 1 |
| PABPC1 | 30 | 29 | -0,027739884 | 0,153195296 | 1 |
| SNCA | 8 | 8 | 0 | 1 | 1 |
| INPP4A | 13 | 13 | 0,022341554 | 0,438251104 | 1 |
| REEP5 | 4 | 4 | 0,09644864 | 0,145700967 | 1 |
| SARS | 30 | 30 | 0,020170631 | 0,423602927 | 1 |
| GLS | 31 | 31 | 0,003477696 | 0,761577121 | 1 |
| RTN2 | 1 | 1 | 0 | 1 | 1 |
| DLST | 21 | 19 | 0 | 1 | 1 |
| GFM2 | 6 | 5 | -3,30571E-17 | 0,999999992 | 1 |
| KIAA1107 | 19 | 19 | -3,37999E-17 | 0,999999983 | 1 |
| FAM217B | 2 | 1 | 0 | 1 | 1 |
| PHPT1 | 8 | 7 | -1,85677E-17 | 0,999999991 | 1 |
| RPS19 | 12 | 12 | -0,044265355 | 0,170257225 | 1 |
| ARPC5 | 6 | 6 | 5,19815E-17 | 0,999999989 | 1 |
| SCRIB | 6 | 6 | -0,030581954 | 0,440458326 | 1 |
| UAP1 | 6 | 6 | -1,85726E-16 | 0,999999957 | 1 |
| ARMC10 | 9 | 8 | 2,15345E-17 | 0,999999995 | 1 |
| GCN1 | 32 | 31 | -0,025041603 | 0,246102299 | 1 |
| SRPRA | 6 | 6 | -4,37527E-18 | 0,999999996 | 1 |
| SLC29A1 | 2 | 2 | 0,023002045 | 0,611725832 | 1 |
| GRPEL1 | 10 | 10 | 0 | 1 | 1 |
| THOP1 | 24 | 24 | -1,72244E-17 | 0,999999994 | 1 |
| CRPPA | 6 | 6 | 2,78031E-17 | 0,999999984 | 1 |
| DYNC1I1 | 16 | 16 | -3,31381E-15 | 0,999999823 | 1 |

| | | | | | |
|---|---|---|---|---|---|
| RAP2A | 7 | 7 | 2,15123E-15 | 0,999999838 | 1 |
| GSS | 18 | 18 | 0 | 1 | 1 |
| RHEB | 14 | 14 | -0,034372218 | 0,273753149 | 1 |
| FAR1 | 4 | 4 | -0,071604446 | 0,23757024 | 1 |
| CSPG5 | 9 | 9 | 7,05397E-18 | 0,999999996 | 1 |
| KLC4 | 7 | 7 | 1,6296E-16 | 0,999999972 | 1 |
| ECHDC2 | 2 | 1 | 0 | 1 | 1 |
| ESYT2 | 10 | 10 | -1,94024E-16 | 0,999999965 | 1 |
| BCS1L | 18 | 17 | 9,95177E-19 | 0,999999999 | 1 |
| SH2B1 | 7 | 7 | -0,030204117 | 0,29997258 | 1 |
| NCSTN | 8 | 8 | 0 | 1 | 1 |
| P4HB | 29 | 29 | 0 | 1 | 1 |
| SPR | 16 | 16 | 0 | 1 | 1 |
| TXNDC5 | 9 | 9 | -2,81981E-17 | 0,999999992 | 1 |
| CTBP1;CTBP2 | 5 | 5 | -0,01505023 | 0,604791305 | 1 |
| GLUD1 | 47 | 45 | 0 | 1 | 1 |
| TSC22D4 | 1 | 1 | 0 | 1 | 1 |
| ATL3 | 6 | 5 | -0,027913581 | 0,52578647 | 1 |
| FAHD2A | 15 | 15 | 3,44657E-20 | 1 | 1 |
| ARPC5L | 6 | 6 | -0,011891885 | 0,588815579 | 1 |
| FAM171A2 | 16 | 16 | 0 | 1 | 1 |
| TAMALIN | 4 | 4 | 5,02248E-17 | 0,999999989 | 1 |
| XPOT | 9 | 9 | -3,25156E-14 | 0,999999248 | 1 |
| TOMM34 | 18 | 18 | -0,044790503 | 0,282565296 | 1 |
| CST3 | 9 | 9 | -1,32039E-17 | 0,999999992 | 1 |
| PRAF2 | 5 | 5 | 1,11396E-05 | 0,988259201 | 1 |
| IKBKB | 2 | 2 | 1,08896E-11 | 0,999995649 | 1 |
| REXO2 | 4 | 3 | -2,01613E-15 | 0,999999909 | 1 |
| SCN2A | 21 | 21 | 1,76446E-17 | 0,999999993 | 1 |
| FCHSD2 | 1 | 1 | 0 | 1 | 1 |
| MAPK3 | 17 | 16 | -0,054931894 | 0,18144365 | 1 |
| UQCRB | 12 | 12 | 1,29743E-16 | 0,999999962 | 1 |
| GUSB | 1 | 1 | 0,112982426 | 0,27477684 | 1 |

| | | | | | |
|---|---|---|---|---|---|
| MINDY3 | 2 | 2 | 7,32485E-18 | 0,999999995 | 1 |
| PFKFB2 | 5 | 5 | -0,082792132 | 0,135066433 | 1 |
| ADGRB1 | 13 | 13 | 5,57849E-17 | 0,999999981 | 1 |
| STIM1 | 6 | 4 | -1,39331E-17 | 0,999999993 | 1 |
| SCRN3 | 16 | 16 | 1,07143E-16 | 0,999999969 | 1 |
| RBP1 | 6 | 6 | -9,25874E-17 | 0,999999983 | 1 |
| APEH | 16 | 16 | 0 | 1 | 1 |
| PCDHB10 | 1 | 1 | 0 | 1 | 1 |
| SPIRE1 | 3 | 3 | 1,22297E-16 | 0,999999985 | 1 |
| INA | 29 | 28 | 1,41525E-16 | 0,999999979 | 1 |
| EXOC7 | 27 | 26 | 3,10985E-16 | 0,999999931 | 1 |
| IDH1 | 31 | 31 | 1,53886E-17 | 0,99999999 | 1 |
| KRT42 | 4 | 4 | 0 | 1 | 1 |
| KRT14;KRT17 | 7 | 5 | 3,95098E-16 | 0,999999984 | 1 |
| RNH1 | 12 | 12 | -4,61868E-16 | 0,999999945 | 1 |
| TPPP3 | 5 | 5 | 4,21919E-18 | 0,999999998 | 1 |
| STMN2 | 4 | 4 | 0 | 1 | 1 |
| STMN3 | 9 | 9 | 0 | 1 | 1 |
| PUM1 | 1 | 1 | 0 | 1 | 1 |
| DYNC2H1 | 3 | 3 | 3,91836E-18 | 0,999999997 | 1 |
| CTPS1 | 12 | 12 | 2,16433E-17 | 0,999999984 | 1 |
| AGL | 43 | 41 | 5,33469E-18 | 0,999999997 | 1 |
| POGK | 1 | 1 | 0 | 1 | 1 |
| CCT7 | 32 | 31 | -2,02546E-16 | 0,999999962 | 1 |
| EPS15L1 | 34 | 34 | -6,56116E-16 | 0,999999916 | 1 |
| STUB1 | 11 | 11 | 0 | 1 | 1 |
| MYH11;MYH10 | 3 | 3 | 0,071850432 | 0,227953786 | 1 |
| PRR7 | 2 | 2 | 0,065152008 | 0,456121322 | 1 |
| GPT | 7 | 7 | -5,49771E-18 | 0,999999996 | 1 |
| MSN | 20 | 20 | 1,16865E-17 | 0,999999996 | 1 |
| SPAG1 | 1 | 1 | 0 | 1 | 1 |
| PEX5 | 3 | 2 | 0 | 1 | 1 |
| NCL | 24 | 24 | 0,015811885 | 0,715477961 | 1 |

| | | | | | |
|---|---|---|---|---|---|
| BRSK1 | 21 | 20 | 0,015840807 | 0,393881296 | 1 |
| LCP1 | 23 | 23 | 1,04275E-15 | 0,99999992 | 1 |
| PRODH | 16 | 13 | 1,65344E-17 | 0,999999991 | 1 |
| HTATSF1 | 5 | 5 | -0,075682944 | 0,132503139 | 1 |
| PDIA3 | 34 | 33 | -1,3472E-17 | 0,999999992 | 1 |
| ZFYVE1 | 10 | 10 | 0 | 1 | 1 |
| AVEN | 1 | 1 | 0 | 1 | 1 |
| SUGT1 | 17 | 17 | -0,041141787 | 0,138525889 | 1 |
| KIF5C | 46 | 46 | 0,007747192 | 0,511482661 | 1 |
| KIF5C;KIF5B | 3 | 3 | -1,78714E-16 | 0,999999968 | 1 |
| RUVBL1 | 12 | 12 | 0 | 1 | 1 |
| PREX1 | 14 | 14 | -0,001313575 | 0,871631803 | 1 |
| RPTOR | 14 | 14 | 0,004171782 | 0,743062146 | 1 |
| GTPBP3 | 2 | 2 | 0 | 1 | 1 |
| CACNB3;CACNB2;C | 4 | 4 | 0 | 1 | 1 |
| CRK | 21 | 20 | -2,09619E-17 | 0,999999991 | 1 |
| ABI2 | 7 | 7 | 0 | 1 | 1 |
| EMC2 | 5 | 5 | -1,49428E-16 | 0,999999963 | 1 |
| SCN11A | 1 | 1 | 0 | 1 | 1 |
| COQ6 | 13 | 13 | 0,027120431 | 0,44222787 | 1 |
| CATSPER1 | 1 | 1 | 0 | 1 | 1 |
| FTL1 | 11 | 10 | 0 | 1 | 1 |
| MRPL41 | 3 | 3 | 2,42119E-19 | 1 | 1 |
| ABCB8 | 21 | 20 | 0,01644532 | 0,38202627 | 1 |
| ETL4 | 31 | 31 | -2,13067E-16 | 0,999999945 | 1 |
| ABHD5 | 2 | 2 | 0,045574559 | 0,50169035 | 1 |
| BCAN | 32 | 32 | -0,011161462 | 0,634714506 | 1 |
| BABAM1 | 3 | 3 | 0 | 1 | 1 |
| PPM1A | 17 | 17 | -0,012067096 | 0,591018542 | 1 |
| PPM1B | 11 | 11 | 1,89951E-17 | 0,999999994 | 1 |
| ZNF428 | 1 | 1 | 0 | 1 | 1 |
| CS | 29 | 28 | -1,03496E-17 | 0,999999995 | 1 |
| PTGS1 | 3 | 3 | 0,047463526 | 0,446309424 | 1 |

| | | | | | |
|---|---|---|---|---|---|
| PI4KA | 68 | 64 | 0,018954346 | 0,350591586 | 1 |
| ACACB | 8 | 8 | 3,86818E-15 | 0,999999779 | 1 |
| SERPINE2 | 10 | 10 | 5,90667E-19 | 1 | 1 |
| GPD1L | 16 | 16 | 0 | 1 | 1 |
| KLC1 | 19 | 18 | -1,43893E-17 | 0,99999999 | 1 |
| FKBP5 | 12 | 12 | 1,33275E-17 | 0,999999989 | 1 |
| ATPAF1 | 9 | 9 | 0 | 1 | 1 |
| SVOP | 6 | 6 | 0,027000958 | 0,352687762 | 1 |
| MYO6 | 29 | 29 | -6,25954E-16 | 0,99999988 | 1 |
| AKR1A1 | 27 | 27 | 3,82418E-16 | 0,999999945 | 1 |
| PPP1R9A | 1 | 1 | 0 | 1 | 1 |
| RAP1GDS1 | 31 | 31 | 2,9153E-17 | 0,999999993 | 1 |
| RAP1GDS1 | 3 | 3 | 0 | 1 | 1 |
| LRP4 | 2 | 2 | 0,066656176 | 0,301633249 | 1 |
| VWA5B2 | 1 | 1 | 0 | 1 | 1 |
| MYL6B | 2 | 2 | 0 | 1 | 1 |
| FABP7 | 8 | 8 | -2,93872E-17 | 0,999999987 | 1 |
| CSL | 3 | 3 | -1,29709E-18 | 1 | 1 |
| AKR1B8 | 2 | 2 | 0,005238991 | 0,81088757 | 1 |
| OTUB2 | 5 | 5 | 2,46737E-16 | 0,999999958 | 1 |
| GLOD4 | 30 | 30 | -2,08792E-18 | 0,999999999 | 1 |
| SLIRP | 3 | 2 | 0 | 1 | 1 |
| DARS1 | 31 | 31 | 7,96737E-18 | 0,999999996 | 1 |
| LRFN1 | 12 | 12 | 4,86547E-17 | 0,999999983 | 1 |
| LRRC8A | 17 | 17 | 1,32902E-18 | 0,999999998 | 1 |
| LRFN5 | 5 | 5 | 0 | 1 | 1 |
| IPO13 | 4 | 4 | -0,031717499 | 0,488309427 | 1 |
| IQCB1 | 5 | 5 | -3,54801E-17 | 0,999999985 | 1 |
| MYRIP | 4 | 3 | 2,57403E-18 | 0,999999999 | 1 |
| CD200 | 10 | 10 | 9,3823E-17 | 0,99999998 | 1 |
| SUCLA2 | 43 | 41 | 3,85513E-17 | 0,999999984 | 1 |
| LYPLA1 | 3 | 3 | -1,64499E-18 | 0,999999998 | 1 |
| sp\|Q8R092\|CA043 | 3 | 3 | -4,60481E-20 | 1 | 1 |

| | | | | | |
|---|---|---|---|---|---|
| LYPLA2 | 9 | 9 | -3,13372E-16 | 0,999999957 | 1 |
| PLCL2 | 19 | 18 | 9,27871E-17 | 0,999999978 | 1 |
| ABI1 | 13 | 13 | 0 | 1 | 1 |
| NME3 | 8 | 8 | -0,051235318 | 0,117501899 | 1 |
| SLC25A11 | 23 | 23 | 4,5907E-18 | 0,999999994 | 1 |
| GMPPB | 8 | 8 | 0,03896958 | 0,256916894 | 1 |
| ENDOD1 | 12 | 11 | 3,83148E-18 | 0,999999996 | 1 |
| GPS1 | 13 | 13 | -0,006348232 | 0,634546342 | 1 |
| SLC25A19 | 6 | 5 | 0,066184587 | 0,225312107 | 1 |
| DOCK3 | 19 | 19 | 0 | 1 | 1 |
| DDX1 | 15 | 15 | -1,61094E-16 | 0,999999961 | 1 |
| NQO1 | 8 | 8 | -5,308E-16 | 0,999999961 | 1 |
| ARMCX3 | 7 | 7 | -0,026452749 | 0,345309403 | 1 |
| POLR2E | 1 | 1 | 0 | 1 | 1 |
| SVIL | 1 | 1 | 0 | 1 | 1 |
| SYNPR | 8 | 8 | 0,049158188 | 0,128653766 | 1 |
| STAMBPL1 | 5 | 5 | 0 | 1 | 1 |
| ACSL5 | 8 | 8 | -0,014521289 | 0,564430202 | 1 |
| HSPA9 | 54 | 52 | 0,055858312 | 0,142131286 | 1 |
| TMEM33 | 4 | 4 | 0 | 1 | 1 |
| SARS2 | 10 | 10 | 7,72941E-18 | 0,999999993 | 1 |
| CAP1 | 36 | 36 | -4,33641E-16 | 0,999999926 | 1 |
| PCDHGC5 | 8 | 7 | 7,16991E-18 | 0,999999996 | 1 |
| NMRAL1 | 15 | 15 | -0,027851016 | 0,506692328 | 1 |
| ADAP1 | 20 | 19 | 0,020021455 | 0,43737224 | 1 |
| HAGH | 11 | 11 | 1,19539E-18 | 0,999999999 | 1 |
| HSP90AA1 | 39 | 39 | 3,39392E-15 | 0,999999825 | 1 |
| FUT11 | 1 | 1 | -6,15037E-18 | 0,999999995 | 1 |
| ELP5 | 2 | 2 | 6,41112E-17 | 0,999999982 | 1 |
| KCNMB4 | 1 | 1 | 0 | 1 | 1 |
| ANLN | 6 | 6 | 0 | 1 | 1 |
| FAM20B | 3 | 1 | 0 | 1 | 1 |
| TPRG1L | 10 | 10 | 0,015361356 | 0,596017428 | 1 |

| | | | | | |
|---|---|---|---|---|---|
| PNMA8B | 1 | 1 | -0,074640574 | 0,274944096 | 1 |
| ERO1A | 11 | 11 | -0,016499148 | 0,543066304 | 1 |
| VTI1B | 8 | 8 | -0,020206137 | 0,41392863 | 1 |
| VPS16 | 11 | 11 | 0 | 1 | 1 |
| PLD3 | 5 | 5 | -1,54014E-16 | 0,999999965 | 1 |
| PDP1 | 15 | 14 | 0,049373322 | 0,175580297 | 1 |
| VPS13D | 15 | 14 | 0,011977825 | 0,640703905 | 1 |
| RO60 | 10 | 10 | 4,41197E-17 | 0,99999998 | 1 |
| DNM1;DNM3 | 5 | 5 | 0,024841343 | 0,541770344 | 1 |
| PLEKHG5 | 7 | 7 | -8,61355E-16 | 0,999999924 | 1 |
| SRP68 | 7 | 7 | -2,21252E-17 | 0,999999986 | 1 |
| HEXA | 2 | 2 | 0 | 1 | 1 |
| CYB5R4 | 1 | 1 | 0 | 1 | 1 |
| ABCA7 | 3 | 3 | 0 | 1 | 1 |
| GAK | 25 | 25 | 0,0286722 | 0,179332283 | 1 |
| IPO11 | 8 | 8 | -0,001289107 | 0,891920904 | 1 |
| ANXA3 | 19 | 19 | 0 | 1 | 1 |
| GBF1 | 6 | 6 | 0 | 1 | 1 |
| ABHD16A | 17 | 17 | 5,25473E-15 | 0,999999761 | 1 |
| UBR1 | 10 | 10 | -1,13899E-17 | 0,999999995 | 1 |
| LIPT1 | 1 | 1 | 0 | 1 | 1 |
| HSPA4L | 52 | 51 | 0 | 1 | 1 |
| ETFDH | 24 | 23 | 2,81014E-17 | 0,999999979 | 1 |
| NADK2 | 23 | 21 | 0,046071736 | 0,113473222 | 1 |
| MAP2K4 | 13 | 13 | 0 | 1 | 1 |
| NDUFAF5 | 6 | 6 | 0,084596561 | 0,132281521 | 1 |
| CYFIP1 | 28 | 28 | 0,025416305 | 0,358584896 | 1 |
| EIF5 | 10 | 10 | -2,3801E-16 | 0,999999956 | 1 |
| TMEM245 | 1 | 1 | 0,029140039 | 0,656127601 | 1 |
| REV3L | 1 | 1 | 0 | 1 | 1 |
| PRDX5 | 15 | 15 | 0 | 1 | 1 |
| TTC5 | 2 | 2 | 0 | 1 | 1 |
| OSBP2 | 4 | 4 | 0 | 1 | 1 |

| | | | | | |
|---|---|---|---|---|---|
| PLCXD3 | 8 | 8 | 3,02474E-17 | 0,999999991 | 1 |
| ADGRA1 | 2 | 2 | -4,81744E-13 | 0,999999413 | 1 |
| IL16 | 1 | 1 | -0,028284182 | 0,666029419 | 1 |
| FAM126B | 18 | 17 | 0,00791678 | 0,629525851 | 1 |
| TMEM35A | 3 | 3 | -8,89428E-15 | 0,999999713 | 1 |
| OCIAD2 | 6 | 6 | 0,039582689 | 0,421685181 | 1 |
| PGAM2 | 7 | 7 | 0 | 1 | 1 |
| PGAM1 | 17 | 17 | 1,19199E-17 | 0,999999994 | 1 |
| RIN1 | 17 | 17 | 0,045795488 | 0,201211463 | 1 |
| CSAD | 8 | 8 | -2,61076E-17 | 0,999999989 | 1 |
| ELMO2 | 21 | 21 | -3,12663E-17 | 0,999999983 | 1 |
| SNX30 | 16 | 16 | -4,37048E-18 | 0,999999996 | 1 |
| sp\|Q9CWB7\|YD28( | 1 | 1 | 0 | 1 | 1 |
| CACNA1E | 18 | 16 | 2,99575E-17 | 0,999999988 | 1 |
| DCDC2B | 1 | 1 | 0 | 1 | 1 |
| GPC1 | 11 | 11 | 0,04745071 | 0,174426666 | 1 |
| LAMP1 | 5 | 5 | 0 | 1 | 1 |
| DDAH2 | 9 | 9 | 2,44222E-16 | 0,999999951 | 1 |
| SCAI | 22 | 21 | 5,82855E-17 | 0,999999969 | 1 |
| GRIPAP1 | 2 | 2 | -6,41038E-18 | 0,999999996 | 1 |
| DST | 41 | 38 | -2,09926E-16 | 0,999999944 | 1 |
| CDC42BPB | 31 | 30 | -2,10174E-17 | 0,999999987 | 1 |
| EXOC1 | 10 | 10 | 2,70718E-17 | 0,999999982 | 1 |
| CARS1 | 27 | 27 | 0 | 1 | 1 |
| CCT6A | 20 | 20 | -1,66864E-17 | 0,999999993 | 1 |
| IGTP | 1 | 1 | 0 | 1 | 1 |
| SLC4A1AP | 1 | 1 | 0 | 1 | 1 |
| CYP2D9 | 1 | 1 | -0,109907631 | 0,380641897 | 1 |
| AP2B1 | 35 | 35 | 1,23864E-16 | 0,999999983 | 1 |
| STAM2 | 5 | 5 | -1,34009E-16 | 0,999999966 | 1 |
| GAPVD1 | 22 | 22 | -0,010040885 | 0,472333373 | 1 |
| COG1 | 4 | 4 | 0 | 1 | 1 |
| HGS | 18 | 18 | -0,007231959 | 0,69492591 | 1 |

| | | | | | |
|---|---|---|---|---|---|
| MVP | 3 | 3 | -1,5081E-16 | 0,999999976 | 1 |
| BORCS6 | 2 | 2 | -2,79652E-16 | 0,999999982 | 1 |
| TMEM205 | 2 | 2 | 8,58433E-17 | 0,999999983 | 1 |
| CUL3 | 38 | 38 | 0 | 1 | 1 |
| IQGAP1 | 16 | 16 | -0,085003511 | 0,166725285 | 1 |
| AKR7A2 | 12 | 12 | 0,030396776 | 0,220565865 | 1 |
| GDE1 | 5 | 5 | 0 | 1 | 1 |
| AP1B1 | 29 | 28 | -0,031399204 | 0,223687753 | 1 |
| RTL8B | 1 | 1 | 0 | 1 | 1 |
| HSDL1 | 9 | 8 | 0,048960109 | 0,121145978 | 1 |
| CBR1 | 22 | 22 | -1,53414E-17 | 0,999999991 | 1 |
| PSMA1 | 19 | 19 | 1,57812E-18 | 0,999999998 | 1 |
| CHMP1A | 4 | 4 | 3,68863E-17 | 0,999999991 | 1 |
| SHMT1 | 2 | 2 | -1,39948E-19 | 1 | 1 |
| SLC18A2 | 1 | 1 | 0 | 1 | 1 |
| FIBCD1 | 4 | 4 | 0 | 1 | 1 |
| LRRC8D | 12 | 11 | 0 | 1 | 1 |
| LRRC8B | 5 | 5 | 5,60128E-19 | 0,999999999 | 1 |
| PRKCE | 39 | 39 | 8,10664E-18 | 0,999999991 | 1 |
| SEMA4A | 7 | 7 | -0,070504798 | 0,236704978 | 1 |
| OTUD6B | 6 | 6 | -0,007410828 | 0,692621125 | 1 |
| CDC23 | 7 | 7 | 1,39364E-17 | 0,999999994 | 1 |
| MCF2L | 5 | 5 | 0 | 1 | 1 |
| PRRC2B | 9 | 9 | -0,027765619 | 0,493622537 | 1 |
| MMS19 | 9 | 9 | 0 | 1 | 1 |
| UBQLN1 | 6 | 6 | -6,87322E-18 | 0,999999993 | 1 |
| UBQLN4 | 4 | 4 | -4,40093E-18 | 0,999999999 | 1 |
| TMEM214 | 1 | 1 | 0 | 1 | 1 |
| PCDH17 | 7 | 6 | 0 | 1 | 1 |
| AKAP5 | 20 | 20 | -0,049132682 | 0,215343087 | 1 |
| LETMD1 | 9 | 9 | 7,20338E-19 | 1 | 1 |
| SNX2 | 24 | 24 | 0 | 1 | 1 |
| GIGYF2 | 3 | 3 | -2,56883E-15 | 0,999999885 | 1 |

| | | | | | |
|---|---|---|---|---|---|
| GIGYF1 | 2 | 2 | -0,077273673 | 0,225184151 | 1 |
| RIPOR1 | 10 | 10 | -1,47804E-16 | 0,999999972 | 1 |
| OXSM | 11 | 11 | 0,01176581 | 0,626656763 | 1 |
| HEATR5B | 11 | 11 | 0 | 1 | 1 |
| ARFIP2 | 14 | 14 | 0 | 1 | 1 |
| ZZEF1 | 18 | 18 | 0 | 1 | 1 |
| FICD | 1 | 1 | 0 | 1 | 1 |
| ESYT1 | 2 | 2 | -2,07746E-15 | 0,999999901 | 1 |
| EPN3 | 5 | 5 | 0 | 1 | 1 |
| EIPR1 | 7 | 7 | 0,044558376 | 0,301601014 | 1 |
| MYADM | 4 | 4 | 0 | 1 | 1 |
| NQO2 | 6 | 6 | 2,29541E-17 | 0,99999999 | 1 |
| SMPD1 | 4 | 4 | 0 | 1 | 1 |
| ME2 | 15 | 14 | 0,041644165 | 0,344786048 | 1 |
| TUBB4B;TUBB6;TU | 2 | 2 | 0 | 1 | 1 |
| TUBB2A;TUBB2B | 4 | 4 | 0 | 1 | 1 |
| TUBB5 | 7 | 7 | -4,41614E-18 | 0,999999997 | 1 |
| FARSB | 21 | 21 | -0,037944427 | 0,189485521 | 1 |
| RPS16 | 10 | 10 | 6,23534E-17 | 0,999999989 | 1 |
| RPL14 | 5 | 5 | 6,59455E-19 | 0,999999999 | 1 |
| CRELD1 | 1 | 1 | -1,74791E-17 | 0,99999999 | 1 |
| MRRF | 9 | 8 | 0 | 1 | 1 |
| MYG1 | 12 | 11 | -1,08872E-16 | 0,999999976 | 1 |
| TTYH3 | 9 | 9 | -1,73127E-17 | 0,999999996 | 1 |
| GUF1 | 3 | 2 | 0 | 1 | 1 |
| VPS13A | 32 | 30 | 6,36001E-17 | 0,999999969 | 1 |
| PIP5K1A | 10 | 10 | -0,067066752 | 0,177575204 | 1 |
| RGS14 | 20 | 20 | 0,010583341 | 0,622440912 | 1 |
| FBXL15 | 3 | 3 | -0,044329861 | 0,45252699 | 1 |
| APOA4 | 11 | 11 | 0 | 1 | 1 |
| ADPRS | 10 | 10 | 7,98985E-18 | 0,999999998 | 1 |
| DDB1 | 54 | 53 | 0 | 1 | 1 |
| ARHGEF17 | 12 | 12 | 1,28864E-15 | 0,999999895 | 1 |

| | | | | | |
|---|---|---|---|---|---|
| CRACDL | 15 | 15 | -0,051146592 | 0,402382327 | 1 |
| SNAP91 | 27 | 26 | -1,32023E-16 | 0,999999973 | 1 |
| SYT11 | 8 | 8 | 0 | 1 | 1 |
| MECR | 10 | 9 | 0,040010921 | 0,382953993 | 1 |
| GNAS | 7 | 7 | -5,50337E-17 | 0,999999973 | 1 |
| SH3GL1 | 17 | 17 | 0 | 1 | 1 |
| SH3GL2 | 24 | 24 | 5,01777E-18 | 0,999999999 | 1 |
| GRM5 | 26 | 26 | -0,006514081 | 0,677747441 | 1 |
| FDPS | 11 | 11 | 4,17942E-16 | 0,999999942 | 1 |
| NEFL | 34 | 32 | -7,15515E-19 | 1 | 1 |
| DAB2IP | 8 | 7 | 0 | 1 | 1 |
| STXBP3 | 9 | 8 | -1,54606E-16 | 0,99999997 | 1 |
| PAK2 | 21 | 19 | -1,43328E-16 | 0,999999948 | 1 |
| PAK3 | 11 | 11 | -0,021398874 | 0,46016664 | 1 |
| RSU1 | 8 | 8 | -7,78107E-16 | 0,999999915 | 1 |
| GPSM2 | 3 | 3 | 0 | 1 | 1 |
| SLC25A3 | 25 | 25 | 0,056394406 | 0,170958565 | 1 |
| OSTF1 | 4 | 4 | 0 | 1 | 1 |
| PHLPP1 | 2 | 2 | -6,71564E-17 | 0,999999995 | 1 |
| NCLN | 13 | 13 | -4,58208E-16 | 0,999999941 | 1 |
| CYFIP2 | 70 | 69 | 0 | 1 | 1 |
| VPS28 | 4 | 4 | -6,75113E-16 | 0,999999938 | 1 |
| SYT3 | 16 | 15 | 0,040817163 | 0,218434431 | 1 |
| OPCML | 15 | 15 | 0 | 1 | 1 |
| COPS3 | 16 | 16 | 0 | 1 | 1 |
| GNAO1 | 22 | 22 | 5,38819E-17 | 0,999999972 | 1 |
| GNAQ | 21 | 18 | 0,004441609 | 0,717351693 | 1 |
| TRIO | 31 | 30 | 0 | 1 | 1 |
| PGAM2;PGAM1 | 6 | 6 | -3,51042E-17 | 0,999999992 | 1 |
| AP1G1 | 19 | 19 | 0,029132749 | 0,159336838 | 1 |
| OSBPL1A | 13 | 13 | -0,006772294 | 0,667950521 | 1 |
| GABBR2 | 21 | 21 | 2,01238E-17 | 0,999999994 | 1 |
| PSMC3 | 28 | 27 | -1,36044E-16 | 0,999999954 | 1 |

| | | | | | |
|---|---|---|---|---|---|
| PFKP | 34 | 33 | 0 | 1 | 1 |
| SORD | 13 | 12 | 0 | 1 | 1 |
| PSPH | 9 | 8 | 1,09769E-18 | 1 | 1 |
| GNAI2 | 14 | 14 | 6,51968E-18 | 0,999999996 | 1 |
| NAA10 | 3 | 3 | 0 | 1 | 1 |
| NAA12 | 1 | 1 | 0 | 1 | 1 |
| YARS | 36 | 36 | -6,14601E-17 | 0,999999987 | 1 |
| ELOC | 9 | 9 | -7,5619E-18 | 0,999999998 | 1 |
| ACY1 | 12 | 12 | -5,84737E-18 | 0,999999998 | 1 |
| VAPB | 13 | 13 | -0,02447237 | 0,300555067 | 1 |
| LARS2 | 10 | 9 | 0 | 1 | 1 |
| TMEM135 | 1 | 1 | 0 | 1 | 1 |
| CCT3 | 40 | 40 | -0,003885661 | 0,717545552 | 1 |
| HSPA2;HSPA8 | 9 | 9 | 8,52965E-17 | 0,999999968 | 1 |
| S100B | 4 | 4 | -4,20991E-17 | 0,999999995 | 1 |
| AK2 | 9 | 8 | 2,25237E-15 | 0,999999854 | 1 |
| EEF1E1 | 8 | 7 | 0 | 1 | 1 |
| LRRFIP1;LRRFIP2 | 2 | 1 | 0 | 1 | 1 |
| HDHD2 | 13 | 12 | 2,57275E-16 | 0,999999965 | 1 |
| NEBL | 16 | 15 | -0,009606777 | 0,668784705 | 1 |
| NDUFB9 | 12 | 11 | 1,51674E-15 | 0,999999893 | 1 |
| ANXA7 | 24 | 24 | -2,92858E-16 | 0,999999946 | 1 |
| IAP | 2 | 2 | -0,091762601 | 0,31677302 | 1 |
| GABRA4 | 5 | 5 | -8,67323E-17 | 0,999999983 | 1 |
| PLPPR4 | 24 | 22 | 0 | 1 | 1 |
| MTX2 | 7 | 7 | 0,001567648 | 0,856509823 | 1 |
| PIK3R4 | 19 | 19 | 0,019490498 | 0,448626712 | 1 |
| NRXN3 | 26 | 25 | 0,033142505 | 0,247512123 | 1 |
| PCDH1 | 23 | 23 | -1,6235E-17 | 0,999999994 | 1 |
| DUSP3 | 8 | 8 | -2,00146E-17 | 0,999999992 | 1 |
| PITPNM3 | 3 | 3 | -0,0372409 | 0,425599163 | 1 |
| PRDX3 | 13 | 13 | 0,028778739 | 0,394161742 | 1 |
| ALPL | 1 | 1 | -7,29941E-19 | 0,999999999 | 1 |

| | | | | | |
|---|---|---|---|---|---|
| CDC16 | 4 | 4 | 0 | 1 | 1 |
| CLIP1 | 35 | 34 | 0 | 1 | 1 |
| TOMM40L | 6 | 5 | 0,032877235 | 0,342370064 | 1 |
| TANC1 | 17 | 17 | -2,23717E-17 | 0,999999992 | 1 |
| AFG1L | 8 | 6 | -1,14179E-19 | 1 | 1 |
| RPL26 | 10 | 9 | -5,54601E-17 | 0,999999983 | 1 |
| SETD7 | 11 | 11 | 0 | 1 | 1 |
| LACTB2 | 4 | 4 | 0 | 1 | 1 |
| POMGNT2 | 2 | 1 | 0 | 1 | 1 |
| PAM | 10 | 9 | -9,50789E-17 | 0,999999969 | 1 |
| RAB8A | 7 | 7 | -3,17876E-16 | 0,999999952 | 1 |
| MBLAC2 | 11 | 10 | -0,016663121 | 0,5645248 | 1 |
| LAP3 | 23 | 23 | -1,46185E-17 | 0,999999995 | 1 |
| PDHX | 20 | 20 | 0,039886512 | 0,218003757 | 1 |
| LNPEP | 16 | 16 | 0 | 1 | 1 |
| EPS8 | 11 | 11 | 1,71166E-17 | 0,999999997 | 1 |
| ARMC8 | 7 | 7 | -1,58026E-16 | 0,99999996 | 1 |
| ATG4B | 7 | 7 | 5,15501E-18 | 0,999999999 | 1 |
| SLC18A3 | 2 | 2 | -6,70886E-17 | 0,999999989 | 1 |
| CKAP5 | 51 | 51 | 2,20472E-17 | 0,999999989 | 1 |
| OSBPL8 | 11 | 11 | 2,03313E-15 | 0,999999868 | 1 |
| SEC22B | 16 | 16 | -0,016366581 | 0,418821491 | 1 |
| CR2 | 1 | 1 | 0 | 1 | 1 |
| NCBP1 | 3 | 3 | -0,062259277 | 0,201221085 | 1 |
| MT-ND3 | 1 | 1 | 0,100063281 | 0,247233741 | 1 |
| MTND5 | 6 | 6 | -1,14677E-17 | 0,999999992 | 1 |
| CALB2 | 22 | 22 | -4,33137E-16 | 0,999999949 | 1 |
| HECW2 | 10 | 10 | 0 | 1 | 1 |
| MAP1S | 16 | 16 | -4,61652E-18 | 0,999999995 | 1 |
| OGA | 17 | 17 | -3,7097E-17 | 0,999999982 | 1 |
| GAD1;GAD2 | 1 | 1 | 0 | 1 | 1 |
| HEXB | 19 | 18 | -0,010312439 | 0,616026093 | 1 |
| GPD1 | 17 | 17 | 0,016369453 | 0,371411834 | 1 |

| | | | | | |
|---|---|---|---|---|---|
| ASTN2 | 7 | 6 | -1,01331E-06 | 0,997167387 | 1 |
| ATP5PD | 18 | 17 | 0,00630451 | 0,730124409 | 1 |
| TBL2 | 3 | 3 | -3,19756E-18 | 0,999999996 | 1 |
| ARHGEF9 | 6 | 6 | 0 | 1 | 1 |
| PLCB4 | 3 | 3 | -2,32468E-18 | 0,999999997 | 1 |
| ACO1 | 26 | 26 | 2,90543E-18 | 0,999999996 | 1 |
| RIMS1 | 9 | 9 | 9,74644E-11 | 0,999969356 | 1 |
| CRYM | 21 | 21 | 1,00909E-17 | 0,999999989 | 1 |
| PGBD5 | 14 | 13 | -2,76652E-17 | 0,999999986 | 1 |
| HCN1 | 13 | 13 | 7,11659E-17 | 0,999999979 | 1 |
| DNM1 | 55 | 55 | 0,012821324 | 0,467229132 | 1 |
| MAN2C1 | 17 | 17 | 8,73966E-16 | 0,999999937 | 1 |
| PTRH2 | 5 | 5 | 6,35319E-17 | 0,999999986 | 1 |
| SLC35B2 | 3 | 3 | 0 | 1 | 1 |
| EZR;MSN | 9 | 9 | 8,07168E-17 | 0,999999993 | 1 |
| GTF2B | 2 | 2 | -1,48857E-18 | 0,999999998 | 1 |
| FADS1 | 4 | 4 | 0 | 1 | 1 |
| ECH1 | 7 | 6 | 0 | 1 | 1 |
| DNAJC5 | 8 | 8 | 0 | 1 | 1 |
| MAPK1 | 35 | 35 | -4,69418E-17 | 0,999999989 | 1 |
| ACTR1B | 10 | 10 | 1,03071E-16 | 0,99999997 | 1 |
| sp\|Q9CYS6\|CB072_ | 2 | 2 | -3,56467E-18 | 0,999999998 | 1 |
| SIRPA | 19 | 18 | 0 | 1 | 1 |
| MAPK8 | 4 | 4 | 0,000681365 | 0,913696932 | 1 |
| PSMB2 | 11 | 11 | 7,60758E-17 | 0,99999998 | 1 |
| SLC4A10 | 16 | 15 | 0 | 1 | 1 |
| TENM4 | 32 | 30 | 4,30235E-18 | 0,999999994 | 1 |
| ADGRG1 | 5 | 5 | 1,84864E-19 | 1 | 1 |
| ALCAM | 24 | 24 | 0 | 1 | 1 |
| FMN2 | 19 | 18 | -1,39896E-16 | 0,999999962 | 1 |
| FXYD6 | 3 | 3 | 1,10032E-16 | 0,999999992 | 1 |
| PAIP1 | 1 | 1 | -0,010989089 | 0,748385593 | 1 |
| JPH4 | 3 | 3 | -1,12032E-17 | 0,999999996 | 1 |

| | | | | | |
|---|---|---|---|---|---|
| LRRC47 | 17 | 17 | -0,041035642 | 0,112857752 | 1 |
| DUSP28 | 1 | 1 | -0,127100066 | 0,115260988 | 1 |
| XPNPEP3 | 5 | 4 | 0 | 1 | 1 |
| ALG2 | 13 | 12 | 0 | 1 | 1 |
| IDH3A | 26 | 26 | 0 | 1 | 1 |
| RPL4 | 18 | 17 | 0,10387149 | 0,185492516 | 1 |
| 2900026A02RIK | 11 | 11 | 7,08787E-17 | 0,999999986 | 1 |
| FN1 | 8 | 7 | 1,95682E-16 | 0,999999965 | 1 |
| CAMSAP3 | 10 | 10 | -4,22224E-17 | 0,999999984 | 1 |
| BTF3L4 | 4 | 4 | -0,081060825 | 0,195773327 | 1 |
| BTF3 | 3 | 3 | -5,55698E-15 | 0,999999873 | 1 |
| PSME1 | 5 | 5 | -4,80205E-17 | 0,999999991 | 1 |
| RFTN2 | 9 | 9 | 3,82849E-18 | 0,999999998 | 1 |
| CSPG4 | 17 | 17 | -3,33957E-17 | 0,999999989 | 1 |
| LYSMD2 | 1 | 1 | 0 | 1 | 1 |
| IARS1 | 36 | 36 | 0 | 1 | 1 |
| CEND1 | 7 | 7 | 6,66223E-14 | 0,999999211 | 1 |
| HMGCS2 | 1 | 1 | 0,308262252 | 0,124727415 | 1 |
| PYGM | 38 | 38 | -2,54571E-17 | 0,999999989 | 1 |
| PLA2G15 | 4 | 4 | 0,056644773 | 0,208095795 | 1 |
| PTPN5 | 4 | 4 | -0,001129743 | 0,915659833 | 1 |
| ENHO | 1 | 1 | 0 | 1 | 1 |
| GOT1 | 36 | 36 | 0 | 1 | 1 |
| CPNE1 | 11 | 11 | 4,8573E-17 | 0,999999986 | 1 |
| ADPGK | 4 | 4 | 0,104718814 | 0,181308979 | 1 |
| CSNK1G1;CSNK1G2 | 5 | 5 | 0 | 1 | 1 |
| MRPL13 | 4 | 3 | 0 | 1 | 1 |
| IGHG2B | 4 | 3 | 1,99565E-18 | 0,999999998 | 1 |
| sp\|P01864\|GCAB_ | 7 | 4 | -8,9373E-19 | 0,999999999 | 1 |
| MICALL1 | 1 | 1 | 0 | 1 | 1 |
| COX7B | 1 | 1 | 0 | 1 | 1 |
| DDHD1 | 6 | 6 | 0 | 1 | 1 |
| BC034090 | 1 | 1 | 0 | 1 | 1 |

| | | | | | |
|---|---|---|---|---|---|
| ATP5MG | 6 | 6 | 0,102985365 | 0,114552892 | 1 |
| ASNS | 19 | 19 | -7,37522E-17 | 0,999999971 | 1 |
| CORO7 | 19 | 19 | 0 | 1 | 1 |
| CRYAB | 12 | 12 | 3,48471E-16 | 0,999999969 | 1 |
| TNC | 15 | 15 | 0 | 1 | 1 |
| FAM131B | 11 | 11 | 0 | 1 | 1 |
| SLC43A2 | 2 | 2 | 0 | 1 | 1 |
| APBA1 | 6 | 6 | -0,075681311 | 0,230441635 | 1 |
| DDN | 2 | 2 | 0 | 1 | 1 |
| ANK2 | 13 | 13 | 2,40301E-16 | 0,99999996 | 1 |
| AGFG1 | 9 | 9 | -1,28825E-15 | 0,999999894 | 1 |
| CBL | 7 | 7 | -0,03453347 | 0,390943976 | 1 |
| LZTS1 | 4 | 4 | -0,044311726 | 0,597181517 | 1 |
| FLOT1 | 29 | 27 | 0,004180575 | 0,671908388 | 1 |
| NRSN1 | 1 | 1 | 0 | 1 | 1 |
| CNOT1 | 6 | 6 | -0,026038286 | 0,415069723 | 1 |
| TBCB | 12 | 12 | 0,007326035 | 0,606855685 | 1 |
| RPL29 | 2 | 2 | 4,57131E-18 | 0,999999997 | 1 |
| NPTN | 6 | 6 | -1,52971E-18 | 0,999999997 | 1 |
| LONP1 | 38 | 36 | 0,020473824 | 0,379624879 | 1 |
| RAB7A | 21 | 21 | -0,033097108 | 0,306421779 | 1 |
| RPS5 | 9 | 8 | -0,074117154 | 0,137545257 | 1 |
| SHFL | 2 | 2 | 0 | 1 | 1 |
| NUDCD2 | 5 | 5 | -5,19918E-17 | 0,999999987 | 1 |
| LZTFL1 | 7 | 7 | -0,059359455 | 0,181322334 | 1 |
| EPB41L3 | 7 | 7 | 6,69114E-18 | 0,999999997 | 1 |
| SMG1 | 3 | 3 | 0 | 1 | 1 |
| ACADL | 24 | 22 | 0 | 1 | 1 |
| SCAMP5 | 2 | 2 | 0 | 1 | 1 |
| COQ3 | 5 | 5 | 0,029052267 | 0,419583715 | 1 |
| CACNA2D1 | 41 | 41 | 0 | 1 | 1 |
| DDHD2 | 14 | 14 | 1,89683E-17 | 0,999999987 | 1 |
| CZIB | 5 | 5 | -5,16463E-17 | 0,999999977 | 1 |

| | | | | | |
|---|---|---|---|---|---|
| XPO4 | 2 | 2 | -1,42646E-16 | 0,999999974 | 1 |
| DIABLO | 2 | 2 | 3,96477E-19 | 1 | 1 |
| BRINP2 | 5 | 5 | 2,04472E-16 | 0,999999976 | 1 |
| FYN | 11 | 11 | 0,047832071 | 0,246801628 | 1 |
| CASK | 11 | 11 | 0 | 1 | 1 |
| EML2 | 16 | 15 | -1,98642E-17 | 0,999999991 | 1 |
| RAB15 | 9 | 9 | 9,83523E-15 | 0,999999725 | 1 |
| ZFAND2B | 1 | 1 | 0 | 1 | 1 |
| PTGDS | 4 | 4 | -4,97272E-17 | 0,999999987 | 1 |
| DNAJA3 | 13 | 12 | 4,29976E-17 | 0,999999986 | 1 |
| TIMM50 | 9 | 9 | 0,043021256 | 0,344847003 | 1 |
| HSPA1B;HSPA1A | 15 | 15 | 2,54707E-17 | 0,999999988 | 1 |
| RTN4 | 39 | 38 | -0,017457486 | 0,551905078 | 1 |
| DBT | 20 | 20 | 0,013765915 | 0,547976476 | 1 |
| TANC2 | 31 | 31 | 0,032435328 | 0,166505875 | 1 |
| KTN1 | 23 | 22 | 0 | 1 | 1 |
| CDKL5 | 14 | 14 | -1,67399E-16 | 0,999999968 | 1 |
| FAM126A | 1 | 1 | 0,094944449 | 0,360367882 | 1 |
| RLBP1 | 5 | 5 | -5,36607E-18 | 0,999999995 | 1 |
| FGD4 | 3 | 3 | 0 | 1 | 1 |
| RAB24 | 6 | 6 | -8,74726E-16 | 0,999999914 | 1 |
| TFG | 13 | 13 | -2,01649E-16 | 0,999999951 | 1 |
| TTC33 | 1 | 1 | 0 | 1 | 1 |
| PPP1R12A | 17 | 16 | 0 | 1 | 1 |
| FN3K | 5 | 5 | 0 | 1 | 1 |
| KCNJ4 | 3 | 3 | 0 | 1 | 1 |
| PNKD | 4 | 4 | 1,92107E-16 | 0,999999974 | 1 |
| DRG2 | 14 | 14 | -0,011835561 | 0,639616151 | 1 |
| L1CAM | 28 | 26 | 4,63571E-16 | 0,999999928 | 1 |
| STIM2 | 11 | 11 | 0,020341268 | 0,491190271 | 1 |
| SIPA1L1 | 50 | 47 | 0 | 1 | 1 |
| STX19 | 1 | 1 | 0 | 1 | 1 |
| PPP4R4 | 7 | 7 | 2,41552E-17 | 0,999999984 | 1 |

| | | | | | |
|---|---|---|---|---|---|
| PTK2 | 9 | 8 | -3,31417E-20 | 1 | 1 |
| SAE1 | 20 | 20 | -0,091247604 | 0,120044814 | 1 |
| DCLK1 | 31 | 29 | -5,27098E-18 | 0,999999996 | 1 |
| ADI1 | 3 | 3 | -2,24066E-18 | 0,999999998 | 1 |
| UBXN6 | 19 | 19 | 0 | 1 | 1 |
| DGKI | 7 | 7 | -6,02139E-06 | 0,989758511 | 1 |
| ATP5PB | 28 | 26 | 7,44544E-17 | 0,999999976 | 1 |
| SCAMP3 | 11 | 11 | 0,061237741 | 0,147556406 | 1 |
| SCAMP1 | 13 | 12 | 0 | 1 | 1 |
| TACC2 | 3 | 3 | -3,96453E-20 | 1 | 1 |
| LZTS3 | 4 | 4 | -0,08794041 | 0,26797897 | 1 |
| AATK | 4 | 4 | -6,41072E-17 | 0,999999994 | 1 |
| FGA | 8 | 7 | 4,28924E-18 | 0,999999998 | 1 |
| CC2D1A | 8 | 8 | 5,09342E-18 | 0,999999999 | 1 |
| CSNK1G3 | 3 | 3 | 1,74904E-17 | 0,999999993 | 1 |
| PLCG1 | 10 | 10 | 4,09796E-17 | 0,99999998 | 1 |
| PSMD2 | 40 | 39 | -0,036802843 | 0,130077551 | 1 |
| GPC5 | 4 | 4 | 5,42159E-17 | 0,999999981 | 1 |
| ATP5F1D | 4 | 4 | 0,074590513 | 0,18661689 | 1 |
| AARS1 | 44 | 44 | -0,005625072 | 0,672866429 | 1 |
| SCAMP4 | 1 | 1 | 0 | 1 | 1 |
| C2CD4C | 6 | 6 | 0 | 1 | 1 |
| SEPTIN10 | 1 | 1 | 0 | 1 | 1 |
| HINT1 | 9 | 9 | 1,14575E-16 | 0,999999977 | 1 |
| LYRM4 | 5 | 5 | 0,056812276 | 0,263081687 | 1 |
| NEFM | 33 | 33 | 0 | 1 | 1 |
| PIGW | 1 | 1 | 0,026978545 | 0,584460253 | 1 |
| SORCS2 | 12 | 12 | 0,047967409 | 0,182277245 | 1 |
| IKBKG | 2 | 2 | 9,99927E-16 | 0,999999938 | 1 |
| CDH11 | 10 | 10 | 0 | 1 | 1 |
| GPLD1 | 2 | 2 | -0,058108702 | 0,294058655 | 1 |
| ACTR1A | 10 | 10 | 5,8572E-18 | 0,999999994 | 1 |
| SETD1A | 1 | 1 | 0 | 1 | 1 |

| | | | | | |
|---|---|---|---|---|---|
| OXR1 | 25 | 25 | -2,2316E-16 | 0,999999948 | 1 |
| PRICKLE2 | 16 | 16 | 0,012783684 | 0,556863036 | 1 |
| NT5M | 7 | 5 | 3,0895E-16 | 0,999999954 | 1 |
| FRRS1L | 10 | 10 | 1,75051E-16 | 0,999999981 | 1 |
| NDUFB5 | 10 | 10 | 5,25376E-17 | 0,999999979 | 1 |
| FKBP3 | 10 | 10 | -0,047405401 | 0,115971801 | 1 |
| RGS7 | 15 | 14 | 0 | 1 | 1 |
| CHKB | 11 | 10 | 3,86115E-18 | 0,999999997 | 1 |
| RBCK1 | 1 | 1 | 0 | 1 | 1 |
| PYGB | 60 | 59 | 0 | 1 | 1 |
| SYT13 | 4 | 4 | 3,39431E-17 | 0,999999991 | 1 |
| EFNB3 | 5 | 5 | -7,58797E-18 | 0,999999996 | 1 |
| TGM2 | 10 | 10 | 0,050497839 | 0,481753173 | 1 |
| CCDC47 | 14 | 14 | -0,006964483 | 0,694078657 | 1 |
| GABARAPL1 | 3 | 3 | 0 | 1 | 1 |
| TPD52 | 9 | 9 | -1,05367E-16 | 0,999999973 | 1 |
| APC | 17 | 17 | 0 | 1 | 1 |
| RASAL2 | 10 | 8 | 0,004126984 | 0,770908222 | 1 |
| C4B | 3 | 3 | 0 | 1 | 1 |
| TMEM47 | 3 | 2 | -3,58209E-19 | 0,999999999 | 1 |
| C2CD5 | 15 | 15 | 0 | 1 | 1 |
| ZNF516 | 2 | 2 | -1,08178E-12 | 0,999998592 | 1 |
| PHACTR2 | 5 | 5 | 1,38922E-16 | 0,999999964 | 1 |
| MRPS30 | 9 | 8 | 0 | 1 | 1 |
| CRYBB1 | 3 | 3 | -2,40193E-18 | 1 | 1 |
| AMER2 | 10 | 10 | -2,65305E-16 | 0,999999949 | 1 |
| ATP6V1C1 | 43 | 43 | 9,23236E-16 | 0,999999917 | 1 |
| CLUH | 18 | 18 | -4,92714E-17 | 0,999999981 | 1 |
| VCAN | 1 | 1 | -3,24474E-18 | 0,999999997 | 1 |
| sp\|Q6PIU9\|YJ005_ | 4 | 4 | -3,35369E-18 | 0,999999997 | 1 |
| TRAPPC8 | 12 | 12 | 0 | 1 | 1 |
| EPHX2 | 12 | 12 | 0 | 1 | 1 |
| RP2 | 2 | 1 | 0 | 1 | 1 |

| | | | | | |
|---|---|---|---|---|---|
| ANKMY2 | 4 | 4 | -7,58834E-16 | 0,999999915 | 1 |
| TRPM3 | 3 | 3 | 3,1308E-18 | 0,999999997 | 1 |
| RCN1 | 2 | 2 | -1,17574E-16 | 0,999999971 | 1 |
| RAPGEF2 | 60 | 59 | 0,011130161 | 0,600784239 | 1 |
| REPS1 | 13 | 13 | -2,68679E-06 | 0,995394248 | 1 |
| TOMM40 | 10 | 10 | -2,3886E-21 | 1 | 1 |
| GDA | 36 | 35 | 1,0218E-16 | 0,999999975 | 1 |
| SLC20A2 | 5 | 5 | 1,57268E-16 | 0,999999982 | 1 |
| GALM | 4 | 4 | 3,56126E-17 | 0,99999999 | 1 |
| COPS6 | 15 | 15 | -2,74473E-17 | 0,999999985 | 1 |
| CSF1R | 4 | 4 | 0 | 1 | 1 |
| SLC2A13 | 8 | 8 | -0,019211214 | 0,495050211 | 1 |
| SRGAP3;SRGAP1 | 4 | 4 | 0 | 1 | 1 |
| CLTB | 2 | 2 | 0 | 1 | 1 |
| MFSD6 | 4 | 4 | 0,024248055 | 0,572341841 | 1 |
| PPP2R5C | 2 | 2 | 0 | 1 | 1 |
| PCDHGA1;PCDHGA | 1 | 1 | 0 | 1 | 1 |
| VPS18 | 14 | 14 | -0,02207952 | 0,443471893 | 1 |
| NOL3 | 6 | 5 | 0,052256161 | 0,29837129 | 1 |
| SEMA6B | 1 | 1 | 0,147500651 | 0,244178018 | 1 |
| EIF4G3 | 1 | 1 | 0 | 1 | 1 |
| OSBPL6 | 13 | 13 | 0,031500175 | 0,311474919 | 1 |
| MRAS | 12 | 12 | 0 | 1 | 1 |
| ECSIT | 7 | 7 | 1,15573E-17 | 0,999999998 | 1 |
| TNIK | 17 | 17 | 5,48559E-14 | 0,999998947 | 1 |
| TXNRD2 | 6 | 5 | 0 | 1 | 1 |
| SACM1L | 28 | 27 | -1,94095E-16 | 0,999999942 | 1 |
| TNPO3 | 10 | 10 | 2,99746E-18 | 0,999999999 | 1 |
| SYN1 | 40 | 40 | 8,5711E-16 | 0,99999991 | 1 |
| NELFB | 2 | 2 | -0,0683703 | 0,487456162 | 1 |
| SLC25A13 | 8 | 7 | 1,43493E-16 | 0,999999987 | 1 |
| CALD1 | 3 | 3 | 0 | 1 | 1 |
| GCLC | 26 | 26 | 0 | 1 | 1 |

| | | | | | |
|---|---|---|---|---|---|
| ENDOG | 7 | 6 | 0 | 1 | 1 |
| NPW | 1 | 1 | -0,159630991 | 0,232303049 | 1 |
| MDN1 | 1 | 1 | 0 | 1 | 1 |
| RPL8 | 9 | 9 | -4,79978E-17 | 0,999999986 | 1 |
| TRIM9 | 3 | 3 | -0,088640275 | 0,183765101 | 1 |
| CD244 | 1 | 1 | 0,189707755 | 0,244289292 | 1 |
| PTCD3 | 13 | 10 | -1,49196E-16 | 0,999999974 | 1 |
| NECAP1 | 10 | 9 | 5,32292E-17 | 0,999999992 | 1 |
| PPM1K | 1 | 1 | -0,012743797 | 0,768453973 | 1 |
| RBBP9 | 10 | 10 | -6,03926E-17 | 0,999999991 | 1 |
| CHL1 | 29 | 29 | -1,04515E-18 | 0,999999999 | 1 |
| SFXN1 | 14 | 13 | 0,053543569 | 0,117707282 | 1 |
| STRADA | 4 | 4 | -0,037283084 | 0,477121787 | 1 |
| C2CD2L | 20 | 19 | 0 | 1 | 1 |
| ARPC2 | 29 | 29 | 0 | 1 | 1 |
| PPP1R21 | 24 | 24 | -0,001502111 | 0,833613958 | 1 |
| CLYBL | 13 | 13 | 0,048102862 | 0,117002249 | 1 |
| SGIP1 | 14 | 14 | -2,74583E-17 | 0,999999994 | 1 |
| DNAH6 | 2 | 2 | 0,05353106 | 0,420137249 | 1 |
| RIC8A | 13 | 13 | 1,53956E-17 | 0,99999999 | 1 |
| SEPTIN2 | 13 | 13 | -5,02494E-16 | 0,999999946 | 1 |
| TRIM2 | 25 | 25 | -2,12903E-17 | 0,999999979 | 1 |
| TAPT1 | 5 | 5 | 0,018293476 | 0,506095372 | 1 |
| MAGI3 | 4 | 4 | 6,35124E-14 | 0,999999356 | 1 |
| RTKN | 2 | 2 | 0 | 1 | 1 |
| ADGRE5 | 1 | 1 | 0 | 1 | 1 |
| KRT16 | 28 | 9 | 8,11804E-16 | 0,999999966 | 1 |
| CPM | 8 | 7 | 5,82433E-18 | 0,999999998 | 1 |
| TSG101 | 7 | 7 | -5,44208E-17 | 0,999999978 | 1 |
| DHX36 | 5 | 5 | 0 | 1 | 1 |
| NEDD4L | 18 | 18 | -6,81104E-18 | 0,999999997 | 1 |
| EEF1B | 10 | 10 | 0 | 1 | 1 |
| UBXN7 | 8 | 8 | -0,013580166 | 0,612534424 | 1 |

| | | | | | |
|---|---|---|---|---|---|
| RNF34 | 4 | 4 | 0 | 1 | 1 |
| IP6K1 | 6 | 6 | -4,47294E-17 | 0,999999981 | 1 |
| ABHD10 | 9 | 6 | 0 | 1 | 1 |
| CHCHD6 | 16 | 15 | 4,24982E-17 | 0,999999981 | 1 |
| PSAT1 | 21 | 21 | 0 | 1 | 1 |
| LRRFIP1 | 5 | 5 | 0 | 1 | 1 |
| NAPG | 24 | 23 | 0 | 1 | 1 |
| KCNMA1 | 26 | 26 | 0 | 1 | 1 |
| COPA | 24 | 24 | 0 | 1 | 1 |
| UBAC2 | 3 | 3 | 3,1576E-16 | 0,999999989 | 1 |
| DPP7 | 4 | 4 | 0 | 1 | 1 |
| EXOC2 | 22 | 22 | 2,49719E-17 | 0,999999995 | 1 |
| ACADVL | 21 | 19 | -9,90579E-18 | 0,999999993 | 1 |
| UNC5C | 3 | 3 | -0,065211801 | 0,365702487 | 1 |
| BIN2 | 1 | 1 | 0 | 1 | 1 |
| ALDH3A2 | 14 | 14 | 0,029495611 | 0,331233231 | 1 |
| AKR1E2 | 7 | 7 | 2,09586E-17 | 0,999999985 | 1 |
| CNPY3 | 3 | 3 | 0 | 1 | 1 |
| CNTN2 | 19 | 19 | 0 | 1 | 1 |
| EPHX1 | 14 | 14 | 1,0295E-16 | 0,999999957 | 1 |
| RGS20 | 3 | 3 | 0 | 1 | 1 |
| PRRT2 | 4 | 4 | 1,13856E-16 | 0,999999986 | 1 |
| NAPEPLD | 5 | 5 | 0,040760424 | 0,369886496 | 1 |
| OGFRL1 | 8 | 8 | -0,060042199 | 0,185789645 | 1 |
| TBC1D10B | 11 | 11 | 4,4995E-17 | 0,999999979 | 1 |
| PPP6R1 | 7 | 7 | 0 | 1 | 1 |
| PDZD11 | 2 | 2 | 2,11009E-18 | 0,999999998 | 1 |
| RAP1GAP2 | 13 | 13 | 0 | 1 | 1 |
| ECI2 | 7 | 7 | -0,070394927 | 0,184036611 | 1 |
| CNOT1 | 20 | 20 | -0,015584739 | 0,552928347 | 1 |
| CAT | 19 | 18 | 3,10322E-17 | 0,999999985 | 1 |
| RIMS2 | 13 | 13 | 1,62944E-15 | 0,999999893 | 1 |
| ADRA2A | 2 | 2 | 0 | 1 | 1 |

| | | | | | |
|---|---|---|---|---|---|
| UBA3 | 16 | 16 | -2,74841E-17 | 0,999999984 | 1 |
| SPECC1 | 4 | 4 | -3,20044E-17 | 0,999999992 | 1 |
| ARPC1B | 3 | 3 | -1,28884E-18 | 0,999999999 | 1 |
| HMGCS1 | 14 | 14 | -0,008737998 | 0,680556417 | 1 |
| MTFR1L | 8 | 8 | -2,80782E-17 | 0,999999987 | 1 |
| RPN1 | 32 | 32 | -3,20103E-16 | 0,999999942 | 1 |
| NPDC1 | 3 | 3 | 1,00578E-15 | 0,999999924 | 1 |
| PKM | 8 | 8 | 0,052148012 | 0,373094415 | 1 |
| OSBPL2 | 10 | 10 | -0,014419077 | 0,490478854 | 1 |
| RGS10 | 4 | 4 | -0,044834857 | 0,403958275 | 1 |
| RABGAP1L | 7 | 7 | -1,2634E-15 | 0,999999883 | 1 |
| NSFL1C | 24 | 24 | -0,026456072 | 0,149167319 | 1 |
| PRKCI | 3 | 3 | 0,063201713 | 0,167393393 | 1 |
| CRIP2 | 4 | 4 | -0,020604047 | 0,579493622 | 1 |
| RAB11FIP2 | 12 | 11 | -0,031056463 | 0,337731504 | 1 |
| GCLM | 7 | 7 | -0,035632421 | 0,235704177 | 1 |
| CHP1 | 11 | 11 | -1,50752E-13 | 0,999998521 | 1 |
| DNAJC10 | 5 | 5 | -0,191168119 | 0,131183867 | 1 |
| LRBA | 11 | 11 | -0,004028487 | 0,749139947 | 1 |
| RNMT | 19 | 19 | -8,1445E-18 | 0,999999997 | 1 |
| CEP170 | 27 | 27 | -3,2282E-17 | 0,999999986 | 1 |
| AKAP10 | 9 | 9 | -0,052048529 | 0,17857096 | 1 |
| DCTN2 | 25 | 25 | -8,07899E-16 | 0,999999906 | 1 |
| PALLD | 1 | 1 | 0 | 1 | 1 |
| GPSM1 | 10 | 10 | -0,065597301 | 0,137724289 | 1 |
| SIPA1L3 | 13 | 12 | 1,33313E-16 | 0,999999974 | 1 |
| RTN4R | 3 | 3 | 0 | 1 | 1 |
| EVL | 8 | 8 | -0,048093994 | 0,201433704 | 1 |
| GLRX3 | 18 | 18 | -1,8081E-17 | 0,999999994 | 1 |
| CADPS2 | 20 | 20 | 0 | 1 | 1 |
| MIF | 7 | 7 | 3,13779E-16 | 0,999999953 | 1 |
| AJAP1 | 3 | 3 | 0 | 1 | 1 |
| DTD1 | 7 | 7 | -0,063947063 | 0,197674376 | 1 |

| | | | | | |
|---|---|---|---|---|---|
| LYPLAL1 | 5 | 5 | 0,063373925 | 0,20289374 | 1 |
| VGF | 10 | 10 | -8,73122E-16 | 0,999999921 | 1 |
| NIT1 | 7 | 7 | -5,88205E-18 | 0,999999997 | 1 |
| RBM14 | 1 | 1 | 0 | 1 | 1 |
| KALRN | 6 | 6 | 0 | 1 | 1 |
| GRB2 | 15 | 15 | 2,6997E-16 | 0,999999944 | 1 |
| ECI1 | 15 | 14 | -0,026757155 | 0,268982765 | 1 |
| RPA1 | 4 | 4 | -0,05455662 | 0,319234216 | 1 |
| PSMB8 | 4 | 4 | 1,93786E-16 | 0,999999964 | 1 |
| EEF1D | 9 | 9 | -0,03085142 | 0,486016688 | 1 |
| BROX | 5 | 5 | 1,60188E-16 | 0,999999973 | 1 |
| COPS7B | 5 | 5 | 0 | 1 | 1 |
| RASA1 | 14 | 13 | 0 | 1 | 1 |
| STK3 | 2 | 2 | 0,110852824 | 0,192508178 | 1 |
| HSPA12A | 39 | 38 | -0,021303344 | 0,257230444 | 1 |
| TMEM119 | 2 | 2 | 7,16237E-18 | 0,999999996 | 1 |
| SH3GLB2 | 20 | 20 | -1,49117E-16 | 0,999999982 | 1 |
| PSMA4 | 11 | 11 | -6,674E-17 | 0,999999984 | 1 |
| NBEA;LRBA | 5 | 5 | 0 | 1 | 1 |
| IDH2 | 27 | 26 | 0 | 1 | 1 |
| CAND2 | 2 | 2 | 5,50221E-17 | 0,999999985 | 1 |
| TSC22D1 | 1 | 1 | 0 | 1 | 1 |
| SALL1 | 1 | 1 | 0 | 1 | 1 |
| TRNT1 | 14 | 14 | -1,9211E-16 | 0,999999961 | 1 |
| TSC22D2 | 4 | 4 | -2,81465E-19 | 0,999999999 | 1 |
| CACNG2 | 2 | 2 | 6,08999E-17 | 0,999999994 | 1 |
| GDPGP1 | 8 | 8 | 9,42689E-18 | 0,999999996 | 1 |
| AP3M2 | 20 | 19 | 0,039898538 | 0,118946242 | 1 |
| OSBP | 15 | 15 | -1,59832E-17 | 0,999999989 | 1 |
| AMDHD2 | 4 | 4 | 0 | 1 | 1 |
| SERPINC1 | 3 | 3 | -1,1465E-16 | 0,999999981 | 1 |
| PSMB7 | 10 | 10 | 5,61335E-17 | 0,999999991 | 1 |
| HNRNPH1;HNRNPF | 1 | 1 | -0,040301863 | 0,527014697 | 1 |

| | | | | | |
|---|---|---|---|---|---|
| USP30 | 2 | 1 | 0 | 1 | 1 |
| PALM3 | 3 | 3 | -2,08883E-17 | 0,999999992 | 1 |
| SNX14 | 2 | 2 | 0,114932777 | 0,238812381 | 1 |
| SUPV3L1 | 17 | 15 | 0 | 1 | 1 |
| PGAM5 | 8 | 8 | 0 | 1 | 1 |
| KIF1A | 3 | 3 | -0,20374881 | 0,164052599 | 1 |
| CYP4X1 | 3 | 2 | -8,31234E-16 | 0,999999969 | 1 |
| RRAGB | 8 | 8 | 0 | 1 | 1 |
| ARHGDIB | 6 | 6 | 0,106437231 | 0,146574981 | 1 |
| NF1 | 19 | 19 | -1,14348E-18 | 0,999999998 | 1 |
| DHODH | 18 | 16 | 2,74928E-17 | 0,999999987 | 1 |
| ATE1 | 8 | 8 | 0 | 1 | 1 |
| COPG1 | 12 | 12 | -0,046331893 | 0,211687514 | 1 |
| COPG2 | 12 | 12 | 2,64764E-18 | 0,999999998 | 1 |
| CACNB1 | 2 | 2 | -8,50745E-19 | 0,999999999 | 1 |
| PRORSD1 | 1 | 1 | -0,069202613 | 0,456982822 | 1 |
| FN3KRP | 6 | 6 | 8,23878E-14 | 0,999999 | 1 |
| DNAJC6;GAK | 3 | 3 | 0 | 1 | 1 |
| CACNB1 | 9 | 9 | 0 | 1 | 1 |
| CACNB4 | 13 | 13 | -3,97997E-16 | 0,999999943 | 1 |
| AGPAT5 | 6 | 6 | 0,058886718 | 0,228068497 | 1 |
| P33MONOX | 3 | 3 | 4,11103E-18 | 0,999999998 | 1 |
| SPTBN5 | 1 | 1 | 0,132384803 | 0,564200337 | 1 |
| HDAC11 | 8 | 8 | 8,82525E-18 | 0,99999999 | 1 |
| NOS1 | 28 | 28 | 0 | 1 | 1 |
| CHMP2B | 3 | 3 | -1,18435E-20 | 1 | 1 |
| PTPRG | 12 | 11 | 0 | 1 | 1 |
| PDE4D | 10 | 10 | 2,80635E-17 | 0,999999983 | 1 |
| NDEL1 | 10 | 10 | -1,32096E-18 | 0,999999998 | 1 |
| GLO1 | 16 | 16 | -1,51287E-16 | 0,999999987 | 1 |
| CTH | 2 | 2 | 6,28189E-16 | 0,999999939 | 1 |
| ATP5C1 | 1 | 1 | 0 | 1 | 1 |
| PPP2R5A | 14 | 14 | -0,030937252 | 0,147308501 | 1 |

| | | | | | |
|---|---|---|---|---|---|
| TST | 14 | 12 | 0 | 1 | 1 |
| SNAP47 | 23 | 22 | 0,00959394 | 0,525870206 | 1 |
| RMC1 | 8 | 8 | 1,59902E-17 | 0,999999994 | 1 |
| NUDT2 | 7 | 7 | 0 | 1 | 1 |
| IGSF8 | 2 | 2 | 0,072795997 | 0,208589441 | 1 |
| ARFGEF3 | 23 | 23 | -0,014825287 | 0,431458353 | 1 |
| PDCD6IP | 37 | 37 | -0,048677737 | 0,11831888 | 1 |
| PRMT1 | 11 | 11 | 0 | 1 | 1 |
| ARHGEF28 | 1 | 1 | 0 | 1 | 1 |
| IFIT3 | 3 | 3 | 1,1362E-15 | 0,999999951 | 1 |
| GABRB1 | 5 | 5 | 4,28704E-17 | 0,999999981 | 1 |
| PAPSS1 | 12 | 12 | -0,022643769 | 0,454894269 | 1 |
| CLINT1 | 2 | 2 | -0,055884848 | 0,244204188 | 1 |
| ZDHHC14 | 1 | 1 | 0 | 1 | 1 |
| RPH3A | 29 | 28 | 0 | 1 | 1 |
| AGAP3 | 6 | 6 | 0 | 1 | 1 |
| NDUFA3 | 2 | 2 | 9,33145E-19 | 0,999999998 | 1 |
| EIF4A1 | 31 | 31 | -0,055431965 | 0,17705951 | 1 |
| CISD3 | 4 | 4 | 1,11932E-16 | 0,999999977 | 1 |
| DPP10 | 29 | 29 | -0,014753457 | 0,484284987 | 1 |
| ZDHHC17 | 1 | 1 | 0,118841092 | 0,220789195 | 1 |
| PPM1E | 21 | 19 | -1,01921E-15 | 0,999999884 | 1 |
| RAPGEF1 | 4 | 4 | 0 | 1 | 1 |
| MYL12B | 14 | 14 | -0,052850142 | 0,364410657 | 1 |
| FAM234B | 6 | 6 | 0,017812799 | 0,618896221 | 1 |
| BRI3BP | 3 | 3 | -0,070306275 | 0,2063653 | 1 |
| ATP8A2 | 8 | 7 | 0 | 1 | 1 |
| DIS3L2 | 7 | 7 | -3,02238E-17 | 0,999999986 | 1 |
| PICALM | 11 | 11 | -0,038284122 | 0,280635613 | 1 |
| RNF220 | 2 | 2 | -0,150586643 | 0,135865598 | 1 |
| RETREG2 | 1 | 1 | 0 | 1 | 1 |
| PEX14 | 6 | 6 | -4,62885E-18 | 0,999999995 | 1 |
| LIN7C;LIN7A | 1 | 1 | 0 | 1 | 1 |

| | | | | | |
|---|---|---|---|---|---|
| LIN7B | 2 | 1 | -0,054987361 | 0,40505216 | 1 |
| TAF9 | 1 | 1 | -1,26383E-18 | 0,999999998 | 1 |
| NAA35 | 3 | 3 | 1,87856E-18 | 0,999999999 | 1 |
| HSPA8 | 39 | 38 | 0,019188906 | 0,419664028 | 1 |
| GRM7 | 15 | 14 | 0,022679671 | 0,330270164 | 1 |
| TNPO1 | 5 | 5 | 0 | 1 | 1 |
| PCYOX1L | 9 | 9 | 6,64975E-17 | 0,999999976 | 1 |
| LINGO2 | 4 | 4 | 0 | 1 | 1 |
| PRPF19 | 5 | 5 | -0,018336142 | 0,453252989 | 1 |
| REEP5 | 4 | 4 | 0 | 1 | 1 |
| GLCCI1 | 3 | 3 | 0 | 1 | 1 |
| BLMH | 17 | 17 | -0,008714111 | 0,614315378 | 1 |
| NUCKS1 | 5 | 5 | -6,72815E-17 | 0,999999982 | 1 |
| YWHAG | 21 | 21 | 1,41078E-16 | 0,999999978 | 1 |
| PRMT7 | 3 | 3 | 0,053986175 | 0,201788019 | 1 |
| CUEDC2 | 3 | 3 | -0,089573165 | 0,177966375 | 1 |
| GABRA3 | 5 | 5 | 2,25453E-15 | 0,999999872 | 1 |
| ELFN1 | 7 | 7 | 4,63006E-13 | 0,999997943 | 1 |
| PPP1R3F | 2 | 2 | 7,45212E-10 | 0,999947492 | 1 |
| FSD1L | 4 | 4 | 0 | 1 | 1 |
| EIF2B2 | 5 | 5 | -0,010066018 | 0,689641266 | 1 |
| PGM2L1 | 40 | 40 | 2,98876E-17 | 0,999999992 | 1 |
| PSMC5 | 26 | 26 | -0,0257208 | 0,202172868 | 1 |
| ABHD14B | 6 | 6 | 0 | 1 | 1 |
| NAV1 | 13 | 12 | 0,028018503 | 0,355729673 | 1 |
| SMYD3 | 1 | 1 | 0 | 1 | 1 |
| CBR4 | 9 | 9 | 4,09883E-17 | 0,999999982 | 1 |
| CCDC127 | 7 | 6 | 0 | 1 | 1 |
| NEFH | 13 | 11 | 0 | 1 | 1 |
| PANK4 | 14 | 14 | 0,016727663 | 0,465826084 | 1 |
| TBPL1 | 2 | 2 | -0,104952812 | 0,272267036 | 1 |
| VPS4B | 8 | 7 | 0 | 1 | 1 |
| SLC7A5 | 6 | 6 | 0 | 1 | 1 |

| | | | | | |
|---|---|---|---|---|---|
| AHSA2 | 3 | 3 | -1,08539E-18 | 0,999999999 | 1 |
| CNTNAP1 | 45 | 44 | 0,038536 | 0,27608925 | 1 |
| ADCK1 | 8 | 8 | 0 | 1 | 1 |
| SLC39A8 | 1 | 1 | 0 | 1 | 1 |
| PSMD14 | 13 | 12 | -4,56776E-14 | 0,999999258 | 1 |
| VAT1 | 18 | 17 | -1,54342E-17 | 0,999999996 | 1 |
| TUBAL3 | 5 | 4 | -4,41521E-18 | 0,999999997 | 1 |
| TENM2 | 21 | 21 | 0,039687421 | 0,286927576 | 1 |
| SUOX | 8 | 8 | 0 | 1 | 1 |
| KIFAP3 | 5 | 5 | -0,027261123 | 0,33002995 | 1 |
| PACSIN2 | 1 | 1 | 0 | 1 | 1 |
| RAB33B | 7 | 6 | 0,034068582 | 0,284505416 | 1 |
| RPL6 | 9 | 9 | 0,018957925 | 0,595935011 | 1 |
| WRNIP1 | 4 | 4 | -1,31809E-18 | 0,999999998 | 1 |
| USP5 | 47 | 47 | -0,029215573 | 0,234237669 | 1 |
| CD47 | 4 | 4 | 0 | 1 | 1 |
| RAB5C | 11 | 11 | -0,01408915 | 0,487960109 | 1 |
| PMPCB | 13 | 12 | 0 | 1 | 1 |
| ABHD11 | 6 | 6 | -0,06250796 | 0,283081542 | 1 |
| TKT | 32 | 32 | 4,35931E-16 | 0,999999963 | 1 |
| RECQL5 | 1 | 1 | -0,050140618 | 0,532197497 | 1 |
| PGP | 15 | 15 | -2,76582E-16 | 0,99999996 | 1 |
| PDCD5 | 6 | 6 | -3,69694E-19 | 1 | 1 |
| PLTP | 6 | 6 | 2,0311E-17 | 0,999999989 | 1 |
| MAEA | 4 | 4 | -1,56956E-17 | 0,99999999 | 1 |
| ASAP2 | 2 | 2 | 0 | 1 | 1 |
| THNSL2 | 4 | 4 | -0,093748539 | 0,158616698 | 1 |
| P2RY12 | 5 | 5 | -3,84699E-16 | 0,999999948 | 1 |
| NANP | 6 | 5 | 0,052762327 | 0,222450765 | 1 |
| SERGEF | 1 | 1 | 0 | 1 | 1 |
| CNDP2 | 27 | 27 | -7,16317E-17 | 0,999999971 | 1 |
| TUBA4A | 10 | 10 | -4,88326E-17 | 0,999999985 | 1 |
| SCFD2 | 4 | 4 | -3,20923E-16 | 0,999999966 | 1 |

| | | | | | |
|---|---|---|---|---|---|
| CPNE8 | 2 | 2 | -4,41698E-19 | 0,999999999 | 1 |
| NTM | 9 | 9 | -1,19391E-17 | 0,999999991 | 1 |
| ZRANB2 | 4 | 4 | -4,63479E-17 | 0,999999983 | 1 |
| BPGM | 8 | 8 | 1,24833E-16 | 0,999999986 | 1 |
| EEPD1 | 3 | 3 | 0 | 1 | 1 |
| F8A1 | 5 | 5 | -4,55446E-14 | 0,999999325 | 1 |
| AVL9 | 12 | 12 | -1,16944E-15 | 0,999999869 | 1 |
| PTPRT | 4 | 4 | 1,46762E-17 | 0,999999999 | 1 |
| SNX1 | 20 | 20 | 1,66763E-18 | 0,999999997 | 1 |
| PCBD1 | 3 | 3 | 1,49812E-17 | 0,999999991 | 1 |
| TTPAL | 4 | 4 | 0 | 1 | 1 |
| PEX11B | 6 | 6 | 4,85583E-17 | 0,999999997 | 1 |
| TTC37 | 8 | 8 | -1,45764E-17 | 0,999999997 | 1 |
| CCDC71 | 1 | 1 | -0,139295219 | 0,27706938 | 1 |
| NDUFB11 | 12 | 11 | 0 | 1 | 1 |
| ATG4C | 2 | 2 | 0 | 1 | 1 |
| AK4 | 14 | 14 | 1,19469E-15 | 0,999999912 | 1 |
| GMPPA | 8 | 8 | 0 | 1 | 1 |
| AK3 | 15 | 15 | 0 | 1 | 1 |
| PRMT3 | 3 | 3 | -3,4736E-17 | 0,999999989 | 1 |
| LAGE3 | 1 | 1 | 0 | 1 | 1 |
| APP | 21 | 20 | -5,34026E-17 | 0,999999976 | 1 |
| ACSS1 | 18 | 16 | -3,05348E-17 | 0,99999998 | 1 |
| SEC24C | 14 | 14 | 0 | 1 | 1 |
| CPD | 11 | 10 | 8,59823E-16 | 0,999999902 | 1 |
| GZF1 | 1 | 1 | 0 | 1 | 1 |
| CPNE2 | 9 | 9 | 3,08162E-14 | 0,999999505 | 1 |
| HDAC6 | 11 | 11 | 2,16718E-15 | 0,999999805 | 1 |
| RAC3;RAC1 | 1 | 1 | 0 | 1 | 1 |
| RAC1 | 5 | 5 | 4,87909E-15 | 0,99999979 | 1 |
| SERPINA3N | 4 | 4 | 3,46016E-16 | 0,999999975 | 1 |
| ROGDI | 17 | 17 | 7,48377E-18 | 0,999999989 | 1 |
| MAT2B | 14 | 14 | 5,95164E-17 | 0,999999982 | 1 |

| | | | | | |
|---|---|---|---|---|---|
| CFL2 | 8 | 8 | 0,002966762 | 0,83774642 | 1 |
| CFL1 | 16 | 16 | 2,4159E-17 | 0,999999994 | 1 |
| ADIPOQ | 2 | 2 | 0,182325285 | 0,484757134 | 1 |
| GYKL1 | 5 | 4 | -7,91166E-19 | 0,999999998 | 1 |
| LYSMD1 | 6 | 6 | -0,050720167 | 0,324202999 | 1 |
| GPI | 48 | 48 | 0,018529862 | 0,542898638 | 1 |
| METTL26 | 5 | 5 | 0 | 1 | 1 |
| IARS2 | 37 | 36 | 0,029328852 | 0,378698037 | 1 |
| TUBG2 | 5 | 5 | 1,73612E-18 | 0,999999998 | 1 |
| RHOG | 8 | 8 | 0 | 1 | 1 |
| TWF2 | 16 | 16 | 1,12611E-17 | 0,99999999 | 1 |
| SIPA1L2 | 5 | 5 | -1,56185E-16 | 0,999999972 | 1 |
| PTDSS1 | 1 | 1 | 0 | 1 | 1 |
| TUBB4B;TUBB4A | 6 | 6 | 0 | 1 | 1 |
| VMN1R87 | 1 | 1 | 0 | 1 | 1 |
| INF2 | 8 | 8 | 0 | 1 | 1 |
| NIPSNAP3B | 9 | 8 | 0,051759156 | 0,178104939 | 1 |
| STX6 | 7 | 7 | -0,05768909 | 0,227963803 | 1 |
| UNC5A | 6 | 6 | 0 | 1 | 1 |
| GFRA1 | 2 | 2 | 0,061674724 | 0,378401387 | 1 |
| ARL15 | 4 | 4 | 1,87482E-19 | 1 | 1 |
| HNRNPA0 | 5 | 5 | -1,15312E-16 | 0,999999973 | 1 |
| SEPSECS | 3 | 3 | 6,16362E-17 | 0,999999985 | 1 |
| COMMD10 | 3 | 3 | -2,58141E-17 | 0,99999999 | 1 |
| VAPA | 11 | 11 | -9,11344E-17 | 0,999999968 | 1 |
| PHYHIP | 15 | 15 | -1,20433E-16 | 0,999999968 | 1 |
| MAP6D1 | 8 | 8 | -6,85587E-17 | 0,999999975 | 1 |
| TUSC2 | 2 | 2 | 4,81514E-17 | 0,999999993 | 1 |
| PDXDC1 | 14 | 14 | -0,050457513 | 0,155892572 | 1 |
| STX16 | 7 | 6 | -7,16965E-17 | 0,999999978 | 1 |
| PRELID1 | 1 | 1 | -0,080118616 | 0,468967229 | 1 |
| DST | 8 | 8 | -2,71819E-17 | 0,999999985 | 1 |
| DNAJC7 | 11 | 11 | -0,01275769 | 0,58514887 | 1 |

| | | | | | |
|---|---|---|---|---|---|
| NDUFA9 | 29 | 28 | 2,4025E-14 | 0,999999481 | 1 |
| FAM120B | 1 | 1 | 0 | 1 | 1 |
| CTDP1 | 3 | 3 | 0 | 1 | 1 |
| CA2 | 16 | 16 | -4,68008E-15 | 0,999999874 | 1 |
| AKAP7 | 3 | 3 | -0,057119615 | 0,316986893 | 1 |
| ARMCX2 | 1 | 1 | 0 | 1 | 1 |
| PCID2 | 1 | 1 | 0,256677688 | 0,151808401 | 1 |
| ZFAND6 | 1 | 1 | -0,119991622 | 0,220672666 | 1 |
| FAM120C | 10 | 10 | -0,015963496 | 0,50562528 | 1 |
| STARD10 | 2 | 2 | -2,99905E-15 | 0,999999867 | 1 |
| PORCN | 1 | 1 | 0 | 1 | 1 |
| SPEG | 6 | 6 | -2,96501E-17 | 0,999999988 | 1 |
| DIP2B | 20 | 20 | -8,52145E-17 | 0,999999967 | 1 |
| SMG8 | 2 | 1 | 0,121537666 | 0,307980961 | 1 |
| TMX2 | 10 | 10 | 0 | 1 | 1 |
| ATOX1 | 4 | 4 | 1,4487E-17 | 0,999999994 | 1 |
| EPHA7 | 5 | 5 | 0 | 1 | 1 |
| WARS2 | 6 | 6 | 1,52788E-17 | 0,999999994 | 1 |
| EHBP1 | 9 | 9 | 0,001689347 | 0,858242853 | 1 |
| YWHAH | 18 | 18 | 5,60824E-17 | 0,999999992 | 1 |
| YWHAQ | 18 | 18 | 2,84405E-17 | 0,999999989 | 1 |
| YWHAB | 10 | 9 | -6,64873E-17 | 0,999999985 | 1 |
| HNRNPLL | 18 | 18 | -0,033846026 | 0,382434356 | 1 |
| HSPB1 | 4 | 4 | 0 | 1 | 1 |
| TSNAX | 15 | 15 | -1,06061E-16 | 0,999999987 | 1 |
| ATP6V1B1 | 1 | 1 | 0 | 1 | 1 |
| AAMP | 4 | 4 | 0 | 1 | 1 |
| CUL2 | 32 | 31 | -1,42657E-17 | 0,999999994 | 1 |
| CUL5 | 33 | 33 | 0 | 1 | 1 |
| EMC7 | 6 | 6 | -9,63398E-19 | 0,999999999 | 1 |
| S100A16 | 1 | 1 | 0 | 1 | 1 |
| ACOT9 | 30 | 28 | 0,020855941 | 0,456124693 | 1 |
| WDR13 | 19 | 18 | -1,1582E-17 | 0,99999999 | 1 |

| | | | | | |
|---|---|---|---|---|---|
| SYNPO | 18 | 18 | -0,008903714 | 0,696872902 | 1 |
| ARL6IP5 | 4 | 4 | 2,39165E-16 | 0,999999955 | 1 |
| FGF14 | 2 | 2 | 3,80104E-19 | 1 | 1 |
| CYRIB | 20 | 20 | -2,88815E-17 | 0,999999992 | 1 |
| MRPS34 | 7 | 7 | 0 | 1 | 1 |
| TPMT | 6 | 6 | 0 | 1 | 1 |
| NSUN3 | 1 | 1 | 0 | 1 | 1 |
| ALDH7A1 | 24 | 24 | -3,69392E-17 | 0,999999996 | 1 |
| CRMP1 | 7 | 7 | 0 | 1 | 1 |
| SEC63 | 4 | 4 | -7,95277E-18 | 0,999999993 | 1 |
| HSD17B12 | 9 | 9 | 4,94495E-17 | 0,999999979 | 1 |
| COASY | 9 | 9 | 0,01988698 | 0,486104039 | 1 |
| APOOL | 7 | 7 | 5,50321E-18 | 0,999999996 | 1 |
| PTPRJ | 11 | 11 | 4,73943E-18 | 0,999999997 | 1 |
| GNAZ | 20 | 19 | 5,28125E-16 | 0,999999916 | 1 |
| NYAP2 | 4 | 4 | -4,67001E-18 | 0,999999998 | 1 |
| CLPP | 13 | 13 | 0,02876903 | 0,323692438 | 1 |
| NDUFB10 | 10 | 10 | 0 | 1 | 1 |
| HCCS | 3 | 3 | 4,15645E-17 | 0,99999999 | 1 |
| SLC38A3 | 6 | 6 | 0 | 1 | 1 |
| GSPT2 | 18 | 16 | -1,4688E-16 | 0,999999973 | 1 |
| PLS1 | 6 | 5 | -0,097944789 | 0,203091887 | 1 |
| MTCO1 | 3 | 3 | 3,45664E-16 | 0,999999982 | 1 |
| TPH2 | 10 | 10 | 9,8453E-17 | 0,99999997 | 1 |
| EIF1A;EIF1AX | 3 | 3 | 0,016746368 | 0,550685742 | 1 |
| TMCC3 | 5 | 4 | 0,17659925 | 0,169142597 | 1 |
| SARDH | 8 | 6 | 0 | 1 | 1 |
| TUBA3B;TUBA4A;T | 2 | 2 | 0 | 1 | 1 |
| ERLEC1 | 5 | 4 | -4,35432E-17 | 0,999999987 | 1 |
| PTPRS;PTPRD | 7 | 7 | 1,34853E-37 | 1 | 1 |
| IRF7 | 1 | 1 | -1,49635E-20 | 1 | 1 |
| RTN4 | 5 | 5 | -1,04965E-17 | 0,999999994 | 1 |
| NUBPL | 5 | 4 | -1,79481E-19 | 1 | 1 |

| | | | | | |
|---|---|---|---|---|---|
| ARPC3 | 11 | 11 | 0 | 1 | 1 |
| PDE10A | 6 | 5 | 0 | 1 | 1 |
| 5031439G07RIK | 4 | 4 | -8,68036E-16 | 0,999999941 | 1 |
| NAXE | 11 | 11 | 0 | 1 | 1 |
| MRTFA | 2 | 2 | -6,97431E-22 | 1 | 1 |
| RPS11 | 12 | 11 | -7,50888E-16 | 0,999999925 | 1 |
| MFF | 5 | 5 | 1,28634E-17 | 0,999999991 | 1 |
| ZNRF2 | 1 | 1 | -0,353982749 | 0,117908317 | 1 |
| KCNJ3 | 5 | 5 | -0,046477997 | 0,30880793 | 1 |
| MDGA1 | 5 | 4 | 2,31883E-14 | 0,999999641 | 1 |
| CAPZA2 | 14 | 14 | 2,18008E-17 | 0,999999988 | 1 |
| GLB1 | 8 | 8 | -0,012274091 | 0,629150185 | 1 |
| VPS50 | 27 | 27 | -4,94128E-16 | 0,999999915 | 1 |
| DIRAS2;DIRAS1 | 2 | 2 | -3,65685E-16 | 0,999999979 | 1 |
| MEMO1 | 5 | 5 | 0 | 1 | 1 |
| DIRAS2 | 12 | 12 | 1,86721E-17 | 0,999999993 | 1 |
| RAP1B | 13 | 13 | 0 | 1 | 1 |
| RAB3D;RAP1B | 1 | 1 | 0 | 1 | 1 |
| RAB3A | 15 | 15 | 0 | 1 | 1 |
| RAB3C | 11 | 10 | -5,80192E-16 | 0,99999994 | 1 |
| ACTBL2 | 8 | 8 | 0 | 1 | 1 |
| SGIP1 | 21 | 21 | -9,76157E-18 | 0,999999996 | 1 |
| SIRT2 | 16 | 16 | -6,07801E-17 | 0,999999984 | 1 |
| NPTX1 | 14 | 14 | 0,042588396 | 0,221865523 | 1 |
| MAPT | 23 | 23 | -0,029347235 | 0,462249791 | 1 |
| THEM6 | 4 | 4 | -5,8778E-18 | 0,999999997 | 1 |
| PIP4K2C | 20 | 20 | -1,22006E-18 | 0,999999999 | 1 |
| OLFM1 | 13 | 13 | 0 | 1 | 1 |
| SLC1A4 | 12 | 12 | -2,23516E-17 | 0,999999992 | 1 |
| ZC3H3 | 1 | 1 | 0 | 1 | 1 |
| DSTN | 15 | 15 | 5,65278E-15 | 0,999999777 | 1 |
| GSTM5 | 25 | 25 | 0 | 1 | 1 |
| SLC1A3 | 14 | 13 | -0,036008657 | 0,439991675 | 1 |

| | | | | | |
|---|---|---|---|---|---|
| SYNE2 | 3 | 3 | 0 | 1 | 1 |
| CPNE6 | 34 | 34 | 0,027876828 | 0,385968297 | 1 |
| TNRC18 | 2 | 2 | 0 | 1 | 1 |
| TXN | 4 | 4 | -3,47638E-20 | 1 | 1 |
| ACTB | 17 | 17 | 0 | 1 | 1 |
| TMEM200C | 1 | 1 | 0 | 1 | 1 |
| PPP6R3 | 16 | 16 | -3,43138E-16 | 0,999999948 | 1 |
| KCTD12 | 13 | 13 | -2,88572E-17 | 0,999999982 | 1 |
| DNAJA1 | 17 | 17 | 2,20957E-17 | 0,99999999 | 1 |
| MTARC2 | 16 | 15 | 9,43681E-16 | 0,999999901 | 1 |
| KIAA0513 | 18 | 18 | -2,88728E-17 | 0,99999998 | 1 |
| PPP2R5B | 9 | 9 | -1,43008E-16 | 0,999999954 | 1 |
| CDC42;RHOG;RAC2 | 1 | 1 | 0 | 1 | 1 |
| PFN1 | 15 | 15 | 5,23743E-18 | 0,999999998 | 1 |
| NAE1 | 26 | 26 | 0 | 1 | 1 |
| RMND1 | 6 | 5 | 0,010262325 | 0,707509751 | 1 |
| S100A1 | 4 | 4 | 0,020523085 | 0,617826581 | 1 |
| FLOT2 | 28 | 28 | 5,91474E-16 | 0,99999988 | 1 |
| GFUS | 8 | 8 | -0,01950068 | 0,647668928 | 1 |
| TGOLN1;TGOLN2 | 1 | 1 | -0,1657974 | 0,130324678 | 1 |
| SRSF2 | 5 | 5 | -1,98323E-16 | 0,999999968 | 1 |
| APOE | 25 | 24 | 0 | 1 | 1 |
| RPS6KA5 | 5 | 5 | -5,62649E-18 | 0,999999998 | 1 |
| CCSAP | 11 | 11 | -0,023677029 | 0,338174328 | 1 |
| MTIF2 | 6 | 6 | 2,82099E-15 | 0,999999871 | 1 |
| AARS2 | 8 | 7 | -2,9756E-16 | 0,99999996 | 1 |
| ARFGEF2;ARFGEF1 | 8 | 8 | 2,95233E-16 | 0,999999966 | 1 |
| TWF1 | 17 | 17 | -6,31653E-17 | 0,999999984 | 1 |
| AGPAT4 | 6 | 6 | 9,31206E-15 | 0,999999706 | 1 |
| TM9SF3 | 6 | 5 | 0 | 1 | 1 |
| CNTNAP2 | 28 | 28 | 1,74371E-16 | 0,999999945 | 1 |
| WDR48 | 22 | 22 | 0,019316929 | 0,289174908 | 1 |
| AKR1C13 | 1 | 1 | 0 | 1 | 1 |

| | | | | | |
|---|---|---|---|---|---|
| HSPA5 | 41 | 40 | 0 | 1 | 1 |
| HSPA2 | 28 | 28 | -2,19511E-17 | 0,99999999 | 1 |
| MRPL39 | 7 | 7 | 7,97974E-17 | 0,99999998 | 1 |
| AGAP1 | 7 | 7 | 1,44013E-16 | 0,999999968 | 1 |
| NUDT9 | 6 | 5 | -0,07422172 | 0,139138459 | 1 |
| MCU | 20 | 19 | 1,92967E-16 | 0,999999954 | 1 |
| ARF6 | 8 | 8 | 0 | 1 | 1 |
| UROS | 7 | 7 | -0,055391151 | 0,212546707 | 1 |
| SOWAHC | 1 | 1 | 0 | 1 | 1 |
| DNAAF3 | 1 | 1 | 0,207092943 | 0,152898936 | 1 |
| PEX19 | 2 | 2 | 0 | 1 | 1 |
| NPL | 5 | 5 | 0 | 1 | 1 |
| TTC38 | 9 | 9 | -1,34666E-17 | 0,999999993 | 1 |
| TRIM23 | 1 | 1 | 0 | 1 | 1 |
| CPSF1 | 1 | 1 | 0 | 1 | 1 |
| sp\|Q8R3C1\|CB042 | 1 | 1 | 0,038360977 | 0,66069314 | 1 |
| AGPAT3 | 8 | 8 | 0 | 1 | 1 |
| CCT6B | 3 | 3 | 6,1523E-18 | 0,999999998 | 1 |
| DAGLA | 15 | 15 | 0 | 1 | 1 |
| PURG | 7 | 7 | 7,34138E-17 | 0,999999982 | 1 |
| RPS4X | 21 | 21 | 4,01165E-17 | 0,999999992 | 1 |
| ELAVL3 | 13 | 12 | -1,92349E-17 | 0,99999999 | 1 |
| PRDX6 | 5 | 5 | -0,023903758 | 0,650940694 | 1 |
| TRAK1 | 1 | 1 | 0,121195779 | 0,287341736 | 1 |
| FAIM2 | 2 | 2 | 2,91137E-16 | 0,999999971 | 1 |
| FHDC1 | 1 | 1 | 0 | 1 | 1 |
| GPRIN3 | 9 | 9 | 9,0912E-18 | 0,999999996 | 1 |
| TCEAL5 | 8 | 8 | -4,57081E-16 | 0,999999954 | 1 |
| HS1BP3 | 5 | 5 | 0 | 1 | 1 |
| UBE4A | 7 | 7 | -1,26623E-14 | 0,999999767 | 1 |
| KPNA4 | 5 | 5 | 0,038793966 | 0,352435911 | 1 |
| ASAH1 | 15 | 15 | 0 | 1 | 1 |
| RAB3B | 11 | 10 | -0,072055697 | 0,128956425 | 1 |

| | | | | | |
|---|---|---|---|---|---|
| ISOC2A | 2 | 2 | 7,98048E-19 | 0,999999999 | 1 |
| CRACD | 6 | 5 | 0 | 1 | 1 |
| EMC8 | 4 | 4 | -0,016973053 | 0,573687304 | 1 |
| DMTF1 | 1 | 1 | 0 | 1 | 1 |
| NMD3 | 1 | 1 | 0 | 1 | 1 |
| DTX3 | 3 | 3 | -0,015776546 | 0,629302641 | 1 |
| NT5C2 | 9 | 9 | -0,047257061 | 0,162889594 | 1 |
| DYNC1LI2 | 13 | 13 | -4,84684E-17 | 0,999999979 | 1 |
| ARF3;ARF1 | 6 | 6 | -4,52405E-16 | 0,999999969 | 1 |
| ARF5 | 15 | 15 | -4,92042E-17 | 0,99999998 | 1 |
| ARF2 | 4 | 4 | -1,01204E-15 | 0,999999935 | 1 |
| SLC25A51 | 11 | 11 | 0,052354032 | 0,144805512 | 1 |
| PSMB3 | 8 | 8 | 1,61687E-17 | 0,999999993 | 1 |
| EIF4E | 6 | 6 | -3,63291E-17 | 0,999999987 | 1 |
| H4F16 | 8 | 5 | -0,016001042 | 0,717270564 | 1 |
| SLC44A2 | 9 | 9 | 5,91826E-18 | 0,999999997 | 1 |
| HSFY2 | 1 | 1 | 0 | 1 | 1 |
| HPCAL1 | 8 | 8 | -0,003826464 | 0,802420237 | 1 |
| HPCA | 7 | 7 | -8,79992E-18 | 0,999999998 | 1 |
| MT3 | 2 | 2 | 0 | 1 | 1 |
| GAL3ST3 | 2 | 1 | 0 | 1 | 1 |
| RPL3 | 12 | 12 | 2,00704E-17 | 0,999999993 | 1 |
| MCF2L | 1 | 1 | 0,062711407 | 0,444377217 | 1 |
| CALR | 21 | 20 | 9,96062E-18 | 0,999999996 | 1 |
| LDHB | 26 | 26 | -2,69096E-15 | 0,999999916 | 1 |
| ATP2B4 | 13 | 13 | -0,038373219 | 0,336445547 | 1 |
| NEDD4 | 12 | 12 | -3,38624E-13 | 0,999998627 | 1 |
| CADM4 | 12 | 12 | -0,026439153 | 0,39323885 | 1 |
| RILPL1 | 13 | 13 | -0,098123232 | 0,15355474 | 1 |
| SUPT5H | 14 | 14 | -0,02647211 | 0,304356636 | 1 |
| ATP6V1F | 9 | 9 | 0 | 1 | 1 |
| TMX3 | 8 | 8 | -2,11116E-17 | 0,999999995 | 1 |
| SAMM50 | 19 | 19 | 0,033211826 | 0,363150261 | 1 |

| | | | | | |
|---|---|---|---|---|---|
| KCNA1 | 10 | 10 | -2,29924E-18 | 1 | 1 |
| RAMAC | 1 | 1 | 0 | 1 | 1 |
| RABGAP1 | 17 | 17 | -3,79996E-17 | 0,999999985 | 1 |
| GRIPAP1 | 32 | 32 | -0,062516242 | 0,153864754 | 1 |
| SLMAP | 7 | 6 | -1,83792E-16 | 0,999999963 | 1 |
| STX17 | 6 | 6 | -1,71438E-17 | 0,999999989 | 1 |
| SUB1 | 11 | 11 | 0 | 1 | 1 |
| PRDX6 | 24 | 24 | 1,35974E-16 | 0,999999984 | 1 |
| SLC25A46 | 11 | 11 | 0 | 1 | 1 |
| PCMT1 | 15 | 15 | 5,52804E-17 | 0,999999986 | 1 |
| CDC42 | 7 | 7 | 0,018857053 | 0,560289793 | 1 |
| HPCAL4 | 19 | 18 | -8,17533E-18 | 0,999999997 | 1 |
| JAM3 | 7 | 7 | -3,61913E-16 | 0,999999956 | 1 |
| TMEM30A | 11 | 11 | 3,3641E-17 | 0,999999986 | 1 |
| ERP44 | 11 | 11 | 0 | 1 | 1 |
| RAB9B | 4 | 4 | 0,070903746 | 0,172410889 | 1 |
| ITM2B | 7 | 7 | 7,58067E-16 | 0,999999911 | 1 |
| GTPBP1 | 8 | 8 | 1,02874E-16 | 0,999999972 | 1 |
| CRTAC1 | 8 | 8 | -4,09939E-17 | 0,999999985 | 1 |
| GRIN2A | 29 | 28 | 0 | 1 | 1 |
| GGT7 | 13 | 13 | 2,1868E-17 | 0,999999988 | 1 |
| ARHGAP21 | 2 | 2 | -5,26114E-19 | 0,999999999 | 1 |
| KIF5B | 28 | 28 | -0,006027418 | 0,604006289 | 1 |
| MBOAT2 | 1 | 1 | -0,157143509 | 0,232673879 | 1 |
| ATG9A | 9 | 9 | 9,39511E-17 | 0,999999977 | 1 |
| NRXN1 | 13 | 13 | 1,0635E-16 | 0,999999981 | 1 |
| FKBP15 | 5 | 5 | 2,34762E-17 | 0,999999995 | 1 |
| PMPCA | 21 | 20 | 0 | 1 | 1 |
| SLC3A2 | 23 | 23 | -6,13216E-18 | 0,999999997 | 1 |
| PPP2R2A | 20 | 20 | 9,76844E-17 | 0,999999968 | 1 |
| FSCN1 | 30 | 30 | 0 | 1 | 1 |
| PARD3B | 1 | 1 | 0 | 1 | 1 |
| DDAH1 | 24 | 24 | -0,028214367 | 0,518448143 | 1 |

| | | | | | |
|---|---|---|---|---|---|
| AKR1D1 | 1 | 1 | 0 | 1 | 1 |
| UBE4B | 15 | 15 | -1,75661E-16 | 0,999999956 | 1 |
| GMPS | 30 | 29 | -6,9501E-17 | 0,999999972 | 1 |
| TTC19 | 7 | 7 | -1,32387E-17 | 0,999999997 | 1 |
| CD9 | 3 | 3 | 0 | 1 | 1 |
| HSDL2 | 6 | 6 | 0 | 1 | 1 |
| YAP1 | 1 | 1 | 0 | 1 | 1 |
| CAPZB | 26 | 26 | 1,41155E-15 | 0,999999851 | 1 |
| AKT1 | 8 | 8 | 6,30666E-17 | 0,99999999 | 1 |
| AKT3 | 14 | 14 | 3,07437E-16 | 0,99999995 | 1 |
| VAC14 | 16 | 16 | 2,32063E-16 | 0,999999952 | 1 |
| TTLL12 | 14 | 14 | 4,30718E-17 | 0,999999981 | 1 |
| KYAT1 | 10 | 10 | -0,002478744 | 0,849853018 | 1 |
| GM382 | 1 | 1 | 0 | 1 | 1 |
| PDLIM1 | 3 | 3 | 2,0741E-16 | 0,999999971 | 1 |
| LGI2 | 8 | 8 | -0,044680914 | 0,261893003 | 1 |
| SERPINF2 | 4 | 4 | -7,55266E-17 | 0,999999987 | 1 |
| NTNG1 | 5 | 5 | 8,23596E-17 | 0,999999976 | 1 |
| KCND2 | 11 | 11 | 3,12846E-17 | 0,999999981 | 1 |
| ACAD9 | 29 | 28 | -1,81167E-17 | 0,999999994 | 1 |
| ATXN7L3B | 1 | 1 | 0 | 1 | 1 |
| VPS33A | 8 | 8 | -2,46761E-17 | 0,999999987 | 1 |
| EIF3M | 8 | 7 | -0,047205657 | 0,334406612 | 1 |
| EIF2B3 | 10 | 10 | -0,00930858 | 0,62336258 | 1 |
| PPP4C | 2 | 2 | 1,88083E-19 | 0,999999999 | 1 |
| TMX1 | 4 | 4 | 0 | 1 | 1 |
| TLR4 | 1 | 1 | -0,184855307 | 0,37141887 | 1 |
| TMEM65 | 6 | 6 | 0 | 1 | 1 |
| EXOC6B | 16 | 16 | -2,57677E-17 | 0,999999987 | 1 |
| PDHB | 26 | 26 | 0,04239144 | 0,210641232 | 1 |
| DDX39B | 8 | 8 | -2,17149E-15 | 0,999999878 | 1 |
| TIMM10B | 3 | 3 | 8,78005E-18 | 0,999999996 | 1 |
| KALRN;ARHGEF25 | 1 | 1 | 0 | 1 | 1 |

| | | | | | |
|---|---|---|---|---|---|
| TIAM1 | 4 | 4 | -1,06185E-15 | 0,999999914 | 1 |
| PPIB | 13 | 13 | 6,59938E-20 | 1 | 1 |
| PPIL1 | 5 | 5 | -5,92851E-20 | 1 | 1 |
| RPL9 | 8 | 8 | -4,39955E-16 | 0,999999959 | 1 |
| PDLIM5 | 3 | 3 | 0 | 1 | 1 |
| STX12 | 10 | 10 | 5,07633E-17 | 0,999999984 | 1 |
| NDUFB6 | 9 | 9 | 0,092700577 | 0,12553543 | 1 |
| PDIA4 | 34 | 34 | 0 | 1 | 1 |
| COPS8 | 7 | 7 | -2,65744E-14 | 0,999999327 | 1 |
| SEPTIN4 | 4 | 4 | 0 | 1 | 1 |
| NDUFS3 | 17 | 16 | 0,051352204 | 0,189497808 | 1 |
| ANK3;ANK2 | 6 | 6 | 0 | 1 | 1 |
| DAB1 | 4 | 4 | 0 | 1 | 1 |
| PDLIM4 | 1 | 1 | 0 | 1 | 1 |
| FEZ1 | 3 | 3 | 0 | 1 | 1 |
| PGRMC2 | 4 | 4 | -1,95517E-18 | 0,999999996 | 1 |
| USP46 | 4 | 4 | -9,30965E-18 | 0,999999993 | 1 |
| HIBADH | 11 | 10 | 0,027257113 | 0,445741344 | 1 |
| GAA | 16 | 16 | -2,73574E-18 | 0,999999997 | 1 |
| HBB-B1 | 13 | 13 | 0,004247904 | 0,836159081 | 1 |
| PGRMC1 | 11 | 11 | -0,000186839 | 0,939148852 | 1 |
| ATP5MF | 5 | 5 | 0 | 1 | 1 |
| EIF4A2 | 11 | 11 | -0,061380367 | 0,156608686 | 1 |
| STARD5 | 2 | 2 | 0,103107917 | 0,162325435 | 1 |
| AFG3L2 | 42 | 41 | 4,10197E-16 | 0,999999931 | 1 |
| PLAA | 24 | 24 | 0 | 1 | 1 |
| SUSD2 | 2 | 2 | 0 | 1 | 1 |
| DCUN1D1 | 8 | 8 | -0,011401882 | 0,603996692 | 1 |
| MCM6 | 1 | 1 | 0 | 1 | 1 |
| CADM2 | 18 | 18 | 0 | 1 | 1 |
| KCNQ2 | 7 | 7 | 0 | 1 | 1 |
| POR | 22 | 19 | 0,037375413 | 0,157893397 | 1 |
| CES1D | 1 | 1 | 0,045496393 | 0,819047613 | 1 |

| | | | | | |
|---|---|---|---|---|---|
| B630019K06RIK | 3 | 3 | 0 | 1 | 1 |
| FSCN2 | 1 | 1 | 0 | 1 | 1 |
| MTX1 | 8 | 7 | 0,054905289 | 0,177084703 | 1 |
| ERP29 | 9 | 9 | 1,56377E-17 | 0,999999989 | 1 |
| NRP2 | 10 | 10 | 2,11803E-17 | 0,999999984 | 1 |
| ARSB | 8 | 8 | -0,092184473 | 0,123001434 | 1 |
| CAPN2 | 22 | 21 | -0,064356507 | 0,246447336 | 1 |
| RASA3 | 14 | 14 | -2,18011E-18 | 0,999999999 | 1 |
| ADPRH | 13 | 13 | 1,63009E-17 | 0,999999995 | 1 |
| TRAPPC1 | 4 | 4 | 0 | 1 | 1 |
| SELENBP1 | 20 | 20 | 3,60148E-18 | 0,999999999 | 1 |
| IGHM | 9 | 7 | -0,265241012 | 0,156208057 | 1 |
| AHCYL2 | 13 | 12 | 0 | 1 | 1 |
| GSTZ1 | 11 | 11 | -4,78428E-17 | 0,999999989 | 1 |
| UBAP2L | 18 | 18 | -0,059603792 | 0,279985085 | 1 |
| SLC12A7 | 1 | 1 | 0 | 1 | 1 |
| WDR37 | 17 | 17 | 9,60915E-18 | 0,999999988 | 1 |
| SYN2 | 31 | 30 | 0 | 1 | 1 |
| ARHGAP5 | 18 | 18 | 0 | 1 | 1 |
| PCDHGA2 | 1 | 1 | 0 | 1 | 1 |
| FUCA1 | 1 | 1 | 0 | 1 | 1 |
| ACOT1 | 3 | 3 | 0 | 1 | 1 |
| G6PDX | 24 | 24 | 0 | 1 | 1 |
| DNAJB11 | 8 | 8 | -1,11969E-16 | 0,999999965 | 1 |
| MDP1 | 5 | 5 | 1,01793E-17 | 0,999999994 | 1 |
| DNAJA2 | 19 | 19 | -0,031579273 | 0,254520435 | 1 |
| CALM1;CALM2;CAL | 9 | 9 | 0 | 1 | 1 |
| S100A11 | 3 | 3 | 0 | 1 | 1 |
| NETO1 | 2 | 1 | 0 | 1 | 1 |
| SOS1 | 2 | 2 | 0 | 1 | 1 |
| RCC2 | 1 | 1 | 0 | 1 | 1 |
| VIM | 13 | 7 | -8,28921E-17 | 0,999999989 | 1 |
| NSUN2 | 12 | 12 | -5,35648E-17 | 0,99999999 | 1 |

| | | | | | |
|---|---|---|---|---|---|
| DCLK2 | 18 | 18 | 0,019563354 | 0,288027505 | 1 |
| DNAJB1 | 7 | 7 | -0,036269453 | 0,334216416 | 1 |
| CPNE5 | 6 | 6 | 0 | 1 | 1 |
| MAP4K3 | 7 | 7 | 6,00645E-17 | 0,999999984 | 1 |
| MCCC1 | 29 | 25 | 0 | 1 | 1 |
| WNK1 | 10 | 10 | -0,057732398 | 0,317750606 | 1 |
| FGFR3 | 2 | 2 | 0 | 1 | 1 |
| SOD1 | 10 | 10 | 0 | 1 | 1 |
| MPP3 | 11 | 10 | 2,91984E-18 | 0,999999997 | 1 |
| ZC3H13 | 1 | 1 | -0,351863168 | 0,189137504 | 1 |
| HMCN1 | 1 | 1 | 0 | 1 | 1 |
| ELP2 | 8 | 8 | -0,029200156 | 0,325594577 | 1 |
| UEVLD | 2 | 2 | -2,33486E-15 | 0,999999889 | 1 |
| MACROD2 | 9 | 9 | 7,17789E-17 | 0,999999985 | 1 |
| RHOA | 5 | 5 | -5,84461E-15 | 0,999999911 | 1 |
| PPP3R1 | 20 | 19 | -0,044465084 | 0,392071957 | 1 |
| AHI1 | 4 | 4 | 0,059637129 | 0,164176015 | 1 |
| CAPRIN1 | 14 | 14 | -0,101200069 | 0,131873042 | 1 |
| RPS21 | 6 | 6 | -3,08937E-17 | 0,999999989 | 1 |
| LMAN2 | 8 | 8 | 0 | 1 | 1 |
| CADM3 | 18 | 17 | 2,38868E-18 | 0,999999997 | 1 |
| SPRED1 | 6 | 5 | 0 | 1 | 1 |
| NUDT16;NUDT16L1 | 1 | 1 | 0 | 1 | 1 |
| ABAT | 38 | 36 | 0,017080548 | 0,541090584 | 1 |
| OAS1F | 1 | 1 | 0 | 1 | 1 |
| RIMBP2 | 19 | 17 | 6,23066E-12 | 0,999990316 | 1 |
| ITIH3 | 4 | 4 | -7,29647E-16 | 0,999999956 | 1 |
| UBE2I | 4 | 4 | -9,53816E-17 | 0,999999996 | 1 |
| SEMA4D | 4 | 4 | 4,82878E-16 | 0,99999996 | 1 |
| PPME1 | 18 | 17 | -9,88328E-17 | 0,999999968 | 1 |
| ATP6AP2 | 4 | 4 | 4,89136E-18 | 0,999999996 | 1 |
| DNAJA4 | 12 | 12 | -0,028001319 | 0,270352827 | 1 |
| DRG1 | 9 | 9 | 7,1188E-17 | 0,999999986 | 1 |

| | | | | | |
|---|---|---|---|---|---|
| GUK1 | 10 | 10 | 0 | 1 | 1 |
| DYNLL1 | 8 | 8 | 0,019032371 | 0,47281189 | 1 |
| SH3BGRL | 9 | 9 | 3,46605E-18 | 0,999999998 | 1 |
| DYNLL2 | 2 | 2 | 0,100860396 | 0,198222573 | 1 |
| TLE2 | 1 | 1 | 0 | 1 | 1 |
| UGT8 | 2 | 2 | 0,038441175 | 0,503767073 | 1 |
| YWHAZ | 27 | 27 | -3,1137E-17 | 0,999999995 | 1 |
| LSMEM1 | 1 | 1 | 0 | 1 | 1 |
| GPX4 | 9 | 9 | 1,11159E-14 | 0,999999558 | 1 |
| GM49601;SEPTIN5 | 2 | 2 | 1,91796E-17 | 0,999999987 | 1 |
| SLITRK1 | 3 | 3 | 0 | 1 | 1 |
| MRPS6 | 1 | 1 | 0 | 1 | 1 |
| NDUFV2 | 15 | 14 | 0,021800746 | 0,424690068 | 1 |
| USP51 | 1 | 1 | 0 | 1 | 1 |
| SUCLG2 | 18 | 16 | 0 | 1 | 1 |
| EXOC6 | 6 | 6 | 0 | 1 | 1 |
| SLC27A4 | 23 | 22 | 3,74971E-17 | 0,999999978 | 1 |
| SULT4A1 | 11 | 11 | -5,0294E-17 | 0,99999998 | 1 |
| RPS29 | 2 | 2 | 0 | 1 | 1 |
| SBDS | 7 | 7 | 0 | 1 | 1 |
| MYO1D | 27 | 21 | 0 | 1 | 1 |
| ELMO2;ELMO1 | 3 | 3 | 0 | 1 | 1 |
| CDH10 | 7 | 7 | 0 | 1 | 1 |
| VARS2 | 11 | 9 | -2,26976E-17 | 0,999999996 | 1 |
| PSMD7 | 14 | 14 | -2,42035E-17 | 0,999999993 | 1 |
| HINT3 | 2 | 2 | 0 | 1 | 1 |
| NEK7 | 2 | 2 | -0,076892647 | 0,258959859 | 1 |
| CSNK1A1 | 9 | 9 | -0,00056174 | 0,909041973 | 1 |
| CHEK1 | 1 | 1 | 0 | 1 | 1 |
| GSK3A | 9 | 9 | -1,0577E-17 | 0,999999992 | 1 |
| CAMKK1 | 17 | 17 | -6,3552E-16 | 0,999999918 | 1 |
| RIN1 | 1 | 1 | 0 | 1 | 1 |
| DOCK2 | 5 | 5 | 5,74358E-15 | 0,999999803 | 1 |

| | | | | | |
|---|---|---|---|---|---|
| CYB5R3 | 17 | 16 | 0,007971998 | 0,65195112 | 1 |
| ZYG11B | 8 | 8 | -0,019736986 | 0,484241907 | 1 |
| DHRS7 | 4 | 4 | 2,60722E-17 | 0,999999991 | 1 |
| TUBGCP3 | 6 | 6 | 0,009227803 | 0,701748479 | 1 |
| TSTD3 | 2 | 2 | 5,44306E-17 | 0,999999984 | 1 |
| HEATR3 | 1 | 1 | -0,13876198 | 0,335452207 | 1 |
| PCSK2 | 5 | 5 | 0 | 1 | 1 |
| UBE2M | 9 | 9 | 0 | 1 | 1 |
| FMR1 | 6 | 6 | -0,050660957 | 0,30723736 | 1 |
| GCC2 | 5 | 5 | -0,084629601 | 0,17463374 | 1 |
| RPS6 | 12 | 12 | 1,52431E-17 | 0,999999995 | 1 |
| NRK | 1 | 1 | 0 | 1 | 1 |
| PLXNC1 | 11 | 9 | 0 | 1 | 1 |
| DLG1 | 6 | 5 | 1,23702E-17 | 0,999999991 | 1 |
| VCAM1 | 15 | 15 | 0 | 1 | 1 |
| PSMD8 | 10 | 10 | -0,011633078 | 0,658366439 | 1 |
| ENTPD1 | 1 | 1 | 0 | 1 | 1 |
| MESD | 5 | 5 | 0 | 1 | 1 |
| GM7356 | 1 | 1 | 3,96909E-21 | 1 | 1 |
| RPS26 | 2 | 2 | 6,43245E-17 | 0,999999993 | 1 |
| SLC6A17 | 20 | 20 | 0,002621886 | 0,833271359 | 1 |
| ANP32B | 4 | 4 | -5,39069E-18 | 0,999999995 | 1 |
| HSP90B1 | 45 | 45 | -9,43851E-18 | 0,999999993 | 1 |
| PITHD1 | 11 | 11 | -1,12057E-16 | 0,999999976 | 1 |
| HP | 8 | 7 | 1,70724E-15 | 0,999999956 | 1 |
| CCNY | 9 | 8 | -1,74593E-14 | 0,999999801 | 1 |
| ACTR3B;ACTR3 | 6 | 5 | 0 | 1 | 1 |
| sp\|P01654\|KV3A1_ | 1 | 1 | 0 | 1 | 1 |
| IGKV5-39 | 1 | 1 | 0 | 1 | 1 |
| CPNE7;CPNE4;CPNI | 1 | 1 | 0 | 1 | 1 |
| RASGEF1A | 3 | 3 | 0 | 1 | 1 |
| VWA8 | 35 | 31 | 4,30403E-18 | 0,999999994 | 1 |
| IGHV1-4;IGHV1-7;I( | 1 | 1 | 0,228487229 | 0,368506409 | 1 |

| | | | | | |
|---|---|---|---|---|---|
| DDX3Y;DDX3X | 17 | 17 | -5,1479E-17 | 0,999999971 | 1 |
| ACTR2 | 22 | 22 | 0 | 1 | 1 |
| HEPACAM | 11 | 11 | -0,03338313 | 0,234631497 | 1 |
| TBC1D13 | 6 | 6 | 0 | 1 | 1 |
| ANO3 | 1 | 1 | 0,2423212 | 0,165000831 | 1 |
| 2310061I04RIK | 8 | 8 | 0,011206234 | 0,62771152 | 1 |
| CUL4B | 17 | 17 | -0,041879762 | 0,233961644 | 1 |
| LRRC59 | 7 | 7 | 4,16393E-16 | 0,999999929 | 1 |
| TMED9 | 4 | 4 | -7,34963E-16 | 0,999999948 | 1 |
| TMED4 | 4 | 4 | 0 | 1 | 1 |
| BRSK1;BRSK2 | 2 | 2 | 0 | 1 | 1 |
| EHD3 | 23 | 23 | 1,71719E-16 | 0,999999965 | 1 |
| PLCB1 | 73 | 72 | 9,61531E-17 | 0,999999953 | 1 |
| MAP2 | 16 | 16 | 0 | 1 | 1 |
| RTN4 | 3 | 3 | 0 | 1 | 1 |
| CCDC25 | 4 | 4 | -6,31578E-19 | 1 | 1 |
| RARS2 | 11 | 10 | 7,2461E-17 | 0,999999986 | 1 |
| LDHA | 26 | 26 | 0,068077489 | 0,181621117 | 1 |
| PTK2B | 44 | 44 | -1,16937E-17 | 0,999999991 | 1 |
| GOLM1 | 2 | 2 | 0,023528083 | 0,680504832 | 1 |
| CNNM4 | 3 | 3 | 0,097208974 | 0,218303676 | 1 |
| ARL8A | 4 | 4 | 3,46852E-16 | 0,999999953 | 1 |
| MOGS | 12 | 12 | -0,026361264 | 0,496074187 | 1 |
| AARSD1 | 12 | 12 | -0,032948075 | 0,218815513 | 1 |
| KMT2C | 1 | 1 | 0 | 1 | 1 |
| RAB4A | 9 | 8 | -0,000330024 | 0,922057471 | 1 |
| NDRG4 | 9 | 9 | -3,3955E-17 | 0,999999991 | 1 |
| SMAP2 | 9 | 8 | 0 | 1 | 1 |
| RAB4B | 9 | 9 | 0 | 1 | 1 |
| MDH1 | 19 | 19 | -6,66751E-18 | 0,999999999 | 1 |
| DNAJB2 | 7 | 7 | 0 | 1 | 1 |
| GJB6 | 2 | 2 | 0 | 1 | 1 |
| CLTB | 10 | 10 | 2,33766E-15 | 0,999999877 | 1 |

| | | | | | |
|---|---|---|---|---|---|
| NDRG3 | 14 | 13 | -2,36032E-19 | 1 | 1 |
| GRHPR | 15 | 15 | 0 | 1 | 1 |
| OGDHL | 49 | 49 | 0,032820304 | 0,284235946 | 1 |
| MAPRE2 | 14 | 14 | 0,011925896 | 0,547642665 | 1 |
| HDGFL3 | 6 | 5 | 2,51887E-17 | 0,999999994 | 1 |
| FIBP | 7 | 7 | 9,0625E-16 | 0,999999907 | 1 |
| TPI1 | 20 | 20 | 6,14269E-18 | 0,999999998 | 1 |
| PLEKHO2 | 2 | 2 | -5,64043E-18 | 0,999999994 | 1 |
| GDPD5 | 1 | 1 | 0 | 1 | 1 |
| DIP2C | 15 | 15 | -2,04917E-16 | 0,999999968 | 1 |
| CEP170;CEP170B | 2 | 2 | -2,10634E-18 | 0,999999997 | 1 |
| GDI2 | 38 | 37 | 0 | 1 | 1 |
| TNRC6B | 3 | 3 | -0,047069528 | 0,565273677 | 1 |
| F3 | 7 | 7 | -0,041958473 | 0,31525548 | 1 |
| RNF187 | 1 | 1 | 1,07252E-18 | 0,999999998 | 1 |
| CLPX | 10 | 8 | 0,035892328 | 0,341293231 | 1 |
| BLVRA | 17 | 17 | 2,79726E-18 | 0,999999997 | 1 |
| HNRNPD | 13 | 12 | -0,10167096 | 0,15242436 | 1 |
| APPL1 | 23 | 22 | -0,005438986 | 0,68715624 | 1 |
| PRKCZ | 2 | 2 | 0,008479674 | 0,742564294 | 1 |
| CARMIL1 | 8 | 8 | -0,041118813 | 0,346005706 | 1 |
| GRK2 | 23 | 23 | 0,005171631 | 0,623560065 | 1 |
| ADRBK2 | 3 | 3 | 1,03995E-15 | 0,99999992 | 1 |
| SMOK3A;SMOK3B | 1 | 1 | 0 | 1 | 1 |
| PHKG1 | 3 | 3 | 0 | 1 | 1 |
| CAMK2B | 19 | 19 | 7,00643E-16 | 0,999999901 | 1 |
| CAMK1 | 8 | 8 | 1,06678E-16 | 0,999999964 | 1 |
| CAMK1D | 15 | 14 | 1,70742E-18 | 0,999999997 | 1 |
| PRKAA2 | 10 | 10 | 1,71273E-18 | 0,999999999 | 1 |
| SNRK | 5 | 5 | 0 | 1 | 1 |
| CELF4 | 4 | 4 | -0,018814348 | 0,542504665 | 1 |
| CDK5 | 13 | 13 | 1,34792E-17 | 0,999999996 | 1 |
| CDK14 | 6 | 6 | 0 | 1 | 1 |

| | | | | | |
|---|---|---|---|---|---|
| PDPR | 16 | 12 | 0 | 1 | 1 |
| TPBG | 3 | 3 | 0 | 1 | 1 |
| ZC3H7B | 2 | 2 | 0 | 1 | 1 |
| RPS27L | 1 | 1 | 0 | 1 | 1 |
| RPS27 | 1 | 1 | 0 | 1 | 1 |
| CCDC22 | 10 | 10 | -1,59127E-16 | 0,999999951 | 1 |
| APBB1 | 9 | 9 | 0 | 1 | 1 |
| FRAS1 | 1 | 1 | 0,034480898 | 0,645105455 | 1 |
| SEC24B | 9 | 9 | 0 | 1 | 1 |
| NDUFAF4 | 8 | 8 | 0 | 1 | 1 |
| LANCL2 | 18 | 18 | -4,73967E-18 | 0,999999997 | 1 |
| TMED10 | 9 | 9 | 0 | 1 | 1 |
| KIF1A | 3 | 3 | -0,04555474 | 0,31483588 | 1 |
| AIDA | 6 | 6 | -0,077561672 | 0,117017109 | 1 |
| HPRT1 | 15 | 15 | 3,36963E-17 | 0,999999993 | 1 |
| PGLS | 10 | 10 | 0 | 1 | 1 |
| CAMSAP3 | 1 | 1 | 0 | 1 | 1 |
| SNAP29 | 9 | 9 | 0 | 1 | 1 |
| CYLD | 11 | 11 | 0 | 1 | 1 |
| SMARCA2 | 2 | 2 | -1,25411E-19 | 1 | 1 |
| CASTOR2 | 6 | 6 | 0 | 1 | 1 |
| CHN1 | 6 | 6 | 2,22224E-17 | 0,999999988 | 1 |
| ARL8B | 7 | 7 | 1,85362E-17 | 0,999999989 | 1 |
| CSDC2 | 2 | 2 | 3,03006E-18 | 0,999999997 | 1 |
| PIP4K2B | 25 | 24 | 0 | 1 | 1 |
| URGCP | 3 | 3 | -0,068893402 | 0,267672175 | 1 |
| FNTA | 9 | 9 | 0 | 1 | 1 |
| SLC2A1 | 7 | 7 | -1,16229E-16 | 0,999999972 | 1 |
| HNMT | 10 | 10 | -1,50573E-17 | 0,999999996 | 1 |
| ASRGL1 | 13 | 13 | -0,069533426 | 0,116993563 | 1 |
| CAMSAP1 | 9 | 9 | -3,18833E-17 | 0,999999988 | 1 |
| CKAP4 | 31 | 30 | -0,002954925 | 0,814133478 | 1 |
| LRBA | 2 | 2 | 2,53304E-20 | 1 | 1 |

| | | | | | |
|---|---|---|---|---|---|
| STMN1;STMN2 | 4 | 4 | -0,049345235 | 0,22674483 | 1 |
| NAA15;NAA16 | 5 | 5 | -3,29042E-17 | 0,999999988 | 1 |
| ERMN | 10 | 10 | -9,19273E-15 | 0,999999763 | 1 |
| ANP32A | 16 | 16 | -0,066611294 | 0,213950862 | 1 |
| ANP32E | 3 | 3 | -4,11844E-16 | 0,999999953 | 1 |
| ARHGAP35 | 26 | 26 | -3,48477E-17 | 0,999999978 | 1 |
| ACTA1;ACTBL2 | 12 | 11 | 0 | 1 | 1 |
| PPP1R9A | 7 | 7 | -0,024937722 | 0,41996272 | 1 |
| WDR54 | 7 | 7 | -0,048934829 | 0,194506187 | 1 |
| SHISA4 | 1 | 1 | 0 | 1 | 1 |
| MLKL | 1 | 1 | 0 | 1 | 1 |
| GALT | 1 | 1 | 0 | 1 | 1 |
| LRRC58 | 1 | 1 | 0 | 1 | 1 |
| SIRT3 | 3 | 3 | -6,70745E-16 | 0,999999927 | 1 |
| HSPB11 | 2 | 2 | 0,031771703 | 0,447983976 | 1 |
| ZMYND10 | 1 | 1 | 0 | 1 | 1 |
| ACTG2 | 2 | 2 | 0 | 1 | 1 |
| SMIM8 | 1 | 1 | 0 | 1 | 1 |
| HNRNPR | 3 | 3 | -0,008057282 | 0,744745881 | 1 |
| VPS16 | 1 | 1 | 0 | 1 | 1 |
| PLPPR2 | 8 | 8 | 0,022592739 | 0,506889705 | 1 |
| TAGLN;TAGLN3 | 1 | 1 | 0 | 1 | 1 |
| DCAF7 | 7 | 7 | -3,25397E-17 | 0,999999984 | 1 |
| RPS15 | 3 | 3 | -7,01904E-16 | 0,99999996 | 1 |
| DDI2 | 8 | 8 | -1,1863E-15 | 0,999999875 | 1 |
| NDUFB8 | 8 | 7 | 0,04817155 | 0,219138985 | 1 |
| MMAA | 5 | 4 | 0 | 1 | 1 |
| CLVS2 | 7 | 7 | -9,87093E-18 | 0,999999991 | 1 |
| PIP4K2A | 17 | 17 | 0 | 1 | 1 |
| RAB12 | 11 | 10 | -0,018167176 | 0,465429151 | 1 |
| UBE2Q1 | 7 | 7 | -1,15203E-16 | 0,999999965 | 1 |
| RAB9A | 6 | 6 | -0,02563689 | 0,42575838 | 1 |
| ARL8A;ARL8B | 6 | 6 | 0 | 1 | 1 |

| | | | | | |
|---|---|---|---|---|---|
| PLEKHA2 | 3 | 3 | -0,039852984 | 0,548954945 | 1 |
| PLEKHA1 | 2 | 2 | 0 | 1 | 1 |
| LYN | 4 | 3 | 0,073254307 | 0,300552266 | 1 |
| TRIM3 | 15 | 15 | 0,007486035 | 0,605419798 | 1 |
| ESPN | 1 | 1 | 0,11458925 | 0,353122297 | 1 |
| THY1 | 8 | 8 | 2,85623E-17 | 0,999999981 | 1 |
| NT5C3A | 7 | 7 | -1,51625E-16 | 0,999999963 | 1 |
| CSF1 | 1 | 1 | 0 | 1 | 1 |
| UBE2E1;UBE2E2 | 2 | 2 | -0,0805476 | 0,312914784 | 1 |
| SNX16 | 6 | 6 | -4,51132E-15 | 0,999999845 | 1 |
| RPS17 | 10 | 10 | -0,02278879 | 0,379988813 | 1 |
| CYTH3 | 3 | 3 | 8,48542E-16 | 0,999999921 | 1 |
| CYTH1 | 3 | 3 | -0,083439729 | 0,282066623 | 1 |
| SHPRH | 1 | 1 | 0 | 1 | 1 |
| MTMR1 | 14 | 14 | -2,12926E-15 | 0,999999871 | 1 |
| TEX2 | 11 | 10 | 0 | 1 | 1 |
| FHIP1A | 1 | 1 | 0 | 1 | 1 |
| GM8251 | 2 | 2 | 9,06504E-18 | 0,999999994 | 1 |
| PALM | 4 | 4 | -1,00937E-16 | 0,999999981 | 1 |
| PSMD12 | 25 | 25 | -2,04103E-15 | 0,99999978 | 1 |
| NSDHL | 3 | 3 | -3,13133E-18 | 0,999999997 | 1 |
| BAG4 | 4 | 4 | -0,096321339 | 0,114565454 | 1 |
| SLC30A3 | 5 | 5 | -5,92573E-17 | 0,999999984 | 1 |
| NEGR1 | 10 | 10 | -9,76371E-18 | 0,999999992 | 1 |
| BRAF | 3 | 3 | 0 | 1 | 1 |
| HARS1;HARS2 | 3 | 2 | 0 | 1 | 1 |
| MYL6 | 8 | 8 | 7,52861E-17 | 0,999999991 | 1 |
| VMP1 | 2 | 2 | 0 | 1 | 1 |
| MAP1LC3A | 4 | 4 | 1,76796E-15 | 0,999999913 | 1 |
| BRAF | 23 | 22 | -3,0279E-16 | 0,999999959 | 1 |
| ATP1B3 | 9 | 9 | -1,53598E-17 | 0,999999996 | 1 |
| AKAP13 | 1 | 1 | 0 | 1 | 1 |
| TSFM | 8 | 6 | 0 | 1 | 1 |

| | | | | | |
|---|---|---|---|---|---|
| KNDC1 | 2 | 2 | -0,08147295 | 0,235155596 | 1 |
| ARHGEF18 | 5 | 5 | 7,50713E-18 | 0,999999995 | 1 |
| SH2D5 | 2 | 2 | 0 | 1 | 1 |
| NLGN3;NLGN1 | 1 | 1 | 0,109970315 | 0,299197014 | 1 |
| JAKMIP2 | 4 | 3 | 0,067934145 | 0,255205315 | 1 |
| DPH7 | 2 | 2 | 0 | 1 | 1 |
| RAD23A | 5 | 5 | -6,78062E-18 | 0,999999998 | 1 |
| C1QA | 5 | 5 | 0 | 1 | 1 |
| HDAC4 | 10 | 10 | -4,09997E-17 | 0,999999989 | 1 |
| MARK2 | 14 | 14 | 0 | 1 | 1 |
| GTF2I | 1 | 1 | 0 | 1 | 1 |
| SERPINA1D | 5 | 5 | -0,068755399 | 0,427941044 | 1 |
| MTR | 8 | 8 | -8,23033E-17 | 0,999999972 | 1 |
| FAM171B | 12 | 12 | -3,45607E-15 | 0,99999982 | 1 |
| ABCD3 | 9 | 9 | -5,56084E-17 | 0,999999977 | 1 |
| CFAP74 | 1 | 1 | -0,018604298 | 0,84877525 | 1 |
| PUM2;PUM1 | 2 | 2 | -0,028367022 | 0,474949718 | 1 |
| MANF | 3 | 3 | 0 | 1 | 1 |
| TTYH2 | 1 | 1 | 0,109847325 | 0,326416256 | 1 |
| PFN2 | 10 | 10 | 1,98842E-17 | 0,999999992 | 1 |
| UFC1 | 6 | 6 | 4,92919E-16 | 0,999999959 | 1 |
| STX1A | 27 | 26 | 1,47031E-17 | 0,999999992 | 1 |
| MCTP1 | 3 | 3 | -7,20358E-17 | 0,999999995 | 1 |
| ELP6 | 1 | 1 | 0 | 1 | 1 |
| NME2 | 6 | 6 | 6,20192E-20 | 1 | 1 |
| NME1 | 18 | 18 | -1,12344E-16 | 0,999999978 | 1 |
| CARHSP1 | 4 | 4 | -1,0898E-13 | 0,999999022 | 1 |
| REEP1 | 2 | 2 | 0 | 1 | 1 |
| GLS | 11 | 11 | 0,059666092 | 0,154441023 | 1 |
| RAB6A | 11 | 11 | -0,026628091 | 0,45290473 | 1 |
| NRIP3 | 3 | 3 | 7,98145E-19 | 0,999999999 | 1 |
| NRIP2 | 3 | 3 | -0,049118159 | 0,382778028 | 1 |
| MRPL4 | 6 | 6 | 0 | 1 | 1 |

| | | | | | |
|---|---|---|---|---|---|
| KLC2 | 4 | 4 | -1,88598E-16 | 0,999999959 | 1 |
| HCN2 | 9 | 9 | 0,014931795 | 0,53677112 | 1 |
| GRID1 | 7 | 6 | 0 | 1 | 1 |
| NRAS;HRAS | 6 | 6 | 0,020240502 | 0,469559108 | 1 |
| CBX3 | 7 | 7 | -0,006668873 | 0,739269324 | 1 |
| RTN4RL2 | 6 | 6 | -1,65866E-16 | 0,999999975 | 1 |
| HSPE1 | 12 | 9 | 3,83453E-17 | 0,999999995 | 1 |
| MLF2 | 6 | 6 | -0,038960672 | 0,319026902 | 1 |
| KCNQ2 | 11 | 11 | 7,72615E-16 | 0,999999915 | 1 |
| KRAS | 13 | 13 | 8,84537E-16 | 0,999999915 | 1 |
| NKAIN4 | 1 | 1 | 0 | 1 | 1 |
| NCK1 | 5 | 5 | -0,055478431 | 0,207160529 | 1 |
| NCK2 | 12 | 12 | 0 | 1 | 1 |
| PPCS | 3 | 3 | -4,0202E-18 | 0,999999996 | 1 |
| CACNA2D1 | 2 | 2 | 7,81681E-19 | 1 | 1 |
| ANK1 | 19 | 19 | 0,001904183 | 0,838908522 | 1 |
| RPS13 | 10 | 10 | -5,90011E-17 | 0,999999978 | 1 |
| USP24 | 26 | 26 | -3,86833E-17 | 0,999999978 | 1 |
| CPOX | 9 | 8 | 0 | 1 | 1 |
| CBARP | 8 | 8 | 8,73183E-17 | 0,999999973 | 1 |
| TBC1D8B | 6 | 6 | -2,04235E-17 | 0,999999991 | 1 |
| MIA3 | 9 | 9 | 0,052808307 | 0,173970111 | 1 |
| SCN3A;SCN2A | 1 | 1 | 0,017564844 | 0,653223232 | 1 |
| THOP1 | 5 | 4 | -0,053750387 | 0,224199546 | 1 |
| CACNB3 | 11 | 11 | 0,060128186 | 0,124933196 | 1 |
| VPS4A | 5 | 5 | -3,07633E-17 | 0,999999984 | 1 |
| PUM1 | 6 | 6 | -0,057255547 | 0,197022425 | 1 |
| RPL36A | 2 | 2 | 9,32814E-19 | 0,999999999 | 1 |
| RAB21 | 10 | 10 | -0,032772397 | 0,18118931 | 1 |
| LGALS1 | 8 | 8 | -5,00852E-16 | 0,999999966 | 1 |
| GSK3B | 9 | 8 | -1,39411E-16 | 0,999999955 | 1 |
| MAP4K4 | 5 | 5 | -5,12522E-18 | 0,999999998 | 1 |
| CNR1 | 9 | 9 | 0,030284713 | 0,42558433 | 1 |

| | | | | | |
|---|---|---|---|---|---|
| CATSPERB | 1 | 1 | 0,069328492 | 0,393779667 | 1 |
| SLC25A10 | 8 | 7 | 1,7209E-18 | 0,999999998 | 1 |
| ARHGAP26 | 11 | 11 | 0 | 1 | 1 |
| KPNA3 | 9 | 9 | 7,66558E-18 | 0,999999992 | 1 |
| GBA2 | 11 | 11 | 3,82341E-17 | 0,999999994 | 1 |
| CCDC88A | 12 | 12 | 1,39922E-17 | 0,999999997 | 1 |
| SPART | 7 | 7 | -1,65794E-17 | 0,999999991 | 1 |
| sp\|Q8BH50\|CR025 | 2 | 2 | -5,33909E-16 | 0,999999981 | 1 |
| SDCBP | 5 | 5 | 1,93081E-16 | 0,999999964 | 1 |
| CAMK1G | 1 | 1 | 0 | 1 | 1 |
| BLVRB | 7 | 7 | -2,26961E-17 | 0,999999991 | 1 |
| CAMSAP2 | 17 | 17 | 0 | 1 | 1 |
| TENM1 | 10 | 10 | -6,20403E-17 | 0,999999976 | 1 |
| SFN;YWHAG;YWHA | 4 | 4 | -4,23427E-19 | 0,999999999 | 1 |
| PPP2R5C | 9 | 9 | 3,45303E-17 | 0,999999983 | 1 |
| BRAP | 3 | 3 | 0 | 1 | 1 |
| RPS6KC1 | 1 | 1 | -0,099745847 | 0,674929729 | 1 |
| CTU1 | 1 | 1 | -0,071712144 | 0,316035176 | 1 |
| IGKC | 7 | 5 | 5,17745E-18 | 0,999999999 | 1 |
| ABCC8 | 7 | 7 | -2,17757E-16 | 0,999999965 | 1 |
| MAPK8IP3 | 24 | 24 | 0 | 1 | 1 |
| PRKAB2 | 7 | 7 | -3,98015E-18 | 0,999999997 | 1 |
| WDR24 | 3 | 3 | 0 | 1 | 1 |
| TMEM106B | 6 | 6 | 7,49329E-18 | 0,999999994 | 1 |
| ACTR10 | 15 | 15 | -1,84403E-17 | 0,999999985 | 1 |
| MICOS13 | 4 | 4 | 0 | 1 | 1 |
| CIRBP | 3 | 3 | 0 | 1 | 1 |
| MAG | 3 | 3 | -9,62391E-17 | 0,99999999 | 1 |
| OXSR1 | 7 | 7 | -0,002445332 | 0,847494887 | 1 |
| TMEM38B | 1 | 1 | -1,29848E-14 | 0,999999804 | 1 |
| CBX5 | 2 | 2 | 6,3893E-19 | 0,999999999 | 1 |
| SEPTIN11;SEPTIN6 | 7 | 7 | -0,017037021 | 0,481214089 | 1 |
| CALML3 | 3 | 3 | 5,01803E-18 | 0,999999996 | 1 |

| | | | | | |
|---|---|---|---|---|---|
| TRIM9 | 3 | 3 | -7,72365E-20 | 1 | 1 |
| RAP1A | 4 | 4 | 0 | 1 | 1 |
| PPM1B | 3 | 3 | -9,67038E-16 | 0,99999994 | 1 |
| MARK2 | 3 | 3 | 0 | 1 | 1 |
| NT5C | 6 | 6 | 0,049512661 | 0,412629637 | 1 |
| RAB2A | 8 | 8 | -0,025337082 | 0,349076113 | 1 |
| MBP | 8 | 8 | -1,76934E-17 | 0,999999993 | 1 |
| RPS20 | 5 | 5 | -0,003445375 | 0,812620934 | 1 |
| PAK1 | 25 | 25 | -9,04052E-17 | 0,999999969 | 1 |
| DDOST | 11 | 11 | -5,98031E-17 | 0,999999984 | 1 |
| TENM1;TENM3 | 1 | 1 | -0,00117306 | 0,947128751 | 1 |
| GNA11;GNAQ | 7 | 7 | 2,31691E-17 | 0,999999987 | 1 |
| PALS2 | 1 | 1 | 0 | 1 | 1 |
| CETN1;CETN2 | 2 | 2 | -6,97702E-16 | 0,999999957 | 1 |
| GNAI1;GNAI2 | 4 | 4 | 6,79169E-17 | 0,999999983 | 1 |
| ANKRD49 | 1 | 1 | 0,105613061 | 0,283414849 | 1 |
| OLFML2B | 1 | 1 | -0,017492656 | 0,738362348 | 1 |
| BIN2 | 2 | 2 | 0 | 1 | 1 |
| YTHDF2 | 3 | 3 | -2,0384E-16 | 0,999999964 | 1 |
| NDUFS4 | 8 | 7 | 2,22344E-16 | 0,999999972 | 1 |
| SRPRB | 5 | 5 | 0,042647449 | 0,356769285 | 1 |
| NAA25 | 11 | 11 | -3,85381E-17 | 0,999999975 | 1 |
| UQCC2 | 4 | 4 | 2,94451E-17 | 0,999999995 | 1 |
| MRPL40 | 3 | 2 | 0 | 1 | 1 |
| EIF2B5 | 4 | 4 | 0 | 1 | 1 |
| PAOX | 10 | 10 | 8,78132E-17 | 0,999999975 | 1 |
| NDUFV3 | 7 | 7 | -2,61826E-18 | 0,999999996 | 1 |
| PLIN4 | 3 | 3 | -2,75059E-17 | 0,999999992 | 1 |
| DIRAS1 | 8 | 8 | 0 | 1 | 1 |
| GGPS1 | 6 | 6 | -5,13191E-16 | 0,999999919 | 1 |
| CAB39 | 16 | 16 | -0,00811752 | 0,666073074 | 1 |
| MYH11 | 4 | 4 | 1,9997E-18 | 0,999999998 | 1 |
| BFSP2 | 1 | 1 | -5,8486E-20 | 1 | 1 |

| | | | | | |
|---|---|---|---|---|---|
| CTBP1 | 12 | 12 | 7,80126E-18 | 0,999999995 | 1 |
| KRT6A;KRT72;KRT5 | 4 | 3 | -5,45837E-17 | 0,999999991 | 1 |
| KRT73 | 5 | 5 | 4,00592E-16 | 0,99999998 | 1 |
| SNX6 | 16 | 16 | 2,29793E-17 | 0,999999989 | 1 |
| ARG1 | 1 | 1 | 0 | 1 | 1 |
| CMTM4 | 2 | 2 | 1,01206E-15 | 0,999999953 | 1 |
| RADIL | 1 | 1 | 0 | 1 | 1 |
| MRPL49 | 3 | 2 | -8,66894E-16 | 0,999999998 | 1 |
| ZC3H15 | 4 | 4 | -0,037590921 | 0,428908597 | 1 |
| PIP4K2A;PIP4K2B | 4 | 4 | 1,02523E-16 | 0,999999971 | 1 |
| PDZD8 | 7 | 7 | 0 | 1 | 1 |
| RPL27 | 6 | 6 | 0 | 1 | 1 |
| ITPR2 | 1 | 1 | 2,51046E-17 | 0,999999989 | 1 |
| FXR1 | 9 | 9 | -3,23259E-17 | 0,999999986 | 1 |
| RPS12 | 3 | 3 | -1,10582E-11 | 0,999988733 | 1 |
| PLG | 6 | 2 | 0 | 1 | 1 |
| CSNK1D;CSNK1E | 10 | 10 | 0 | 1 | 1 |
| STK38L | 5 | 5 | 2,67008E-15 | 0,999999842 | 1 |
| ROCK1;ROCK2 | 8 | 8 | 7,47656E-18 | 0,999999995 | 1 |
| CSNK1G1;CSNK1G3 | 1 | 1 | 0 | 1 | 1 |
| CSNK2A1 | 18 | 18 | 0 | 1 | 1 |
| CSNK2A2 | 18 | 18 | 2,65855E-17 | 0,999999983 | 1 |
| MAP2K7 | 5 | 5 | -0,063455938 | 0,135476891 | 1 |
| NAA50 | 9 | 9 | -7,82847E-15 | 0,999999664 | 1 |
| MCEE | 5 | 5 | 1,80079E-16 | 0,999999971 | 1 |
| GMPR2;GMPR | 2 | 2 | 0 | 1 | 1 |
| PARK7 | 20 | 18 | 0 | 1 | 1 |
| CDH9 | 4 | 4 | -0,115463271 | 0,230089912 | 1 |
| MT-CYB | 1 | 1 | 0 | 1 | 1 |
| MAOA | 19 | 18 | 0 | 1 | 1 |
| SLC25A22 | 13 | 13 | 0,011606647 | 0,593194108 | 1 |
| QTRT1 | 2 | 2 | -2,39679E-19 | 0,999999999 | 1 |
| EPB41 | 2 | 2 | 3,6886E-16 | 0,999999969 | 1 |

| | | | | | |
|---|---|---|---|---|---|
| TMED8 | 4 | 4 | 2,51701E-16 | 0,999999965 | 1 |
| CFAP44 | 1 | 1 | 0,171831354 | 0,132842459 | 1 |
| CAPZA1 | 8 | 8 | 8,24604E-18 | 0,999999995 | 1 |
| SRP54 | 5 | 5 | -1,37402E-17 | 0,999999991 | 1 |
| CLCC1 | 5 | 5 | 0 | 1 | 1 |
| GDAP1L1 | 16 | 16 | 1,07907E-16 | 0,999999965 | 1 |
| PPP2R2C | 2 | 2 | 2,97674E-16 | 0,99999997 | 1 |
| MTHFSD | 1 | 1 | 0 | 1 | 1 |
| TRAPPC9 | 17 | 17 | 0,041676581 | 0,13672401 | 1 |
| MGAT4B | 1 | 1 | 0 | 1 | 1 |
| WASF2 | 2 | 2 | -2,86185E-19 | 1 | 1 |
| BCAS3 | 15 | 15 | 5,97848E-18 | 0,999999992 | 1 |
| FDX2 | 5 | 3 | 0,003904703 | 0,841526385 | 1 |
| NDUFAF6 | 1 | 1 | 0 | 1 | 1 |
| NTRK2;NTRK3 | 3 | 3 | 1,04779E-14 | 0,999999763 | 1 |
| DOCK7 | 8 | 8 | 0 | 1 | 1 |
| DUS3L | 4 | 4 | -4,55036E-12 | 0,99999462 | 1 |
| CALU | 10 | 10 | 1,2031E-17 | 0,999999996 | 1 |
| UNC5CL | 1 | 1 | 0 | 1 | 1 |
| PDE1A | 25 | 24 | -3,4313E-18 | 0,999999998 | 1 |
| PSMD6 | 20 | 20 | -0,045858225 | 0,147652827 | 1 |
| GDI1 | 49 | 47 | 4,38938E-15 | 0,999999854 | 1 |
| PEX5L | 2 | 2 | 0 | 1 | 1 |
| ACSL4 | 9 | 9 | 1,2766E-16 | 0,999999961 | 1 |
| CDH2 | 13 | 12 | 0 | 1 | 1 |
| GLRX5 | 3 | 3 | 5,82412E-20 | 1 | 1 |
| RAB22A | 5 | 5 | -4,23443E-18 | 0,999999995 | 1 |
| RAB35 | 14 | 14 | -1,54411E-17 | 0,999999992 | 1 |
| RALGAPB | 14 | 14 | 0 | 1 | 1 |
| TUBA1A | 2 | 2 | 0 | 1 | 1 |
| TRAPPC3 | 6 | 6 | 0 | 1 | 1 |
| HPX | 18 | 17 | -2,61403E-16 | 0,999999993 | 1 |
| DPP6 | 28 | 28 | 0,024198171 | 0,31956998 | 1 |

| | | | | | |
|---|---|---|---|---|---|
| PPTC7 | 5 | 4 | 0,087017588 | 0,168320961 | 1 |
| PRDX4 | 6 | 6 | 1,88017E-16 | 0,999999977 | 1 |
| MAPK8;MAPK9 | 1 | 1 | 0 | 1 | 1 |
| MTRES1 | 2 | 2 | -8,5633E-18 | 0,999999995 | 1 |
| B3GLCT | 2 | 2 | -1,14386E-15 | 0,999999911 | 1 |
| PURA | 16 | 16 | -0,010325166 | 0,593239434 | 1 |
| APMAP | 14 | 14 | 0 | 1 | 1 |
| EMD | 6 | 6 | 9,27888E-17 | 0,999999969 | 1 |
| MAPRE3 | 16 | 16 | -2,119E-15 | 0,999999834 | 1 |
| TECR | 11 | 11 | -0,000207291 | 0,944548328 | 1 |
| AGO3;AGO1 | 2 | 2 | -0,059565204 | 0,203057069 | 1 |
| GOLGA7B | 3 | 3 | -4,04629E-16 | 0,99999996 | 1 |
| TMCC1 | 7 | 5 | -1,67669E-18 | 0,999999999 | 1 |
| GRAMD1B | 5 | 4 | 4,90424E-18 | 0,999999996 | 1 |
| PTBP2 | 2 | 2 | 0 | 1 | 1 |
| MTPN | 10 | 10 | 2,30549E-14 | 0,999999865 | 1 |
| EML6 | 5 | 5 | 1,76284E-17 | 0,999999996 | 1 |
| SGSM1 | 1 | 1 | 0 | 1 | 1 |
| UBTD2 | 3 | 3 | 0 | 1 | 1 |
| PYGL | 4 | 4 | 1,36871E-16 | 0,999999973 | 1 |
| CAD | 18 | 18 | 0 | 1 | 1 |
| ATP6V1G1 | 3 | 3 | -6,44117E-15 | 0,999999778 | 1 |
| AGRN | 2 | 2 | 1,49748E-14 | 0,999999751 | 1 |
| RPL13 | 9 | 9 | -5,44953E-17 | 0,999999987 | 1 |
| PEX3 | 1 | 1 | 0 | 1 | 1 |
| APEX1 | 5 | 5 | -9,6518E-20 | 1 | 1 |
| KRIT1 | 2 | 2 | 0,100421241 | 0,231345725 | 1 |
| NYAP1 | 1 | 1 | 0 | 1 | 1 |
| VANGL2 | 1 | 1 | 0 | 1 | 1 |
| GSTP2 | 1 | 1 | 0 | 1 | 1 |
| NAP1L4 | 11 | 11 | -2,41446E-18 | 0,999999999 | 1 |
| KPNA6 | 4 | 4 | -3,38599E-17 | 0,999999991 | 1 |
| TRAPPC5 | 8 | 8 | 1,14388E-16 | 0,999999985 | 1 |

| | | | | | |
|---|---|---|---|---|---|
| EXOC4 | 25 | 25 | 0,003643851 | 0,698382037 | 1 |
| HEBP1 | 7 | 7 | -7,17514E-17 | 0,999999979 | 1 |
| PKM | 4 | 4 | -1,05396E-19 | 1 | 1 |
| CPLX1 | 5 | 5 | -0,023747134 | 0,617593442 | 1 |
| AKT1S1 | 2 | 2 | -5,91189E-16 | 0,999999949 | 1 |
| KCTD8 | 6 | 5 | -3,30923E-17 | 0,999999984 | 1 |
| SCN1A;SCN3A;SCN: | 7 | 7 | 0,060326727 | 0,139796244 | 1 |
| ANK2 | 3 | 3 | 2,88035E-15 | 0,999999863 | 1 |
| PNPO | 6 | 6 | 3,8388E-16 | 0,999999949 | 1 |
| KCTD16 | 12 | 12 | 0,045732297 | 0,151938123 | 1 |
| CETN2 | 2 | 2 | 4,10836E-18 | 0,999999999 | 1 |
| NUDT19 | 1 | 1 | 0 | 1 | 1 |
| ARVCF | 13 | 13 | 0 | 1 | 1 |
| BCL2L1 | 2 | 2 | -0,053422569 | 0,326956632 | 1 |
| PCDH10 | 6 | 5 | -5,21412E-17 | 0,999999986 | 1 |
| SPTAN1 | 5 | 5 | -2,56589E-16 | 0,999999977 | 1 |
| DBH | 1 | 1 | 2,90723E-20 | 1 | 1 |
| EEF1AKMT1 | 2 | 2 | 0 | 1 | 1 |
| NEB | 1 | 1 | 0 | 1 | 1 |
| DNPEP | 16 | 15 | 0,000885581 | 0,855029809 | 1 |
| STXBP5L | 25 | 24 | 0,027297618 | 0,199372688 | 1 |
| EPB41;EPB41L3 | 4 | 4 | 0 | 1 | 1 |
| FAM228A | 1 | 1 | -0,076130346 | 0,5034118 | 1 |
| SLC10A4 | 2 | 2 | 0 | 1 | 1 |
| GSTO1 | 16 | 16 | 0 | 1 | 1 |
| IMPA1 | 15 | 15 | 0 | 1 | 1 |
| DOHH | 1 | 1 | 0 | 1 | 1 |
| GULO | 1 | 1 | 0 | 1 | 1 |
| FRMPD3 | 7 | 6 | 0,032036027 | 0,508401796 | 1 |
| ALG5 | 1 | 1 | -0,071432411 | 0,368680955 | 1 |
| PLSCR3 | 3 | 3 | 0 | 1 | 1 |
| IPO9 | 18 | 18 | -3,42756E-15 | 0,999999788 | 1 |
| KLHL26 | 4 | 4 | 0,069776706 | 0,303163162 | 1 |

| | | | | | |
|---|---|---|---|---|---|
| PPAT | 8 | 8 | -3,74247E-16 | 0,999999945 | 1 |
| DKK3 | 5 | 5 | -3,80587E-13 | 0,999998316 | 1 |
| ACHE | 6 | 6 | 2,08925E-17 | 0,999999989 | 1 |
| TUBGCP4 | 2 | 2 | 5,26501E-17 | 0,999999983 | 1 |
| LASP1 | 18 | 17 | -5,91141E-14 | 0,999999041 | 1 |
| CEP104 | 1 | 1 | 0 | 1 | 1 |
| EARS2 | 10 | 9 | 1,28801E-14 | 0,999999817 | 1 |
| STOM | 4 | 4 | 1,53896E-16 | 0,999999975 | 1 |
| SLC49A4 | 1 | 1 | 0 | 1 | 1 |
| GLDC | 15 | 13 | -0,025868784 | 0,353568829 | 1 |
| CLINT1 | 12 | 12 | -0,086128327 | 0,180067397 | 1 |
| RAB11FIP3 | 1 | 1 | 0 | 1 | 1 |
| VPS4B;VPS4A | 2 | 2 | 1,75012E-18 | 0,999999998 | 1 |
| SLC44A1 | 9 | 9 | -1,08162E-16 | 0,999999987 | 1 |
| ETHE1 | 5 | 5 | 0,049922951 | 0,283615315 | 1 |
| SFXN5 | 12 | 11 | 0,023842855 | 0,292835943 | 1 |
| TSN | 15 | 14 | -0,04340814 | 0,465197379 | 1 |
| CDC42BPA | 15 | 15 | 0,036686257 | 0,199698872 | 1 |
| PITRM1 | 32 | 27 | 0 | 1 | 1 |
| STT3B | 8 | 7 | 1,23792E-16 | 0,999999973 | 1 |
| PSD | 11 | 11 | 0,063812393 | 0,212250736 | 1 |
| STT3A | 11 | 11 | 4,95966E-19 | 1 | 1 |
| INPP1 | 17 | 17 | 0 | 1 | 1 |
| MOB4 | 8 | 8 | -0,011176699 | 0,603326976 | 1 |
| MBOAT7 | 3 | 3 | 0 | 1 | 1 |
| CHGB | 3 | 3 | -6,54308E-19 | 0,999999999 | 1 |
| GPM6B | 1 | 1 | -2,84631E-19 | 0,999999999 | 1 |
| UBR4 | 8 | 8 | -3,36179E-16 | 0,999999948 | 1 |
| ACP1 | 4 | 4 | 2,1681E-18 | 0,999999998 | 1 |
| CR1L | 5 | 4 | 0,091313723 | 0,294537699 | 1 |
| TTC39C | 7 | 7 | 0 | 1 | 1 |
| TMX4 | 6 | 6 | -4,3297E-16 | 0,999999926 | 1 |
| RABGGTB | 5 | 5 | -1,39538E-18 | 1 | 1 |

| | | | | | |
|---|---|---|---|---|---|
| NARS1 | 26 | 24 | 0 | 1 | 1 |
| DARS2 | 10 | 10 | 0,049051045 | 0,198267709 | 1 |
| RWDD2B | 1 | 1 | 0 | 1 | 1 |
| NUDCD1 | 7 | 7 | 0 | 1 | 1 |
| ILF3 | 8 | 8 | -0,010772992 | 0,652154957 | 1 |
| HAX1 | 5 | 3 | 0 | 1 | 1 |
| LARS1 | 39 | 39 | 0 | 1 | 1 |
| RALGPS1 | 1 | 1 | 0 | 1 | 1 |
| GPR37 | 2 | 2 | 0 | 1 | 1 |
| SLITRK5 | 6 | 6 | -2,43304E-18 | 0,999999999 | 1 |
| RAB23 | 10 | 10 | -0,016755695 | 0,469686789 | 1 |
| EIF5A | 13 | 13 | -7,5638E-17 | 0,999999989 | 1 |
| RGS12 | 8 | 7 | -1,05638E-18 | 0,999999998 | 1 |
| GNG2 | 2 | 2 | -2,80783E-18 | 0,999999997 | 1 |
| CNKSR2 | 2 | 2 | 1,62024E-18 | 1 | 1 |
| HNRNPA1 | 12 | 12 | -0,028395303 | 0,369611002 | 1 |
| SERPINB9 | 7 | 7 | 0 | 1 | 1 |
| GPT2 | 6 | 6 | 0 | 1 | 1 |
| C2CD3 | 1 | 1 | 0 | 1 | 1 |
| FRYL | 3 | 3 | 0 | 1 | 1 |
| FREM1 | 1 | 1 | -0,175467199 | 0,147948248 | 1 |
| MTDH | 15 | 15 | -1,8884E-16 | 0,999999958 | 1 |
| PIN1 | 6 | 6 | -1,51185E-17 | 0,999999996 | 1 |
| TMUB1 | 2 | 2 | 0 | 1 | 1 |
| ATP6V1G2 | 12 | 11 | 2,51032E-16 | 0,999999944 | 1 |
| CD82 | 5 | 5 | -4,17228E-18 | 0,999999999 | 1 |
| WASHC1 | 4 | 4 | 0 | 1 | 1 |
| ZNF445 | 1 | 1 | 0,241565133 | 0,177441023 | 1 |
| UQCRH | 4 | 4 | 0,048883633 | 0,399501868 | 1 |
| SLITRK3 | 4 | 3 | 0 | 1 | 1 |
| UBN1 | 1 | 1 | 0,01370653 | 0,735658041 | 1 |
| FAM162A | 9 | 9 | 0,011611547 | 0,593707133 | 1 |
| KCNA2 | 5 | 5 | 5,82355E-18 | 0,999999995 | 1 |

| | | | | | |
|---|---|---|---|---|---|
| UBQLN2 | 17 | 15 | -0,013029541 | 0,705845747 | 1 |
| TUBB4A | 14 | 14 | -5,11611E-17 | 0,999999986 | 1 |
| RAB2A;RAB2B | 9 | 9 | -7,4355E-17 | 0,999999983 | 1 |
| MTCH2 | 12 | 12 | 0,04350449 | 0,178353276 | 1 |
| PNPLA6 | 2 | 2 | 0 | 1 | 1 |
| SLC6A7 | 13 | 13 | 2,3091E-17 | 0,99999999 | 1 |
| ALDH1A2 | 5 | 5 | 9,2578E-15 | 0,999999891 | 1 |
| NDRG1 | 9 | 9 | 1,24522E-17 | 0,999999993 | 1 |
| MARK4 | 7 | 7 | 0,022016515 | 0,548609222 | 1 |
| DVL2 | 2 | 2 | 0 | 1 | 1 |
| MED7 | 1 | 1 | -0,210947205 | 0,16999931 | 1 |
| RABGAP1L | 3 | 3 | 1,29919E-18 | 0,999999999 | 1 |
| MLC1 | 6 | 6 | 0 | 1 | 1 |
| PCDH9 | 14 | 14 | -0,028060453 | 0,420134602 | 1 |
| COMMD8 | 2 | 2 | 0 | 1 | 1 |
| FKBPL | 1 | 1 | 0 | 1 | 1 |
| NDUFAF2 | 10 | 10 | -1,48909E-17 | 0,999999995 | 1 |
| GPM6A | 7 | 7 | -4,37771E-17 | 0,999999978 | 1 |
| TYW5 | 1 | 1 | 0 | 1 | 1 |
| FIS1 | 7 | 7 | 0 | 1 | 1 |
| TMEM151B | 1 | 1 | 0 | 1 | 1 |
| PYCARD | 1 | 1 | 0 | 1 | 1 |
| NES | 1 | 1 | 0 | 1 | 1 |
| SPOPL | 1 | 1 | 0,143335086 | 0,333476867 | 1 |
| ATP6V1B2;ATP6V1 | 8 | 8 | 0,030417525 | 0,241317864 | 1 |
| TUBB4B;TUBB5;TU | 1 | 1 | 7,05858E-20 | 1 | 1 |
| MIEN1 | 2 | 2 | -1,72692E-18 | 0,999999998 | 1 |
| CFAP95 | 1 | 1 | -0,056486553 | 0,538464359 | 1 |
| LSM14A | 3 | 3 | -0,026601868 | 0,512488402 | 1 |
| RAB1A | 14 | 14 | 1,77749E-18 | 0,999999999 | 1 |
| BRINP1 | 15 | 14 | 1,43414E-17 | 0,999999988 | 1 |
| KCND3 | 7 | 7 | 0 | 1 | 1 |
| TTL | 7 | 7 | -2,26069E-17 | 0,99999999 | 1 |

| | | | | | |
|---|---|---|---|---|---|
| NTNG2 | 3 | 3 | 4,39222E-16 | 0,999999945 | 1 |
| DDAH2;DDAH1 | 1 | 1 | -2,11277E-18 | 0,999999999 | 1 |
| PRXL2A | 4 | 4 | 0 | 1 | 1 |
| STX7 | 9 | 9 | 2,51708E-17 | 0,999999983 | 1 |
| CPEB4;CPEB3;CPEB | 3 | 3 | -0,090176302 | 0,211781052 | 1 |
| SET | 9 | 9 | 0 | 1 | 1 |
| HPCAL1;NCALD | 6 | 6 | -1,10249E-16 | 0,999999979 | 1 |
| BEST3 | 1 | 1 | 0 | 1 | 1 |
| GBE1 | 15 | 15 | -6,1129E-17 | 0,999999979 | 1 |
| UBE2K | 8 | 8 | 2,58916E-17 | 0,999999985 | 1 |
| UBL3 | 2 | 2 | 0,020003579 | 0,58236605 | 1 |
| RPL5 | 18 | 16 | 1,01414E-17 | 0,999999996 | 1 |
| EXD2 | 6 | 4 | 0 | 1 | 1 |
| MAGEE2 | 3 | 3 | -3,7452E-19 | 0,999999999 | 1 |
| GNA11 | 12 | 12 | -1,72111E-17 | 0,999999988 | 1 |
| TRPV2 | 6 | 6 | 0 | 1 | 1 |
| HSCB | 7 | 6 | 0 | 1 | 1 |
| UCHL1 | 17 | 17 | 0 | 1 | 1 |
| PTPDC1 | 2 | 2 | 0,055544279 | 0,50123991 | 1 |
| MAIP1 | 5 | 5 | 1,18652E-17 | 0,999999991 | 1 |
| NFASC | 4 | 4 | -0,049753262 | 0,308013937 | 1 |
| FBXL16 | 15 | 15 | -2,12364E-16 | 0,999999955 | 1 |
| ATP1B2 | 14 | 14 | -0,023343341 | 0,339788447 | 1 |
| MAG | 11 | 11 | 1,68319E-16 | 0,999999972 | 1 |
| PYCR2 | 15 | 15 | 0,069210081 | 0,116499637 | 1 |
| PPT1 | 7 | 7 | -1,57884E-17 | 0,999999991 | 1 |
| ATP8A1 | 38 | 37 | 0,029977212 | 0,231374211 | 1 |
| SEZ6L2 | 12 | 12 | 0 | 1 | 1 |
| TMEM132B | 16 | 16 | 0 | 1 | 1 |
| RPS28 | 6 | 6 | 9,39385E-17 | 0,999999979 | 1 |
| RANBP6;IPO5 | 2 | 2 | -4,59415E-17 | 0,999999985 | 1 |
| SLC25A16 | 3 | 3 | 0 | 1 | 1 |
| PRDX2 | 10 | 10 | -6,17825E-18 | 0,999999998 | 1 |

| | | | | | |
|---|---|---|---|---|---|
| HERC4 | 8 | 8 | 8,57339E-18 | 0,999999994 | 1 |
| ASPM | 1 | 1 | 0 | 1 | 1 |
| ATG13 | 1 | 1 | 0 | 1 | 1 |
| SLC7A10 | 1 | 1 | 0 | 1 | 1 |
| SLC7A8 | 5 | 5 | 0 | 1 | 1 |
| FBXO21 | 7 | 7 | 0 | 1 | 1 |
| FAIM | 4 | 4 | 0 | 1 | 1 |
| UBB;UBC;RPS27A;U | 8 | 8 | 0,05173285 | 0,142707953 | 1 |
| NEDD8 | 4 | 4 | 2,34373E-12 | 0,999995814 | 1 |
| MTERF2 | 3 | 2 | -5,78026E-16 | 0,999999941 | 1 |
| CLCN3 | 2 | 2 | -2,32149E-18 | 0,999999998 | 1 |
| CCDC51 | 8 | 6 | 1,12684E-16 | 0,999999973 | 1 |
| CTNNB1;JUP | 2 | 2 | 1,65914E-17 | 0,999999993 | 1 |
| ERI3 | 2 | 2 | 5,01478E-18 | 0,999999997 | 1 |
| CLASP1;CLASP2 | 4 | 4 | 4,82306E-18 | 0,999999997 | 1 |
| TSPAN9 | 1 | 1 | 0,139276106 | 0,170740403 | 1 |
| PPIA | 19 | 19 | -3,60074E-16 | 0,999999976 | 1 |
| ARAF | 4 | 4 | 0 | 1 | 1 |
| SQOR | 8 | 7 | -1,33163E-18 | 0,999999998 | 1 |
| MYL1;MYL3 | 2 | 2 | 1,01573E-17 | 0,999999998 | 1 |
| GRIP2 | 3 | 3 | 0,086227747 | 0,258392336 | 1 |
| LIMCH1 | 8 | 8 | -5,84169E-14 | 0,99999952 | 1 |
| STK39 | 7 | 7 | -0,052385554 | 0,342403889 | 1 |
| MCTS1 | 5 | 5 | -2,20487E-17 | 0,999999993 | 1 |
| SLC45A1 | 1 | 1 | 0 | 1 | 1 |
| EMC6 | 1 | 1 | 0 | 1 | 1 |
| PPP2R5C | 7 | 7 | 8,05453E-15 | 0,999999738 | 1 |
| OPCML | 2 | 2 | 4,64729E-15 | 0,99999984 | 1 |
| SMG9 | 1 | 1 | 0 | 1 | 1 |
| MSRA | 10 | 10 | 5,19095E-18 | 0,999999994 | 1 |
| LSM12 | 3 | 3 | -1,25617E-18 | 0,999999998 | 1 |
| KCNJ10 | 5 | 5 | 3,7426E-16 | 0,999999957 | 1 |
| CNOT2 | 2 | 2 | -0,106348346 | 0,271730975 | 1 |

| | | | | | |
|---|---|---|---|---|---|
| TIMM17B | 2 | 2 | 0,074983819 | 0,211077829 | 1 |
| HACD3 | 8 | 8 | -1,48063E-17 | 0,99999999 | 1 |
| DBN1 | 1 | 1 | 0 | 1 | 1 |
| MT-CO3 | 1 | 1 | 0 | 1 | 1 |
| SASH1 | 3 | 3 | -0,125153345 | 0,208266631 | 1 |
| SNCB | 6 | 6 | -0,165321762 | 0,212176153 | 1 |
| RHOC | 1 | 1 | -0,042590427 | 0,640047698 | 1 |
| ARFIP1 | 8 | 8 | 4,04346E-17 | 0,999999992 | 1 |
| PYM1 | 3 | 3 | -3,0022E-14 | 0,999999522 | 1 |
| EEF1A1;EEF1A2 | 14 | 13 | -0,024507633 | 0,41915317 | 1 |
| CFAP300 | 1 | 1 | 0 | 1 | 1 |
| UBXN2B | 4 | 4 | -0,11698253 | 0,143043106 | 1 |
| GOPC | 4 | 4 | 0 | 1 | 1 |
| SPOPL | 1 | 1 | 0 | 1 | 1 |
| NMT2 | 11 | 11 | 0 | 1 | 1 |
| YWHAG;YWHAH | 3 | 3 | -1,42818E-15 | 0,999999928 | 1 |
| EHD1 | 20 | 19 | -2,17919E-16 | 0,99999997 | 1 |
| MRPL48 | 1 | 1 | 0 | 1 | 1 |
| RPL10 | 9 | 9 | 2,27909E-17 | 0,999999994 | 1 |
| AGAP3 | 11 | 11 | -1,90592E-16 | 0,999999951 | 1 |
| HIP1 | 7 | 7 | -2,7369E-16 | 0,999999964 | 1 |
| REEP2 | 4 | 4 | 0 | 1 | 1 |
| CRYL1 | 14 | 14 | 1,82254E-17 | 0,999999993 | 1 |
| DYNLRB1 | 4 | 4 | 1,25202E-17 | 0,999999993 | 1 |
| RNF213 | 1 | 1 | 0 | 1 | 1 |
| UVRAG | 4 | 3 | -4,28583E-17 | 0,999999982 | 1 |
| RAP2B;RAP2A;RAP: | 8 | 8 | 3,5293E-16 | 0,99999994 | 1 |
| EML4 | 9 | 9 | 0 | 1 | 1 |
| GFPT1 | 5 | 5 | -2,34384E-18 | 0,999999999 | 1 |
| SAR1A | 5 | 5 | 0 | 1 | 1 |
| DENND10 | 7 | 7 | 0,033511477 | 0,270795744 | 1 |
| ABI2;ABI1 | 9 | 8 | -7,81769E-16 | 0,999999915 | 1 |
| PPFIA3;PPFIA2 | 1 | 1 | 0 | 1 | 1 |

| | | | | | |
|---|---|---|---|---|---|
| APOL10A | 1 | 1 | 0 | 1 | 1 |
| KIFBP | 12 | 12 | -0,004659442 | 0,738569368 | 1 |
| USP32 | 18 | 18 | 0,05006887 | 0,126935125 | 1 |
| NMD3 | 1 | 1 | 0 | 1 | 1 |
| MYOZ2 | 1 | 1 | 0 | 1 | 1 |
| ARFGEF1 | 14 | 13 | 0 | 1 | 1 |
| GALNT2 | 2 | 2 | -1,31996E-15 | 0,999999929 | 1 |
| MAP9 | 1 | 1 | 0 | 1 | 1 |
| INMT | 3 | 3 | -2,18956E-16 | 0,999999976 | 1 |
| RPL12 | 6 | 6 | -0,03150288 | 0,336579413 | 1 |
| MAOB | 20 | 18 | 0 | 1 | 1 |
| ACSL3 | 14 | 13 | -0,056177358 | 0,112420514 | 1 |
| SETD3 | 3 | 3 | -1,97049E-18 | 0,999999998 | 1 |
| DTNB;DTNA | 1 | 1 | 0 | 1 | 1 |
| PLPP3 | 9 | 8 | -2,86436E-18 | 0,999999996 | 1 |
| MTO1 | 7 | 7 | -4,84922E-18 | 0,999999995 | 1 |
| RAP1GAP;RAP1GAI | 2 | 2 | -1,29026E-17 | 0,999999995 | 1 |
| CTSB | 12 | 12 | 7,91954E-17 | 0,999999977 | 1 |
| LRRC4C | 5 | 5 | 0,041019091 | 0,333829734 | 1 |
| PLXDC2 | 4 | 4 | -7,27219E-19 | 0,999999998 | 1 |
| SHMT2 | 15 | 14 | 0,009031494 | 0,672022131 | 1 |
| HSD17B10 | 12 | 12 | 3,94399E-18 | 0,999999999 | 1 |
| RDH11 | 6 | 6 | 0,008011452 | 0,670701507 | 1 |
| OLFR332 | 1 | 1 | 0 | 1 | 1 |
| PRKCA | 18 | 18 | 5,91706E-17 | 0,999999984 | 1 |
| ETNPPL | 3 | 3 | -0,144475817 | 0,207132354 | 1 |
| EGFR | 7 | 6 | -0,004559929 | 0,833198013 | 1 |
| B4GALNT1 | 2 | 2 | 0 | 1 | 1 |
| VPS26A | 12 | 12 | 0 | 1 | 1 |
| HAP1 | 3 | 3 | 5,11447E-19 | 0,999999999 | 1 |
| D7H11ORF16 | 1 | 1 | 0 | 1 | 1 |
| AASS | 3 | 3 | 0 | 1 | 1 |
| HSPA13 | 5 | 5 | 2,09058E-19 | 0,999999999 | 1 |

| | | | | | |
|---|---|---|---|---|---|
| INPP5A | 12 | 11 | 0 | 1 | 1 |
| MAP7 | 8 | 8 | -1,55377E-14 | 0,999999631 | 1 |
| MRPS27 | 11 | 9 | -7,81631E-19 | 1 | 1 |
| PRKCSH | 11 | 11 | -1,14336E-17 | 0,999999991 | 1 |
| TIGAR | 5 | 5 | -2,35611E-16 | 0,999999957 | 1 |
| PXMP4 | 1 | 1 | 0 | 1 | 1 |
| TTC21B | 1 | 1 | 0 | 1 | 1 |
| NDUFA11 | 5 | 5 | 0,019841643 | 0,538901647 | 1 |
| ALDH3B1 | 6 | 6 | 9,17375E-18 | 0,999999996 | 1 |
| MAGI2 | 24 | 23 | 9,3933E-14 | 0,999998519 | 1 |
| CFDP1 | 4 | 4 | -0,002892772 | 0,812136624 | 1 |
| CACYBP | 12 | 11 | -0,045734961 | 0,265127583 | 1 |
| MTX1 | 3 | 3 | -1,30263E-16 | 0,999999986 | 1 |
| IMPDH2 | 14 | 14 | 0 | 1 | 1 |
| CHRM1 | 1 | 1 | 0 | 1 | 1 |
| ATP5ME | 8 | 8 | 0,059613419 | 0,138929534 | 1 |
| UBE2E2 | 2 | 2 | 0 | 1 | 1 |
| KCNJ6 | 3 | 3 | -0,003970358 | 0,825553658 | 1 |
| COPS7A | 9 | 9 | 4,10023E-18 | 0,999999997 | 1 |
| NECTIN4 | 1 | 1 | 0 | 1 | 1 |
| LAMTOR3 | 2 | 2 | -2,41958E-18 | 0,999999998 | 1 |
| ABHD17C | 1 | 1 | 0 | 1 | 1 |
| RUFY3 | 4 | 4 | 0 | 1 | 1 |
| SERPINA1B | 5 | 5 | -1,42742E-18 | 0,999999998 | 1 |
| PPP2CA | 2 | 2 | 0,076455164 | 0,217351079 | 1 |
| PPP2CB | 2 | 2 | 1,92575E-12 | 0,999997308 | 1 |
| NENF | 2 | 2 | 1,20867E-14 | 0,999999783 | 1 |
| MAP2K2 | 5 | 5 | -8,06142E-15 | 0,999999795 | 1 |
| RAB11FIP2;RAB11F | 1 | 1 | 0 | 1 | 1 |
| ABHD6 | 17 | 15 | 0 | 1 | 1 |
| CLEC2L | 1 | 1 | 0 | 1 | 1 |
| LRRC57 | 12 | 12 | -9,57595E-17 | 0,999999974 | 1 |
| ARHGEF7 | 24 | 23 | 0 | 1 | 1 |

| | | | | | |
|---|---|---|---|---|---|
| ELAVL2;ELAVL4 | 3 | 3 | -0,034832513 | 0,34290806 | 1 |
| SYNRG | 17 | 17 | -0,074651067 | 0,329655615 | 1 |
| NIPBL | 1 | 1 | 0 | 1 | 1 |
| PMM1 | 9 | 9 | 1,17212E-15 | 0,999999862 | 1 |
| CA14 | 3 | 3 | -5,99641E-18 | 0,999999999 | 1 |
| METTL7A1 | 2 | 2 | 1,08921E-17 | 0,999999998 | 1 |
| PAFAH1B2 | 5 | 5 | 0 | 1 | 1 |
| FTH1 | 18 | 17 | -9,73061E-18 | 0,999999999 | 1 |
| FABP5 | 11 | 11 | -1,07783E-16 | 0,999999985 | 1 |
| UBLCP1 | 4 | 4 | 8,82949E-16 | 0,999999937 | 1 |
| UFSP2 | 7 | 7 | -1,75727E-18 | 0,999999999 | 1 |
| KCTD17 | 3 | 3 | -1,50818E-18 | 0,999999999 | 1 |
| TFAM | 13 | 13 | 0,054305108 | 0,131315455 | 1 |
| SH3GLB1 | 11 | 11 | 2,53308E-14 | 0,999999602 | 1 |
| SLC25A20 | 3 | 3 | 9,73661E-18 | 0,999999996 | 1 |
| HSP90B1;HSP90AB | 2 | 2 | -1,83414E-16 | 0,999999981 | 1 |
| CEP131 | 1 | 1 | -0,045255737 | 0,544921461 | 1 |
| RIMOC1 | 4 | 4 | -0,016858831 | 0,564858656 | 1 |
| UBE2W | 1 | 1 | -0,051740899 | 0,563241729 | 1 |
| ARR3 | 1 | 1 | -0,147998976 | 0,307390182 | 1 |
| OGFOD1 | 5 | 5 | 1,40326E-16 | 0,999999981 | 1 |
| RPS9 | 16 | 16 | -3,82725E-18 | 0,999999995 | 1 |
| PFDN5 | 7 | 6 | 0 | 1 | 1 |
| PEX5L | 3 | 3 | 0 | 1 | 1 |
| STAU1 | 3 | 3 | -4,29067E-16 | 0,999999974 | 1 |
| ATP6V1H | 37 | 37 | 0 | 1 | 1 |
| OXSR1;STK39 | 1 | 1 | 0 | 1 | 1 |
| UBE2D2 | 2 | 2 | -1,00818E-16 | 0,999999975 | 1 |
| PDE6D | 6 | 5 | 0 | 1 | 1 |
| SCCPDH | 13 | 13 | 0,05469564 | 0,127359455 | 1 |
| GNB4 | 5 | 5 | -1,20997E-18 | 0,999999998 | 1 |
| LEMD2 | 1 | 1 | 0 | 1 | 1 |
| RANBP9 | 8 | 8 | -0,008605815 | 0,617673802 | 1 |

| | | | | | |
|---|---|---|---|---|---|
| SKI | 1 | 1 | 0,017985224 | 0,780467218 | 1 |
| SCAMP2 | 1 | 1 | 0 | 1 | 1 |
| ARL2 | 7 | 7 | -8,93122E-15 | 0,999999677 | 1 |
| DPM3 | 1 | 1 | 0,114509773 | 0,329187032 | 1 |
| PPP1R1A | 2 | 2 | 0 | 1 | 1 |
| SYT5 | 8 | 8 | 0,026619245 | 0,439127215 | 1 |
| MRPL43 | 2 | 2 | 0 | 1 | 1 |
| NDUFB1 | 2 | 2 | 1,66016E-15 | 0,999999902 | 1 |
| STAT1 | 8 | 8 | -6,58633E-16 | 0,999999956 | 1 |
| MVB12B | 5 | 5 | 0 | 1 | 1 |
| sp\|Q8CCC3\|CL056_ | 1 | 1 | -0,12635428 | 0,220734343 | 1 |
| APOD | 4 | 4 | 2,46207E-18 | 0,999999998 | 1 |
| RWDD4 | 2 | 2 | -2,85792E-18 | 0,999999996 | 1 |
| TEX264 | 2 | 2 | -6,36689E-20 | 1 | 1 |
| RAB6B | 17 | 16 | -0,011085204 | 0,592999052 | 1 |
| SLC4A10;SLC4A7;Sl | 1 | 1 | 0,003051408 | 0,882783075 | 1 |
| NT5C3B | 8 | 8 | 5,34414E-18 | 0,999999997 | 1 |
| GPCPD1 | 16 | 16 | -3,37545E-18 | 0,999999995 | 1 |
| PDE4A | 8 | 8 | 0 | 1 | 1 |
| MYH13 | 2 | 2 | 6,32138E-17 | 0,999999988 | 1 |
| LRRC73 | 2 | 2 | 1,01961E-12 | 0,999997839 | 1 |
| FBXO2 | 8 | 7 | -5,80255E-17 | 0,999999981 | 1 |
| YIF1B | 2 | 2 | -4,88875E-18 | 0,999999998 | 1 |
| RNF14 | 5 | 5 | 7,78984E-17 | 0,999999983 | 1 |
| MTIF3 | 1 | 1 | 0,230206377 | 0,268141243 | 1 |
| ACADS | 7 | 7 | -1,95289E-16 | 0,999999971 | 1 |
| MYO7A | 2 | 2 | 0 | 1 | 1 |
| ZNF106 | 1 | 1 | 0 | 1 | 1 |
| HDDC2 | 6 | 6 | 0 | 1 | 1 |
| TAMM41 | 4 | 3 | 0,048866676 | 0,435865843 | 1 |
| WDR20 | 4 | 4 | 0 | 1 | 1 |
| TRAPPC2L | 5 | 5 | 0,010666893 | 0,617459537 | 1 |
| CCZ1 | 7 | 7 | 0 | 1 | 1 |

| | | | | | |
|---|---|---|---|---|---|
| FAM98B | 6 | 6 | -3,88106E-16 | 0,999999956 | 1 |
| REXO1 | 1 | 1 | 0 | 1 | 1 |
| CMPK1 | 18 | 18 | 7,61566E-18 | 0,999999995 | 1 |
| ALPK2 | 1 | 1 | 0 | 1 | 1 |
| LYSMD3 | 1 | 1 | 3,31926E-19 | 0,999999999 | 1 |
| SYAP1 | 8 | 8 | -9,01781E-17 | 0,999999984 | 1 |
| SAR1B | 8 | 8 | 2,83523E-17 | 0,999999993 | 1 |
| RPL31 | 5 | 5 | 0 | 1 | 1 |
| COX15 | 4 | 4 | -4,50444E-15 | 0,999999945 | 1 |
| BPNT2 | 6 | 6 | 0 | 1 | 1 |
| H60C | 1 | 1 | 0 | 1 | 1 |
| DSG1A;DSG1B | 8 | 2 | -2,41775E-16 | 0,999999987 | 1 |
| GAS7 | 15 | 15 | -1,90259E-16 | 0,99999995 | 1 |
| RBBP7 | 3 | 2 | -1,74018E-19 | 1 | 1 |
| PHAF1 | 5 | 4 | -0,026900458 | 0,463154358 | 1 |
| EIF3K | 4 | 4 | -2,11058E-06 | 0,995577769 | 1 |
| PMFBP1 | 1 | 1 | 0 | 1 | 1 |
| ITGAM | 10 | 10 | 0,036993429 | 0,429120673 | 1 |
| TESC | 8 | 8 | 2,00685E-15 | 0,99999987 | 1 |
| CTSL | 1 | 1 | 0 | 1 | 1 |
| MUP2 | 2 | 2 | 5,14191E-18 | 0,999999998 | 1 |
| PRPS1 | 6 | 6 | 3,29925E-16 | 0,999999971 | 1 |
| NDUFA13 | 20 | 18 | 5,25501E-06 | 0,991467454 | 1 |
| GSPT1 | 11 | 10 | -1,53686E-16 | 0,999999966 | 1 |
| STAT5A;STAT5B | 1 | 1 | 0 | 1 | 1 |
| UBE2V1 | 5 | 5 | -2,33399E-17 | 0,999999989 | 1 |
| UBE2V2 | 7 | 6 | 0 | 1 | 1 |
| KCTD4 | 6 | 6 | -0,027166811 | 0,4396848 | 1 |
| SKP1 | 14 | 14 | -0,005457583 | 0,699892609 | 1 |
| SLC4A10;SLC4A7 | 1 | 1 | -0,096788525 | 0,170337888 | 1 |
| FLAD1 | 11 | 11 | 0 | 1 | 1 |
| CRLF3 | 3 | 3 | 0 | 1 | 1 |
| CLPB | 15 | 14 | 3,06933E-16 | 0,99999995 | 1 |

| | | | | | |
|---|---|---|---|---|---|
| ARCN1 | 17 | 17 | -0,041566888 | 0,177357497 | 1 |
| CRYZL1 | 7 | 7 | -6,99353E-17 | 0,999999975 | 1 |
| UBA6 | 24 | 22 | 1,95595E-18 | 0,999999998 | 1 |
| ZFP64 | 1 | 1 | 0 | 1 | 1 |
| NDUFS8 | 17 | 17 | 0,026587149 | 0,35901828 | 1 |
| SLITRK2 | 2 | 2 | -0,100755289 | 0,372118737 | 1 |
| ZAR1 | 1 | 1 | 0 | 1 | 1 |
| NDUFV3 | 5 | 4 | 0 | 1 | 1 |
| UFD1 | 7 | 7 | 0 | 1 | 1 |
| B4GAT1 | 3 | 3 | 0 | 1 | 1 |
| RBM39 | 2 | 2 | -0,063622673 | 0,368191594 | 1 |
| RNF25 | 4 | 4 | 0 | 1 | 1 |
| OCIAD1 | 1 | 1 | 0 | 1 | 1 |
| FAM53C | 1 | 1 | 0 | 1 | 1 |
| GSAP | 1 | 1 | 0 | 1 | 1 |
| DPP6 | 5 | 5 | -0,063779156 | 0,2255305 | 1 |
| CTU2 | 5 | 5 | -0,023733264 | 0,504949747 | 1 |
| PI4KB | 2 | 2 | 1,65733E-18 | 0,999999999 | 1 |
| MRPS22 | 8 | 8 | 0 | 1 | 1 |
| CYB5A | 4 | 4 | 0 | 1 | 1 |
| SLC7A14 | 13 | 13 | 0,010610334 | 0,628370332 | 1 |
| OTUD7B;OTUD7A | 2 | 2 | 0 | 1 | 1 |
| GBP9 | 1 | 1 | 0 | 1 | 1 |
| HSPBP1 | 6 | 6 | -0,053253324 | 0,115799361 | 1 |
| STX5 | 2 | 2 | 7,21571E-17 | 0,999999995 | 1 |
| LARP1 | 14 | 14 | 0 | 1 | 1 |
| RNF170 | 3 | 3 | -0,026296834 | 0,543149506 | 1 |
| MEAK7 | 2 | 2 | 0 | 1 | 1 |
| SLC25A31 | 3 | 1 | 0 | 1 | 1 |
| TXLNA | 1 | 1 | 0 | 1 | 1 |
| MAP1LC3B | 3 | 3 | -3,17372E-17 | 0,999999994 | 1 |
| ZKSCAN16 | 1 | 1 | -0,325834698 | 0,179128278 | 1 |
| FCSK | 7 | 7 | 0,005146242 | 0,707746311 | 1 |

| | | | | | |
|---|---|---|---|---|---|
| RAB33A | 3 | 3 | 0 | 1 | 1 |
| RPL17 | 10 | 10 | 6,95386E-16 | 0,999999942 | 1 |
| TBK1 | 9 | 9 | 0,04625856 | 0,227420102 | 1 |
| FAM50A | 1 | 1 | 0 | 1 | 1 |
| MTPAP | 6 | 5 | 0 | 1 | 1 |
| DENND1A | 4 | 4 | 1,84452E-16 | 0,999999968 | 1 |
| GABRA5 | 5 | 5 | 7,06944E-17 | 0,999999976 | 1 |
| BMPR2 | 6 | 6 | 1,42407E-16 | 0,999999978 | 1 |
| HERC2 | 2 | 2 | 0 | 1 | 1 |
| TFCP2 | 3 | 3 | -1,70072E-17 | 0,99999999 | 1 |
| P4HTM | 1 | 1 | 1,62677E-20 | 1 | 1 |
| LMTK2 | 4 | 3 | 8,06617E-19 | 0,999999999 | 1 |
| MAPRE1 | 11 | 11 | -0,035062142 | 0,233895276 | 1 |
| C2CD6 | 2 | 2 | 0 | 1 | 1 |
| CSTB | 4 | 4 | 2,09777E-18 | 0,999999999 | 1 |
| FNTB | 7 | 7 | 0 | 1 | 1 |
| PI4K2A | 11 | 11 | 3,56937E-16 | 0,999999969 | 1 |
| PIK3R2 | 3 | 3 | -1,83602E-17 | 0,999999994 | 1 |
| CMTM5 | 2 | 2 | 0 | 1 | 1 |
| IMMT | 6 | 6 | 0,022579863 | 0,537943807 | 1 |
| CHRM4 | 1 | 1 | 0 | 1 | 1 |
| GCKR | 1 | 1 | 0 | 1 | 1 |
| PTPN11 | 32 | 32 | -1,15911E-16 | 0,999999947 | 1 |
| MPP1 | 12 | 12 | 0,001677136 | 0,838102717 | 1 |
| IL22RA1 | 1 | 1 | 0 | 1 | 1 |
| UNC13A;UNC13B | 9 | 9 | 0,045187779 | 0,163746795 | 1 |
| CRP | 2 | 2 | 1,91377E-16 | 0,999999989 | 1 |
| HBS1L | 3 | 3 | -5,93494E-19 | 0,999999999 | 1 |
| MYDGF | 6 | 6 | 7,98362E-17 | 0,999999978 | 1 |
| GPRASP2 | 4 | 4 | 6,80519E-16 | 0,999999963 | 1 |
| YPEL5 | 2 | 2 | 5,38929E-14 | 0,999999377 | 1 |
| ABL2 | 9 | 9 | -6,73228E-17 | 0,999999984 | 1 |
| SORBS2 | 1 | 1 | 2,18663E-15 | 0,999999919 | 1 |

| | | | | | |
|---|---|---|---|---|---|
| CYP2J9 | 4 | 3 | 0 | 1 | 1 |
| JMY | 1 | 1 | -1,61227E-17 | 0,999999992 | 1 |
| RNF141 | 2 | 2 | 0 | 1 | 1 |
| MAP3K5 | 7 | 7 | -1,98863E-17 | 0,999999994 | 1 |
| AQP4 | 6 | 6 | 0 | 1 | 1 |
| NR3C1 | 6 | 6 | -1,06127E-16 | 0,999999976 | 1 |
| CACNG3 | 4 | 4 | -0,118660936 | 0,147276287 | 1 |
| NHLRC2 | 13 | 13 | 0 | 1 | 1 |
| RIPOR2 | 3 | 3 | -7,91209E-18 | 0,999999995 | 1 |
| MYL4 | 2 | 2 | -6,40125E-17 | 0,999999983 | 1 |
| SLC1A2 | 11 | 11 | -0,036907068 | 0,185794611 | 1 |
| CCDC136 | 5 | 5 | -2,16779E-18 | 0,999999997 | 1 |
| KNG1 | 4 | 4 | 0,032402509 | 0,58693074 | 1 |
| SH3BGRL2 | 8 | 8 | -4,36542E-18 | 0,999999999 | 1 |
| HARS2 | 10 | 9 | 1,02976E-16 | 0,999999974 | 1 |
| FNBP1L | 2 | 2 | 0 | 1 | 1 |
| ANAPC4 | 5 | 5 | -0,021908087 | 0,485279042 | 1 |
| PRNP | 10 | 10 | 0 | 1 | 1 |
| NECTIN3 | 1 | 1 | -0,194102755 | 0,177422368 | 1 |
| PEAK1 | 6 | 6 | 1,3695E-16 | 0,999999983 | 1 |
| SCAF1 | 1 | 1 | -0,097009739 | 0,262737002 | 1 |
| TSPAN2 | 3 | 3 | 0 | 1 | 1 |
| WWP2 | 2 | 2 | -2,56861E-15 | 0,999999905 | 1 |
| SIKE1 | 1 | 1 | 0 | 1 | 1 |
| UBAP2 | 2 | 2 | -5,98607E-17 | 0,999999992 | 1 |
| FGF12 | 2 | 2 | 8,62147E-18 | 0,999999993 | 1 |
| MPI | 19 | 19 | 0 | 1 | 1 |
| GET3 | 12 | 12 | 0 | 1 | 1 |
| SPATA45 | 1 | 1 | 0 | 1 | 1 |
| WDR4 | 1 | 1 | 0 | 1 | 1 |
| VPS29 | 10 | 10 | 0,023685921 | 0,362037122 | 1 |
| OCIAD1 | 4 | 4 | 1,39675E-16 | 0,99999997 | 1 |
| TPM1 | 7 | 7 | -0,016459896 | 0,596831255 | 1 |

| | | | | | |
|---|---|---|---|---|---|
| ST3GAL5 | 1 | 1 | 0 | 1 | 1 |
| PPP4R2 | 2 | 2 | 8,00128E-16 | 0,999999958 | 1 |
| PBXIP1 | 3 | 3 | 3,68601E-16 | 0,999999961 | 1 |
| CLCN4 | 1 | 1 | 0 | 1 | 1 |
| KCNA6 | 6 | 6 | 0,062108521 | 0,234804016 | 1 |
| SDC4 | 3 | 3 | -5,28027E-18 | 0,999999997 | 1 |
| EIF3J1;EIF3J2 | 7 | 7 | -0,036783499 | 0,473845082 | 1 |
| AP3S2 | 5 | 5 | -0,022971372 | 0,456253841 | 1 |
| RAB2B | 3 | 3 | 0 | 1 | 1 |
| AAMDC | 7 | 7 | 3,24089E-17 | 0,999999986 | 1 |
| SEZ6L | 8 | 8 | -1,70938E-17 | 0,999999987 | 1 |
| SYP | 9 | 9 | 2,13329E-16 | 0,999999974 | 1 |
| GMFB | 8 | 8 | 0,040813119 | 0,159616977 | 1 |
| UHRF1 | 1 | 1 | 0 | 1 | 1 |
| CDH13 | 10 | 10 | -4,36526E-17 | 0,999999994 | 1 |
| SLC17A6 | 7 | 7 | -1,86338E-18 | 0,999999997 | 1 |
| PFDN6 | 2 | 2 | 0 | 1 | 1 |
| PRPF39 | 1 | 1 | 0 | 1 | 1 |
| ANO10 | 2 | 2 | -1,29031E-17 | 0,999999997 | 1 |
| AW551984 | 1 | 1 | 0,059268593 | 0,374054965 | 1 |
| MMS19 | 2 | 2 | 6,6265E-11 | 0,999977967 | 1 |
| KRT18 | 1 | 1 | 8,67486E-18 | 0,999999996 | 1 |
| SLC7A6OS | 2 | 2 | -8,85903E-18 | 0,999999997 | 1 |
| HIGD1A | 1 | 1 | 0 | 1 | 1 |
| sp\|Q9D9H8\|CB069 | 4 | 4 | 3,46105E-16 | 0,999999969 | 1 |
| LIG1 | 1 | 1 | 0 | 1 | 1 |
| sp\|Q9D1K7\|CT027 | 4 | 4 | 1,32494E-20 | 1 | 1 |
| EIF4A3 | 3 | 3 | -0,014424002 | 0,696826223 | 1 |
| PEX16 | 1 | 1 | 0 | 1 | 1 |
| VEZT | 3 | 2 | -5,09868E-17 | 0,999999989 | 1 |
| PPFIA1 | 4 | 4 | 0 | 1 | 1 |
| TMEM109 | 2 | 2 | 1,56994E-20 | 1 | 1 |
| SYPL1 | 2 | 2 | 0 | 1 | 1 |

| | | | | | |
|---|---|---|---|---|---|
| KIDINS220 | 9 | 9 | -2,0838E-14 | 0,999999573 | 1 |
| SYT7 | 3 | 3 | -3,68172E-25 | 1 | 1 |
| LAMTOR2 | 3 | 3 | 0 | 1 | 1 |
| L1CAM | 4 | 4 | 0 | 1 | 1 |
| DPM1 | 7 | 7 | 0 | 1 | 1 |
| EPN2 | 1 | 1 | -0,09216581 | 0,308813629 | 1 |
| NUDT3 | 9 | 9 | 5,47375E-18 | 0,999999998 | 1 |
| DHX15 | 8 | 8 | -1,43759E-16 | 0,999999984 | 1 |
| TUBB3 | 15 | 15 | -0,005280979 | 0,738048645 | 1 |
| TUBB4B;TUBB5;TU | 6 | 6 | 2,67809E-16 | 0,999999968 | 1 |
| DNM2;DNM3 | 8 | 8 | 0 | 1 | 1 |
| RAB18 | 7 | 7 | 0 | 1 | 1 |
| HECTD1 | 3 | 3 | 0 | 1 | 1 |
| CAND1;CAND2 | 3 | 3 | -4,22358E-21 | 1 | 1 |
| GNPAT | 2 | 2 | 0 | 1 | 1 |
| RHOC;RHOA | 5 | 5 | 0 | 1 | 1 |
| PDCD6IP | 1 | 1 | 0 | 1 | 1 |
| AKR1B1 | 17 | 16 | 0 | 1 | 1 |
| SDF2 | 2 | 2 | -4,04827E-17 | 0,999999988 | 1 |
| NXN | 2 | 2 | 0 | 1 | 1 |
| PFDN1 | 4 | 4 | -0,02233275 | 0,572165994 | 1 |
| TIPRL | 13 | 12 | -2,60839E-17 | 0,999999996 | 1 |
| NUP98 | 1 | 1 | -0,016530422 | 0,718826587 | 1 |
| DAZL | 1 | 1 | 0 | 1 | 1 |
| ENTPD3 | 2 | 2 | -8,8282E-17 | 0,999999995 | 1 |
| TRIM46 | 7 | 7 | 0 | 1 | 1 |
| MTAP | 7 | 7 | -9,0956E-17 | 0,999999985 | 1 |
| TIMM22 | 2 | 2 | 2,77193E-18 | 0,999999998 | 1 |
| MRPL22 | 3 | 3 | 0 | 1 | 1 |
| KARS1 | 26 | 26 | -0,007500757 | 0,628312833 | 1 |
| STAT2 | 1 | 1 | 0 | 1 | 1 |
| DNAJC3 | 5 | 5 | -8,99711E-18 | 0,999999994 | 1 |
| ATP6AP1 | 7 | 7 | -2,35857E-17 | 0,999999994 | 1 |

| | | | | | |
|---|---|---|---|---|---|
| NDUFS6 | 7 | 7 | 0,069876754 | 0,141150887 | 1 |
| ABCF2 | 7 | 7 | -5,79767E-18 | 0,999999994 | 1 |
| RBBP8 | 1 | 1 | -0,013984539 | 0,793384245 | 1 |
| sp\|P06330\|HVM51 | 3 | 1 | 3,46444E-17 | 0,999999992 | 1 |
| HCLS1 | 3 | 3 | 0,113319488 | 0,121953524 | 1 |
| SRBD1 | 1 | 1 | 0 | 1 | 1 |
| EEF1A2 | 14 | 14 | -5,84207E-16 | 0,99999992 | 1 |
| SH2D3C | 3 | 2 | 5,46523E-18 | 0,999999997 | 1 |
| AIF1 | 2 | 2 | 1,90768E-18 | 0,999999998 | 1 |
| EEF1A1 | 14 | 14 | -0,038557836 | 0,230838054 | 1 |
| HMGCL | 9 | 9 | 4,46135E-16 | 0,99999995 | 1 |
| RAB4A;RAB4B | 2 | 2 | -7,53907E-18 | 0,999999997 | 1 |
| URB1 | 1 | 1 | 0 | 1 | 1 |
| ETNK1 | 5 | 5 | 2,0984E-16 | 0,999999967 | 1 |
| SNRPA | 1 | 1 | 0 | 1 | 1 |
| MCAM | 4 | 3 | 0 | 1 | 1 |
| FGGY | 1 | 1 | 0 | 1 | 1 |
| METAP2 | 9 | 9 | -1,99728E-18 | 0,999999998 | 1 |
| LACTB | 16 | 15 | 0 | 1 | 1 |
| KLHL22 | 6 | 6 | 2,36079E-16 | 0,999999957 | 1 |
| NUDT4 | 4 | 4 | -1,23082E-17 | 0,999999998 | 1 |
| LIN7C | 6 | 5 | 0 | 1 | 1 |
| GNAI3 | 7 | 7 | 9,8673E-18 | 0,999999996 | 1 |
| CRBN | 5 | 5 | -1,37119E-18 | 0,999999999 | 1 |
| PEBP1 | 16 | 16 | 1,15139E-15 | 0,999999906 | 1 |
| RAB31 | 4 | 4 | -7,60405E-16 | 0,999999925 | 1 |
| CSNK1G2;CSNK1G3 | 3 | 3 | 0 | 1 | 1 |
| BRK1 | 3 | 3 | -2,09633E-17 | 0,999999991 | 1 |
| CNTFR | 4 | 4 | 5,68426E-16 | 0,999999934 | 1 |
| MRPS15 | 4 | 4 | 3,20728E-16 | 0,99999996 | 1 |
| SNX4 | 15 | 15 | 1,09093E-17 | 0,999999993 | 1 |
| NDUFB4 | 7 | 7 | 0,030106598 | 0,410848519 | 1 |
| FAF1 | 9 | 9 | -0,03238696 | 0,292563935 | 1 |

| | | | | | |
|---|---|---|---|---|---|
| MAGI1 | 7 | 7 | -4,21827E-16 | 0,999999936 | 1 |
| QKI | 4 | 4 | -1,9119E-15 | 0,999999898 | 1 |
| GRIA4 | 4 | 4 | 3,10347E-17 | 0,999999992 | 1 |
| SENP6 | 1 | 1 | 0 | 1 | 1 |
| NEFL;NEFH;VIM;IN/ | 1 | 1 | 0 | 1 | 1 |
| KRT5 | 37 | 12 | -3,89841E-16 | 0,999999992 | 1 |
| CELSR1 | 1 | 1 | 0 | 1 | 1 |
| ERGIC1 | 5 | 5 | -4,57323E-17 | 0,999999984 | 1 |
| CSNK1G1 | 2 | 2 | 4,39999E-17 | 0,999999994 | 1 |
| DNAJC8 | 3 | 3 | -0,041506086 | 0,397212414 | 1 |
| UBASH3B | 4 | 4 | 0 | 1 | 1 |
| ATAD1 | 11 | 10 | 0,043568592 | 0,174715394 | 1 |
| TALDO1 | 25 | 25 | 0 | 1 | 1 |
| SHC2 | 2 | 2 | 1,96132E-15 | 0,999999925 | 1 |
| ILKAP | 1 | 1 | -1,39737E-18 | 0,999999998 | 1 |
| HSD17B11 | 5 | 5 | 4,05466E-17 | 0,99999998 | 1 |
| RAB1A;RAB1B | 5 | 5 | -1,55534E-17 | 0,999999996 | 1 |
| BPHL | 14 | 13 | 6,38283E-17 | 0,999999979 | 1 |
| SLC22A4 | 2 | 2 | 0 | 1 | 1 |
| THG1L | 2 | 2 | 0 | 1 | 1 |
| ARL4C | 1 | 1 | -0,196671754 | 0,244753653 | 1 |
| MLEC | 8 | 8 | 1,04876E-16 | 0,99999997 | 1 |
| MRPL38 | 3 | 3 | 2,13847E-19 | 1 | 1 |
| TRAPPC2 | 2 | 2 | -5,44485E-17 | 0,999999987 | 1 |
| STON2 | 7 | 7 | 3,93596E-16 | 0,999999954 | 1 |
| PSMB10 | 1 | 1 | 0,188690182 | 0,25421502 | 1 |
| WASHC5 | 8 | 7 | 0,0322785 | 0,337531869 | 1 |
| ZYX | 7 | 7 | -1,77014E-15 | 0,999999881 | 1 |
| SNF8 | 4 | 4 | -0,006816282 | 0,749391811 | 1 |
| ACADSB | 11 | 11 | -9,50613E-18 | 0,999999995 | 1 |
| VSNL1;HPCAL4 | 4 | 4 | -0,075015339 | 0,222318343 | 1 |
| RAB39B | 11 | 11 | 0,031930143 | 0,265735281 | 1 |
| HSPA14 | 1 | 1 | 0,114554078 | 0,318782689 | 1 |

| | | | | | |
|---|---|---|---|---|---|
| RANBP1 | 5 | 5 | 0 | 1 | 1 |
| KRT1;KRT6A;KRT2;I | 1 | 1 | 0 | 1 | 1 |
| USP13 | 2 | 2 | 0 | 1 | 1 |
| PRL8A8 | 1 | 1 | 0 | 1 | 1 |
| LRRC8A;LRRC8C | 1 | 1 | 0 | 1 | 1 |
| GPR20 | 1 | 1 | -0,150135877 | 0,144776794 | 1 |
| LCMT1 | 12 | 12 | -1,52567E-17 | 0,999999995 | 1 |
| CYP51A1 | 4 | 4 | -0,034424565 | 0,401211255 | 1 |
| GLG1 | 9 | 9 | -8,80993E-18 | 0,999999993 | 1 |
| MSRB2 | 5 | 5 | 2,39919E-06 | 0,993971331 | 1 |
| USP9Y | 11 | 11 | -4,47209E-18 | 0,999999998 | 1 |
| SPAST | 6 | 6 | 0,042990149 | 0,241970108 | 1 |
| SPCS2 | 6 | 6 | 2,87318E-17 | 0,999999985 | 1 |
| KATNAL1 | 6 | 6 | -3,12772E-19 | 0,999999999 | 1 |
| WNK2;WNK3 | 1 | 1 | 0 | 1 | 1 |
| AP3M1 | 7 | 7 | 0 | 1 | 1 |
| NAP1L5 | 1 | 1 | 0 | 1 | 1 |
| MRS2 | 8 | 7 | 9,93568E-18 | 0,999999994 | 1 |
| TUBA1C;TUBAL3;TU | 2 | 2 | 0 | 1 | 1 |
| TENM3 | 10 | 9 | 0 | 1 | 1 |
| HSPA1L;HSPA2 | 2 | 2 | 0 | 1 | 1 |
| USP4 | 10 | 10 | -7,47471E-18 | 0,999999995 | 1 |
| CPNE5;CPNE8 | 3 | 3 | -0,023575768 | 0,5868288 | 1 |
| ILK | 2 | 2 | 1,78855E-18 | 0,999999998 | 1 |
| SCN3A;SCN2A;SCN9 | 4 | 4 | 1,32318E-18 | 0,999999998 | 1 |
| SCN1A | 15 | 15 | 1,72759E-16 | 0,999999978 | 1 |
| NDUFA8 | 8 | 8 | 5,75852E-16 | 0,999999945 | 1 |
| DAP3 | 9 | 9 | -1,40617E-17 | 0,999999996 | 1 |
| PRKAG1;PRKAG2 | 1 | 1 | 0 | 1 | 1 |
| CWF19L1 | 2 | 2 | 0 | 1 | 1 |
| RPS15A | 9 | 9 | -1,04743E-17 | 0,999999995 | 1 |
| NIBAN2 | 3 | 3 | -4,72185E-17 | 0,999999991 | 1 |
| KCNC3 | 4 | 4 | 0 | 1 | 1 |

| | | | | | |
|---|---|---|---|---|---|
| SMCR8 | 14 | 14 | 0 | 1 | 1 |
| SNX15 | 4 | 4 | 7,76187E-17 | 0,999999988 | 1 |
| VTN | 1 | 1 | 0 | 1 | 1 |
| MRPS11 | 1 | 1 | 0,102180686 | 0,284133915 | 1 |
| SLC4A10;SLC4A8 | 2 | 2 | 0,017648441 | 0,695424853 | 1 |
| OGDH | 8 | 6 | 1,06252E-07 | 0,999191813 | 1 |
| EEF1D | 1 | 1 | 0 | 1 | 1 |
| KDSR | 2 | 2 | 1,46029E-16 | 0,999999973 | 1 |
| PBDC1 | 4 | 4 | 0 | 1 | 1 |
| BOLA2 | 3 | 3 | -2,03073E-15 | 0,9999999 | 1 |
| B3GAT3 | 4 | 4 | 0 | 1 | 1 |
| VPS26B | 12 | 12 | 0 | 1 | 1 |
| RAB5C;RAB5B;RAB! | 4 | 4 | -0,088755235 | 0,118252875 | 1 |
| CBFB | 1 | 1 | 0 | 1 | 1 |
| POLDIP2 | 9 | 9 | 3,06196E-15 | 0,999999838 | 1 |
| AGAP1 | 3 | 3 | 0 | 1 | 1 |
| 1700109H08RIK | 1 | 1 | 0 | 1 | 1 |
| ATP5PF | 5 | 5 | 0 | 1 | 1 |
| PTPRF | 6 | 6 | 7,53201E-19 | 0,999999999 | 1 |
| TM9SF4 | 4 | 4 | -6,74718E-18 | 0,999999995 | 1 |
| TRPC4 | 3 | 3 | 0 | 1 | 1 |
| TH | 2 | 2 | -1,19032E-15 | 0,99999994 | 1 |
| TIMELESS | 1 | 1 | 2,10644E-19 | 0,999999999 | 1 |
| AP4B1 | 1 | 1 | 0 | 1 | 1 |
| MAPRE1;MAPRE3 | 2 | 2 | 0 | 1 | 1 |
| SSR4 | 6 | 6 | -7,78383E-17 | 0,999999971 | 1 |
| PPP2R2B | 1 | 1 | 0 | 1 | 1 |
| PUM2 | 1 | 1 | -0,10632549 | 0,135646031 | 1 |
| RPL15 | 8 | 8 | -4,05191E-17 | 0,999999992 | 1 |
| PURB | 14 | 14 | 0 | 1 | 1 |
| GNAO1 | 6 | 6 | -1,2656E-18 | 1 | 1 |
| NCAM1 | 6 | 6 | 0 | 1 | 1 |
| TOMM20 | 4 | 4 | 0 | 1 | 1 |

| | | | | | |
|---|---|---|---|---|---|
| DST;MACF1 | 3 | 3 | -0,046679855 | 0,320421567 | 1 |
| TMEM43 | 6 | 6 | 0 | 1 | 1 |
| N6AMT1 | 3 | 3 | 0 | 1 | 1 |
| DDT | 13 | 13 | 4,71454E-16 | 0,99999994 | 1 |
| JMJD6 | 1 | 1 | 0 | 1 | 1 |
| SLC25A1 | 10 | 10 | 6,00141E-17 | 0,999999981 | 1 |
| LPGAT1 | 4 | 4 | 1,3271E-18 | 0,999999998 | 1 |
| SLC25A40 | 3 | 3 | 0 | 1 | 1 |
| 1700012B07RIK | 1 | 1 | 0 | 1 | 1 |
| DOCK8 | 2 | 2 | 6,64251E-17 | 0,999999984 | 1 |
| SORBS1 | 5 | 5 | -6,55576E-18 | 0,999999995 | 1 |
| NDRG2 | 13 | 13 | -0,064400912 | 0,182303079 | 1 |
| SCN2B | 9 | 8 | 0 | 1 | 1 |
| INPP4A | 4 | 4 | -1,67761E-16 | 0,999999973 | 1 |
| MDGA2 | 4 | 4 | 0 | 1 | 1 |
| PDCL3 | 2 | 2 | -0,015603777 | 0,643474449 | 1 |
| HNRNPAB;HNRNPD | 1 | 1 | -0,249144298 | 0,117426017 | 1 |
| PCSK1 | 1 | 1 | 0 | 1 | 1 |
| MAPKAP1 | 1 | 1 | 0 | 1 | 1 |
| MARS1 | 1 | 1 | -3,28635E-18 | 0,999999996 | 1 |
| RIT2 | 3 | 3 | 3,30246E-17 | 0,999999989 | 1 |
| CPNE3 | 5 | 5 | 3,85564E-18 | 0,999999998 | 1 |
| IST1 | 6 | 6 | -1,53702E-16 | 0,999999965 | 1 |
| POFUT1 | 1 | 1 | 0 | 1 | 1 |
| COQ10A | 2 | 1 | 0,114615442 | 0,409704607 | 1 |
| U2SURP | 1 | 1 | 0 | 1 | 1 |
| CPEB3;CPEB2 | 1 | 1 | 0 | 1 | 1 |
| SCFD1 | 5 | 5 | -0,03907473 | 0,351829571 | 1 |
| PTPA | 14 | 14 | -6,86881E-15 | 0,999999727 | 1 |
| FIG4 | 2 | 2 | 4,91945E-16 | 0,999999953 | 1 |
| PDE3A | 1 | 1 | 0,012924227 | 0,763551665 | 1 |
| LIMA1 | 2 | 2 | -0,098924049 | 0,1683591 | 1 |
| NARS2 | 5 | 4 | 0 | 1 | 1 |

| | | | | | |
|---|---|---|---|---|---|
| ABRACL | 1 | 1 | 0 | 1 | 1 |
| PDE4B | 9 | 9 | 3,44594E-17 | 0,999999982 | 1 |
| IGSF9B | 2 | 1 | 0 | 1 | 1 |
| COMMD7 | 4 | 4 | 0 | 1 | 1 |
| TMOD1 | 9 | 9 | -0,010162367 | 0,713206516 | 1 |
| HIF1AN | 1 | 1 | 0 | 1 | 1 |
| NDUFAF7 | 9 | 7 | 1,51145E-16 | 0,999999967 | 1 |
| RPL22L1 | 3 | 3 | -2,36469E-14 | 0,999999645 | 1 |
| DYNC1I1;DYNC1I2 | 2 | 2 | -3,82478E-19 | 0,999999999 | 1 |
| ETF1 | 12 | 12 | -0,020193105 | 0,368695433 | 1 |
| PGK2;PGK1 | 8 | 8 | -1,62594E-16 | 0,999999983 | 1 |
| PTPRA;PTPRE | 1 | 1 | 0 | 1 | 1 |
| WWP1 | 3 | 2 | 0 | 1 | 1 |
| TIMM8A2;TIMM8A | 1 | 1 | 0,001246403 | 0,924901172 | 1 |
| PTP4A1 | 3 | 3 | 1,22286E-17 | 0,999999997 | 1 |
| GLRX2 | 1 | 1 | 0,038835503 | 0,547647578 | 1 |
| GAREM2 | 1 | 1 | 0 | 1 | 1 |
| SMG5 | 1 | 1 | 0 | 1 | 1 |
| AP2S1 | 11 | 11 | 0 | 1 | 1 |
| WDR81 | 4 | 4 | -1,56804E-17 | 0,999999989 | 1 |
| WDR61 | 6 | 6 | -7,71793E-17 | 0,999999968 | 1 |
| PACS2 | 6 | 6 | -2,6998E-17 | 0,999999991 | 1 |
| NFU1 | 6 | 6 | 1,73721E-19 | 1 | 1 |
| TFB1M | 3 | 2 | 0 | 1 | 1 |
| ARHGEF1 | 3 | 3 | 3,44941E-16 | 0,99999996 | 1 |
| SNAP91;PICALM | 2 | 2 | 0 | 1 | 1 |
| TBC1D7 | 1 | 1 | 0 | 1 | 1 |
| SPIRE1 | 5 | 5 | -1,24263E-17 | 0,999999994 | 1 |
| ABHD17A | 2 | 1 | 0,006554167 | 0,882890087 | 1 |
| KIAA0100 | 1 | 1 | -0,004898527 | 0,869091767 | 1 |
| RAB27B;RAB27A | 2 | 2 | -5,6057E-18 | 0,999999997 | 1 |
| KIAA1109 | 15 | 13 | 6,22633E-20 | 1 | 1 |
| MCRIP1 | 1 | 1 | 0 | 1 | 1 |

| | | | | | |
|---|---|---|---|---|---|
| DYRK1A | 4 | 4 | -7,80777E-15 | 0,999999797 | 1 |
| HMGB1 | 9 | 9 | -0,053944538 | 0,338276047 | 1 |
| WNK1;WNK2;WNK | 2 | 2 | 1,69789E-18 | 0,999999999 | 1 |
| NLRP9C | 1 | 1 | 0 | 1 | 1 |
| GSTM1;GSTM2;GST | 3 | 3 | -0,053961706 | 0,274809162 | 1 |
| GID4 | 1 | 1 | 0 | 1 | 1 |
| VPS41 | 6 | 6 | -1,57014E-17 | 0,999999991 | 1 |
| SH3BP1 | 5 | 5 | 0 | 1 | 1 |
| HBA-A1 | 9 | 9 | -1,81195E-15 | 0,999999922 | 1 |
| CNNM2 | 6 | 6 | 0 | 1 | 1 |
| H2-EB2 | 1 | 1 | -0,116314403 | 0,551514246 | 1 |
| BAG2 | 2 | 2 | -7,89725E-12 | 0,999994511 | 1 |
| COA3 | 2 | 2 | 0 | 1 | 1 |
| MAGEE1 | 1 | 1 | 0 | 1 | 1 |
| PTPN3 | 3 | 3 | 1,03211E-16 | 0,99999998 | 1 |
| EPHB3 | 7 | 7 | 0,003904245 | 0,769758486 | 1 |
| UCHL3 | 13 | 13 | 0 | 1 | 1 |
| WASHC4 | 11 | 11 | -3,39234E-17 | 0,999999984 | 1 |
| CD109 | 6 | 6 | -2,27778E-18 | 0,999999998 | 1 |
| ANXA1 | 17 | 6 | 0,047774762 | 0,547294864 | 1 |
| AMZ2 | 2 | 2 | 0,00882776 | 0,681977013 | 1 |
| VIM;KRT6A;KRT76; | 1 | 1 | 0 | 1 | 1 |
| KRT72 | 1 | 1 | 0 | 1 | 1 |
| KRT1;KRT77;KRT73 | 1 | 1 | 0 | 1 | 1 |
| RGS7;RGS6 | 4 | 4 | 3,06373E-17 | 0,999999989 | 1 |
| UPF3B | 2 | 2 | 2,99064E-17 | 0,999999991 | 1 |
| RAB8B | 6 | 6 | 0 | 1 | 1 |
| PROX1 | 1 | 1 | 0 | 1 | 1 |
| SUMO1 | 2 | 2 | 6,767E-17 | 0,999999988 | 1 |
| SLC4A8 | 5 | 5 | 1,03185E-18 | 0,999999999 | 1 |
| SLC4A7 | 8 | 6 | 1,17724E-16 | 0,999999981 | 1 |
| AIMP2 | 6 | 5 | 0,0119896 | 0,624728745 | 1 |
| ANK2 | 6 | 6 | 9,86682E-18 | 0,999999991 | 1 |

| | | | | | |
|---|---|---|---|---|---|
| TXNRD1 | 20 | 20 | 2,16645E-15 | 0,999999855 | 1 |
| SNRPD3 | 3 | 3 | -0,04898858 | 0,381479783 | 1 |
| PTP4A2;PTP4A1 | 3 | 3 | -9,7369E-16 | 0,999999915 | 1 |
| NDST2 | 1 | 1 | 0 | 1 | 1 |
| FADS2 | 3 | 3 | 1,81559E-17 | 0,999999994 | 1 |
| NUP133 | 1 | 1 | 0,206790824 | 0,169694283 | 1 |
| WDR26 | 9 | 9 | -0,041617442 | 0,183615521 | 1 |
| AP1G2;AP1G1 | 2 | 2 | 0 | 1 | 1 |
| SKIV2L | 1 | 1 | 0 | 1 | 1 |
| CMIP | 3 | 3 | 0 | 1 | 1 |
| ASCC3 | 1 | 1 | 0 | 1 | 1 |
| GM10717 | 1 | 1 | 0 | 1 | 1 |
| LIN52 | 1 | 1 | -0,200695999 | 0,234107279 | 1 |
| PPP1CB | 6 | 6 | 1,14523E-15 | 0,999999901 | 1 |
| RGS17 | 1 | 1 | 0 | 1 | 1 |
| MAPK8IP1 | 2 | 2 | 0,068932713 | 0,351922237 | 1 |
| VPS35L | 7 | 7 | -3,37568E-15 | 0,999999824 | 1 |
| EFNB2 | 2 | 2 | -1,11182E-17 | 0,999999995 | 1 |
| NPC1 | 3 | 3 | 0,063332204 | 0,385113712 | 1 |
| GSTM7 | 14 | 14 | -8,57149E-17 | 0,999999985 | 1 |
| KCNN2 | 3 | 3 | 0,050634083 | 0,447269317 | 1 |
| MAN2B1 | 3 | 3 | 0,095637627 | 0,244983109 | 1 |
| RNF123 | 6 | 6 | -6,44381E-17 | 0,999999987 | 1 |
| HID1 | 10 | 10 | -8,93473E-17 | 0,999999972 | 1 |
| ABCC9 | 1 | 1 | 0 | 1 | 1 |
| ZFP735 | 1 | 1 | 0 | 1 | 1 |
| ITPR1;ITPR2 | 2 | 2 | -5,29807E-19 | 1 | 1 |
| NOL7 | 1 | 1 | 0 | 1 | 1 |
| MCMBP | 2 | 1 | 0 | 1 | 1 |
| UST | 2 | 2 | 3,06698E-20 | 1 | 1 |
| MRPL19 | 7 | 6 | 0 | 1 | 1 |
| MRPL16 | 3 | 2 | -2,02828E-16 | 0,999999987 | 1 |
| HKDC1 | 8 | 8 | 0,03500866 | 0,336751592 | 1 |

| | | | | | |
|---|---|---|---|---|---|
| ACSF3 | 13 | 13 | 6,13363E-17 | 0,999999975 | 1 |
| COQ5 | 5 | 5 | 0 | 1 | 1 |
| RGS19 | 1 | 1 | 0 | 1 | 1 |
| INPP5F | 10 | 10 | 0 | 1 | 1 |
| SSR1 | 3 | 3 | -0,042691455 | 0,425106506 | 1 |
| RPL32 | 1 | 1 | 0 | 1 | 1 |
| TNIK;MAP4K4;MIN | 9 | 9 | 0 | 1 | 1 |
| NDST3;NDST4 | 1 | 1 | -0,213315796 | 0,402484485 | 1 |
| MARK3 | 12 | 12 | -6,69417E-17 | 0,999999985 | 1 |
| PLCH2;PLCH1 | 2 | 2 | 0 | 1 | 1 |
| CABLES2 | 1 | 1 | 0,112025601 | 0,43103577 | 1 |
| PPOX | 5 | 5 | 0,051001676 | 0,319347896 | 1 |
| PRKCE;PRKCH | 1 | 1 | 0 | 1 | 1 |
| SLC25A23 | 22 | 21 | -1,47633E-17 | 0,999999991 | 1 |
| MRPS2 | 5 | 5 | 1,00178E-16 | 0,99999998 | 1 |
| TATDN1 | 3 | 3 | -0,001586832 | 0,898856081 | 1 |
| LRRC20 | 2 | 2 | 0 | 1 | 1 |
| ACYP1 | 5 | 5 | 2,22287E-17 | 0,999999991 | 1 |
| SNX7 | 4 | 4 | 9,50114E-17 | 0,99999997 | 1 |
| NRXN3;NRXN1 | 3 | 3 | 0 | 1 | 1 |
| SLC13A5 | 2 | 2 | 0 | 1 | 1 |
| PARL | 2 | 2 | 0 | 1 | 1 |
| FGF1 | 2 | 2 | 0 | 1 | 1 |
| CSNK2B | 5 | 5 | -2,07263E-16 | 0,999999958 | 1 |
| UBR2 | 1 | 1 | 0 | 1 | 1 |
| UBE2Z | 9 | 9 | -0,03313703 | 0,349558172 | 1 |
| BMPR1A | 2 | 2 | 0 | 1 | 1 |
| D3ERTD751E | 2 | 2 | 1,67307E-16 | 0,999999988 | 1 |
| XRCC5 | 2 | 2 | 3,53665E-15 | 0,999999848 | 1 |
| CASK | 1 | 1 | 0 | 1 | 1 |
| C1QC | 4 | 4 | 1,12728E-18 | 0,999999999 | 1 |
| RAN | 12 | 12 | 0 | 1 | 1 |
| PPIF | 6 | 6 | 0 | 1 | 1 |

| | | | | | |
|---|---|---|---|---|---|
| TTN | 7 | 5 | -0,05789378 | 0,171798684 | 1 |
| DNM1;DNM2 | 8 | 8 | 0 | 1 | 1 |
| TUBB1;TUBB4B;TU | 4 | 4 | -3,38199E-17 | 0,999999984 | 1 |
| NT5E | 2 | 2 | 0 | 1 | 1 |
| HAPLN2 | 8 | 8 | -0,013576898 | 0,796356727 | 1 |
| RPL23A | 7 | 7 | 6,57952E-17 | 0,999999982 | 1 |
| COX6B1 | 7 | 7 | 2,96828E-18 | 0,999999996 | 1 |
| DHRS4 | 8 | 8 | 0 | 1 | 1 |
| CAMK2G | 3 | 3 | 1,61282E-15 | 0,999999912 | 1 |
| SLC6A15;SLC6A17 | 1 | 1 | -1,35553E-21 | 1 | 1 |
| SLC6A6;SLC6A2;SL( | 1 | 1 | 0 | 1 | 1 |
| PON2 | 5 | 5 | -2,51392E-15 | 0,999999877 | 1 |
| GADD45GIP1 | 2 | 2 | 0 | 1 | 1 |
| ITPK1 | 5 | 5 | -6,06782E-16 | 0,999999939 | 1 |
| SCN1A;SCN3A | 1 | 1 | 0 | 1 | 1 |
| TXNL1 | 18 | 18 | -1,00389E-16 | 0,99999997 | 1 |
| BLES03 | 9 | 9 | -2,45396E-17 | 0,999999991 | 1 |
| AP3B1 | 14 | 14 | 0 | 1 | 1 |
| CC2D2B | 1 | 1 | 0 | 1 | 1 |
| SRP14 | 3 | 3 | -3,41393E-17 | 0,999999986 | 1 |
| HAGHL | 5 | 5 | -9,02722E-17 | 0,999999982 | 1 |
| SBNO1 | 3 | 3 | 0,060434353 | 0,261197819 | 1 |
| SKT | 3 | 3 | 0 | 1 | 1 |
| ATP2B4 | 5 | 5 | -0,060227925 | 0,213333406 | 1 |
| CELF3 | 1 | 1 | 0 | 1 | 1 |
| GNAS;GNAL | 5 | 5 | 0 | 1 | 1 |
| MRPL1 | 10 | 9 | 0 | 1 | 1 |
| COMMD4 | 1 | 1 | 0 | 1 | 1 |
| DYNC2I2 | 1 | 1 | 0 | 1 | 1 |
| DGKA | 1 | 1 | 0 | 1 | 1 |
| SPOCK3 | 1 | 1 | 0 | 1 | 1 |
| FHIT | 3 | 3 | 1,08831E-16 | 0,999999981 | 1 |
| SEC14L3 | 1 | 1 | 0 | 1 | 1 |

| | | | | | |
|---|---|---|---|---|---|
| GRIP1 | 9 | 9 | 2,36125E-17 | 0,999999994 | 1 |
| CORO1B | 17 | 16 | 0 | 1 | 1 |
| CLCN6 | 8 | 8 | 0,014043449 | 0,522548789 | 1 |
| TPP1 | 4 | 4 | -2,8775E-19 | 0,999999999 | 1 |
| LGALSL | 7 | 7 | -0,014782454 | 0,670242836 | 1 |
| ZBTB8OS | 1 | 1 | 0 | 1 | 1 |
| SRPK1 | 4 | 4 | 0 | 1 | 1 |
| ARHGAP32 | 12 | 12 | 7,91699E-15 | 0,999999645 | 1 |
| CREBZF | 1 | 1 | -0,007764086 | 0,837719596 | 1 |
| PHACTR3 | 2 | 2 | 1,47437E-16 | 0,999999973 | 1 |
| SCD2;SCD1;SCD4;S( | 2 | 2 | -8,4071E-13 | 0,999998759 | 1 |
| PIN4 | 1 | 1 | 0 | 1 | 1 |
| GFER | 4 | 4 | 0 | 1 | 1 |
| sp\|P01631\|KV2A7_ | 1 | 1 | 0,23268689 | 0,320238564 | 1 |
| IGKV12-41 | 1 | 1 | 0,12253048 | 0,431502569 | 1 |
| GMPR2 | 5 | 5 | 0 | 1 | 1 |
| PRPS1L1 | 4 | 4 | 5,63142E-17 | 0,999999976 | 1 |
| PRPS2 | 13 | 13 | -3,46154E-17 | 0,999999986 | 1 |
| RPS6KB1 | 3 | 3 | 0 | 1 | 1 |
| ATP5PO | 18 | 18 | 0 | 1 | 1 |
| PDK1;PDK2 | 1 | 1 | 0,06771442 | 0,316819151 | 1 |
| RRM2B | 3 | 3 | 0 | 1 | 1 |
| KPTN | 1 | 1 | 0 | 1 | 1 |
| GC | 5 | 5 | 0,023937443 | 0,654518726 | 1 |
| MARS2 | 8 | 8 | 5,3479E-19 | 0,999999999 | 1 |
| RPL10A | 1 | 1 | 0 | 1 | 1 |
| NUDT7 | 1 | 1 | 0 | 1 | 1 |
| SLC39A10 | 9 | 9 | 2,50945E-15 | 0,999999835 | 1 |
| MBP | 3 | 3 | 3,52603E-19 | 0,999999999 | 1 |
| SEC23A;SEC23B | 3 | 3 | -0,118052862 | 0,113591269 | 1 |
| GRIN2C | 1 | 1 | 0 | 1 | 1 |
| PTPN1 | 4 | 4 | -2,5632E-16 | 0,999999959 | 1 |
| ITGA6 | 5 | 4 | -8,19388E-17 | 0,999999987 | 1 |

| | | | | | |
|---|---|---|---|---|---|
| BCCIP | 2 | 2 | -1,78242E-18 | 0,999999998 | 1 |
| FHL1 | 2 | 2 | -1,68193E-16 | 0,999999987 | 1 |
| CAMK2G | 4 | 4 | -0,057807972 | 0,288106506 | 1 |
| EPHA6 | 5 | 5 | 0 | 1 | 1 |
| KCNJ13 | 4 | 4 | 0 | 1 | 1 |
| PTP4A2 | 3 | 3 | 0 | 1 | 1 |
| SLC24A4 | 1 | 1 | 2,78208E-20 | 1 | 1 |
| TAS2R41 | 1 | 1 | -0,20114883 | 0,187073307 | 1 |
| PMPCB;UQCRC1 | 1 | 1 | 0,094475031 | 0,156445303 | 1 |
| TTN | 1 | 1 | 0 | 1 | 1 |
| NAAA | 4 | 4 | -0,073328208 | 0,25500533 | 1 |
| SLC6A2 | 2 | 2 | 8,79023E-18 | 0,999999997 | 1 |
| ELL | 1 | 1 | 0 | 1 | 1 |
| sp\|Q8BN57\|CC033 | 2 | 1 | 0 | 1 | 1 |
| SNX12 | 6 | 6 | -9,58334E-18 | 0,999999993 | 1 |
| ITCH | 7 | 7 | 0 | 1 | 1 |
| CDK14 | 5 | 5 | -7,93756E-18 | 0,999999996 | 1 |
| BRCC3 | 6 | 6 | 0 | 1 | 1 |
| PMM2 | 13 | 13 | -1,91412E-15 | 0,999999835 | 1 |
| SLC25A44 | 7 | 6 | -4,76499E-17 | 0,999999994 | 1 |
| CD101 | 2 | 2 | 0,003615107 | 0,863647452 | 1 |
| CD47 | 5 | 5 | 0 | 1 | 1 |
| COQ4 | 4 | 4 | -0,032063367 | 0,516391414 | 1 |
| PLXNA1;PLXNA4 | 3 | 3 | 0 | 1 | 1 |
| SESTD1 | 4 | 4 | 0 | 1 | 1 |
| TTR | 6 | 4 | -0,073662272 | 0,523076935 | 1 |
| DOCK11 | 6 | 6 | 0 | 1 | 1 |
| MRPS25 | 4 | 4 | 0 | 1 | 1 |
| PAN2 | 1 | 1 | 0 | 1 | 1 |
| TXN2 | 2 | 2 | 0 | 1 | 1 |
| MTMR9 | 5 | 5 | 0 | 1 | 1 |
| GRN | 1 | 1 | 0 | 1 | 1 |
| GLS2 | 2 | 2 | -0,266719852 | 0,178712103 | 1 |

| | | | | | |
|---|---|---|---|---|---|
| UCKL1 | 6 | 6 | 0 | 1 | 1 |
| HERC1 | 4 | 4 | 2,335E-16 | 0,999999961 | 1 |
| FRS2 | 1 | 1 | 0 | 1 | 1 |
| TCF25 | 4 | 4 | -6,39366E-17 | 0,999999983 | 1 |
| ZC3HAV1 | 1 | 1 | -0,088386449 | 0,462790111 | 1 |
| SDHC | 4 | 4 | 5,64415E-15 | 0,999999806 | 1 |
| FAU | 1 | 1 | 0 | 1 | 1 |
| KCNJ11 | 2 | 2 | 0 | 1 | 1 |
| CPEB2 | 3 | 3 | 0 | 1 | 1 |
| CPEB3 | 2 | 2 | -7,47733E-16 | 0,999999937 | 1 |
| PSME3IP1 | 2 | 2 | 0 | 1 | 1 |
| ECHDC1 | 8 | 8 | 0 | 1 | 1 |
| SLC37A4 | 1 | 1 | 0,224986477 | 0,363284046 | 1 |
| ACP2 | 3 | 3 | -0,024597595 | 0,618560443 | 1 |
| JAKMIP3 | 6 | 6 | 0 | 1 | 1 |
| MAN2A2 | 2 | 2 | 0 | 1 | 1 |
| GABARAP | 1 | 1 | 0 | 1 | 1 |
| PLXNB2;PLXND1;PL | 1 | 1 | 0 | 1 | 1 |
| PLXNA1;PLXNA2 | 2 | 2 | 2,08315E-18 | 0,999999998 | 1 |
| NT5C1B | 1 | 1 | 4,64479E-20 | 1 | 1 |
| DET1 | 1 | 1 | 0 | 1 | 1 |
| RUNDC3A | 7 | 7 | 3,85702E-16 | 0,999999943 | 1 |
| NAP1L1;NAP1L4 | 2 | 2 | 0 | 1 | 1 |
| PELO | 2 | 2 | -4,65769E-17 | 0,999999985 | 1 |
| ALDH3B2;ALDH3B1 | 2 | 2 | -4,96559E-19 | 0,999999999 | 1 |
| MMP14 | 1 | 1 | 0 | 1 | 1 |
| UBE2H | 5 | 5 | 0 | 1 | 1 |
| SLC6A6 | 1 | 1 | 0 | 1 | 1 |
| SLC6A11 | 13 | 13 | 0 | 1 | 1 |
| MYCBP2 | 13 | 12 | -0,060850933 | 0,327967588 | 1 |
| CNRIP1 | 8 | 8 | 0,053580703 | 0,204629391 | 1 |
| GMDS | 9 | 9 | 2,61568E-18 | 0,999999998 | 1 |
| KCNA3 | 2 | 2 | 1,46684E-19 | 1 | 1 |

| | | | | | |
|---|---|---|---|---|---|
| SEC61B | 3 | 3 | 0 | 1 | 1 |
| WASF2;WASF1 | 1 | 1 | 0 | 1 | 1 |
| NSG2 | 1 | 1 | 0 | 1 | 1 |
| CLDN11 | 3 | 3 | 0 | 1 | 1 |
| ANKLE2 | 1 | 1 | 0,148158936 | 0,157970159 | 1 |
| HK2;HK1 | 3 | 3 | 0,093115001 | 0,13142436 | 1 |
| GM3839 | 30 | 30 | -6,9108E-16 | 0,999999953 | 1 |
| EGR4 | 1 | 1 | 0 | 1 | 1 |
| SLC25A22;SLC25A1 | 3 | 3 | 2,70819E-18 | 0,999999997 | 1 |
| MSTO1 | 3 | 3 | -0,035380862 | 0,337933991 | 1 |
| HTRA1 | 5 | 5 | 0 | 1 | 1 |
| CADM1 | 3 | 3 | -6,72869E-18 | 0,999999993 | 1 |
| EIF2B4 | 5 | 5 | -0,036137154 | 0,315351634 | 1 |
| PIKFYVE | 8 | 8 | 6,53313E-19 | 0,999999999 | 1 |
| DDX3Y | 5 | 5 | -0,045381621 | 0,39140953 | 1 |
| ATP8A1;ATP8B1 | 1 | 1 | 0 | 1 | 1 |
| GYS1 | 8 | 8 | 1,62354E-17 | 0,999999988 | 1 |
| ANKRD17 | 2 | 2 | 5,27301E-18 | 0,999999997 | 1 |
| TMEM151A | 1 | 1 | 0 | 1 | 1 |
| YBX3 | 1 | 1 | 0 | 1 | 1 |
| MBP | 1 | 1 | 0 | 1 | 1 |
| SLC12A5;SLC12A7 | 1 | 1 | 0 | 1 | 1 |
| GNL2 | 1 | 1 | 0 | 1 | 1 |
| TBC1D5 | 5 | 5 | 1,53218E-18 | 0,999999999 | 1 |
| SHF | 2 | 2 | 0 | 1 | 1 |
| TOPBP1 | 1 | 1 | 0 | 1 | 1 |
| SDSL | 3 | 3 | -4,90865E-09 | 0,999814433 | 1 |
| DTYMK | 6 | 5 | 6,82196E-17 | 0,999999989 | 1 |
| CSTF2 | 1 | 1 | -6,40798E-20 | 1 | 1 |
| PTPRD | 2 | 2 | -1,34946E-17 | 0,999999991 | 1 |
| GFPT2 | 4 | 4 | 0 | 1 | 1 |
| GYG1 | 4 | 4 | 0 | 1 | 1 |
| RPS6KA3 | 3 | 3 | 8,46895E-18 | 0,999999994 | 1 |

| | | | | | |
|---|---|---|---|---|---|
| RAB8A;RAB8B | 3 | 3 | 0 | 1 | 1 |
| MAP4K5 | 3 | 3 | -0,127264181 | 0,24583897 | 1 |
| CSNK1E | 4 | 4 | -1,91254E-15 | 0,999999917 | 1 |
| SIRT5 | 11 | 11 | -2,58842E-16 | 0,999999951 | 1 |
| ACOT13 | 7 | 7 | 0,098106768 | 0,13251146 | 1 |
| SHC3 | 3 | 3 | -0,077177872 | 0,224111602 | 1 |
| LUZP1 | 8 | 8 | 0 | 1 | 1 |
| TDRP | 2 | 2 | -2,26028E-16 | 0,999999971 | 1 |
| MYO1C | 4 | 4 | 0 | 1 | 1 |
| RAB5B | 6 | 6 | -2,91563E-17 | 0,999999986 | 1 |
| CRYZL2 | 7 | 5 | 0 | 1 | 1 |
| UCHL5 | 6 | 6 | 0 | 1 | 1 |
| SDC3 | 2 | 2 | -1,59657E-17 | 0,999999997 | 1 |
| SEPHS2 | 3 | 3 | -4,68711E-16 | 0,999999946 | 1 |
| EPS8L1 | 1 | 1 | 0 | 1 | 1 |
| TAGLN3;TAGLN2 | 4 | 4 | -0,103000544 | 0,47009307 | 1 |
| STUM | 1 | 1 | 0,07418157 | 0,258379115 | 1 |
| SLC25A4;SLC25A5 | 6 | 6 | 0,059230394 | 0,334388255 | 1 |
| MBP | 2 | 2 | -3,56482E-19 | 1 | 1 |
| CELF1;CELF2 | 2 | 2 | -1,01551E-15 | 0,999999936 | 1 |
| FERMT3 | 6 | 6 | 0,027510389 | 0,47075167 | 1 |
| UHRF1BP1 | 1 | 1 | 0 | 1 | 1 |
| EIF1;EIF1B | 3 | 3 | 1,05779E-19 | 1 | 1 |
| GABRB2 | 10 | 10 | -0,008101198 | 0,719496861 | 1 |
| TSR2 | 4 | 4 | -1,40323E-17 | 0,999999994 | 1 |
| KCNQ5 | 2 | 2 | 0 | 1 | 1 |
| LSM3 | 2 | 2 | 0 | 1 | 1 |
| SEC11A | 3 | 3 | 0 | 1 | 1 |
| PRRT1 | 4 | 4 | 2,29841E-18 | 0,999999996 | 1 |
| CAMK4 | 2 | 2 | 0 | 1 | 1 |
| SLC25A25 | 15 | 14 | 0,009499246 | 0,625050882 | 1 |
| ARPIN | 4 | 4 | 0 | 1 | 1 |
| IMMT | 2 | 2 | 0 | 1 | 1 |

| | | | | | |
|---|---|---|---|---|---|
| FBXO3 | 10 | 10 | -7,48207E-17 | 0,999999978 | 1 |
| GMPR | 11 | 11 | 4,39163E-14 | 0,999999203 | 1 |
| MTX3 | 9 | 9 | -1,85554E-16 | 0,999999963 | 1 |
| IL33 | 1 | 1 | 0,208230338 | 0,233590064 | 1 |
| DDX3X | 4 | 4 | 0,06033333 | 0,336251734 | 1 |
| TBCC | 5 | 5 | -0,079507138 | 0,236056405 | 1 |
| RUFY3 | 19 | 19 | -1,33064E-17 | 0,999999989 | 1 |
| DNPH1 | 1 | 1 | 0 | 1 | 1 |
| RTN4IP1 | 9 | 8 | 5,78678E-18 | 0,999999997 | 1 |
| CSK | 6 | 6 | 3,10148E-18 | 0,999999999 | 1 |
| ADCY3 | 5 | 5 | 0,059016335 | 0,389701929 | 1 |
| ZDHHC15 | 1 | 1 | 0 | 1 | 1 |
| PIGT | 3 | 3 | -2,92081E-18 | 0,999999996 | 1 |
| AK1 | 17 | 17 | 0 | 1 | 1 |
| ADCY1 | 10 | 10 | -0,001001722 | 0,888181847 | 1 |
| CD99L2 | 2 | 2 | -0,040260931 | 0,585964014 | 1 |
| CLEC16A | 2 | 2 | 8,93489E-18 | 0,999999995 | 1 |
| SLC9A3R2 | 6 | 6 | 1,79716E-17 | 0,999999996 | 1 |
| SLC22A23 | 6 | 6 | -2,99314E-18 | 0,999999997 | 1 |
| PTGES3 | 6 | 6 | -5,56081E-16 | 0,999999973 | 1 |
| ANAPC1 | 4 | 4 | 3,09645E-17 | 0,999999985 | 1 |
| GNG3 | 2 | 2 | 0 | 1 | 1 |
| USP19 | 8 | 8 | 0,053938464 | 0,229041706 | 1 |
| PARS2 | 8 | 6 | 0 | 1 | 1 |
| COMTD1 | 2 | 2 | 0 | 1 | 1 |
| RNF150 | 1 | 1 | 0 | 1 | 1 |
| ACOT1;ACOT2 | 10 | 10 | 3,78329E-16 | 0,999999934 | 1 |
| SEPTIN4;SEPTIN5 | 4 | 4 | 0,084193771 | 0,215000391 | 1 |
| LSM8 | 2 | 2 | 5,77277E-17 | 0,999999982 | 1 |
| FKBP1B | 3 | 3 | 0,055654335 | 0,25621844 | 1 |
| EPPK1;PLEC | 3 | 3 | 0 | 1 | 1 |
| HNRNPD;HNRNPDL | 1 | 1 | -3,0085E-18 | 0,999999996 | 1 |
| MSI2 | 4 | 4 | -0,025593603 | 0,556031985 | 1 |

| | | | | | |
|---|---|---|---|---|---|
| RBFOX1 | 2 | 2 | 0 | 1 | 1 |
| RBFOX3 | 2 | 2 | -7,81803E-18 | 0,999999996 | 1 |
| GBA | 4 | 4 | 0 | 1 | 1 |
| VKORC1L1 | 1 | 1 | 0 | 1 | 1 |
| PREX2 | 6 | 6 | -7,67296E-19 | 0,999999999 | 1 |
| ITGAM;GM49368 | 1 | 1 | 0 | 1 | 1 |
| PTBP1 | 4 | 4 | 0,082257066 | 0,317883482 | 1 |
| PPIL3 | 2 | 2 | 0 | 1 | 1 |
| INPP5J | 6 | 6 | 0,070833222 | 0,163789151 | 1 |
| MMAB | 6 | 6 | 2,06258E-16 | 0,999999966 | 1 |
| CLIC1 | 6 | 6 | 1,15006E-17 | 0,999999993 | 1 |
| ANK3;ANK1;ANK2 | 2 | 2 | 4,33465E-16 | 0,999999952 | 1 |
| CCT6A;CCT6B | 5 | 5 | 1,33534E-18 | 0,999999999 | 1 |
| GATD3 | 13 | 13 | 3,65431E-17 | 0,99999999 | 1 |
| COX7A2 | 3 | 3 | 0,095972783 | 0,203033831 | 1 |
| CKM | 2 | 2 | -7,00318E-17 | 0,999999984 | 1 |
| CUTC | 5 | 5 | -1,2419E-17 | 0,999999997 | 1 |
| ATP4A | 2 | 2 | 1,50061E-15 | 0,999999905 | 1 |
| H2-D1 | 4 | 2 | 0,262501099 | 0,377433467 | 1 |
| ACBD5 | 5 | 5 | 0 | 1 | 1 |
| ADAM23 | 9 | 9 | 3,18017E-17 | 0,999999984 | 1 |
| PTER | 3 | 3 | -7,77834E-19 | 0,999999998 | 1 |
| UNC119 | 1 | 1 | 0 | 1 | 1 |
| PBX1 | 1 | 1 | 0 | 1 | 1 |
| MTMR2 | 12 | 12 | 0 | 1 | 1 |
| MERTK | 2 | 2 | 0 | 1 | 1 |
| AKAP8 | 2 | 2 | 0,056507484 | 0,374954274 | 1 |
| HSD17B10 | 5 | 5 | 1,88325E-16 | 0,999999964 | 1 |
| CNOT7 | 3 | 3 | -0,064852181 | 0,180819549 | 1 |
| DAG1 | 6 | 6 | -1,68342E-19 | 1 | 1 |
| PHYHD1 | 4 | 4 | 0,007296878 | 0,761732598 | 1 |
| CADPS | 1 | 1 | 1,30519E-19 | 1 | 1 |
| CNOT11 | 3 | 3 | -0,014064032 | 0,642986805 | 1 |

| | | | | | |
|---|---|---|---|---|---|
| EFNB1 | 2 | 2 | -2,78235E-16 | 0,999999988 | 1 |
| PLPBP | 13 | 12 | 6,4916E-18 | 0,999999997 | 1 |
| SYBU | 3 | 2 | 0,265914983 | 0,158170862 | 1 |
| PRKAR1B | 9 | 9 | -0,012090163 | 0,533422745 | 1 |
| HBEGF | 1 | 1 | 0 | 1 | 1 |
| TMEM163 | 5 | 5 | -3,17366E-17 | 0,999999991 | 1 |
| LMAN1 | 2 | 2 | -7,69207E-17 | 0,999999977 | 1 |
| SCRN2 | 5 | 5 | 2,82828E-16 | 0,999999967 | 1 |
| TUBB4B;TUBB5;TU | 2 | 2 | -3,10299E-16 | 0,999999981 | 1 |
| FLRT2 | 4 | 4 | 1,20854E-17 | 0,999999996 | 1 |
| DGUOK | 3 | 3 | 0 | 1 | 1 |
| DCHS1 | 1 | 1 | 0 | 1 | 1 |
| CBR1;CBR3 | 1 | 1 | -8,62542E-21 | 1 | 1 |
| NIPA1 | 3 | 3 | 0 | 1 | 1 |
| HECW1 | 4 | 4 | 0 | 1 | 1 |
| ETFB | 8 | 8 | 0,050458689 | 0,158521886 | 1 |
| CTPS2 | 4 | 4 | -3,96058E-16 | 0,999999949 | 1 |
| SERAC1 | 5 | 4 | 0 | 1 | 1 |
| PIK3C3 | 10 | 10 | -0,0275261 | 0,354127489 | 1 |
| CYTH3;CYTH2;CYTH | 1 | 1 | 0 | 1 | 1 |
| SLC25A35 | 4 | 4 | -1,66228E-17 | 0,999999998 | 1 |
| SVIP | 2 | 2 | 3,11426E-16 | 0,999999969 | 1 |
| YME1L1 | 10 | 10 | 0,04145152 | 0,309164049 | 1 |
| CDS1 | 2 | 2 | -1,55831E-15 | 0,999999921 | 1 |
| CYTH2 | 4 | 4 | 1,34099E-14 | 0,999999718 | 1 |
| ABCA5 | 3 | 3 | -0,117295856 | 0,146463347 | 1 |
| NAP1L1 | 10 | 9 | 0 | 1 | 1 |
| SCYL1 | 2 | 2 | 4,67707E-18 | 0,999999997 | 1 |
| TKTL2 | 1 | 1 | 0 | 1 | 1 |
| CDHR4 | 1 | 1 | 0 | 1 | 1 |
| SFPQ | 2 | 2 | 0 | 1 | 1 |
| NPEPL1 | 3 | 3 | 1,8288E-15 | 0,999999883 | 1 |
| AKAP1 | 2 | 2 | 0 | 1 | 1 |

| | | | | | |
|---|---|---|---|---|---|
| IMPDH1 | 3 | 3 | 0 | 1 | 1 |
| UBFD1 | 8 | 7 | -0,062524383 | 0,177485315 | 1 |
| GALNT17 | 7 | 7 | 0 | 1 | 1 |
| PGRMC1;PGRMC2 | 1 | 1 | 0 | 1 | 1 |
| ARHGAP33 | 3 | 2 | 0 | 1 | 1 |
| NEDD4;NEDD4L | 1 | 1 | 0 | 1 | 1 |
| PLBD2 | 5 | 5 | 0 | 1 | 1 |
| MRPL47 | 2 | 2 | 2,09416E-18 | 0,999999999 | 1 |
| RMND5A | 3 | 3 | 0 | 1 | 1 |
| SLC25A33 | 2 | 2 | 0 | 1 | 1 |
| SMYD5 | 2 | 2 | -1,94891E-18 | 0,999999998 | 1 |
| DAPK1 | 2 | 1 | 0 | 1 | 1 |
| THA1 | 3 | 2 | 0,043330107 | 0,547714257 | 1 |
| ZDHHC5 | 7 | 7 | 0,026275265 | 0,541352076 | 1 |
| CLNS1A | 2 | 2 | -4,65494E-16 | 0,999999976 | 1 |
| ENOX1 | 1 | 1 | 1,18625E-18 | 0,999999998 | 1 |
| CORO2B | 19 | 19 | -7,29236E-17 | 0,999999987 | 1 |
| PLGRKT | 3 | 3 | 0,032818363 | 0,388117277 | 1 |
| RAPGEF6 | 3 | 3 | 0 | 1 | 1 |
| RAE1 | 3 | 3 | 0 | 1 | 1 |
| NUDT18 | 1 | 1 | 0 | 1 | 1 |
| EMC10 | 1 | 1 | 0,200658803 | 0,247651945 | 1 |
| SPATA5;KATNAL2 | 1 | 1 | 1,22406E-17 | 0,999999995 | 1 |
| METTL3 | 1 | 1 | 0 | 1 | 1 |
| BORCS5 | 4 | 4 | 1,25056E-14 | 0,999999643 | 1 |
| MPDU1 | 2 | 2 | -3,49492E-18 | 0,999999996 | 1 |
| C1QL3 | 2 | 2 | 0 | 1 | 1 |
| TRUB2 | 1 | 1 | -2,56596E-19 | 0,999999999 | 1 |
| ELAVL2 | 7 | 7 | -1,48766E-17 | 0,99999999 | 1 |
| PTPRK | 4 | 4 | 0 | 1 | 1 |
| COL11A2 | 1 | 1 | 0,171571462 | 0,369467403 | 1 |
| CTNND2;PKP4 | 4 | 4 | 0 | 1 | 1 |
| MMP17 | 2 | 2 | -5,77441E-16 | 0,999999965 | 1 |

| | | | | | |
|---|---|---|---|---|---|
| MRTFB | 8 | 7 | -0,01523247 | 0,622691251 | 1 |
| CDC34 | 2 | 2 | 0,049078156 | 0,406040905 | 1 |
| WNK1;WNK3 | 1 | 1 | -0,043213638 | 0,504218872 | 1 |
| WNK1;WNK2;WNK | 1 | 1 | -0,029904327 | 0,545071228 | 1 |
| KCNA4 | 5 | 5 | 4,20103E-16 | 0,999999944 | 1 |
| KCNA10;KCNA1;KC | 1 | 1 | 0 | 1 | 1 |
| CNNM3 | 8 | 8 | -0,040188154 | 0,330957107 | 1 |
| EVI5L | 9 | 9 | 4,4785E-18 | 0,999999997 | 1 |
| MVD | 4 | 4 | 0 | 1 | 1 |
| NDUFB3 | 6 | 6 | 0,041602645 | 0,339431247 | 1 |
| PLP1 | 8 | 8 | -2,83239E-17 | 0,999999993 | 1 |
| ALDH3A1 | 3 | 2 | 0 | 1 | 1 |
| LRRC4 | 2 | 2 | 1,06207E-18 | 0,999999999 | 1 |
| STXBP5L | 6 | 5 | 5,59828E-18 | 0,999999998 | 1 |
| SLIT1 | 3 | 3 | -7,41621E-18 | 0,999999995 | 1 |
| SHANK1 | 1 | 1 | 0 | 1 | 1 |
| PPP3CC | 2 | 2 | 0,009622887 | 0,767642175 | 1 |
| SLC25A36 | 1 | 1 | 0 | 1 | 1 |
| ALS2CL | 1 | 1 | 0 | 1 | 1 |
| RPAP3 | 2 | 2 | -1,66201E-15 | 0,999999906 | 1 |
| VPS37B | 3 | 3 | 0 | 1 | 1 |
| SCARB2 | 4 | 4 | -3,28791E-19 | 1 | 1 |
| BRSK2 | 4 | 4 | 3,20071E-17 | 0,999999995 | 1 |
| STK11 | 5 | 5 | -1,17383E-14 | 0,999999862 | 1 |
| COX5A | 7 | 7 | 0,031998748 | 0,380166489 | 1 |
| TEX26 | 1 | 1 | 0 | 1 | 1 |
| FMR1 | 2 | 2 | -0,002684421 | 0,860182628 | 1 |
| EFTUD2 | 5 | 5 | -8,22482E-18 | 0,999999994 | 1 |
| SLC36A1 | 1 | 1 | 0 | 1 | 1 |
| IL1RAP | 3 | 3 | -0,007728086 | 0,757607727 | 1 |
| SLC25A4;SLC25A5;S | 4 | 3 | 0 | 1 | 1 |
| SHISA9 | 2 | 2 | -0,035150627 | 0,470088224 | 1 |
| MATK | 4 | 4 | -1,81413E-17 | 0,999999992 | 1 |

| | | | | | |
|---|---|---|---|---|---|
| LSM6 | 4 | 4 | -1,12154E-15 | 0,999999924 | 1 |
| SLC44A2 | 3 | 3 | 0 | 1 | 1 |
| ENO1;ENO2 | 1 | 1 | 0,084246813 | 0,220861887 | 1 |
| SSB | 10 | 10 | 0 | 1 | 1 |
| EDIL3 | 4 | 4 | -7,07002E-16 | 0,999999931 | 1 |
| FBXO22 | 10 | 10 | -0,005438271 | 0,764880317 | 1 |
| NLGN4L;NLGN2 | 2 | 2 | 2,09036E-18 | 0,999999998 | 1 |
| GIPC1 | 9 | 9 | -0,013611184 | 0,514120571 | 1 |
| SYNJ1 | 1 | 1 | 0 | 1 | 1 |
| TNFRSF21 | 6 | 6 | 0,032673484 | 0,407500938 | 1 |
| PPIH | 3 | 3 | -0,016735178 | 0,611331496 | 1 |
| PDLIM5 | 4 | 4 | -2,05267E-18 | 0,999999998 | 1 |
| CPPED1 | 5 | 5 | -0,042992298 | 0,338723257 | 1 |
| KRT77 | 4 | 2 | -0,141248399 | 0,392623444 | 1 |
| OSGEP | 1 | 1 | -5,95235E-20 | 1 | 1 |
| NRGN | 2 | 2 | 1,10288E-18 | 0,999999999 | 1 |
| PEX6 | 1 | 1 | 0 | 1 | 1 |
| BCKDHA | 3 | 2 | -0,070863305 | 0,281053328 | 1 |
| MAGED2 | 1 | 1 | -0,095688565 | 0,335691616 | 1 |
| LPIN2 | 4 | 4 | 0 | 1 | 1 |
| HCFC1 | 6 | 6 | 0 | 1 | 1 |
| MTND1 | 7 | 7 | 4,0124E-18 | 0,999999998 | 1 |
| UBE2J1 | 3 | 3 | 6,93435E-20 | 1 | 1 |
| RNF24 | 1 | 1 | 0,075926046 | 0,299561561 | 1 |
| SYNJ2BP | 6 | 6 | 1,90889E-17 | 0,999999996 | 1 |
| RPRD2 | 1 | 1 | 0 | 1 | 1 |
| GRIA1 | 1 | 1 | 2,40137E-15 | 0,999999887 | 1 |
| LRFN4 | 4 | 4 | 0 | 1 | 1 |
| TTC9 | 1 | 1 | -2,84935E-22 | 1 | 1 |
| TMED2 | 4 | 4 | 0 | 1 | 1 |
| P2RX7 | 2 | 2 | 0 | 1 | 1 |
| CFAP298 | 1 | 1 | 0 | 1 | 1 |
| OPCML | 2 | 2 | 0,08384313 | 0,132681971 | 1 |

| | | | | | |
|---|---|---|---|---|---|
| CSTF2 | 1 | 1 | -0,080038475 | 0,437651277 | 1 |
| COMT | 5 | 5 | 0 | 1 | 1 |
| NUDT17 | 2 | 2 | -2,44958E-15 | 0,999999917 | 1 |
| GSTT3 | 1 | 1 | -6,64697E-20 | 1 | 1 |
| FKBP1A | 8 | 8 | 0 | 1 | 1 |
| SDK2 | 9 | 7 | 4,1426E-16 | 0,99999994 | 1 |
| ITPA | 8 | 8 | -3,37739E-16 | 0,999999966 | 1 |
| KRT14;KRT42 | 6 | 4 | 4,26057E-18 | 0,999999998 | 1 |
| RUNDC3B | 4 | 4 | 0 | 1 | 1 |
| GOLGA5 | 5 | 3 | -0,268654931 | 0,537433755 | 1 |
| KRT6A;KRT5 | 3 | 2 | 5,3701E-15 | 0,999999926 | 1 |
| DRG1;DRG2 | 1 | 1 | 0 | 1 | 1 |
| BOLA1 | 3 | 3 | -1,73056E-18 | 0,999999998 | 1 |
| RPL23 | 4 | 4 | 7,171E-19 | 0,999999999 | 1 |
| CNTN4 | 5 | 5 | -0,031629156 | 0,506737944 | 1 |
| PES1 | 1 | 1 | 0 | 1 | 1 |
| NANS | 8 | 8 | 2,02377E-14 | 0,999999507 | 1 |
| S100A1 | 1 | 1 | 0 | 1 | 1 |
| FGL1 | 1 | 1 | 0 | 1 | 1 |
| SEMA6C | 1 | 1 | 0 | 1 | 1 |
| RRAS2 | 3 | 3 | 1,42682E-16 | 0,999999972 | 1 |
| RERG | 1 | 1 | -0,057421586 | 0,433371973 | 1 |
| PLPP6 | 2 | 2 | -0,009946162 | 0,765156548 | 1 |
| DCAF11 | 5 | 5 | -0,007681588 | 0,732971805 | 1 |
| NAT14 | 1 | 1 | 0 | 1 | 1 |
| EMC4 | 2 | 2 | -1,07454E-18 | 1 | 1 |
| PPP1R3D | 2 | 2 | -4,01974E-19 | 0,999999999 | 1 |
| SNTB1 | 2 | 2 | -5,96458E-18 | 0,999999996 | 1 |
| DCHS2 | 1 | 1 | -0,304442367 | 0,216167048 | 1 |
| EFR3A | 15 | 14 | -9,26569E-17 | 0,999999974 | 1 |
| FERMT2 | 1 | 1 | 7,32267E-20 | 1 | 1 |
| IFT22 | 4 | 4 | 0 | 1 | 1 |
| PTAR1 | 3 | 3 | -0,049575295 | 0,406840572 | 1 |

| | | | | | |
|---|---|---|---|---|---|
| SRC;FYN | 3 | 3 | 1,36314E-17 | 0,999999993 | 1 |
| S100A13 | 2 | 2 | -1,71432E-18 | 0,999999998 | 1 |
| VPS36 | 9 | 9 | -1,33554E-12 | 0,999995491 | 1 |
| UBE3C | 11 | 11 | -5,55286E-16 | 0,999999928 | 1 |
| OPALIN | 2 | 2 | -4,73992E-17 | 0,999999992 | 1 |
| RFT1 | 1 | 1 | -0,171372362 | 0,283939039 | 1 |
| CDK9 | 1 | 1 | 0 | 1 | 1 |
| AHCYL2 | 1 | 1 | 0 | 1 | 1 |
| TECPR2 | 7 | 7 | 4,64296E-19 | 1 | 1 |
| SGPL1 | 5 | 5 | -0,041159782 | 0,473936437 | 1 |
| CAMK2G | 6 | 6 | 2,21345E-17 | 0,999999991 | 1 |
| MYCBP2 | 4 | 4 | -3,75323E-18 | 0,999999997 | 1 |
| MRPS36 | 4 | 4 | 1,38612E-19 | 1 | 1 |
| PLA2G7 | 5 | 5 | -9,55939E-18 | 0,999999996 | 1 |
| SCP2 | 6 | 5 | -0,043472519 | 0,274918197 | 1 |
| TPCN1 | 1 | 1 | 0 | 1 | 1 |
| AGFG2 | 5 | 5 | -3,53378E-05 | 0,980285817 | 1 |
| MAP4K4;MINK1 | 1 | 1 | 0 | 1 | 1 |
| F11R | 2 | 2 | 2,43711E-14 | 0,999999665 | 1 |
| HACD2 | 2 | 2 | -0,14315678 | 0,170852186 | 1 |
| PISD | 2 | 2 | 0 | 1 | 1 |
| GAS8 | 2 | 2 | 0 | 1 | 1 |
| TMEM254 | 1 | 1 | 0 | 1 | 1 |
| FKBP2 | 1 | 1 | 0 | 1 | 1 |
| DDX17 | 4 | 4 | -0,098919739 | 0,175817008 | 1 |
| SCN1B | 6 | 6 | 0,006619968 | 0,710177179 | 1 |
| HBB-BS | 4 | 4 | 0,058354522 | 0,222043193 | 1 |
| GPRASP1 | 1 | 1 | 0 | 1 | 1 |
| EMB | 4 | 3 | 0,026603258 | 0,444973471 | 1 |
| SFT2D3 | 1 | 1 | 0 | 1 | 1 |
| ELP3 | 5 | 5 | -2,61353E-18 | 0,999999999 | 1 |
| DNAAF10 | 1 | 1 | -0,017405362 | 0,735992326 | 1 |
| SYT17 | 13 | 13 | 0,063783649 | 0,192579849 | 1 |

| | | | | | |
|---|---|---|---|---|---|
| SLC30A6 | 1 | 1 | 0 | 1 | 1 |
| MARK3;MARK2 | 1 | 1 | -0,128952973 | 0,402745603 | 1 |
| ABI2 | 1 | 1 | 0 | 1 | 1 |
| SHISA7 | 13 | 13 | 1,33572E-17 | 0,99999999 | 1 |
| MRPL24 | 2 | 1 | 0 | 1 | 1 |
| MELK | 2 | 2 | -0,048454559 | 0,388526384 | 1 |
| KIF13A | 1 | 1 | 0 | 1 | 1 |
| GTF2F1 | 1 | 1 | 0 | 1 | 1 |
| ACVR2A | 1 | 1 | -0,039186669 | 0,619182092 | 1 |
| SERPINB6B | 1 | 1 | 0 | 1 | 1 |
| GAD2 | 11 | 10 | 0 | 1 | 1 |
| NCOA7 | 8 | 8 | 0 | 1 | 1 |
| FAM81A | 9 | 7 | -4,27756E-18 | 0,999999996 | 1 |
| PHF24 | 4 | 4 | 1,06271E-15 | 0,999999961 | 1 |
| ACYP2 | 4 | 4 | 1,36075E-20 | 1 | 1 |
| EEF1AKMT2 | 3 | 3 | -3,13428E-17 | 0,999999988 | 1 |
| TMEM178B | 4 | 4 | 0 | 1 | 1 |
| KIF3C | 1 | 1 | -0,072701717 | 0,373548101 | 1 |
| ABHD3 | 4 | 3 | -0,129519847 | 0,247365419 | 1 |
| DHPS | 4 | 4 | 0 | 1 | 1 |
| HNRNPM | 1 | 1 | 0 | 1 | 1 |
| SNRPF | 1 | 1 | 0 | 1 | 1 |
| PDAP1 | 9 | 9 | -1,45466E-16 | 0,999999979 | 1 |
| UBE2V1;UBE2V2 | 8 | 8 | -6,79361E-17 | 0,999999989 | 1 |
| FAHD1 | 9 | 7 | 0,067784926 | 0,192039417 | 1 |
| PLLP | 2 | 2 | -1,1614E-17 | 0,999999997 | 1 |
| TNFRSF14 | 1 | 1 | 0 | 1 | 1 |
| PLEKHA5 | 4 | 3 | 7,68924E-19 | 0,999999999 | 1 |
| PTPMT1 | 7 | 7 | 0,062780583 | 0,209377617 | 1 |
| FER1L4 | 1 | 1 | 0 | 1 | 1 |
| ANAPC2 | 2 | 2 | -1,62329E-19 | 1 | 1 |
| TPD52L2 | 9 | 8 | 0 | 1 | 1 |
| NUDT8 | 1 | 1 | 0 | 1 | 1 |

| | | | | | |
|---|---|---|---|---|---|
| SBF1 | 3 | 3 | 0,04070796 | 0,507135492 | 1 |
| FABP1 | 1 | 1 | 0 | 1 | 1 |
| PPP1CA | 7 | 7 | -9,39506E-18 | 0,999999995 | 1 |
| MGRN1 | 3 | 3 | -1,16582E-17 | 0,999999994 | 1 |
| MRPL15 | 4 | 3 | -4,4874E-17 | 0,999999987 | 1 |
| CLSTN2 | 1 | 1 | 0 | 1 | 1 |
| SERPINH1 | 2 | 2 | -2,53613E-17 | 0,999999991 | 1 |
| CCP110 | 2 | 2 | -0,108301036 | 0,316802149 | 1 |
| AP1G2 | 1 | 1 | 0 | 1 | 1 |
| CAB39L | 5 | 5 | 0 | 1 | 1 |
| TOR1A | 2 | 1 | 0,009754545 | 0,832981682 | 1 |
| TIA1 | 1 | 1 | 0 | 1 | 1 |
| CALM3 | 1 | 1 | 0 | 1 | 1 |
| APOO | 7 | 6 | -6,39809E-17 | 0,999999986 | 1 |
| SEMA4B | 4 | 4 | -0,008226545 | 0,72559631 | 1 |
| SLC5A7 | 5 | 5 | 2,46899E-16 | 0,999999956 | 1 |
| AHSG | 5 | 5 | 2,44476E-16 | 0,999999966 | 1 |
| AK9 | 1 | 1 | -0,063953371 | 0,293282533 | 1 |
| NUMB | 5 | 5 | 0,016515898 | 0,549869115 | 1 |
| CNTNAP5A | 3 | 3 | 0 | 1 | 1 |
| CNTNAP5C | 1 | 1 | 0 | 1 | 1 |
| PRXL2B | 6 | 6 | -8,76392E-18 | 0,999999994 | 1 |
| NEK9 | 5 | 5 | -0,065556202 | 0,169243853 | 1 |
| DLG3;DLG1 | 2 | 2 | 0 | 1 | 1 |
| SYT2 | 7 | 7 | -3,48344E-17 | 0,999999981 | 1 |
| ASPHD2 | 4 | 4 | -0,008430886 | 0,765716179 | 1 |
| TSPAN7 | 3 | 3 | 2,15007E-18 | 0,999999999 | 1 |
| ERCC5 | 1 | 1 | 0 | 1 | 1 |
| MRPS17 | 2 | 2 | 0,135365945 | 0,22790493 | 1 |
| MAPK14 | 2 | 2 | 0 | 1 | 1 |
| CAMKMT | 1 | 1 | 0 | 1 | 1 |
| WASF1 | 13 | 13 | -1,3625E-16 | 0,999999981 | 1 |
| NOCT | 2 | 1 | 0 | 1 | 1 |

| | | | | | |
|---|---|---|---|---|---|
| EIF5B | 3 | 3 | 0 | 1 | 1 |
| LY6H | 8 | 8 | -1,954E-16 | 0,999999982 | 1 |
| SORBS1 | 14 | 13 | -0,022066504 | 0,376779167 | 1 |
| CUZD1 | 1 | 1 | -0,283077037 | 0,238139423 | 1 |
| MKNK2 | 1 | 1 | 0 | 1 | 1 |
| CD38 | 3 | 2 | 0,04890262 | 0,269916651 | 1 |
| PRL8A1 | 1 | 1 | 0 | 1 | 1 |
| BCKDHA | 10 | 10 | 0,002532028 | 0,795363011 | 1 |
| CYB5B | 8 | 8 | 7,79834E-15 | 0,999999685 | 1 |
| POMT1 | 1 | 1 | 0 | 1 | 1 |
| PABIR1 | 3 | 3 | 2,6485E-18 | 0,999999997 | 1 |
| EIF6 | 3 | 3 | -1,50852E-16 | 0,999999974 | 1 |
| ACAT2 | 1 | 1 | 0 | 1 | 1 |
| NRXN2 | 1 | 1 | -1,68859E-21 | 1 | 1 |
| RPL21 | 6 | 6 | -2,90384E-16 | 0,999999958 | 1 |
| RAC2 | 5 | 5 | -5,43393E-17 | 0,999999981 | 1 |
| AKR1B10 | 4 | 4 | 2,62079E-17 | 0,999999989 | 1 |
| PRMT9 | 1 | 1 | 0 | 1 | 1 |
| SYNJ1 | 3 | 3 | 7,39505E-19 | 0,999999999 | 1 |
| MRPS14 | 1 | 1 | 0 | 1 | 1 |
| BID | 1 | 1 | 0,107038963 | 0,320037554 | 1 |
| CACNA1C;CACNA1S | 1 | 1 | 0 | 1 | 1 |
| SCLT1 | 1 | 1 | 0,026121622 | 0,689828461 | 1 |
| FUT8 | 1 | 1 | 0 | 1 | 1 |
| CWF19L2 | 1 | 1 | 0 | 1 | 1 |
| CCDC68 | 1 | 1 | 0 | 1 | 1 |
| VAMP4 | 2 | 2 | 0 | 1 | 1 |
| POU2F3 | 1 | 1 | 0 | 1 | 1 |
| SLC7A9 | 1 | 1 | 0 | 1 | 1 |
| SV2C | 2 | 1 | 0 | 1 | 1 |
| VIM;INA | 1 | 1 | 0,127066172 | 0,447564655 | 1 |
| OSBPL10 | 6 | 6 | -0,075508581 | 0,192991277 | 1 |
| EBP | 1 | 1 | 0 | 1 | 1 |

| | | | | | |
|---|---|---|---|---|---|
| VCPKMT | 1 | 1 | 0 | 1 | 1 |
| NUDT5 | 4 | 4 | -0,038378261 | 0,328818809 | 1 |
| CLCN7 | 2 | 2 | 6,46722E-18 | 0,999999995 | 1 |
| VPS26B | 7 | 7 | 2,26716E-19 | 1 | 1 |
| DLG2 | 8 | 7 | -9,8973E-14 | 0,999999079 | 1 |
| CISD2 | 3 | 3 | -6,70135E-15 | 0,999999869 | 1 |
| CAB39;CAB39L | 4 | 4 | -1,93591E-18 | 0,999999997 | 1 |
| GSTM2 | 6 | 6 | 3,6659E-17 | 0,999999991 | 1 |
| FOXK1 | 2 | 2 | -0,06319163 | 0,301336907 | 1 |
| SETD2 | 1 | 1 | -0,042316185 | 0,51560848 | 1 |
| PTK7 | 3 | 2 | -3,22641E-19 | 0,999999999 | 1 |
| PDS5B | 1 | 1 | -0,060983697 | 0,566715444 | 1 |
| ZCCHC2 | 1 | 1 | 0 | 1 | 1 |
| AKT1;AKT2 | 1 | 1 | 0 | 1 | 1 |
| KCNC4 | 2 | 2 | -1,68965E-18 | 0,999999997 | 1 |
| TBCE | 8 | 7 | -0,03940844 | 0,405742558 | 1 |
| CAMK2G;CAMK2B | 5 | 5 | 2,92957E-16 | 0,99999996 | 1 |
| ATG16L1 | 6 | 6 | -1,74253E-16 | 0,999999963 | 1 |
| MOBP | 2 | 2 | 0 | 1 | 1 |
| KCNIP4 | 2 | 2 | -0,051778131 | 0,395238783 | 1 |
| PLCB3 | 6 | 6 | -0,053402345 | 0,249469366 | 1 |
| GM45623 | 1 | 1 | 0 | 1 | 1 |
| STAM2;STAM | 1 | 1 | 0 | 1 | 1 |
| HNRNPF | 4 | 4 | -5,28615E-17 | 0,999999985 | 1 |
| FST | 1 | 1 | 5,26456E-17 | 0,999999988 | 1 |
| AGPAT1 | 4 | 4 | 6,1526E-17 | 0,999999986 | 1 |
| MRPS28 | 2 | 2 | 0 | 1 | 1 |
| TRP53I11 | 4 | 4 | 0 | 1 | 1 |
| ULK2 | 2 | 2 | -0,009360162 | 0,769008424 | 1 |
| TNFRSF11B | 1 | 1 | 0 | 1 | 1 |
| PCDHAC2 | 3 | 3 | 0 | 1 | 1 |
| ARHGAP12 | 1 | 1 | 0 | 1 | 1 |
| TRMT61A | 2 | 2 | -3,58595E-17 | 0,999999988 | 1 |

| | | | | | |
|---|---|---|---|---|---|
| VMN2R98;VMN2R1 | 1 | 1 | -1,65832E-19 | 1 | 1 |
| PLP2 | 1 | 1 | 0 | 1 | 1 |
| ARHGEF26 | 3 | 3 | 3,85593E-18 | 0,999999999 | 1 |
| XIRP2 | 1 | 1 | -2,79598E-15 | 0,999999894 | 1 |
| M6PR | 4 | 4 | -0,019251564 | 0,582516656 | 1 |
| TRPM1 | 1 | 1 | 0 | 1 | 1 |
| HELZ2 | 2 | 2 | 0,004401708 | 0,847123941 | 1 |
| DCAF6 | 1 | 1 | -0,01940889 | 0,67284226 | 1 |
| TMEM63C | 4 | 4 | 0,009769237 | 0,71751489 | 1 |
| AMPD3 | 6 | 6 | -2,40044E-17 | 0,999999994 | 1 |
| LRFN3 | 4 | 4 | -2,25367E-16 | 0,999999977 | 1 |
| PIGK | 4 | 4 | -8,15915E-19 | 0,999999999 | 1 |
| SCAPER | 2 | 2 | 0 | 1 | 1 |
| FETUB | 1 | 1 | 0,113429607 | 0,166511326 | 1 |
| RHOT1 | 1 | 1 | 0 | 1 | 1 |
| CTSO | 1 | 1 | 0 | 1 | 1 |
| MIPEP | 6 | 6 | 0,038502289 | 0,391478296 | 1 |
| INPP4A | 9 | 9 | -7,15602E-18 | 0,999999993 | 1 |
| FOXRED2 | 1 | 1 | 0 | 1 | 1 |
| KLHL14 | 1 | 1 | 0,044439487 | 0,613862851 | 1 |
| KIF14 | 1 | 1 | 0 | 1 | 1 |
| DNAH1 | 1 | 1 | -9,52383E-19 | 0,999999999 | 1 |
| ELMO1 | 10 | 10 | -0,003009701 | 0,8015689 | 1 |
| PTDSS2 | 2 | 2 | -9,14747E-18 | 0,999999994 | 1 |
| DCTN3 | 7 | 7 | -0,025437634 | 0,503330466 | 1 |
| FAM53B | 1 | 1 | 0 | 1 | 1 |
| SAMD9L | 1 | 1 | -0,129768573 | 0,1531993 | 1 |
| SLC25A24 | 5 | 3 | 0 | 1 | 1 |
| AGTPBP1 | 8 | 8 | -0,028348141 | 0,396013719 | 1 |
| NF2 | 2 | 2 | 0 | 1 | 1 |
| IMPACT | 16 | 16 | 5,4574E-16 | 0,999999922 | 1 |
| MFN1 | 3 | 3 | 0,307335798 | 0,999419808 | 1 |
| COMMD2 | 4 | 4 | 0 | 1 | 1 |

| | | | | | |
|---|---|---|---|---|---|
| CLDND1 | 4 | 4 | -0,040858022 | 0,323618264 | 1 |
| TOP3B | 1 | 1 | 0 | 1 | 1 |
| KBTBD2 | 3 | 3 | -0,04225003 | 0,42363646 | 1 |
| USP46;USP12 | 3 | 3 | -1,74786E-18 | 0,999999999 | 1 |
| CD81 | 5 | 5 | 3,6415E-15 | 0,999999925 | 1 |
| MYO1E | 1 | 1 | 0 | 1 | 1 |
| PCYT1A | 5 | 5 | -3,65887E-16 | 0,99999995 | 1 |
| KAZN | 2 | 2 | 7,72889E-16 | 0,999999983 | 1 |
| PBSN | 1 | 1 | -0,173794145 | 0,167862332 | 1 |
| RBSN | 1 | 1 | 0 | 1 | 1 |
| FBXL20 | 1 | 1 | 0,042990503 | 0,510016314 | 1 |
| CACNA2D1;CACNA: | 1 | 1 | 0 | 1 | 1 |
| CLCN2 | 2 | 2 | -6,83286E-18 | 0,999999996 | 1 |
| TRIP12 | 6 | 6 | -1,05815E-17 | 0,999999996 | 1 |
| PCDH7 | 14 | 14 | -0,066105343 | 0,123630117 | 1 |
| ACP1 | 1 | 1 | 0 | 1 | 1 |
| NRXN3 | 12 | 12 | 2,40095E-16 | 0,999999947 | 1 |
| TJP1 | 1 | 1 | 0 | 1 | 1 |
| GNAL | 4 | 4 | 0 | 1 | 1 |
| CCDC90B | 3 | 2 | 0,298304591 | 0,999938406 | 1 |
| FAM185A | 2 | 2 | -0,072927102 | 0,719581497 | 1 |
| EPHA3;EPHA6 | 1 | 1 | 0 | 1 | 1 |
| AKTIP | 2 | 2 | 0 | 1 | 1 |
| RAB3IP | 3 | 3 | 0 | 1 | 1 |
| SIDT1 | 4 | 4 | 0,067879316 | 0,187241032 | 1 |
| FAM98A | 2 | 2 | 1,13267E-17 | 0,999999995 | 1 |
| sp\|Q8BHB7\|CP046 | 1 | 1 | 0 | 1 | 1 |
| ZKSCAN8 | 1 | 1 | 0 | 1 | 1 |
| LIFR | 1 | 1 | -0,122656591 | 0,347287083 | 1 |
| BECN1 | 3 | 3 | 0 | 1 | 1 |
| TAOK3 | 3 | 3 | 1,51124E-17 | 0,99999999 | 1 |
| NUP85 | 1 | 1 | -0,066128638 | 0,677632333 | 1 |
| SNAP25 | 8 | 8 | 0,010975685 | 0,61322958 | 1 |

| | | | | | |
|---|---|---|---|---|---|
| RPL38 | 4 | 4 | -3,60725E-18 | 0,999999995 | 1 |
| EPHB1 | 8 | 7 | 0 | 1 | 1 |
| SELENOT | 3 | 3 | 7,93489E-15 | 0,999999757 | 1 |
| TMEM63B | 4 | 4 | -3,3841E-17 | 0,999999983 | 1 |
| INO80 | 1 | 1 | 0 | 1 | 1 |
| ANKRD13D | 4 | 4 | 0 | 1 | 1 |
| GAREM1 | 3 | 3 | 0 | 1 | 1 |
| RPL37A | 3 | 3 | 0 | 1 | 1 |
| NDUFAF3 | 5 | 5 | 0,011317415 | 0,635879523 | 1 |
| GABRB1;GABRB2 | 5 | 5 | -0,041260366 | 0,269946215 | 1 |
| GABRB3 | 5 | 5 | 0,04275996 | 0,340507759 | 1 |
| TGFB2 | 1 | 1 | 0 | 1 | 1 |
| CRYBG3 | 2 | 2 | 0,329462057 | 0,122615971 | 1 |
| PPIH | 1 | 1 | 0 | 1 | 1 |
| TSC22D1;TSC22D4 | 1 | 1 | 0 | 1 | 1 |
| ULK1 | 2 | 2 | -7,71724E-18 | 0,999999995 | 1 |
| COG6 | 2 | 2 | -4,32308E-17 | 0,999999988 | 1 |
| POMT2 | 1 | 1 | 0 | 1 | 1 |
| NLRP4A | 1 | 1 | -1,39926E-17 | 0,999999993 | 1 |
| FOCAD | 1 | 1 | 0 | 1 | 1 |
| WDR45B | 4 | 4 | -1,36623E-18 | 0,999999998 | 1 |
| PON1 | 2 | 2 | 0 | 1 | 1 |
| PCK2;PCK1 | 1 | 1 | 0 | 1 | 1 |
| SNRPN | 3 | 3 | 0 | 1 | 1 |
| MAP3K15;MAP3K5 | 2 | 2 | 0,102427314 | 0,132833368 | 1 |
| SERPINA1E | 6 | 6 | -0,575763701 | 0,225059738 | 1 |
| FGFR1OP2 | 3 | 3 | -0,02109627 | 0,576182923 | 1 |
| NRBP2 | 9 | 9 | 0,027006513 | 0,251658904 | 1 |
| PAPOLA;PAPOLB | 1 | 1 | 0 | 1 | 1 |
| NAT8L | 3 | 3 | 0 | 1 | 1 |
| SLC14A1 | 3 | 3 | -6,63255E-16 | 0,999999946 | 1 |
| PPP2R3D | 1 | 1 | 0,033868195 | 0,62241766 | 1 |
| NOS1AP | 2 | 2 | 0 | 1 | 1 |

| | | | | | |
|---|---|---|---|---|---|
| BAX | 8 | 8 | -0,02698279 | 0,373742502 | 1 |
| NTRK2;NTRK1;NTRI | 2 | 2 | -0,012016372 | 0,750563536 | 1 |
| INSR;INSRR | 1 | 1 | 0 | 1 | 1 |
| KCNJ16 | 1 | 1 | 0 | 1 | 1 |
| WNK1;WNK2;WNK | 2 | 2 | 1,24457E-17 | 0,999999998 | 1 |
| TUSC3 | 3 | 3 | 0 | 1 | 1 |
| ITGB2 | 15 | 15 | 0,033630874 | 0,335209809 | 1 |
| PRKAA1 | 4 | 4 | 0,080037905 | 0,133792634 | 1 |
| STN1 | 2 | 2 | 7,93806E-16 | 0,999999947 | 1 |
| SLC25A14 | 2 | 2 | 0 | 1 | 1 |
| OLFR552 | 1 | 1 | 0 | 1 | 1 |
| ABCC1 | 1 | 1 | 2,20847E-17 | 0,999999989 | 1 |
| RNF11 | 2 | 2 | -2,73537E-18 | 0,999999998 | 1 |
| UTRN | 1 | 1 | 0 | 1 | 1 |
| DNAJB14 | 2 | 2 | -2,21796E-18 | 0,999999997 | 1 |
| NGLY1 | 4 | 4 | 0 | 1 | 1 |
| FGFR1 | 2 | 1 | 0 | 1 | 1 |
| GRM4 | 2 | 2 | 0,03339744 | 0,695015166 | 1 |
| CTBP2 | 3 | 3 | -0,026954444 | 0,55890278 | 1 |
| TPRKB | 3 | 3 | 0 | 1 | 1 |
| SLC18A3 | 2 | 2 | -4,89726E-17 | 1 | 1 |
| KIF2B | 1 | 1 | 0 | 1 | 1 |
| CHD4 | 1 | 1 | 0 | 1 | 1 |
| PNKD | 1 | 1 | -4,09468E-20 | 1 | 1 |
| PPP2R3C | 1 | 1 | 0 | 1 | 1 |
| CCDC92 | 6 | 6 | -0,047388074 | 0,256665222 | 1 |
| KIF16B | 2 | 2 | 1,72239E-14 | 0,999999795 | 1 |
| SLC35G2 | 4 | 3 | 0 | 1 | 1 |
| MAGOH;MAGOHB | 1 | 1 | 0 | 1 | 1 |
| MYLK3;MYLK4;MYL | 1 | 1 | -1,0447E-22 | 1 | 1 |
| UQCC1 | 6 | 6 | -0,059487541 | 0,154344741 | 1 |
| HCK | 1 | 1 | 0 | 1 | 1 |
| APBA2 | 7 | 7 | 1,13056E-15 | 0,999999897 | 1 |

| | | | | | |
|---|---|---|---|---|---|
| XPNPEP1 | 1 | 1 | 0 | 1 | 1 |
| SEC14L1 | 2 | 2 | -3,18401E-15 | 0,99999989 | 1 |
| CBX1 | 6 | 6 | 0 | 1 | 1 |
| CAVIN1 | 1 | 1 | 0 | 1 | 1 |
| GNA14 | 1 | 1 | -0,142279149 | 0,153587527 | 1 |
| LOXHD1 | 1 | 1 | 0 | 1 | 1 |
| RASGRP1 | 4 | 4 | 7,83823E-17 | 0,999999983 | 1 |
| EHD4;EHD3;EHD1 | 4 | 4 | 0 | 1 | 1 |
| CNOT10 | 2 | 2 | -1,39729E-17 | 0,999999995 | 1 |
| SH3BP4 | 1 | 1 | 0 | 1 | 1 |
| RPA3 | 1 | 1 | -0,060897202 | 0,519075912 | 1 |
| HSPA1L;HSPA2;HSF | 2 | 2 | 1,28479E-14 | 0,999999735 | 1 |
| IPO8 | 2 | 2 | 0 | 1 | 1 |
| PIP4P2 | 2 | 2 | 0 | 1 | 1 |
| IGSF10 | 2 | 2 | -5,68115E-11 | 0,99999354 | 1 |
| LIN7A | 7 | 7 | -0,043881767 | 0,36031034 | 1 |
| RBM12B1 | 1 | 1 | 0 | 1 | 1 |
| STX8 | 3 | 3 | 0 | 1 | 1 |
| APBA3 | 1 | 1 | 0,046177444 | 0,444621594 | 1 |
| JADE3 | 1 | 1 | 0 | 1 | 1 |
| PLCL1;PLCL2 | 1 | 1 | 0 | 1 | 1 |
| RANBP6 | 2 | 2 | -2,60968E-17 | 0,999999989 | 1 |
| DYM | 2 | 2 | 1,2131E-10 | 0,999972043 | 1 |
| SPG21 | 2 | 2 | 0 | 1 | 1 |
| RAB33B;RAB33A | 1 | 1 | 0 | 1 | 1 |
| PSMG2 | 3 | 3 | -4,94036E-17 | 0,999999992 | 1 |
| MRPL21 | 4 | 4 | 2,70655E-18 | 0,999999997 | 1 |
| CPQ | 1 | 1 | 0 | 1 | 1 |
| TAF6 | 1 | 1 | 0 | 1 | 1 |
| CWC25 | 1 | 1 | -8,43758E-18 | 0,999999995 | 1 |
| NAB1 | 1 | 1 | -0,226623318 | 0,177774588 | 1 |
| MRPL14 | 4 | 3 | 0,085079789 | 0,379858884 | 1 |
| RAB8A;RAB10;RAB[illegible] | 1 | 1 | 0 | 1 | 1 |

| | | | | | |
|---|---|---|---|---|---|
| sp\|Q8K207\|CA021 | 1 | 1 | 0 | 1 | 1 |
| TOP1 | 1 | 1 | 0 | 1 | 1 |
| ERAP1 | 1 | 1 | 0 | 1 | 1 |
| AP4S1 | 1 | 1 | 0,094275895 | 0,398498836 | 1 |
| RICTOR | 1 | 1 | -0,011203988 | 0,777406845 | 1 |
| GDAP1 | 17 | 16 | 2,83075E-18 | 0,999999995 | 1 |
| CHMP6 | 2 | 2 | -0,061457078 | 0,314552725 | 1 |
| B3GALT9 | 2 | 2 | 6,83747E-18 | 0,999999996 | 1 |
| RPLP2 | 4 | 4 | -0,014657252 | 0,734048697 | 1 |
| HMBS | 4 | 4 | 0 | 1 | 1 |
| NFXL1 | 1 | 1 | 0 | 1 | 1 |
| NAA15 | 12 | 12 | -0,013895396 | 0,517563615 | 1 |
| METTL8 | 1 | 1 | 0 | 1 | 1 |
| NSL1 | 1 | 1 | 0 | 1 | 1 |
| GSDME | 8 | 8 | -0,064299124 | 0,138967783 | 1 |
| CHMP3 | 3 | 3 | 0 | 1 | 1 |
| MAPK8 | 5 | 5 | 6,47016E-15 | 0,999999772 | 1 |
| UBE3B | 3 | 3 | 2,30026E-12 | 0,999996174 | 1 |
| CFD | 1 | 1 | 0 | 1 | 1 |
| DNAJB5 | 2 | 2 | 1,70078E-18 | 0,999999999 | 1 |
| DNAJC24 | 1 | 1 | 0 | 1 | 1 |
| SCN4B | 1 | 1 | 0 | 1 | 1 |
| UPRT | 2 | 2 | -5,80509E-19 | 0,999999999 | 1 |
| ILF2 | 4 | 4 | 7,71396E-18 | 0,999999997 | 1 |
| ISOC2B | 2 | 2 | 0 | 1 | 1 |
| STRIP1;STRIP2 | 2 | 2 | -0,130780641 | 0,180587967 | 1 |
| POLR1C | 2 | 2 | 0 | 1 | 1 |
| NDRG2 | 1 | 1 | 0 | 1 | 1 |
| GM21698;GM2166 | 1 | 1 | 0,164314187 | 0,183114808 | 1 |
| TYW3 | 1 | 1 | 0 | 1 | 1 |
| GNA12;GNA13 | 1 | 1 | 0 | 1 | 1 |
| EFCAB5 | 1 | 1 | 0,170090929 | 0,170422675 | 1 |
| PHLPP2 | 1 | 1 | 0 | 1 | 1 |

| | | | | | |
|---|---|---|---|---|---|
| ERCC6L | 1 | 1 | 0,138744323 | 0,236120545 | 1 |
| SGO2 | 1 | 1 | 0 | 1 | 1 |
| KCNA10 | 2 | 2 | 2,24495E-17 | 0,999999994 | 1 |
| DCTN6 | 6 | 6 | 0 | 1 | 1 |
| ELP4 | 5 | 5 | -3,68804E-17 | 0,999999988 | 1 |
| RALGAPA1 | 14 | 14 | -1,08389E-16 | 0,99999998 | 1 |
| PKNOX2 | 1 | 1 | 0 | 1 | 1 |
| TMEM177 | 1 | 1 | 0 | 1 | 1 |
| GALNS | 1 | 1 | 0 | 1 | 1 |
| NBAS | 4 | 3 | 0,02368028 | 0,607387576 | 1 |
| TPM3 | 6 | 6 | -0,017443056 | 0,5639509 | 1 |
| LIRE1 | 1 | 1 | 0 | 1 | 1 |
| CEP97 | 2 | 2 | 0 | 1 | 1 |
| MRPS21 | 2 | 2 | 0,133499903 | 0,173912623 | 1 |
| IGSF21 | 8 | 8 | 3,66753E-17 | 0,999999987 | 1 |
| ASB6 | 1 | 1 | 0 | 1 | 1 |
| DENND4A | 2 | 2 | -4,32695E-14 | 0,999999743 | 1 |
| UBAC1 | 3 | 3 | -0,044754533 | 0,458565776 | 1 |
| SEC31B | 1 | 1 | 0 | 1 | 1 |
| DNAL1 | 4 | 4 | 2,14527E-16 | 0,999999965 | 1 |
| ABCA1 | 2 | 2 | 0 | 1 | 1 |
| LENG8 | 1 | 1 | 0 | 1 | 1 |
| PRPF31 | 2 | 2 | -0,060169853 | 0,414830944 | 1 |
| TUBB2A;TUBB6;TU | 3 | 3 | -5,1227E-18 | 0,999999998 | 1 |
| TUBB4B;TUBB5;TU | 2 | 2 | -1,00485E-18 | 0,999999999 | 1 |
| CARNMT1 | 1 | 1 | 0 | 1 | 1 |
| DRAP1 | 1 | 1 | 0,021905787 | 0,624053937 | 1 |
| PKP2 | 9 | 9 | 0,049678202 | 0,232522974 | 1 |
| VAV1 | 1 | 1 | 0 | 1 | 1 |
| GCA | 2 | 2 | -1,71317E-18 | 0,999999998 | 1 |
| DCAKD | 6 | 6 | 0,038012828 | 0,305547354 | 1 |
| COQ9 | 12 | 12 | 0 | 1 | 1 |
| ADH1 | 1 | 1 | 0 | 1 | 1 |

| | | | | | |
|---|---|---|---|---|---|
| SMAP | 4 | 4 | 0 | 1 | 1 |
| GANC | 7 | 7 | 4,27019E-15 | 0,999999871 | 1 |
| CEP152 | 1 | 1 | 0 | 1 | 1 |
| LRRTM2 | 4 | 3 | -7,42084E-18 | 0,999999995 | 1 |
| RCAN1 | 6 | 6 | -0,003703377 | 0,804569853 | 1 |
| UBE2D3 | 1 | 1 | 0 | 1 | 1 |
| PHYHIPL | 4 | 4 | -0,0405276 | 0,245464634 | 1 |
| MAP2K3;MAP2K6 | 1 | 1 | 0 | 1 | 1 |
| NR3C2 | 1 | 1 | 0,301208337 | 0,206854793 | 1 |
| TRMT112 | 2 | 2 | 0 | 1 | 1 |
| NAGA | 3 | 3 | 2,57178E-19 | 1 | 1 |
| EFNA3 | 1 | 1 | 0 | 1 | 1 |
| GUCY1A1 | 2 | 2 | -1,62723E-15 | 0,999999901 | 1 |
| TUBB2A | 2 | 2 | 9,562E-16 | 0,999999936 | 1 |
| TUBB4B | 1 | 1 | 0 | 1 | 1 |
| TUBB2B | 2 | 2 | 0 | 1 | 1 |
| DCAF5 | 3 | 3 | 2,67611E-16 | 0,999999978 | 1 |
| MARCHF5 | 6 | 5 | 0 | 1 | 1 |
| 1700014D04RIK | 1 | 1 | 0,039015814 | 0,530255136 | 1 |
| ACTL6B | 4 | 4 | 0 | 1 | 1 |
| TANC2 | 3 | 2 | 0 | 1 | 1 |
| SHOC2 | 4 | 4 | 5,12917E-16 | 0,999999951 | 1 |
| SPAG17 | 1 | 1 | 0 | 1 | 1 |
| SERINC1 | 3 | 3 | -0,075914419 | 0,28106059 | 1 |
| DNAH14 | 2 | 2 | 6,64625E-15 | 0,999999866 | 1 |
| NEU1 | 1 | 1 | 0,040763559 | 0,508481273 | 1 |
| ZFP407 | 1 | 1 | 0,104016887 | 0,396741197 | 1 |
| ELAC1 | 1 | 1 | 0 | 1 | 1 |
| TSHR | 2 | 2 | 0 | 1 | 1 |
| MAP2 | 1 | 1 | -0,053529712 | 0,558866666 | 1 |
| KMT2D | 1 | 1 | 0 | 1 | 1 |
| CX3CL1 | 2 | 2 | -0,036106102 | 0,586411812 | 1 |
| NTRK2 | 3 | 3 | 0 | 1 | 1 |

| | | | | | |
|---|---|---|---|---|---|
| MAP1LC3A;MAP1L | 2 | 2 | 1,81552E-18 | 0,999999999 | 1 |
| CBWD1 | 2 | 2 | 0 | 1 | 1 |
| P2YR13 | 1 | 1 | 0 | 1 | 1 |
| GSK3B | 5 | 5 | 0 | 1 | 1 |
| ADHFE1 | 4 | 3 | 2,48743E-18 | 0,999999999 | 1 |
| GABRA2 | 5 | 5 | -0,082123404 | 0,262513248 | 1 |
| R3HDM1 | 1 | 1 | -0,015063908 | 0,745913537 | 1 |
| DDX19B | 1 | 1 | 0 | 1 | 1 |
| HMG20B | 1 | 1 | 0 | 1 | 1 |
| SEC62 | 2 | 2 | 0 | 1 | 1 |
| SDAD1 | 1 | 1 | -0,139702514 | 0,243014085 | 1 |
| COQ8B | 2 | 1 | 0 | 1 | 1 |
| ZNFX1 | 1 | 1 | 0 | 1 | 1 |
| XRN1 | 1 | 1 | -0,036172943 | 0,614920654 | 1 |
| TPM3;TPM1;TPM2 | 4 | 4 | -0,04184086 | 0,345788815 | 1 |
| PPM1B;PPM1A | 5 | 5 | 0 | 1 | 1 |
| FBXW5 | 1 | 1 | 0 | 1 | 1 |
| SNX19 | 2 | 2 | 0 | 1 | 1 |
| WASHC3 | 1 | 1 | 0 | 1 | 1 |
| CSPG4B | 1 | 1 | 0 | 1 | 1 |
| VMN2R16 | 1 | 1 | 0 | 1 | 1 |
| SPRYD7 | 3 | 3 | 0 | 1 | 1 |
| ATP2B2 | 4 | 4 | 0 | 1 | 1 |
| ATP2B3 | 5 | 5 | 9,16438E-18 | 0,999999992 | 1 |
| FAM184A | 1 | 1 | 0 | 1 | 1 |
| SH3BGRL3 | 5 | 5 | 0,068974198 | 0,22829102 | 1 |
| ATCAY | 7 | 7 | -0,015546503 | 0,5502792 | 1 |
| KIF5A | 5 | 5 | -8,1427E-16 | 0,99999991 | 1 |
| KRT14;KRT16 | 8 | 3 | 1,88136E-18 | 0,999999999 | 1 |
| TUBB1;TUBB4B;TU | 1 | 1 | 0 | 1 | 1 |
| CLASP1;CLASP2 | 1 | 1 | -0,155859645 | 0,30821146 | 1 |
| CCDC77 | 1 | 1 | 0 | 1 | 1 |
| PLEKHH1;PLEKHH2 | 1 | 1 | 0 | 1 | 1 |

| | | | | | |
|---|---|---|---|---|---|
| DNM1 | 6 | 6 | -1,01618E-17 | 0,999999995 | 1 |
| TCEAL1 | 1 | 1 | -0,050243957 | 0,483891881 | 1 |
| TBCK | 4 | 4 | -3,9493E-13 | 0,999997843 | 1 |
| TNS2 | 2 | 1 | 0 | 1 | 1 |
| SHMT1;SHMT2 | 1 | 1 | 0 | 1 | 1 |
| OXNAD1 | 3 | 3 | 0 | 1 | 1 |
| CRHBP | 2 | 2 | -6,23086E-17 | 0,999999985 | 1 |
| ZFP804B | 1 | 1 | -8,96497E-15 | 0,999999802 | 1 |
| TIAM2 | 2 | 2 | 5,47715E-17 | 0,99999999 | 1 |
| ERLIN2;ERLIN1 | 3 | 3 | 0,071169904 | 0,184949852 | 1 |
| PSME1 | 6 | 6 | 0,034969026 | 0,46067206 | 1 |
| TUBB4B;TUBB5;TU | 6 | 6 | 9,81534E-16 | 0,999999935 | 1 |
| LAMA5 | 1 | 1 | -1,66296E-14 | 0,999999752 | 1 |
| ACOT1 | 1 | 1 | 0 | 1 | 1 |
| SCRG1 | 1 | 1 | 0 | 1 | 1 |
| MAG | 1 | 1 | 9,24101E-15 | 0,999999807 | 1 |
| CDK1 | 1 | 1 | 0,104187603 | 0,511378182 | 1 |
| FAM210B | 2 | 2 | -0,10373367 | 0,34851457 | 1 |
| HIVEP2 | 4 | 4 | 6,70774E-15 | 0,999999905 | 1 |
| PSD3 | 2 | 2 | 5,86879E-14 | 0,999999826 | 1 |
| MESP2 | 1 | 1 | 0,236878664 | 0,300287438 | 1 |
| TGOLN1 | 2 | 2 | 0 | 1 | 1 |
| DCN | 4 | 2 | 1,07692E-17 | 0,999999998 | 1 |
| SZRD1 | 1 | 1 | 0 | 1 | 1 |
| TAF5 | 1 | 1 | 0 | 1 | 1 |
| ANKH | 3 | 3 | 0 | 1 | 1 |
| SLC2A4 | 1 | 1 | 0,035807336 | 0,6991018 | 1 |
| KRTCAP2 | 1 | 1 | 0 | 1 | 1 |
| USP33 | 3 | 2 | 4,07194E-17 | 0,999999994 | 1 |
| RPH3A;DOC2A | 1 | 1 | 0 | 1 | 1 |
| MAPK9 | 6 | 6 | 0 | 1 | 1 |
| SSBP1 | 10 | 9 | 2,21314E-15 | 0,99999992 | 1 |
| ATP9A | 11 | 11 | 0,012586234 | 0,537197551 | 1 |

| | | | | | |
|---|---|---|---|---|---|
| MRPL20 | 1 | 1 | -1,75555E-18 | 0,999999998 | 1 |
| OSBPL9 | 3 | 3 | 9,39897E-19 | 0,999999999 | 1 |
| TXNDC17 | 3 | 3 | 0,107717033 | 0,14798191 | 1 |
| AKT1;AKT2;AKT3 | 3 | 3 | -5,14248E-16 | 0,999999943 | 1 |
| DSTYK | 1 | 1 | -0,202325725 | 0,113803976 | 1 |
| VPS72 | 1 | 1 | 0 | 1 | 1 |
| CRYGE;CRYGF | 1 | 1 | 0 | 1 | 1 |
| SURF1 | 3 | 2 | 1,09056E-17 | 0,999999996 | 1 |
| 4930512M02RIK | 1 | 1 | 0 | 1 | 1 |
| POLR2I | 1 | 1 | 0 | 1 | 1 |
| HSPA1L;HSPA2;HSF | 3 | 3 | 0 | 1 | 1 |
| CDH8 | 2 | 2 | -0,16694203 | 0,11631689 | 1 |
| ABCA9 | 4 | 4 | -6,9697E-16 | 0,99999996 | 1 |
| UFM1 | 2 | 2 | 1,74341E-16 | 0,999999963 | 1 |
| SLC27A2 | 1 | 1 | -0,035814638 | 0,637237567 | 1 |
| AFAP1 | 1 | 1 | 0,210830421 | 0,294716897 | 1 |
| ADGB | 2 | 2 | -4,2888E-17 | 0,999999991 | 1 |
| FAM107A | 1 | 1 | 0 | 1 | 1 |
| CDC27 | 2 | 2 | -0,166351192 | 0,192634997 | 1 |
| PCDHA3 | 1 | 1 | 0 | 1 | 1 |
| SCN7A | 1 | 1 | 0 | 1 | 1 |
| PPP6R2 | 1 | 1 | 0 | 1 | 1 |
| POLR2B | 4 | 4 | 0 | 1 | 1 |
| DNAJB3 | 1 | 1 | 1,11935E-21 | 1 | 1 |
| DNAJB6 | 1 | 1 | 0,019462433 | 0,642319101 | 1 |
| COG4 | 2 | 2 | -7,54167E-18 | 0,999999994 | 1 |
| GARIN2 | 1 | 1 | 0 | 1 | 1 |
| PABPC1L | 1 | 1 | 0 | 1 | 1 |
| NAA10;NAA12 | 3 | 3 | -0,086239517 | 0,217333557 | 1 |
| MIX23 | 3 | 3 | 0 | 1 | 1 |
| EPB41L3 | 2 | 2 | 6,20703E-19 | 1 | 1 |
| TMPPE | 2 | 1 | 0 | 1 | 1 |
| PRSS33 | 1 | 1 | 0 | 1 | 1 |

| | | | | | |
|---|---|---|---|---|---|
| ARHGEF11 | 13 | 13 | -2,50502E-17 | 0,999999992 | 1 |
| LMF1 | 1 | 1 | 0,056257313 | 0,445338517 | 1 |
| IER5 | 1 | 1 | 0 | 1 | 1 |
| GAPDH | 2 | 2 | 0 | 1 | 1 |
| STAMBP | 3 | 3 | -1,24177E-17 | 0,999999995 | 1 |
| ANO6 | 1 | 1 | 0 | 1 | 1 |
| GRIA2;GRIA4 | 1 | 1 | 8,72036E-22 | 1 | 1 |
| GABRA2;GABRA1;G | 3 | 3 | -7,16157E-18 | 0,999999996 | 1 |
| GOLGA7 | 2 | 2 | 0 | 1 | 1 |
| TMEM192 | 1 | 1 | 0 | 1 | 1 |
| UBE2D1 | 2 | 2 | 0,026839623 | 0,5073794 | 1 |
| UBE2D2B | 2 | 2 | -0,014218496 | 0,66018161 | 1 |
| RNF14 | 1 | 1 | 0 | 1 | 1 |
| MTATP8 | 3 | 3 | 0 | 1 | 1 |
| FLRT3 | 4 | 4 | -1,88381E-18 | 0,999999998 | 1 |
| ABLIM3 | 1 | 1 | 0 | 1 | 1 |
| WDR45 | 3 | 3 | -1,15476E-16 | 0,999999977 | 1 |
| DNAJB7 | 2 | 2 | 0 | 1 | 1 |
| PHF14 | 1 | 1 | -0,084117287 | 0,32671167 | 1 |
| MYC | 1 | 1 | 0,10172443 | 0,181458259 | 1 |
| SARG | 1 | 1 | 0 | 1 | 1 |
| SNX10 | 2 | 2 | -9,97847E-18 | 0,999999994 | 1 |
| KHDRBS1 | 1 | 1 | 0 | 1 | 1 |
| ADRM1 | 8 | 7 | -2,88453E-17 | 0,999999988 | 1 |
| ODAD2 | 1 | 1 | -9,04388E-15 | 0,999999824 | 1 |
| ODAD4 | 1 | 1 | 0 | 1 | 1 |
| GNAI1;GNAI2;GNAI | 4 | 4 | 0 | 1 | 1 |
| HNRNPDL | 2 | 2 | -0,049593917 | 0,370282608 | 1 |
| GUCY2C | 1 | 1 | 0 | 1 | 1 |
| CEACAM13 | 1 | 1 | 0 | 1 | 1 |
| CYP3A16 | 1 | 1 | 0 | 1 | 1 |
| PRKCH | 1 | 1 | -0,079374765 | 0,478689515 | 1 |
| PLPP1 | 1 | 1 | 0,015585915 | 0,741459426 | 1 |

| | | | | | |
|---|---|---|---|---|---|
| SMAP1 | 4 | 4 | 5,39613E-17 | 0,999999984 | 1 |
| SLC35F3 | 1 | 1 | 0 | 1 | 1 |
| FAM210A | 5 | 4 | 0 | 1 | 1 |
| AGAP3;AGAP1 | 2 | 2 | 3,61901E-17 | 0,999999988 | 1 |
| KCNQ2 | 1 | 1 | -0,177586795 | 0,345232625 | 1 |
| ZC3HAV1 | 1 | 1 | 0 | 1 | 1 |
| TMCO1 | 3 | 3 | 0 | 1 | 1 |
| SARS1 | 1 | 1 | 1,04131E-20 | 1 | 1 |
| PTMA | 4 | 4 | -7,72044E-17 | 0,999999988 | 1 |
| SPAG9;MAPK8IP3 | 6 | 6 | -2,82891E-17 | 0,999999985 | 1 |
| LIG3 | 1 | 1 | 0 | 1 | 1 |
| DOCK4;DOCK3 | 3 | 3 | 0 | 1 | 1 |
| ATP6V0A4;ATP6V0 | 1 | 1 | 0 | 1 | 1 |
| ANO1 | 1 | 1 | 0 | 1 | 1 |
| FKBP2 | 4 | 4 | 0 | 1 | 1 |
| GSTT1 | 2 | 2 | 0 | 1 | 1 |
| BSDC1 | 4 | 4 | 0 | 1 | 1 |
| RP1 | 2 | 2 | 1,58883E-18 | 0,999999998 | 1 |
| CCDC106 | 1 | 1 | 0 | 1 | 1 |
| A630010A05RIK | 1 | 1 | 0,013173528 | 0,784399695 | 1 |
| LIMCH1 | 1 | 1 | 0 | 1 | 1 |
| NUP205 | 2 | 2 | -3,73242E-16 | 0,999999958 | 1 |
| GRM1 | 8 | 8 | -0,001011493 | 0,884146093 | 1 |
| NEFH;KRT6A;KRT2; | 2 | 2 | -6,19087E-20 | 1 | 1 |
| NEFL;INA | 2 | 1 | 0 | 1 | 1 |
| GPAA1 | 3 | 3 | 1,00849E-14 | 0,99999972 | 1 |
| GJC3 | 2 | 2 | -9,60294E-18 | 0,999999998 | 1 |
| SNX17 | 5 | 5 | -0,028206139 | 0,342262943 | 1 |
| MAPK10;MAPK8;M | 1 | 1 | 0,13735788 | 0,129961582 | 1 |
| GNG7 | 3 | 3 | 0 | 1 | 1 |
| MRPL12 | 5 | 5 | 0 | 1 | 1 |
| CEP164 | 1 | 1 | 0 | 1 | 1 |
| SYT1;SYT5 | 5 | 5 | 3,74176E-17 | 0,999999983 | 1 |

| | | | | | |
|---|---|---|---|---|---|
| CAMTA2 | 1 | 1 | 0 | 1 | 1 |
| GTF3C4 | 1 | 1 | 0 | 1 | 1 |
| CHD2;CHD1 | 1 | 1 | 0 | 1 | 1 |
| BCAT2 | 3 | 3 | 0,084064094 | 0,331808902 | 1 |
| RPL28 | 3 | 3 | 2,37875E-19 | 0,999999999 | 1 |
| BSG | 9 | 9 | 0 | 1 | 1 |
| UGT2B1 | 1 | 1 | 0 | 1 | 1 |
| MAP7D3 | 1 | 1 | 0 | 1 | 1 |
| GPT2;GPT | 1 | 1 | 0 | 1 | 1 |
| FHL2 | 1 | 1 | 0 | 1 | 1 |
| WNT7A | 1 | 1 | 0 | 1 | 1 |
| MACF1 | 2 | 2 | 3,58834E-12 | 0,999999949 | 1 |
| RGS2 | 1 | 1 | 0 | 1 | 1 |
| CAST | 1 | 1 | -2,04941E-17 | 0,999999991 | 1 |
| GTF3C1 | 2 | 2 | -0,011328191 | 0,70839701 | 1 |
| TACC2 | 1 | 1 | 0 | 1 | 1 |
| LASP1;NEBL | 1 | 1 | 0 | 1 | 1 |
| OSBPL8;OSBPL5 | 1 | 1 | 0 | 1 | 1 |
| HOOK1 | 2 | 2 | 0 | 1 | 1 |
| PPP1R13B | 9 | 9 | -1,21048E-18 | 1 | 1 |
| RFK | 3 | 3 | 0,007662774 | 0,733111221 | 1 |
| SYCP1 | 1 | 1 | 0,049972176 | 0,568883359 | 1 |
| RPS24 | 3 | 3 | 8,5196E-17 | 0,999999985 | 1 |
| GNL3 | 1 | 1 | 0 | 1 | 1 |
| RPL30 | 8 | 8 | -1,37127E-16 | 0,999999961 | 1 |
| SYN2;SYN3 | 2 | 2 | -3,01016E-17 | 0,999999994 | 1 |
| PLAG1 | 1 | 1 | 0 | 1 | 1 |
| ZGRF1 | 1 | 1 | 0 | 1 | 1 |
| MPC1 | 4 | 4 | 0 | 1 | 1 |
| ANKS1B | 2 | 2 | 0,002220072 | 0,86001484 | 1 |
| MMS22L | 1 | 1 | -0,154045825 | 0,220867797 | 1 |
| CDC14A | 1 | 1 | 0 | 1 | 1 |
| GLRX | 3 | 3 | 0 | 1 | 1 |

| | | | | | |
|---|---|---|---|---|---|
| DNM1 | 3 | 3 | 0 | 1 | 1 |
| TICAM1 | 1 | 1 | 0 | 1 | 1 |
| ARXES1;ARXES2 | 3 | 3 | 0 | 1 | 1 |
| DAO | 1 | 1 | 0 | 1 | 1 |
| CALR3 | 1 | 1 | -0,297255217 | 0,125515123 | 1 |
| COL28A1 | 1 | 1 | 0 | 1 | 1 |
| KIF21A | 1 | 1 | 0 | 1 | 1 |
| ARRB2 | 3 | 3 | -0,050541698 | 0,409973887 | 1 |
| PDE12 | 7 | 7 | -7,10539E-17 | 0,999999975 | 1 |
| ISCU | 5 | 5 | 0 | 1 | 1 |
| RALA;RALB | 6 | 6 | 0,049484808 | 0,242313607 | 1 |
| EP400 | 1 | 1 | 0 | 1 | 1 |
| ABL1;ABL2 | 3 | 3 | 0 | 1 | 1 |
| AAK1;BMP2K | 2 | 2 | 2,65666E-17 | 0,999999986 | 1 |
| PURB;PURG | 2 | 2 | -8,04362E-17 | 0,999999983 | 1 |
| PPP1R2 | 5 | 4 | 0 | 1 | 1 |
| KRT24;KRT36;KRT1 | 1 | 1 | 0 | 1 | 1 |
| PSMB6 | 4 | 4 | 7,44996E-20 | 1 | 1 |
| MINPP1 | 1 | 1 | 0 | 1 | 1 |
| ZFYVE19 | 3 | 3 | -0,086561723 | 0,132195057 | 1 |
| ELOA | 1 | 1 | 0 | 1 | 1 |
| PDIA6 | 2 | 2 | 1,76008E-07 | 0,998870808 | 1 |
| RAB3D | 3 | 3 | -5,5767E-16 | 0,999999948 | 1 |
| CDK14;CDK1;CDK5; | 1 | 1 | 0 | 1 | 1 |
| STK24;STK25 | 3 | 3 | -3,71355E-06 | 0,994699677 | 1 |
| KCNH1 | 1 | 1 | 0,094189097 | 0,433866412 | 1 |
| WWOX | 4 | 4 | -1,69472E-16 | 0,99999998 | 1 |
| OXA1L | 2 | 2 | 1,99757E-19 | 1 | 1 |
| SDS | 1 | 1 | 0,330302124 | 0,142468948 | 1 |
| NUBP2 | 3 | 3 | 0 | 1 | 1 |
| RPL13A | 3 | 3 | 0 | 1 | 1 |
| ALG11 | 1 | 1 | 0 | 1 | 1 |
| CARD14 | 1 | 1 | 0 | 1 | 1 |

| | | | | | |
|---|---|---|---|---|---|
| SZT2 | 3 | 3 | 3,75689E-18 | 0,999999996 | 1 |
| KRT6A;KRT76;KRT5 | 2 | 2 | 5,92568E-15 | 0,999999999 | 1 |
| VPS37C | 2 | 2 | 0 | 1 | 1 |
| PCNX3 | 2 | 2 | -0,041372939 | 0,522228767 | 1 |
| CROCC | 2 | 1 | -0,098292372 | 0,315077736 | 1 |
| VPS8 | 3 | 3 | -3,12911E-18 | 0,999999998 | 1 |
| DIAPH2 | 3 | 3 | -0,068375701 | 0,179541835 | 1 |
| GATC | 1 | 1 | 0 | 1 | 1 |
| TMEM63B | 1 | 1 | 0 | 1 | 1 |
| RDH12 | 1 | 1 | 0 | 1 | 1 |
| FBXL18 | 1 | 1 | 0 | 1 | 1 |
| RAC3 | 2 | 2 | -4,40247E-19 | 0,999999999 | 1 |
| USP25 | 3 | 3 | -0,088118348 | 0,189154655 | 1 |
| LANCL3 | 1 | 1 | -2,35803E-21 | 1 | 1 |
| sp\|Q922C1\|CS044_ | 1 | 1 | 0 | 1 | 1 |
| DNAH2 | 1 | 1 | 0 | 1 | 1 |
| TRPM3 | 1 | 1 | 0 | 1 | 1 |
| HIKESHI | 3 | 3 | 0 | 1 | 1 |
| TBL1XR1 | 1 | 1 | 0,087576949 | 0,332275032 | 1 |
| TCAIM | 1 | 1 | 0 | 1 | 1 |
| PTK2 | 7 | 6 | -1,39595E-16 | 0,999999961 | 1 |
| ENO1;ENO3 | 2 | 2 | 8,0602E-13 | 0,999998 | 1 |
| TBPL2 | 1 | 1 | 0 | 1 | 1 |
| CCDC32 | 1 | 1 | 0 | 1 | 1 |
| GFRA2 | 2 | 2 | 0 | 1 | 1 |
| UBP1 | 4 | 4 | 0 | 1 | 1 |
| MINDY1 | 3 | 3 | -0,063394743 | 0,159231696 | 1 |
| PSD2 | 2 | 2 | 0 | 1 | 1 |
| HAS1 | 1 | 1 | 0 | 1 | 1 |
| ZSWIM8 | 1 | 1 | 0 | 1 | 1 |
| COMMD5 | 1 | 1 | 0,065981297 | 0,476581994 | 1 |
| NCOR2 | 1 | 1 | 0 | 1 | 1 |
| MAPK15 | 1 | 1 | 0 | 1 | 1 |

| | | | | | |
|---|---|---|---|---|---|
| DENR | 5 | 5 | 0 | 1 | 1 |
| NLGN1 | 2 | 2 | 0,05288404 | 0,441013675 | 1 |
| DCUN1D2 | 2 | 2 | -0,096052647 | 0,116657135 | 1 |
| ULK3 | 1 | 1 | 0 | 1 | 1 |
| NR2C2 | 1 | 1 | 0 | 1 | 1 |
| RRAS;RRAS2 | 3 | 3 | 0 | 1 | 1 |
| OGDH | 2 | 2 | 0 | 1 | 1 |
| SMC1A | 1 | 1 | 0 | 1 | 1 |
| MACO1 | 2 | 2 | 1,09027E-17 | 0,999999999 | 1 |
| DNM1;DNM2;DNM | 7 | 7 | 6,73004E-15 | 0,999999679 | 1 |
| LRTM2 | 2 | 2 | -1,22115E-16 | 0,999999994 | 1 |
| FXN | 2 | 2 | 0 | 1 | 1 |
| LRFN2;LRFN4 | 1 | 1 | 0 | 1 | 1 |
| TSC22D1 | 1 | 1 | 0 | 1 | 1 |
| KBTBD4 | 1 | 1 | 0 | 1 | 1 |
| MAP3K10 | 1 | 1 | 0,001766538 | 0,905178124 | 1 |
| TIMM13 | 3 | 3 | -8,4002E-17 | 0,999999989 | 1 |
| TXNDC9 | 4 | 4 | -1,96315E-15 | 0,999999877 | 1 |
| TBCEL | 7 | 7 | -0,047362194 | 0,218200818 | 1 |
| IRGM1 | 5 | 5 | 9,86601E-19 | 0,999999999 | 1 |
| SYNGAP1 | 1 | 1 | 0 | 1 | 1 |
| DCLK2;DCLK1 | 1 | 1 | 0,060343768 | 0,359368231 | 1 |
| SPO11 | 1 | 1 | 0 | 1 | 1 |
| NCEH1 | 15 | 13 | -2,08737E-17 | 0,999999988 | 1 |
| STMN3;STMN1;STM | 1 | 1 | -1,96728E-18 | 0,999999997 | 1 |
| HOOK1;HOOK3 | 1 | 1 | 0 | 1 | 1 |
| ODF2 | 1 | 1 | 0 | 1 | 1 |
| ZNF608 | 1 | 1 | 0 | 1 | 1 |
| GRM2;GRM4 | 2 | 2 | 0 | 1 | 1 |
| GNG12 | 5 | 5 | 4,13763E-16 | 0,999999938 | 1 |
| OBSCN | 1 | 1 | 0 | 1 | 1 |
| TMED5 | 1 | 1 | 0 | 1 | 1 |
| LRG1 | 2 | 2 | 3,32848E-18 | 0,999999999 | 1 |

| | | | | | |
|---|---|---|---|---|---|
| ADD1 | 1 | 1 | 0 | 1 | 1 |
| ADD1 | 1 | 1 | -9,09264E-19 | 0,999999999 | 1 |
| MYH11 | 1 | 1 | 0 | 1 | 1 |
| ARHGAP22 | 1 | 1 | 0 | 1 | 1 |
| ATPAF2 | 2 | 2 | 3,08519E-18 | 0,999999996 | 1 |
| SLC12A5;SLC12A6 | 3 | 3 | -2,81971E-16 | 0,999999942 | 1 |
| BBS9 | 1 | 1 | -1,1649E-20 | 1 | 1 |
| SPECC1 | 1 | 1 | 0,0052768 | 0,844007511 | 1 |
| ARHGAP36 | 1 | 1 | 0 | 1 | 1 |
| GPC6 | 1 | 1 | 0 | 1 | 1 |
| GSE1 | 1 | 1 | 0 | 1 | 1 |
| LAMC3 | 1 | 1 | 0 | 1 | 1 |
| PEX1 | 2 | 2 | 3,39523E-16 | 0,999999975 | 1 |
| TAX1BP1 | 4 | 4 | 0 | 1 | 1 |
| APOL7B;APOL7E | 1 | 1 | 0 | 1 | 1 |
| GOLGA4 | 1 | 1 | 0 | 1 | 1 |
| BBS7 | 1 | 1 | 0 | 1 | 1 |
| MCUR1 | 4 | 4 | -5,63421E-16 | 0,999999944 | 1 |
| KIF1B | 3 | 3 | 0 | 1 | 1 |
| DR1 | 1 | 1 | -0,016446535 | 0,675735537 | 1 |
| 4931408C20RIK | 2 | 2 | -2,35531E-16 | 0,999999969 | 1 |
| YTHDF1;YTHDF3 | 1 | 1 | 0 | 1 | 1 |
| ARC | 1 | 1 | 0 | 1 | 1 |
| LMBRD1 | 1 | 1 | 0 | 1 | 1 |
| DMAC2 | 1 | 1 | 0 | 1 | 1 |
| TRIM58 | 1 | 1 | 0,14732092 | 0,149523209 | 1 |
| KCNT1 | 1 | 1 | 0 | 1 | 1 |
| DMD | 6 | 6 | 5,79767E-14 | 0,999999746 | 1 |
| PFKFB2;PFKFB4 | 1 | 1 | 0 | 1 | 1 |
| C1QB | 4 | 4 | 0 | 1 | 1 |
| KRT14;KRT42;KRT1 | 3 | 3 | 1,7408E-15 | 0,999999947 | 1 |
| TOR1AIP1 | 1 | 1 | 0 | 1 | 1 |
| SCYL2 | 13 | 12 | 0 | 1 | 1 |

| | | | | | |
|---|---|---|---|---|---|
| HDAC5 | 4 | 4 | 1,46513E-15 | 0,999999931 | 1 |
| PPP1R12B | 2 | 2 | 0 | 1 | 1 |
| TRAPPC6B | 5 | 4 | 0,000792747 | 0,906000025 | 1 |
| FASTKD1 | 1 | 1 | 0 | 1 | 1 |
| ANKS1B | 1 | 1 | 0 | 1 | 1 |
| PRC1 | 1 | 1 | 0,112374402 | 0,471475675 | 1 |
| TRIM6 | 1 | 1 | 0 | 1 | 1 |
| PCDH1 | 6 | 6 | 0 | 1 | 1 |
| HMGCS2;HMGCS1 | 1 | 1 | 0 | 1 | 1 |
| PDILT | 1 | 1 | 0 | 1 | 1 |
| TLR13 | 1 | 1 | 0 | 1 | 1 |
| SEZ6L2 | 2 | 2 | 0 | 1 | 1 |
| PLXNA2;PLXNA4 | 5 | 5 | 0,003289021 | 0,771004888 | 1 |
| ARMC8 | 3 | 3 | -7,49055E-17 | 0,999999981 | 1 |
| COG5 | 2 | 2 | 2,20334E-20 | 1 | 1 |
| HSPB1 | 1 | 1 | 0 | 1 | 1 |
| GNAI1;GNAI2;GNA( | 5 | 5 | 0,060769532 | 0,185754407 | 1 |
| PAPOLA | 1 | 1 | -0,091694295 | 0,31382069 | 1 |
| TBC1D14 | 1 | 1 | 0 | 1 | 1 |
| PPP2R2A;PPP2R2C | 2 | 2 | -4,75452E-19 | 0,999999999 | 1 |
| NCF2 | 1 | 1 | 0 | 1 | 1 |
| MARK4;MARK1 | 1 | 1 | 4,53369E-18 | 0,999999995 | 1 |
| TMEM9B | 1 | 1 | 0 | 1 | 1 |
| CHKA | 1 | 1 | 0 | 1 | 1 |
| LARP6 | 3 | 3 | 0 | 1 | 1 |
| IL1RAPL1 | 2 | 1 | 0,094126857 | 0,399904352 | 1 |
| PTPRC | 1 | 1 | 0 | 1 | 1 |
| SNX24 | 1 | 1 | 0 | 1 | 1 |
| WDR6 | 2 | 2 | 0 | 1 | 1 |
| RHBDD2 | 1 | 1 | 0 | 1 | 1 |
| CYB5D2 | 2 | 2 | 1,2788E-18 | 0,999999999 | 1 |
| TP53RKB | 1 | 1 | -0,141799468 | 0,230559103 | 1 |
| NUDT21 | 2 | 1 | 0 | 1 | 1 |

| | | | | | |
|---|---|---|---|---|---|
| RIOX1 | 1 | 1 | 0 | 1 | 1 |
| COQ10B | 1 | 1 | 0 | 1 | 1 |
| USP12 | 1 | 1 | 0 | 1 | 1 |
| PDK1;PDK3 | 1 | 1 | 0 | 1 | 1 |
| MAPK10 | 4 | 4 | 0 | 1 | 1 |
| AKR1C14 | 1 | 1 | 0 | 1 | 1 |
| AMIGO2 | 1 | 1 | 0,094553161 | 0,409716853 | 1 |
| RIMS3 | 2 | 2 | 1,0751E-18 | 0,999999998 | 1 |
| COX7A1 | 1 | 1 | 0 | 1 | 1 |
| CEPT1 | 1 | 1 | 0,00886386 | 0,759523585 | 1 |
| GSTM4 | 4 | 4 | -6,85862E-16 | 0,999999972 | 1 |
| RRAS | 3 | 3 | 4,01188E-17 | 0,999999989 | 1 |
| RBP3 | 1 | 1 | 0 | 1 | 1 |
| ITIH4 | 5 | 5 | 7,19392E-17 | 0,99999999 | 1 |
| DAZAP1 | 2 | 2 | 0 | 1 | 1 |
| TMEM126B | 1 | 1 | 0 | 1 | 1 |
| VLDLR | 2 | 1 | -0,084305506 | 0,456293476 | 1 |
| FAM241B | 2 | 2 | -0,012679468 | 0,687172428 | 1 |
| MIGA1 | 2 | 2 | 0 | 1 | 1 |
| ATP7A | 1 | 1 | 0 | 1 | 1 |
| ATP4A;ATP1A4 | 1 | 1 | 0 | 1 | 1 |
| TMEM256 | 2 | 2 | -1,09408E-17 | 0,999999997 | 1 |
| ACOX3 | 5 | 5 | 0 | 1 | 1 |
| RRAGA | 1 | 1 | 0 | 1 | 1 |
| COA7 | 3 | 3 | 0,047327534 | 0,412409457 | 1 |
| GRSF1 | 2 | 2 | -0,012235448 | 0,749787358 | 1 |
| NEK2;IRAK1 | 1 | 1 | 0 | 1 | 1 |
| PTK2;PTK2B | 1 | 1 | 0 | 1 | 1 |
| NRN1 | 4 | 4 | 8,16226E-18 | 0,999999996 | 1 |
| CDK16;CDK17 | 2 | 2 | -2,17293E-20 | 1 | 1 |
| PRSS1 | 1 | 1 | -0,005567182 | 0,815856327 | 1 |
| ITGB5 | 1 | 1 | 0 | 1 | 1 |
| ITGB6 | 1 | 1 | 0 | 1 | 1 |

| | | | | | |
|---|---|---|---|---|---|
| DHX9 | 1 | 1 | -6,52118E-19 | 0,999999999 | 1 |
| PPM1L | 3 | 3 | 0 | 1 | 1 |
| KCNC2 | 3 | 2 | 0 | 1 | 1 |
| CD34 | 2 | 2 | 0 | 1 | 1 |
| CD2AP | 1 | 1 | 0 | 1 | 1 |
| MRPL17 | 1 | 1 | 0 | 1 | 1 |
| CDC5L | 1 | 1 | 0 | 1 | 1 |
| SLC25A27 | 1 | 1 | 0 | 1 | 1 |
| MOCS3 | 2 | 2 | -1,31073E-15 | 0,999999896 | 1 |
| MOCS2 | 1 | 1 | 0 | 1 | 1 |
| ICOSLG | 1 | 1 | 0 | 1 | 1 |
| FAM171A1 | 7 | 7 | -6,10821E-16 | 0,999999958 | 1 |
| TCAF1 | 2 | 2 | -0,09590079 | 0,260947014 | 1 |
| PCDH1 | 5 | 5 | 9,79409E-17 | 0,999999984 | 1 |
| RER1 | 4 | 4 | -4,31002E-18 | 0,999999996 | 1 |
| LYRM7 | 2 | 1 | 0 | 1 | 1 |
| WWC1 | 2 | 2 | 0,116297334 | 0,222275378 | 1 |
| YRDC | 2 | 2 | -1,64273E-15 | 0,999999902 | 1 |
| SLC30A5 | 2 | 1 | 0 | 1 | 1 |
| SOGA1 | 2 | 2 | -2,73903E-17 | 0,999999993 | 1 |
| LSAMP | 1 | 1 | -3,33145E-20 | 1 | 1 |
| PIBF1 | 1 | 1 | 0 | 1 | 1 |
| KIF5C;KIF5A | 6 | 5 | 0 | 1 | 1 |
| CCAR1 | 1 | 1 | 0 | 1 | 1 |
| NUCB2 | 2 | 2 | 0 | 1 | 1 |
| TFCP2L1 | 1 | 1 | 0 | 1 | 1 |
| TUBB6;TUBB3 | 4 | 4 | -2,37651E-17 | 0,999999993 | 1 |
| TUBB4B;TUBB5;TU | 7 | 7 | 0 | 1 | 1 |
| CLSTN3 | 2 | 2 | 5,35661E-19 | 0,999999999 | 1 |
| KIF3A;KIF4 | 1 | 1 | -0,013144871 | 0,731114036 | 1 |
| ZFP101 | 1 | 1 | 0 | 1 | 1 |
| MTATP6 | 1 | 1 | 0 | 1 | 1 |
| PTS | 5 | 5 | 0,042064312 | 0,187331924 | 1 |

| | | | | | |
|---|---|---|---|---|---|
| ATP6V0A2;ATP6V0. | 1 | 1 | 0 | 1 | 1 |
| SCFD1 | 7 | 7 | -9,15692E-16 | 0,999999907 | 1 |
| TKTL1 | 2 | 2 | 0,040673704 | 0,454265929 | 1 |
| ANK2 | 3 | 3 | -8,82278E-15 | 0,999999876 | 1 |
| ATP1A2;ATP1A1;AT | 6 | 6 | -2,64453E-20 | 1 | 1 |
| IQGAP2;IQGAP1 | 2 | 2 | -6,3316E-16 | 0,999999957 | 1 |
| TUBA3B;TUBA1A | 1 | 1 | 0 | 1 | 1 |
| GNB4;GNB1 | 2 | 2 | -4,25685E-20 | 1 | 1 |
| UQCR11 | 1 | 1 | 0 | 1 | 1 |
| OLFR141 | 1 | 1 | 0 | 1 | 1 |
| STING1 | 1 | 1 | 0 | 1 | 1 |
| FABP3 | 10 | 10 | -7,23923E-18 | 0,999999997 | 1 |
| GID8 | 3 | 3 | 8,02461E-16 | 0,999999942 | 1 |
| PIK3CA | 1 | 1 | 0 | 1 | 1 |
| NTF3 | 1 | 1 | 0 | 1 | 1 |
| D130040H23RIK | 1 | 1 | 0 | 1 | 1 |
| CTSS | 4 | 4 | 2,17665E-17 | 0,999999992 | 1 |
| NEDD4 | 14 | 14 | -1,00449E-15 | 0,999999905 | 1 |
| SRSF5;SRSF6;SRSF4 | 1 | 1 | 0 | 1 | 1 |
| MT-ND2 | 2 | 2 | 0 | 1 | 1 |
| TMEM70 | 2 | 2 | 0 | 1 | 1 |
| ST6GALNAC1 | 1 | 1 | 0 | 1 | 1 |
| CNTRL | 1 | 1 | -0,160267357 | 0,37551531 | 1 |
| PPP1R12B;PPP1R1. | 2 | 2 | -1,18938E-17 | 0,999999994 | 1 |
| STK16 | 1 | 1 | 0 | 1 | 1 |
| NAA16 | 2 | 2 | 0,093935175 | 0,116866158 | 1 |
| LRRTM4;LRRTM3 | 1 | 1 | 0 | 1 | 1 |
| DNM2 | 1 | 1 | 0 | 1 | 1 |
| THSD7A | 5 | 5 | -4,536E-19 | 0,999999999 | 1 |
| CAPG | 4 | 4 | 5,45081E-16 | 0,999999958 | 1 |
| RPL19 | 3 | 3 | 0,095203228 | 0,130650972 | 1 |
| PRKCD | 2 | 2 | -0,066700603 | 0,327148631 | 1 |
| PODXL2 | 1 | 1 | 0 | 1 | 1 |

| | | | | | |
|---|---|---|---|---|---|
| sp\|Q8WUR0\|CS01: | 1 | 1 | 0 | 1 | 1 |
| SPRY3 | 1 | 1 | 0,098268534 | 0,388096779 | 1 |
| SYNGAP1 | 1 | 1 | 0 | 1 | 1 |
| SELENOF | 1 | 1 | 0 | 1 | 1 |
| TBC1D22B | 4 | 4 | 0 | 1 | 1 |
| TBC1D22A | 7 | 7 | 0 | 1 | 1 |
| GABRA2;GABRA1 | 2 | 2 | 3,96248E-16 | 0,99999999 | 1 |
| KIF23 | 1 | 1 | 3,11025E-21 | 1 | 1 |
| TTC27 | 1 | 1 | 0 | 1 | 1 |
| ARL10 | 1 | 1 | 0 | 1 | 1 |
| STXBP1 | 2 | 2 | 0 | 1 | 1 |
| PIK3R1 | 3 | 3 | -0,055890311 | 0,326573895 | 1 |
| LCT | 2 | 2 | -0,046122159 | 0,609407466 | 1 |
| MRPL28 | 4 | 4 | -1,118E-17 | 0,999999991 | 1 |
| DNAJC16 | 3 | 3 | -0,001059074 | 0,927742777 | 1 |
| ANKRD46 | 1 | 1 | 0,097269027 | 0,261937597 | 1 |
| GLT1D1 | 1 | 1 | -0,00654317 | 0,85041918 | 1 |
| KPNA2 | 1 | 1 | 0 | 1 | 1 |
| PET117 | 1 | 1 | 0 | 1 | 1 |
| DPP6 | 1 | 1 | 0 | 1 | 1 |
| GIPC2 | 2 | 1 | 0 | 1 | 1 |
| SCO2 | 3 | 3 | 0 | 1 | 1 |
| SLC4A9 | 1 | 1 | 0 | 1 | 1 |
| TELO2 | 2 | 2 | -0,062364075 | 0,477554735 | 1 |
| SPRY2 | 2 | 2 | 0 | 1 | 1 |
| ARHGEF25 | 2 | 2 | -3,8373E-17 | 0,999999987 | 1 |
| HDAC4 | 1 | 1 | 0 | 1 | 1 |
| NUDT10;NUDT11 | 7 | 7 | 4,18309E-18 | 0,999999999 | 1 |
| PIP5KL1 | 1 | 1 | 0 | 1 | 1 |
| RGS7 | 1 | 1 | 0 | 1 | 1 |
| NRCAM | 1 | 1 | 0 | 1 | 1 |
| PLXNA2 | 7 | 7 | -6,40026E-17 | 0,999999994 | 1 |
| TMEM143 | 3 | 2 | 1,04282E-16 | 0,999999977 | 1 |

| | | | | | |
|---|---|---|---|---|---|
| PON3 | 1 | 1 | 0 | 1 | 1 |
| ELK3 | 1 | 1 | 0 | 1 | 1 |
| BCAP29 | 3 | 3 | 0,038721724 | 0,451500768 | 1 |
| CAMK2A;CAMK2D; | 1 | 1 | 3,57907E-17 | 0,999999988 | 1 |
| ITPR3 | 3 | 3 | -4,29902E-19 | 1 | 1 |
| MRM1 | 1 | 1 | 0 | 1 | 1 |
| MTMR3 | 4 | 4 | 1,40463E-17 | 0,999999992 | 1 |
| CASP1 | 1 | 1 | -0,162962364 | 0,165183496 | 1 |
| DZANK1 | 2 | 2 | 8,03453E-18 | 0,999999998 | 1 |
| RAB8A;RAB10;RAB | 1 | 1 | -0,064843975 | 0,395219149 | 1 |
| RILP | 1 | 1 | -0,069444211 | 0,517249838 | 1 |
| CHADL | 1 | 1 | 0 | 1 | 1 |
| YES1 | 1 | 1 | 0 | 1 | 1 |
| RAPGEFL1 | 5 | 5 | -9,93003E-17 | 0,999999971 | 1 |
| H2BC15;H2BC14;H | 5 | 2 | -2,60945E-16 | 0,999999988 | 1 |
| NTMT1 | 2 | 2 | -2,54876E-16 | 0,999999963 | 1 |
| MPP4 | 1 | 1 | 0 | 1 | 1 |
| DEF8 | 1 | 1 | 0 | 1 | 1 |
| DPH1 | 1 | 1 | 0 | 1 | 1 |
| STARD3NL | 2 | 2 | 0 | 1 | 1 |
| ENPP4 | 1 | 1 | 0 | 1 | 1 |
| AKR1B7;AKR1B8 | 1 | 1 | 0 | 1 | 1 |
| SLC13A1 | 1 | 1 | 0 | 1 | 1 |
| MROH1 | 1 | 1 | -0,187230552 | 0,300746814 | 1 |
| TP53BP2 | 1 | 1 | 0 | 1 | 1 |
| GFOD2 | 1 | 1 | 0 | 1 | 1 |
| PIP5K1C;PIP5K1B;P | 2 | 2 | 4,33696E-18 | 0,999999996 | 1 |
| JAK3 | 1 | 1 | 0,09414206 | 0,383664325 | 1 |
| HGH1 | 1 | 1 | 0 | 1 | 1 |
| GDAP2 | 3 | 3 | 6,13635E-18 | 0,999999996 | 1 |
| AMFR | 3 | 3 | -0,096828742 | 0,218709512 | 1 |
| CALB1;CALB2 | 1 | 1 | 0 | 1 | 1 |
| DNAH9 | 1 | 1 | -0,087594107 | 0,347216896 | 1 |

| | | | | | |
|---|---|---|---|---|---|
| SORCS3 | 4 | 4 | 0 | 1 | 1 |
| SURF6 | 1 | 1 | 0 | 1 | 1 |
| SMURF1 | 1 | 1 | 0 | 1 | 1 |
| SCG2 | 4 | 4 | -9,79541E-21 | 1 | 1 |
| HSD11B2 | 1 | 1 | 0 | 1 | 1 |
| SAMD10 | 1 | 1 | 0 | 1 | 1 |
| IGHV4-1 | 1 | 1 | 0 | 1 | 1 |
| LCMT2;SEPTIN5 | 1 | 1 | 0 | 1 | 1 |
| EXOC1 | 3 | 3 | 3,55816E-16 | 0,999999962 | 1 |
| NUP93 | 2 | 2 | 0 | 1 | 1 |
| METAP1 | 2 | 2 | 0 | 1 | 1 |
| PRELID3A | 2 | 1 | -9,34004E-17 | 0,999999984 | 1 |
| 2410002F23RIK | 3 | 3 | -3,42919E-18 | 0,999999996 | 1 |
| UBR5 | 1 | 1 | 0 | 1 | 1 |
| OLFR1180 | 1 | 1 | 0 | 1 | 1 |
| RNF112 | 1 | 1 | 0 | 1 | 1 |
| ARHGAP33;ARHGA | 1 | 1 | -0,067690702 | 0,432078843 | 1 |
| IYD | 1 | 1 | 0 | 1 | 1 |
| HAT1 | 1 | 1 | -0,137822756 | 0,264542314 | 1 |
| SLC33A1 | 2 | 2 | 0 | 1 | 1 |
| NDUFAB1 | 5 | 4 | 0,090393923 | 0,225211827 | 1 |
| VTA1 | 6 | 6 | 0 | 1 | 1 |
| NAA20 | 2 | 2 | -0,165799958 | 0,16114404 | 1 |
| KCNB1 | 2 | 2 | -5,26039E-18 | 0,999999995 | 1 |
| LRFN2 | 4 | 4 | -2,7938E-18 | 1 | 1 |
| SLC25A32 | 3 | 2 | 0 | 1 | 1 |
| SNN | 1 | 1 | 0 | 1 | 1 |
| EFCAB8 | 1 | 1 | -1,16783E-18 | 0,999999998 | 1 |
| TSPAN15 | 1 | 1 | 0 | 1 | 1 |
| QSOX1 | 1 | 1 | -0,114390499 | 0,287320475 | 1 |
| DOK6 | 1 | 1 | 0 | 1 | 1 |
| NRXN2;NRXN3 | 1 | 1 | 0 | 1 | 1 |
| ALG10B | 1 | 1 | 0 | 1 | 1 |

| | | | | | |
|---|---|---|---|---|---|
| LRRTM3 | 1 | 1 | 0 | 1 | 1 |
| BUB3 | 3 | 3 | -0,069679725 | 0,261788669 | 1 |
| REPS1;REPS2 | 1 | 1 | 0 | 1 | 1 |
| PARVA | 2 | 2 | -5,25781E-18 | 0,999999999 | 1 |
| KCTD2 | 1 | 1 | 0 | 1 | 1 |
| DUSP23 | 1 | 1 | 0 | 1 | 1 |
| YEATS2 | 1 | 1 | 0 | 1 | 1 |
| CDK16 | 4 | 4 | -1,01648E-16 | 0,999999981 | 1 |
| PRKDC | 1 | 1 | 0 | 1 | 1 |
| LONRF2 | 1 | 1 | 0 | 1 | 1 |
| MRPL9 | 3 | 2 | 0 | 1 | 1 |
| VMN2R11 | 1 | 1 | 0 | 1 | 1 |
| AKT1;AKT3 | 1 | 1 | 0 | 1 | 1 |
| MTM1 | 2 | 2 | 2,84613E-17 | 1 | 1 |
| GPATCH2L | 1 | 1 | 0 | 1 | 1 |
| UGCG | 1 | 1 | 0 | 1 | 1 |
| RHOT2 | 1 | 1 | 0 | 1 | 1 |
| EPG5 | 4 | 4 | -0,040385114 | 0,359875573 | 1 |
| TMEM11 | 3 | 3 | 3,16266E-18 | 0,999999996 | 1 |
| PCMTD1 | 2 | 2 | -4,41474E-14 | 0,999999643 | 1 |
| ANKMY1 | 1 | 1 | 0 | 1 | 1 |
| EPHB6 | 2 | 2 | 7,78596E-19 | 0,999999999 | 1 |
| ZBTB18 | 2 | 2 | -0,101474577 | 0,211707826 | 1 |
| LHFPL4 | 1 | 1 | 0 | 1 | 1 |
| PANX2 | 2 | 2 | -0,019312448 | 0,660092034 | 1 |
| ITSN2 | 1 | 1 | 0 | 1 | 1 |
| NCF1 | 1 | 1 | 0 | 1 | 1 |
| SLC35F6 | 1 | 1 | 0 | 1 | 1 |
| CHAC2 | 3 | 3 | -0,151407142 | 0,191773075 | 1 |
| SLC25A29 | 3 | 3 | 0,113508411 | 0,200703299 | 1 |
| KTN1 | 1 | 1 | -0,011529826 | 0,724200047 | 1 |
| S100A6 | 1 | 1 | 0 | 1 | 1 |
| ACO2 | 1 | 1 | 0,095668989 | 0,325787136 | 1 |

| | | | | | |
|---|---|---|---|---|---|
| P4HA1 | 2 | 2 | 4,59632E-18 | 0,999999996 | 1 |
| RBL2 | 1 | 1 | 0 | 1 | 1 |
| DDX10 | 1 | 1 | 0,005296669 | 0,833497615 | 1 |
| PARP14 | 1 | 1 | 0 | 1 | 1 |
| HDAC9;HDAC5 | 1 | 1 | -3,17625E-19 | 0,999999999 | 1 |
| ARHGAP10 | 1 | 1 | 0 | 1 | 1 |
| OS9 | 1 | 1 | 0,156303362 | 0,339424243 | 1 |
| CNTN3;CNTN4 | 1 | 1 | 0 | 1 | 1 |
| HLCS | 2 | 2 | -0,060560103 | 0,498354043 | 1 |
| RDH13 | 3 | 2 | 3,65993E-15 | 0,999999927 | 1 |
| TTC3 | 2 | 2 | -5,29032E-19 | 0,999999999 | 1 |
| SPRY4 | 2 | 2 | 0 | 1 | 1 |
| ERBIN | 3 | 3 | 0 | 1 | 1 |
| GPATCH11 | 2 | 2 | 5,60867E-19 | 0,999999999 | 1 |
| RAB8A;RAB4A;RAB | 1 | 1 | 0 | 1 | 1 |
| KIFC1 | 1 | 1 | -0,010499046 | 0,846188752 | 1 |
| RAPH1 | 7 | 7 | 1,24745E-18 | 0,999999999 | 1 |
| GABRD | 1 | 1 | -0,030981271 | 0,631555147 | 1 |
| MMP25 | 1 | 1 | 0 | 1 | 1 |
| CCDC190 | 1 | 1 | 0,078252135 | 0,266296297 | 1 |
| ANGPTL6 | 1 | 1 | 0 | 1 | 1 |
| DEF6 | 1 | 1 | 0 | 1 | 1 |
| SALL2 | 1 | 1 | 0 | 1 | 1 |
| LSP1 | 1 | 1 | 0 | 1 | 1 |
| STAU1 | 1 | 1 | 0 | 1 | 1 |
| SPTAN1 | 2 | 2 | 7,02352E-16 | 0,999999947 | 1 |
| TMEM175 | 2 | 2 | 0 | 1 | 1 |
| DNAL4 | 1 | 1 | 0 | 1 | 1 |
| FBXL2 | 1 | 1 | -0,1721806 | 0,306640917 | 1 |
| MKRN2 | 1 | 1 | 0 | 1 | 1 |
| TMEM9;TMEM9B | 1 | 1 | 0 | 1 | 1 |
| XKR4 | 2 | 2 | 1,04281E-05 | 0,991559402 | 1 |
| MAP3K6 | 1 | 1 | 0 | 1 | 1 |

| | | | | | |
|---|---|---|---|---|---|
| SAMD14 | 3 | 3 | 0 | 1 | 1 |
| TBX15 | 1 | 1 | 0 | 1 | 1 |
| RGS7BP | 4 | 4 | 2,92821E-17 | 0,999999988 | 1 |
| UNC80 | 2 | 2 | -7,24084E-17 | 0,999999991 | 1 |
| ITGB1 | 4 | 4 | 2,69574E-19 | 0,999999999 | 1 |
| GABRA1;GABRA5 | 2 | 2 | 0 | 1 | 1 |
| MPC2 | 7 | 7 | -3,94853E-20 | 1 | 1 |
| SELENOS | 1 | 1 | 1,79381E-19 | 1 | 1 |
| ARHGEF4 | 2 | 2 | 0 | 1 | 1 |
| CLDN12 | 1 | 1 | 0 | 1 | 1 |
| SLC23A2 | 3 | 2 | 0 | 1 | 1 |
| IGHV1-62-1 | 1 | 1 | 0,090917812 | 0,201027887 | 1 |
| PRELP | 2 | 2 | 3,99222E-14 | 0,999999864 | 1 |
| USP38 | 2 | 1 | -0,285789272 | 0,281122078 | 1 |
| JAK1 | 3 | 3 | -2,51099E-17 | 0,999999994 | 1 |
| TOMM34 | 1 | 1 | 0 | 1 | 1 |
| MRPS36 | 1 | 1 | 0 | 1 | 1 |
| NSG1 | 1 | 1 | 0 | 1 | 1 |
| GRIA1;GRIA2 | 2 | 2 | 0 | 1 | 1 |
| GM11639 | 2 | 2 | 2,63944E-16 | 0,999999968 | 1 |
| COMMD1 | 2 | 2 | 0 | 1 | 1 |
| B4GALNT3 | 1 | 1 | 0 | 1 | 1 |
| IFT122 | 1 | 1 | 0 | 1 | 1 |
| PMVK | 2 | 2 | 5,20922E-17 | 0,999999992 | 1 |
| PDSS2 | 1 | 1 | 0 | 1 | 1 |
| 4930438A08RIK | 1 | 1 | 0,370841947 | 0,16088248 | 1 |
| SLC22A6 | 1 | 1 | 0 | 1 | 1 |
| MLIP | 2 | 1 | -0,176733129 | 0,253032534 | 1 |
| KIF13B | 1 | 1 | 0 | 1 | 1 |
| UNC5D | 1 | 1 | 0 | 1 | 1 |
| PLS1;LCP1 | 1 | 1 | -8,04912E-20 | 1 | 1 |
| HK2 | 2 | 2 | 0 | 1 | 1 |
| MYO16 | 1 | 1 | -0,415821384 | 0,156227392 | 1 |

| | | | | | |
|---|---|---|---|---|---|
| SLC29A2 | 1 | 1 | -0,104063816 | 0,208206748 | 1 |
| PHACTR1;PHACTR4 | 1 | 1 | 0,0170445 | 0,66392457 | 1 |
| WASF3 | 8 | 7 | -7,48675E-17 | 0,999999987 | 1 |
| DVL1 | 2 | 2 | 0 | 1 | 1 |
| CNST | 2 | 2 | 0 | 1 | 1 |
| ZFP189 | 1 | 1 | 0 | 1 | 1 |
| CPEB2 | 1 | 1 | 0 | 1 | 1 |
| LRRFIP1 | 1 | 1 | 0 | 1 | 1 |
| RFC2 | 1 | 1 | 0 | 1 | 1 |
| 1810024B03RIK | 1 | 1 | 0 | 1 | 1 |
| LSM2 | 3 | 3 | 0 | 1 | 1 |
| AGO4;AGO3;AGO1 | 1 | 1 | 0 | 1 | 1 |
| CTTNBP2NL | 1 | 1 | 0 | 1 | 1 |
| MAP3K20 | 1 | 1 | 0 | 1 | 1 |
| RAB3GAP1 | 1 | 1 | 0 | 1 | 1 |
| ATP6V0A2;ATP6V0 | 1 | 1 | 0 | 1 | 1 |
| GAB1 | 2 | 2 | -3,47638E-17 | 0,999999994 | 1 |
| CSMD1 | 2 | 2 | -0,170379631 | 0,137234003 | 1 |
| OLFM2 | 2 | 2 | -2,87534E-16 | 0,999999981 | 1 |
| HM13 | 4 | 4 | -3,098E-16 | 0,999999971 | 1 |
| UBE2A | 1 | 1 | 0 | 1 | 1 |
| MORC3 | 1 | 1 | 0 | 1 | 1 |
| ATRN | 1 | 1 | 0 | 1 | 1 |
| MYO5A | 4 | 4 | -0,035511845 | 0,357949023 | 1 |
| ARFGAP3 | 3 | 3 | 5,19025E-14 | 0,999999524 | 1 |
| DCP1A | 1 | 1 | 0 | 1 | 1 |
| VPS26C | 3 | 3 | -4,40578E-17 | 0,999999993 | 1 |
| LGI4 | 2 | 1 | 0 | 1 | 1 |
| MRPS18B | 2 | 2 | -1,17187E-17 | 0,999999993 | 1 |
| KLC1 | 1 | 1 | 0 | 1 | 1 |
| EMC3 | 5 | 5 | 4,02426E-16 | 0,999999982 | 1 |
| CAMK2G;CAMK2B | 2 | 2 | 0 | 1 | 1 |
| CNOT9 | 3 | 3 | -5,0157E-18 | 0,999999998 | 1 |

| | | | | | |
|---|---|---|---|---|---|
| ANKRD13A | 2 | 2 | -1,28962E-17 | 0,999999997 | 1 |
| KIF21A | 1 | 1 | 0 | 1 | 1 |
| MRPS16 | 2 | 1 | 0 | 1 | 1 |
| LRIT1 | 1 | 1 | 0 | 1 | 1 |
| PRKG1;PRKG2 | 1 | 1 | 0 | 1 | 1 |
| GNPDA2 | 6 | 6 | -6,62276E-17 | 0,999999988 | 1 |
| CD2BP2 | 1 | 1 | 0 | 1 | 1 |
| TMOD3 | 2 | 1 | 0 | 1 | 1 |
| INPP5B | 3 | 3 | 2,99391E-17 | 0,999999988 | 1 |
| PLAAT3 | 2 | 1 | 0,00013452 | 0,974313903 | 1 |
| UBE2B | 1 | 1 | 0,112774218 | 0,204889268 | 1 |
| LDHC;LDHA | 1 | 1 | 0 | 1 | 1 |
| TPM2 | 1 | 1 | 0 | 1 | 1 |
| MOXD1 | 1 | 1 | 0 | 1 | 1 |
| SERPINB9C;SERPINI | 1 | 1 | 0 | 1 | 1 |
| SERPINB8 | 1 | 1 | 0 | 1 | 1 |
| SLC5A3 | 2 | 2 | 2,55354E-14 | 0,999999796 | 1 |
| HDHD5 | 5 | 3 | -0,034379803 | 0,561470736 | 1 |
| CARS2 | 2 | 2 | 5,21819E-18 | 0,999999998 | 1 |
| NCOA1 | 2 | 2 | 0 | 1 | 1 |
| RPS27;RPS27L | 2 | 2 | 0,048104555 | 0,295886952 | 1 |
| CACNA1B;CACNA1/ | 1 | 1 | 0 | 1 | 1 |
| SNX14 | 1 | 1 | 0 | 1 | 1 |
| CD47 | 3 | 3 | 0 | 1 | 1 |
| HS2ST1 | 2 | 1 | 0 | 1 | 1 |
| NUCB1 | 3 | 3 | 7,52869E-19 | 0,999999999 | 1 |
| PCBP3;PCBP2 | 1 | 1 | -0,040964629 | 0,448297995 | 1 |
| FEZF1 | 1 | 1 | 0 | 1 | 1 |
| FCGR1 | 3 | 3 | 0 | 1 | 1 |
| MFHAS1 | 1 | 1 | -4,47547E-21 | 1 | 1 |
| RPL27A | 4 | 4 | -0,054096117 | 0,181573962 | 1 |
| KLHL3 | 2 | 2 | -4,4145E-19 | 0,999999999 | 1 |
| GDF10 | 1 | 1 | 0 | 1 | 1 |

| | | | | | |
|---|---|---|---|---|---|
| NAA30 | 3 | 3 | -2,06651E-16 | 0,999999975 | 1 |
| TAF1C | 1 | 1 | 0 | 1 | 1 |
| MRPL44 | 1 | 1 | 8,14455E-20 | 1 | 1 |
| IVNS1ABP | 1 | 1 | -0,050369318 | 0,430373611 | 1 |
| FAM91A1 | 4 | 4 | 0 | 1 | 1 |
| CAMSAP3;CAMSAP | 1 | 1 | 0 | 1 | 1 |
| PCNT | 1 | 1 | -0,023235265 | 0,802539512 | 1 |
| CFAP20 | 2 | 2 | 5,19013E-17 | 0,999999995 | 1 |
| CDK3 | 1 | 1 | 0,247013229 | 0,254840138 | 1 |
| SESN1 | 1 | 1 | 0 | 1 | 1 |
| MRPL55 | 3 | 2 | 0 | 1 | 1 |
| WDR73 | 1 | 1 | 0 | 1 | 1 |
| CAMK2G | 2 | 2 | 0 | 1 | 1 |
| ZFP386 | 1 | 1 | 0,107112623 | 0,252790579 | 1 |
| GSTM1;GSTM2 | 1 | 1 | 0 | 1 | 1 |
| UFSP1 | 1 | 1 | 0 | 1 | 1 |
| WDR12 | 1 | 1 | 0,008086272 | 0,780851058 | 1 |
| USP35 | 3 | 3 | 4,51377E-19 | 0,999999999 | 1 |
| DESI1 | 1 | 1 | 0 | 1 | 1 |
| ABLIM2 | 1 | 1 | 0,050145565 | 0,398168657 | 1 |
| NPR2 | 1 | 1 | -0,029565222 | 0,704742375 | 1 |
| NEFL;NEFH;INA | 1 | 1 | 0 | 1 | 1 |
| ATP6V1C2 | 1 | 1 | 0 | 1 | 1 |
| FAM3C | 2 | 2 | -1,22827E-16 | 0,999999976 | 1 |
| JPT1 | 3 | 3 | -2,00955E-17 | 0,999999995 | 1 |
| CDC42EP1 | 1 | 1 | 0 | 1 | 1 |
| RTN1 | 3 | 3 | 7,71023E-19 | 0,999999999 | 1 |
| TENM4;TENM1 | 2 | 2 | 0,038331285 | 0,607050427 | 1 |
| NPTXR;NPCD | 2 | 2 | 0,04246052 | 0,418925571 | 1 |
| DPYSL2;CRMP1 | 3 | 3 | -9,92798E-18 | 0,999999993 | 1 |
| TXNL4A | 1 | 1 | 0 | 1 | 1 |
| FAT3 | 2 | 2 | 0 | 1 | 1 |
| LMNTD1 | 1 | 1 | 0 | 1 | 1 |

| | | | | | |
|---|---|---|---|---|---|
| MYZAP | 1 | 1 | -0,191555144 | 0,354298698 | 1 |
| GOLGA1 | 1 | 1 | -0,002327393 | 0,879148022 | 1 |
| GRIK3;GRIK2 | 1 | 1 | 0 | 1 | 1 |
| ATP5MJ | 2 | 2 | 1,90788E-17 | 0,999999997 | 1 |
| TRMT6 | 3 | 3 | -0,094645731 | 0,217398902 | 1 |
| CDIPT | 6 | 6 | 0 | 1 | 1 |
| CCDC159 | 1 | 1 | 0 | 1 | 1 |
| NFYC | 2 | 2 | -0,106145184 | 0,177931495 | 1 |
| HK2;HK1;HK3 | 3 | 3 | 0,079261477 | 0,405103864 | 1 |
| MAP3K7 | 3 | 3 | 1,54331E-16 | 0,999999968 | 1 |
| VASP | 1 | 1 | 0 | 1 | 1 |
| ARF3;ARF1;ARF2 | 7 | 7 | -0,076245453 | 0,227288419 | 1 |
| TK2 | 3 | 3 | 0 | 1 | 1 |
| STRN;STRN3 | 1 | 1 | 0 | 1 | 1 |
| DTL | 1 | 1 | 0 | 1 | 1 |
| STXBP6 | 1 | 1 | -0,08430676 | 0,419308067 | 1 |
| HIPK3 | 1 | 1 | 0 | 1 | 1 |
| ROBO1;ROBO2 | 2 | 2 | 1,53276E-17 | 0,999999998 | 1 |
| GRCC10 | 3 | 3 | 0 | 1 | 1 |
| MAPK10 | 4 | 3 | -3,51822E-20 | 1 | 1 |
| DPH5 | 2 | 2 | 0 | 1 | 1 |
| ASB3 | 1 | 1 | 0 | 1 | 1 |
| SCG3 | 3 | 3 | -1,78079E-17 | 0,999999991 | 1 |
| RBM8A | 2 | 2 | 0 | 1 | 1 |
| INSR | 3 | 3 | -1,35512E-16 | 0,999999988 | 1 |
| TUBB4B;TUBB2A;TI | 4 | 4 | -1,28345E-15 | 0,999999932 | 1 |
| ACO2 | 2 | 2 | -6,37121E-18 | 0,999999998 | 1 |
| CYRIA;CYRIB | 5 | 5 | -1,95052E-16 | 0,999999968 | 1 |
| RRAGD | 1 | 1 | 0 | 1 | 1 |
| MYLK4 | 1 | 1 | 0 | 1 | 1 |
| CSNK1D | 1 | 1 | 0 | 1 | 1 |
| KRT14;KRT42;KRT1 | 3 | 2 | 7,46542E-16 | 0,999999974 | 1 |
| H1F10 | 2 | 2 | 0 | 1 | 1 |

| | | | | | |
|---|---|---|---|---|---|
| STX18 | 2 | 2 | -1,01307E-17 | 0,999999995 | 1 |
| VNN1 | 1 | 1 | 0 | 1 | 1 |
| GCSH | 3 | 3 | 2,77907E-15 | 0,999999914 | 1 |
| CCDC113 | 1 | 1 | 0 | 1 | 1 |
| PIR | 4 | 4 | -1,36611E-14 | 0,999999798 | 1 |
| GNAZ;GNA11;GNA( | 2 | 2 | 0 | 1 | 1 |
| BBOF1 | 1 | 1 | 0,086825086 | 0,457273646 | 1 |
| MLST8 | 5 | 5 | 0 | 1 | 1 |
| MLH1 | 1 | 1 | -0,01444543 | 0,749509744 | 1 |
| ROBO1 | 1 | 1 | 0,101517018 | 0,408443285 | 1 |
| CETN4 | 1 | 1 | -2,03565E-17 | 0,999999992 | 1 |
| PATL1 | 1 | 1 | -0,060207031 | 0,315002149 | 1 |
| PLA2R1 | 1 | 1 | 0,149076258 | 0,208175331 | 1 |
| CFL1;CFL2;DSTN | 1 | 1 | 0 | 1 | 1 |
| YWHAQ;YWHAB | 1 | 1 | 1,90806E-20 | 1 | 1 |
| YWHAG;YWHAZ;YV | 2 | 2 | -0,12908333 | 0,212615011 | 1 |
| OLFR1188 | 1 | 1 | -0,31658579 | 0,132761931 | 1 |
| MUC16 | 2 | 1 | 0 | 1 | 1 |
| APBA1;APBA2 | 2 | 1 | -0,378415944 | 0,141257933 | 1 |
| BPIFB9A | 1 | 1 | 0 | 1 | 1 |
| TERB1 | 1 | 1 | 0 | 1 | 1 |
| UMODL1 | 1 | 1 | 0,005541845 | 0,812176695 | 1 |
| ZP3R | 1 | 1 | 0 | 1 | 1 |
| ATP12A | 1 | 1 | 0 | 1 | 1 |
| SNX11 | 2 | 2 | 0 | 1 | 1 |
| VPS9D1 | 1 | 1 | 0 | 1 | 1 |
| TARS3;TARS1 | 2 | 2 | -1,94075E-14 | 0,999999747 | 1 |
| AGO1 | 4 | 4 | -1,11935E-15 | 0,99999992 | 1 |
| SMN1 | 1 | 1 | 0 | 1 | 1 |
| GRIA3;GRIA4;GRIA: | 1 | 1 | 0 | 1 | 1 |
| PKIG | 1 | 1 | 0 | 1 | 1 |
| RIPK2 | 1 | 1 | -0,343746044 | 0,201389409 | 1 |
| ITFG1 | 7 | 7 | 6,28786E-17 | 0,999999982 | 1 |

| | | | | | |
|---|---|---|---|---|---|
| ARHGEF10 | 1 | 1 | -3,32381E-18 | 0,999999997 | 1 |
| CAPZB | 3 | 3 | -4,17294E-18 | 0,999999995 | 1 |
| STAC3 | 2 | 1 | 0 | 1 | 1 |
| ATF2 | 1 | 1 | -0,074715443 | 0,470483356 | 1 |
| SFR1 | 3 | 3 | -4,77696E-12 | 0,99999436 | 1 |
| KCNIP2 | 1 | 1 | 0 | 1 | 1 |
| ZMPSTE24 | 4 | 4 | 0 | 1 | 1 |
| PLEKHA5 | 4 | 4 | 0 | 1 | 1 |
| LPCAT2 | 1 | 1 | 0 | 1 | 1 |
| LPAR1 | 1 | 1 | -0,044140521 | 0,669320403 | 1 |
| ANKS1B | 2 | 2 | 0 | 1 | 1 |
| UBR7 | 2 | 2 | -0,059482203 | 0,345415296 | 1 |
| CPLX3 | 1 | 1 | 0 | 1 | 1 |
| KCNA10;KCNA1;KC | 1 | 1 | 0,121691959 | 0,23026527 | 1 |
| U2AF2 | 1 | 1 | 0 | 1 | 1 |
| DERL1 | 1 | 1 | 0 | 1 | 1 |
| NEUROD2 | 1 | 1 | 0 | 1 | 1 |
| NEO1 | 2 | 2 | 2,24077E-16 | 0,999999974 | 1 |
| TRIM2;TRIM3 | 1 | 1 | -9,85376E-15 | 0,99999979 | 1 |
| SGPP1 | 1 | 1 | 0 | 1 | 1 |
| KRT42;KRT17 | 3 | 2 | 0 | 1 | 1 |
| MMGT1 | 2 | 2 | -2,17398E-19 | 0,999999999 | 1 |
| GPM6B | 1 | 1 | 0 | 1 | 1 |
| DNLZ | 1 | 1 | 0 | 1 | 1 |
| PGM5 | 2 | 2 | 0 | 1 | 1 |
| TBC1D23 | 3 | 3 | 0 | 1 | 1 |
| MAPK10;MAPK9 | 2 | 2 | 7,0886E-16 | 0,999999954 | 1 |
| IDH1;IDH2 | 2 | 2 | 1,2482E-17 | 0,999999992 | 1 |
| TRIM9 | 3 | 3 | 6,72545E-17 | 0,999999978 | 1 |
| IL18 | 2 | 2 | 0 | 1 | 1 |
| IGSF3 | 1 | 1 | 0 | 1 | 1 |
| MTMR10 | 1 | 1 | 0 | 1 | 1 |
| RTF1 | 3 | 3 | -1,29111E-15 | 0,999999892 | 1 |

| | | | | | |
|---|---|---|---|---|---|
| RHOT1;RHOT2 | 1 | 1 | 0 | 1 | 1 |
| GPD1 | 2 | 2 | 0 | 1 | 1 |
| PUS7 | 1 | 1 | 0 | 1 | 1 |
| CHMP5 | 1 | 1 | 0 | 1 | 1 |
| KRT77;KRT79;KRT5 | 1 | 1 | 0 | 1 | 1 |
| FMNL2;FMNL3;FMI | 1 | 1 | 0 | 1 | 1 |
| ARHGAP30 | 2 | 2 | 0,008864647 | 0,755844507 | 1 |
| TNFAIP8L3 | 3 | 3 | 2,1293E-16 | 0,999999962 | 1 |
| ARHGAP31 | 2 | 2 | 0 | 1 | 1 |
| ATP7B | 1 | 1 | 0 | 1 | 1 |
| LCLAT1 | 5 | 5 | -6,79985E-18 | 0,999999993 | 1 |
| KRT79 | 1 | 1 | 0 | 1 | 1 |
| MRPL18 | 1 | 1 | -0,17949916 | 0,155064547 | 1 |
| SMAP2;SMAP1 | 1 | 1 | -0,012918798 | 0,70630492 | 1 |
| ZW10 | 2 | 2 | 0 | 1 | 1 |
| LRRC7;ERBIN | 2 | 2 | -3,31787E-16 | 0,999999985 | 1 |
| COX7C | 2 | 2 | 0 | 1 | 1 |
| INPP4A | 1 | 1 | 0 | 1 | 1 |
| AMBRA1 | 1 | 1 | -0,154248853 | 0,33424706 | 1 |
| SLC7A2 | 4 | 4 | -5,9035E-17 | 0,99999998 | 1 |
| LARP4B | 1 | 1 | -0,073906994 | 0,30271897 | 1 |
| IGFALS | 1 | 1 | 0 | 1 | 1 |
| PSMG3 | 2 | 2 | -2,06163E-16 | 0,999999981 | 1 |
| FAM163B | 1 | 1 | 0 | 1 | 1 |
| FBL | 1 | 1 | 0 | 1 | 1 |
| ATL2 | 4 | 4 | -4,04251E-16 | 0,999999967 | 1 |
| ENSA | 2 | 2 | -5,19831E-19 | 1 | 1 |
| PALS1 | 2 | 2 | -0,111137637 | 0,18454587 | 1 |
| DTX4 | 1 | 1 | 0,145431645 | 0,183232568 | 1 |
| CNTD1 | 1 | 1 | 0 | 1 | 1 |
| TUBB4B;TUBB5;TU | 4 | 4 | -1,24702E-15 | 0,999999968 | 1 |
| PSME4 | 2 | 2 | 1,7842E-19 | 1 | 1 |
| FMO3 | 1 | 1 | 0 | 1 | 1 |

| | | | | | |
|---|---|---|---|---|---|
| FAN1 | 2 | 2 | 0 | 1 | 1 |
| KHDC1A;KHDC1C | 1 | 1 | 0 | 1 | 1 |
| STK32B | 1 | 1 | 0 | 1 | 1 |
| LRRC4B;LRRC4C;LR | 1 | 1 | 0,069121383 | 0,268489635 | 1 |
| STK26 | 1 | 1 | 0 | 1 | 1 |
| CDYL | 1 | 1 | 0 | 1 | 1 |
| NAA15 | 1 | 1 | -1,81297E-18 | 0,999999997 | 1 |
| CNIH2 | 2 | 2 | -0,062550977 | 0,34614692 | 1 |
| GM14151 | 1 | 1 | 0,010301396 | 0,756228505 | 1 |
| LIAS | 4 | 4 | 0 | 1 | 1 |
| IGHA | 1 | 1 | 0 | 1 | 1 |
| CHMP1B2 | 1 | 1 | 0 | 1 | 1 |
| HEBP2 | 1 | 1 | -0,152094041 | 0,343194888 | 1 |
| CHMP1B1 | 1 | 1 | 0 | 1 | 1 |
| SLC6A13 | 1 | 1 | 0 | 1 | 1 |
| PEX5L | 3 | 2 | 7,01966E-18 | 0,999999996 | 1 |
| AQP1 | 2 | 2 | 0 | 1 | 1 |
| PECR | 1 | 1 | 0,116191324 | 0,267570343 | 1 |
| MOCS2 | 2 | 2 | -7,43647E-20 | 1 | 1 |
| C1QTNF4 | 4 | 4 | 0,055690505 | 0,230050954 | 1 |
| SNRPD1 | 1 | 1 | 0 | 1 | 1 |
| KCNA1;KCNA3;KCN | 1 | 1 | 0 | 1 | 1 |
| LRRFIP2 | 1 | 1 | 0 | 1 | 1 |
| SENP8 | 2 | 2 | 0 | 1 | 1 |
| RABIF | 3 | 3 | 0,013798436 | 0,632485689 | 1 |
| SLC24A2 | 4 | 4 | 0,066914773 | 0,266159944 | 1 |
| MOV10L1 | 1 | 1 | 0 | 1 | 1 |
| ATAD5 | 1 | 1 | -0,073949743 | 0,567181291 | 1 |
| SCPEP1 | 1 | 1 | 0 | 1 | 1 |
| UBXN4 | 1 | 1 | 0 | 1 | 1 |
| TMSB15B1 | 1 | 1 | 0 | 1 | 1 |
| CC2D1A | 1 | 1 | 0 | 1 | 1 |
| PPP1R37 | 2 | 2 | 1,14394E-15 | 0,999999947 | 1 |

| | | | | | |
|---|---|---|---|---|---|
| IGHG1 | 1 | 1 | 0 | 1 | 1 |
| DDX20 | 1 | 1 | 0 | 1 | 1 |
| PHOSPHO2 | 1 | 1 | 0 | 1 | 1 |
| AMT | 2 | 1 | 0 | 1 | 1 |
| CDC42 | 2 | 2 | 0 | 1 | 1 |
| CDC42 | 2 | 2 | -0,06179204 | 0,329150095 | 1 |
| EVA1A | 1 | 1 | 0 | 1 | 1 |
| EFCAB9 | 1 | 1 | -3,90732E-20 | 1 | 1 |
| SLC25A17 | 1 | 1 | 0 | 1 | 1 |
| FGFR1;FGFR3 | 1 | 1 | 0 | 1 | 1 |
| MRPL23 | 4 | 4 | 0 | 1 | 1 |
| CTSZ | 2 | 2 | 2,89538E-16 | 0,999999969 | 1 |
| DCX;DCLK1 | 1 | 1 | 0 | 1 | 1 |
| DNM1L | 3 | 3 | -0,01480054 | 0,640968596 | 1 |
| CTSC | 1 | 1 | 0 | 1 | 1 |
| TRIM9 | 1 | 1 | 0 | 1 | 1 |
| EPB41L2;EPB41L3 | 1 | 1 | 0 | 1 | 1 |
| SINHCAF | 1 | 1 | 0 | 1 | 1 |
| GRB10;GRB14 | 1 | 1 | 0 | 1 | 1 |
| RNF31 | 3 | 3 | 8,75071E-17 | 0,999999992 | 1 |
| sp\|P0C913\|OCC1_ | 1 | 1 | 0 | 1 | 1 |
| D6WSU163E | 2 | 2 | 0 | 1 | 1 |
| RUFY1;RUFY3 | 1 | 1 | 0 | 1 | 1 |
| MSANTD4 | 2 | 2 | 0,106947863 | 0,221394519 | 1 |
| GM35060 | 1 | 1 | 0 | 1 | 1 |
| SEC23B | 1 | 1 | 0 | 1 | 1 |
| IKZF2 | 1 | 1 | 0 | 1 | 1 |
| CRTC1 | 9 | 9 | -0,105859657 | 0,128891545 | 1 |
| ZC3H12B | 1 | 1 | 0 | 1 | 1 |
| SYN1;SYN2 | 7 | 7 | 0,001901371 | 0,862308544 | 1 |
| OAF | 1 | 1 | -3,92504E-21 | 1 | 1 |
| GZMK | 1 | 1 | 0 | 1 | 1 |
| ZMYND15 | 1 | 1 | 0,170955286 | 0,322116545 | 1 |

| | | | | | |
|---|---|---|---|---|---|
| FOXL2 | 1 | 1 | 0 | 1 | 1 |
| ZHX1 | 1 | 1 | 0 | 1 | 1 |
| ADAMTS4 | 1 | 1 | -3,53596E-17 | 0,999999992 | 1 |
| TMPO | 3 | 3 | 3,19724E-19 | 0,999999999 | 1 |
| PINX1 | 1 | 1 | 0 | 1 | 1 |
| CMYA5 | 1 | 1 | 0 | 1 | 1 |
| POU4F3 | 1 | 1 | -0,024558379 | 0,652108451 | 1 |
| ZNF609 | 1 | 1 | 0 | 1 | 1 |
| FER1L6 | 1 | 1 | 0 | 1 | 1 |
| SOX15;SOX16 | 1 | 1 | 0 | 1 | 1 |
| EGF | 1 | 1 | -7,44846E-22 | 1 | 1 |
| SLC25A44 | 1 | 1 | 0 | 1 | 1 |
| COX17 | 1 | 1 | 0 | 1 | 1 |
| RELCH | 1 | 1 | -0,117332556 | 0,259954465 | 1 |
| ECI3 | 1 | 1 | 0 | 1 | 1 |
| KDM4B | 1 | 1 | 0 | 1 | 1 |
| LATS2 | 1 | 1 | -0,137907676 | 0,154425537 | 1 |
| ARHGAP6 | 1 | 1 | -0,156825107 | 0,223849857 | 1 |
| OLFR1257 | 1 | 1 | 0 | 1 | 1 |
| OLFR1228;OLFR122 | 1 | 1 | 0 | 1 | 1 |
| RANBP17;XPO7 | 1 | 1 | 0 | 1 | 1 |
| DGKI;DGKZ | 2 | 2 | 4,35169E-18 | 0,999999997 | 1 |
| BAIAP2 | 3 | 3 | 0 | 1 | 1 |
| GAL3ST4 | 1 | 1 | 6,36504E-22 | 1 | 1 |
| GSTM2;GSTM4 | 3 | 3 | 0 | 1 | 1 |
| FMNL3 | 1 | 1 | 0 | 1 | 1 |
| SH2D7 | 1 | 1 | 0 | 1 | 1 |
| ACVRL1 | 1 | 1 | 0 | 1 | 1 |
| IRX2 | 1 | 1 | 0 | 1 | 1 |
| TMEM134 | 1 | 1 | 0 | 1 | 1 |
| LPP | 1 | 1 | -2,80224E-17 | 0,999999988 | 1 |
| CHRNB1 | 1 | 1 | 1,39136E-18 | 0,999999998 | 1 |
| MPLKIP | 1 | 1 | 0 | 1 | 1 |

| | | | | | |
|---|---|---|---|---|---|
| SYN1 | 3 | 3 | -6,36314E-18 | 0,999999998 | 1 |
| PARP1 | 1 | 1 | 0 | 1 | 1 |
| MORC2A | 1 | 1 | 0 | 1 | 1 |
| CHD1 | 1 | 1 | 0 | 1 | 1 |
| PCBP2 | 1 | 1 | 0 | 1 | 1 |
| PCBP2 | 1 | 1 | -0,134026538 | 0,119927667 | 1 |
| COL6A3 | 1 | 1 | 0 | 1 | 1 |
| NCKAP5 | 1 | 1 | 0,047059916 | 0,435543795 | 1 |
| NECAP2 | 1 | 1 | 0,095338379 | 0,549923299 | 1 |
| TONSL | 2 | 2 | -0,093562228 | 0,251943503 | 1 |
| PLCH1 | 1 | 1 | 0 | 1 | 1 |
| EPHA1 | 1 | 1 | 0 | 1 | 1 |
| MYO1B | 1 | 1 | 0 | 1 | 1 |
| BSN;PCLO | 3 | 3 | 0 | 1 | 1 |
| ADNP | 1 | 1 | 0 | 1 | 1 |
| KLC1;KLC2 | 4 | 4 | 5,29008E-18 | 0,999999996 | 1 |
| CPLX1;CPLX2 | 5 | 5 | -4,64947E-17 | 0,999999986 | 1 |
| RECQL4 | 1 | 1 | 0 | 1 | 1 |
| SNX25 | 2 | 1 | -0,185528124 | 0,114666153 | 1 |
| DRGX | 1 | 1 | 0 | 1 | 1 |
| DBI | 4 | 4 | 2,62194E-15 | 0,999999854 | 1 |
| DGKB | 1 | 1 | 0 | 1 | 1 |
| TMCC1;TMCC2 | 1 | 1 | 0 | 1 | 1 |
| SGTA | 1 | 1 | -0,177340185 | 0,213549756 | 1 |
| 4930402K13RIK | 1 | 1 | 0 | 1 | 1 |
| CAMK2A;CAMK2D | 2 | 2 | 7,86205E-16 | 0,999999947 | 1 |
| MACF1 | 2 | 1 | -0,009278931 | 0,831434504 | 1 |
| STXBP5L | 1 | 1 | 0 | 1 | 1 |
| MATN4 | 5 | 5 | -4,94102E-18 | 0,999999998 | 1 |
| CYTH3;CYTH2;CYTH | 1 | 1 | 0 | 1 | 1 |
| NUDT14 | 3 | 3 | 0 | 1 | 1 |
| SNX3;SNX12 | 2 | 2 | 0 | 1 | 1 |
| TRUB1 | 2 | 2 | 0 | 1 | 1 |

| | | | | | |
|---|---|---|---|---|---|
| MAPRE3 | 1 | 1 | -0,12252342 | 0,173208979 | 1 |
| SPTAN1 | 2 | 2 | 0,074129534 | 0,462241765 | 1 |
| SETD6 | 1 | 1 | 0 | 1 | 1 |
| SNCB;SNCG | 2 | 2 | -3,03782E-16 | 0,999999968 | 1 |
| RNF157 | 1 | 1 | 0 | 1 | 1 |
| MAP3K7CL | 1 | 1 | 0 | 1 | 1 |
| TMEM129 | 1 | 1 | 0 | 1 | 1 |
| CDR2 | 1 | 1 | 0,085717349 | 0,69373205 | 1 |
| MANF | 2 | 2 | -3,3015E-16 | 0,99999996 | 1 |
| TRIM33 | 1 | 1 | 0 | 1 | 1 |
| HROB | 1 | 1 | 0,01696302 | 0,674533406 | 1 |
| MAP3K2 | 1 | 1 | 0 | 1 | 1 |
| MYO5C | 1 | 1 | 0 | 1 | 1 |
| SGSM2;SGSM1 | 1 | 1 | 0 | 1 | 1 |
| CRY1 | 1 | 1 | -0,013959664 | 0,765683286 | 1 |
| WWC2 | 2 | 2 | 0 | 1 | 1 |
| ASIC1 | 1 | 1 | 0 | 1 | 1 |
| STX1A;STX1B | 1 | 1 | 0 | 1 | 1 |
| ACTN4 | 2 | 2 | 0 | 1 | 1 |
| MAK | 2 | 2 | 2,11842E-16 | 0,999999987 | 1 |
| BCKDK | 4 | 4 | 0 | 1 | 1 |
| CYTH4 | 2 | 1 | -0,090848961 | 0,260428959 | 1 |
| CLN6 | 1 | 1 | 0 | 1 | 1 |
| CDH2;CDH4 | 2 | 2 | 0 | 1 | 1 |
| NRBF2 | 2 | 2 | 0 | 1 | 1 |
| RBM15B | 1 | 1 | 0 | 1 | 1 |
| SNX3 | 1 | 1 | 0 | 1 | 1 |
| SNX12 | 1 | 1 | 0 | 1 | 1 |
| RABEPK | 3 | 3 | 0,055292985 | 0,274327635 | 1 |
| RIPOR3 | 1 | 1 | 0 | 1 | 1 |
| KRT34 | 1 | 1 | 0,044344724 | 0,509978855 | 1 |
| ACTG1 | 1 | 1 | 0 | 1 | 1 |
| TCP11L2 | 1 | 1 | 0 | 1 | 1 |

| | | | | | |
|---|---|---|---|---|---|
| TRPC6 | 1 | 1 | 0 | 1 | 1 |
| TMEFF1 | 2 | 2 | 0 | 1 | 1 |
| RPL39 | 1 | 1 | 0 | 1 | 1 |
| DLGAP1 | 1 | 1 | 0 | 1 | 1 |
| PCDHGC5;PCDHGA | 2 | 2 | 0 | 1 | 1 |
| ATG4B | 1 | 1 | 0 | 1 | 1 |
| QRICH2 | 1 | 1 | 5,53277E-15 | 0,999999852 | 1 |
| ARAF | 1 | 1 | 0 | 1 | 1 |
| KSR2 | 2 | 2 | 0 | 1 | 1 |
| SNU13 | 1 | 1 | 0 | 1 | 1 |
| GCNT3 | 1 | 1 | 0 | 1 | 1 |
| CUBN | 1 | 1 | -0,159080765 | 0,408259885 | 1 |
| SSR3 | 1 | 1 | 0 | 1 | 1 |
| ARFGAP1 | 1 | 1 | 0 | 1 | 1 |
| MMTAG2 | 2 | 2 | -0,09594197 | 0,369877263 | 1 |
| TGM4 | 1 | 1 | 0 | 1 | 1 |
| SCAF4 | 1 | 1 | 0 | 1 | 1 |
| EEF1D | 1 | 1 | 0 | 1 | 1 |
| CLASP1 | 2 | 2 | 1,67931E-10 | 0,999990784 | 1 |
| KRT10;KRT13 | 2 | 2 | -3,64676E-17 | 0,999999997 | 1 |
| TIA1;TIAL1 | 3 | 3 | -0,026608731 | 0,509972398 | 1 |
| TMEM186 | 1 | 1 | 0 | 1 | 1 |
| PPP3CA | 2 | 2 | 0 | 1 | 1 |
| CDCA2 | 1 | 1 | -0,050155044 | 0,565918036 | 1 |
| NSD2 | 1 | 1 | -0,070574749 | 0,496111872 | 1 |
| KRT12 | 2 | 1 | 0 | 1 | 1 |
| RBX1 | 1 | 1 | -6,85419E-19 | 0,999999999 | 1 |
| MYO10 | 1 | 1 | 0 | 1 | 1 |
| YARS1 | 2 | 2 | -1,86469E-15 | 0,999999933 | 1 |
| RHOB;RHOC;RHOA | 1 | 1 | 0 | 1 | 1 |
| HCN2;HCN1;HCN3 | 1 | 1 | 0 | 1 | 1 |
| TEX28 | 1 | 1 | 0 | 1 | 1 |
| TLE5 | 1 | 1 | 0 | 1 | 1 |

| | | | | | |
|---|---|---|---|---|---|
| MFGE8 | 4 | 4 | 1,48093E-15 | 0,999999899 | 1 |
| KLC3 | 1 | 1 | 0 | 1 | 1 |
| TRIML1 | 1 | 1 | 0 | 1 | 1 |
| KCNA1;KCNA3;KCN | 1 | 1 | 0 | 1 | 1 |
| SURF4 | 2 | 1 | 0 | 1 | 1 |
| CNBD1 | 1 | 1 | 0 | 1 | 1 |
| CHCHD2 | 1 | 1 | 0 | 1 | 1 |
| IFT81 | 1 | 1 | 0 | 1 | 1 |
| MYH3 | 1 | 1 | -0,092652649 | 0,297905422 | 1 |
| NGLY1 | 1 | 1 | 0 | 1 | 1 |
| CCDC150 | 1 | 1 | 0 | 1 | 1 |
| CHD7;CHD9 | 1 | 1 | 0 | 1 | 1 |
| LRRC75A | 1 | 1 | 0,295292522 | 0,191925774 | 1 |
| SNRPD2 | 3 | 3 | -2,6855E-17 | 0,999999989 | 1 |
| HIPK2 | 1 | 1 | 0,085269673 | 0,267636606 | 1 |
| CSRNP1 | 1 | 1 | 0 | 1 | 1 |
| DENND11 | 3 | 3 | 0,046073894 | 0,247802231 | 1 |
| EBAG9 | 1 | 1 | 0 | 1 | 1 |
| CC2D1B | 1 | 1 | 0,158058462 | 0,404196343 | 1 |
| PURB;PURA;PURG | 1 | 1 | 0 | 1 | 1 |
| SELENOW | 1 | 1 | 0 | 1 | 1 |
| ABCD2 | 2 | 1 | 5,22918E-21 | 1 | 1 |
| ESF1 | 1 | 1 | 0 | 1 | 1 |
| L1TD1 | 1 | 1 | 2,71926E-17 | 0,999999991 | 1 |
| ENPP2 | 1 | 1 | 0,033469087 | 0,641029527 | 1 |
| TMPRSS11B | 1 | 1 | 0 | 1 | 1 |
| MFAP3;MFAP3L | 1 | 1 | 0 | 1 | 1 |
| TAF1 | 1 | 1 | 0,063283759 | 0,352745604 | 1 |
| PHKA2 | 2 | 2 | 0 | 1 | 1 |
| LRRC4B;LRRC4C | 1 | 1 | 0,050524618 | 0,4531025 | 1 |
| SARAF | 1 | 1 | 0 | 1 | 1 |
| SPARC | 1 | 1 | 0 | 1 | 1 |
| TEKT2 | 1 | 1 | 0 | 1 | 1 |

| | | | | | |
|---|---|---|---|---|---|
| RBFOX2 | 1 | 1 | -0,045666654 | 0,540057174 | 1 |
| ADSS1;ADSS2 | 5 | 5 | 3,17396E-17 | 0,999999994 | 1 |
| COX18 | 1 | 1 | 0 | 1 | 1 |
| SYNGAP1 | 1 | 1 | 0 | 1 | 1 |
| HLTF | 2 | 2 | -1,62823E-19 | 1 | 1 |
| ANGPTL8 | 1 | 1 | 0 | 1 | 1 |
| SERPINA16 | 1 | 1 | 1,29649E-19 | 1 | 1 |
| CUX1 | 1 | 1 | 0 | 1 | 1 |
| NPTN | 2 | 2 | 0 | 1 | 1 |
| ORM2;ORM1 | 1 | 1 | 3,55453E-17 | 0,999999992 | 1 |
| AGAP3 | 1 | 1 | 0,038069381 | 0,499651123 | 1 |
| CCM2 | 2 | 2 | 0 | 1 | 1 |
| UBE2J1 | 1 | 1 | 2,28739E-20 | 1 | 1 |
| CACNB1 | 1 | 1 | 0 | 1 | 1 |
| QTRT2 | 2 | 2 | 4,08622E-17 | 0,999999987 | 1 |
| RECQL | 1 | 1 | -0,111712984 | 0,183785456 | 1 |
| DCAF8 | 2 | 1 | -0,094304939 | 0,297025744 | 1 |
| NIN | 1 | 1 | 0 | 1 | 1 |
| MEIS3 | 1 | 1 | 0 | 1 | 1 |
| HSPA12A;HSPA12B | 2 | 2 | -1,18945E-18 | 0,999999999 | 1 |
| RALGAPA2 | 1 | 1 | 0,005112144 | 0,820790177 | 1 |
| CACNB2;CACNB4;C | 4 | 4 | 0 | 1 | 1 |
| PTMS | 4 | 4 | 0 | 1 | 1 |
| ADAMTS17 | 1 | 1 | 0 | 1 | 1 |
| KRT1 | 34 | 4 | 0 | 1 | 1 |
| TMC7 | 1 | 1 | 0 | 1 | 1 |
| ARHGAP25 | 1 | 1 | 0 | 1 | 1 |
| SMAD1;SMAD5 | 2 | 1 | 0 | 1 | 1 |
| LDAH | 1 | 1 | 0 | 1 | 1 |
| GOLT1B | 2 | 2 | -3,89032E-19 | 0,999999999 | 1 |
| ALDH1A1;ALDH1A2 | 1 | 1 | 0,094772799 | 0,218866662 | 1 |
| CHCHD3 | 2 | 2 | -5,72145E-17 | 0,999999993 | 1 |
| SNX32 | 3 | 3 | 0 | 1 | 1 |

| | | | | | |
|---|---|---|---|---|---|
| PIP5K1B | 2 | 2 | -0,106914664 | 0,158988307 | 1 |
| DNAJC19 | 1 | 1 | -0,332934698 | 0,160421142 | 1 |
| SNX7 | 1 | 1 | 0 | 1 | 1 |
| ZC3H14 | 1 | 1 | 0 | 1 | 1 |
| MAL2 | 2 | 2 | 1,19444E-15 | 0,999999907 | 1 |
| HCN3 | 1 | 1 | 0 | 1 | 1 |
| KEAP1 | 1 | 1 | 0 | 1 | 1 |
| RAP2A;RAP2C | 1 | 1 | 0,122447713 | 0,144841147 | 1 |
| SST | 1 | 1 | 0 | 1 | 1 |
| SPATA2 | 3 | 3 | -0,103440977 | 0,166323349 | 1 |
| ENTPD6 | 1 | 1 | 0 | 1 | 1 |
| SSBP2;SSBP3 | 1 | 1 | 0 | 1 | 1 |
| FOXO4 | 1 | 1 | 0 | 1 | 1 |
| ATP2B3 | 2 | 2 | 0 | 1 | 1 |
| DMAC1 | 1 | 1 | 0 | 1 | 1 |
| SFN | 14 | 2 | 0 | 1 | 1 |
| RTN3 | 4 | 4 | 0 | 1 | 1 |
| GOLPH3 | 1 | 1 | 0 | 1 | 1 |
| NTAQ1 | 1 | 1 | -0,075833386 | 0,344055711 | 1 |
| ERICH6B | 1 | 1 | 0 | 1 | 1 |
| XKR7 | 1 | 1 | 0 | 1 | 1 |
| DAAM2;DAAM1 | 3 | 3 | 0 | 1 | 1 |
| FER | 2 | 2 | 0 | 1 | 1 |
| FAM107B | 1 | 1 | 0 | 1 | 1 |
| LEXM | 1 | 1 | 0 | 1 | 1 |
| CTNND1 | 1 | 1 | 0 | 1 | 1 |
| PPCDC | 2 | 2 | 7,68465E-18 | 0,999999996 | 1 |
| NACAD | 1 | 1 | -0,00414309 | 0,866441773 | 1 |
| NCBP2 | 1 | 1 | 0 | 1 | 1 |
| PIK3IP1 | 1 | 1 | 0,032188211 | 0,662836418 | 1 |
| RB1 | 2 | 2 | 8,84211E-12 | 0,99999209 | 1 |
| GRAMD4 | 1 | 1 | 0 | 1 | 1 |
| ADAMTS16 | 1 | 1 | 0,059046321 | 0,570862031 | 1 |

| | | | | | |
|---|---|---|---|---|---|
| ATG14 | 1 | 1 | 0 | 1 | 1 |
| RASSF2 | 2 | 2 | 0,01904875 | 0,647151937 | 1 |
| TMEM132E | 2 | 2 | -8,78658E-18 | 0,999999997 | 1 |
| PDLIM5 | 1 | 1 | 0 | 1 | 1 |
| SPOCK2 | 2 | 2 | 5,27267E-16 | 0,999999961 | 1 |
| AP3S1 | 1 | 1 | -6,26794E-19 | 0,999999999 | 1 |
| INSRR | 1 | 1 | 0 | 1 | 1 |
| DCTN5 | 1 | 1 | 0,025273801 | 0,595665599 | 1 |
| NRAS | 1 | 1 | 0 | 1 | 1 |
| HRAS | 3 | 3 | 0,029639782 | 0,410695716 | 1 |
| LSM14B | 1 | 1 | -0,009111069 | 0,766279747 | 1 |
| RNPEPL1 | 1 | 1 | 0 | 1 | 1 |
| TAB1 | 3 | 2 | 2,6258E-14 | 0,999999832 | 1 |
| INPP5K | 2 | 2 | -8,9454E-18 | 0,999999994 | 1 |
| GM21876 | 1 | 1 | 0 | 1 | 1 |
| CD63 | 1 | 1 | 0 | 1 | 1 |
| OSBPL11 | 2 | 2 | 0 | 1 | 1 |
| INPP5E | 1 | 1 | 0 | 1 | 1 |
| PHACTR4 | 1 | 1 | -2,13009E-18 | 0,999999997 | 1 |
| SMDT1 | 1 | 1 | 0 | 1 | 1 |
| CDIP1 | 1 | 1 | 0 | 1 | 1 |
| TMOD4 | 1 | 1 | 0 | 1 | 1 |
| SLCO3A1 | 1 | 1 | 0 | 1 | 1 |
| SRP19 | 1 | 1 | 0 | 1 | 1 |
| PRRC1 | 1 | 1 | -0,179606281 | 0,125782085 | 1 |
| 6330409D20RIK | 1 | 1 | 0 | 1 | 1 |
| NR1H4 | 1 | 1 | 0 | 1 | 1 |
| CMTM6 | 1 | 1 | 0 | 1 | 1 |
| CROT | 1 | 1 | 1,22623E-13 | 0,999999166 | 1 |
| HEPH | 1 | 1 | 0 | 1 | 1 |
| EGLN1 | 2 | 2 | 0 | 1 | 1 |
| CSMD3 | 1 | 1 | 0,156137052 | 0,541508628 | 1 |
| CASP3 | 1 | 1 | 0 | 1 | 1 |

| | | | | | |
|---|---|---|---|---|---|
| ATP6V0C | 4 | 4 | -6,43039E-18 | 0,999999998 | 1 |
| PABPC4L;PABPC4;F | 1 | 1 | -0,282467354 | 0,27482354 | 1 |
| SLC12A3 | 1 | 1 | -4,00174E-21 | 1 | 1 |
| H1-4;H1-3 | 3 | 1 | 0 | 1 | 1 |
| ITSN1 | 1 | 1 | -0,187841689 | 0,182002261 | 1 |
| SLC5A6 | 1 | 1 | 0 | 1 | 1 |
| SLC38A2 | 1 | 1 | 0 | 1 | 1 |
| OLFM3 | 1 | 1 | 0 | 1 | 1 |
| TPM1 | 1 | 1 | 0 | 1 | 1 |
| DEPDC5 | 1 | 1 | -8,47018E-18 | 0,999999994 | 1 |
| AGBL1 | 1 | 1 | 0,011068438 | 0,733825187 | 1 |
| DNER | 1 | 1 | 0 | 1 | 1 |
| LEKR1 | 1 | 1 | 0 | 1 | 1 |
| SYNGAP1 | 1 | 1 | 0 | 1 | 1 |
| RYR3;RYR1;RYR2 | 3 | 3 | 2,1231E-11 | 0,999990949 | 1 |
| GARNL3 | 1 | 1 | 0 | 1 | 1 |
| CHCHD1 | 1 | 1 | 0 | 1 | 1 |
| AASDHPPT | 1 | 1 | 0 | 1 | 1 |
| SLC6A4 | 1 | 1 | 0,031037297 | 0,595878076 | 1 |
| BACE1 | 1 | 1 | 0 | 1 | 1 |
| RGSL1 | 1 | 1 | 0,126300316 | 0,243095162 | 1 |
| SYTL5 | 1 | 1 | 0 | 1 | 1 |
| SNUPN | 1 | 1 | 0 | 1 | 1 |
| CCNYL1 | 2 | 2 | 0 | 1 | 1 |
| BET1L | 1 | 1 | 0 | 1 | 1 |
| PKN1 | 1 | 1 | 0 | 1 | 1 |
| ARPP21 | 3 | 3 | 0 | 1 | 1 |
| CTNND1 | 1 | 1 | 0,158551758 | 0,429285889 | 1 |
| SNX13 | 1 | 1 | 0,128347263 | 0,261562431 | 1 |
| TC2N | 1 | 1 | 0 | 1 | 1 |
| TPM3;TPM1 | 1 | 1 | 0 | 1 | 1 |
| OSBPL7 | 1 | 1 | 0 | 1 | 1 |
| SCMH1 | 1 | 1 | 0 | 1 | 1 |

| | | | | | |
|---|---|---|---|---|---|
| IL19 | 1 | 1 | 0 | 1 | 1 |
| FHIP2B | 1 | 1 | 0 | 1 | 1 |
| DACT3 | 1 | 1 | 2,14511E-15 | 0,999999897 | 1 |
| ATP11C | 1 | 1 | 0 | 1 | 1 |
| CYP2D10 | 1 | 1 | 0 | 1 | 1 |
| PHKG2 | 1 | 1 | 0 | 1 | 1 |
| SPOCK1 | 1 | 1 | 0,012224918 | 0,730762506 | 1 |
| OVOS | 1 | 1 | -0,112302144 | 0,386091969 | 1 |
| TTC23L | 1 | 1 | 0 | 1 | 1 |
| FRMPD4 | 3 | 3 | -0,102823296 | 0,158034563 | 1 |
| KRT78 | 5 | 2 | 0 | 1 | 1 |
| SLIT1;SLIT2 | 1 | 1 | 0 | 1 | 1 |
| PTPRR | 3 | 3 | 2,63353E-18 | 0,999999997 | 1 |
| MVK | 5 | 5 | -1,5606E-16 | 0,999999968 | 1 |
| UBE2R2 | 2 | 1 | 1,36422E-17 | 0,999999993 | 1 |
| ARL14EP | 1 | 1 | 1,39243E-20 | 1 | 1 |
| ARIH2 | 4 | 4 | 0 | 1 | 1 |
| PLEKHA6 | 2 | 2 | -6,3275E-18 | 0,999999996 | 1 |
| BET1 | 1 | 1 | -1,15119E-17 | 0,999999992 | 1 |
| SRA1 | 1 | 1 | 0 | 1 | 1 |
| SFXN4 | 1 | 1 | 0 | 1 | 1 |
| GM2A | 2 | 2 | 9,887E-18 | 0,999999995 | 1 |
| PRSS16 | 1 | 1 | 0,33712138 | 0,119272129 | 1 |
| CCS | 2 | 2 | -1,62415E-17 | 0,999999994 | 1 |
| FBXL12 | 1 | 1 | 0 | 1 | 1 |
| GLI3 | 1 | 1 | 0 | 1 | 1 |
| ADD1 | 1 | 1 | 0 | 1 | 1 |
| SMPDL3A | 1 | 1 | 0 | 1 | 1 |
| TUBGCP5 | 1 | 1 | 0,192725413 | 0,155156477 | 1 |
| TGS1 | 1 | 1 | 0 | 1 | 1 |
| MYH11;MYH14 | 1 | 1 | 0 | 1 | 1 |
| RAP2C | 5 | 4 | 0 | 1 | 1 |
| POGLUT2 | 1 | 1 | 0 | 1 | 1 |

| | | | | | |
|---|---|---|---|---|---|
| BAG1 | 1 | 1 | 0 | 1 | 1 |
| IL6ST | 1 | 1 | 0 | 1 | 1 |
| ANKRD50 | 1 | 1 | 0 | 1 | 1 |
| IRS2 | 2 | 2 | 2,76119E-17 | 0,99999999 | 1 |
| 1700009N14RIK | 1 | 1 | 0 | 1 | 1 |
| DHX29 | 2 | 2 | 0,039814673 | 0,481049455 | 1 |
| MBP | 2 | 2 | -5,10646E-14 | 0,99999949 | 1 |
| SMARCD1 | 1 | 1 | 0 | 1 | 1 |
| HTR3A | 1 | 1 | 0 | 1 | 1 |
| PKP1 | 5 | 1 | 0 | 1 | 1 |
| PAQR9 | 1 | 1 | -0,211820547 | 0,132799766 | 1 |
| RETREG3 | 2 | 2 | 2,05822E-16 | 0,999999984 | 1 |
| SYNDIG1 | 1 | 1 | 0 | 1 | 1 |
| CCDC141 | 1 | 1 | 0,024052875 | 0,601956968 | 1 |
| TNFRSF13B | 1 | 1 | -0,038363602 | 0,581774336 | 1 |
| FAM234A | 1 | 1 | 0 | 1 | 1 |
| AAGAB | 2 | 2 | 1,19334E-17 | 0,999999998 | 1 |
| SPX | 1 | 1 | 0 | 1 | 1 |
| CFAP99 | 1 | 1 | 0 | 1 | 1 |
| PTBP3 | 2 | 2 | -3,53317E-19 | 0,999999999 | 1 |
| AFTPH | 3 | 3 | 0,049219225 | 0,239226573 | 1 |
| IFT27 | 1 | 1 | 0 | 1 | 1 |
| HYDIN | 1 | 1 | 0,144964063 | 0,190063628 | 1 |
| TRIM15 | 1 | 1 | -0,009636593 | 0,835869945 | 1 |
| SH3D21 | 1 | 1 | -0,0398645 | 0,455872824 | 1 |
| CX3CR1 | 1 | 1 | 0 | 1 | 1 |
| ZFP646 | 1 | 1 | 0,137825388 | 0,309465706 | 1 |
| AMOT | 1 | 1 | 0 | 1 | 1 |
| FRS3 | 1 | 1 | 0 | 1 | 1 |
| LAMA3 | 2 | 1 | 0,132561212 | 0,169568593 | 1 |
| CSNK1G2 | 1 | 1 | 0,119355783 | 0,238409592 | 1 |
| INF2 | 2 | 2 | -3,18429E-13 | 0,999998909 | 1 |
| TRMT10C | 3 | 1 | 0 | 1 | 1 |

| | | | | | |
|---|---|---|---|---|---|
| CYFIP1 | 2 | 1 | 0 | 1 | 1 |
| TMF1 | 1 | 1 | 0 | 1 | 1 |
| ZFYVE27 | 1 | 1 | 0 | 1 | 1 |
| SYNJ1 | 1 | 1 | 0 | 1 | 1 |
| SYNJ1 | 1 | 1 | 0 | 1 | 1 |
| TYRO3 | 1 | 1 | 0 | 1 | 1 |
| CAMK2A | 2 | 2 | 1,48278E-17 | 0,999999996 | 1 |
| ARL2BP | 2 | 2 | 0 | 1 | 1 |
| NAV3 | 2 | 2 | 0 | 1 | 1 |
| TMEM200A | 1 | 1 | 0 | 1 | 1 |
| FHIP2A | 1 | 1 | 0 | 1 | 1 |
| PCDHGA6 | 1 | 1 | 0 | 1 | 1 |
| ARHGEF6;ARHGEF7 | 2 | 2 | 3,38372E-16 | 0,999999954 | 1 |
| YTHDC2 | 1 | 1 | 0,069084653 | 0,571979829 | 1 |
| CCDC28A | 1 | 1 | 0 | 1 | 1 |
| HOXA13 | 1 | 1 | 0 | 1 | 1 |
| FAM117B | 2 | 2 | -0,090282974 | 0,213192898 | 1 |
| POLR2M | 1 | 1 | 0,155005688 | 0,199117676 | 1 |
| CLPTM1L | 3 | 3 | 0 | 1 | 1 |
| FEM1B | 1 | 1 | 0 | 1 | 1 |
| RNF146 | 1 | 1 | 0 | 1 | 1 |
| ANKS1B | 1 | 1 | 0 | 1 | 1 |
| ICA | 1 | 1 | 0 | 1 | 1 |
| DMAP1 | 1 | 1 | -0,01380045 | 0,751338611 | 1 |
| FASTKD2 | 1 | 1 | -0,173932275 | 0,364626077 | 1 |
| GTF2A1 | 1 | 1 | -0,06900574 | 0,430313998 | 1 |
| CDC123 | 1 | 1 | 0 | 1 | 1 |
| GNAT2 | 1 | 1 | 0,498235034 | 0,248418116 | 1 |
| SEPTIN7 | 3 | 3 | -2,71103E-19 | 1 | 1 |
| TRIM45 | 1 | 1 | 0 | 1 | 1 |
| AKAP2 | 3 | 2 | 2,10204E-19 | 1 | 1 |
| ANKS1B | 3 | 3 | 0 | 1 | 1 |
| SUSD5 | 1 | 1 | 0 | 1 | 1 |

| | | | | | |
|---|---|---|---|---|---|
| PPP1R16A | 1 | 1 | 0 | 1 | 1 |
| TRAK2 | 1 | 1 | 0 | 1 | 1 |
| B3GNT7 | 1 | 1 | 0 | 1 | 1 |
| RPRD1A | 1 | 1 | 0 | 1 | 1 |
| GPR17 | 1 | 1 | 9,97586E-20 | 1 | 1 |
| PTK2 | 1 | 1 | 0 | 1 | 1 |
| MDM1 | 1 | 1 | 0 | 1 | 1 |
| TSPAN14 | 1 | 1 | 0 | 1 | 1 |
| TENM4;TENM2;TEI | 1 | 1 | 0 | 1 | 1 |
| MSI1 | 1 | 1 | 0,044479151 | 0,623537827 | 1 |
| DPH2 | 1 | 1 | 0 | 1 | 1 |
| PLEKHB1 | 3 | 3 | 1,68682E-18 | 0,999999998 | 1 |
| SDF4 | 2 | 1 | 0 | 1 | 1 |
| MAPK10;MAPK8 | 5 | 4 | -3,17851E-19 | 0,999999999 | 1 |
| DCX;DCLK2 | 1 | 1 | 0 | 1 | 1 |
| PIK3R2;PIK3R1 | 1 | 1 | 0 | 1 | 1 |
| ABCC3 | 1 | 1 | 0 | 1 | 1 |
| CDH24 | 1 | 1 | 5,41119E-18 | 0,999999995 | 1 |
| TMEM63A | 1 | 1 | 0 | 1 | 1 |
| DNAJC30 | 2 | 1 | 4,38208E-18 | 0,999999997 | 1 |
| TSHZ1 | 1 | 1 | 0,055743618 | 0,515565962 | 1 |
| MEIOC | 1 | 1 | 0,050137248 | 0,586053902 | 1 |
| D17H6S53E | 1 | 1 | -0,151525512 | 0,142613159 | 1 |
| RHOB;RHOA | 1 | 1 | 0 | 1 | 1 |
| TMEM151A | 1 | 1 | 0 | 1 | 1 |
| CLUH | 1 | 1 | -0,079696538 | 0,240688428 | 1 |
| SDHAF2 | 1 | 1 | 0 | 1 | 1 |
| ADAL | 1 | 1 | 0 | 1 | 1 |
| MAP4 | 1 | 1 | 0 | 1 | 1 |
| TERF2 | 1 | 1 | 0 | 1 | 1 |
| POGLUT1 | 3 | 3 | -0,018128591 | 0,591475026 | 1 |
| CCDC134 | 2 | 2 | 0 | 1 | 1 |
| ANGEL2 | 1 | 1 | 0 | 1 | 1 |

| | | | | | |
|---|---|---|---|---|---|
| B3GALNT1 | 1 | 1 | 0 | 1 | 1 |
| RFXAP | 1 | 1 | 0,036691024 | 0,494441 | 1 |
| CHGA | 1 | 1 | 0 | 1 | 1 |
| VAMP8 | 1 | 1 | 0,100798768 | 0,561004844 | 1 |
| TCEA3 | 1 | 1 | 0 | 1 | 1 |
| SLC25A37 | 1 | 1 | 0 | 1 | 1 |
| DNM3 | 1 | 1 | 0 | 1 | 1 |
| SUMO3 | 1 | 1 | 0 | 1 | 1 |
| NPTX2 | 1 | 1 | 0 | 1 | 1 |
| ARHGAP17 | 1 | 1 | 0 | 1 | 1 |
| FCHO2 | 1 | 1 | -0,035051645 | 0,595924706 | 1 |
| PRKCG;PRKCB | 1 | 1 | 0,090617431 | 0,347848375 | 1 |
| SCN1A;SCN2A;SCN! | 1 | 1 | 0 | 1 | 1 |
| RAPGEF5 | 1 | 1 | -0,056504362 | 0,502570172 | 1 |
| TMEM68 | 2 | 2 | -0,014430126 | 0,634934163 | 1 |
| EI24 | 1 | 1 | 0 | 1 | 1 |
| MRPL53 | 2 | 1 | 0 | 1 | 1 |
| EPN2 | 1 | 1 | 0 | 1 | 1 |
| NLGN4L | 1 | 1 | 0 | 1 | 1 |
| POLR3G | 1 | 1 | -0,153130455 | 0,327208322 | 1 |
| FXYD1 | 1 | 1 | -1,65077E-17 | 0,999999992 | 1 |
| SEPTIN12 | 2 | 2 | 0 | 1 | 1 |
| 2010109A12RIK | 1 | 1 | -2,12949E-15 | 0,999999891 | 1 |
| CD300LF | 1 | 1 | 0 | 1 | 1 |
| TMEM19 | 1 | 1 | 0 | 1 | 1 |
| ACTN1 | 2 | 2 | 3,32065E-18 | 0,999999998 | 1 |
| CAMK2G | 1 | 1 | -0,085580019 | 0,219124573 | 1 |
| CAPN7 | 1 | 1 | 0 | 1 | 1 |
| FAM189A1 | 1 | 1 | 0 | 1 | 1 |
| TNIK;MINK1 | 2 | 2 | 0 | 1 | 1 |
| DAB2 | 1 | 1 | -1,75893E-19 | 1 | 1 |
| RELL2 | 1 | 1 | 0 | 1 | 1 |
| ATP5F1A | 3 | 2 | 0,023592522 | 0,472693987 | 1 |

| | | | | | |
|---|---|---|---|---|---|
| KMT2A | 1 | 1 | -3,66024E-18 | 0,999999996 | 1 |
| STAT1 | 1 | 1 | 0 | 1 | 1 |
| CTC1 | 2 | 2 | -6,61986E-18 | 0,999999998 | 1 |
| APOA2 | 1 | 1 | -0,245115153 | 0,152071314 | 1 |
| NDRG2 | 1 | 1 | 0 | 1 | 1 |
| RPS5 | 1 | 1 | 0 | 1 | 1 |
| ANKRD52 | 1 | 1 | 0 | 1 | 1 |
| TRMU | 4 | 3 | 0 | 1 | 1 |
| ARAF | 1 | 1 | 0 | 1 | 1 |
| PLCG2 | 1 | 1 | 0,142820264 | 0,31773075 | 1 |
| PRKD1;PRKD3 | 1 | 1 | 0,069318898 | 0,415131751 | 1 |
| SCN1A;SCN3A;SCN: | 1 | 1 | 0 | 1 | 1 |
| RAB3D;RAB8A;RAB | 1 | 1 | 0 | 1 | 1 |
| SCN1A;SCN3A;SCN: | 2 | 2 | 0,066017304 | 0,302527855 | 1 |
| ACVR1B | 1 | 1 | 5,54946E-19 | 0,999999999 | 1 |
| KCNT2 | 1 | 1 | 0 | 1 | 1 |
| CLIP1 | 1 | 1 | 0 | 1 | 1 |
| CACNB2;CACNB1 | 1 | 1 | 0 | 1 | 1 |
| FAT1 | 1 | 1 | 0 | 1 | 1 |
| RTN4RL1 | 1 | 1 | 1,33432E-17 | 0,999999993 | 1 |
| HEATR1 | 1 | 1 | 0 | 1 | 1 |
| MGAM | 1 | 1 | 0 | 1 | 1 |
| MBTPS1 | 1 | 1 | 0 | 1 | 1 |
| TRY10 | 2 | 2 | -0,058523003 | 0,782757176 | 1 |
| ABCA8B | 1 | 1 | 0 | 1 | 1 |
| SLC2A8 | 1 | 1 | 0,007747487 | 0,82893758 | 1 |
| FGD5 | 1 | 1 | 0 | 1 | 1 |
| RPS6KL1 | 1 | 1 | 0 | 1 | 1 |
| MTURN | 1 | 1 | -0,020740202 | 0,673727295 | 1 |
| FCGR2 | 1 | 1 | 0 | 1 | 1 |
| ABHD4 | 3 | 3 | -1,70241E-20 | 1 | 1 |
| NKIRAS1 | 1 | 1 | 0 | 1 | 1 |
| MOSPD2 | 1 | 1 | 0 | 1 | 1 |

| | | | | | |
|---|---|---|---|---|---|
| SCML2 | 1 | 1 | 0,019108761 | 0,69250953 | 1 |
| BNIP2 | 1 | 1 | 0 | 1 | 1 |
| CUL9 | 1 | 1 | -0,156524584 | 0,24591631 | 1 |
| DDHD1 | 1 | 1 | 0 | 1 | 1 |
| NDUFS2 | 1 | 1 | 0,072104428 | 0,300116466 | 1 |
| FBXO6 | 2 | 2 | 8,40439E-16 | 0,999999936 | 1 |
| PTPRS | 1 | 1 | 0 | 1 | 1 |
| BLOC1S1 | 1 | 1 | 0 | 1 | 1 |
| FZR1 | 1 | 1 | 0 | 1 | 1 |
| YIPF5 | 1 | 1 | -0,026063619 | 0,573770089 | 1 |
| ST8SIA3 | 2 | 2 | 0,201192196 | 0,38106619 | 1 |
| SH3RF1 | 2 | 2 | 0,094485755 | 0,227204397 | 1 |
| SNAP25 | 2 | 2 | 0 | 1 | 1 |
| PSAP | 1 | 1 | 0 | 1 | 1 |
| CACNA1A;CACNA1I | 1 | 1 | 0 | 1 | 1 |
| TMEM9 | 1 | 1 | 0 | 1 | 1 |
| SLC1A1 | 1 | 1 | 0 | 1 | 1 |
| CA10 | 1 | 1 | 0 | 1 | 1 |
| KIF5A;KIF5B | 2 | 2 | -0,387703393 | 0,869684146 | 1 |
| ERGIC3 | 1 | 1 | 0 | 1 | 1 |
| PHAX | 1 | 1 | 0 | 1 | 1 |
| ARPC3 | 1 | 1 | 0,000580294 | 0,95200492 | 1 |
| KCNQ3 | 1 | 1 | -1,93471E-19 | 1 | 1 |
| B2M | 1 | 1 | 0 | 1 | 1 |
| CSPG5 | 1 | 1 | -0,050673862 | 0,376500937 | 1 |
| RBBP4;RBBP7 | 3 | 2 | 0 | 1 | 1 |
| FOXRED1 | 2 | 2 | 0 | 1 | 1 |
| DENND6A | 1 | 1 | 0 | 1 | 1 |
| LRP11 | 1 | 1 | 0 | 1 | 1 |
| PARK7 | 1 | 1 | 0 | 1 | 1 |
| ZFP442 | 1 | 1 | 0 | 1 | 1 |
| COL6A5 | 1 | 1 | 0 | 1 | 1 |
| CUTA | 1 | 1 | 0 | 1 | 1 |

| | | | | | |
|---|---|---|---|---|---|
| FAM151B | 1 | 1 | 0 | 1 | 1 |
| KCTD6 | 3 | 3 | 0,051030554 | 0,335464795 | 1 |
| CNBP | 1 | 1 | 0 | 1 | 1 |
| SCN3A | 1 | 1 | 0 | 1 | 1 |
| PABPN1 | 1 | 1 | 0 | 1 | 1 |
| SLC12A6;GM21985 | 2 | 1 | 0,204606342 | 0,182880677 | 1 |
| FBXL4 | 2 | 1 | 0 | 1 | 1 |
| ADAMTS10 | 1 | 1 | 0 | 1 | 1 |
| SFI1 | 1 | 1 | 0 | 1 | 1 |
| TMEM145 | 1 | 1 | 0 | 1 | 1 |
| KALRN | 1 | 1 | 0 | 1 | 1 |
| DBNDD2 | 1 | 1 | -0,087607641 | 0,312782583 | 1 |
| SATB2 | 1 | 1 | -0,129449316 | 0,217456417 | 1 |
| TRANK1 | 1 | 1 | 0 | 1 | 1 |
| RASSF2;RASSF4 | 1 | 1 | -4,95157E-20 | 1 | 1 |
| ABCA3 | 1 | 1 | -2,33351E-19 | 1 | 1 |
| CHRM3 | 1 | 1 | -0,007876453 | 0,814944654 | 1 |
| ERMAP | 1 | 1 | 0 | 1 | 1 |
| ACSM3 | 1 | 1 | -0,235482662 | 0,136657365 | 1 |
| NHSL1 | 1 | 1 | 0 | 1 | 1 |
| SLC7A1 | 1 | 1 | 0 | 1 | 1 |
| FRMPD4 | 1 | 1 | 0 | 1 | 1 |
| DNM1L | 1 | 1 | 0 | 1 | 1 |
| CAPG | 2 | 2 | 3,15467E-14 | 0,999999629 | 1 |
| CYSTM1 | 1 | 1 | 0 | 1 | 1 |
| IRF3 | 2 | 2 | -4,90616E-19 | 0,999999999 | 1 |
| NEURL1 | 1 | 1 | 0 | 1 | 1 |
| SNCG | 1 | 1 | 0 | 1 | 1 |
| ARID2 | 1 | 1 | 0 | 1 | 1 |
| IMPA1 | 1 | 1 | 0 | 1 | 1 |
| SMG6 | 2 | 2 | 0,171444454 | 0,17257799 | 1 |
| IGF1R | 2 | 1 | 4,99271E-19 | 0,999999999 | 1 |
| ACKR1 | 1 | 1 | 0 | 1 | 1 |

| | | | | | |
|---|---|---|---|---|---|
| PRELID3B | 1 | 1 | 0 | 1 | 1 |
| NUP214 | 1 | 1 | 0 | 1 | 1 |
| APOC3 | 1 | 1 | 0 | 1 | 1 |
| CDA | 1 | 1 | 0 | 1 | 1 |
| SLC4A4 | 1 | 1 | 0 | 1 | 1 |
| IQSEC2 | 1 | 1 | 0 | 1 | 1 |
| CD47 | 1 | 1 | 0 | 1 | 1 |
| PTN | 1 | 1 | -0,036560197 | 0,564854554 | 1 |
| CPLANE1 | 1 | 1 | 0 | 1 | 1 |
| PIP5K1C;PIP5K1A | 1 | 1 | 0 | 1 | 1 |
| LYNX1 | 1 | 1 | -0,16472357 | 0,30815869 | 1 |
| FBXO42 | 1 | 1 | -0,076331853 | 0,259031859 | 1 |
| KIAA0319 | 1 | 1 | 0,08518841 | 0,401806789 | 1 |
| KIFC2 | 1 | 1 | 0 | 1 | 1 |
| DPH6 | 2 | 2 | 0 | 1 | 1 |
| CXADR | 2 | 2 | 1,28899E-16 | 0,999999973 | 1 |
| ZHX2 | 1 | 1 | 0 | 1 | 1 |
| RELCH | 1 | 1 | 0 | 1 | 1 |
| ATP1A2;ATP12A | 1 | 1 | 0 | 1 | 1 |
| DECR2 | 1 | 1 | 0 | 1 | 1 |
| YIPF3 | 1 | 1 | -0,17206223 | 0,251411991 | 1 |
| TOR1AIP2 | 1 | 1 | 0 | 1 | 1 |
| SUMO2;SUMO3 | 1 | 1 | 0 | 1 | 1 |
| BCKDHB | 1 | 1 | 0 | 1 | 1 |
| CYP4F3;CYP4F14 | 1 | 1 | 0 | 1 | 1 |
| MIB1 | 1 | 1 | 0,017774145 | 0,667988249 | 1 |
| GM1043 | 1 | 1 | 0 | 1 | 1 |
| CSNK1D | 1 | 1 | 0 | 1 | 1 |
| PDE4DIP | 1 | 1 | -0,015958556 | 0,73626504 | 1 |
| RGN | 1 | 1 | 0 | 1 | 1 |
| ATP8A1 | 1 | 1 | 0 | 1 | 1 |
| LGALS3 | 5 | 1 | 0 | 1 | 1 |
| PTPRF;PTPRS;PTPR | 2 | 2 | -1,29423E-16 | 0,999999978 | 1 |

| | | | | | |
|---|---|---|---|---|---|
| DCLK1 | 2 | 2 | -1,69533E-17 | 0,999999995 | 1 |
| TMEM181A | 1 | 1 | 0 | 1 | 1 |
| KLK10 | 1 | 1 | 0 | 1 | 1 |
| TG | 1 | 1 | 0 | 1 | 1 |
| DPY30 | 1 | 1 | 0 | 1 | 1 |
| STAC | 1 | 1 | 0 | 1 | 1 |
| DIP2C | 1 | 1 | 0 | 1 | 1 |
| MRPS31 | 3 | 1 | 0 | 1 | 1 |
| MAST2 | 1 | 1 | 0 | 1 | 1 |
| BCL11A | 1 | 1 | 0 | 1 | 1 |
| MECP2 | 1 | 1 | 0 | 1 | 1 |
| SMAD2 | 1 | 1 | 0 | 1 | 1 |
| GSC2;GSC | 1 | 1 | 0 | 1 | 1 |
| MGST1 | 1 | 1 | 0 | 1 | 1 |
| ZFP958 | 1 | 1 | 0 | 1 | 1 |
| DNAH8 | 1 | 1 | 0 | 1 | 1 |
| NFKB1 | 1 | 1 | 0 | 1 | 1 |
| APPBP2 | 1 | 1 | 0 | 1 | 1 |
| HPDL | 1 | 1 | 0 | 1 | 1 |
| FUNDC1 | 1 | 1 | 0 | 1 | 1 |
| GABBR1 | 1 | 1 | 4,59107E-17 | 0,999999984 | 1 |
| CFL1;CFL2 | 3 | 3 | -7,39121E-17 | 0,999999985 | 1 |
| DEPDC5 | 1 | 1 | 0,006893229 | 0,835362456 | 1 |
| MRPL10 | 2 | 1 | 0 | 1 | 1 |
| MRPL30 | 1 | 1 | 0 | 1 | 1 |
| GRB10 | 1 | 1 | 0 | 1 | 1 |
| FBXW15 | 1 | 1 | 0 | 1 | 1 |
| MAP4K2 | 1 | 1 | 0,004874122 | 0,840579747 | 1 |
| GAS2L1 | 1 | 1 | 0,024423028 | 0,73390973 | 1 |
| NBEAL2 | 2 | 2 | -3,18632E-16 | 0,99999998 | 1 |
| RANBP10 | 2 | 2 | 0 | 1 | 1 |
| SAAL1 | 1 | 1 | -0,110481397 | 0,27581122 | 1 |
| APCS | 1 | 1 | 0,362354499 | 0,112764848 | 1 |

| | | | | | |
|---|---|---|---|---|---|
| TMEM14C | 1 | 1 | 0 | 1 | 1 |
| SLC47A2 | 1 | 1 | 0 | 1 | 1 |
| STK11IP | 1 | 1 | 0 | 1 | 1 |
| GM14692 | 1 | 1 | 0 | 1 | 1 |
| BTBD11 | 1 | 1 | 0 | 1 | 1 |
| QRICH1 | 1 | 1 | 0 | 1 | 1 |
| FEM1AA | 1 | 1 | 0 | 1 | 1 |
| LIMS1 | 1 | 1 | 0 | 1 | 1 |
| TDRD3 | 1 | 1 | 0 | 1 | 1 |
| AGBL4 | 1 | 1 | 0 | 1 | 1 |
| TBC1D9 | 1 | 1 | 0 | 1 | 1 |
| LDHC;LDHA;LDHB | 1 | 1 | 0 | 1 | 1 |
| MATR3 | 1 | 1 | 0 | 1 | 1 |
| PCDHGA8 | 1 | 1 | 0 | 1 | 1 |
| RAB19 | 1 | 1 | -0,114367352 | 0,378637762 | 1 |
| SCYL3 | 1 | 1 | 0 | 1 | 1 |
| FER1L5 | 1 | 1 | 0,147672233 | 0,37415652 | 1 |
| SIPA1L3;SIPA1L1 | 1 | 1 | 0 | 1 | 1 |
| OVCA2 | 1 | 1 | 0 | 1 | 1 |
| ABCB9 | 2 | 1 | 0 | 1 | 1 |
| TGM3 | 7 | 1 | 0 | 1 | 1 |
| CRIPT | 1 | 1 | 0 | 1 | 1 |
| FBXW11 | 1 | 1 | 0 | 1 | 1 |
| CFAP46 | 1 | 1 | 0 | 1 | 1 |
| AGFG2;AGFG1 | 1 | 1 | -1,68366E-19 | 1 | 1 |
| AUP1 | 1 | 1 | 0,096136184 | 0,301538871 | 1 |
| SH2D1A | 1 | 1 | 0 | 1 | 1 |
| MYH11;MYH10;MY | 1 | 1 | 0 | 1 | 1 |
| CRAT | 1 | 1 | -0,067069139 | 0,404126536 | 1 |
| LDAH | 1 | 1 | -0,060728087 | 0,372036191 | 1 |
| sp\|Q8BHN7\|CL029 | 1 | 1 | 0 | 1 | 1 |
| TRMT1 | 1 | 1 | 0 | 1 | 1 |
| MPV17 | 1 | 1 | 0 | 1 | 1 |

| | | | | | |
|---|---|---|---|---|---|
| MTHFD1L;MTHFD1 | 1 | 1 | 0 | 1 | 1 |
| CCDC85A | 1 | 1 | 0 | 1 | 1 |
| EPHA3;EPHA4;EPH/ | 1 | 1 | 0 | 1 | 1 |
| LIN7B;LIN7A | 2 | 2 | -9,90316E-19 | 0,999999999 | 1 |
| PCDHGA9;PCDHGA | 1 | 1 | 0,382890326 | 0,117504758 | 1 |
| TMEM94 | 1 | 1 | 0,103919659 | 0,666765771 | 1 |
| GLT6D1 | 1 | 1 | 0 | 1 | 1 |
| GSTT2 | 1 | 1 | -9,67549E-19 | 0,999999999 | 1 |
| GIMAP8 | 1 | 1 | 0 | 1 | 1 |
| VMN2R3 | 1 | 1 | 0,210925362 | 0,302109829 | 1 |
| KCNA5 | 1 | 1 | 0 | 1 | 1 |
| CPVL | 1 | 1 | -0,29959793 | 0,121315327 | 1 |
| ACO1;IREB2 | 1 | 1 | 0 | 1 | 1 |
| SKOR2 | 1 | 1 | 0 | 1 | 1 |
| IL23R | 1 | 1 | 0,207973709 | 0,231376693 | 1 |
| ADCY10 | 1 | 1 | 0 | 1 | 1 |
| ZBTB48 | 1 | 1 | 0 | 1 | 1 |
| SFT2D2 | 1 | 1 | 0 | 1 | 1 |
| FBXO33 | 1 | 1 | 0 | 1 | 1 |
| MTG1 | 1 | 1 | 0 | 1 | 1 |
| ODR4 | 1 | 1 | 0 | 1 | 1 |
| RPL35 | 1 | 1 | 0 | 1 | 1 |
| GRAMD1A | 1 | 1 | 0,109661248 | 0,238607799 | 1 |
| RABL3 | 1 | 1 | 0 | 1 | 1 |
| ENPP5 | 1 | 1 | 0 | 1 | 1 |
| DEGS1 | 2 | 2 | 0,131683893 | 0,238166224 | 1 |
| CA11 | 1 | 1 | 0 | 1 | 1 |
| OLFR1458 | 1 | 1 | 0,010018733 | 0,788815445 | 1 |
| PPM1J;PPM1H | 2 | 1 | 0 | 1 | 1 |
| ANK3;ANK1 | 1 | 1 | 0 | 1 | 1 |
| RTL8C | 1 | 1 | 0 | 1 | 1 |
| GATM | 4 | 3 | 3,69945E-18 | 0,999999997 | 1 |
| SNRPE | 1 | 1 | 0 | 1 | 1 |

| | | | | | |
|---|---|---|---|---|---|
| SHC1 | 1 | 1 | 0,089781722 | 0,686021847 | 1 |
| CYP1A2 | 1 | 1 | 0 | 1 | 1 |
| TRPC3 | 1 | 1 | 0 | 1 | 1 |
| NEURL4 | 1 | 1 | -3,33778E-22 | 1 | 1 |
| RTP3 | 1 | 1 | 0 | 1 | 1 |
| ENO3 | 3 | 3 | 0,020192379 | 0,608268439 | 1 |
| IGHG1 | 3 | 1 | 0 | 1 | 1 |
| TMEM223 | 1 | 1 | 0 | 1 | 1 |
| PLEKHA6 | 1 | 1 | -0,167009447 | 0,184332402 | 1 |
| PRX | 1 | 1 | -0,18452447 | 0,348814332 | 1 |
| OSTC | 1 | 1 | 0,19026107 | 0,140938328 | 1 |
| RPP25L | 1 | 1 | 0 | 1 | 1 |
| ANKZF1 | 1 | 1 | 0 | 1 | 1 |
| IGHG2C | 3 | 1 | 0 | 1 | 1 |
| METTL1 | 2 | 2 | 4,34946E-18 | 0,999999996 | 1 |
| ABCA17 | 1 | 1 | 0 | 1 | 1 |
| IDS | 2 | 2 | 0 | 1 | 1 |
| GABBR1 | 1 | 1 | 0 | 1 | 1 |
| TYMP | 1 | 1 | -0,314528498 | 0,180677821 | 1 |
| ICMT | 1 | 1 | 0 | 1 | 1 |
| WDFY2 | 1 | 1 | 0 | 1 | 1 |
| TICAM2 | 1 | 1 | 0 | 1 | 1 |
| CCHCR1 | 1 | 1 | 0 | 1 | 1 |
| FMNL2;FMNL1 | 1 | 1 | 0 | 1 | 1 |
| ABHD14B | 1 | 1 | 0 | 1 | 1 |
| ORAI2 | 1 | 1 | 0 | 1 | 1 |
| SLC25A53 | 1 | 1 | -3,10147E-20 | 1 | 1 |
| TEX19.2 | 1 | 1 | 0 | 1 | 1 |
| SGSH | 1 | 1 | 0 | 1 | 1 |
| MRPS26 | 2 | 1 | 0 | 1 | 1 |
| NDUFA1 | 2 | 2 | 0,092284949 | 0,181596886 | 1 |
| MRPL50 | 1 | 1 | -3,63911E-18 | 0,999999997 | 1 |
| ADAMTS20 | 1 | 1 | 0 | 1 | 1 |

| | | | | | |
|---|---|---|---|---|---|
| PSMB9 | 1 | 1 | 0 | 1 | 1 |
| TNS3;TNS2 | 1 | 1 | 0 | 1 | 1 |
| LARP7 | 1 | 1 | 1,09294E-19 | 1 | 1 |
| ADORA1 | 1 | 1 | 0 | 1 | 1 |
| ACP1 | 1 | 1 | 0 | 1 | 1 |
| TOMM34 | 1 | 1 | 0 | 1 | 1 |
| TAX1BP3 | 1 | 1 | 0 | 1 | 1 |
| STXBP1 | 3 | 3 | 4,26657E-17 | 0,999999992 | 1 |
| TGOLN2 | 1 | 1 | 0 | 1 | 1 |
| ROBO4 | 1 | 1 | 0 | 1 | 1 |
| SEZ6 | 1 | 1 | 0 | 1 | 1 |
| DNAJB4;DNAJB1 | 1 | 1 | 0 | 1 | 1 |
| ARMC8 | 1 | 1 | 0 | 1 | 1 |
| UBAP1 | 1 | 1 | 0,121439115 | 0,240478928 | 1 |
| SUPT4H1A;SUPT4H | 1 | 1 | 0 | 1 | 1 |
| PNKD | 1 | 1 | 0 | 1 | 1 |
| ABCG2;ABCG3 | 1 | 1 | 0 | 1 | 1 |
| ALDH1L2 | 1 | 1 | 0,03062262 | 0,780017002 | 1 |
| SCN3A;SCN2A;SCN! | 2 | 2 | 0,035778297 | 0,699334585 | 1 |
| sp\|Q8VE95\|CH082 | 1 | 1 | 0 | 1 | 1 |
| ATG2A | 1 | 1 | -0,079976221 | 0,362923021 | 1 |
| CYP26B1 | 1 | 1 | 0 | 1 | 1 |
| HNRNPH1 | 1 | 1 | 0 | 1 | 1 |
| CUL9;CUL7 | 1 | 1 | 0 | 1 | 1 |
| UBXN4 | 1 | 1 | 0 | 1 | 1 |
| SAXO2 | 1 | 1 | 0 | 1 | 1 |
| COPS7A | 1 | 1 | 0 | 1 | 1 |
| MAVS | 1 | 1 | 0 | 1 | 1 |
| SBF1 | 1 | 1 | 0 | 1 | 1 |
| JAM2 | 1 | 1 | 0 | 1 | 1 |
| PDF | 1 | 1 | -0,386873621 | 0,152398819 | 1 |
| FGF13 | 1 | 1 | 0 | 1 | 1 |
| RAF1 | 2 | 2 | 3,57332E-16 | 0,999999963 | 1 |

| | | | | | |
|---|---|---|---|---|---|
| DISC1 | 1 | 1 | 0 | 1 | 1 |
| MEP1B | 1 | 1 | 0 | 1 | 1 |
| ATP13A3 | 1 | 1 | 0 | 1 | 1 |
| PANK1 | 1 | 1 | 0 | 1 | 1 |
| KCNA1;KCNA3;KCN | 2 | 2 | 3,34945E-14 | 0,999999733 | 1 |
| PTRHD1 | 1 | 1 | 0 | 1 | 1 |
| NDUFS1 | 1 | 1 | 0 | 1 | 1 |
| SNAPIN | 1 | 1 | 0 | 1 | 1 |
| ACOT8 | 1 | 1 | 0 | 1 | 1 |
| PLAT | 1 | 1 | 0,05537994 | 0,460463773 | 1 |
| NOLC1 | 1 | 1 | -3,06628E-14 | 0,999999629 | 1 |
| IGHG;sp\|P01864\|G | 3 | 2 | 0 | 1 | 1 |
| PKP3 | 1 | 1 | 0 | 1 | 1 |
| ARFRP1 | 1 | 1 | -0,09786151 | 0,47329849 | 1 |
| SLC27A6 | 1 | 1 | 0 | 1 | 1 |
| SSU72 | 1 | 1 | 0 | 1 | 1 |
| UNC13B | 1 | 1 | 0 | 1 | 1 |
| EPHB3;EPHB4 | 1 | 1 | 0,223209088 | 0,128614135 | 1 |
| FRMPD1 | 1 | 1 | 0 | 1 | 1 |
| ACAD10 | 2 | 1 | 0 | 1 | 1 |
| RGMA | 1 | 1 | 0 | 1 | 1 |
| TOR1B | 1 | 1 | -0,173654569 | 0,198809462 | 1 |
| PPIP5K1;PPIP5K2 | 1 | 1 | 0,180922628 | 0,137962353 | 1 |
| LRP2 | 1 | 1 | 0,041732944 | 0,570521113 | 1 |
| MSL3 | 1 | 1 | 0,040698935 | 0,570832312 | 1 |
| SLC17A8;SLC17A6 | 1 | 1 | 0 | 1 | 1 |
| STXBP5L;STXBP5 | 1 | 1 | 0 | 1 | 1 |
| KATNAL2 | 1 | 1 | 0,022522648 | 0,670669847 | 1 |
| CSF2RB2;CSF2RB | 1 | 1 | 0,170096734 | 0,156453686 | 1 |
| EPHB2;EPHA7 | 1 | 1 | 0 | 1 | 1 |
| ARL6IP1 | 1 | 1 | -1,72066E-19 | 1 | 1 |
| HSPA12B | 1 | 1 | 1,77783E-23 | 1 | 1 |
| RAB3D;RAB3A | 1 | 1 | 0 | 1 | 1 |

| | | | | | |
|---|---|---|---|---|---|
| IGHV9-4 | 1 | 1 | 0 | 1 | 1 |
| CD44 | 3 | 2 | 3,07438E-17 | 0,99999999 | 1 |
| PXN | 1 | 1 | 0,079953547 | 0,358461328 | 1 |
| LRRN1 | 1 | 1 | 0 | 1 | 1 |
| VMN2R70 | 1 | 1 | -0,093010749 | 0,314574965 | 1 |
| RBFOX1 | 1 | 1 | -2,19247E-15 | 0,999999897 | 1 |
| HSD17B7 | 1 | 1 | 0 | 1 | 1 |
| UGT1A6 | 1 | 1 | 0 | 1 | 1 |
| METTL5 | 1 | 1 | 0 | 1 | 1 |
| GOSR2 | 1 | 1 | 0 | 1 | 1 |
| KRT1;KRT6A;KRT2;I | 2 | 2 | 3,78787E-17 | 0,999999996 | 1 |
| KRT86;KRT81 | 1 | 1 | 0 | 1 | 1 |
| ORM1 | 2 | 2 | 4,14432E-16 | 0,999999982 | 1 |
| HMGB2 | 3 | 1 | 8,87864E-20 | 1 | 1 |
| ABHD17B | 2 | 2 | 6,38863E-19 | 0,999999999 | 1 |
| CCDC158 | 1 | 1 | 0 | 1 | 1 |
| MYO5B | 1 | 1 | 0,120009226 | 0,177692641 | 1 |
| B4GALT4 | 1 | 1 | 0 | 1 | 1 |
| MBNL2 | 1 | 1 | -1,80599E-20 | 1 | 1 |
| GNA12 | 1 | 1 | 0 | 1 | 1 |
| SCN1A;SCN3A;SCN: | 1 | 1 | 0 | 1 | 1 |
| PDCD6IP | 1 | 1 | 0 | 1 | 1 |
| CHST14 | 1 | 1 | -0,20204599 | 0,15468307 | 1 |
| DOP1B | 1 | 1 | 0 | 1 | 1 |
| HDAC1;HDAC2 | 1 | 1 | 0 | 1 | 1 |
| HDAC2 | 1 | 1 | -0,11891376 | 0,204147663 | 1 |
| OTUD7B | 1 | 1 | 0 | 1 | 1 |
| KCTD21 | 1 | 1 | 0,028249061 | 0,56026592 | 1 |
| GOLGA7;GOLGA7B | 1 | 1 | 0 | 1 | 1 |
| CEP85L | 1 | 1 | -0,006007877 | 0,802932659 | 1 |
| SPRED2 | 1 | 1 | 0 | 1 | 1 |
| EHD4;EHD3 | 2 | 2 | 4,08087E-16 | 0,99999997 | 1 |
| TBC1D25 | 1 | 1 | 0 | 1 | 1 |

| | | | | | |
|---|---|---|---|---|---|
| RLN1 | 1 | 1 | 0 | 1 | 1 |
| TUBB4B;TUBB5;TU | 1 | 1 | 0 | 1 | 1 |
| ATAD2 | 1 | 1 | 0 | 1 | 1 |
| TIMM23 | 2 | 1 | 0 | 1 | 1 |
| TAT | 1 | 1 | 0 | 1 | 1 |
| CACNB4;CACNB1 | 1 | 1 | 0 | 1 | 1 |
| XPR1 | 1 | 1 | 0 | 1 | 1 |
| HDGFL2;HDGFL3 | 1 | 1 | 0 | 1 | 1 |
| ARHGEF9;SPATA13 | 1 | 1 | 0 | 1 | 1 |
| TMCO5A | 1 | 1 | 0 | 1 | 1 |
| UNC119B;UNC119 | 1 | 1 | 0 | 1 | 1 |
| CLDN10 | 1 | 1 | 0 | 1 | 1 |
| CRK | 1 | 1 | -1,49223E-17 | 0,999999991 | 1 |
| RIMS1;RIMS2 | 1 | 1 | 0 | 1 | 1 |
| ACAP2 | 1 | 1 | 0 | 1 | 1 |
| ADGRB3;ADGRB2 | 1 | 1 | 0 | 1 | 1 |
| SMIM26 | 1 | 1 | 0 | 1 | 1 |
| POLR2A | 1 | 1 | -0,043632948 | 0,485908864 | 1 |
| MSTN | 1 | 1 | 0,016605654 | 0,761996349 | 1 |
| RALGPS2 | 1 | 1 | 0 | 1 | 1 |
| MDGA2 | 1 | 1 | -0,02752266 | 0,662450571 | 1 |
| SLC6A21 | 1 | 1 | -0,087032716 | 0,520817559 | 1 |
| SERPINA3I | 1 | 1 | 0 | 1 | 1 |
| WDR82 | 1 | 1 | 0 | 1 | 1 |
| HDAC9 | 1 | 1 | 0 | 1 | 1 |
| RAPGEF3;RAPGEF4 | 1 | 1 | 0 | 1 | 1 |
| UPP1 | 1 | 1 | 0,139727631 | 0,293787515 | 1 |
| sp\|E0CYV9\|CD054_ | 1 | 1 | 0 | 1 | 1 |
| PTPRA | 1 | 1 | -1,27662E-17 | 0,999999992 | 1 |
| SNRNP25 | 1 | 1 | 0 | 1 | 1 |
| GPX3 | 1 | 1 | 0 | 1 | 1 |
| ETFBKMT | 1 | 1 | -0,080451662 | 0,321689447 | 1 |
| TAF6L | 1 | 1 | 0 | 1 | 1 |

| | | |
|---|---|---|
| FAM193B | 1 | 1 |
| PCDHB1 | 1 | 1 |
| TXLNG | 1 | 1 |
| PGAP4 | 1 | 1 |
| PDIA5 | 1 | 1 |
| H2-Q10;H2-D1 | 1 | 1 |
| ECEL1 | 1 | 1 |
| COA6 | 1 | 1 |
| NACC1 | 1 | 1 |
| SH2D6 | 1 | 1 |
| GLYCTK | 1 | 1 |
| THEMIS3 | 1 | 1 |
| ADAMTSL2 | 1 | 1 |
| TOGARAM1 | 1 | 1 |
| SELENOI | 1 | 1 |
| sp\|Q8K1L6\|CP074_ | 1 | 1 |
| EID2 | 1 | 1 |
| sp\|Q9CPS8\|SMAKA | 1 | 1 |
| DNAH10 | 1 | 1 |
| MYO1F | 1 | 1 |
| EPHA5 | 1 | 1 |
| EPPK1 | 13 | 1 |
| sp\|P04945\|KV6AB_ | 1 | 1 |
| IGKV6-17 | 2 | 1 |
| sp\|P01644\|KV5AB_ | 4 | 1 |
| DNAAF2 | 1 | 1 |
| ZDHHC20 | 1 | 1 |
| PRSS44 | 1 | 1 |
| ARL4A | 1 | 1 |
| PTK6 | 1 | 1 |
| ANXA8 | 5 | 1 |
| WDCP | 1 | 1 |
| TLR5 | 1 | 1 |

| | | |
|---|---|---|
| AFF4 | 1 | 1 |
| ATRX | 1 | 1 |
| GLI1 | 1 | 1 |
| H1-4 | 1 | 1 |
| TPGS1 | 1 | 1 |
| ANO8 | 1 | 1 |
| CRK | 1 | 1 |
| OCIAD1 | 1 | 1 |
| H3-5;H3C11;H3C15 | 2 | 1 |
| ERBB4 | 1 | 1 |
| MMP23 | 1 | 0 |
| BCKDHB | 1 | 0 |
| BPTF | 1 | 0 |
| TMBIM1 | 1 | 0 |
| MRPS5 | 3 | 0 |
| CEMIP2 | 1 | 0 |
| LRRC74B | 1 | 0 |
| SLC41A3 | 1 | 0 |
| CPEB1 | 1 | 0 |
| H1-3 | 3 | 0 |
| SHANK3 | 1 | 0 |
| DBR1 | 1 | 0 |
| PSIP1 | 1 | 0 |
| 1700029H14RIK | 1 | 0 |
| KIAA0895L | 1 | 0 |
| MYBPC3 | 1 | 0 |
| CALM4 | 2 | 0 |
| HEG1 | 1 | 0 |
| SIX6 | 1 | 0 |
| ULK4 | 1 | 0 |
| IFI202 | 5 | 0 |
| TRO | 1 | 0 |
| MMRN1 | 1 | 0 |

| | | |
|---|---|---|
| KRT15 | 19 | 0 |
| KRT6A;KRT76 | 1 | 0 |
| KRT1;KRT6A;KRT2;I | 1 | 0 |
| CLU | 1 | 0 |
| ZFX | 1 | 0 |
| HRNR | 5 | 0 |
| ZCCHC18 | 1 | 0 |
| TMEM237 | 1 | 0 |
| TIMM21 | 2 | 0 |
| PAN3 | 1 | 0 |
| ADCY6 | 1 | 0 |
| CD177 | 2 | 0 |
| COL1A2 | 7 | 0 |
| NRXN1 | 1 | 0 |
| SLC8A3 | 1 | 0 |
| KCTD12 | 1 | 0 |
| PPP1R15A | 1 | 0 |
| YIPF2 | 1 | 0 |
| MME | 1 | 0 |
| PTCD2 | 1 | 0 |
| TUBB1 | 1 | 0 |
| DSC1 | 1 | 0 |
| KCNJ15 | 1 | 0 |
| HERC3 | 1 | 0 |
| CACFD1 | 1 | 0 |
| RBM26 | 1 | 0 |
| PRSS58 | 1 | 0 |
| MTFMT | 1 | 0 |
| GTPBP8 | 1 | 0 |
| TENM4;TENM2 | 1 | 0 |
| TULP4 | 1 | 0 |
| AMN | 1 | 0 |
| RHOT2 | 1 | 0 |

| | | |
|---|---|---|
| YIPF4 | 1 | 0 |
| UCP3 | 1 | 0 |
| PGK2 | 1 | 0 |
| DNAJC28 | 1 | 0 |
| ALDH3B2 | 1 | 0 |
| LIPT2 | 1 | 0 |
| ARMC9 | 1 | 0 |
| OLFR1141 | 1 | 0 |
| SLC17A8 | 1 | 0 |
| SMC1B | 1 | 0 |
| TOMM6 | 1 | 0 |
| CAPS2 | 1 | 0 |
| SLIT3 | 1 | 0 |
| PVR | 1 | 0 |
| OAS2 | 1 | 0 |
| TTC16 | 1 | 0 |
| SPTLC2 | 1 | 0 |
| PRPH | 1 | 0 |
| RPL36 | 1 | 0 |
| SMIM4 | 1 | 0 |
| S100A14 | 1 | 0 |
| SPECC1L | 1 | 0 |
| S100A9 | 5 | 0 |
| KRT80 | 4 | 0 |
| TM9SF1 | 1 | 0 |
| CHD6 | 1 | 0 |
| PTCD1 | 1 | 0 |
| GRP | 1 | 0 |
| STOX2 | 1 | 0 |
| ARAP1 | 1 | 0 |
| ARG2 | 1 | 0 |
| CDH4 | 1 | 0 |
| HBS1L | 1 | 0 |

| | | |
|---|---|---|
| CLCN3;CLCN4;CLCN | 1 | 0 |
| NCOR1 | 1 | 0 |
| UQCC3 | 1 | 0 |
| TEX9 | 1 | 0 |
| LTF | 14 | 0 |
| ZBTB4 | 1 | 0 |
| MACF1;DST | 1 | 0 |
| KAT6A | 1 | 0 |
| PLA2G4C | 1 | 0 |
| MUL1 | 2 | 0 |
| APC2 | 1 | 0 |
| ATP8A1 | 1 | 0 |
| ACAD11 | 1 | 0 |
| EFEMP2 | 1 | 0 |
| FDX1 | 1 | 0 |
| SGSM2 | 1 | 0 |
| ITGAE | 1 | 0 |
| NRDE2 | 1 | 0 |
| FAXC | 1 | 0 |
| IFIH1 | 1 | 0 |
| MEP1A | 1 | 0 |
| FLNC | 1 | 0 |
| TIFAB | 1 | 0 |
| OLFR576 | 1 | 0 |
| NGP | 6 | |
| SERPINB5 | 11 | |
| IGKV4-63 | 1 | |
| MSH2 | 1 | |
| DSG1B | 1 | |
| ITPR1 | 1 | |
| CHIL3 | 7 | |
| WTAP | 1 | |
| NEU2 | 1 | |

| | |
|---|---|
| INTS7 | 1 |
| URAH | 2 |
| GM32742 | 1 |
| POLQ | 1 |
| LCN6 | 1 |
| DOLK | 1 |
| MPO | 12 |
| FLG2 | 3 |
| NCCRP1 | 1 |
| ABCA6 | 1 |
| KRT13 | 2 |
| CFAP47 | 1 |
| TRIM66 | 1 |
| NKTR | 1 |
| FBXL5 | 1 |
| CCHCR1 | 1 |
| TMEM119 | 1 |
| RUFY2 | 1 |
| TACR3 | 1 |
| S100A8 | 1 |
| KRT84 | 3 |
| A1BG | 2 |
| HELQ | 1 |
| SAA1 | 1 |
| ENDOU | 7 |
| KRT33B | 1 |
| sp\|P01654\|KV3A1_ | 1 |
| H2-D1;H2-T23 | 1 |
| IGKV4-57 | 1 |
| LYZ2 | 1 |
| MPZ | 1 |
| POF1B | 6 |
| ORM2 | 1 |

| | |
|---|---|
| CFAP57 | 1 |
| KRT6B | 5 |
| AHNAK2 | 1 |
| KRT36 | 1 |
| KRT35;KRT34 | 1 |
| PKHD1L1 | 1 |
| FLG | 3 |
| LCN2 | 3 |
| KRT6A;KRT79 | 1 |
| IGHV1-18;IGHV1-2[illegible] | 1 |
| GM5478;GM5414 | 1 |
| SERPINB3B | 1 |
| LMNB1 | 3 |
| ZFP60 | 1 |
| H1-5 | 2 |
| KRT24 | 6 |
| TREX2 | 3 |
| IGHV9-3 | 3 |
| OBP1B | 1 |
| CAMP | 2 |
| LGALS7 | 6 |
| SERPINB2 | 4 |
| TRIM29 | 4 |
| DSG1A | 1 |
| SBSN | 2 |
| IGKV1-135;IGKV1-1 | 1 |
| COL1A1 | 6 |
| MMP9 | 2 |
| FLG | 2 |
| CDH22 | 1 |
| SERPINB6D | 1 |
| IGHV1-20;IGHV1-3[illegible] | 1 |
| GM1553 | 1 |

| | |
|---|---|
| IGHG2B | 2 |
| SUN2 | 1 |
| ITGB2L | 1 |
| PRTN3 | 1 |
| CYBB | 1 |
| IGKV17-127;IGKV1[illegible] | 1 |
| IGHV5-9-1 | 1 |
| IGHV5-12 | 2 |
| KRT1;KRT2;KRT77 | 1 |
| ELANE | 1 |
| DSC3 | 1 |
| PPL | 2 |
| KRT10 | 1 |
| KRT19;KRT15 | 1 |
| OLFM4 | 1 |
| CSTDC5;CSTDC6 | 1 |
| OBP1A | 1 |
| PIP | 2 |
| LY6D | 1 |
| DSG3 | 1 |
| KRT13;KRT15 | 1 |
| CDSN | 1 |
