## Supplementary material for "SFRP1 upregulation causes hippocampal synaptic dysfunction and memory impairment": Suppl Table 2

| gene_symbols_or_id | unique_peptides | peptides_used_for_dea_contrast: Control_Syn vs SFRP1-TG_Syn | foldchange.log2_msqrob_contrast: Control_Syn vs SFRP1-TG_Syn | pvalue_msqrob_contrast: Control_Syn vs SFRP1-TG_Syn | qvalue_msqrob_contrast: Control_Syn vs SFRP1-TG_Syn |
|---|---|---|---|---|---|
| GPC1 | 11 | 11 | 0,152786143 | 2,50337E-07 | 0,001810186 |
| ANPEP | 3 | 2 | 1,036356018 | 6,58809E-07 | 0,002381922 |
| KRT14 | 18 | 4 | 3,164606308 | 3,54893E-06 | 0,005769103 |
| TJP2 | 18 | 18 | -0,169088495 | 4,16777E-06 | 0,005769103 |
| ANKS1B | 15 | 15 | 0,141736087 | 4,78698E-06 | 0,005769103 |
| WIPF2 | 7 | 7 | -0,173425562 | 4,56567E-06 | 0,005769103 |
| WDR7 | 66 | 66 | 0,100375279 | 7,99724E-06 | 0,008261149 |
| IRS2 | 2 | 2 | -0,459283281 | 1,37751E-05 | 0,012450958 |
| HOMER3 | 17 | 17 | 0,196654375 | 2,25319E-05 | 0,018103134 |
| SYT12 | 13 | 13 | 0,135300488 | 2,89903E-05 | 0,01905716 |
| RAB39B | 11 | 11 | 0,144366806 | 2,73857E-05 | 0,01905716 |
| EFR3B | 24 | 24 | 0,085336927 | 4,43398E-05 | 0,022901504 |
| EPHA4 | 26 | 25 | 0,0890485 | 4,04442E-05 | 0,022901504 |
| TPRKB | 3 | 1 | 0,943223848 | 4,36203E-05 | 0,022901504 |
| ROGDI | 17 | 17 | 0,096886292 | 5,70832E-05 | 0,025798045 |
| TIAM2 | 2 | 1 | -0,447529831 | 5,38125E-05 | 0,025798045 |
| ARHGAP39 | 27 | 27 | 0,084990671 | 7,94929E-05 | 0,028150672 |
| PIP5K1C | 25 | 25 | 0,1052462 | 9,81263E-05 | 0,028150672 |
| NRP1 | 16 | 16 | 0,155108333 | 9,93171E-05 | 0,028150672 |
| ATP6V0A1 | 43 | 43 | 0,068593823 | 9,78176E-05 | 0,028150672 |
| HSPB1 | 4 | 1 | 2,618497127 | 8,74277E-05 | 0,028150672 |
| CA4 | 7 | 7 | 0,180842521 | 7,14437E-05 | 0,028150672 |
| PPP2R5C | 9 | 9 | 0,140217261 | 7,14373E-05 | 0,028150672 |
| ACTR10 | 15 | 15 | 0,105230943 | 0,000105112 | 0,028150672 |
| PBXIP1 | 3 | 3 | -0,237400525 | 7,5682E-05 | 0,028150672 |
| SLC44A2 | 3 | 3 | 0,327262783 | 0,000102834 | 0,028150672 |
| GARIN2 | 1 | 1 | -1,387520798 | 8,76585E-05 | 0,028150672 |
| SCAI | 22 | 22 | 0,092115657 | 0,000123391 | 0,031865757 |

| | | | | | |
|---|---|---|---|---|---|
| PPP4C | 2 | 2 | 0,31067145 | 0,000144582 | 0,03592955 |
| CNNM4 | 3 | 3 | 0,240099871 | 0,000149065 | 0,03592955 |
| sp|Q8C3W1|CA198_ | 9 | 9 | -0,14069992 | 0,000175581 | 0,040955733 |
| CADM3 | 18 | 18 | 0,163400154 | 0,000199984 | 0,045075834 |
| RAB2A;RAB2B | 9 | 9 | 0,194747959 | 0,000205712 | 0,045075834 |
| SDC3 | 2 | 2 | -0,348606093 | 0,000212088 | 0,045106201 |
| NLN | 13 | 13 | -0,202098548 | 0,000221901 | 0,045844792 |
| AP2A2 | 54 | 54 | 0,088091473 | 0,000239435 | 0,048093138 |
| GM382 | 1 | 1 | -1,683590491 | 0,000247882 | 0,048444108 |
| NUCB1 | 11 | 11 | -0,218720815 | 0,000255612 | 0,048640238 |
| CACNA2D3 | 31 | 31 | 0,089605173 | 0,000273439 | 0,050698451 |
| **CPNE4** | **18** | **18** | **0,17838917** | **0,000281079** | **0,050811977** |
| NRXN3 | 26 | 26 | 0,088866938 | 0,00030548 | 0,053876233 |
| CNKSR2 | 35 | 35 | 0,065176218 | 0,000349335 | 0,056436789 |
| PFKL | 37 | 37 | 0,181248902 | 0,000335854 | 0,056436789 |
| RAPGEF4 | 24 | 24 | 0,088161727 | 0,000351218 | 0,056436789 |
| NUP205 | 2 | 2 | 0,336911953 | 0,000348298 | 0,056436789 |
| FAM81A | 9 | 9 | -0,140943577 | 0,000360265 | 0,056632105 |
| NRP2 | 10 | 10 | 0,128254186 | 0,000383082 | 0,058937602 |
| COX18 | 1 | 1 | -1,159185641 | 0,000442972 | 0,066731961 |
| VNN1 | 1 | 1 | -1,256474775 | 0,00045355 | 0,066931089 |
| FAAH | 14 | 14 | 0,13337254 | 0,000468336 | 0,067730745 |
| TJP1 | 30 | 30 | -0,098534433 | 0,000485275 | 0,067955931 |
| NPTX1 | 14 | 14 | 0,136175788 | 0,000496103 | 0,067955931 |
| PRL8A8 | 1 | 1 | -0,958549589 | 0,000498087 | 0,067955931 |
| ARL8B | 7 | 7 | 0,207457886 | 0,000532828 | 0,071349568 |
| CSNK1D | 1 | 1 | -0,446212335 | 0,000545457 | 0,071712689 |
| D7H11ORF16 | 1 | 1 | -0,927861261 | 0,000577003 | 0,074505534 |
| SPOCK3 | 1 | 1 | -0,530199294 | 0,000653144 | 0,082857617 |
| SH3GL3 | 12 | 12 | 0,096185821 | 0,000674015 | 0,084031093 |
| CYFIP2 | 70 | 68 | 0,093083692 | 0,000697174 | 0,08544518 |
| RAB8A | 7 | 7 | 0,1560117 | 0,000717916 | 0,086520798 |
| CACNA1A | 13 | 13 | 0,146275732 | 0,000742788 | 0,087165379 |

| | | | | | |
|---|---|---|---|---|---|
| L1CAM | 4 | 4 | 0,203101597 | 0,000747373 | 0,087165379 |
| EPHA6 | 5 | 5 | 0,165780933 | 0,00078292 | 0,089861775 |
| SLC17A7 | 14 | 14 | 0,160087982 | 0,000843769 | 0,095332724 |
| PRICKLE2 | 16 | 16 | 0,083086025 | 0,000933679 | 0,10386825 |
| CCT3 | 40 | 39 | 0,072125167 | 0,000999493 | 0,109505048 |
| SFRP1 | 2 | 1 | 0,972223664 | 0,001021743 | 0,11027196 |
| USO1 | 36 | 30 | 0,070558965 | 0,00106739 | 0,111859422 |
| CPD | 11 | 10 | -0,113899215 | 0,001059749 | 0,111859422 |
| MACF1 | 72 | 72 | -0,077693315 | 0,001101457 | 0,1137805 |
| FAM120C | 10 | 10 | -0,119613645 | 0,001195736 | 0,121779774 |
| MYO16 | 1 | 1 | -0,478305078 | 0,001245569 | 0,125093229 |
| GMPPB | 8 | 8 | 0,132553739 | 0,001276988 | 0,12584759 |
| CAMK2D | 14 | 13 | -0,114534883 | 0,001287888 | 0,12584759 |
| LRRTM1 | 9 | 9 | 0,130560572 | 0,001321766 | 0,127435907 |
| SYNPR | 8 | 8 | 0,19473612 | 0,001369287 | 0,130280424 |
| EHD3;EHD1 | 11 | 11 | 0,116650814 | 0,001393751 | 0,1308859 |
| FIBCD1 | 4 | 4 | 0,162780194 | 0,001422214 | 0,131699363 |
| AKAP7 | 3 | 3 | -0,315977513 | 0,001457053 | 0,131699363 |
| RPS27 | 1 | 1 | 0,713531752 | 0,001455782 | 0,131699363 |
| MRPL24 | 2 | 2 | -0,265376724 | 0,001541396 | 0,137602864 |
| DMD | 6 | 6 | -0,156650656 | 0,001695094 | 0,149478388 |
| PPP2R5A | 14 | 14 | 0,072144954 | 0,001840002 | 0,160301891 |
| NEURL4 | 1 | 1 | -0,786272323 | 0,001881817 | 0,161993047 |
| ABCF1 | 8 | 6 | 0,208605077 | 0,001960127 | 0,166399587 |
| EIF3F | 15 | 13 | 0,102750421 | 0,00197903 | 0,166399587 |
| RAB9A | 6 | 6 | 0,135285022 | 0,002022471 | 0,168097586 |
| KCNMA1 | 26 | 26 | 0,087147538 | 0,00215052 | 0,176709235 |
| FAM184A | 1 | 1 | 0,420865663 | 0,002213701 | 0,177858573 |
| KRT1;KRT6A;KRT2;KI | 1 | 1 | 2,331009639 | 0,002207806 | 0,177858573 |
| USP12 | 1 | 1 | 0,394282283 | 0,002265382 | 0,180010725 |
| CYP2D22 | 6 | 6 | -0,20791185 | 0,002314692 | 0,180659647 |
| DDX3X | 4 | 4 | 0,165824413 | 0,002323516 | 0,180659647 |
| EPB41L2 | 37 | 37 | -0,092000829 | 0,002435095 | 0,182080177 |

| | | | | | |
|---|---|---|---|---|---|
| SYT3 | 16 | 16 | 0,092788007 | 0,002442508 | 0,182080177 |
| TUBAL3 | 5 | 4 | 0,223457965 | 0,002406533 | 0,182080177 |
| TMEM181A | 1 | 1 | -0,473476379 | 0,002400066 | 0,182080177 |
| SMPD3 | 11 | 11 | 0,097089781 | 0,002507183 | 0,184994279 |
| EEF1G | 20 | 19 | 0,090706554 | 0,002700624 | 0,188658369 |
| SV2B | 20 | 20 | 0,110183731 | 0,002810267 | 0,188658369 |
| LIPT1 | 1 | 1 | -0,331579487 | 0,002697081 | 0,188658369 |
| KTN1 | 23 | 21 | -0,112593674 | 0,002817744 | 0,188658369 |
| HBB-B1 | 13 | 11 | 0,379414977 | 0,002761041 | 0,188658369 |
| RAB4B | 9 | 9 | 0,084644395 | 0,002780318 | 0,188658369 |
| SLC6A7 | 13 | 13 | 0,10928876 | 0,002777453 | 0,188658369 |
| RAB1A | 14 | 14 | 0,118494853 | 0,002585663 | 0,188658369 |
| ARL4C | 1 | 1 | 0,336826257 | 0,002733418 | 0,188658369 |
| CPEB3;CPEB2 | 1 | 1 | 0,446782988 | 0,002714846 | 0,188658369 |
| CYFIP1 | 28 | 28 | 0,092407014 | 0,002960309 | 0,196385239 |
| FGFR1;FGFR3 | 1 | 1 | -0,264500952 | 0,00303084 | 0,198368799 |
| PIP5K1B | 2 | 2 | -0,334451483 | 0,00306713 | 0,198368799 |
| RAF1 | 2 | 2 | 0,244228364 | 0,003072508 | 0,198368799 |
| ACSL3 | 14 | 14 | -0,08340412 | 0,00324006 | 0,207335144 |
| ATP6V0D1 | 19 | 19 | 0,069802898 | 0,003334886 | 0,211531239 |
| ZER1 | 10 | 9 | 0,153088731 | 0,003369627 | 0,211876314 |
| RIN1 | 1 | 1 | 0,497634476 | 0,003653388 | 0,223878373 |
| CCDC136 | 5 | 5 | -0,135530977 | 0,003622553 | 0,223878373 |
| RBX1 | 1 | 1 | 0,31008394 | 0,003623299 | 0,223878373 |
| sp\|Q9CYI0\|NJMU_M | 1 | 1 | -0,569415845 | 0,003775261 | 0,227649783 |
| ADI1 | 3 | 3 | -0,18590936 | 0,003777897 | 0,227649783 |
| FLAD1 | 11 | 10 | -0,093845797 | 0,003818353 | 0,228186031 |
| DGKB | 16 | 16 | 0,156342415 | 0,003891647 | 0,230659822 |
| PSEN1 | 1 | 1 | 0,371702768 | 0,003949268 | 0,232172026 |
| ARHGEF2 | 12 | 12 | -0,120862501 | 0,004001026 | 0,233158793 |
| VKORC1L1 | 1 | 1 | 0,48714163 | 0,004030542 | 0,233158793 |
| TRIM28 | 15 | 7 | 0,375646213 | 0,004075716 | 0,233900821 |
| SERGEF | 1 | 1 | -0,533623846 | 0,004147002 | 0,23611789 |

| | | | | | |
|---|---|---|---|---|---|
| SENP6 | 1 | 1 | -0,242214218 | 0,004265209 | 0,239737855 |
| GPATCH11 | 2 | 1 | -1,068240542 | 0,004276889 | 0,239737855 |
| SLC16A1 | 5 | 5 | 0,152985427 | 0,004345519 | 0,240453822 |
| RAB10 | 11 | 11 | 0,096625948 | 0,004366773 | 0,240453822 |
| ITGAM;GM49368 | 1 | 1 | 0,469931076 | 0,004389421 | 0,240453822 |
| DNM1 | 55 | 55 | 0,090190789 | 0,004580991 | 0,249061236 |
| SCN3A;SCN2A;SCN9A | 4 | 4 | 0,152768205 | 0,004621592 | 0,249393543 |
| CCDC88A | 12 | 12 | -0,140456505 | 0,004684581 | 0,250826707 |
| LAMTOR2 | 3 | 3 | 0,178880134 | 0,004717526 | 0,250826707 |
| PDIA5 | 1 | 1 | -0,787282203 | 0,004780372 | 0,251638551 |
| AP4S1 | 1 | 1 | 0,474170786 | 0,004802395 | 0,251638551 |
| WDR20 | 4 | 4 | -0,15356437 | 0,004987499 | 0,259457579 |
| ST3GAL5 | 1 | 1 | 0,310849295 | 0,005162466 | 0,26664138 |
| TPRG1L | 10 | 10 | 0,104675408 | 0,005261332 | 0,269820533 |
| CPE | 22 | 22 | 0,070971218 | 0,005360273 | 0,270809994 |
| SF3B3 | 11 | 7 | 0,227359495 | 0,005377748 | 0,270809994 |
| S100A1 | 1 | 1 | 1,283458299 | 0,00539298 | 0,270809994 |
| ACOX1 | 15 | 15 | 0,095982048 | 0,005497695 | 0,272286524 |
| CETN2 | 2 | 2 | 0,296325394 | 0,005484107 | 0,272286524 |
| RIMS1 | 12 | 12 | 0,070145153 | 0,005594294 | 0,273326609 |
| HPCA | 7 | 7 | 0,221212158 | 0,005577924 | 0,273326609 |
| PRPSAP1 | 11 | 10 | 0,114705983 | 0,005762047 | 0,277769073 |
| AP2B1 | 35 | 34 | 0,06484113 | 0,005752212 | 0,277769073 |
| AGAP3 | 11 | 11 | 0,089904902 | 0,005945828 | 0,28392721 |
| THA1 | 3 | 3 | -0,201611162 | 0,005968322 | 0,28392721 |
| DPY19L4 | 1 | 1 | -0,294329887 | 0,006342177 | 0,290254938 |
| PITPNM2 | 21 | 21 | -0,084505291 | 0,006311186 | 0,290254938 |
| RAB27A | 2 | 2 | 0,142098726 | 0,006337061 | 0,290254938 |
| JAKMIP3 | 6 | 6 | -0,113125782 | 0,006299192 | 0,290254938 |
| RPL21 | 6 | 5 | 0,276957573 | 0,006243214 | 0,290254938 |
| TTC16 | 1 | 1 | -0,659020808 | 0,006312433 | 0,290254938 |
| NEDD4L | 18 | 18 | 0,064130971 | 0,006456385 | 0,293623417 |
| RAB14 | 20 | 20 | 0,085765403 | 0,00656608 | 0,296299798 |

| | | | | | |
|---|---|---|---|---|---|
| EPHA3;EPHA6 | 1 | 1 | 0,44432434 | 0,006597188 | 0,296299798 |
| FAM163B | 1 | 1 | -0,443801649 | 0,006755772 | 0,301549312 |
| VPS25 | 5 | 5 | 0,141001079 | 0,0068013 | 0,301719024 |
| HPCAL1;NCALD | 6 | 6 | 0,090336798 | 0,007007192 | 0,30895736 |
| ARAP2 | 1 | 1 | -0,319047195 | 0,007106048 | 0,310665524 |
| EIPR1 | 7 | 7 | 0,146236288 | 0,00713186 | 0,310665524 |
| SPHKAP | 15 | 15 | -0,104336203 | 0,007282873 | 0,313466982 |
| SERINC5 | 1 | 1 | -0,277190582 | 0,00726428 | 0,313466982 |
| HPCAL4 | 19 | 17 | 0,179659966 | 0,007421863 | 0,317559112 |
| GABRA5 | 5 | 5 | 0,156976784 | 0,007948927 | 0,334178454 |
| NFXL1 | 1 | 1 | 0,546361252 | 0,007861798 | 0,334178454 |
| RALGAPA1 | 14 | 14 | -0,116073209 | 0,007931243 | 0,334178454 |
| SLC39A12 | 6 | 6 | 0,120247504 | 0,008285662 | 0,346021708 |
| ARHGEF9;SPATA13;A | 1 | 1 | 0,412698847 | 0,008326342 | 0,346021708 |
| NHSL2 | 10 | 10 | -0,087432395 | 0,008569531 | 0,352081117 |
| SNRPF | 1 | 1 | 1,17862729 | 0,00855588 | 0,352081117 |
| AK9 | 1 | 1 | -0,176096271 | 0,008673107 | 0,354323377 |
| HIP1 | 7 | 7 | -0,139524069 | 0,008886011 | 0,358907622 |
| CRBN | 5 | 5 | 0,210597088 | 0,008982674 | 0,358907622 |
| MRPS14 | 1 | 1 | -0,249340431 | 0,008865227 | 0,358907622 |
| VMN2R11 | 1 | 1 | 0,593121304 | 0,008983858 | 0,358907622 |
| PRKAG1 | 6 | 6 | 0,132208902 | 0,009635984 | 0,377241427 |
| RGS14 | 20 | 20 | 0,113407451 | 0,00969083 | 0,377241427 |
| AKR1C13 | 1 | 1 | 0,19275623 | 0,009703624 | 0,377241427 |
| CDK12;CDK13 | 2 | 2 | -0,439633376 | 0,009506434 | 0,377241427 |
| LMNTD1 | 1 | 1 | -0,389187647 | 0,009649027 | 0,377241427 |
| CCT4 | 38 | 38 | 0,080630015 | 0,009769723 | 0,37778002 |
| CSNK2A2 | 18 | 18 | 0,067683896 | 0,009941932 | 0,380370936 |
| AASS | 3 | 3 | -0,342654269 | 0,009940182 | 0,380370936 |
| RPS13 | 10 | 9 | 0,231947507 | 0,010003996 | 0,380731017 |
| KIAA1549 | 17 | 17 | 0,083274744 | 0,010076102 | 0,381467491 |
| VPS26B | 7 | 7 | 0,099435642 | 0,010198024 | 0,381603453 |
| NPTN | 2 | 2 | 0,233848665 | 0,010238013 | 0,381603453 |

| | | | | | |
|---|---|---|---|---|---|
| TRMU | 4 | 4 | -0,179828971 | 0,010144029 | 0,381603453 |
| CDV3 | 6 | 6 | -0,134908008 | 0,010665333 | 0,395492431 |
| PCDHGC5 | 8 | 8 | 0,093702136 | 0,010791281 | 0,398121186 |
| FLOT1 | 29 | 29 | 0,063103007 | 0,01106354 | 0,404042726 |
| SPTAN1 | 2 | 2 | -0,228934895 | 0,011031538 | 0,404042726 |
| SERPINF2 | 4 | 1 | 1,24304504 | 0,011223954 | 0,406314333 |
| MAPK10;MAPK9 | 2 | 2 | 0,190350635 | 0,011238123 | 0,406314333 |
| DBH | 1 | 1 | 0,273935557 | 0,011324798 | 0,407411009 |
| SLC25A18 | 11 | 11 | -0,233778029 | 0,01147406 | 0,410737276 |
| PACSIN1 | 43 | 42 | 0,070423853 | 0,011532487 | 0,410795158 |
| GPD1 | 17 | 17 | 0,079436957 | 0,011658185 | 0,413236944 |
| TRAPPC2 | 2 | 2 | 0,163835973 | 0,01180065 | 0,416246332 |
| DST | 41 | 41 | -0,088625323 | 0,011933793 | 0,418899289 |
| SHISA4 | 1 | 1 | 0,350995923 | 0,012006187 | 0,419404549 |
| MTERF2 | 3 | 3 | -0,195755044 | 0,012068958 | 0,419570378 |
| SYP | 9 | 9 | 0,129589062 | 0,012230836 | 0,423163506 |
| ARHGEF12 | 19 | 19 | -0,057619531 | 0,01230138 | 0,423577529 |
| AP3S1 | 7 | 7 | 0,145850074 | 0,012493431 | 0,428151641 |
| C9ORF72 | 3 | 3 | 0,20643437 | 0,01276277 | 0,431250407 |
| NAPG | 24 | 24 | 0,055848395 | 0,012708642 | 0,431250407 |
| TRIM3 | 15 | 15 | 0,058330968 | 0,012697229 | 0,431250407 |
| SQOR | 8 | 8 | -0,188907551 | 0,012989429 | 0,436867732 |
| KLC2 | 15 | 15 | -0,060585741 | 0,013481734 | 0,442835672 |
| RPS23 | 5 | 5 | 0,24553799 | 0,013278135 | 0,442835672 |
| KIT | 17 | 17 | 0,10278335 | 0,013434881 | 0,442835672 |
| OSBP2 | 4 | 4 | 0,13143984 | 0,013534322 | 0,442835672 |
| YWHAB | 10 | 9 | 0,084926418 | 0,013416734 | 0,442835672 |
| ATP2B3 | 5 | 5 | 0,099792457 | 0,013532518 | 0,442835672 |
| RCC2 | 1 | 1 | 0,743837294 | 0,013752378 | 0,445934733 |
| USP46;USP12 | 3 | 3 | 0,137196535 | 0,013741631 | 0,445934733 |
| CPOX | 9 | 9 | -0,129413513 | 0,013852689 | 0,447182107 |
| LSM14A | 3 | 1 | 0,331538182 | 0,014173415 | 0,45550207 |
| GRIA2 | 36 | 36 | 0,093509203 | 0,014380237 | 0,456067941 |

| | | | | | |
|---|---|---|---|---|---|
| VCAN | 1 | 1 | -0,201322518 | 0,014375542 | 0,456067941 |
| HBB-BS | 4 | 2 | 0,356056749 | 0,014275013 | 0,456067941 |
| RHOT1 | 1 | 1 | -0,197575088 | 0,014531007 | 0,458837167 |
| RAP2A | 7 | 7 | 0,109899792 | 0,014662862 | 0,458992008 |
| SMN1 | 1 | 1 | -0,522090053 | 0,0146181 | 0,458992008 |
| SEPTIN8 | 30 | 30 | 0,072579256 | 0,014818256 | 0,461856945 |
| TNIK;MAP4K4;MINK | 9 | 9 | 0,083540468 | 0,015024644 | 0,463443257 |
| ABHD3 | 4 | 3 | -0,217324237 | 0,015061425 | 0,463443257 |
| TOMM34 | 1 | 1 | 0,26375213 | 0,015046721 | 0,463443257 |
| SPECC1 | 15 | 15 | -0,079881517 | 0,01514975 | 0,464185767 |
| HAPLN4 | 10 | 10 | 0,095982319 | 0,015595268 | 0,471799093 |
| FARSB | 21 | 16 | 0,165261009 | 0,015464823 | 0,471799093 |
| SPATA45 | 1 | 1 | -0,177750449 | 0,015629133 | 0,471799093 |
| FAM117B | 2 | 2 | -0,264579042 | 0,015686863 | 0,471799093 |
| ZFX | 1 | 1 | 0,585405863 | 0,015724462 | 0,471799093 |
| RPL18A | 6 | 6 | 0,433484938 | 0,015801099 | 0,472139436 |
| CLVS1 | 12 | 11 | 0,077418536 | 0,016135353 | 0,47428755 |
| 2900026A02RIK | 11 | 10 | 0,104921681 | 0,016078285 | 0,47428755 |
| SPIRE1 | 5 | 5 | 0,106258329 | 0,01609257 | 0,47428755 |
| ITGA6 | 5 | 5 | 0,279985094 | 0,016057477 | 0,47428755 |
| TLCD4 | 2 | 2 | 0,149924276 | 0,016413871 | 0,479236378 |
| SPART | 7 | 7 | -0,116337869 | 0,016436264 | 0,479236378 |
| GPX1 | 13 | 13 | 0,110014168 | 0,016523183 | 0,479686182 |
| SRP68 | 7 | 7 | 0,154714359 | 0,01668062 | 0,479686182 |
| ELOC | 9 | 9 | 0,094059174 | 0,01671704 | 0,479686182 |
| MRPS31 | 3 | 3 | -0,215122449 | 0,016664282 | 0,479686182 |
| DNAJC13 | 29 | 29 | 0,051245189 | 0,016940695 | 0,481344594 |
| MPST | 15 | 15 | -0,177523095 | 0,017343538 | 0,481344594 |
| EPHB2 | 10 | 10 | 0,13953185 | 0,016843118 | 0,481344594 |
| NTNG1 | 5 | 5 | -0,094996355 | 0,01716822 | 0,481344594 |
| RAP2B;RAP2A;RAP2( | 8 | 8 | 0,100144132 | 0,017096365 | 0,481344594 |
| UBE2V2 | 7 | 6 | 0,119331968 | 0,017380274 | 0,481344594 |
| TXNDC12 | 2 | 2 | -0,130615706 | 0,017071304 | 0,481344594 |

| | | | | | |
|---|---|---|---|---|---|
| DCTN6 | 6 | 6 | 0,114776943 | 0,017440504 | 0,481344594 |
| DR1 | 1 | 1 | 0,677139106 | 0,017396248 | 0,481344594 |
| ANK3;ANK1 | 1 | 1 | -0,426601752 | 0,017219364 | 0,481344594 |
| DNAJC6 | 24 | 24 | 0,042019134 | 0,017536837 | 0,481355569 |
| RILPL1 | 13 | 13 | -0,118226426 | 0,017640607 | 0,481355569 |
| NLGN4L | 1 | 1 | -0,306105458 | 0,017633442 | 0,481355569 |
| SCD2;SCD1;SCD4;SCI | 2 | 2 | 0,340205575 | 0,017780813 | 0,481547025 |
| CA11 | 1 | 1 | 0,198697729 | 0,017759247 | 0,481547025 |
| BORCS6 | 2 | 2 | -0,311437444 | 0,017949382 | 0,482498079 |
| NDRG3 | 14 | 12 | 0,086239318 | 0,017892595 | 0,482498079 |
| UNC5CL | 1 | 1 | -0,267023137 | 0,018130041 | 0,483757663 |
| CAPS2 | 1 | 1 | -0,725037888 | 0,018070146 | 0,483757663 |
| PLXNA1 | 40 | 40 | 0,084121851 | 0,018444039 | 0,488720433 |
| ADGRE5 | 1 | 1 | 0,525329894 | 0,018451207 | 0,488720433 |
| CMPK2 | 14 | 14 | -0,090889386 | 0,01857888 | 0,49030613 |
| VPS41 | 6 | 6 | -0,109521107 | 0,018676468 | 0,491089224 |
| ZNRF2 | 1 | 1 | 0,403337704 | 0,018800369 | 0,492556031 |
| ATP8A2 | 8 | 8 | 0,090161683 | 0,018966054 | 0,495103012 |
| PPM1K | 1 | 1 | -0,283396066 | 0,019190482 | 0,497370532 |
| BRCC3 | 6 | 6 | 0,152967127 | 0,019129147 | 0,497370532 |
| TRAPPC8 | 12 | 12 | 0,068909829 | 0,019301568 | 0,498463006 |
| SBF1 | 58 | 58 | 0,037581229 | 0,019820936 | 0,500681558 |
| SUGT1 | 17 | 17 | 0,07218434 | 0,020062926 | 0,500681558 |
| MAPK1 | 35 | 34 | 0,104052356 | 0,019889054 | 0,500681558 |
| DIRAS2 | 12 | 12 | 0,159972377 | 0,02010955 | 0,500681558 |
| LANCL2 | 18 | 18 | 0,096067261 | 0,020218367 | 0,500681558 |
| CSNK1G1;CSNK1G3 | 1 | 1 | 0,184884885 | 0,020013018 | 0,500681558 |
| RAB35 | 14 | 14 | 0,061230043 | 0,020197441 | 0,500681558 |
| GRAMD1B | 5 | 5 | 0,105649141 | 0,019985153 | 0,500681558 |
| BRINP1 | 15 | 15 | 0,068574596 | 0,01950883 | 0,500681558 |
| QKI | 4 | 4 | -0,109908448 | 0,019608522 | 0,500681558 |
| MAPK8IP1 | 2 | 2 | -0,189867882 | 0,020163373 | 0,500681558 |
| INPP4A | 9 | 9 | 0,077940687 | 0,02018842 | 0,500681558 |

| | | | | | |
|---|---|---|---|---|---|
| ZNF148 | 1 | 1 | -0,241123526 | 0,020291365 | 0,500774271 |
| RAB15 | 9 | 9 | 0,105811799 | 0,020540387 | 0,503822151 |
| DCLK2 | 18 | 18 | 0,064343399 | 0,020582785 | 0,503822151 |
| PRKAB2 | 7 | 7 | 0,120025935 | 0,020623891 | 0,503822151 |
| AMN | 1 | 1 | 0,720635484 | 0,020896359 | 0,508759507 |
| GSK3A | 9 | 9 | 0,080761345 | 0,021150127 | 0,510287805 |
| PTP4A1 | 3 | 3 | 0,148943628 | 0,021134834 | 0,510287805 |
| CELSR2 | 3 | 3 | -0,153503359 | 0,02117084 | 0,510287805 |
| RPL22 | 4 | 4 | 0,174402908 | 0,021537449 | 0,517399647 |
| ABCF3 | 7 | 7 | -0,098859162 | 0,021644277 | 0,51820835 |
| SHISA9 | 2 | 2 | -0,145150194 | 0,021714442 | 0,51820835 |
| CAPZA2 | 14 | 14 | 0,093063193 | 0,021923407 | 0,521245807 |
| EIF5A | 13 | 13 | 0,152880982 | 0,02198589 | 0,521245807 |
| ITM2C | 5 | 5 | 0,11120589 | 0,022345081 | 0,528027384 |
| FAM241B | 2 | 2 | -0,108269631 | 0,022417979 | 0,528027384 |
| PFDN4 | 4 | 3 | 0,138651408 | 0,022561004 | 0,528601883 |
| SYNE1 | 43 | 43 | -0,08228481 | 0,022807881 | 0,528601883 |
| ALDH1A1 | 19 | 18 | 0,087048538 | 0,022754891 | 0,528601883 |
| RPS6 | 12 | 9 | 0,096022012 | 0,022804866 | 0,528601883 |
| FRYL | 3 | 3 | -0,174332052 | 0,022682 | 0,528601883 |
| GNS | 5 | 5 | -0,208959475 | 0,022882352 | 0,528633513 |
| ACTR3B | 15 | 15 | 0,070009737 | 0,023027713 | 0,528947493 |
| TSHZ1 | 1 | 1 | -0,316186478 | 0,023042243 | 0,528947493 |
| PTBP2 | 2 | 2 | 0,284421554 | 0,023149348 | 0,529724481 |
| ACAP2 | 13 | 13 | 0,090559572 | 0,023392638 | 0,532051697 |
| LRFN1 | 12 | 12 | 0,09088923 | 0,023398208 | 0,532051697 |
| SRCIN1 | 10 | 10 | -0,078948665 | 0,02379371 | 0,535515392 |
| RPL10A | 14 | 12 | 0,1401705 | 0,023722355 | 0,535515392 |
| PLXNB1 | 28 | 28 | -0,041651085 | 0,023994881 | 0,535515392 |
| RAB1A;RAB1B | 5 | 5 | 0,081249631 | 0,023925106 | 0,535515392 |
| SLC1A7 | 1 | 1 | -0,827725346 | 0,023958798 | 0,535515392 |
| IGHG;sp\|P01864\|GC | 3 | 2 | 1,821950923 | 0,023977417 | 0,535515392 |
| ADCY9 | 31 | 31 | 0,07588402 | 0,024380576 | 0,538823907 |

| | | | | | |
|---|---|---|---|---|---|
| RPL10A | 1 | 1 | 0,230833608 | 0,024292066 | 0,538823907 |
| LIAS | 4 | 4 | -0,125952597 | 0,024363875 | 0,538823907 |
| TPM1 | 1 | 1 | -0,1999848 | 0,02444119 | 0,538823907 |
| GALE | 8 | 7 | 0,158464314 | 0,024621118 | 0,541140742 |
| GSTM2;GSTM4 | 3 | 2 | -0,193311331 | 0,024697493 | 0,541174471 |
| SLC8A1 | 30 | 29 | 0,069668329 | 0,024997717 | 0,544900106 |
| RNF123 | 6 | 6 | 0,094642385 | 0,025018232 | 0,544900106 |
| RAB2B | 3 | 3 | 0,143439202 | 0,025201202 | 0,547236918 |
| SPTLC2 | 1 | 1 | -0,314639063 | 0,025348404 | 0,548785357 |
| HGS | 18 | 18 | -0,058189066 | 0,025589792 | 0,549511758 |
| MADD | 47 | 47 | 0,044919654 | 0,025609938 | 0,549511758 |
| AP2S1 | 11 | 11 | 0,105508695 | 0,025477371 | 0,549511758 |
| SPTBN4 | 44 | 44 | -0,108368672 | 0,026224156 | 0,549802579 |
| LRRC8D | 12 | 12 | 0,083205547 | 0,026359116 | 0,549802579 |
| ACTR1A | 10 | 10 | 0,075084799 | 0,025953256 | 0,549802579 |
| HSPA8 | 39 | 38 | 0,061626985 | 0,025933048 | 0,549802579 |
| INF2 | 8 | 7 | -0,144713612 | 0,026298243 | 0,549802579 |
| CAMK2B | 19 | 19 | 0,058305853 | 0,026459867 | 0,549802579 |
| SLC9A7 | 7 | 7 | 0,069622478 | 0,026284878 | 0,549802579 |
| TMOD1 | 9 | 9 | -0,108242299 | 0,026452804 | 0,549802579 |
| YBX3 | 1 | 1 | 0,546997516 | 0,026031633 | 0,549802579 |
| PRKAR1A | 17 | 17 | -0,051021669 | 0,026252562 | 0,549802579 |
| LSM2 | 3 | 1 | 0,359076399 | 0,025992963 | 0,549802579 |
| GABRB3 | 5 | 5 | 0,090928483 | 0,02670382 | 0,553281725 |
| NSUN2 | 12 | 3 | 0,578939869 | 0,026857471 | 0,553294501 |
| GSPT1 | 11 | 9 | 0,095028153 | 0,026805971 | 0,553294501 |
| MINAR1 | 1 | 1 | -0,653297302 | 0,027156261 | 0,556381409 |
| NOVA2 | 4 | 3 | 0,362862612 | 0,0271612 | 0,556381409 |
| STMN1;STMN2 | 4 | 4 | -0,072973124 | 0,027384132 | 0,556385036 |
| HBA-A1 | 9 | 9 | 0,302029391 | 0,027379113 | 0,556385036 |
| ATP6V0A2;ATP6V0A | 1 | 1 | 0,182189816 | 0,02739221 | 0,556385036 |
| COPG1 | 12 | 12 | -0,094109832 | 0,027745298 | 0,560126527 |
| KIAA1109 | 15 | 15 | -0,067274204 | 0,027808799 | 0,560126527 |

| | | | | | |
|---|---|---|---|---|---|
| SPOCK2 | 2 | 2 | 0,158447763 | 0,027696829 | 0,560126527 |
| 4930438A08RIK | 1 | 1 | 0,432863987 | 0,027957432 | 0,561556095 |
| METTL8 | 1 | 1 | -0,442432611 | 0,028145203 | 0,56376168 |
| DDX1 | 15 | 14 | 0,062937758 | 0,02835187 | 0,566332511 |
| SLC6A17 | 20 | 20 | 0,081084773 | 0,028636217 | 0,569069589 |
| DNPH1 | 1 | 1 | 0,675172419 | 0,028646291 | 0,569069589 |
| WFS1 | 19 | 19 | -0,113557285 | 0,029143457 | 0,569159258 |
| GATD1 | 5 | 5 | 0,134647737 | 0,029260887 | 0,569159258 |
| CUL3 | 38 | 37 | 0,045993313 | 0,029170931 | 0,569159258 |
| YWHAH | 18 | 18 | 0,078266469 | 0,029359204 | 0,569159258 |
| PARD3B | 1 | 1 | -0,191457925 | 0,029295145 | 0,569159258 |
| RERG | 1 | 1 | 0,787165911 | 0,02893155 | 0,569159258 |
| TMEM70 | 2 | 2 | -0,188441412 | 0,02882011 | 0,569159258 |
| PRELID3A | 2 | 2 | -0,194483847 | 0,028814864 | 0,569159258 |
| RHOT2 | 1 | 1 | -0,222140967 | 0,029170671 | 0,569159258 |
| SMG5 | 1 | 1 | -0,403584399 | 0,029475051 | 0,569877257 |
| RAB7A | 21 | 21 | 0,072429073 | 0,029671501 | 0,571953089 |
| PAN2 | 1 | 1 | 0,168621711 | 0,029740611 | 0,571953089 |
| CORO1A | 22 | 22 | 0,056919484 | 0,029970893 | 0,572509421 |
| ADD3 | 25 | 24 | -0,051583767 | 0,030026284 | 0,572509421 |
| CD47 | 5 | 5 | 0,127189551 | 0,030022784 | 0,572509421 |
| SLC25A17 | 1 | 1 | 0,156886509 | 0,030086237 | 0,572509421 |
| TUBA8 | 10 | 9 | 0,141164878 | 0,030709928 | 0,577600086 |
| CSDC2 | 2 | 2 | -0,200616991 | 0,030728977 | 0,577600086 |
| MIGA1 | 2 | 2 | -0,197524384 | 0,030753151 | 0,577600086 |
| SNRPD2 | 3 | 3 | 0,393793506 | 0,030742163 | 0,577600086 |
| CNBP | 1 | 1 | 0,336143322 | 0,030469683 | 0,577600086 |
| GPR37L1 | 3 | 3 | 0,171313739 | 0,03094086 | 0,579620095 |
| ARF6 | 8 | 8 | 0,199892056 | 0,031191069 | 0,582797466 |
| GOLM2 | 3 | 3 | -0,154762348 | 0,031530826 | 0,587627328 |
| SCN2B | 9 | 9 | 0,078749342 | 0,031716203 | 0,589562636 |
| MAP1B | 109 | 105 | -0,080765705 | 0,03179874 | 0,58958125 |
| SGTB | 6 | 5 | -0,061654225 | 0,032143848 | 0,592701868 |

| | | | | | |
|---|---|---|---|---|---|
| DLGAP1 | 9 | 9 | 0,07418487 | 0,03218351 | 0,592701868 |
| POMT1 | 1 | 1 | -0,243659737 | 0,032212949 | 0,592701868 |
| AP1S1 | 7 | 7 | 0,098569336 | 0,032459109 | 0,595715271 |
| CYTH3 | 3 | 3 | 0,126913517 | 0,03281015 | 0,600633399 |
| GIT1 | 35 | 34 | -0,038612023 | 0,033128197 | 0,602349101 |
| PLPPR2 | 8 | 8 | 0,100954747 | 0,033135108 | 0,602349101 |
| ARL8A;ARL8B | 6 | 6 | 0,19818433 | 0,033153774 | 0,602349101 |
| NCKAP1 | 69 | 68 | 0,078904246 | 0,033484085 | 0,604161308 |
| NRDC | 24 | 24 | 0,071791035 | 0,033504174 | 0,604161308 |
| TTC7B | 29 | 28 | 0,070347943 | 0,033481507 | 0,604161308 |
| CNIH2 | 2 | 2 | 0,129786066 | 0,033666191 | 0,605572704 |
| RPL24 | 5 | 5 | 0,341618608 | 0,033906006 | 0,608373018 |
| PDE1A | 25 | 23 | 0,11209023 | 0,033997047 | 0,608496657 |
| GNAI1 | 9 | 9 | 0,101421382 | 0,034113613 | 0,60907539 |
| MAP7 | 1 | 1 | -0,22833328 | 0,034835023 | 0,610165802 |
| ARFGAP2 | 8 | 8 | 0,114913603 | 0,034814253 | 0,610165802 |
| TGOLN1;TGOLN2 | 1 | 1 | -0,33778751 | 0,034631831 | 0,610165802 |
| CALU | 10 | 10 | -0,079801006 | 0,034849741 | 0,610165802 |
| RBFOX3;RBFOX1 | 5 | 2 | 0,197407215 | 0,03484634 | 0,610165802 |
| MIPEP | 6 | 6 | -0,135785677 | 0,034847534 | 0,610165802 |
| PRKAA1 | 4 | 4 | 0,116271239 | 0,034396182 | 0,610165802 |
| SLIT1;SLIT2 | 1 | 1 | -0,209653411 | 0,034774101 | 0,610165802 |
| PDPK1 | 17 | 17 | 0,062889054 | 0,035298694 | 0,614618559 |
| DLG2 | 28 | 28 | 0,052551404 | 0,03552905 | 0,614618559 |
| GRIN2B | 39 | 38 | 0,083482943 | 0,035481042 | 0,614618559 |
| CLDND1 | 4 | 4 | 0,103466452 | 0,03549002 | 0,614618559 |
| BACE1 | 1 | 1 | 0,193860337 | 0,035245519 | 0,614618559 |
| TESC | 8 | 8 | 0,075974089 | 0,035979822 | 0,619452602 |
| RAP2C | 5 | 4 | 0,097914538 | 0,035899499 | 0,619452602 |
| MARK1 | 18 | 18 | 0,059421188 | 0,03631136 | 0,62072682 |
| PALM | 4 | 4 | -0,118798125 | 0,036291969 | 0,62072682 |
| SPCS2 | 6 | 6 | 0,087699111 | 0,036156474 | 0,62072682 |
| CPNE6 | 34 | 34 | 0,104090403 | 0,036448291 | 0,621598097 |

| | | | | | |
|---|---|---|---|---|---|
| SAMHD1 | 2 | 1 | 0,444663876 | 0,036930759 | 0,625047517 |
| FXR1 | 9 | 9 | -0,080167883 | 0,036762573 | 0,625047517 |
| RAC3 | 2 | 2 | 0,111263015 | 0,036996313 | 0,625047517 |
| BUB3 | 3 | 2 | 0,232294524 | 0,036825084 | 0,625047517 |
| SNUPN | 1 | 1 | -0,2768942 | 0,037495716 | 0,632008212 |
| PPP2CB;PPP2CA | 14 | 14 | 0,11232655 | 0,037739942 | 0,634645395 |
| PRKAG2 | 16 | 16 | 0,065103466 | 0,037975858 | 0,63713093 |
| ANKFY1 | 13 | 13 | 0,087728985 | 0,038687203 | 0,637334649 |
| KPNB1 | 33 | 33 | 0,055952372 | 0,038210434 | 0,637334649 |
| TBC1D10A | 3 | 3 | 0,105197503 | 0,03850072 | 0,637334649 |
| RAP2B | 7 | 7 | 0,09059717 | 0,038515753 | 0,637334649 |
| ZNF516 | 2 | 2 | -0,183988106 | 0,038693114 | 0,637334649 |
| YWHAQ | 18 | 18 | 0,090876082 | 0,038603588 | 0,637334649 |
| GJA1 | 19 | 18 | -0,130995541 | 0,038539747 | 0,637334649 |
| BPTF | 1 | 1 | -0,261330877 | 0,038304201 | 0,637334649 |
| NCSTN | 8 | 8 | 0,088662242 | 0,039012223 | 0,639339796 |
| RAB26 | 4 | 4 | 0,124043466 | 0,039030847 | 0,639339796 |
| CYTH3;CYTH2;CYTH4 | 1 | 1 | 0,135791409 | 0,039080098 | 0,639339796 |
| PCDHGA9 | 1 | 1 | 0,375412096 | 0,039447835 | 0,641005154 |
| SUSD2 | 2 | 2 | 0,122806998 | 0,039406141 | 0,641005154 |
| TTC9 | 1 | 1 | -0,205213458 | 0,039325857 | 0,641005154 |
| MRTFA | 2 | 2 | -0,306539065 | 0,039625858 | 0,641016964 |
| MACO1 | 2 | 2 | -0,221303997 | 0,039580982 | 0,641016964 |
| ARHGAP25 | 1 | 1 | 0,454495211 | 0,039721798 | 0,64113464 |
| SIPA1L3 | 13 | 13 | -0,081286019 | 0,039862538 | 0,641973302 |
| C1QL3 | 2 | 2 | -0,156809263 | 0,040016773 | 0,643025074 |
| MANF | 3 | 3 | 0,217884946 | 0,040172865 | 0,644101971 |
| FST | 1 | 1 | -0,770518662 | 0,040365265 | 0,64575493 |
| LYSMD3 | 1 | 1 | -0,345905485 | 0,040464119 | 0,645907376 |
| MYH14 | 42 | 42 | -0,069494121 | 0,040925067 | 0,651826346 |
| SRGAP3 | 43 | 43 | 0,038119071 | 0,041587715 | 0,654676586 |
| RAB11FIP3 | 1 | 1 | -0,536749151 | 0,041247394 | 0,654676586 |
| TRPV2 | 6 | 6 | 0,066126123 | 0,041647245 | 0,654676586 |

| | | | | | |
|---|---|---|---|---|---|
| SEC11A | 3 | 3 | 0,129683336 | 0,041618841 | 0,654676586 |
| SF3B1 | 2 | 1 | 0,290470467 | 0,041608631 | 0,654676586 |
| UNC80 | 2 | 2 | -0,1690287 | 0,041301002 | 0,654676586 |
| ROBO1;ROBO2 | 2 | 2 | 0,262944642 | 0,041785945 | 0,655432028 |
| DLGAP3 | 21 | 21 | 0,066651516 | 0,041989578 | 0,655625456 |
| ARL6 | 7 | 7 | 0,135952241 | 0,042070282 | 0,655625456 |
| IMMT | 2 | 2 | -0,153338444 | 0,04195129 | 0,655625456 |
| ME1 | 29 | 29 | 0,107360843 | 0,042546726 | 0,657532704 |
| DNAJC7 | 11 | 10 | 0,083112501 | 0,042542417 | 0,657532704 |
| FBXL16 | 15 | 15 | 0,061183629 | 0,042582773 | 0,657532704 |
| IGFALS | 1 | 1 | 0,184378961 | 0,042647329 | 0,657532704 |
| EFEMP2 | 1 | 1 | -0,322460593 | 0,042293363 | 0,657532704 |
| VWA5A | 18 | 18 | 0,097625609 | 0,042810291 | 0,658640887 |
| RPL26 | 10 | 9 | 0,147811649 | 0,042902502 | 0,658658152 |
| KYAT3 | 12 | 12 | -0,201377094 | 0,043065567 | 0,65976083 |
| ALDH3A2 | 1 | 1 | 0,157702707 | 0,043405177 | 0,663557793 |
| NAP1L5 | 1 | 1 | -0,279492957 | 0,043633947 | 0,665647821 |
| USP4 | 10 | 10 | 0,092773994 | 0,043967845 | 0,66792329 |
| MFGE8 | 4 | 4 | -0,100944775 | 0,043939415 | 0,66792329 |
| RPS27;RPS27L | 2 | 1 | 0,135762906 | 0,04418584 | 0,669827695 |
| OLA1 | 25 | 21 | 0,041245479 | 0,044826933 | 0,675235965 |
| LRRC4B | 12 | 12 | 0,053532748 | 0,044840271 | 0,675235965 |
| SLITRK1 | 3 | 3 | 0,093248178 | 0,044706843 | 0,675235965 |
| ACADS | 7 | 7 | -0,173001047 | 0,044916125 | 0,675235965 |
| GM14151 | 1 | 1 | -0,160052453 | 0,045032433 | 0,675579924 |
| MTMR1 | 14 | 14 | 0,074892348 | 0,045385836 | 0,678068134 |
| KCNA2 | 5 | 5 | 0,082713976 | 0,045348792 | 0,678068134 |
| CC2D2B | 1 | 1 | 0,243263711 | 0,045667091 | 0,680863364 |
| SOD1 | 10 | 9 | -0,068986469 | 0,046115825 | 0,686138956 |
| TSPAN15 | 1 | 1 | -0,262535943 | 0,046253716 | 0,686777454 |
| ALDOA | 21 | 21 | 0,071374066 | 0,046540811 | 0,686809401 |
| DNAJC16 | 3 | 3 | 0,167159926 | 0,046352583 | 0,686809401 |
| IFT81 | 1 | 1 | -0,388063921 | 0,046514023 | 0,686809401 |

| | | | | | |
|---|---|---|---|---|---|
| PPP1CA;PPP1CB | 9 | 9 | 0,086079411 | 0,046842229 | 0,689849613 |
| STX4 | 7 | 7 | -0,121889302 | 0,047065898 | 0,690331653 |
| USP19 | 8 | 8 | 0,096587709 | 0,04701677 | 0,690331653 |
| REM2 | 7 | 7 | 0,142194008 | 0,047559345 | 0,694750751 |
| GABRB1 | 5 | 5 | 0,126107309 | 0,047546337 | 0,694750751 |
| PRR36 | 14 | 14 | -0,12943072 | 0,047948973 | 0,697411615 |
| YKT6 | 10 | 10 | 0,09646827 | 0,048105986 | 0,697411615 |
| FMN2 | 19 | 19 | -0,058865642 | 0,048035408 | 0,697411615 |
| PHYHD1 | 4 | 4 | 0,155831561 | 0,048127285 | 0,697411615 |
| ATP1A2;ATP1A1 | 19 | 19 | 0,059769799 | 0,048302578 | 0,698551877 |
| SRR | 12 | 12 | 0,112541593 | 0,048454834 | 0,699355095 |
| LINGO1 | 16 | 16 | 0,104287012 | 0,049281685 | 0,708460969 |
| MRPL9 | 3 | 3 | -0,158337544 | 0,049207487 | 0,708460969 |
| SLITRK3 | 4 | 4 | 0,103850561 | 0,049460336 | 0,709014571 |
| CTPS2 | 4 | 4 | 0,115774141 | 0,049516299 | 0,709014571 |
| PCDH1 | 23 | 23 | 0,061992499 | 0,049894164 | 0,710206104 |
| CLPX | 10 | 10 | -0,123295238 | 0,049779937 | 0,710206104 |
| ACOT1 | 1 | 1 | -0,363557058 | 0,049823376 | 0,710206104 |
| RPS8 | 5 | 5 | 0,348530136 | 0,050335541 | 0,715081133 |
| **DECR2** | **1** | **1** | **0,190838108** | **0,050766565** | **0,719790254** |
| SH3GL1 | 17 | 17 | 0,035963709 | 0,051031457 | 0,720719666 |
| RPL15 | 8 | 8 | 0,208484313 | 0,050939315 | 0,720719666 |
| CDC42EP4 | 10 | 10 | -0,082879333 | 0,051806163 | 0,721346188 |
| YARS2 | 9 | 9 | -0,122365309 | 0,051871343 | 0,721346188 |
| SPAG1 | 1 | 1 | 0,289131085 | 0,052173151 | 0,721346188 |
| RAB5A | 14 | 14 | 0,052072258 | 0,051702122 | 0,721346188 |
| 5031439G07RIK | 4 | 4 | 0,094566056 | 0,052124807 | 0,721346188 |
| TXLNA | 1 | 1 | 0,600730162 | 0,052049247 | 0,721346188 |
| RAB8A;RAB8B | 3 | 3 | 0,120612493 | 0,051978811 | 0,721346188 |
| ABCC3 | 1 | 1 | -0,44150404 | 0,051636365 | 0,721346188 |
| DNM3 | 1 | 1 | 0,534523934 | 0,051613875 | 0,721346188 |
| ZHX2 | 1 | 1 | -0,190860743 | 0,051572416 | 0,721346188 |
| ERICH5 | 1 | 1 | 0,182932493 | 0,051679898 | 0,721346188 |

| | | | | | |
|---|---|---|---|---|---|
| SLC2A8 | 1 | 1 | -0,214164867 | 0,052381714 | 0,722847662 |
| ZFPL1 | 1 | 1 | -0,245415409 | 0,05270745 | 0,725720506 |
| TMEM30A | 11 | 11 | 0,078868827 | 0,052790622 | 0,725720506 |
| CDS2 | 7 | 7 | -0,071357897 | 0,05310647 | 0,726508835 |
| FHL1 | 2 | 2 | 0,269687276 | 0,053149381 | 0,726508835 |
| H1-4;H1-3 | 3 | 2 | 0,840304753 | 0,053090133 | 0,726508835 |
| ESF1 | 1 | 1 | -0,13643341 | 0,05330968 | 0,727325084 |
| MARCKSL1 | 2 | 2 | -0,14686803 | 0,053535087 | 0,729024885 |
| NFASC | 4 | 4 | -0,232259147 | 0,053648609 | 0,72919754 |
| KPNA3 | 9 | 9 | 0,056049604 | 0,05431246 | 0,734623207 |
| EPS8L1 | 1 | 1 | 0,518293164 | 0,054170031 | 0,734623207 |
| NHSL1 | 1 | 1 | -0,189466077 | 0,054352567 | 0,734623207 |
| IL16 | 1 | 1 | 0,267456071 | 0,054773443 | 0,736183582 |
| IGHV1-4;IGHV1-7;IGI | 1 | 1 | -0,251956869 | 0,054638333 | 0,736183582 |
| AUP1 | 1 | 1 | -0,150756888 | 0,054720492 | 0,736183582 |
| TIMM17B | 2 | 2 | -0,121095989 | 0,055061408 | 0,738680964 |
| MYO18A | 70 | 70 | -0,036879365 | 0,055175782 | 0,738844585 |
| RPL11 | 6 | 6 | 0,105847436 | 0,055501991 | 0,741469835 |
| RELN | 4 | 3 | -0,404444509 | 0,055576912 | 0,741469835 |
| ARPC5L | 6 | 6 | 0,073357107 | 0,055760455 | 0,742548529 |
| TRNT1 | 14 | 14 | -0,075989169 | 0,056023303 | 0,743311019 |
| PRPS1L1 | 4 | 4 | 0,087952435 | 0,05598101 | 0,743311019 |
| UFC1 | 6 | 6 | -0,112030322 | 0,056293055 | 0,745522121 |
| TEX19.2 | 1 | 1 | 0,161129375 | 0,056467729 | 0,746468281 |
| SYNJ1 | 51 | 51 | 0,051968325 | 0,056634082 | 0,747301175 |
| TOMM34 | 18 | 17 | 0,072687276 | 0,057001037 | 0,750773226 |
| ATP2B3 | 33 | 33 | 0,063089779 | 0,057115297 | 0,750910389 |
| KCNQ2 | 7 | 7 | 0,095709522 | 0,057548774 | 0,755236267 |
| PPP4R4 | 7 | 7 | 0,086734643 | 0,058154351 | 0,758594603 |
| VTN | 1 | 1 | 0,533442312 | 0,05843413 | 0,758594603 |
| ACYP2 | 4 | 4 | -0,106688013 | 0,05826735 | 0,758594603 |
| ERCC5 | 1 | 1 | -0,332512716 | 0,058419821 | 0,758594603 |
| MAPK10;MAPK8;MA | 1 | 1 | 0,202257003 | 0,058389562 | 0,758594603 |

| | | | | | |
|---|---|---|---|---|---|
| IL6ST | 1 | 1 | -0,138010346 | 0,058269409 | 0,758594603 |
| PSME2 | 8 | 8 | 0,074356973 | 0,058755153 | 0,759648751 |
| RIMBP2 | 19 | 19 | 0,041284753 | 0,05869178 | 0,759648751 |
| STAU1 | 3 | 2 | 0,266970048 | 0,058830494 | 0,759648751 |
| EHD1 | 20 | 20 | 0,058818184 | 0,059116154 | 0,760967778 |
| HSD17B11 | 5 | 5 | 0,065315735 | 0,059143119 | 0,760967778 |
| MYO6 | 29 | 29 | -0,047379848 | 0,059625417 | 0,764452825 |
| PLXNA1;PLXNA2;PLX | 7 | 7 | 0,06105727 | 0,059625241 | 0,764452825 |
| SPAG9 | 29 | 27 | -0,054928987 | 0,059858029 | 0,766076824 |
| ARL8A | 4 | 4 | 0,129891682 | 0,060170573 | 0,767360517 |
| BCAS3 | 15 | 15 | 0,100093098 | 0,060079357 | 0,767360517 |
| RASAL1 | 31 | 31 | 0,07813006 | 0,060812976 | 0,77418773 |
| CUL5 | 33 | 32 | 0,043811004 | 0,061069391 | 0,776085701 |
| CASKIN1 | 52 | 52 | -0,070728078 | 0,061733107 | 0,779771136 |
| NQO1 | 8 | 7 | -0,079362973 | 0,061790743 | 0,779771136 |
| CSNK1D;CSNK1E | 10 | 10 | 0,061566315 | 0,061669057 | 0,779771136 |
| TJP1 | 1 | 1 | -0,187899589 | 0,061634249 | 0,779771136 |
| RCN2 | 14 | 14 | -0,075411906 | 0,062181918 | 0,780620567 |
| CLIC6;CLIC4 | 1 | 1 | -0,598446886 | 0,062120667 | 0,780620567 |
| DBR1 | 1 | 1 | -0,273353911 | 0,062143437 | 0,780620567 |
| LRRC4B;LRRC4C;LRR( | 1 | 1 | 0,12946392 | 0,062446837 | 0,782587653 |
| MYH11;MYH10 | 3 | 3 | -0,074511146 | 0,062703484 | 0,783089625 |
| SPX | 1 | 1 | -0,277144445 | 0,062601518 | 0,783089625 |
| CALB1 | 26 | 18 | -0,096905322 | 0,063745248 | 0,786590259 |
| ENHO | 1 | 1 | -0,16720777 | 0,063624876 | 0,786590259 |
| UBQLN4;UBQLN2 | 1 | 1 | 0,185353089 | 0,063654047 | 0,786590259 |
| RPL10 | 9 | 9 | 0,220618031 | 0,06323269 | 0,786590259 |
| WASHC4 | 11 | 11 | 0,05875237 | 0,063306921 | 0,786590259 |
| DEPDC5 | 1 | 1 | -0,227876348 | 0,063419467 | 0,786590259 |
| KCTD6 | 3 | 3 | 0,229903286 | 0,063354704 | 0,786590259 |
| PALLD | 1 | 1 | -0,199807234 | 0,063906554 | 0,787237295 |
| CAMK1D;CAMK1 | 5 | 5 | 0,105574608 | 0,06422511 | 0,789815932 |
| ELAVL4 | 4 | 3 | 0,194207889 | 0,064825971 | 0,795851611 |

| | | | | | |
|---|---|---|---|---|---|
| CAMTA2 | 1 | 1 | -0,33500992 | 0,06494646 | 0,795979411 |
| NCK2 | 12 | 12 | 0,049402427 | 0,065319588 | 0,799197873 |
| HNRNPK | 22 | 19 | 0,105517408 | 0,065607146 | 0,799997361 |
| PITPNM3 | 3 | 3 | -0,140958915 | 0,065530394 | 0,799997361 |
| RAB2A | 8 | 8 | 0,086248352 | 0,065827469 | 0,799997361 |
| ARMC9 | 1 | 1 | 0,410719807 | 0,065755612 | 0,799997361 |
| VPS39 | 6 | 6 | 0,076188975 | 0,066035077 | 0,8011739 |
| GPRIN1 | 45 | 44 | -0,11393436 | 0,066311101 | 0,80119067 |
| IDH2 | 27 | 27 | -0,229124956 | 0,066638745 | 0,80119067 |
| ARHGAP32 | 12 | 12 | -0,045551833 | 0,066376957 | 0,80119067 |
| RAB3IP | 3 | 3 | 0,105979178 | 0,066487554 | 0,80119067 |
| ZBTB18 | 2 | 1 | 0,28021413 | 0,066701256 | 0,80119067 |
| LRRN1 | 1 | 1 | -0,124829631 | 0,066202207 | 0,80119067 |
| GPM6B | 5 | 5 | 0,077444016 | 0,067108339 | 0,801863985 |
| GJB6 | 2 | 2 | -0,160694963 | 0,067154582 | 0,801863985 |
| CSNK1G1 | 2 | 2 | 0,133311862 | 0,066926936 | 0,801863985 |
| PKP1 | 5 | 2 | 0,75201413 | 0,067200882 | 0,801863985 |
| CALCOCO1 | 14 | 14 | -0,045277781 | 0,067422025 | 0,803177371 |
| NUDT9 | 6 | 4 | -0,094735741 | 0,067669807 | 0,804803248 |
| GPX4 | 9 | 9 | 0,051882055 | 0,068045668 | 0,807944537 |
| YWHAG;YWHAH | 3 | 3 | 0,112327098 | 0,068252066 | 0,809066704 |
| RAPGEF2 | 60 | 59 | 0,052533546 | 0,068570745 | 0,811514006 |
| PDZD11 | 2 | 2 | 0,127543685 | 0,069082271 | 0,812250252 |
| ATP9A | 11 | 11 | 0,062709274 | 0,069033449 | 0,812250252 |
| XPR1 | 1 | 1 | -0,140503695 | 0,069067993 | 0,812250252 |
| ACAP2 | 1 | 1 | 0,153391653 | 0,069058011 | 0,812250252 |
| VPS13C | 63 | 63 | 0,03997807 | 0,070103539 | 0,821624727 |
| YTHDF1;YTHDF3 | 1 | 1 | 0,255976118 | 0,070106826 | 0,821624727 |
| ATP6V1E1 | 28 | 26 | 0,058787065 | 0,070338845 | 0,823010017 |
| RRBP1 | 20 | 20 | -0,041019399 | 0,070796606 | 0,827027875 |
| AP3B2 | 50 | 49 | 0,039767352 | 0,071397526 | 0,829924773 |
| RPL29 | 2 | 2 | 0,301927158 | 0,071503683 | 0,829924773 |
| VPS18 | 14 | 14 | -0,060568752 | 0,071423323 | 0,829924773 |

| | | | | | |
|---|---|---|---|---|---|
| GM35060 | 1 | 1 | -0,380968762 | 0,071249729 | 0,829924773 |
| USP30 | 2 | 2 | -0,127408689 | 0,071791705 | 0,830439464 |
| NRAS;HRAS | 6 | 6 | 0,112169537 | 0,071707041 | 0,830439464 |
| DISC1 | 1 | 1 | -0,200132974 | 0,07189256 | 0,830439464 |
| STK4 | 4 | 4 | 0,103687567 | 0,07226994 | 0,830817071 |
| NPTN | 6 | 6 | 0,099690996 | 0,072129404 | 0,830817071 |
| HIVEP2 | 4 | 3 | -0,119209698 | 0,072248622 | 0,830817071 |
| PRKAR2A | 1 | 1 | -0,1123308 | 0,072683521 | 0,831945481 |
| HDAC11 | 8 | 8 | -0,068836729 | 0,072713255 | 0,831945481 |
| ALDH7A1 | 24 | 24 | -0,097604319 | 0,072688218 | 0,831945481 |
| LIMCH1 | 8 | 8 | -0,081383019 | 0,073088452 | 0,834917218 |
| BPNT1 | 1 | 1 | -0,245766582 | 0,073343919 | 0,836513998 |
| COPA | 24 | 24 | -0,103145415 | 0,073490333 | 0,836863929 |
| PSD3 | 34 | 34 | 0,049819329 | 0,073810083 | 0,838408258 |
| SYNGAP1 | 1 | 1 | -0,129672013 | 0,073857843 | 0,838408258 |
| SYN1 | 40 | 40 | 0,054611651 | 0,074162224 | 0,840543955 |
| ALB | 40 | 21 | 0,251960552 | 0,074858689 | 0,84184009 |
| LRPAP1 | 17 | 17 | -0,067415034 | 0,074629869 | 0,84184009 |
| PLPPR3 | 8 | 8 | -0,068609652 | 0,074750317 | 0,84184009 |
| TCP1 | 34 | 34 | 0,06875354 | 0,074447948 | 0,84184009 |
| TMF1 | 1 | 1 | -0,197482712 | 0,074669529 | 0,84184009 |
| AIFM3 | 10 | 10 | -0,067136435 | 0,075616178 | 0,843816121 |
| GALK1 | 9 | 6 | -0,069506275 | 0,075505697 | 0,843816121 |
| CACNA1C | 7 | 7 | 0,080104539 | 0,075617874 | 0,843816121 |
| RPL3 | 12 | 11 | 0,243346798 | 0,075194175 | 0,843816121 |
| PNKD | 1 | 1 | -0,154968866 | 0,075449393 | 0,843816121 |
| DMXL2 | 123 | 123 | 0,052547814 | 0,075807041 | 0,843941698 |
| SDS | 1 | 1 | -0,178562338 | 0,075862551 | 0,843941698 |
| TRHDE | 9 | 9 | 0,061713257 | 0,07671071 | 0,843949156 |
| CLMN | 14 | 14 | -0,064815964 | 0,076740258 | 0,843949156 |
| RPL35A | 9 | 9 | 0,089223688 | 0,076500946 | 0,843949156 |
| DCDC2B | 1 | 1 | 0,381225253 | 0,076913635 | 0,843949156 |
| GABARAPL1 | 3 | 3 | 0,102023413 | 0,076883873 | 0,843949156 |

| | | | | | |
|---|---|---|---|---|---|
| CD99L2 | 2 | 2 | 0,12513289 | 0,076657419 | 0,843949156 |
| SLC7A11 | 4 | 4 | -0,152474896 | 0,076625579 | 0,843949156 |
| CLNS1A | 2 | 2 | 0,127060147 | 0,076880025 | 0,843949156 |
| ATRN | 1 | 1 | 0,160182499 | 0,076048072 | 0,843949156 |
| MAOA | 19 | 18 | -0,084308394 | 0,077306428 | 0,84569256 |
| RANBP10 | 2 | 2 | -0,101666973 | 0,077219684 | 0,84569256 |
| SGTA | 1 | 1 | 0,257308181 | 0,07747929 | 0,846303239 |
| EFTUD2 | 5 | 3 | 0,360737023 | 0,077682809 | 0,847246445 |
| ITPR2 | 1 | 1 | 0,156994827 | 0,078471919 | 0,854563924 |
| ACSF2 | 20 | 20 | -0,194175773 | 0,079063508 | 0,854571337 |
| ELOB | 11 | 11 | 0,070342198 | 0,079036218 | 0,854571337 |
| EEF1D | 1 | 1 | -0,258579428 | 0,078709625 | 0,854571337 |
| FERMT3 | 6 | 6 | 0,176602771 | 0,078927679 | 0,854571337 |
| GOSR2 | 1 | 1 | -0,165642685 | 0,078976859 | 0,854571337 |
| PPP3CB | 17 | 17 | 0,077743399 | 0,079462808 | 0,85692867 |
| DMTN | 24 | 22 | -0,064613855 | 0,07951862 | 0,85692867 |
| CHMP1B1;CHMP1B2 | 2 | 2 | 0,101994215 | 0,079712182 | 0,857736293 |
| PABPC4 | 12 | 10 | -0,085553849 | 0,080079025 | 0,860403317 |
| PIP5K1C;PIP5K1B;PIF | 2 | 2 | 0,103562387 | 0,081018656 | 0,869207568 |
| PKM | 52 | 51 | 0,083795312 | 0,081537021 | 0,869789829 |
| TPP2 | 46 | 45 | 0,075599388 | 0,081554073 | 0,869789829 |
| VCP | 73 | 71 | 0,062182243 | 0,081386254 | 0,869789829 |
| ATP2B4 | 13 | 13 | -0,145861273 | 0,081435901 | 0,869789829 |
| TMED5 | 1 | 1 | 0,399347887 | 0,081788708 | 0,871007583 |
| RTN2 | 1 | 1 | -0,230164935 | 0,081944807 | 0,871232757 |
| SNRPN | 3 | 2 | 0,476108455 | 0,082050824 | 0,871232757 |
| TSFM | 8 | 8 | -0,113236108 | 0,082557545 | 0,875327876 |
| SYNGR1 | 3 | 3 | 0,072460641 | 0,083069351 | 0,879464827 |
| RPL27A | 4 | 4 | 0,306789006 | 0,083366719 | 0,881322729 |
| RPL14 | 5 | 4 | 0,261062231 | 0,083952136 | 0,884924049 |
| RTN4RL2 | 6 | 6 | 0,064828815 | 0,083880583 | 0,884924049 |
| STMN1 | 7 | 7 | -0,055715451 | 0,084218987 | 0,885156245 |
| RAB9B | 4 | 4 | 0,092725591 | 0,08419785 | 0,885156245 |

| | | | | | |
|---|---|---|---|---|---|
| RPLP0 | 13 | 13 | 0,08179128 | 0,084352446 | 0,885272191 |
| REXO2 | 4 | 4 | -0,153607908 | 0,084725598 | 0,885333526 |
| sp\|Q8BH50\|CR025_ | 2 | 2 | -0,139369077 | 0,084721929 | 0,885333526 |
| RIPOR2 | 3 | 3 | -0,153398079 | 0,084645553 | 0,885333526 |
| MRPL46 | 5 | 5 | -0,09823767 | 0,085051737 | 0,885985996 |
| MRPL13 | 4 | 4 | -0,085770051 | 0,085068609 | 0,885985996 |
| MAP3K6 | 1 | 1 | -0,154571477 | 0,085155617 | 0,885985996 |
| DGKZ | 8 | 8 | 0,079827181 | 0,085935731 | 0,888986086 |
| ARPC2 | 29 | 29 | 0,05438101 | 0,085845421 | 0,888986086 |
| CAPZB | 26 | 26 | 0,038936498 | 0,085809838 | 0,888986086 |
| PPP2CA | 2 | 2 | 0,123580463 | 0,085643178 | 0,888986086 |
| DOCK2 | 5 | 5 | 0,103087649 | 0,086314199 | 0,891118476 |
| EIF3I | 11 | 7 | 0,097510119 | 0,086388335 | 0,891118476 |
| CSNK1G1;CSNK1G2;( | 5 | 5 | 0,077108961 | 0,087133092 | 0,894970723 |
| AKT1 | 8 | 6 | 0,16983768 | 0,087122201 | 0,894970723 |
| GCKR | 1 | 1 | -0,382776362 | 0,086895388 | 0,894970723 |
| APBB1 | 9 | 9 | 0,059848546 | 0,087430514 | 0,896751843 |
| PPP2R5E | 14 | 14 | 0,045609878 | 0,087639609 | 0,897623252 |
| FRMPD1 | 1 | 1 | 0,603532203 | 0,087790062 | 0,897892417 |
| CEP170B | 35 | 35 | -0,070650395 | 0,088444863 | 0,898629163 |
| PDCD10 | 7 | 7 | 0,077071771 | 0,088914366 | 0,898629163 |
| STRN4 | 16 | 16 | -0,051242435 | 0,088980567 | 0,898629163 |
| GJC2 | 2 | 2 | 0,145517602 | 0,088835662 | 0,898629163 |
| PEX14 | 6 | 6 | -0,077777439 | 0,088636587 | 0,898629163 |
| CPNE2 | 9 | 9 | 0,160920205 | 0,088346226 | 0,898629163 |
| PPP2R5C | 7 | 7 | 0,130503919 | 0,088902184 | 0,898629163 |
| ALS2CL | 1 | 1 | -0,188656519 | 0,088147527 | 0,898629163 |
| SYT17 | 13 | 13 | 0,086820929 | 0,088157807 | 0,898629163 |
| RALA | 7 | 7 | 0,05586439 | 0,089310753 | 0,898800848 |
| PNPO | 6 | 6 | -0,088599771 | 0,089370462 | 0,898800848 |
| PVR | 1 | 1 | 0,224603447 | 0,089200489 | 0,898800848 |
| EXD2 | 6 | 6 | -0,100979715 | 0,089782858 | 0,899196463 |
| THEMIS3 | 1 | 1 | 0,502029633 | 0,089691684 | 0,899196463 |

| | | | | | |
|---|---|---|---|---|---|
| KIFC1 | 1 | 1 | -0,159598799 | 0,089557725 | 0,899196463 |
| TBC1D14 | 1 | 1 | 0,245032943 | 0,089922874 | 0,899353118 |
| ALYREF;ALYREF2 | 1 | 1 | 0,44166031 | 0,090319641 | 0,902073655 |
| NAPB | 26 | 26 | 0,038094688 | 0,090827564 | 0,905895329 |
| MIA2 | 6 | 6 | -0,087806903 | 0,091428576 | 0,910536541 |
| TAMALIN | 4 | 4 | 0,080404918 | 0,091569712 | 0,910536541 |
| HBS1L | 3 | 3 | 0,148118352 | 0,091670668 | 0,910536541 |
| KIF21A | 37 | 35 | 0,026663025 | 0,092511085 | 0,91714037 |
| ANKS1B | 3 | 3 | 0,087269813 | 0,092589195 | 0,91714037 |
| GNAZ | 20 | 20 | 0,076453702 | 0,092758099 | 0,917556514 |
| IGFBP5 | 1 | 1 | 0,274003749 | 0,092903087 | 0,917735278 |
| DOCK3 | 19 | 19 | -0,067804337 | 0,093050497 | 0,917937437 |
| CAPN5 | 22 | 22 | 0,047766324 | 0,093301828 | 0,919162829 |
| ANK3 | 42 | 42 | -0,087295095 | 0,093605212 | 0,920896991 |
| MACF1 | 41 | 41 | -0,058216859 | 0,093750841 | 0,921076537 |
| STAC | 1 | 1 | 0,201938246 | 0,09396355 | 0,921913742 |
| BAG3 | 3 | 3 | -0,117668927 | 0,094285402 | 0,922725546 |
| STXBP5 | 23 | 22 | 0,051731427 | 0,094437066 | 0,922725546 |
| EIF4G1 | 1 | 1 | 0,201739024 | 0,09430248 | 0,922725546 |
| KRT80 | 4 | 1 | 0,443450845 | 0,094556718 | 0,922725546 |
| RASGRF2 | 15 | 15 | -0,078038997 | 0,094741941 | 0,923072628 |
| PCDH7 | 14 | 14 | -0,055292008 | 0,094847595 | 0,923072628 |
| C1QTNF4 | 4 | 4 | -0,113994147 | 0,095021997 | 0,923526963 |
| ABHD12 | 17 | 17 | 0,035075718 | 0,095530675 | 0,927224576 |
| CAMKV | 25 | 25 | 0,048472695 | 0,096800217 | 0,929953877 |
| EPB41L1 | 62 | 62 | -0,025951675 | 0,096520867 | 0,929953877 |
| AP1B1 | 29 | 29 | 0,04072996 | 0,096155945 | 0,929953877 |
| TRIM2 | 25 | 25 | 0,034387698 | 0,096263787 | 0,929953877 |
| DOCK10 | 11 | 11 | -0,076230257 | 0,096840723 | 0,929953877 |
| NUDT18 | 1 | 1 | 0,207540174 | 0,096791263 | 0,929953877 |
| STXBP5L | 6 | 6 | 0,054140677 | 0,096305466 | 0,929953877 |
| SORBS1 | 14 | 13 | -0,06040087 | 0,096663926 | 0,929953877 |
| CIT | 16 | 16 | -0,076729648 | 0,09842262 | 0,942641015 |

| | | | | | |
|---|---|---|---|---|---|
| SIRPA | 19 | 19 | 0,06742607 | 0,098303671 | 0,942641015 |
| PPP3R1 | 20 | 20 | 0,091103295 | 0,09856411 | 0,942747464 |
| BIN2 | 1 | 1 | -0,151624452 | 0,099196755 | 0,9450484 |
| CCDC77 | 1 | 1 | 0,205788616 | 0,099096277 | 0,9450484 |
| PRSS16 | 1 | 1 | 0,430144395 | 0,099036317 | 0,9450484 |
| ALDH3A2 | 14 | 14 | 0,046944113 | 0,099747545 | 0,949045389 |
| MAP6 | 68 | 67 | -0,127034711 | 0,100180605 | 0,949285886 |
| TRAPPC4 | 4 | 4 | 0,114162176 | 0,100429222 | 0,949285886 |
| CNTNAP2 | 28 | 28 | 0,043344456 | 0,100243946 | 0,949285886 |
| TRMT112 | 2 | 2 | 0,118717935 | 0,099970358 | 0,949285886 |
| ZMYND15 | 1 | 1 | -0,253505969 | 0,100338002 | 0,949285886 |
| KIF16B | 2 | 1 | 0,212643611 | 0,10073626 | 0,950945037 |
| ALDH6A1 | 31 | 31 | -0,156429176 | 0,101584142 | 0,952730134 |
| RPL12 | 6 | 5 | 0,100449972 | 0,101553269 | 0,952730134 |
| TFB1M | 3 | 3 | -0,112448706 | 0,101484863 | 0,952730134 |
| FBXL18 | 1 | 1 | 0,156007513 | 0,101204132 | 0,952730134 |
| GFRA2 | 2 | 2 | -0,126111441 | 0,101431456 | 0,952730134 |
| VPS13A | 32 | 32 | -0,031782457 | 0,101771574 | 0,95325162 |
| PSPH | 9 | 8 | 0,065153896 | 0,102221502 | 0,954991839 |
| SENP8 | 2 | 2 | -0,166919221 | 0,10214405 | 0,954991839 |
| MPP3 | 8 | 8 | 0,046945152 | 0,102743523 | 0,958029164 |
| GAA | 16 | 15 | -0,072325701 | 0,102811593 | 0,958029164 |
| UNC45B | 3 | 3 | -0,13816555 | 0,102977931 | 0,958344169 |
| FAM126B | 18 | 18 | 0,048732646 | 0,103246877 | 0,95838019 |
| POU2F3 | 1 | 1 | 0,120870351 | 0,103233121 | 0,95838019 |
| EHD3 | 23 | 23 | 0,027607359 | 0,103477559 | 0,959290035 |
| PANK4 | 14 | 13 | 0,053036973 | 0,103884017 | 0,960462838 |
| PTK2B | 44 | 38 | 0,073484811 | 0,104135371 | 0,960462838 |
| SYNGAP1;DAB2IP | 2 | 2 | 0,105189948 | 0,104102791 | 0,960462838 |
| GRN | 1 | 1 | -0,282964799 | 0,103878045 | 0,960462838 |
| PI4K2A | 11 | 11 | 0,047872929 | 0,104417239 | 0,960944945 |
| STAT1 | 1 | 1 | -0,365538408 | 0,104453426 | 0,960944945 |
| DTNB;DTNA | 1 | 1 | -0,100871586 | 0,105487993 | 0,96922958 |

| | | | | | |
|---|---|---|---|---|---|
| GPT | 7 | 7 | -0,096235461 | 0,1058065 | 0,969691765 |
| MAP3K2 | 1 | 1 | -0,345894772 | 0,105787037 | 0,969691765 |
| VARS1 | 31 | 28 | 0,043598064 | 0,107099881 | 0,977827327 |
| DAAM1 | 18 | 18 | 0,046046618 | 0,106968458 | 0,977827327 |
| MRPL39 | 7 | 7 | -0,101070003 | 0,107083813 | 0,977827327 |
| MAP1A | 126 | 113 | -0,103343587 | 0,107642506 | 0,979072907 |
| SRSF1 | 3 | 2 | 0,216792422 | 0,107635663 | 0,979072907 |
| MIA3 | 9 | 9 | -0,059567668 | 0,107423362 | 0,979072907 |
| PSMG2 | 3 | 3 | 0,179471358 | 0,107808274 | 0,979348777 |
| UST | 2 | 2 | -0,106494721 | 0,108292301 | 0,982511451 |
| KAT6A | 1 | 1 | 0,263995622 | 0,108466514 | 0,982858849 |
| MAP4 | 16 | 14 | -0,068465237 | 0,108730821 | 0,983248632 |
| NOCT | 2 | 2 | -0,148512173 | 0,108906385 | 0,983248632 |
| GABRA2;GABRA1;GA | 3 | 3 | 0,152387329 | 0,10891746 | 0,983248632 |
| WDR77 | 8 | 8 | 0,058783111 | 0,109615522 | 0,984633339 |
| ENTPD2 | 9 | 9 | -0,109338276 | 0,109348866 | 0,984633339 |
| COL11A2 | 1 | 1 | 0,287594022 | 0,109579934 | 0,984633339 |
| LY6H | 8 | 8 | 0,157323188 | 0,109564675 | 0,984633339 |
| ARHGEF9 | 6 | 6 | 0,069285485 | 0,110164157 | 0,98833377 |
| UNC13A | 35 | 35 | 0,051596611 | 0,110634094 | 0,988869139 |
| KCNJ10 | 5 | 5 | 0,088158344 | 0,110612117 | 0,988869139 |
| NMD3 | 1 | 1 | 0,178509755 | 0,110383762 | 0,988869139 |
| DNAJB3 | 1 | 1 | -0,157024784 | 0,110812942 | 0,989244916 |
| RPLP1 | 2 | 2 | 0,123960177 | 0,111893039 | 0,990094001 |
| SLC9A6 | 8 | 8 | -0,054335854 | 0,112081983 | 0,990094001 |
| PFKM | 40 | 36 | 0,066354618 | 0,111615446 | 0,990094001 |
| NAPA | 20 | 20 | 0,048318139 | 0,111519265 | 0,990094001 |
| COPS8 | 7 | 7 | 0,077495922 | 0,112039734 | 0,990094001 |
| PEX5L | 2 | 2 | 0,096761265 | 0,112058531 | 0,990094001 |
| CD109 | 6 | 6 | 0,11955594 | 0,112183884 | 0,990094001 |
| RPL23A | 7 | 5 | 0,203810549 | 0,112235689 | 0,990094001 |
| EBP | 1 | 1 | 0,131294488 | 0,111570193 | 0,990094001 |
| ATP2B1 | 6 | 6 | 0,085732783 | 0,11227729 | 0,990094001 |

| | | | | | |
|---|---|---|---|---|---|
| VPS28 | 4 | 4 | 0,075141491 | 0,112800374 | 0,9922865 |
| TRY10 | 2 | 2 | -0,194687963 | 0,11279718 | 0,9922865 |
| SPARC | 1 | 1 | -0,124722611 | 0,113150978 | 0,994161263 |
| BCAM | 1 | 1 | 0,17446247 | 0,113472852 | 0,995779357 |
| AKAP5 | 20 | 20 | -0,071322632 | 0,113860737 | 0,995950076 |
| DNAJC5 | 8 | 8 | 0,074431268 | 0,113960261 | 0,995950076 |
| RAB12 | 11 | 11 | 0,065247071 | 0,114043239 | 0,995950076 |
| PIP4K2A;PIP4K2B | 4 | 4 | 0,055881907 | 0,113849107 | 0,995950076 |
| HSP90AA1;HSP90AB | 16 | 15 | 0,049759371 | 0,114626555 | 0,999803901 |
| CRACDL | 20 | 20 | -0,06171964 | 0,1152185 | 0,999803901 |
| TACC1 | 8 | 8 | -0,065194832 | 0,115105166 | 0,999803901 |
| ETF1 | 12 | 12 | 0,083870283 | 0,115080838 | 0,999803901 |
| RPL23 | 4 | 4 | 0,184108281 | 0,115314127 | 0,999803901 |
| GRCC10 | 3 | 3 | 0,119926573 | 0,114829941 | 0,999803901 |
| IPO5 | 35 | 33 | 0 | 1 | 1 |
| DDRGK1 | 3 | 3 | -0,059136542 | 0,338839713 | 1 |
| UAP1L1 | 18 | 17 | -4,57474E-15 | 0,999999759 | 1 |
| PSMD4 | 3 | 2 | 0 | 1 | 1 |
| PSMD4 | 1 | 1 | 0,01753524 | 0,75025932 | 1 |
| MRPS7 | 5 | 5 | -0,069570523 | 0,300147389 | 1 |
| EDC4 | 9 | 9 | -0,065846334 | 0,157646198 | 1 |
| SEPTIN6 | 10 | 10 | 0 | 1 | 1 |
| NCKIPSD | 34 | 32 | 0 | 1 | 1 |
| FH | 29 | 29 | -5,05464E-18 | 0,999999997 | 1 |
| TOMM70 | 28 | 28 | 1,23943E-17 | 0,999999997 | 1 |
| PIGS | 7 | 7 | 4,39088E-16 | 0,999999934 | 1 |
| EFHD2 | 20 | 20 | -1,2091E-15 | 0,999999864 | 1 |
| RUFY1 | 7 | 7 | -1,97608E-15 | 0,999999977 | 1 |
| PCLO | 170 | 170 | -0,013012916 | 0,574147641 | 1 |
| TARS3 | 24 | 16 | -2,25686E-18 | 0,999999996 | 1 |
| CCDC177 | 19 | 19 | -0,003737496 | 0,736076091 | 1 |
| ACBD6 | 6 | 6 | -6,10162E-16 | 0,999999936 | 1 |
| SEMA7A | 8 | 8 | -0,015292178 | 0,489057173 | 1 |

| | | | | | |
|---|---|---|---|---|---|
| STIP1 | 55 | 52 | 8,88858E-17 | 0,999999961 | 1 |
| TSC1 | 17 | 17 | -0,049151145 | 0,14020907 | 1 |
| ANK3 | 38 | 38 | -0,021380472 | 0,496119089 | 1 |
| CTNND1 | 18 | 18 | -4,50568E-18 | 0,999999993 | 1 |
| CHCHD3 | 14 | 14 | -3,30217E-16 | 0,99999996 | 1 |
| GOLGB1 | 4 | 4 | -1,45609E-17 | 0,999999992 | 1 |
| SEPTIN11 | 11 | 11 | 0 | 1 | 1 |
| PNP | 22 | 21 | 1,83305E-15 | 0,999999906 | 1 |
| PKP4 | 34 | 34 | 1,95585E-17 | 0,999999985 | 1 |
| TTC4 | 2 | 1 | 0 | 1 | 1 |
| RAB3GAP2 | 32 | 32 | 1,27013E-17 | 0,99999999 | 1 |
| CAMSAP3 | 14 | 14 | -1,92109E-18 | 0,999999998 | 1 |
| PDHA1 | 37 | 37 | -5,23445E-16 | 0,999999951 | 1 |
| NDUFA6 | 1 | 1 | -0,066241876 | 0,456807854 | 1 |
| SND1 | 37 | 37 | -6,22798E-18 | 0,999999991 | 1 |
| PANK2 | 5 | 5 | 0,042509269 | 0,211696806 | 1 |
| BCR | 22 | 22 | -7,47035E-17 | 0,999999965 | 1 |
| RAB3GAP1 | 23 | 22 | 0,019290873 | 0,471367286 | 1 |
| EXOC8 | 20 | 20 | 0,00201038 | 0,799676575 | 1 |
| IDH3B | 36 | 35 | -8,40287E-16 | 0,999999946 | 1 |
| PSMD9 | 9 | 8 | -4,14684E-17 | 0,999999991 | 1 |
| FMNL2;FMNL3 | 4 | 4 | -2,74745E-18 | 0,999999998 | 1 |
| HDLBP | 21 | 20 | 4,95248E-16 | 0,999999938 | 1 |
| AGK | 26 | 26 | -7,86704E-16 | 0,999999918 | 1 |
| UBE3A | 26 | 24 | -1,52211E-16 | 0,999999963 | 1 |
| MICU1 | 15 | 15 | 0 | 1 | 1 |
| TPM3 | 3 | 3 | 0 | 1 | 1 |
| ADK | 17 | 15 | -3,76847E-18 | 0,999999996 | 1 |
| PACSIN2 | 15 | 15 | -6,79699E-17 | 0,99999997 | 1 |
| VPS52 | 24 | 24 | 0,021117171 | 0,379927113 | 1 |
| CSE1L | 36 | 31 | 1,81764E-17 | 0,99999999 | 1 |
| RABL6 | 17 | 17 | 0,044688019 | 0,443730811 | 1 |
| LMTK3 | 9 | 9 | -0,069241661 | 0,121871398 | 1 |

| | | | | | |
|---|---|---|---|---|---|
| VDAC3 | 12 | 12 | 4,79741E-17 | 0,999999983 | 1 |
| DGLUCY | 8 | 8 | -0,029610609 | 0,536092093 | 1 |
| CAND1 | 44 | 42 | 0,013592371 | 0,42886589 | 1 |
| ATP2C1 | 8 | 8 | 0 | 1 | 1 |
| SORL1 | 23 | 23 | 2,04296E-17 | 0,999999992 | 1 |
| SRPK2 | 11 | 11 | 0 | 1 | 1 |
| AP2A1 | 45 | 44 | 1,15658E-16 | 0,999999953 | 1 |
| ADD2 | 36 | 36 | -0,035407637 | 0,239220458 | 1 |
| PLXNA4 | 41 | 41 | 9,54119E-15 | 0,999999647 | 1 |
| ALDH4A1 | 13 | 12 | -0,097084721 | 0,292617228 | 1 |
| PPP1R12C | 10 | 10 | -0,007733877 | 0,696565779 | 1 |
| YBX1 | 3 | 3 | -3,35613E-17 | 0,999999989 | 1 |
| PAG1 | 2 | 2 | 6,94491E-18 | 0,999999997 | 1 |
| PRKRA | 6 | 6 | 0 | 1 | 1 |
| SPG7 | 12 | 12 | -1,3707E-17 | 0,999999991 | 1 |
| AAK1 | 37 | 37 | 2,34992E-15 | 0,999999851 | 1 |
| TAGLN | 10 | 1 | 0 | 1 | 1 |
| PLEC | 194 | 194 | -0,06996064 | 0,247113635 | 1 |
| CLTA | 1 | 1 | 0 | 1 | 1 |
| MTHFD1 | 34 | 34 | 3,13163E-18 | 0,999999995 | 1 |
| MAP7D1 | 15 | 15 | -0,006503439 | 0,737964376 | 1 |
| CENPE | 1 | 1 | 0 | 1 | 1 |
| MARCHF6 | 1 | 1 | 0 | 1 | 1 |
| ACACA | 27 | 27 | 0 | 1 | 1 |
| MAP4 | 24 | 24 | -1,81836E-17 | 0,999999997 | 1 |
| CNNM1 | 16 | 16 | 2,18998E-16 | 0,999999943 | 1 |
| RELCH | 2 | 2 | 0 | 1 | 1 |
| SPTB | 51 | 51 | -0,107102676 | 0,136324638 | 1 |
| PIK3CB | 1 | 1 | 0 | 1 | 1 |
| PHB1 | 22 | 22 | -5,75313E-18 | 0,999999995 | 1 |
| PPP1R16B | 1 | 1 | -0,00376085 | 0,863746611 | 1 |
| 2210016F16RIK | 8 | 2 | -1,70203E-13 | 0,999998971 | 1 |
| SPTBN2 | 114 | 113 | -0,038317095 | 0,532767119 | 1 |

| | | | | | |
|---|---|---|---|---|---|
| PDE2A | 39 | 39 | 0,01251386 | 0,521336987 | 1 |
| ECI3;ECI2 | 4 | 4 | -0,031385203 | 0,485518346 | 1 |
| FKBP4 | 32 | 29 | 0 | 1 | 1 |
| PCDHB14;PCDHB6 | 1 | 1 | 0 | 1 | 1 |
| KSR1 | 20 | 20 | 6,9718E-16 | 0,999999907 | 1 |
| NEMF | 4 | 4 | -1,62835E-18 | 1 | 1 |
| TXNDC15 | 2 | 2 | -2,05273E-16 | 0,999999989 | 1 |
| CLTC | 130 | 129 | 0 | 1 | 1 |
| COPB2 | 16 | 16 | -2,73391E-13 | 0,999998499 | 1 |
| NDUFA7 | 16 | 16 | -1,71143E-16 | 0,999999973 | 1 |
| IQSEC3 | 16 | 16 | 0,013379982 | 0,523870728 | 1 |
| MRPS35 | 6 | 6 | -1,34231E-15 | 0,999999924 | 1 |
| RFTN1 | 5 | 5 | 3,78314E-17 | 0,99999999 | 1 |
| TOM1L2 | 23 | 23 | 4,09781E-17 | 0,999999989 | 1 |
| ABCB10 | 10 | 10 | -0,010574104 | 0,718948794 | 1 |
| MAP4 | 21 | 21 | -0,079483684 | 0,160670783 | 1 |
| CLIP2 | 42 | 42 | -7,50068E-14 | 0,999998965 | 1 |
| TAGLN3 | 18 | 18 | 0 | 1 | 1 |
| YWHAE | 35 | 35 | 0,067861788 | 0,150410623 | 1 |
| GRM2 | 15 | 15 | -0,003775157 | 0,807529472 | 1 |
| SDHA | 29 | 28 | -1,02415E-16 | 0,999999973 | 1 |
| ITGA3 | 1 | 1 | 0 | 1 | 1 |
| VPS51 | 32 | 32 | 0,034458871 | 0,275741049 | 1 |
| METTL13 | 3 | 3 | 0 | 1 | 1 |
| ALDH2 | 27 | 27 | -0,084324194 | 0,311890642 | 1 |
| COMMD3 | 3 | 3 | 0,003629489 | 0,818194343 | 1 |
| NWD2 | 23 | 23 | 0,027897384 | 0,14762283 | 1 |
| CCDC124 | 3 | 3 | -1,21534E-17 | 0,999999993 | 1 |
| CTSA | 6 | 4 | 1,24104E-17 | 0,999999994 | 1 |
| SRM | 12 | 11 | -6,74182E-17 | 0,999999974 | 1 |
| IQGAP2 | 45 | 45 | -0,029016988 | 0,355724763 | 1 |
| CNP | 34 | 34 | -0,092343868 | 0,273853653 | 1 |
| FRMD4A | 6 | 6 | 0 | 1 | 1 |

| | | | | | |
|---|---|---|---|---|---|
| PPP1R9B | 35 | 35 | -6,57247E-15 | 0,999999764 | 1 |
| NNT | 30 | 30 | 1,55701E-16 | 0,999999985 | 1 |
| DCXR | 3 | 3 | 0 | 1 | 1 |
| BAG6 | 22 | 20 | -9,00917E-15 | 0,999999573 | 1 |
| BCL2L2 | 3 | 3 | -8,43832E-16 | 0,999999938 | 1 |
| ATP6V1D | 21 | 20 | 0,014140478 | 0,560439049 | 1 |
| GART | 26 | 26 | 2,05131E-16 | 0,999999956 | 1 |
| OTUD7A | 4 | 4 | 0 | 1 | 1 |
| FBXO41 | 18 | 18 | 5,86561E-15 | 0,999999695 | 1 |
| RAP1GAP | 7 | 7 | 0,058225883 | 0,153126315 | 1 |
| RAP1GAP | 16 | 16 | -5,0885E-18 | 0,999999995 | 1 |
| PICALM | 6 | 5 | 0 | 1 | 1 |
| IGBP1 | 4 | 4 | -3,28728E-16 | 0,999999944 | 1 |
| AJM1 | 20 | 20 | -3,82614E-17 | 0,999999985 | 1 |
| CTNND2 | 40 | 39 | -8,26912E-16 | 0,999999915 | 1 |
| OPA1 | 62 | 62 | -3,44588E-17 | 0,999999988 | 1 |
| MACROD1 | 4 | 4 | -3,34968E-19 | 0,999999999 | 1 |
| JAGN1 | 2 | 2 | 9,06215E-16 | 0,999999942 | 1 |
| CNOT3 | 3 | 3 | -0,045827117 | 0,436491509 | 1 |
| ITPKA | 27 | 26 | 7,216E-17 | 0,99999998 | 1 |
| OXCT1 | 31 | 31 | -0,051610888 | 0,402634527 | 1 |
| RUVBL2 | 21 | 19 | 2,65714E-16 | 0,999999967 | 1 |
| PCBD2 | 2 | 2 | 1,94101E-15 | 0,999999866 | 1 |
| DUT | 2 | 2 | 1,08424E-18 | 0,999999998 | 1 |
| RAB11B | 3 | 3 | 2,31482E-18 | 0,999999997 | 1 |
| NCAM1 | 35 | 35 | 1,07988E-13 | 0,999998675 | 1 |
| IQSEC2 | 32 | 32 | -8,65966E-17 | 0,999999965 | 1 |
| CCT2 | 47 | 46 | 0,042739165 | 0,150455371 | 1 |
| STXBP1 | 60 | 59 | 0,04050133 | 0,303817876 | 1 |
| ANKIB1 | 2 | 2 | 3,63258E-18 | 0,999999997 | 1 |
| FUS | 9 | 5 | -1,03503E-18 | 0,999999999 | 1 |
| MAT2A | 20 | 18 | -1,9425E-17 | 0,999999989 | 1 |
| TNR | 42 | 39 | 0 | 1 | 1 |

| | | | | | |
|---|---|---|---|---|---|
| TDRKH | 15 | 15 | -0,005446649 | 0,788053913 | 1 |
| NPRL3 | 2 | 2 | 3,29525E-17 | 0,999999989 | 1 |
| PRMT5 | 11 | 11 | 1,33409E-17 | 0,999999997 | 1 |
| ERC2;ERC1 | 14 | 14 | 0 | 1 | 1 |
| TTC39B | 3 | 3 | -5,84864E-19 | 0,999999999 | 1 |
| MIEF1 | 1 | 1 | -0,022146242 | 0,627507796 | 1 |
| EMC1 | 22 | 21 | -0,025371154 | 0,21238188 | 1 |
| EXOC3 | 13 | 13 | 0,029769012 | 0,227865103 | 1 |
| RYR2 | 6 | 6 | -7,12838E-17 | 0,99999998 | 1 |
| ACP6 | 6 | 6 | 0 | 1 | 1 |
| GPR158 | 36 | 35 | 0,009878013 | 0,441768232 | 1 |
| SLC27A1;SLC27A4 | 2 | 2 | -1,76979E-18 | 0,999999998 | 1 |
| ITPR1 | 39 | 39 | 0 | 1 | 1 |
| NCAN | 34 | 24 | 0 | 1 | 1 |
| DPYSL3 | 31 | 29 | 0,007743921 | 0,712888531 | 1 |
| CRMP1 | 31 | 31 | 0,005360528 | 0,747917665 | 1 |
| DPYSL2 | 44 | 44 | 0 | 1 | 1 |
| CARS2 | 6 | 6 | 6,92322E-17 | 0,999999988 | 1 |
| SHANK3 | 46 | 46 | 6,46029E-17 | 0,99999997 | 1 |
| RYR2 | 77 | 76 | -3,56709E-15 | 0,999999768 | 1 |
| KIF1A | 49 | 47 | 0 | 1 | 1 |
| QDPR | 12 | 12 | -1,2531E-17 | 0,999999991 | 1 |
| CELF2 | 10 | 7 | 0,025619528 | 0,526669612 | 1 |
| CELF1 | 5 | 2 | 0 | 1 | 1 |
| HARS1 | 16 | 16 | -8,18509E-17 | 0,999999972 | 1 |
| APLP1 | 12 | 9 | -0,086850161 | 0,145753796 | 1 |
| EEA1 | 52 | 49 | -0,039583688 | 0,137646409 | 1 |
| CPLX2 | 6 | 6 | 5,92473E-17 | 0,999999981 | 1 |
| MPP2 | 22 | 22 | 0 | 1 | 1 |
| MTOR | 45 | 44 | 5,77409E-18 | 0,999999998 | 1 |
| PRXL2A | 9 | 9 | -1,41533E-17 | 0,999999989 | 1 |
| PTPRF;PTPRD | 5 | 5 | -1,73873E-18 | 0,999999997 | 1 |
| CCDC43 | 5 | 4 | 0 | 1 | 1 |

| | | | | | |
|---|---|---|---|---|---|
| WASL | 10 | 10 | 0 | 1 | 1 |
| UBE2O | 36 | 35 | -1,66578E-15 | 0,999999862 | 1 |
| EPRS1 | 53 | 45 | -0,010430347 | 0,428388724 | 1 |
| ELAC2 | 5 | 3 | -0,141115454 | 0,130498077 | 1 |
| VMA21 | 1 | 1 | 0 | 1 | 1 |
| AMPH | 21 | 21 | 1,74796E-17 | 0,999999983 | 1 |
| HUWE1 | 54 | 48 | -5,79974E-17 | 0,99999997 | 1 |
| HTT | 51 | 51 | 0 | 1 | 1 |
| PTPN23 | 20 | 20 | 0 | 1 | 1 |
| FOXO3 | 1 | 1 | 0 | 1 | 1 |
| CHMP4B | 9 | 8 | 0 | 1 | 1 |
| APOB | 1 | 1 | 0 | 1 | 1 |
| KIF2A | 29 | 29 | 1,84826E-14 | 0,99999947 | 1 |
| RPS25 | 5 | 5 | 0,028760898 | 0,521860511 | 1 |
| ASPSCR1 | 9 | 9 | 1,43327E-17 | 0,999999988 | 1 |
| WDR41 | 6 | 6 | -7,40335E-16 | 0,999999924 | 1 |
| PITPNM1 | 25 | 24 | -1,35674E-16 | 0,99999996 | 1 |
| PPP5C | 25 | 21 | 0,054321837 | 0,137552249 | 1 |
| DFFA | 1 | 1 | 0,039108913 | 0,696542164 | 1 |
| LRP1 | 52 | 52 | -0,03186027 | 0,169153262 | 1 |
| HADHB | 26 | 26 | -1,00748E-18 | 0,999999998 | 1 |
| XPO7 | 34 | 33 | 0 | 1 | 1 |
| MAST1 | 13 | 13 | 0 | 1 | 1 |
| COPS2 | 31 | 31 | 0 | 1 | 1 |
| EIF4G1 | 22 | 21 | 0 | 1 | 1 |
| ISCA1 | 4 | 4 | -0,046904102 | 0,480507937 | 1 |
| SRCIN1 | 68 | 68 | -0,013288381 | 0,533033123 | 1 |
| TUBB6 | 7 | 7 | -1,08887E-19 | 1 | 1 |
| SLC25A27 | 10 | 10 | -1,04601E-17 | 0,999999991 | 1 |
| PRMT8 | 10 | 9 | 0 | 1 | 1 |
| ELP1 | 15 | 15 | -0,003812003 | 0,716508296 | 1 |
| TLN1 | 40 | 40 | 0 | 1 | 1 |
| ILVBL | 5 | 5 | 0,023339255 | 0,584473985 | 1 |

| | | | | | |
|---|---|---|---|---|---|
| JUP | 16 | 16 | 3,1518E-17 | 0,999999992 | 1 |
| AP1B1;AP2B1 | 33 | 33 | 0,033080193 | 0,24737452 | 1 |
| FASN | 83 | 83 | 1,34401E-18 | 0,999999998 | 1 |
| SPTBN1 | 208 | 205 | -0,032305259 | 0,385653876 | 1 |
| CPT1C | 11 | 11 | -8,91541E-17 | 0,999999983 | 1 |
| WARS1 | 33 | 32 | 0,037329282 | 0,183886383 | 1 |
| ACAA2 | 27 | 27 | -0,061696837 | 0,402608308 | 1 |
| LSAMP | 14 | 14 | 2,77594E-17 | 0,999999982 | 1 |
| VCL | 42 | 39 | -0,030922465 | 0,302689744 | 1 |
| CAMK4 | 13 | 13 | -4,28194E-17 | 0,999999989 | 1 |
| PSMA2 | 12 | 12 | 0,067844369 | 0,202920293 | 1 |
| ITGB8 | 8 | 8 | -0,009742786 | 0,623026626 | 1 |
| WIPI2 | 6 | 6 | 0 | 1 | 1 |
| CYC1 | 17 | 17 | -1,03217E-16 | 0,999999983 | 1 |
| THUMPD3 | 3 | 3 | 0 | 1 | 1 |
| HYOU1 | 36 | 36 | -0,048384267 | 0,120386285 | 1 |
| STK24 | 8 | 8 | -1,04685E-14 | 0,999999806 | 1 |
| STK25 | 4 | 4 | 0 | 1 | 1 |
| DLAT | 32 | 32 | 0 | 1 | 1 |
| GAMT | 5 | 5 | 0,025755301 | 0,514373102 | 1 |
| PRKCG | 50 | 48 | 0,072363986 | 0,136330081 | 1 |
| MTUS2 | 5 | 5 | -8,09573E-16 | 0,999999909 | 1 |
| NPTN | 10 | 10 | 0,062521431 | 0,269648247 | 1 |
| PPA1 | 19 | 17 | -8,75455E-17 | 0,999999981 | 1 |
| HINT2 | 5 | 5 | -1,76881E-17 | 0,999999992 | 1 |
| PHOSPHO1 | 4 | 4 | 0 | 1 | 1 |
| PRRT3 | 14 | 14 | 1,14989E-15 | 0,999999889 | 1 |
| PYGB;PYGL | 11 | 11 | -1,43614E-17 | 0,999999988 | 1 |
| MRPL37 | 9 | 9 | -7,73637E-18 | 0,999999993 | 1 |
| SV2A | 22 | 22 | 1,24504E-16 | 0,999999959 | 1 |
| PDK3 | 17 | 17 | -3,67143E-16 | 0,999999958 | 1 |
| UBR4 | 61 | 61 | -0,000577497 | 0,859481954 | 1 |
| SPON1 | 7 | 5 | -0,035357935 | 0,518347743 | 1 |

| | | | | | |
|---|---|---|---|---|---|
| TKFC | 7 | 6 | -8,53095E-17 | 0,999999976 | 1 |
| NFASC | 45 | 44 | 1,07253E-16 | 0,999999975 | 1 |
| CRYZ | 13 | 13 | 1,14133E-17 | 0,999999993 | 1 |
| ZNRD2 | 2 | 2 | 0,107041872 | 0,180690764 | 1 |
| TTC9B | 5 | 5 | 0,05898774 | 0,256165693 | 1 |
| HK1 | 61 | 61 | 0 | 1 | 1 |
| ACAD8 | 16 | 16 | -0,054083153 | 0,235372037 | 1 |
| GAP43 | 18 | 18 | -3,04271E-16 | 0,999999952 | 1 |
| CSNK1A1 | 1 | 1 | 0,064959324 | 0,393307926 | 1 |
| CSNK1A1 | 1 | 1 | 0 | 1 | 1 |
| SORBS1 | 2 | 2 | 0 | 1 | 1 |
| NSF | 72 | 70 | 0,043185377 | 0,157218424 | 1 |
| VPS53 | 17 | 17 | 0,036549127 | 0,154803595 | 1 |
| TBCD | 16 | 13 | -0,02631089 | 0,390553649 | 1 |
| AGPS | 7 | 7 | -5,99031E-18 | 0,999999997 | 1 |
| SHANK1 | 52 | 52 | 0 | 1 | 1 |
| ACOT7 | 26 | 24 | 0 | 1 | 1 |
| PRKAR2B | 15 | 15 | -1,20234E-17 | 0,999999998 | 1 |
| TUBGCP2 | 9 | 9 | 0 | 1 | 1 |
| CLSTN1 | 12 | 10 | 0 | 1 | 1 |
| ITSN1 | 34 | 33 | -1,52823E-17 | 0,999999985 | 1 |
| DMAC2L | 7 | 7 | 0 | 1 | 1 |
| ANXA4 | 6 | 6 | -3,70118E-19 | 0,999999999 | 1 |
| TAOK2 | 11 | 11 | -0,042886712 | 0,128096391 | 1 |
| IAH1 | 2 | 2 | -0,013470447 | 0,710104628 | 1 |
| COX5B | 8 | 7 | 0 | 1 | 1 |
| DLG1 | 29 | 29 | 0 | 1 | 1 |
| ATXN2 | 8 | 7 | 0 | 1 | 1 |
| EIF4B | 14 | 14 | 1,47535E-16 | 0,999999966 | 1 |
| ACSBG1 | 34 | 34 | 0 | 1 | 1 |
| SDHD | 1 | 1 | 0 | 1 | 1 |
| BEGAIN | 18 | 18 | 4,84887E-17 | 0,999999976 | 1 |
| STRIP1 | 6 | 6 | 0 | 1 | 1 |

| | | | | | |
|---|---|---|---|---|---|
| SH3PXD2A | 6 | 6 | 0 | 1 | 1 |
| SEL1L | 6 | 6 | -0,03961373 | 0,282564956 | 1 |
| ARHGAP21 | 25 | 25 | -9,53368E-17 | 0,999999974 | 1 |
| LXN | 6 | 6 | -7,93535E-16 | 0,99999992 | 1 |
| SHANK2 | 47 | 47 | -1,394E-17 | 0,999999997 | 1 |
| GEMIN5 | 7 | 4 | 1,0172E-17 | 0,999999998 | 1 |
| STRAP | 19 | 19 | -3,13648E-16 | 0,999999946 | 1 |
| PDIA6 | 9 | 9 | -3,19597E-17 | 0,999999988 | 1 |
| LTBP2 | 1 | 1 | -0,03092408 | 0,759630162 | 1 |
| GPAM | 1 | 1 | 0,010309451 | 0,817042427 | 1 |
| OGT | 37 | 37 | 0,030987201 | 0,17215623 | 1 |
| MCCC2 | 21 | 21 | -6,61934E-17 | 0,999999987 | 1 |
| TMOD2 | 20 | 20 | -8,06704E-17 | 0,999999966 | 1 |
| RUSF1 | 3 | 3 | -1,8084E-15 | 0,999999889 | 1 |
| PRKAR2A | 16 | 16 | -4,23065E-17 | 0,999999981 | 1 |
| ADGRB3 | 16 | 16 | 8,07641E-16 | 0,999999881 | 1 |
| MYO5A | 90 | 90 | -4,82968E-17 | 0,999999982 | 1 |
| ADGRL3 | 25 | 25 | 0 | 1 | 1 |
| PYCR3 | 6 | 6 | 0,058574708 | 0,247021436 | 1 |
| DCUN1D2;DCUN1D1 | 2 | 2 | 0,00054304 | 0,952309717 | 1 |
| GUCY1A2 | 9 | 9 | 0 | 1 | 1 |
| GALK2 | 8 | 5 | 0 | 1 | 1 |
| DYNC1H1 | 275 | 266 | -1,32623E-16 | 0,999999963 | 1 |
| NCAM1 | 15 | 15 | -0,170286886 | 0,135607307 | 1 |
| BCAR1 | 5 | 5 | 0 | 1 | 1 |
| ATAT1 | 8 | 8 | 0 | 1 | 1 |
| UBA1 | 60 | 59 | 0 | 1 | 1 |
| HSD17B4 | 17 | 17 | -5,08207E-17 | 0,999999988 | 1 |
| ADH5 | 12 | 12 | 5,18873E-17 | 0,999999977 | 1 |
| ETFA | 18 | 18 | -0,07674641 | 0,303447644 | 1 |
| HSPD1 | 52 | 52 | -1,56324E-17 | 0,999999992 | 1 |
| MICAL1 | 4 | 4 | -1,41579E-17 | 0,999999991 | 1 |
| MTHFD1L | 39 | 39 | -1,71254E-15 | 0,999999881 | 1 |

| | | | | | |
|---|---|---|---|---|---|
| PEX5L | 7 | 7 | 4,83005E-17 | 0,999999982 | 1 |
| PSMD1 | 42 | 39 | 2,50571E-16 | 0,999999942 | 1 |
| EWSR1 | 5 | 3 | 0,108020859 | 0,35017722 | 1 |
| EDC3 | 2 | 2 | -3,89767E-18 | 0,999999998 | 1 |
| C3 | 18 | 3 | 3,09968E-20 | 1 | 1 |
| APIP | 5 | 3 | -0,105158123 | 0,212309124 | 1 |
| CTNNB1 | 27 | 27 | 0 | 1 | 1 |
| ENAH | 13 | 12 | -0,068629358 | 0,146080047 | 1 |
| ENO2;ENO3 | 2 | 2 | 0 | 1 | 1 |
| ENO1 | 33 | 33 | 0 | 1 | 1 |
| ENO1;ENO2;ENO3 | 3 | 3 | 0 | 1 | 1 |
| CBR3 | 15 | 15 | -0,049564682 | 0,185775302 | 1 |
| MMUT | 21 | 21 | -0,028933651 | 0,555829519 | 1 |
| RAP1GAP2 | 6 | 6 | 0 | 1 | 1 |
| ADD1 | 34 | 34 | -0,026851007 | 0,298190383 | 1 |
| ATRX | 1 | 1 | 0 | 1 | 1 |
| OPA3 | 3 | 3 | -1,92722E-17 | 0,999999994 | 1 |
| WDR47 | 31 | 31 | 3,21821E-16 | 0,99999992 | 1 |
| IBA57 | 6 | 6 | -3,96892E-16 | 0,99999997 | 1 |
| ALDH5A1 | 28 | 28 | -0,09769441 | 0,196884467 | 1 |
| RALBP1 | 6 | 6 | 1,99172E-16 | 0,999999971 | 1 |
| LMNA | 31 | 10 | 7,23793E-16 | 0,999999957 | 1 |
| SLC25A5 | 12 | 12 | 1,87385E-17 | 0,999999989 | 1 |
| SLC25A4 | 18 | 17 | 1,26603E-16 | 0,999999984 | 1 |
| MYO15A | 2 | 2 | 0 | 1 | 1 |
| NDUFV1 | 26 | 26 | -5,38531E-16 | 0,999999951 | 1 |
| RABGGTA | 20 | 19 | 7,19278E-17 | 0,999999983 | 1 |
| COTL1 | 16 | 15 | 0 | 1 | 1 |
| RARS1 | 33 | 29 | 9,84393E-18 | 0,999999988 | 1 |
| JAKMIP1 | 2 | 2 | 0 | 1 | 1 |
| RIDA | 9 | 9 | -8,55121E-15 | 0,99999978 | 1 |
| USP15;USP11 | 1 | 1 | 0 | 1 | 1 |
| GNB1 | 15 | 14 | 0,105577784 | 0,247502137 | 1 |

| | | | | | |
|---|---|---|---|---|---|
| IVD | 17 | 17 | -0,133158416 | 0,118562479 | 1 |
| NUB1 | 3 | 2 | 5,15342E-16 | 0,999999954 | 1 |
| L2HGDH | 18 | 18 | -4,73284E-16 | 0,999999936 | 1 |
| GNB2 | 9 | 8 | 0,048893118 | 0,539767282 | 1 |
| ARRB1 | 16 | 16 | 0 | 1 | 1 |
| IQSEC1 | 43 | 43 | 4,72995E-16 | 0,999999894 | 1 |
| SLC39A6 | 5 | 5 | 0 | 1 | 1 |
| KBTBD11 | 18 | 17 | -3,18946E-16 | 0,999999964 | 1 |
| GLUL | 25 | 23 | -3,68061E-17 | 0,999999988 | 1 |
| RPS3A | 19 | 16 | 0,043852405 | 0,519168662 | 1 |
| NISCH | 15 | 15 | -2,09532E-17 | 0,999999986 | 1 |
| ATP1A3 | 49 | 49 | 0,036115536 | 0,195594533 | 1 |
| ATP1A1 | 43 | 43 | 2,86739E-17 | 0,999999985 | 1 |
| ATP1A2 | 56 | 56 | 0,008507898 | 0,644345996 | 1 |
| ACTR1A;ACTR1B | 12 | 12 | 1,22561E-17 | 0,999999985 | 1 |
| JPH1 | 2 | 2 | -0,048394624 | 0,309830349 | 1 |
| SACS | 12 | 12 | 0 | 1 | 1 |
| ERLIN2 | 15 | 15 | -0,007247327 | 0,636034951 | 1 |
| DYNC1I2 | 9 | 9 | -0,024519677 | 0,348602673 | 1 |
| LMBRD2 | 7 | 7 | -2,32956E-17 | 0,999999984 | 1 |
| CACNA2D2 | 11 | 11 | -0,008434999 | 0,65118932 | 1 |
| ANXA5 | 25 | 22 | 0 | 1 | 1 |
| ERLIN1 | 2 | 2 | 2,27632E-17 | 0,999999998 | 1 |
| CASK | 21 | 21 | 0 | 1 | 1 |
| ATG3 | 6 | 6 | 0 | 1 | 1 |
| ACOT2 | 8 | 8 | 2,02815E-17 | 0,999999994 | 1 |
| SNTA1 | 8 | 7 | 0 | 1 | 1 |
| CNTN3 | 7 | 7 | 0 | 1 | 1 |
| GABRG2 | 9 | 9 | 2,8627E-16 | 0,999999942 | 1 |
| VAMP1 | 6 | 6 | 0,109882425 | 0,199581219 | 1 |
| VAMP3 | 6 | 6 | -4,85709E-17 | 0,999999987 | 1 |
| ARHGAP44 | 15 | 15 | 1,33745E-17 | 0,99999999 | 1 |
| CCDC115 | 1 | 1 | 0 | 1 | 1 |

| | | | | | |
|---|---|---|---|---|---|
| GORASP2 | 9 | 8 | 2,22175E-17 | 0,999999989 | 1 |
| AHSA1 | 25 | 24 | 6,87395E-12 | 0,999988122 | 1 |
| ADAM22 | 25 | 25 | 1,56292E-17 | 0,999999992 | 1 |
| TUFM | 34 | 34 | -6,44321E-16 | 0,999999948 | 1 |
| PPA2 | 20 | 20 | -3,07281E-17 | 0,999999988 | 1 |
| SEPTIN5 | 29 | 28 | 0,032780617 | 0,299347034 | 1 |
| TLN2 | 7 | 7 | 9,06397E-19 | 1 | 1 |
| ALDOART2 | 14 | 14 | 1,25715E-17 | 0,99999999 | 1 |
| MICU3 | 21 | 21 | -7,41146E-18 | 0,999999996 | 1 |
| MAP2K6 | 5 | 5 | 1,53468E-17 | 0,999999999 | 1 |
| DGKG | 16 | 15 | 5,52587E-17 | 0,99999998 | 1 |
| PC | 57 | 57 | -1,65184E-15 | 0,999999923 | 1 |
| COX10 | 1 | 1 | -7,44887E-22 | 1 | 1 |
| PITPNB | 12 | 11 | -5,42723E-18 | 0,999999996 | 1 |
| NUTF2 | 8 | 7 | 5,08776E-17 | 0,999999993 | 1 |
| KCNC1 | 6 | 5 | 0,066664343 | 0,18566914 | 1 |
| PRDX1 | 20 | 20 | 3,30167E-16 | 0,999999965 | 1 |
| BIN1 | 33 | 33 | -1,61282E-16 | 0,999999947 | 1 |
| PALM | 15 | 15 | 0 | 1 | 1 |
| MYH9 | 99 | 99 | 0,017325824 | 0,603455215 | 1 |
| GSR | 10 | 10 | 2,61523E-17 | 0,999999992 | 1 |
| NIF3L1 | 11 | 11 | 6,17261E-17 | 0,999999972 | 1 |
| DAD1 | 2 | 2 | 0,001409896 | 0,907512614 | 1 |
| TTYH1 | 6 | 4 | 0 | 1 | 1 |
| OCIAD1 | 10 | 10 | -1,43845E-17 | 0,999999994 | 1 |
| SERPINB6 | 25 | 23 | 1,01573E-17 | 0,999999989 | 1 |
| FKBP8 | 10 | 10 | -7,36089E-19 | 1 | 1 |
| ICAM1 | 1 | 1 | 0 | 1 | 1 |
| SPHK2 | 6 | 6 | 0,054746513 | 0,190147773 | 1 |
| SMS | 16 | 13 | -3,57995E-17 | 0,999999984 | 1 |
| COX7A2L | 5 | 5 | -0,034062989 | 0,454257382 | 1 |
| IPCEF1 | 2 | 2 | -1,03408E-16 | 0,999999977 | 1 |
| FUOM | 3 | 3 | -6,03473E-24 | 1 | 1 |

| | | | | | |
|---|---|---|---|---|---|
| DLD | 22 | 22 | -6,60173E-17 | 0,999999985 | 1 |
| MBLAC1 | 3 | 2 | -0,086475859 | 0,473088352 | 1 |
| CAMK2B | 3 | 3 | 0 | 1 | 1 |
| PSMC6 | 23 | 20 | 3,73684E-16 | 0,999999977 | 1 |
| HSP90AB1 | 36 | 32 | 0,045232725 | 0,148156442 | 1 |
| RPSA | 11 | 11 | -2,97218E-17 | 0,999999994 | 1 |
| RAB11FIP5 | 16 | 16 | -0,067634961 | 0,161319707 | 1 |
| GRIA3 | 23 | 23 | 0 | 1 | 1 |
| OTUD4 | 2 | 2 | 0 | 1 | 1 |
| TM9SF2 | 7 | 7 | 7,02276E-18 | 0,999999997 | 1 |
| ABCE1 | 14 | 11 | 0 | 1 | 1 |
| ATP5F1C | 20 | 20 | -1,58771E-16 | 0,999999973 | 1 |
| ASPA | 9 | 8 | -0,094233077 | 0,182261256 | 1 |
| HNRNPM | 4 | 4 | 1,00139E-15 | 0,999999942 | 1 |
| PAIP1 | 6 | 6 | -5,93819E-17 | 0,999999978 | 1 |
| MRPS23 | 7 | 7 | -0,049458418 | 0,429861084 | 1 |
| GLTP | 4 | 4 | -0,068833815 | 0,325046869 | 1 |
| GSTM1 | 28 | 25 | -0,000398874 | 0,935801334 | 1 |
| BRSK2 | 14 | 14 | 0 | 1 | 1 |
| RAB28 | 4 | 3 | 3,60367E-16 | 0,999999976 | 1 |
| MARS1 | 32 | 26 | -0,020901917 | 0,350480475 | 1 |
| SLC30A9 | 13 | 13 | -0,002119886 | 0,86071956 | 1 |
| ENPP6 | 11 | 11 | -7,73415E-17 | 0,99999998 | 1 |
| UBE2L3 | 8 | 8 | 0 | 1 | 1 |
| MPDZ | 10 | 10 | -0,038479426 | 0,22055584 | 1 |
| SPATA2L | 3 | 3 | -3,00936E-17 | 0,999999987 | 1 |
| GRIK2 | 3 | 3 | 3,57066E-17 | 0,999999994 | 1 |
| OXR1 | 24 | 23 | 0 | 1 | 1 |
| SORT1 | 9 | 9 | 3,36626E-06 | 0,992266271 | 1 |
| CYCS | 14 | 14 | -0,033871952 | 0,499782879 | 1 |
| HECTD4 | 28 | 28 | -6,0904E-16 | 0,999999901 | 1 |
| MLYCD | 8 | 8 | 0 | 1 | 1 |
| RAB27B | 8 | 8 | 0,047462119 | 0,188970497 | 1 |

| | | | | | |
|---|---|---|---|---|---|
| SORBS2 | 11 | 11 | -0,073295896 | 0,159215408 | 1 |
| ADSL | 13 | 13 | 0 | 1 | 1 |
| PACS1 | 31 | 31 | 0,010520621 | 0,556345753 | 1 |
| SERPINB1A | 21 | 21 | -4,33326E-17 | 0,999999995 | 1 |
| FLII | 8 | 8 | -1,27963E-17 | 0,999999994 | 1 |
| EIF2B1 | 5 | 5 | 0 | 1 | 1 |
| NIT2 | 14 | 14 | -9,91633E-17 | 0,99999998 | 1 |
| APLP2 | 9 | 9 | -0,024214915 | 0,371388169 | 1 |
| TARS1 | 35 | 31 | 1,66246E-17 | 0,999999984 | 1 |
| BSN | 157 | 157 | -8,57828E-16 | 0,999999925 | 1 |
| HADHA | 36 | 36 | -0,033956213 | 0,502682357 | 1 |
| MYH10 | 106 | 105 | -0,029424555 | 0,344525545 | 1 |
| ABCB7 | 18 | 17 | -3,51136E-16 | 0,999999958 | 1 |
| PTEN | 9 | 9 | 1,56587E-17 | 0,999999992 | 1 |
| PGM1 | 43 | 43 | -7,02157E-16 | 0,999999937 | 1 |
| ISYNA1 | 12 | 11 | 5,96516E-17 | 0,999999981 | 1 |
| GATB | 4 | 4 | -2,53773E-17 | 0,999999995 | 1 |
| UBA2 | 23 | 1 | 0 | 1 | 1 |
| RUFY3 | 1 | 1 | 0,014673107 | 0,779569119 | 1 |
| ALDH1L1 | 44 | 41 | -5,07531E-16 | 0,999999931 | 1 |
| NCS1 | 11 | 11 | 0,076856668 | 0,176961046 | 1 |
| RPS6KA2 | 10 | 7 | 4,61149E-20 | 1 | 1 |
| USP47 | 26 | 23 | -0,040059233 | 0,21510089 | 1 |
| SLC30A1 | 8 | 8 | 0,01320903 | 0,550731568 | 1 |
| POLR2H | 7 | 5 | -0,050515855 | 0,393399598 | 1 |
| MAP2 | 103 | 93 | -0,029928676 | 0,279428151 | 1 |
| SNAP25 | 26 | 25 | 1,11032E-16 | 0,999999964 | 1 |
| EHD4 | 11 | 11 | 6,75899E-17 | 0,999999976 | 1 |
| RPS14 | 8 | 8 | 0,002456789 | 0,909213808 | 1 |
| TF | 38 | 24 | 0,003274903 | 0,832871131 | 1 |
| ATAD3 | 27 | 27 | -0,020046912 | 0,662573692 | 1 |
| GNB5 | 10 | 10 | 0,066615316 | 0,255700674 | 1 |
| ACE | 12 | 9 | -0,000272809 | 0,967528419 | 1 |

| | | | | | |
|---|---|---|---|---|---|
| SRGAP1 | 4 | 4 | 1,98834E-18 | 0,999999998 | 1 |
| DLGAP2 | 20 | 19 | 1,89576E-16 | 0,999999946 | 1 |
| ZNF365 | 2 | 2 | -0,018188704 | 0,679040645 | 1 |
| NEO1 | 21 | 21 | 0 | 1 | 1 |
| PACSIN3 | 7 | 7 | 0 | 1 | 1 |
| TBC1D9B | 10 | 10 | 0 | 1 | 1 |
| PTPRN2 | 24 | 24 | 0 | 1 | 1 |
| BCAS1 | 16 | 16 | -0,115146581 | 0,16378205 | 1 |
| LNPK | 12 | 12 | -0,020187061 | 0,505976393 | 1 |
| MAGED1 | 4 | 4 | -2,95591E-17 | 0,999999989 | 1 |
| PSMC4 | 18 | 17 | -0,021548573 | 0,314171712 | 1 |
| SEPTIN9 | 30 | 30 | 0,005860469 | 0,687855345 | 1 |
| SEPTIN7 | 38 | 38 | 0,036690989 | 0,141255928 | 1 |
| SEPTIN4 | 13 | 13 | -0,002224057 | 0,808690682 | 1 |
| SEPTIN3 | 26 | 26 | 1,03015E-16 | 0,999999958 | 1 |
| ERH | 1 | 1 | 0 | 1 | 1 |
| NUFIP2 | 2 | 2 | 0 | 1 | 1 |
| GRIA1 | 37 | 37 | 0,045871497 | 0,151756551 | 1 |
| ATP2B1 | 62 | 62 | 0,022223828 | 0,390289128 | 1 |
| BTBD17 | 11 | 11 | 0 | 1 | 1 |
| GPHN | 25 | 25 | 2,41529E-16 | 0,999999956 | 1 |
| AMACR | 8 | 8 | -0,067392037 | 0,147956825 | 1 |
| SPTAN1 | 235 | 234 | -0,036128933 | 0,391715673 | 1 |
| SPTA1 | 2 | 2 | 0 | 1 | 1 |
| COX4I1 | 17 | 17 | 2,10633E-17 | 0,999999991 | 1 |
| ATP6V1A | 58 | 58 | 0,064841935 | 0,123252462 | 1 |
| MTMR7 | 6 | 6 | 1,40612E-13 | 0,999998909 | 1 |
| FNBP1 | 9 | 9 | -6,98588E-17 | 0,999999971 | 1 |
| ADCY2 | 8 | 8 | -1,99556E-16 | 0,999999953 | 1 |
| AGAP2 | 31 | 31 | -4,74626E-17 | 0,99999998 | 1 |
| JPH3 | 4 | 4 | -0,014085349 | 0,566510579 | 1 |
| DCUN1D3 | 3 | 3 | 2,23739E-19 | 1 | 1 |
| DCPS | 15 | 3 | 0,059316998 | 0,449278839 | 1 |

| | | | | | |
|---|---|---|---|---|---|
| KRT76 | 4 | 3 | -1,01449E-16 | 0,999999993 | 1 |
| TOMM22 | 6 | 6 | -1,05396E-17 | 0,999999992 | 1 |
| ASL | 16 | 16 | 0,01626182 | 0,519187315 | 1 |
| PAPSS2 | 2 | 2 | -0,06901069 | 0,308625545 | 1 |
| RPS10 | 9 | 7 | 0 | 1 | 1 |
| PDXK | 16 | 15 | -1,35312E-14 | 0,999999496 | 1 |
| WBP2 | 9 | 9 | -1,42389E-17 | 0,999999989 | 1 |
| ARMC1 | 8 | 8 | 0 | 1 | 1 |
| ANAPC5 | 3 | 3 | 0 | 1 | 1 |
| WASHC2 | 5 | 5 | -0,034445179 | 0,37654629 | 1 |
| IGF2R | 4 | 4 | 0 | 1 | 1 |
| NUDCD3 | 18 | 17 | 0,04262989 | 0,263582463 | 1 |
| COMMD9 | 3 | 3 | 0,081601705 | 0,226226418 | 1 |
| PCSK1N | 6 | 6 | -0,086908465 | 0,140357354 | 1 |
| GOT2 | 39 | 39 | -2,66783E-17 | 0,999999989 | 1 |
| TARS2 | 12 | 12 | -2,60853E-17 | 0,999999994 | 1 |
| USP8 | 9 | 8 | -3,72997E-17 | 0,999999979 | 1 |
| CHMP2A | 5 | 5 | 0 | 1 | 1 |
| EIF3B | 22 | 19 | 1,01905E-17 | 0,999999994 | 1 |
| ICAM5 | 24 | 24 | 0 | 1 | 1 |
| SLC12A5 | 32 | 32 | 6,36016E-18 | 0,999999992 | 1 |
| UHRF1BP1L | 26 | 26 | 0 | 1 | 1 |
| MARCKS | 11 | 11 | -0,066142374 | 0,284965598 | 1 |
| TBCA | 6 | 6 | -0,037341882 | 0,390567102 | 1 |
| ANXA2 | 16 | 11 | 0,284055465 | 0,293226183 | 1 |
| RPS2 | 9 | 8 | 0,110270774 | 0,302332871 | 1 |
| ADO | 9 | 9 | -0,049605061 | 0,146710309 | 1 |
| PCNA | 3 | 1 | 0,313895874 | 0,170815985 | 1 |
| PPP1R3G | 4 | 4 | 0,013473313 | 0,667153753 | 1 |
| USP15 | 25 | 22 | 9,74467E-18 | 0,999999995 | 1 |
| HSPH1 | 51 | 49 | 0,010985669 | 0,588640085 | 1 |
| CHID1 | 10 | 10 | -8,47442E-18 | 0,999999995 | 1 |
| ROCK2 | 52 | 52 | 0,017891401 | 0,347437876 | 1 |

| | | | | | |
|---|---|---|---|---|---|
| RDH14 | 8 | 8 | 0 | 1 | 1 |
| SCN3B | 4 | 4 | 4,55184E-20 | 1 | 1 |
| STRN3 | 14 | 14 | 1,3968E-16 | 0,999999965 | 1 |
| MAPT | 1 | 1 | -0,077299764 | 0,211025744 | 1 |
| EPB41L3 | 31 | 31 | -3,88696E-16 | 0,99999992 | 1 |
| CANX | 25 | 25 | 0 | 1 | 1 |
| PITPNA | 27 | 26 | 0,019570202 | 0,42740038 | 1 |
| ATPIF1 | 6 | 6 | 0 | 1 | 1 |
| CDC37 | 20 | 20 | 1,45292E-16 | 0,999999968 | 1 |
| DBNL | 23 | 23 | 0 | 1 | 1 |
| CDC37L1 | 3 | 2 | 0 | 1 | 1 |
| BIN1 | 1 | 1 | -0,238481264 | 0,203495181 | 1 |
| RAB30 | 8 | 8 | 1,4953E-15 | 0,999999863 | 1 |
| RHOT2 | 8 | 8 | -6,31483E-16 | 0,999999941 | 1 |
| RHOT1 | 15 | 15 | 0 | 1 | 1 |
| SETD3 | 10 | 6 | -6,11718E-17 | 0,999999995 | 1 |
| ACAA1A | 18 | 18 | 1,90574E-17 | 0,999999988 | 1 |
| RABEP2 | 2 | 2 | 0 | 1 | 1 |
| DNM1L | 40 | 39 | 0,022354245 | 0,403923412 | 1 |
| RMDN3 | 14 | 14 | -1,95196E-16 | 0,999999975 | 1 |
| TMEM132A | 4 | 4 | 9,20955E-14 | 0,999999169 | 1 |
| C1QBP | 7 | 7 | 0 | 1 | 1 |
| TPM1 | 8 | 8 | -4,78444E-17 | 0,99999999 | 1 |
| VAT1L | 18 | 18 | -3,26308E-17 | 0,999999994 | 1 |
| CES1C | 13 | 1 | 0 | 1 | 1 |
| RMDN1 | 8 | 8 | 0 | 1 | 1 |
| FUNDC2 | 5 | 5 | 0 | 1 | 1 |
| RALB | 4 | 4 | 1,3594E-19 | 1 | 1 |
| ARHGDIA | 15 | 15 | -2,28926E-16 | 0,999999931 | 1 |
| PA2G4 | 20 | 17 | 0,049670769 | 0,236553781 | 1 |
| ENOPH1 | 8 | 8 | -3,91378E-16 | 0,999999975 | 1 |
| DHDH | 8 | 6 | -0,073664627 | 0,204530736 | 1 |
| RNPEP | 21 | 17 | -1,07466E-16 | 0,999999977 | 1 |

| | | | | | |
|---|---|---|---|---|---|
| MAST3 | 9 | 9 | 4,64095E-14 | 0,999999498 | 1 |
| BCL2L13 | 6 | 6 | -0,026475398 | 0,490482652 | 1 |
| BASP1 | 16 | 16 | 1,02889E-16 | 0,999999974 | 1 |
| TPM4 | 8 | 8 | -1,15638E-17 | 0,999999997 | 1 |
| ATXN10 | 23 | 18 | 2,87945E-07 | 0,998023727 | 1 |
| ALS2 | 8 | 8 | 4,81735E-18 | 0,999999998 | 1 |
| ARHGAP23 | 21 | 21 | -1,32129E-17 | 0,99999999 | 1 |
| ANK2 | 150 | 150 | -0,046038597 | 0,154870222 | 1 |
| VCPIP1 | 29 | 29 | -2,61836E-16 | 0,999999941 | 1 |
| PTPRA | 21 | 21 | 4,39945E-14 | 0,999999195 | 1 |
| RYR2 | 19 | 19 | -0,048733259 | 0,130838661 | 1 |
| PTPRE | 12 | 12 | 0,041477412 | 0,298010048 | 1 |
| ATXN2L | 18 | 17 | 1,91962E-17 | 0,999999992 | 1 |
| AHNAK | 4 | 4 | -4,84797E-18 | 0,999999996 | 1 |
| PTPRN | 11 | 11 | 2,82909E-17 | 0,999999995 | 1 |
| LRPPRC | 62 | 62 | -3,52015E-16 | 0,999999959 | 1 |
| AP3D1 | 31 | 31 | 0,052304192 | 0,56914873 | 1 |
| RTN1 | 27 | 27 | -0,011736881 | 0,657997741 | 1 |
| ATG7 | 13 | 13 | 4,17909E-16 | 0,999999933 | 1 |
| PPP2R2D | 2 | 2 | -0,066967079 | 0,232449155 | 1 |
| RELCH | 32 | 31 | 0,002025229 | 0,774408846 | 1 |
| EXTL2 | 4 | 4 | -0,047612514 | 0,405310763 | 1 |
| TLN2 | 75 | 75 | 7,47798E-17 | 0,999999983 | 1 |
| RTN3 | 40 | 39 | -0,057142858 | 0,235902266 | 1 |
| MAP7D2 | 13 | 12 | -4,90226E-16 | 0,999999925 | 1 |
| SCO1 | 2 | 2 | 0 | 1 | 1 |
| BNIP3 | 4 | 4 | -0,076744799 | 0,323343823 | 1 |
| SRGAP2 | 25 | 25 | 0 | 1 | 1 |
| KIF3B | 15 | 15 | 0 | 1 | 1 |
| THNSL1 | 21 | 20 | -1,2059E-15 | 0,999999897 | 1 |
| RABEP1 | 32 | 30 | -1,23674E-17 | 0,999999985 | 1 |
| TRAPPC10 | 13 | 13 | -4,69478E-17 | 0,99999999 | 1 |
| RNF214 | 14 | 14 | -0,032283198 | 0,222674577 | 1 |

| | | | | | |
|---|---|---|---|---|---|
| EML1 | 7 | 7 | 0 | 1 | 1 |
| KATNB1 | 10 | 9 | 0 | 1 | 1 |
| DCTN1 | 57 | 57 | 1,56618E-12 | 0,999994231 | 1 |
| GHITM | 2 | 2 | -6,80393E-16 | 0,999999948 | 1 |
| EPS15 | 14 | 14 | -0,008589806 | 0,595925125 | 1 |
| GANAB | 39 | 39 | 0,02210245 | 0,164108768 | 1 |
| TMEM165 | 3 | 3 | -1,48041E-16 | 0,999999968 | 1 |
| VPS35 | 33 | 31 | 0,027412439 | 0,18271364 | 1 |
| TRAPPC14 | 6 | 6 | 0,033088761 | 0,455458595 | 1 |
| SOGA3 | 34 | 34 | -1,89786E-16 | 0,999999956 | 1 |
| RB1CC1 | 7 | 7 | -0,033328152 | 0,342515929 | 1 |
| ABCG2 | 6 | 6 | 7,64578E-17 | 0,999999971 | 1 |
| CTTNBP2 | 25 | 25 | -8,90622E-16 | 0,999999907 | 1 |
| CCDC9 | 2 | 2 | 0,169340641 | 0,205940181 | 1 |
| APRT | 7 | 6 | -1,45163E-18 | 0,999999999 | 1 |
| CCDC6 | 12 | 11 | 2,0068E-17 | 0,999999988 | 1 |
| DENND4B | 4 | 4 | 0 | 1 | 1 |
| IDI1 | 7 | 6 | 0 | 1 | 1 |
| STRIP2 | 1 | 1 | 0 | 1 | 1 |
| ACSS2 | 11 | 9 | 0,005066113 | 0,781742644 | 1 |
| DSP | 53 | 39 | 9,53099E-15 | 0,999999874 | 1 |
| KCNH3 | 1 | 1 | 0 | 1 | 1 |
| TPD52L1 | 3 | 3 | 0 | 1 | 1 |
| CCDC93 | 10 | 10 | 0 | 1 | 1 |
| NPTXR | 13 | 13 | 0,022754666 | 0,529802441 | 1 |
| BNIP1 | 2 | 2 | -1,7769E-19 | 0,999999999 | 1 |
| RPS3 | 24 | 23 | 0 | 1 | 1 |
| UMPS | 7 | 5 | 0,033172483 | 0,416507639 | 1 |
| STRN | 27 | 27 | 0 | 1 | 1 |
| GOLGA3 | 13 | 13 | -1,00058E-16 | 0,999999966 | 1 |
| GOLGA2 | 5 | 5 | 0 | 1 | 1 |
| PSMD11 | 28 | 27 | 0 | 1 | 1 |
| WDR11 | 9 | 9 | 0 | 1 | 1 |

| | | | | | |
|---|---|---|---|---|---|
| ACSL1 | 22 | 22 | 3,96583E-17 | 0,999999988 | 1 |
| MPRIP | 20 | 20 | 0 | 1 | 1 |
| PALM2 | 13 | 13 | 0 | 1 | 1 |
| GPM6B | 9 | 9 | 0,062813918 | 0,293224834 | 1 |
| USP20 | 1 | 1 | 0 | 1 | 1 |
| DPYSL5 | 30 | 29 | 1,34745E-17 | 0,99999999 | 1 |
| NCALD | 13 | 13 | -8,91185E-18 | 0,999999995 | 1 |
| GFAP | 29 | 29 | 0 | 1 | 1 |
| ALDOC | 37 | 37 | -0,017766471 | 0,50086589 | 1 |
| ATP13A1 | 12 | 12 | -7,08641E-18 | 0,999999995 | 1 |
| ADCK2 | 1 | 1 | 0 | 1 | 1 |
| BCAP31 | 8 | 8 | -0,055736344 | 0,26208702 | 1 |
| ARPC4 | 7 | 7 | 0,023948481 | 0,43449148 | 1 |
| SIK3 | 16 | 15 | -1,15483E-17 | 0,999999994 | 1 |
| LPCAT4 | 5 | 5 | 5,9378E-17 | 0,999999984 | 1 |
| HMOX2 | 16 | 16 | -0,029700691 | 0,12683009 | 1 |
| PRRC2A | 11 | 10 | -0,061130791 | 0,195800532 | 1 |
| UBE2N | 11 | 11 | 0,020856433 | 0,572148588 | 1 |
| ACTR3 | 19 | 19 | 0,043029978 | 0,20154917 | 1 |
| AKAP12 | 16 | 16 | -0,055830685 | 0,229269244 | 1 |
| PTPRD | 33 | 33 | -0,004014986 | 0,687163081 | 1 |
| CHORDC1 | 13 | 12 | 0,017150474 | 0,510214528 | 1 |
| BMERB1 | 3 | 3 | -1,30047E-16 | 0,999999971 | 1 |
| RACK1 | 20 | 15 | 0,017707384 | 0,625306936 | 1 |
| CADPS | 81 | 80 | 0,054037076 | 0,13718221 | 1 |
| HOMER1 | 23 | 23 | 0 | 1 | 1 |
| DOCK9 | 31 | 31 | 0,038205012 | 0,192748089 | 1 |
| LRRFIP2 | 2 | 2 | 3,34444E-16 | 0,999999955 | 1 |
| OGDH | 50 | 50 | -2,68577E-18 | 0,999999997 | 1 |
| STOML2 | 12 | 12 | -3,00831E-16 | 0,999999959 | 1 |
| PDXP | 16 | 16 | -1,23319E-16 | 0,999999971 | 1 |
| DNAH7C;DNAH7B;DI | 1 | 1 | -0,044262326 | 0,420858937 | 1 |
| CYP46A1 | 21 | 21 | 0 | 1 | 1 |

| | | | | | |
|---|---|---|---|---|---|
| IDE | 7 | 7 | 2,00223E-14 | 0,999999574 | 1 |
| LRP1B | 10 | 10 | 6,05933E-17 | 0,999999986 | 1 |
| AK5 | 19 | 18 | 0,021653582 | 0,280364812 | 1 |
| CXXC5 | 1 | 1 | -2,32048E-15 | 0,99999989 | 1 |
| PPP1R9A | 18 | 18 | -0,02687234 | 0,540546301 | 1 |
| KRT2 | 43 | 12 | 0,28174803 | 0,370201584 | 1 |
| HSPA4 | 74 | 73 | 0 | 1 | 1 |
| RPGR | 5 | 4 | 0 | 1 | 1 |
| FARP1 | 21 | 21 | -2,97652E-17 | 0,999999981 | 1 |
| FECH | 14 | 14 | 0 | 1 | 1 |
| PLS3 | 29 | 28 | -0,009694715 | 0,60634085 | 1 |
| WDR44 | 27 | 26 | 0 | 1 | 1 |
| GPS1 | 9 | 9 | 1,52466E-18 | 0,999999999 | 1 |
| SGIP1 | 2 | 2 | 0 | 1 | 1 |
| RNF14 | 1 | 1 | 0,058464007 | 0,402592702 | 1 |
| OPTC | 1 | 1 | 0 | 1 | 1 |
| KRT10 | 26 | 9 | 0,516777636 | 0,243613722 | 1 |
| SEC23A | 14 | 14 | 0,025296634 | 0,306523861 | 1 |
| SHPK | 3 | 3 | -2,05759E-16 | 0,999999981 | 1 |
| SEH1L | 4 | 4 | 0 | 1 | 1 |
| PPFIA3 | 52 | 49 | 0 | 1 | 1 |
| PPFIA2 | 25 | 25 | 0,008470598 | 0,618099104 | 1 |
| LLGL1 | 14 | 14 | -1,10962E-17 | 0,999999996 | 1 |
| NUDC | 26 | 23 | -6,12729E-17 | 0,999999964 | 1 |
| FAM114A2 | 11 | 10 | -4,24173E-18 | 0,999999999 | 1 |
| CLMP | 1 | 1 | 0 | 1 | 1 |
| NRCAM | 32 | 31 | 0,037900218 | 0,125983639 | 1 |
| GET4 | 7 | 7 | 0,054665855 | 0,284517336 | 1 |
| HNRNPU | 16 | 13 | 0,132227557 | 0,156015284 | 1 |
| MOG | 13 | 13 | -0,108090558 | 0,30876192 | 1 |
| CLIP3 | 6 | 6 | -5,92391E-16 | 0,999999927 | 1 |
| KCNAB1 | 13 | 12 | 2,55692E-16 | 0,999999949 | 1 |
| UQCRFS1 | 16 | 16 | -1,06048E-16 | 0,999999987 | 1 |

| | | | | | |
|---|---|---|---|---|---|
| BZW1 | 12 | 11 | 3,33278E-17 | 0,999999987 | 1 |
| NF1 | 13 | 13 | 0 | 1 | 1 |
| THTPA | 7 | 5 | -0,017472417 | 0,626890383 | 1 |
| MICAL3 | 27 | 27 | -0,04673832 | 0,143965262 | 1 |
| NTAN1 | 1 | 1 | 0 | 1 | 1 |
| GARS1 | 30 | 26 | 0 | 1 | 1 |
| LHPP | 4 | 3 | 4,11321E-17 | 0,999999991 | 1 |
| KCNAB2 | 21 | 21 | 0 | 1 | 1 |
| HABP4 | 6 | 5 | 0,116198045 | 0,266938878 | 1 |
| GABRA1 | 10 | 10 | 0,00877375 | 0,660827161 | 1 |
| DNM3 | 36 | 36 | 2,76636E-17 | 0,999999982 | 1 |
| PAICS | 18 | 18 | -1,38574E-16 | 0,999999976 | 1 |
| PEA15 | 11 | 11 | -3,32912E-16 | 0,999999962 | 1 |
| NSMF | 8 | 8 | 0,007692437 | 0,669661013 | 1 |
| HACE1 | 10 | 10 | -1,26946E-17 | 0,999999996 | 1 |
| NDUFS7 | 9 | 9 | -1,43677E-17 | 0,999999995 | 1 |
| WDFY3 | 27 | 27 | 0 | 1 | 1 |
| ECHS1 | 17 | 17 | -0,050120647 | 0,334082018 | 1 |
| RDX | 16 | 16 | 0 | 1 | 1 |
| CCT5 | 36 | 35 | 0 | 1 | 1 |
| OCRL | 22 | 21 | 2,51449E-16 | 0,999999954 | 1 |
| PLCD1 | 15 | 15 | 0,02102019 | 0,377134751 | 1 |
| TARDBP | 6 | 6 | 2,63893E-16 | 0,999999967 | 1 |
| MTMR11 | 1 | 1 | 0 | 1 | 1 |
| PZP | 33 | 5 | -1,7271E-17 | 0,999999997 | 1 |
| SAG | 2 | 2 | -0,004317098 | 0,815709098 | 1 |
| IPO7 | 22 | 22 | -2,21069E-17 | 0,999999995 | 1 |
| HIBCH | 24 | 24 | -0,067210106 | 0,34184926 | 1 |
| QRSL1 | 9 | 9 | -0,008749889 | 0,738251312 | 1 |
| CLASP1 | 27 | 27 | 0 | 1 | 1 |
| PGM2 | 7 | 3 | -0,094516853 | 0,190699423 | 1 |
| GDPD1 | 14 | 14 | 0,01371755 | 0,538297317 | 1 |
| PCCB | 28 | 28 | 6,47835E-18 | 0,999999999 | 1 |

| | | | | | |
|---|---|---|---|---|---|
| UQCR10 | 3 | 3 | -2,78528E-18 | 0,999999997 | 1 |
| ACLY | 56 | 56 | 0,02418968 | 0,373011134 | 1 |
| ARFGAP1 | 17 | 16 | -2,02925E-18 | 0,999999997 | 1 |
| CPNE7 | 22 | 20 | 0,026992252 | 0,565665832 | 1 |
| SLC2A3 | 14 | 14 | 0,00914536 | 0,662148218 | 1 |
| CSMD2 | 7 | 7 | 1,29177E-14 | 0,999999655 | 1 |
| PSMA3 | 15 | 14 | 0,044201693 | 0,217230953 | 1 |
| ALDH9A1 | 19 | 19 | -1,08264E-16 | 0,999999967 | 1 |
| HCN2;HCN1 | 4 | 4 | 0,04058018 | 0,310765395 | 1 |
| PGAP1 | 8 | 7 | 0 | 1 | 1 |
| RYR3 | 4 | 4 | 0 | 1 | 1 |
| NPEPPS | 49 | 47 | -3,56324E-16 | 0,999999968 | 1 |
| SLC32A1 | 13 | 13 | -1,16197E-16 | 0,99999996 | 1 |
| NPLOC4 | 7 | 7 | 0 | 1 | 1 |
| PICK1 | 2 | 2 | -2,39438E-17 | 0,999999991 | 1 |
| GM49486 | 6 | 6 | 0,025428252 | 0,404028296 | 1 |
| PPP1R11 | 8 | 8 | 0,012198401 | 0,695010707 | 1 |
| OTUB1 | 18 | 16 | 0,018916942 | 0,534957325 | 1 |
| FSD1 | 21 | 20 | -5,30359E-18 | 0,999999997 | 1 |
| FLNA | 20 | 20 | 6,30996E-16 | 0,999999964 | 1 |
| PRRC2C | 12 | 12 | -0,006650118 | 0,733993389 | 1 |
| SRSF3 | 2 | 2 | 6,28275E-17 | 0,999999996 | 1 |
| PFAS | 27 | 16 | 9,3815E-22 | 1 | 1 |
| ACBD3 | 6 | 6 | 0 | 1 | 1 |
| IGSF11 | 3 | 3 | 0 | 1 | 1 |
| FOLH1 | 13 | 13 | -0,07705238 | 0,19129812 | 1 |
| MUG1 | 22 | 1 | 2,70591E-19 | 1 | 1 |
| ERMP1 | 8 | 8 | -0,042623684 | 0,252348881 | 1 |
| IDE | 18 | 17 | 2,68254E-16 | 0,999999959 | 1 |
| EIF3A | 45 | 38 | 0 | 1 | 1 |
| DTNA | 9 | 9 | -2,95792E-16 | 0,999999967 | 1 |
| B3GALT6 | 1 | 1 | 0 | 1 | 1 |
| PTGES2 | 13 | 13 | -2,1263E-14 | 0,999999522 | 1 |

| | | | | | |
|---|---|---|---|---|---|
| TRAP1 | 22 | 22 | -6,15639E-18 | 0,999999996 | 1 |
| VPS11 | 18 | 18 | 0 | 1 | 1 |
| RRAGC | 16 | 14 | 0,046337864 | 0,175805517 | 1 |
| NELL2 | 3 | 3 | 0 | 1 | 1 |
| RPL34 | 5 | 5 | 5,67171E-17 | 0,999999985 | 1 |
| CAP2 | 31 | 31 | 2,42759E-17 | 0,999999989 | 1 |
| ZFP933 | 1 | 1 | 0 | 1 | 1 |
| GSTP1 | 14 | 13 | 0 | 1 | 1 |
| ACSL6 | 40 | 40 | 0 | 1 | 1 |
| FRY | 12 | 12 | -0,009660658 | 0,57055054 | 1 |
| TRAPPC12 | 12 | 12 | 2,07711E-18 | 0,999999999 | 1 |
| NBEA | 65 | 61 | 0 | 1 | 1 |
| LYST | 10 | 10 | 1,78392E-17 | 0,999999987 | 1 |
| PNCK | 5 | 4 | 0 | 1 | 1 |
| DCC | 11 | 11 | 0,034376917 | 0,282538039 | 1 |
| ARFGEF2 | 16 | 15 | 0 | 1 | 1 |
| DNAJC2 | 2 | 2 | 0 | 1 | 1 |
| GUCY1B1 | 18 | 18 | 0,007793219 | 0,659918249 | 1 |
| YJEFN3 | 2 | 2 | -0,030990362 | 0,604331491 | 1 |
| LANCL1 | 10 | 10 | 0,029682835 | 0,301609408 | 1 |
| UQCRQ | 9 | 9 | -2,79676E-16 | 0,999999971 | 1 |
| BAIAP2 | 34 | 34 | 0,014247898 | 0,444419443 | 1 |
| SLC9A1 | 11 | 11 | 1,07482E-17 | 0,999999991 | 1 |
| NAPRT | 8 | 7 | -0,081425606 | 0,155510263 | 1 |
| LAMP2 | 5 | 5 | 0 | 1 | 1 |
| AKR1B7 | 2 | 2 | -1,8591E-19 | 0,999999999 | 1 |
| OPLAH | 30 | 29 | -4,53287E-17 | 0,999999979 | 1 |
| GFOD1 | 9 | 9 | -0,014964024 | 0,381070072 | 1 |
| PROM1 | 4 | 4 | -4,49724E-19 | 0,999999999 | 1 |
| UBXN1 | 5 | 5 | 0 | 1 | 1 |
| VAMP7 | 9 | 9 | 0,039757379 | 0,21631557 | 1 |
| ESD | 13 | 13 | 0 | 1 | 1 |
| TOLLIP | 12 | 12 | 1,26832E-17 | 0,99999999 | 1 |

| | | | | | |
|---|---|---|---|---|---|
| GPRC5B | 3 | 3 | -0,057578642 | 0,199844818 | 1 |
| SEC61A2 | 1 | 1 | 0 | 1 | 1 |
| SEC61A1 | 2 | 2 | -2,4104E-16 | 0,999999967 | 1 |
| FARSA | 19 | 13 | 0 | 1 | 1 |
| G3BP2 | 16 | 13 | 0 | 1 | 1 |
| CLASP2 | 6 | 6 | -6,81453E-17 | 0,99999998 | 1 |
| ATG2B | 12 | 12 | -1,7403E-18 | 0,999999998 | 1 |
| VIPAS39 | 11 | 11 | 2,73748E-17 | 0,999999984 | 1 |
| ACADM | 16 | 16 | -0,094325037 | 0,260716535 | 1 |
| ATP2A2 | 56 | 56 | 5,17484E-12 | 0,999988236 | 1 |
| ATIC | 29 | 26 | -5,75978E-18 | 0,999999998 | 1 |
| TGFBRAP1 | 5 | 5 | 6,01191E-17 | 0,999999983 | 1 |
| RPP14 | 1 | 1 | -0,191284865 | 0,229021025 | 1 |
| FAH | 9 | 8 | -0,065901971 | 0,22083419 | 1 |
| TFRC | 21 | 21 | 2,55318E-18 | 0,999999998 | 1 |
| RTCA | 7 | 5 | 0 | 1 | 1 |
| SLC12A9 | 7 | 7 | -6,53627E-18 | 0,999999996 | 1 |
| PGM3 | 18 | 15 | 0 | 1 | 1 |
| ABR | 23 | 23 | -2,02648E-16 | 0,999999951 | 1 |
| PREP | 44 | 40 | 0 | 1 | 1 |
| VPS45 | 20 | 20 | 1,61231E-15 | 0,99999981 | 1 |
| EEF1D | 6 | 5 | 0 | 1 | 1 |
| TIAM1 | 1 | 1 | 0 | 1 | 1 |
| RGS1 | 1 | 1 | 6,09897E-18 | 0,999999994 | 1 |
| TUBA1C;TUBA8 | 17 | 17 | 1,1057E-16 | 0,999999975 | 1 |
| TUBA1C | 15 | 15 | 4,52021E-17 | 0,99999999 | 1 |
| UPF2 | 7 | 7 | -1,79707E-17 | 0,999999989 | 1 |
| XPNPEP1 | 17 | 16 | -1,09355E-16 | 0,999999975 | 1 |
| SLC12A2 | 24 | 23 | -4,05954E-16 | 0,99999994 | 1 |
| OAT | 17 | 17 | -0,181765153 | 0,118528958 | 1 |
| CLTB | 2 | 2 | 0 | 1 | 1 |
| ARHGAP12 | 4 | 4 | 0 | 1 | 1 |
| ENO2 | 29 | 28 | 6,76381E-17 | 0,999999991 | 1 |

| | | | | | |
|---|---|---|---|---|---|
| SYNGR3 | 4 | 4 | 9,77615E-19 | 0,999999998 | 1 |
| PCNP | 4 | 4 | 3,11038E-16 | 0,999999958 | 1 |
| GAD1 | 13 | 13 | 3,77415E-16 | 0,99999992 | 1 |
| PLCL1 | 14 | 13 | 2,46374E-17 | 0,999999993 | 1 |
| ALAD | 12 | 10 | 1,41572E-17 | 0,999999988 | 1 |
| UBR3 | 3 | 3 | 2,53507E-18 | 0,999999999 | 1 |
| ATP5MK | 4 | 4 | 1,31072E-15 | 0,999999979 | 1 |
| NUMBL | 11 | 11 | 5,24701E-16 | 0,999999912 | 1 |
| MRI1 | 5 | 4 | 0,003946572 | 0,803924023 | 1 |
| MDH2 | 24 | 24 | -1,66208E-18 | 0,999999997 | 1 |
| ANKRD63 | 3 | 3 | -4,65014E-13 | 0,999997965 | 1 |
| DCTN4 | 13 | 13 | 3,79211E-17 | 0,999999978 | 1 |
| TMEM160 | 4 | 4 | -1,40089E-18 | 0,999999999 | 1 |
| DYNC1LI1 | 26 | 26 | 1,39576E-16 | 0,999999965 | 1 |
| LGI3 | 11 | 11 | 0 | 1 | 1 |
| DHRS1 | 10 | 10 | 0 | 1 | 1 |
| TNKS1BP1 | 36 | 36 | -9,31361E-18 | 0,999999994 | 1 |
| SEC11C | 4 | 4 | 0,066467921 | 0,355756033 | 1 |
| DIAPH1 | 12 | 12 | 5,54205E-16 | 0,999999958 | 1 |
| HADH | 7 | 7 | -0,002939971 | 0,883416406 | 1 |
| SQSTM1 | 6 | 6 | 0 | 1 | 1 |
| ETFB | 6 | 6 | -0,1046078 | 0,182759418 | 1 |
| SLC6A1 | 11 | 11 | 0 | 1 | 1 |
| MRPL2 | 2 | 2 | -4,07615E-18 | 0,999999999 | 1 |
| LYSMD2 | 3 | 3 | 0,00920924 | 0,629937454 | 1 |
| PCCA | 41 | 41 | -7,23114E-18 | 0,999999995 | 1 |
| HDGF | 12 | 2 | -1,18071E-18 | 0,999999999 | 1 |
| LMO7 | 4 | 4 | -1,41005E-16 | 0,999999979 | 1 |
| DDC | 14 | 13 | -0,05390116 | 0,238194926 | 1 |
| STAM | 14 | 14 | 0 | 1 | 1 |
| PDCL | 3 | 3 | -1,82206E-17 | 0,999999989 | 1 |
| NRAP | 1 | 1 | 0 | 1 | 1 |
| RPS18 | 12 | 11 | 0,154590465 | 0,189818877 | 1 |

| | | | | | |
|---|---|---|---|---|---|
| MRPL58 | 2 | 2 | 0 | 1 | 1 |
| SEC14L2 | 18 | 17 | 0 | 1 | 1 |
| PALS2 | 22 | 21 | 2,48331E-17 | 0,999999987 | 1 |
| ECPAS | 20 | 19 | 0 | 1 | 1 |
| USP11 | 16 | 15 | 0 | 1 | 1 |
| DMWD | 7 | 7 | -0,023958545 | 0,437656709 | 1 |
| MTFP1 | 6 | 6 | -7,65407E-18 | 0,999999996 | 1 |
| ACTA1 | 23 | 19 | -8,0394E-18 | 0,999999994 | 1 |
| DNAJC11 | 22 | 21 | -4,8707E-15 | 0,999999811 | 1 |
| USP7 | 29 | 28 | 6,6604E-15 | 0,999999614 | 1 |
| IPO4 | 9 | 8 | -2,77649E-17 | 0,999999984 | 1 |
| PCYOX1 | 14 | 14 | 0 | 1 | 1 |
| LGI1 | 24 | 24 | 3,74052E-16 | 0,999999927 | 1 |
| CLASP2 | 42 | 41 | 0 | 1 | 1 |
| MIB2 | 2 | 2 | 0 | 1 | 1 |
| EIF2S3X | 16 | 13 | 0,035319054 | 0,347137577 | 1 |
| PGK1 | 33 | 29 | 1,14308E-15 | 0,999999915 | 1 |
| XPO5 | 14 | 13 | -0,04877577 | 0,324666084 | 1 |
| CACNG8 | 11 | 11 | 0 | 1 | 1 |
| USP31 | 8 | 8 | -1,64403E-16 | 0,999999962 | 1 |
| PSMB1 | 12 | 12 | 0,012389516 | 0,640906147 | 1 |
| ARMC6 | 7 | 7 | 1,40805E-16 | 0,999999961 | 1 |
| TANGO2 | 2 | 2 | 1,23009E-13 | 0,999999523 | 1 |
| ACAT3 | 11 | 11 | 2,34592E-17 | 0,999999988 | 1 |
| PADI2 | 10 | 10 | -2,02881E-17 | 0,999999986 | 1 |
| PAM16 | 7 | 7 | 0 | 1 | 1 |
| GAPDHS | 1 | 1 | -0,036796029 | 0,521205623 | 1 |
| COX6C | 11 | 11 | -3,0171E-17 | 0,999999988 | 1 |
| IRGQ | 13 | 13 | -2,8991E-17 | 0,999999992 | 1 |
| TRAPPC11 | 12 | 12 | 0 | 1 | 1 |
| NCDN | 35 | 33 | 4,1384E-16 | 0,999999941 | 1 |
| ACAT1 | 27 | 27 | -4,8784E-18 | 0,999999996 | 1 |
| NECTIN1 | 8 | 8 | 0 | 1 | 1 |

| | | | | | |
|---|---|---|---|---|---|
| AHCY | 28 | 24 | 4,74675E-18 | 0,999999998 | 1 |
| CUL1 | 27 | 27 | 0 | 1 | 1 |
| TMCC2 | 6 | 6 | -0,020217957 | 0,489935199 | 1 |
| TECPR1 | 20 | 20 | 0,047209531 | 0,278554901 | 1 |
| SMIM20 | 1 | 1 | 0 | 1 | 1 |
| GSN | 20 | 20 | 1,27544E-15 | 0,999999938 | 1 |
| DPP8 | 10 | 9 | 2,03436E-17 | 0,999999989 | 1 |
| DPP9 | 13 | 12 | 5,63873E-18 | 0,999999998 | 1 |
| DGKH | 18 | 18 | -0,053037944 | 0,279970848 | 1 |
| UROD | 9 | 9 | 0,013391318 | 0,611636086 | 1 |
| SPARCL1 | 15 | 14 | -0,053722874 | 0,121416093 | 1 |
| DPP3 | 31 | 31 | 3,15893E-17 | 0,999999988 | 1 |
| CPT2 | 11 | 11 | -0,052348805 | 0,296452494 | 1 |
| TIMM9 | 4 | 4 | 0 | 1 | 1 |
| ARL5A | 2 | 2 | 4,58039E-18 | 0,999999997 | 1 |
| FDXR | 11 | 11 | -0,052553418 | 0,416599486 | 1 |
| VPS33B | 10 | 10 | 9,28659E-18 | 0,999999994 | 1 |
| EIF2S1 | 19 | 18 | 0 | 1 | 1 |
| PPP2R5D | 24 | 24 | -4,18522E-16 | 0,999999909 | 1 |
| ADGRL1 | 25 | 25 | -5,81712E-17 | 0,999999985 | 1 |
| HECTD3 | 11 | 10 | 8,10735E-17 | 0,999999983 | 1 |
| MTCH1 | 13 | 13 | 0 | 1 | 1 |
| CP | 9 | 6 | 0,099667247 | 0,444911379 | 1 |
| AMPD2 | 22 | 22 | 5,45967E-14 | 0,999999018 | 1 |
| H2AC20;HIST2H2AA: | 4 | 3 | 0,466919049 | 0,26654816 | 1 |
| TOM1 | 12 | 12 | -1,88572E-17 | 0,999999987 | 1 |
| HTRA2 | 9 | 9 | -0,039298873 | 0,400774671 | 1 |
| SORBS1 | 8 | 8 | -0,070153309 | 0,150586909 | 1 |
| UPF1 | 26 | 25 | -1,57102E-16 | 0,999999961 | 1 |
| MON2 | 12 | 12 | 0 | 1 | 1 |
| PRPSAP2 | 18 | 18 | 0 | 1 | 1 |
| SNTB2 | 4 | 3 | 0,01378138 | 0,621258077 | 1 |
| MRPL45 | 3 | 3 | -2,45058E-18 | 0,999999998 | 1 |

| | | | | | |
|---|---|---|---|---|---|
| LRRC8C | 5 | 5 | 0,026477474 | 0,36725503 | 1 |
| SLC27A1 | 8 | 8 | -1,45486E-17 | 0,99999999 | 1 |
| DGKD | 4 | 4 | -0,016517908 | 0,665358842 | 1 |
| GSTK1 | 15 | 15 | -1,26874E-16 | 0,999999975 | 1 |
| RPL7 | 16 | 15 | 0,079404907 | 0,213577531 | 1 |
| SLK | 20 | 19 | 0 | 1 | 1 |
| TAOK1 | 15 | 14 | 8,77802E-17 | 0,999999964 | 1 |
| CUL4A | 15 | 15 | 0,003403104 | 0,756363219 | 1 |
| CBS | 8 | 8 | -0,019664591 | 0,540213863 | 1 |
| CUL9 | 2 | 2 | -3,56488E-17 | 0,999999988 | 1 |
| NF1 | 10 | 10 | 0 | 1 | 1 |
| CPT1A | 9 | 9 | -9,54719E-18 | 0,999999997 | 1 |
| CNTNAP4 | 11 | 11 | 1,42953E-17 | 0,999999989 | 1 |
| SRRT | 5 | 1 | 0 | 1 | 1 |
| CAMKK2 | 14 | 13 | 3,34178E-16 | 0,999999971 | 1 |
| STAU2 | 4 | 3 | 7,96239E-17 | 0,999999992 | 1 |
| COG7 | 6 | 6 | 2,37894E-12 | 0,999997499 | 1 |
| ANKRD34B | 3 | 3 | -2,37763E-16 | 0,999999967 | 1 |
| MTSS2 | 6 | 6 | 0 | 1 | 1 |
| SYNCRIP | 21 | 19 | -1,63603E-17 | 0,999999994 | 1 |
| SRC | 18 | 18 | 2,32683E-18 | 0,999999996 | 1 |
| PHACTR1 | 15 | 14 | 1,84296E-15 | 0,999999851 | 1 |
| NIPSNAP1 | 16 | 16 | 0 | 1 | 1 |
| MAP2K5 | 2 | 2 | 0 | 1 | 1 |
| ATL1 | 26 | 26 | 0 | 1 | 1 |
| MRPL11 | 6 | 6 | -0,096262445 | 0,180678805 | 1 |
| PGD | 27 | 25 | 0 | 1 | 1 |
| LMAN2L | 5 | 5 | -2,74594E-15 | 0,999999856 | 1 |
| TBC1D24 | 21 | 21 | 3,84603E-18 | 0,999999999 | 1 |
| ITSN2 | 14 | 14 | -0,028419186 | 0,335074603 | 1 |
| MIA3 | 3 | 3 | 0 | 1 | 1 |
| CNN3 | 10 | 8 | -4,18481E-17 | 0,999999979 | 1 |
| DLG3 | 44 | 42 | 0,005973265 | 0,634098889 | 1 |

| | | | | | |
|---|---|---|---|---|---|
| SLC25A12 | 38 | 38 | -6,27907E-16 | 0,999999961 | 1 |
| MYO5B;MYO5A | 6 | 6 | 0,032084903 | 0,321573958 | 1 |
| ATP2B2 | 37 | 37 | 4,56494E-17 | 0,99999999 | 1 |
| MAP2K1 | 20 | 20 | 0,041722425 | 0,241897225 | 1 |
| TPPP | 14 | 14 | -3,12916E-16 | 0,999999958 | 1 |
| RASGRF1 | 18 | 17 | 0 | 1 | 1 |
| ASTN1 | 25 | 25 | 0,023872295 | 0,159298133 | 1 |
| CARMIL2 | 15 | 15 | 0,040478627 | 0,312783888 | 1 |
| CLIC4 | 9 | 9 | -0,020788241 | 0,600400622 | 1 |
| TMEM229A | 1 | 1 | 0 | 1 | 1 |
| SLC8A2 | 31 | 31 | 0,002331678 | 0,800364593 | 1 |
| LRRC7 | 46 | 46 | 5,167E-15 | 0,999999612 | 1 |
| WNK2 | 20 | 20 | -5,76981E-17 | 0,999999973 | 1 |
| MYO6 | 11 | 11 | 0 | 1 | 1 |
| PPIP5K1 | 7 | 7 | 0 | 1 | 1 |
| UQCRC2 | 25 | 25 | -9,80685E-17 | 0,999999976 | 1 |
| FMNL1 | 17 | 17 | 0 | 1 | 1 |
| DPY19L1 | 1 | 1 | 0 | 1 | 1 |
| TTBK1 | 4 | 4 | 0 | 1 | 1 |
| PSMG1 | 3 | 3 | 0 | 1 | 1 |
| PHGDH | 18 | 17 | -0,048106019 | 0,151878629 | 1 |
| PPM1H | 20 | 20 | 0,0299058 | 0,401127054 | 1 |
| FAM13C | 7 | 7 | -0,016148054 | 0,549537359 | 1 |
| CAPN1 | 15 | 15 | 0 | 1 | 1 |
| ACTN4 | 16 | 16 | -0,010868956 | 0,649210733 | 1 |
| ACTN1 | 48 | 48 | -0,036695926 | 0,423111401 | 1 |
| ACTN2 | 31 | 31 | -3,33646E-18 | 0,999999998 | 1 |
| ACTN1;ACTN2 | 13 | 13 | 0 | 1 | 1 |
| ITGAV | 17 | 17 | 3,63291E-18 | 0,999999996 | 1 |
| PCDH19 | 6 | 6 | 4,52607E-18 | 0,999999997 | 1 |
| TTC9C | 4 | 4 | 0 | 1 | 1 |
| SFXN3 | 15 | 15 | -3,35113E-17 | 0,999999985 | 1 |
| PRKCB | 38 | 38 | 0,065666808 | 0,120240544 | 1 |

| | | | | | |
|---|---|---|---|---|---|
| NAGK | 10 | 10 | 0 | 1 | 1 |
| CKB | 30 | 28 | -2,14611E-16 | 0,999999971 | 1 |
| CKMT1 | 28 | 28 | -2,81848E-17 | 0,999999989 | 1 |
| HSD17B8 | 10 | 10 | 4,11246E-17 | 0,999999993 | 1 |
| BIRC6 | 27 | 27 | 0 | 1 | 1 |
| RTRAF | 10 | 10 | 0 | 1 | 1 |
| PDCD6 | 9 | 9 | -3,42519E-17 | 0,999999987 | 1 |
| RPL7A | 9 | 9 | 0,330867601 | 0,153052016 | 1 |
| CMBL | 5 | 4 | -1,04228E-16 | 0,99999998 | 1 |
| CLIC6 | 16 | 6 | 7,81291E-17 | 0,999999989 | 1 |
| PAK5 | 7 | 7 | 0,022724285 | 0,467673153 | 1 |
| FXR2 | 10 | 10 | -4,68878E-18 | 0,999999994 | 1 |
| SYNGAP1 | 74 | 72 | 0,059936083 | 0,120166555 | 1 |
| AIP | 9 | 9 | 1,27497E-17 | 0,999999994 | 1 |
| WDR1 | 33 | 32 | 9,58466E-17 | 0,99999999 | 1 |
| PPP3CA | 29 | 29 | 0,044093918 | 0,116100485 | 1 |
| TRAF3 | 14 | 14 | 0 | 1 | 1 |
| ARPC1A | 18 | 18 | 0,032836607 | 0,359103495 | 1 |
| ARSA | 4 | 4 | -1,53674E-17 | 0,999999995 | 1 |
| AP1M1 | 17 | 17 | 2,93525E-17 | 0,999999981 | 1 |
| ILDR2 | 7 | 7 | 0,056215403 | 0,205520316 | 1 |
| ATP5F1B | 43 | 43 | -0,001332585 | 0,887405535 | 1 |
| NTRK3 | 5 | 5 | 1,21659E-16 | 0,999999959 | 1 |
| HDHD3 | 7 | 7 | -0,086720519 | 0,181670794 | 1 |
| PCYT2 | 14 | 12 | -9,65777E-15 | 0,999999559 | 1 |
| GGACT | 3 | 2 | 0 | 1 | 1 |
| sp\|Q91V76\|CK054_ | 9 | 8 | -0,011752037 | 0,625600718 | 1 |
| ADSS2 | 17 | 17 | 2,8523E-18 | 0,999999996 | 1 |
| ATP9B | 3 | 3 | 1,99661E-16 | 0,999999968 | 1 |
| ARMT1 | 17 | 17 | -1,05019E-17 | 0,999999994 | 1 |
| VTI1A | 6 | 6 | -5,93641E-18 | 0,999999992 | 1 |
| FBXO7 | 5 | 5 | -8,95624E-18 | 0,999999997 | 1 |
| ANXA11 | 18 | 18 | 4,12338E-18 | 0,999999997 | 1 |

| | | | | | |
|---|---|---|---|---|---|
| ADSS1 | 17 | 8 | 0 | 1 | 1 |
| PRCP | 2 | 2 | -1,83323E-15 | 0,999999914 | 1 |
| TPD52L2 | 3 | 3 | -3,97556E-18 | 0,999999997 | 1 |
| PLD2 | 4 | 4 | 0 | 1 | 1 |
| NDUFA2 | 5 | 5 | 2,15243E-18 | 0,999999999 | 1 |
| FAF2 | 10 | 10 | 0 | 1 | 1 |
| CRAT | 17 | 17 | -1,71738E-15 | 0,999999888 | 1 |
| PLCH2 | 27 | 25 | 0 | 1 | 1 |
| PPP6C | 8 | 8 | 0,072635886 | 0,158249269 | 1 |
| IMMT | 55 | 55 | -3,10536E-12 | 0,999996101 | 1 |
| MGST3 | 5 | 5 | 0,003727101 | 0,850936672 | 1 |
| SNAP23 | 3 | 3 | -5,98514E-16 | 0,999999935 | 1 |
| CNTN1 | 68 | 68 | 0,041888923 | 0,224006543 | 1 |
| CADM1 | 11 | 11 | 5,69065E-17 | 0,999999984 | 1 |
| MINK1 | 22 | 22 | 0 | 1 | 1 |
| CSDE1 | 32 | 31 | -5,93896E-17 | 0,999999975 | 1 |
| ELFN2 | 18 | 18 | 0,0481133 | 0,165648458 | 1 |
| SEC23IP | 18 | 12 | -1,93723E-18 | 0,999999997 | 1 |
| DIP2A | 14 | 14 | -2,34334E-17 | 0,999999996 | 1 |
| UGP2 | 24 | 24 | -4,16137E-17 | 0,999999971 | 1 |
| MTND4 | 10 | 10 | 1,542E-16 | 0,99999998 | 1 |
| HOMER2 | 14 | 14 | 0 | 1 | 1 |
| OMG | 10 | 10 | -0,038885107 | 0,195410037 | 1 |
| CTNNA1 | 21 | 21 | -0,032481265 | 0,222505619 | 1 |
| CTNNA2 | 47 | 47 | 7,07377E-18 | 0,999999989 | 1 |
| DAAM2 | 9 | 9 | -0,040110115 | 0,34964853 | 1 |
| ZC2HC1A | 13 | 13 | -2,35496E-12 | 0,999993271 | 1 |
| ADCY5 | 14 | 14 | 0,035902209 | 0,248603828 | 1 |
| PAFAH1B1 | 30 | 29 | 0 | 1 | 1 |
| BPNT1 | 17 | 16 | -0,021517933 | 0,297455266 | 1 |
| CCT8 | 50 | 47 | 0,027524044 | 0,276997075 | 1 |
| NAPB;NAPA | 4 | 4 | 0,045014475 | 0,180087638 | 1 |
| NLGN2 | 15 | 15 | 4,7045E-16 | 0,9999999 | 1 |

| | | | | | |
|---|---|---|---|---|---|
| AIFM1 | 28 | 27 | -1,4836E-15 | 0,999999901 | 1 |
| SLC4A4 | 23 | 23 | 8,32427E-18 | 0,999999996 | 1 |
| LRP5 | 1 | 1 | -0,104600141 | 0,181642026 | 1 |
| MRPS9 | 5 | 5 | -1,03391E-17 | 0,999999993 | 1 |
| NCAM2 | 27 | 27 | 1,11166E-16 | 0,999999975 | 1 |
| ST13 | 16 | 13 | 0 | 1 | 1 |
| USP9X | 63 | 63 | -0,013432827 | 0,238247441 | 1 |
| ADAM10 | 6 | 6 | -1,31909E-17 | 0,999999992 | 1 |
| DLGAP4 | 17 | 17 | -1,00035E-17 | 0,999999992 | 1 |
| SELENOO | 4 | 4 | -1,56911E-16 | 0,999999984 | 1 |
| NT5DC3 | 30 | 30 | 0 | 1 | 1 |
| DHCR7 | 4 | 4 | 0,086535833 | 0,158540072 | 1 |
| GNPDA1 | 12 | 11 | -1,40306E-14 | 0,999999522 | 1 |
| EPHA4;EPHA7 | 1 | 1 | 0 | 1 | 1 |
| MTMR6 | 6 | 6 | 4,12195E-18 | 0,999999998 | 1 |
| TTC22 | 1 | 1 | 0 | 1 | 1 |
| NMT1 | 17 | 14 | 0,008114325 | 0,67292676 | 1 |
| SRP72 | 7 | 6 | 0,019024417 | 0,568187036 | 1 |
| SGTA | 15 | 15 | 0,047665765 | 0,130331164 | 1 |
| OTULIN | 3 | 3 | 0 | 1 | 1 |
| ACOT11 | 6 | 6 | 1,88727E-15 | 0,999999908 | 1 |
| PPP1R7 | 32 | 30 | 0 | 1 | 1 |
| GGCT | 10 | 9 | 1,42468E-18 | 0,999999997 | 1 |
| BAG5 | 12 | 12 | 1,48378E-16 | 0,999999952 | 1 |
| AP1S2 | 2 | 2 | -1,15141E-15 | 0,999999943 | 1 |
| EIF2A | 9 | 9 | 0 | 1 | 1 |
| STX1B | 24 | 24 | 0,015941001 | 0,273506624 | 1 |
| ASPH | 3 | 3 | 1,32192E-20 | 1 | 1 |
| PNPLA8 | 17 | 17 | -1,31099E-17 | 0,999999993 | 1 |
| CORO1C | 23 | 22 | 0 | 1 | 1 |
| GABARAPL2 | 6 | 6 | 0,013801162 | 0,669353724 | 1 |
| DHX30 | 11 | 11 | 0 | 1 | 1 |
| TCP11L1 | 6 | 6 | 3,92562E-17 | 0,999999985 | 1 |

| | | | | | |
|---|---|---|---|---|---|
| PVALB | 10 | 8 | -0,112486093 | 0,158606036 | 1 |
| CTSD | 19 | 18 | 2,10954E-16 | 0,99999997 | 1 |
| PPM1F | 9 | 9 | 1,07311E-17 | 0,999999997 | 1 |
| EXOG | 16 | 16 | -7,4629E-17 | 0,99999998 | 1 |
| AUH | 17 | 17 | -6,88249E-18 | 0,999999995 | 1 |
| DGKQ | 14 | 14 | 0,03110909 | 0,349851767 | 1 |
| PSMA5 | 12 | 11 | 3,24273E-18 | 0,999999996 | 1 |
| NFS1 | 20 | 20 | -0,000225368 | 0,954002004 | 1 |
| GK | 19 | 19 | -8,09502E-17 | 0,999999987 | 1 |
| APPL2;APPL1 | 1 | 1 | 0 | 1 | 1 |
| PSMB4 | 13 | 13 | -1,61318E-17 | 0,999999998 | 1 |
| FMNL2 | 9 | 9 | -0,063371393 | 0,129859542 | 1 |
| XPO1 | 35 | 33 | 2,25632E-16 | 0,999999936 | 1 |
| PTPRZ1 | 34 | 34 | 0,02899196 | 0,437202814 | 1 |
| HOOK3 | 21 | 17 | 4,68479E-17 | 0,999999992 | 1 |
| ASAP1 | 20 | 20 | -9,91407E-17 | 0,999999962 | 1 |
| RPS7 | 11 | 11 | 2,49439E-17 | 0,999999986 | 1 |
| ABCA2 | 3 | 3 | 0 | 1 | 1 |
| NLGN3 | 13 | 13 | 1,52359E-17 | 0,999999995 | 1 |
| CLTA | 10 | 9 | 0 | 1 | 1 |
| PTGR2 | 10 | 9 | -1,06866E-18 | 0,999999999 | 1 |
| PTPN9 | 16 | 16 | 0,060075253 | 0,123588793 | 1 |
| COPZ1 | 5 | 5 | -2,02038E-17 | 0,999999988 | 1 |
| BDH1 | 23 | 23 | -3,23079E-17 | 0,99999999 | 1 |
| LTA4H | 41 | 37 | 0 | 1 | 1 |
| AIMP1 | 13 | 10 | -1,35492E-17 | 0,999999987 | 1 |
| ESRRB | 1 | 1 | 0 | 1 | 1 |
| CARM1 | 7 | 7 | -1,49632E-14 | 0,9999996 | 1 |
| TBC1D1 | 2 | 2 | 4,69909E-19 | 1 | 1 |
| KALRN | 35 | 34 | 0,025316482 | 0,185024096 | 1 |
| DMXL1;DMXL2 | 2 | 2 | 1,83354E-15 | 0,999999926 | 1 |
| STAT3 | 4 | 4 | 0 | 1 | 1 |
| GSTA4 | 13 | 13 | 0 | 1 | 1 |

| | | | | | |
|---|---|---|---|---|---|
| NAXD | 15 | 15 | 8,44374E-16 | 0,99999994 | 1 |
| PPP6R2 | 13 | 13 | 1,69715E-14 | 0,99999964 | 1 |
| TUBB5;TUBB2A;TUB | 5 | 5 | 6,08204E-17 | 0,999999981 | 1 |
| ARL1 | 4 | 4 | 0,053813218 | 0,450989971 | 1 |
| SDR39U1 | 10 | 10 | -2,72342E-16 | 0,999999953 | 1 |
| GPD2 | 56 | 56 | 0 | 1 | 1 |
| SYT7 | 23 | 23 | 0,027976542 | 0,298485222 | 1 |
| ANXA6 | 52 | 50 | 0,055354972 | 0,130374001 | 1 |
| QARS1 | 25 | 24 | 1,85886E-16 | 0,999999964 | 1 |
| ASAP2 | 14 | 14 | -1,23146E-17 | 0,99999999 | 1 |
| ARHGAP1 | 18 | 18 | 0 | 1 | 1 |
| ROBO2 | 21 | 21 | 0 | 1 | 1 |
| PSMA6 | 13 | 11 | 0,021927239 | 0,497815763 | 1 |
| KHSRP | 9 | 5 | -1,1947E-17 | 0,999999996 | 1 |
| HIP1R | 24 | 24 | -0,010482073 | 0,552888951 | 1 |
| IGSF8 | 18 | 18 | 0 | 1 | 1 |
| EFL1 | 6 | 6 | -1,38937E-17 | 0,99999999 | 1 |
| EDF1 | 4 | 1 | 0 | 1 | 1 |
| DNM2 | 13 | 13 | 0,041950385 | 0,169211819 | 1 |
| PPID | 21 | 19 | 0,006878895 | 0,625334615 | 1 |
| ABCB6 | 3 | 2 | -3,91107E-19 | 0,999999999 | 1 |
| THEM4 | 9 | 9 | 8,73758E-17 | 0,99999998 | 1 |
| MIOS | 9 | 9 | 0 | 1 | 1 |
| PSMD3 | 34 | 32 | 0 | 1 | 1 |
| TTC1 | 8 | 8 | 5,82778E-17 | 0,999999982 | 1 |
| ANAPC7 | 2 | 2 | 0 | 1 | 1 |
| COPS4 | 30 | 29 | 0,027983025 | 0,278180512 | 1 |
| ATXN3 | 3 | 2 | 0 | 1 | 1 |
| AKT2 | 3 | 3 | 0 | 1 | 1 |
| DNAJB12 | 4 | 4 | -2,81103E-18 | 0,999999996 | 1 |
| REPS2 | 15 | 15 | 0 | 1 | 1 |
| SNPH | 22 | 22 | 0 | 1 | 1 |
| FGG | 10 | 4 | 1,22438E-15 | 0,999999981 | 1 |

| | | | | | |
|---|---|---|---|---|---|
| NDUFAF1 | 9 | 9 | -1,13571E-15 | 0,9999999 | 1 |
| D430041D05RIK | 15 | 15 | 1,55666E-16 | 0,999999964 | 1 |
| ICA1L | 3 | 3 | 0 | 1 | 1 |
| ICA1 | 11 | 11 | 5,44606E-17 | 0,999999988 | 1 |
| TMEM121B | 4 | 4 | 1,59346E-19 | 1 | 1 |
| ZPR1 | 5 | 3 | 0 | 1 | 1 |
| WIPF3 | 13 | 12 | -2,7207E-17 | 0,999999986 | 1 |
| PPP2R1A | 35 | 35 | 0,028857957 | 0,206940297 | 1 |
| SHISA6 | 21 | 20 | 0,007841173 | 0,681774977 | 1 |
| OLFR1135 | 1 | 1 | 0 | 1 | 1 |
| TIMM44 | 25 | 25 | -9,01186E-16 | 0,999999943 | 1 |
| LRSAM1 | 7 | 7 | 9,06155E-17 | 0,999999979 | 1 |
| TSC2 | 20 | 20 | -0,018063626 | 0,347533855 | 1 |
| ACAN | 22 | 17 | -0,048566518 | 0,387849186 | 1 |
| TBRG4 | 12 | 12 | 0 | 1 | 1 |
| YTHDF3 | 5 | 4 | 1,21601E-14 | 0,999999798 | 1 |
| RABGEF1 | 6 | 6 | 0,007283549 | 0,723121041 | 1 |
| PLXNB2 | 16 | 16 | 0 | 1 | 1 |
| GCDH | 10 | 10 | -0,036590256 | 0,482456644 | 1 |
| HNRNPL | 3 | 3 | 0,158024012 | 0,133096032 | 1 |
| PCBP1 | 12 | 12 | 0,052622417 | 0,335643383 | 1 |
| RAB11B;RAB11A | 14 | 14 | 0,023858812 | 0,378480782 | 1 |
| GRIN1 | 29 | 28 | 0,0447424 | 0,191929696 | 1 |
| UGGT1 | 21 | 21 | -0,01356672 | 0,417597986 | 1 |
| PSMA7 | 13 | 13 | 0 | 1 | 1 |
| NRXN2 | 19 | 19 | -2,58058E-16 | 0,999999938 | 1 |
| FHL1 | 3 | 3 | 0 | 1 | 1 |
| CHM | 10 | 8 | 0,01875369 | 0,532393505 | 1 |
| ADGRL2 | 7 | 7 | 1,60364E-15 | 0,999999896 | 1 |
| WNK3 | 2 | 2 | 3,65926E-18 | 0,999999996 | 1 |
| HNRNPUL1 | 9 | 4 | -1,19428E-19 | 1 | 1 |
| PAFAH1B3 | 9 | 7 | -3,31925E-16 | 0,999999949 | 1 |
| NCKAP1L | 8 | 7 | 4,95255E-16 | 0,999999958 | 1 |

| | | | | | |
|---|---|---|---|---|---|
| SOD2 | 13 | 13 | 0 | 1 | 1 |
| ABLIM1 | 17 | 17 | -0,034862066 | 0,354909065 | 1 |
| PEPD | 15 | 15 | -1,20325E-17 | 0,999999997 | 1 |
| GFM1 | 21 | 21 | -6,37099E-16 | 0,999999942 | 1 |
| ABLIM2 | 14 | 14 | -1,58467E-17 | 0,999999988 | 1 |
| PSMB5 | 14 | 14 | 4,40735E-18 | 0,999999994 | 1 |
| ABRAXAS2 | 8 | 8 | 6,82776E-17 | 0,999999973 | 1 |
| SCRN1 | 19 | 14 | -0,008900662 | 0,701227721 | 1 |
| ACO2 | 53 | 53 | -0,023288553 | 0,61273372 | 1 |
| FERMT2 | 15 | 14 | -3,19958E-17 | 0,999999986 | 1 |
| EZR | 25 | 25 | -0,012491135 | 0,54363871 | 1 |
| PRKACA | 21 | 19 | 0 | 1 | 1 |
| PRKACB | 8 | 7 | 0 | 1 | 1 |
| ATP1B1 | 19 | 19 | 0,044310232 | 0,140657109 | 1 |
| TMEM38A | 1 | 1 | 0 | 1 | 1 |
| 4932415D10RIK | 3 | 3 | 0 | 1 | 1 |
| PABPC1 | 30 | 28 | -5,32438E-15 | 0,999999618 | 1 |
| SNCA | 8 | 8 | 0,081098979 | 0,316443683 | 1 |
| INPP4A | 13 | 13 | 0,02390729 | 0,285667274 | 1 |
| EIF3C | 28 | 22 | 2,82048E-18 | 0,999999999 | 1 |
| REEP5 | 4 | 4 | -0,003836199 | 0,799588752 | 1 |
| SARS | 30 | 30 | 0,033994124 | 0,247765633 | 1 |
| GLS | 31 | 31 | 0 | 1 | 1 |
| DLST | 21 | 20 | -1,39622E-15 | 0,999999903 | 1 |
| GFM2 | 6 | 6 | -0,033415714 | 0,363024818 | 1 |
| KIAA1107 | 19 | 19 | 0 | 1 | 1 |
| FAM217B | 2 | 2 | 0 | 1 | 1 |
| AS3MT | 7 | 7 | 0 | 1 | 1 |
| SUCLG1 | 14 | 14 | 0 | 1 | 1 |
| PHPT1 | 8 | 5 | 0 | 1 | 1 |
| RPS19 | 12 | 11 | 0,048673285 | 0,2735187 | 1 |
| ARPC5 | 6 | 6 | 4,64432E-17 | 0,999999993 | 1 |
| SCRIB | 6 | 6 | 1,60548E-17 | 0,999999992 | 1 |

| | | | | | |
|---|---|---|---|---|---|
| UAP1 | 6 | 6 | 0 | 1 | 1 |
| ARMC10 | 9 | 9 | 2,78816E-18 | 0,999999998 | 1 |
| GCN1 | 32 | 27 | 6,77488E-16 | 0,999999933 | 1 |
| SRPRA | 6 | 6 | 0,044938589 | 0,314725946 | 1 |
| SLC29A1 | 2 | 2 | 0,094873509 | 0,165052604 | 1 |
| GRPEL1 | 10 | 10 | -1,78942E-16 | 0,999999979 | 1 |
| THOP1 | 24 | 23 | 0 | 1 | 1 |
| CRPPA | 6 | 6 | 0 | 1 | 1 |
| DYNC1I1 | 16 | 16 | 0 | 1 | 1 |
| GSS | 18 | 17 | -3,89822E-18 | 0,999999994 | 1 |
| RHEB | 14 | 13 | 1,04026E-16 | 0,99999997 | 1 |
| FAR1 | 4 | 4 | 0 | 1 | 1 |
| CSPG5 | 9 | 9 | 0 | 1 | 1 |
| KLC4 | 7 | 7 | 3,1415E-17 | 0,999999985 | 1 |
| ECHDC2 | 2 | 2 | 1,01677E-16 | 0,999999982 | 1 |
| ESYT2 | 10 | 10 | 2,28076E-17 | 0,999999985 | 1 |
| BCS1L | 18 | 18 | -9,19598E-17 | 0,999999976 | 1 |
| SH2B1 | 7 | 6 | 0 | 1 | 1 |
| P4HB | 29 | 29 | -9,48798E-05 | 0,955228701 | 1 |
| SPR | 16 | 16 | 2,29475E-14 | 0,999999328 | 1 |
| TXNDC5 | 9 | 9 | 1,12301E-17 | 0,999999993 | 1 |
| CTBP1;CTBP2 | 5 | 5 | 6,51481E-17 | 0,999999979 | 1 |
| GLUD1 | 47 | 47 | -0,10871101 | 0,209306475 | 1 |
| TSC22D4 | 1 | 1 | 0 | 1 | 1 |
| ATL3 | 6 | 6 | 0 | 1 | 1 |
| FAHD2A | 15 | 15 | -1,33427E-16 | 0,999999972 | 1 |
| FAM171A2 | 16 | 16 | -0,010751245 | 0,499582907 | 1 |
| XPOT | 9 | 8 | -3,07203E-17 | 0,999999989 | 1 |
| CST3 | 9 | 7 | -2,46841E-14 | 0,999999602 | 1 |
| PRAF2 | 5 | 5 | 0 | 1 | 1 |
| IKBKB | 2 | 2 | 0 | 1 | 1 |
| PSMC2 | 27 | 26 | -4,65642E-16 | 0,999999922 | 1 |
| SCN2A | 21 | 21 | 1,34544E-16 | 0,999999968 | 1 |

| | | | | | |
|---|---|---|---|---|---|
| MAPK3 | 17 | 16 | 1,39602E-17 | 0,999999991 | 1 |
| UQCRB | 12 | 12 | 0 | 1 | 1 |
| GUSB | 1 | 1 | 0 | 1 | 1 |
| MINDY3 | 2 | 2 | 0,006290695 | 0,842280811 | 1 |
| PFKFB2 | 5 | 5 | 0,022727037 | 0,590717896 | 1 |
| ADGRB1 | 13 | 13 | -0,048285291 | 0,230655039 | 1 |
| STIM1 | 6 | 6 | 1,50103E-17 | 0,999999993 | 1 |
| SCRN3 | 16 | 15 | 2,39485E-18 | 0,999999996 | 1 |
| RBP1 | 6 | 5 | -0,038911576 | 0,325682794 | 1 |
| APEH | 16 | 16 | 0 | 1 | 1 |
| PCDHB10 | 1 | 1 | 0 | 1 | 1 |
| SPIRE1 | 3 | 3 | 6,00105E-17 | 0,999999986 | 1 |
| INA | 29 | 29 | -0,079860278 | 0,353698851 | 1 |
| EXOC7 | 27 | 27 | 1,32676E-16 | 0,999999949 | 1 |
| IDH1 | 31 | 31 | 0,012910389 | 0,581831075 | 1 |
| KRT42 | 4 | 4 | 2,14839E-17 | 0,999999996 | 1 |
| KRT14;KRT17 | 7 | 5 | 3,86508E-18 | 0,999999998 | 1 |
| RNH1 | 12 | 12 | 6,22837E-17 | 0,999999981 | 1 |
| TPPP3 | 5 | 5 | -2,23886E-18 | 0,999999996 | 1 |
| STMN2 | 4 | 4 | -0,054244722 | 0,184325539 | 1 |
| STMN3 | 9 | 9 | -0,031076984 | 0,382410366 | 1 |
| PUM1 | 1 | 1 | 0 | 1 | 1 |
| DYNC2H1 | 3 | 3 | -0,035637211 | 0,458750849 | 1 |
| CTPS1 | 12 | 12 | 0,045006856 | 0,203166607 | 1 |
| AGL | 43 | 42 | -0,020944413 | 0,328551078 | 1 |
| POGK | 1 | 1 | 0 | 1 | 1 |
| CCT7 | 32 | 30 | 0,02659091 | 0,28110586 | 1 |
| EPS15L1 | 34 | 34 | -0,039721992 | 0,322499495 | 1 |
| STUB1 | 11 | 11 | -2,72751E-17 | 0,999999981 | 1 |
| PRR7 | 2 | 2 | 0 | 1 | 1 |
| MSN | 20 | 20 | 2,21603E-16 | 0,999999975 | 1 |
| PEX5 | 3 | 3 | -1,78025E-17 | 0,999999991 | 1 |
| NCL | 24 | 14 | 3,11258E-16 | 0,999999972 | 1 |

| | | | | | |
|---|---|---|---|---|---|
| BRSK1 | 21 | 21 | 0,01952749 | 0,420868737 | 1 |
| LCP1 | 23 | 23 | 1,59207E-17 | 0,999999993 | 1 |
| PRODH | 16 | 16 | -0,10158875 | 0,268238601 | 1 |
| PDIA3 | 34 | 33 | 3,70346E-17 | 0,999999985 | 1 |
| ZFYVE1 | 10 | 10 | 0,023969769 | 0,409933678 | 1 |
| AVEN | 1 | 1 | 0 | 1 | 1 |
| KIF5C | 46 | 46 | -1,56918E-17 | 0,999999989 | 1 |
| KIF5C;KIF5B | 3 | 3 | 1,66931E-18 | 0,999999997 | 1 |
| RUVBL1 | 12 | 12 | 3,47809E-17 | 0,999999979 | 1 |
| PREX1 | 14 | 14 | -0,064861173 | 0,152237406 | 1 |
| RPTOR | 14 | 14 | 8,45849E-18 | 0,999999991 | 1 |
| NUMA1 | 1 | 1 | -0,044075003 | 0,536900701 | 1 |
| GTPBP3 | 2 | 2 | 3,55664E-15 | 0,999999861 | 1 |
| CIAO2A | 1 | 1 | 0 | 1 | 1 |
| SART3 | 17 | 1 | 0 | 1 | 1 |
| CACNB3;CACNB2;CA | 4 | 4 | 1,49409E-17 | 0,99999999 | 1 |
| CRK | 21 | 20 | 5,58616E-17 | 0,99999998 | 1 |
| ABI2 | 7 | 7 | 0 | 1 | 1 |
| EMC2 | 5 | 5 | 8,8066E-17 | 0,999999965 | 1 |
| SCN11A | 1 | 1 | 0 | 1 | 1 |
| COQ6 | 13 | 13 | -3,09E-18 | 1 | 1 |
| CATSPER1 | 1 | 1 | 0 | 1 | 1 |
| FTL1 | 11 | 10 | 0 | 1 | 1 |
| MRPL41 | 3 | 3 | -0,059848212 | 0,369003637 | 1 |
| ABCB8 | 21 | 21 | -0,008189678 | 0,719415441 | 1 |
| ETL4 | 31 | 31 | -0,024944142 | 0,391110353 | 1 |
| ABHD5 | 2 | 2 | 0,118027211 | 0,123501889 | 1 |
| BCAN | 32 | 23 | -6,36265E-05 | 0,970967125 | 1 |
| BABAM1 | 3 | 3 | -5,6893E-16 | 0,999999943 | 1 |
| PPM1A | 17 | 17 | 2,40195E-16 | 0,999999956 | 1 |
| PPM1B | 11 | 10 | -0,040544005 | 0,315728401 | 1 |
| CS | 29 | 29 | -3,87677E-16 | 0,999999973 | 1 |
| GCAT | 6 | 6 | -0,073444332 | 0,232719757 | 1 |

| | | | | | |
|---|---|---|---|---|---|
| GPR162 | 3 | 3 | -0,003109798 | 0,809730715 | 1 |
| PTGS1 | 3 | 3 | 0 | 1 | 1 |
| PI4KA | 68 | 68 | 0,03673251 | 0,186821803 | 1 |
| ACACB | 8 | 8 | 0,066733681 | 0,139278981 | 1 |
| SERPINE2 | 10 | 10 | 3,70884E-18 | 0,999999997 | 1 |
| GPD1L | 16 | 16 | -0,004663059 | 0,696193542 | 1 |
| KLC1 | 19 | 19 | 0 | 1 | 1 |
| FKBP5 | 12 | 11 | 0 | 1 | 1 |
| ATPAF1 | 9 | 9 | -0,015450788 | 0,486935389 | 1 |
| SVOP | 6 | 6 | 1,06859E-17 | 0,999999995 | 1 |
| AKR1A1 | 27 | 24 | -3,4405E-18 | 0,999999995 | 1 |
| PPP1R9A | 1 | 1 | 0 | 1 | 1 |
| RAP1GDS1 | 31 | 31 | -2,32926E-17 | 0,999999986 | 1 |
| RAP1GDS1 | 3 | 3 | -2,09385E-17 | 0,999999997 | 1 |
| LRP4 | 2 | 2 | 0 | 1 | 1 |
| GNL1 | 13 | 13 | 1,38331E-14 | 0,99999977 | 1 |
| VWA5B2 | 1 | 1 | -0,001543004 | 0,924060057 | 1 |
| NOMO1 | 27 | 27 | 4,262E-17 | 0,999999985 | 1 |
| MYL6B | 2 | 2 | -1,00071E-10 | 0,999969535 | 1 |
| FABP7 | 8 | 8 | -0,067448052 | 0,393445302 | 1 |
| CSL | 3 | 3 | 0 | 1 | 1 |
| AKR1B8 | 2 | 2 | -0,036339098 | 0,396684761 | 1 |
| LETM1 | 31 | 31 | -1,25118E-15 | 0,999999921 | 1 |
| SERPINI1 | 6 | 6 | 0 | 1 | 1 |
| OTUB2 | 5 | 2 | 0 | 1 | 1 |
| GLOD4 | 30 | 30 | -0,030451551 | 0,302235428 | 1 |
| SLIRP | 3 | 3 | -5,33196E-19 | 0,999999999 | 1 |
| DARS1 | 31 | 28 | 9,44899E-17 | 0,999999975 | 1 |
| LRRC8A | 17 | 17 | 0 | 1 | 1 |
| LRFN5 | 5 | 5 | 0,035206449 | 0,446306689 | 1 |
| IPO13 | 4 | 2 | -1,03172E-18 | 0,999999999 | 1 |
| IQCB1 | 5 | 5 | 0 | 1 | 1 |
| MYRIP | 4 | 4 | -1,93251E-16 | 0,999999973 | 1 |

| | | | | | |
|---|---|---|---|---|---|
| CD200 | 10 | 10 | 6,24272E-18 | 0,999999995 | 1 |
| SUCLA2 | 43 | 43 | 0 | 1 | 1 |
| LYPLA1 | 3 | 3 | 0 | 1 | 1 |
| sp\|Q8R092\|CA043_ | 3 | 3 | -1,48221E-15 | 0,999999927 | 1 |
| SNX5 | 14 | 13 | 0 | 1 | 1 |
| LYPLA2 | 9 | 9 | 0,045696554 | 0,27890087 | 1 |
| PLCL2 | 19 | 19 | 0,043790163 | 0,206569804 | 1 |
| ABI1 | 13 | 13 | 0,017264854 | 0,374415148 | 1 |
| MGLL | 20 | 20 | 0,039019163 | 0,444433412 | 1 |
| NME3 | 8 | 8 | 1,95866E-16 | 0,999999963 | 1 |
| SLC25A11 | 23 | 23 | 0 | 1 | 1 |
| ENDOD1 | 12 | 11 | -4,77229E-17 | 0,999999972 | 1 |
| GPS1 | 13 | 13 | 0,041908216 | 0,218545366 | 1 |
| SLC25A19 | 6 | 6 | -0,050653896 | 0,291747285 | 1 |
| ARMCX3 | 7 | 7 | -0,005521656 | 0,712321871 | 1 |
| SVIL | 1 | 1 | 0 | 1 | 1 |
| STAMBPL1 | 5 | 5 | 4,41189E-18 | 0,999999995 | 1 |
| ACSL5 | 8 | 8 | 4,51914E-17 | 0,999999986 | 1 |
| HSPA9 | 54 | 54 | -3,58167E-15 | 0,999999883 | 1 |
| TMEM33 | 4 | 4 | 1,03073E-17 | 0,999999994 | 1 |
| SARS2 | 10 | 10 | -1,99157E-17 | 0,999999995 | 1 |
| CAP1 | 36 | 34 | 1,4221E-16 | 0,999999983 | 1 |
| CHAT | 6 | 6 | 1,7932E-16 | 0,999999967 | 1 |
| NMRAL1 | 15 | 14 | -1,13859E-17 | 0,999999993 | 1 |
| ADAP1 | 20 | 19 | 0,024142534 | 0,424510527 | 1 |
| EEF2 | 46 | 42 | 0,067065476 | 0,220721961 | 1 |
| DMXL1 | 15 | 15 | 0 | 1 | 1 |
| HAGH | 11 | 11 | -0,003927199 | 0,725924758 | 1 |
| HSP90AA1 | 39 | 37 | 0,045317012 | 0,150659299 | 1 |
| FUT11 | 1 | 1 | 0 | 1 | 1 |
| MFN2 | 31 | 31 | -1,03722E-16 | 0,999999974 | 1 |
| ELP5 | 2 | 2 | 1,42859E-18 | 0,999999999 | 1 |
| KCNMB4 | 1 | 1 | 0 | 1 | 1 |

| | | | | | |
|---|---|---|---|---|---|
| ANLN | 6 | 6 | -0,038162263 | 0,535915344 | 1 |
| FAM20B | 3 | 2 | -0,078814679 | 0,166381582 | 1 |
| ACTN4 | 23 | 22 | -0,050905227 | 0,196196272 | 1 |
| PNMA8B | 1 | 1 | 0 | 1 | 1 |
| ERO1A | 11 | 11 | -8,24221E-16 | 0,999999883 | 1 |
| VTI1B | 8 | 7 | -3,52272E-17 | 0,99999999 | 1 |
| VPS16 | 11 | 11 | 0 | 1 | 1 |
| PLD3 | 5 | 5 | 0 | 1 | 1 |
| HNRNPUL2 | 29 | 6 | 0 | 1 | 1 |
| PDP1 | 15 | 15 | -1,01653E-16 | 0,999999977 | 1 |
| DHRS7B | 11 | 11 | 4,54753E-17 | 0,999999974 | 1 |
| PHF24 | 15 | 15 | 0,054496297 | 0,273726012 | 1 |
| VPS13D | 15 | 15 | 0 | 1 | 1 |
| RO60 | 10 | 10 | -9,92294E-17 | 0,999999981 | 1 |
| DNM1;DNM3 | 5 | 5 | 1,95397E-15 | 0,999999838 | 1 |
| PLEKHG5 | 7 | 7 | -1,17598E-16 | 0,999999973 | 1 |
| HEXA | 2 | 2 | -3,72801E-20 | 1 | 1 |
| CYB5R4 | 1 | 1 | 0 | 1 | 1 |
| ABCA7 | 3 | 3 | 2,34481E-16 | 0,999999976 | 1 |
| GAK | 25 | 25 | 0 | 1 | 1 |
| IPO11 | 8 | 7 | 2,87624E-12 | 0,999997819 | 1 |
| ANXA3 | 19 | 19 | -2,15506E-17 | 0,999999995 | 1 |
| GBF1 | 6 | 4 | 0 | 1 | 1 |
| PDE1B | 18 | 17 | -2,17341E-17 | 0,999999989 | 1 |
| ABHD16A | 17 | 17 | -3,46506E-18 | 0,999999999 | 1 |
| UBR1 | 10 | 9 | 0,006647395 | 0,713255454 | 1 |
| HSPA4L | 52 | 51 | -0,009756486 | 0,338156657 | 1 |
| ETFDH | 24 | 24 | -0,001511593 | 0,893620132 | 1 |
| NADK2 | 23 | 23 | -2,58588E-15 | 0,99999988 | 1 |
| ANK2 | 3 | 3 | -0,073493527 | 0,169415671 | 1 |
| MAP2K4 | 13 | 13 | 0 | 1 | 1 |
| NDUFAF5 | 6 | 6 | -3,87352E-17 | 0,999999989 | 1 |
| COG2 | 2 | 1 | 1,42918E-21 | 1 | 1 |

| | | | | | |
|---|---|---|---|---|---|
| EIF5 | 10 | 10 | 0 | 1 | 1 |
| TMEM245 | 1 | 1 | 0 | 1 | 1 |
| REV3L | 1 | 1 | 0 | 1 | 1 |
| PRDX5 | 15 | 15 | -1,08463E-14 | 0,999999682 | 1 |
| TTC5 | 2 | 2 | -2,53728E-16 | 0,999999981 | 1 |
| PLCXD3 | 8 | 6 | 0,090779633 | 0,127734027 | 1 |
| ADGRA1 | 2 | 2 | -3,52937E-18 | 0,999999997 | 1 |
| TMEM35A | 3 | 3 | -1,48367E-17 | 0,999999995 | 1 |
| OCIAD2 | 6 | 6 | 0 | 1 | 1 |
| PGAM2 | 7 | 5 | 1,28934E-17 | 0,999999989 | 1 |
| PGAM1 | 17 | 17 | 0 | 1 | 1 |
| RIN1 | 17 | 17 | 0,065800306 | 0,134804842 | 1 |
| CSAD | 8 | 7 | 0 | 1 | 1 |
| ELMO2 | 21 | 21 | 0 | 1 | 1 |
| SNX30 | 16 | 16 | 0 | 1 | 1 |
| SLC4A3 | 9 | 9 | 0,037913023 | 0,417444934 | 1 |
| sp\|Q9CWB7\|YD286_ | 1 | 1 | -2,17258E-19 | 1 | 1 |
| CACNA1E | 18 | 18 | 3,10833E-18 | 0,999999996 | 1 |
| LAMP1 | 5 | 5 | 5,27426E-15 | 0,999999807 | 1 |
| DDAH2 | 9 | 9 | 0 | 1 | 1 |
| GRIPAP1 | 2 | 2 | 4,21753E-15 | 0,999999906 | 1 |
| CDC42BPB | 31 | 31 | 0 | 1 | 1 |
| EXOC1 | 10 | 10 | 2,11461E-17 | 0,999999982 | 1 |
| CARS1 | 27 | 21 | 0 | 1 | 1 |
| CCT6A | 20 | 20 | 6,72494E-18 | 0,999999994 | 1 |
| IGTP | 1 | 1 | 0,001479954 | 0,922552556 | 1 |
| SLC4A1AP | 1 | 1 | 0 | 1 | 1 |
| CYP2D9 | 1 | 1 | 0 | 1 | 1 |
| STAM2 | 5 | 5 | 0 | 1 | 1 |
| GAPVD1 | 22 | 22 | 8,43698E-15 | 0,999999855 | 1 |
| COG1 | 4 | 3 | 1,34869E-16 | 0,999999987 | 1 |
| TMEM126A | 5 | 5 | 0 | 1 | 1 |
| MVP | 3 | 3 | 0 | 1 | 1 |

| | | | | | |
|---|---|---|---|---|---|
| TMEM205 | 2 | 2 | -1,21483E-15 | 0,999999941 | 1 |
| IQGAP1 | 16 | 14 | 0 | 1 | 1 |
| PSMD5 | 15 | 14 | 0 | 1 | 1 |
| EIF4G3 | 24 | 21 | -1,22534E-16 | 0,999999966 | 1 |
| AKR7A2 | 12 | 11 | -3,15694E-17 | 0,999999985 | 1 |
| ERC2 | 45 | 45 | 0,005691076 | 0,702940952 | 1 |
| GDE1 | 5 | 5 | -7,53623E-17 | 0,999999977 | 1 |
| RTL8B | 1 | 1 | -1,6237E-19 | 1 | 1 |
| HSDL1 | 9 | 9 | 0 | 1 | 1 |
| CBR1 | 22 | 22 | -1,55412E-16 | 0,999999952 | 1 |
| PSMA1 | 19 | 18 | 0,008477887 | 0,586622198 | 1 |
| CHMP1A | 4 | 4 | 5,12192E-18 | 0,999999996 | 1 |
| SHMT1 | 2 | 2 | -1,09335E-18 | 0,999999999 | 1 |
| SLC18A2 | 1 | 1 | 0,036104859 | 0,477555274 | 1 |
| NDUFS1 | 55 | 55 | 0 | 1 | 1 |
| LRRC8B | 5 | 5 | -1,06184E-14 | 0,999999885 | 1 |
| PRKCE | 39 | 39 | 0,0283082 | 0,465251128 | 1 |
| SEMA4A | 7 | 7 | 0 | 1 | 1 |
| OTUD6B | 6 | 5 | -5,76475E-18 | 0,999999994 | 1 |
| CDC23 | 7 | 6 | 0,072228275 | 0,1698065 | 1 |
| MCF2L | 5 | 5 | -0,006151876 | 0,772618972 | 1 |
| PRRC2B | 9 | 5 | -9,50224E-17 | 0,999999982 | 1 |
| MMS19 | 9 | 8 | 0 | 1 | 1 |
| UBQLN1 | 6 | 6 | 0 | 1 | 1 |
| UBQLN4 | 4 | 4 | 0 | 1 | 1 |
| TMEM214 | 1 | 1 | 0 | 1 | 1 |
| PCDH17 | 7 | 7 | 9,24728E-19 | 0,999999999 | 1 |
| LETMD1 | 9 | 9 | -0,006649908 | 0,78631258 | 1 |
| SNX2 | 24 | 23 | -0,010313374 | 0,531723539 | 1 |
| GIGYF2 | 3 | 2 | -1,03808E-18 | 0,999999999 | 1 |
| GIGYF1 | 2 | 2 | 1,53694E-14 | 0,9999997 | 1 |
| RIPOR1 | 10 | 10 | 0 | 1 | 1 |
| OXSM | 11 | 11 | -2,70284E-16 | 0,999999964 | 1 |

| | | | | | |
|---|---|---|---|---|---|
| PDK2 | 12 | 12 | 0 | 1 | 1 |
| PDK1 | 10 | 10 | 0 | 1 | 1 |
| HEATR5B | 11 | 11 | 0 | 1 | 1 |
| EEFSEC | 3 | 3 | 0,067121131 | 0,22143099 | 1 |
| ARFIP2 | 14 | 14 | -1,83682E-15 | 0,99999986 | 1 |
| ZZEF1 | 18 | 14 | 1,00974E-05 | 0,984694124 | 1 |
| FICD | 1 | 1 | 0 | 1 | 1 |
| ESYT1 | 2 | 2 | 0 | 1 | 1 |
| EPN3 | 5 | 5 | -7,06973E-17 | 0,99999999 | 1 |
| MYADM | 4 | 4 | 0,05689904 | 0,253879676 | 1 |
| NQO2 | 6 | 5 | 7,63272E-15 | 0,999999738 | 1 |
| SMPD1 | 4 | 4 | 3,24713E-16 | 0,999999956 | 1 |
| SRI | 6 | 4 | -6,15354E-19 | 0,999999999 | 1 |
| ME2 | 15 | 15 | -2,30663E-16 | 0,999999972 | 1 |
| TUBB4B;TUBB6;TUBI | 2 | 2 | -1,34445E-14 | 0,999999849 | 1 |
| TUBB2A;TUBB2B | 4 | 4 | 3,19717E-16 | 0,999999947 | 1 |
| TUBB5 | 7 | 6 | -3,20354E-17 | 0,999999991 | 1 |
| COX6A1 | 3 | 3 | 2,77934E-17 | 0,999999996 | 1 |
| RPS16 | 10 | 9 | 0,03286989 | 0,597858417 | 1 |
| CNPY2 | 5 | 5 | 4,02089E-17 | 0,999999979 | 1 |
| CRELD1 | 1 | 1 | 0 | 1 | 1 |
| MRRF | 9 | 9 | 0 | 1 | 1 |
| MYG1 | 12 | 9 | 1,63599E-18 | 0,999999998 | 1 |
| TTYH3 | 9 | 9 | -3,01828E-17 | 0,999999991 | 1 |
| GUF1 | 3 | 3 | -0,02112668 | 0,623074765 | 1 |
| PIP5K1A | 10 | 9 | 0 | 1 | 1 |
| TNPO1;TNPO2 | 5 | 5 | -1,71998E-17 | 0,999999992 | 1 |
| FBXL15 | 3 | 3 | 7,88797E-18 | 0,999999994 | 1 |
| APOA4 | 11 | 2 | -8,67412E-17 | 0,999999997 | 1 |
| ADPRS | 10 | 9 | 0 | 1 | 1 |
| DDB1 | 54 | 52 | 2,59704E-16 | 0,999999944 | 1 |
| ARHGEF17 | 12 | 12 | -1,12553E-17 | 0,999999995 | 1 |
| EPDR1 | 8 | 8 | 0,012828658 | 0,646460778 | 1 |

| | | | | | |
|---|---|---|---|---|---|
| CRACDL | 15 | 14 | -6,01328E-05 | 0,973579296 | 1 |
| SNAP91 | 27 | 27 | -1,51252E-17 | 0,999999997 | 1 |
| SYT11 | 8 | 8 | -0,022430165 | 0,408840204 | 1 |
| MECR | 10 | 10 | -1,60669E-17 | 0,999999991 | 1 |
| GNAS | 7 | 7 | 0 | 1 | 1 |
| GNA13 | 19 | 18 | -1,59337E-17 | 0,999999989 | 1 |
| PTPRS | 36 | 36 | 0,03402672 | 0,121200963 | 1 |
| SH3GL2 | 24 | 24 | 0,047357556 | 0,2679826 | 1 |
| GRM5 | 26 | 26 | 1,03766E-16 | 0,999999984 | 1 |
| FDPS | 11 | 11 | 0 | 1 | 1 |
| NEFL | 34 | 34 | -0,008672902 | 0,797738794 | 1 |
| DAB2IP | 8 | 8 | -2,06832E-16 | 0,999999961 | 1 |
| STXBP3 | 9 | 9 | 2,3839E-17 | 0,999999993 | 1 |
| PAK2 | 21 | 18 | 3,31165E-17 | 0,999999985 | 1 |
| PAK3 | 11 | 10 | 0,060221284 | 0,175604992 | 1 |
| RSU1 | 8 | 8 | -1,9079E-16 | 0,999999963 | 1 |
| GPSM2 | 3 | 3 | -0,036548672 | 0,461817556 | 1 |
| SERPINA3K | 23 | 8 | 6,13763E-16 | 0,999999957 | 1 |
| SLC25A3 | 25 | 25 | 0 | 1 | 1 |
| OSTF1 | 4 | 4 | 1,20648E-16 | 0,999999983 | 1 |
| PHLPP1 | 2 | 2 | 0,055897833 | 0,40878581 | 1 |
| NCLN | 13 | 13 | -1,94362E-16 | 0,999999958 | 1 |
| GRM3 | 22 | 22 | 0 | 1 | 1 |
| OPCML | 15 | 15 | 0,046173748 | 0,184416452 | 1 |
| COPS3 | 16 | 16 | 1,23468E-15 | 0,999999845 | 1 |
| PLXND1 | 9 | 9 | -2,74661E-17 | 0,999999983 | 1 |
| GNAO1 | 22 | 22 | 0,018813125 | 0,561666187 | 1 |
| GNAQ | 21 | 20 | 1,14435E-16 | 0,999999967 | 1 |
| TRIO | 31 | 31 | 0 | 1 | 1 |
| PGAM2;PGAM1 | 6 | 5 | 0 | 1 | 1 |
| AP1G1 | 19 | 19 | 9,22703E-18 | 0,999999994 | 1 |
| OSBPL1A | 13 | 11 | -1,86664E-21 | 1 | 1 |
| GABBR2 | 21 | 21 | 6,42902E-16 | 0,999999888 | 1 |

| | | | | | |
|---|---|---|---|---|---|
| PSMC3 | 28 | 25 | 9,25567E-17 | 0,999999967 | 1 |
| PFKP | 34 | 32 | 0,054033013 | 0,163131863 | 1 |
| SORD | 13 | 12 | -0,022320956 | 0,468573452 | 1 |
| DLGAP1 | 16 | 16 | 3,2254E-18 | 0,999999994 | 1 |
| GNAI2 | 14 | 14 | 0,074849587 | 0,158109203 | 1 |
| NAA10 | 3 | 2 | 0,116295711 | 0,236630842 | 1 |
| NAA12 | 1 | 1 | 0,141045848 | 0,301627676 | 1 |
| YARS | 36 | 35 | 1,51574E-15 | 0,999999842 | 1 |
| RBM3 | 6 | 6 | 0 | 1 | 1 |
| ACY1 | 12 | 12 | -2,13891E-17 | 0,999999987 | 1 |
| VAPB | 13 | 13 | 0 | 1 | 1 |
| LARS2 | 10 | 10 | -0,079104618 | 0,186104646 | 1 |
| TMEM135 | 1 | 1 | 0 | 1 | 1 |
| HSPA2;HSPA8 | 9 | 9 | 1,27788E-06 | 0,994712999 | 1 |
| EPN1 | 15 | 15 | 0 | 1 | 1 |
| DECR1 | 11 | 11 | -0,096329562 | 0,145072658 | 1 |
| S100B | 4 | 3 | 0 | 1 | 1 |
| AK2 | 9 | 9 | -2,33123E-16 | 0,999999974 | 1 |
| CISD1 | 9 | 9 | 1,30905E-17 | 0,99999999 | 1 |
| EEF1E1 | 8 | 7 | 0,026210733 | 0,546939325 | 1 |
| LRRFIP1;LRRFIP2 | 2 | 1 | 0,074236354 | 0,29872929 | 1 |
| HDHD2 | 13 | 12 | 0,020873383 | 0,446483329 | 1 |
| NEBL | 16 | 15 | 6,20574E-18 | 0,999999997 | 1 |
| NDUFB9 | 12 | 12 | -3,12289E-16 | 0,999999948 | 1 |
| ANXA7 | 24 | 23 | 0,007545628 | 0,573804454 | 1 |
| IAP | 2 | 2 | -0,084867628 | 0,269923421 | 1 |
| GABRA4 | 5 | 5 | 2,11671E-17 | 0,999999988 | 1 |
| PLPPR4 | 24 | 23 | 1,93984E-17 | 0,99999999 | 1 |
| MTX2 | 7 | 7 | 0 | 1 | 1 |
| PIK3R4 | 19 | 19 | 1,07086E-14 | 0,999999585 | 1 |
| DLG4 | 33 | 33 | 0,012642764 | 0,499734546 | 1 |
| TBC1D15 | 8 | 8 | -2,49269E-17 | 0,999999984 | 1 |
| DUSP3 | 8 | 8 | 1,89936E-15 | 0,999999908 | 1 |

| | | | | | |
|---|---|---|---|---|---|
| PRDX3 | 13 | 13 | 0 | 1 | 1 |
| ALPL | 1 | 1 | 0,067640603 | 0,304070958 | 1 |
| CDC16 | 4 | 4 | 0 | 1 | 1 |
| ANKRD29 | 4 | 4 | -1,41664E-19 | 1 | 1 |
| CLIP1 | 35 | 34 | -2,47172E-16 | 0,999999975 | 1 |
| TOMM40L | 6 | 6 | 1,79977E-16 | 0,999999971 | 1 |
| TANC1 | 17 | 17 | 3,95877E-17 | 0,999999977 | 1 |
| AFG1L | 8 | 8 | -0,077477019 | 0,266980643 | 1 |
| SETD7 | 11 | 9 | 0,008462384 | 0,666126873 | 1 |
| LACTB2 | 4 | 4 | 0,056513548 | 0,297935303 | 1 |
| POMGNT2 | 2 | 2 | 5,84395E-16 | 0,999999977 | 1 |
| PAM | 10 | 10 | -7,96872E-17 | 0,999999978 | 1 |
| MBLAC2 | 11 | 11 | -6,14493E-17 | 0,999999976 | 1 |
| LAP3 | 23 | 23 | -0,008501368 | 0,639343137 | 1 |
| PDHX | 20 | 20 | -4,36851E-18 | 0,999999995 | 1 |
| LNPEP | 16 | 16 | -9,07629E-18 | 0,99999999 | 1 |
| EPS8 | 11 | 11 | -0,017255663 | 0,585142539 | 1 |
| ARMC8 | 7 | 7 | 0 | 1 | 1 |
| ATG4B | 7 | 7 | 0,095440933 | 0,319272047 | 1 |
| SLC18A3 | 2 | 2 | 3,7965E-18 | 0,999999996 | 1 |
| CKAP5 | 51 | 51 | -0,005717689 | 0,621686484 | 1 |
| OSBPL8 | 11 | 11 | 0 | 1 | 1 |
| SEC22B | 16 | 15 | 8,4505E-18 | 0,999999992 | 1 |
| CR2 | 1 | 1 | -0,164615588 | 0,133169393 | 1 |
| MT-ND3 | 1 | 1 | 0 | 1 | 1 |
| MTND5 | 6 | 6 | 0 | 1 | 1 |
| CALB2 | 22 | 20 | -0,000178828 | 0,959130648 | 1 |
| HECW2 | 10 | 10 | 0 | 1 | 1 |
| MAP1S | 16 | 16 | -3,398E-16 | 0,999999938 | 1 |
| OGA | 17 | 17 | 0,063373214 | 0,133986175 | 1 |
| GAD1;GAD2 | 1 | 1 | -0,012143511 | 0,709491586 | 1 |
| HEXB | 19 | 18 | 0 | 1 | 1 |
| SYN3 | 14 | 14 | -4,82363E-16 | 0,999999925 | 1 |

| | | | | | |
|---|---|---|---|---|---|
| ASTN2 | 7 | 7 | -3,68714E-18 | 0,999999999 | 1 |
| ATP5PD | 18 | 18 | -2,1523E-18 | 0,999999997 | 1 |
| TBL2 | 3 | 3 | 0 | 1 | 1 |
| PLCB4 | 3 | 3 | -6,4708E-19 | 0,999999999 | 1 |
| ACO1 | 26 | 26 | 0 | 1 | 1 |
| RIMS1 | 9 | 9 | 0 | 1 | 1 |
| CRYM | 21 | 20 | 0,048545242 | 0,199013095 | 1 |
| PGBD5 | 14 | 12 | 0 | 1 | 1 |
| HCN1 | 13 | 13 | 1,19067E-17 | 0,999999996 | 1 |
| MAN2C1 | 17 | 17 | -1,28195E-18 | 0,999999999 | 1 |
| PTRH2 | 5 | 5 | 0 | 1 | 1 |
| SLC35B2 | 3 | 3 | -1,44998E-18 | 0,999999998 | 1 |
| EZR;MSN | 9 | 9 | 0,023212474 | 0,496379423 | 1 |
| FADS1 | 4 | 4 | -3,63628E-17 | 0,999999986 | 1 |
| ECH1 | 7 | 7 | -7,83647E-17 | 0,999999978 | 1 |
| ACTR1B | 10 | 10 | 0,048157334 | 0,132243269 | 1 |
| sp\|Q9CYS6\|CB072_I | 2 | 2 | -8,28443E-18 | 0,999999997 | 1 |
| BMP2K | 2 | 2 | 0,047680858 | 0,415932271 | 1 |
| MAPK8 | 4 | 4 | 5,3107E-18 | 0,999999994 | 1 |
| PSMB2 | 11 | 11 | 0,03410544 | 0,353069465 | 1 |
| SLC4A10 | 16 | 15 | 1,49419E-17 | 0,999999994 | 1 |
| TENM4 | 32 | 32 | 1,21343E-16 | 0,999999972 | 1 |
| ADGRG1 | 5 | 5 | 0 | 1 | 1 |
| ALCAM | 24 | 24 | 3,58336E-16 | 0,999999932 | 1 |
| PREB | 6 | 6 | 0 | 1 | 1 |
| FXYD6 | 3 | 3 | 0,09188909 | 0,182474804 | 1 |
| HECTD1 | 21 | 21 | -1,43532E-15 | 0,999999878 | 1 |
| NDUFS2 | 27 | 27 | -1,90206E-16 | 0,999999986 | 1 |
| PAIP1 | 1 | 1 | 0 | 1 | 1 |
| JPH4 | 3 | 3 | -1,77998E-17 | 0,999999998 | 1 |
| LRRC47 | 17 | 17 | 0 | 1 | 1 |
| RPL18 | 6 | 5 | 0,155909309 | 0,354397382 | 1 |
| DUSP28 | 1 | 1 | 0 | 1 | 1 |

| | | | | | |
|---|---|---|---|---|---|
| XPNPEP3 | 5 | 5 | 2,7802E-17 | 0,999999988 | 1 |
| ALG2 | 13 | 13 | -7,7578E-17 | 0,999999983 | 1 |
| DDX17;DDX5 | 1 | 1 | 1,32499E-21 | 1 | 1 |
| IDH3A | 26 | 25 | -3,19214E-17 | 0,999999987 | 1 |
| RPL4 | 18 | 15 | 0,196747155 | 0,187771941 | 1 |
| FN1 | 8 | 3 | -8,4055E-18 | 0,999999998 | 1 |
| CAMSAP3 | 10 | 10 | -3,89372E-14 | 0,999999366 | 1 |
| BTF3L4 | 4 | 3 | 4,41119E-17 | 0,999999992 | 1 |
| BTF3 | 3 | 2 | 0,253454114 | 0,310402509 | 1 |
| PSME1 | 5 | 5 | 9,00499E-18 | 0,999999993 | 1 |
| RFTN2 | 9 | 9 | 1,36133E-17 | 0,999999993 | 1 |
| CSPG4 | 17 | 17 | -2,44721E-17 | 0,999999992 | 1 |
| LYSMD2 | 1 | 1 | 0,042237088 | 0,411776322 | 1 |
| IARS1 | 36 | 28 | 0,005202301 | 0,627346286 | 1 |
| MAST4 | 2 | 2 | -0,019646482 | 0,606909236 | 1 |
| CEND1 | 7 | 7 | -8,76999E-18 | 0,999999994 | 1 |
| HMGCS2 | 1 | 1 | 3,596E-20 | 1 | 1 |
| NECAB2 | 15 | 15 | 0 | 1 | 1 |
| PYGM | 38 | 38 | 2,13441E-16 | 0,999999947 | 1 |
| PLA2G15 | 4 | 4 | -8,94295E-17 | 0,999999979 | 1 |
| ASS1 | 15 | 15 | -7,82579E-17 | 0,999999989 | 1 |
| PTPN5 | 4 | 4 | -9,16472E-18 | 0,999999996 | 1 |
| RANBP3 | 15 | 9 | 0 | 1 | 1 |
| GOT1 | 36 | 35 | -2,34415E-16 | 0,999999968 | 1 |
| CPNE1 | 11 | 11 | 3,02658E-17 | 0,999999985 | 1 |
| COPE | 4 | 4 | -8,55159E-21 | 1 | 1 |
| TBC1D17 | 11 | 11 | 0,012507664 | 0,585355655 | 1 |
| ADPGK | 4 | 4 | 5,70297E-17 | 0,999999985 | 1 |
| PSMC1 | 29 | 28 | -6,87052E-17 | 0,99999998 | 1 |
| PDCD4 | 3 | 3 | 0 | 1 | 1 |
| IGHG2B | 4 | 1 | 0 | 1 | 1 |
| sp\|P01864\|GCAB_M | 7 | 3 | 0,892480116 | 0,352198319 | 1 |
| MICALL1 | 1 | 1 | 0 | 1 | 1 |

| | | | | | |
|---|---|---|---|---|---|
| COX7B | 1 | 1 | 0 | 1 | 1 |
| DDHD1 | 6 | 6 | 2,43117E-17 | 0,999999984 | 1 |
| DGKE | 9 | 9 | -0,000113093 | 0,956958813 | 1 |
| ANKRD34A | 6 | 6 | -1,1703E-15 | 0,999999925 | 1 |
| BC034090 | 1 | 1 | 0 | 1 | 1 |
| SEC16A | 11 | 8 | 0,085907252 | 0,131649798 | 1 |
| ATP5MG | 6 | 6 | 2,3946E-17 | 0,999999991 | 1 |
| ASNS | 19 | 14 | 3,42875E-18 | 0,999999997 | 1 |
| CORO7 | 19 | 17 | 0 | 1 | 1 |
| TCEA1 | 13 | 5 | 1,69412E-17 | 0,999999994 | 1 |
| NACA | 8 | 6 | 0,058745106 | 0,384671995 | 1 |
| CRYAB | 12 | 12 | -2,91556E-16 | 0,999999957 | 1 |
| TNC | 15 | 15 | 0,020271952 | 0,592546816 | 1 |
| FAM131B | 11 | 11 | 8,58621E-18 | 0,999999995 | 1 |
| SLC43A2 | 2 | 2 | 0,027515405 | 0,546665552 | 1 |
| APBA1 | 6 | 6 | 0 | 1 | 1 |
| DDN | 2 | 2 | -1,31843E-14 | 0,999999758 | 1 |
| ANK2 | 13 | 13 | -1,34046E-17 | 0,999999994 | 1 |
| AGFG1 | 9 | 9 | 0,043389355 | 0,122503185 | 1 |
| CBL | 7 | 7 | 0 | 1 | 1 |
| LZTS1 | 4 | 4 | -0,065311248 | 0,449238906 | 1 |
| NRSN1 | 1 | 1 | 0 | 1 | 1 |
| CNOT1 | 6 | 6 | -0,058039958 | 0,184068054 | 1 |
| TBCB | 12 | 12 | 0 | 1 | 1 |
| CLPTM1 | 12 | 12 | 0 | 1 | 1 |
| LONP1 | 38 | 38 | -1,97475E-16 | 0,999999968 | 1 |
| VCAN | 19 | 18 | -3,39059E-17 | 0,999999987 | 1 |
| RPS5 | 9 | 7 | -4,76067E-16 | 0,999999941 | 1 |
| SHFL | 2 | 2 | -0,009543695 | 0,731300277 | 1 |
| SEC31A | 30 | 27 | 0,034317551 | 0,222391701 | 1 |
| NUDCD2 | 5 | 4 | -4,59826E-16 | 0,999999973 | 1 |
| LZTFL1 | 7 | 7 | 0 | 1 | 1 |
| EPB41L3 | 7 | 7 | -0,070841191 | 0,118914661 | 1 |

| | | | | | |
|---|---|---|---|---|---|
| SMG1 | 3 | 3 | 5,52782E-17 | 0,999999984 | 1 |
| ACADL | 24 | 24 | -0,068559956 | 0,36485386 | 1 |
| HDGFL2 | 8 | 1 | 0 | 1 | 1 |
| SCAMP5 | 2 | 2 | 2,80357E-16 | 0,999999957 | 1 |
| COQ3 | 5 | 5 | 0 | 1 | 1 |
| CACNA2D1 | 41 | 41 | 4,31532E-16 | 0,999999949 | 1 |
| DDHD2 | 14 | 12 | 2,53287E-16 | 0,999999947 | 1 |
| OSCP1 | 9 | 8 | 5,06688E-20 | 1 | 1 |
| CZIB | 5 | 5 | -3,9727E-18 | 0,999999996 | 1 |
| XPO4 | 2 | 2 | 0 | 1 | 1 |
| PLIN3 | 8 | 7 | -0,084771296 | 0,13902293 | 1 |
| DIABLO | 2 | 2 | 0 | 1 | 1 |
| BRINP2 | 5 | 5 | 0,085506928 | 0,118849328 | 1 |
| FYN | 11 | 11 | 0,02757907 | 0,347208623 | 1 |
| CASK | 11 | 11 | 7,19674E-18 | 0,999999992 | 1 |
| PHB2 | 20 | 20 | 0 | 1 | 1 |
| EML2 | 16 | 15 | -0,018656853 | 0,389329526 | 1 |
| AHCYL1 | 39 | 38 | -3,06269E-17 | 0,999999992 | 1 |
| ZFAND2B | 1 | 1 | 0 | 1 | 1 |
| PTGDS | 4 | 3 | 1,68622E-17 | 0,999999996 | 1 |
| DNAJA3 | 13 | 13 | -7,64111E-17 | 0,999999978 | 1 |
| TIMM50 | 9 | 9 | -1,51509E-16 | 0,999999972 | 1 |
| HSPA1B;HSPA1A | 15 | 15 | 0,031767406 | 0,306551841 | 1 |
| RTN4 | 39 | 39 | -0,046164955 | 0,29209913 | 1 |
| DBT | 20 | 20 | -8,34103E-15 | 0,99999982 | 1 |
| TANC2 | 31 | 31 | -1,33117E-16 | 0,999999955 | 1 |
| CDKL5 | 14 | 14 | -3,74896E-16 | 0,999999946 | 1 |
| FAM126A | 1 | 1 | 0,111360053 | 0,266265987 | 1 |
| RLBP1 | 5 | 5 | -0,01160018 | 0,757950552 | 1 |
| FGD4 | 3 | 3 | -1,68909E-15 | 0,999999929 | 1 |
| USP14 | 27 | 27 | -4,43753E-18 | 0,999999997 | 1 |
| RAB24 | 6 | 6 | -1,21052E-17 | 0,999999991 | 1 |
| TFG | 13 | 13 | 0 | 1 | 1 |

| | | | | | |
|---|---|---|---|---|---|
| PPP1R12A | 17 | 16 | -0,028487115 | 0,367690822 | 1 |
| FN3K | 5 | 5 | 0 | 1 | 1 |
| KCNJ4 | 3 | 3 | -4,71576E-18 | 0,999999997 | 1 |
| PNKD | 4 | 4 | 2,71273E-19 | 1 | 1 |
| DRG2 | 14 | 14 | 7,70689E-18 | 0,999999994 | 1 |
| L1CAM | 28 | 28 | 0,034230221 | 0,235215916 | 1 |
| STIM2 | 11 | 11 | 4,32494E-17 | 0,999999982 | 1 |
| SIPA1L1 | 50 | 50 | -1,94027E-16 | 0,999999964 | 1 |
| STX19 | 1 | 1 | -0,199046357 | 0,147476154 | 1 |
| PTK2 | 9 | 8 | 1,00443E-15 | 0,999999941 | 1 |
| SAE1 | 20 | 2 | 0 | 1 | 1 |
| DCLK1 | 31 | 30 | 1,27091E-13 | 0,999998037 | 1 |
| UBXN6 | 19 | 19 | 0,035789142 | 0,132609834 | 1 |
| DGKI | 7 | 7 | -8,07337E-17 | 0,999999985 | 1 |
| ATP5PB | 28 | 28 | 0 | 1 | 1 |
| SCAMP3 | 11 | 11 | 0 | 1 | 1 |
| SCAMP1 | 13 | 12 | 0 | 1 | 1 |
| KIF3A | 17 | 17 | -6,95453E-19 | 0,999999999 | 1 |
| TACC2 | 3 | 3 | -7,50472E-17 | 0,999999978 | 1 |
| LZTS3 | 4 | 4 | -4,56959E-19 | 0,999999999 | 1 |
| AATK | 4 | 4 | 0 | 1 | 1 |
| CC2D1A | 8 | 8 | 0 | 1 | 1 |
| CSNK1G3 | 3 | 3 | 3,49296E-16 | 0,999999962 | 1 |
| PLCG1 | 10 | 10 | 0,003819832 | 0,786124081 | 1 |
| PSMD2 | 40 | 37 | 0,014609553 | 0,433305994 | 1 |
| GPC5 | 4 | 4 | 0 | 1 | 1 |
| ATP5F1D | 4 | 4 | -0,046875089 | 0,430453308 | 1 |
| AACS | 11 | 9 | 2,84089E-16 | 0,999999976 | 1 |
| AARS1 | 44 | 41 | -8,28318E-17 | 0,999999962 | 1 |
| SCAMP4 | 1 | 1 | 0 | 1 | 1 |
| MAPT | 6 | 6 | -0,003524288 | 0,814474054 | 1 |
| C2CD4C | 6 | 6 | 1,03278E-17 | 0,999999996 | 1 |
| SEPTIN10 | 1 | 1 | 0 | 1 | 1 |

| | | | | | |
|---|---|---|---|---|---|
| HINT1 | 9 | 9 | 0 | 1 | 1 |
| LYRM4 | 5 | 5 | -1,50139E-17 | 0,99999999 | 1 |
| NEFM | 33 | 33 | -0,055529878 | 0,501065726 | 1 |
| PIGW | 1 | 1 | 0 | 1 | 1 |
| SORCS2 | 12 | 12 | 0 | 1 | 1 |
| IKBKG | 2 | 2 | 0,033786581 | 0,512147809 | 1 |
| CDH11 | 10 | 10 | -4,05892E-18 | 0,999999999 | 1 |
| GPLD1 | 2 | 2 | -2,52923E-19 | 0,999999999 | 1 |
| SETD1A | 1 | 1 | 1,08671E-18 | 0,999999998 | 1 |
| OXR1 | 25 | 25 | -1,10704E-16 | 0,999999948 | 1 |
| TPM3 | 11 | 11 | 2,87572E-17 | 0,999999985 | 1 |
| NT5M | 7 | 7 | -0,108440629 | 0,206419993 | 1 |
| FRRS1L | 10 | 10 | 2,62836E-14 | 0,999999468 | 1 |
| NDUFB5 | 10 | 10 | 0 | 1 | 1 |
| FAM193B | 1 | 1 | 0 | 1 | 1 |
| FKBP3 | 10 | 8 | 4,62383E-18 | 0,999999998 | 1 |
| RGS7 | 15 | 15 | 0,010360062 | 0,620431126 | 1 |
| STK32C | 5 | 5 | -0,023140929 | 0,595853132 | 1 |
| APOA1 | 18 | 4 | 0,054355211 | 0,638780625 | 1 |
| CHKB | 11 | 9 | 2,5941E-16 | 0,999999954 | 1 |
| RBCK1 | 1 | 1 | 0 | 1 | 1 |
| PYGB | 60 | 58 | -1,91293E-16 | 0,999999949 | 1 |
| ROBO1 | 14 | 14 | 0 | 1 | 1 |
| SYT13 | 4 | 4 | -3,72717E-16 | 0,999999973 | 1 |
| EFNB3 | 5 | 5 | -0,053304151 | 0,289474982 | 1 |
| KIF2A | 2 | 2 | -3,3408E-17 | 0,999999987 | 1 |
| TGM2 | 10 | 7 | 1,01957E-16 | 0,999999977 | 1 |
| CCDC47 | 14 | 14 | -1,24728E-16 | 0,99999996 | 1 |
| TPD52 | 9 | 9 | 0,038639531 | 0,215043318 | 1 |
| APC | 17 | 17 | -9,22418E-17 | 0,999999978 | 1 |
| RASAL2 | 10 | 10 | -0,039380018 | 0,259876142 | 1 |
| DDX6 | 14 | 12 | -1,3713E-17 | 0,999999997 | 1 |
| TMEM47 | 3 | 2 | 0,06885838 | 0,290521749 | 1 |

| | | | | | |
|---|---|---|---|---|---|
| C2CD5 | 15 | 15 | 1,1688E-17 | 0,999999991 | 1 |
| ISOC1 | 9 | 9 | -1,43843E-16 | 0,999999968 | 1 |
| GLIPR2 | 2 | 2 | 0,065979417 | 0,469989887 | 1 |
| PHACTR2 | 5 | 5 | -3,30074E-17 | 0,999999985 | 1 |
| MRPS30 | 9 | 9 | -0,033516683 | 0,442532146 | 1 |
| CRYBB1 | 3 | 3 | -0,057612777 | 0,337251877 | 1 |
| AMER2 | 10 | 10 | 0 | 1 | 1 |
| ATP6V1C1 | 43 | 43 | 8,95297E-19 | 0,999999998 | 1 |
| CLUH | 18 | 16 | 0 | 1 | 1 |
| sp\|Q6PIU9\|YJ005_M | 4 | 4 | -0,036074593 | 0,48803673 | 1 |
| EPHX2 | 12 | 12 | -5,16803E-16 | 0,999999937 | 1 |
| RP2 | 2 | 2 | 2,72562E-16 | 0,999999975 | 1 |
| ANKMY2 | 4 | 4 | 0 | 1 | 1 |
| TRPM3 | 3 | 3 | -3,20373E-18 | 0,999999996 | 1 |
| RCN1 | 2 | 2 | -1,16957E-14 | 0,999999723 | 1 |
| REPS1 | 13 | 13 | 0 | 1 | 1 |
| TOMM40 | 10 | 10 | -6,47961E-15 | 0,999999768 | 1 |
| DBN1 | 34 | 34 | -0,026706578 | 0,456176326 | 1 |
| GDA | 36 | 32 | 0 | 1 | 1 |
| SLC20A2 | 5 | 5 | -2,41998E-17 | 0,999999987 | 1 |
| GALM | 4 | 3 | 0 | 1 | 1 |
| COPS6 | 15 | 14 | 0,050914004 | 0,187802436 | 1 |
| CSF1R | 4 | 4 | 0,02110646 | 0,640935806 | 1 |
| SLC2A13 | 8 | 8 | -0,033031784 | 0,285462186 | 1 |
| SRGAP3;SRGAP1 | 4 | 4 | 1,54688E-18 | 0,999999997 | 1 |
| CLTB | 2 | 2 | 4,21587E-16 | 0,999999973 | 1 |
| ARHGEF6 | 6 | 6 | -0,004261674 | 0,762978237 | 1 |
| MFSD6 | 3 | 3 | 5,67729E-16 | 0,99999994 | 1 |
| MFSD6 | 4 | 4 | 0,000892528 | 0,900206866 | 1 |
| PPP2R5C | 2 | 2 | 6,59546E-17 | 0,999999982 | 1 |
| AP2M1 | 35 | 35 | 0,045987585 | 0,129393687 | 1 |
| PCDHGA1;PCDHGA2 | 1 | 1 | -3,56829E-19 | 0,999999999 | 1 |
| NOL3 | 6 | 5 | -0,059939786 | 0,244226379 | 1 |

| | | | | | |
|---|---|---|---|---|---|
| SEMA6B | 1 | 1 | 0 | 1 | 1 |
| EIF4G3 | 1 | 1 | -0,061127404 | 0,608108157 | 1 |
| OSBPL6 | 13 | 13 | -4,48683E-17 | 0,999999979 | 1 |
| MRAS | 12 | 12 | 0,042475293 | 0,267816386 | 1 |
| ECSIT | 7 | 7 | -0,008273568 | 0,762900219 | 1 |
| RAD23B | 18 | 14 | -2,89663E-17 | 0,999999983 | 1 |
| TNIK | 17 | 16 | -9,62099E-17 | 0,999999968 | 1 |
| TXNRD2 | 6 | 6 | 0 | 1 | 1 |
| SACM1L | 28 | 28 | 2,70619E-16 | 0,999999928 | 1 |
| TNPO3 | 10 | 10 | 0 | 1 | 1 |
| NELFB | 2 | 2 | 4,96204E-15 | 1 | 1 |
| SLC25A13 | 8 | 8 | -0,002837677 | 0,848269099 | 1 |
| CALD1 | 3 | 3 | 0,107756146 | 0,250455629 | 1 |
| GCLC | 26 | 19 | -2,46377E-17 | 0,999999989 | 1 |
| CLU | 16 | 16 | 0 | 1 | 1 |
| ENDOG | 7 | 7 | -0,047459042 | 0,221825383 | 1 |
| NPW | 1 | 1 | 0 | 1 | 1 |
| MDN1 | 1 | 1 | 0,12924747 | 0,34635903 | 1 |
| RPL8 | 9 | 8 | 0,071668343 | 0,413511908 | 1 |
| TRIM9 | 3 | 3 | -4,13181E-15 | 0,99999986 | 1 |
| CD244 | 1 | 1 | 0,229679004 | 0,127297293 | 1 |
| PTCD3 | 13 | 13 | -9,64717E-18 | 0,999999992 | 1 |
| NECAP1 | 10 | 9 | 0 | 1 | 1 |
| RBBP9 | 10 | 10 | 4,38645E-16 | 0,99999997 | 1 |
| CHL1 | 29 | 27 | 6,62188E-16 | 0,999999904 | 1 |
| SFXN1 | 14 | 14 | -2,44644E-16 | 0,999999978 | 1 |
| STRADA | 4 | 4 | 2,65449E-18 | 0,999999998 | 1 |
| C2CD2L | 20 | 20 | -0,020105081 | 0,295287342 | 1 |
| PPP1R21 | 24 | 23 | 0 | 1 | 1 |
| CLYBL | 13 | 13 | -5,86619E-14 | 0,999999374 | 1 |
| SGIP1 | 14 | 14 | -1,19585E-16 | 0,999999968 | 1 |
| DNAH6 | 2 | 2 | -6,36973E-18 | 0,999999998 | 1 |
| RIC8A | 13 | 13 | 0,039822455 | 0,278588182 | 1 |

| | | | | | |
|---|---|---|---|---|---|
| SEPTIN2 | 13 | 13 | 0 | 1 | 1 |
| EIF3H | 12 | 10 | 4,92606E-19 | 0,999999999 | 1 |
| TAPT1 | 5 | 5 | 0 | 1 | 1 |
| MAGI3 | 4 | 4 | 0 | 1 | 1 |
| RTKN | 2 | 2 | -6,16641E-16 | 0,999999983 | 1 |
| KRT16 | 28 | 9 | 0 | 1 | 1 |
| CPM | 8 | 8 | -0,090320509 | 0,149615823 | 1 |
| TSG101 | 7 | 7 | 2,40842E-17 | 0,999999991 | 1 |
| DHX36 | 5 | 4 | 0 | 1 | 1 |
| EEF1B | 10 | 9 | 0,059169237 | 0,218941837 | 1 |
| UBXN7 | 8 | 4 | 0 | 1 | 1 |
| RNF34 | 4 | 4 | 2,32676E-16 | 0,999999962 | 1 |
| IP6K1 | 6 | 6 | 0 | 1 | 1 |
| ABHD10 | 9 | 9 | -0,04599144 | 0,365579105 | 1 |
| CHCHD6 | 16 | 16 | -1,10648E-16 | 0,999999981 | 1 |
| PSAT1 | 21 | 21 | -1,81249E-17 | 0,999999987 | 1 |
| LRRFIP1 | 5 | 5 | -1,02498E-16 | 0,999999969 | 1 |
| UBAC2 | 3 | 3 | -0,056731619 | 0,356408028 | 1 |
| DPP7 | 4 | 4 | -0,057094561 | 0,320521383 | 1 |
| EXOC2 | 22 | 22 | 0,036497634 | 0,115813577 | 1 |
| ACADVL | 21 | 21 | -0,049222807 | 0,372694979 | 1 |
| UNC5C | 3 | 3 | 7,12692E-17 | 0,999999981 | 1 |
| AKR1E2 | 7 | 7 | -1,49439E-17 | 0,999999992 | 1 |
| CNPY3 | 3 | 3 | -3,86764E-15 | 0,999999829 | 1 |
| CNTN2 | 19 | 19 | -2,79428E-17 | 0,999999983 | 1 |
| EPHX1 | 14 | 14 | -1,97431E-17 | 0,999999996 | 1 |
| RGS20 | 3 | 3 | -8,46699E-17 | 0,999999984 | 1 |
| PRRT2 | 4 | 4 | -3,02571E-17 | 0,999999992 | 1 |
| PNPT1 | 17 | 17 | 3,17747E-16 | 0,999999971 | 1 |
| NAPEPLD | 5 | 5 | 0,05644819 | 0,273361027 | 1 |
| OGFRL1 | 8 | 8 | 0,035277296 | 0,309247392 | 1 |
| TBC1D10B | 11 | 11 | 0,045873517 | 0,118567266 | 1 |
| PPP6R1 | 7 | 7 | -0,072053434 | 0,151535503 | 1 |

| | | | | | |
|---|---|---|---|---|---|
| RAP1GAP2 | 13 | 13 | 2,44515E-16 | 0,999999947 | 1 |
| ECI2 | 7 | 7 | -3,86077E-17 | 0,999999981 | 1 |
| CNOT1 | 20 | 20 | -1,47769E-15 | 0,999999851 | 1 |
| CAT | 19 | 19 | 1,34891E-17 | 0,999999996 | 1 |
| RIMS2 | 13 | 13 | 4,51365E-18 | 0,999999999 | 1 |
| ADRA2A | 2 | 2 | 0 | 1 | 1 |
| UBA3 | 16 | 13 | -0,030681772 | 0,265561631 | 1 |
| SPECC1 | 4 | 4 | -1,24479E-18 | 0,999999998 | 1 |
| ARPC1B | 3 | 3 | 6,72708E-17 | 0,999999988 | 1 |
| HMGCS1 | 14 | 10 | 3,15944E-19 | 0,999999999 | 1 |
| MTFR1L | 8 | 8 | 0 | 1 | 1 |
| R3HDM2 | 5 | 5 | 1,10494E-14 | 0,999999942 | 1 |
| RPN1 | 32 | 32 | 0 | 1 | 1 |
| NPDC1 | 3 | 3 | 2,89853E-18 | 0,999999997 | 1 |
| PKM | 8 | 8 | 0,008816854 | 0,711413781 | 1 |
| EIF4G2 | 22 | 21 | 2,49124E-17 | 0,999999984 | 1 |
| SH3KBP1 | 15 | 15 | -0,036032874 | 0,288640145 | 1 |
| OSBPL2 | 10 | 10 | 1,12698E-17 | 0,999999994 | 1 |
| RGS10 | 4 | 4 | -4,71638E-17 | 0,999999984 | 1 |
| RABGAP1L | 7 | 7 | 0,056218772 | 0,236700561 | 1 |
| NSFL1C | 24 | 24 | 6,36546E-17 | 0,999999972 | 1 |
| PRKCI | 3 | 3 | 0,062505732 | 0,335708796 | 1 |
| TUB | 2 | 2 | 0 | 1 | 1 |
| CRIP2 | 4 | 4 | 2,47093E-18 | 0,999999999 | 1 |
| RAB11FIP2 | 12 | 12 | -1,63078E-16 | 0,999999959 | 1 |
| GCLM | 7 | 7 | 0 | 1 | 1 |
| CHP1 | 11 | 11 | 0,043864653 | 0,125899964 | 1 |
| DNAJC10 | 5 | 5 | 1,31434E-16 | 0,999999976 | 1 |
| LRBA | 11 | 8 | 0 | 1 | 1 |
| RNMT | 19 | 3 | -3,70222E-17 | 0,999999989 | 1 |
| CEP170 | 27 | 27 | -2,69198E-16 | 0,999999964 | 1 |
| AKAP10 | 9 | 9 | 0 | 1 | 1 |
| DCTN2 | 25 | 24 | 0 | 1 | 1 |

| | | | | | |
|---|---|---|---|---|---|
| GPSM1 | 10 | 9 | -0,048602888 | 0,253875113 | 1 |
| RTN4R | 3 | 3 | 0 | 1 | 1 |
| EVL | 8 | 8 | -7,42202E-20 | 1 | 1 |
| GLRX3 | 18 | 17 | 9,54552E-17 | 0,999999963 | 1 |
| CADPS2 | 20 | 20 | 5,15619E-17 | 0,999999991 | 1 |
| MIF | 7 | 7 | 4,06358E-16 | 0,999999956 | 1 |
| AJAP1 | 3 | 3 | 1,00744E-17 | 0,999999996 | 1 |
| DTD1 | 7 | 6 | 5,19871E-17 | 0,999999974 | 1 |
| LYPLAL1 | 5 | 5 | 0,065882554 | 0,185869629 | 1 |
| VGF | 10 | 10 | 1,52774E-16 | 0,999999977 | 1 |
| NIT1 | 7 | 7 | 0 | 1 | 1 |
| RBM14 | 1 | 1 | 0,280176693 | 0,180032998 | 1 |
| KALRN | 6 | 6 | 0 | 1 | 1 |
| GRB2 | 15 | 14 | -6,86293E-17 | 0,999999983 | 1 |
| ECI1 | 15 | 15 | -0,084939741 | 0,334234232 | 1 |
| RPA1 | 4 | 3 | 0 | 1 | 1 |
| HSPA1L | 11 | 10 | 0,010601476 | 0,650711464 | 1 |
| PSMB8 | 4 | 3 | 0,041564042 | 0,521849276 | 1 |
| EEF1D | 9 | 8 | 6,98318E-17 | 0,999999981 | 1 |
| BROX | 5 | 5 | 0 | 1 | 1 |
| COPS7B | 5 | 5 | -8,72361E-18 | 0,999999995 | 1 |
| RASA1 | 14 | 14 | 2,93372E-18 | 0,999999997 | 1 |
| STK3 | 2 | 2 | 0 | 1 | 1 |
| HSPA12A | 39 | 38 | 0,031762731 | 0,223967702 | 1 |
| TMEM119 | 2 | 2 | 1,3394E-19 | 1 | 1 |
| SH3GLB2 | 20 | 20 | -3,53385E-16 | 0,999999931 | 1 |
| PSMA4 | 11 | 11 | 0,044602951 | 0,282777044 | 1 |
| NBEA;LRBA | 5 | 5 | 0 | 1 | 1 |
| PCDHB1 | 1 | 1 | 0 | 1 | 1 |
| CAND2 | 2 | 2 | -0,136217044 | 0,116465009 | 1 |
| TSC22D1 | 1 | 1 | -0,033198649 | 0,533600483 | 1 |
| SALL1 | 1 | 1 | 0,118060175 | 0,269550546 | 1 |
| TSC22D2 | 4 | 4 | 0 | 1 | 1 |

| | | | | | |
|---|---|---|---|---|---|
| CACNG2 | 2 | 2 | 0,044436824 | 0,465906021 | 1 |
| GDPGP1 | 8 | 8 | -4,70329E-19 | 1 | 1 |
| AP3M2 | 20 | 20 | 0,031517992 | 0,321171061 | 1 |
| OSBP | 15 | 15 | 2,71925E-18 | 0,999999997 | 1 |
| AMDHD2 | 4 | 3 | 0 | 1 | 1 |
| SERPINC1 | 3 | 3 | 9,62222E-16 | 0,999999968 | 1 |
| PSMB7 | 10 | 10 | 5,69295E-17 | 0,999999988 | 1 |
| HNRNPH3 | 1 | 1 | 0 | 1 | 1 |
| HNRNPH2 | 6 | 6 | 3,62391E-17 | 0,999999986 | 1 |
| HNRNPH1;HNRNPF | 1 | 1 | 0 | 1 | 1 |
| PALM3 | 3 | 3 | -4,38911E-17 | 0,999999993 | 1 |
| SNX14 | 2 | 2 | 2,56652E-17 | 0,999999994 | 1 |
| SUPV3L1 | 17 | 17 | -6,6662E-16 | 0,99999995 | 1 |
| PGAM5 | 8 | 8 | -2,45358E-16 | 0,999999961 | 1 |
| KIF1A | 3 | 3 | 0,116318287 | 0,315045575 | 1 |
| CYP4X1 | 3 | 3 | -9,59372E-19 | 0,999999999 | 1 |
| RRAGB | 8 | 8 | 1,48606E-17 | 0,999999987 | 1 |
| ARHGDIB | 6 | 6 | 3,02902E-18 | 0,999999997 | 1 |
| NF1 | 19 | 19 | -7,19776E-17 | 0,999999971 | 1 |
| DHODH | 18 | 18 | -0,006226518 | 0,720164752 | 1 |
| ATE1 | 8 | 7 | 4,15713E-18 | 0,999999998 | 1 |
| COPG2 | 12 | 11 | -0,000821215 | 0,923557564 | 1 |
| CACNB1 | 2 | 2 | 0 | 1 | 1 |
| PRORSD1 | 1 | 1 | 0,336708486 | 0,130203378 | 1 |
| FN3KRP | 6 | 6 | 0 | 1 | 1 |
| DNAJC6;GAK | 3 | 3 | 3,79299E-18 | 0,999999997 | 1 |
| CACNB2 | 8 | 8 | 5,60345E-17 | 0,999999979 | 1 |
| CACNB1 | 9 | 9 | 0 | 1 | 1 |
| CACNB4 | 13 | 13 | -6,27529E-17 | 0,999999989 | 1 |
| AGPAT5 | 6 | 6 | -2,40319E-17 | 0,999999987 | 1 |
| P33MONOX | 3 | 3 | 9,78037E-15 | 0,999999732 | 1 |
| TNPO2 | 9 | 8 | -1,1296E-14 | 0,999999693 | 1 |
| NOS1 | 28 | 28 | 3,3082E-16 | 0,999999961 | 1 |

| | | | | | |
|---|---|---|---|---|---|
| CHMP2B | 3 | 3 | 0 | 1 | 1 |
| PTPRG | 12 | 12 | 0,020071282 | 0,484977374 | 1 |
| PDE4D | 10 | 10 | 2,2048E-17 | 0,999999987 | 1 |
| NDEL1 | 10 | 10 | -1,98303E-17 | 0,999999993 | 1 |
| GLO1 | 16 | 15 | -2,70912E-17 | 0,999999989 | 1 |
| CACNA1B | 14 | 14 | 0,019237518 | 0,484986611 | 1 |
| CTH | 2 | 2 | -6,35555E-18 | 0,999999995 | 1 |
| ATP5C1 | 1 | 1 | -2,42826E-18 | 0,999999997 | 1 |
| TST | 14 | 14 | -0,163632563 | 0,146347978 | 1 |
| SNAP47 | 23 | 23 | 2,99697E-17 | 0,999999982 | 1 |
| RMC1 | 8 | 8 | 0 | 1 | 1 |
| NUDT2 | 7 | 7 | 0 | 1 | 1 |
| IGSF8 | 2 | 2 | -1,08682E-16 | 0,999999996 | 1 |
| ARFGEF3 | 23 | 23 | 0 | 1 | 1 |
| PDCD6IP | 37 | 36 | -0,025113657 | 0,118654952 | 1 |
| PRMT1 | 11 | 9 | 0,033696804 | 0,341142749 | 1 |
| ARHGEF28 | 1 | 1 | 0 | 1 | 1 |
| KRT17 | 9 | 6 | 0,333378658 | 0,31638676 | 1 |
| IFIT3 | 3 | 1 | 0 | 1 | 1 |
| PAPSS1 | 12 | 5 | 0 | 1 | 1 |
| CLINT1 | 2 | 2 | 0 | 1 | 1 |
| ZDHHC14 | 1 | 1 | 0 | 1 | 1 |
| RPH3A | 29 | 29 | 0 | 1 | 1 |
| AGAP3 | 6 | 6 | 2,16316E-18 | 0,999999996 | 1 |
| NDUFA3 | 2 | 2 | -3,035E-17 | 0,999999986 | 1 |
| EIF4A1 | 31 | 29 | 0,070953136 | 0,162632211 | 1 |
| CISD3 | 4 | 4 | 0 | 1 | 1 |
| DPP10 | 29 | 29 | 0 | 1 | 1 |
| ZDHHC17 | 1 | 1 | 0 | 1 | 1 |
| TXLNG | 1 | 1 | 0 | 1 | 1 |
| PPM1E | 21 | 16 | 0,057394134 | 0,199218911 | 1 |
| RAPGEF1 | 4 | 4 | 0 | 1 | 1 |
| MYL12B | 14 | 14 | 0,044477239 | 0,26587522 | 1 |

| | | | | | |
|---|---|---|---|---|---|
| FAM234B | 6 | 6 | 0,078726702 | 0,141792226 | 1 |
| BRI3BP | 3 | 3 | -4,318E-19 | 1 | 1 |
| DIS3L2 | 7 | 6 | 6,10892E-18 | 0,999999999 | 1 |
| TAB3 | 1 | 1 | 0 | 1 | 1 |
| PICALM | 11 | 11 | 0 | 1 | 1 |
| RIMS1 | 29 | 29 | 2,46687E-16 | 0,999999952 | 1 |
| RETREG2 | 1 | 1 | 0 | 1 | 1 |
| LIN7C;LIN7A | 1 | 1 | 0 | 1 | 1 |
| LIN7B | 2 | 1 | 0 | 1 | 1 |
| TAF9 | 1 | 1 | 0 | 1 | 1 |
| NAA35 | 3 | 3 | 0 | 1 | 1 |
| GRM7 | 15 | 15 | 0,003378449 | 0,729250185 | 1 |
| TNPO1 | 5 | 5 | 0 | 1 | 1 |
| PCYOX1L | 9 | 9 | 0,004409766 | 0,729126409 | 1 |
| LINGO2 | 4 | 4 | -4,50277E-15 | 0,999999803 | 1 |
| PRPF19 | 5 | 5 | 4,94252E-19 | 0,999999999 | 1 |
| REEP5 | 4 | 4 | 0 | 1 | 1 |
| GLCCI1 | 3 | 3 | -3,17324E-17 | 0,999999992 | 1 |
| BLMH | 17 | 17 | 9,82648E-17 | 0,999999962 | 1 |
| NUCKS1 | 5 | 1 | 0 | 1 | 1 |
| YWHAG | 21 | 20 | 1,49523E-16 | 0,999999969 | 1 |
| PRMT7 | 3 | 2 | -2,45018E-15 | 0,999999899 | 1 |
| CUEDC2 | 3 | 3 | 0 | 1 | 1 |
| ADGRB2 | 8 | 7 | 0,039754247 | 0,233374242 | 1 |
| GABRA3 | 5 | 5 | 0,042231319 | 0,438568521 | 1 |
| ELFN1 | 7 | 7 | -0,022720408 | 0,457256861 | 1 |
| PPP1R3F | 2 | 2 | -8,89739E-16 | 0,999999984 | 1 |
| THBS4 | 6 | 2 | -3,82149E-18 | 0,999999997 | 1 |
| FSD1L | 4 | 4 | 0 | 1 | 1 |
| EIF2B2 | 5 | 5 | -1,04178E-15 | 0,999999935 | 1 |
| PGM2L1 | 40 | 38 | 0 | 1 | 1 |
| PSMC5 | 26 | 25 | 0,018293579 | 0,418739455 | 1 |
| ABHD14B | 6 | 4 | -3,17115E-18 | 0,999999999 | 1 |

| | | | | | |
|---|---|---|---|---|---|
| NAV1 | 13 | 13 | -5,03152E-17 | 0,999999979 | 1 |
| SMYD3 | 1 | 1 | 0 | 1 | 1 |
| CBR4 | 9 | 9 | -1,45459E-17 | 0,999999995 | 1 |
| CCDC127 | 7 | 7 | -2,68262E-19 | 0,999999999 | 1 |
| NEFH | 13 | 13 | -0,083924213 | 0,286492045 | 1 |
| VPS4B | 8 | 7 | -0,025320564 | 0,436847778 | 1 |
| SLC7A5 | 6 | 6 | 0,049860885 | 0,122809625 | 1 |
| AHSA2 | 3 | 3 | -2,19905E-19 | 1 | 1 |
| CNTNAP1 | 45 | 45 | 0 | 1 | 1 |
| ADCK1 | 8 | 8 | -0,080273649 | 0,125886138 | 1 |
| SLC39A8 | 1 | 1 | 0 | 1 | 1 |
| PSMD14 | 13 | 12 | 0,050536696 | 0,193079572 | 1 |
| VAT1 | 18 | 18 | 6,01556E-16 | 0,999999928 | 1 |
| TENM2 | 21 | 21 | -4,63966E-17 | 0,999999991 | 1 |
| SUOX | 8 | 8 | -0,09565888 | 0,260557263 | 1 |
| KIFAP3 | 5 | 5 | 0 | 1 | 1 |
| PACSIN2 | 1 | 1 | 0 | 1 | 1 |
| RAB33B | 7 | 7 | 0 | 1 | 1 |
| RPL6 | 9 | 9 | 0,203280304 | 0,161824774 | 1 |
| SLC9A3R1 | 19 | 18 | -9,56593E-17 | 0,99999997 | 1 |
| MINDY2 | 3 | 3 | -0,068952964 | 0,18193594 | 1 |
| ATP5A1 | 50 | 50 | -1,79622E-16 | 0,999999966 | 1 |
| USP5 | 47 | 45 | 0,004511906 | 0,738924556 | 1 |
| CD47 | 4 | 4 | 0 | 1 | 1 |
| RAB5C | 11 | 11 | 3,89236E-17 | 0,999999981 | 1 |
| PMPCB | 13 | 13 | -0,092744366 | 0,17951747 | 1 |
| ABHD11 | 6 | 6 | -2,57801E-17 | 0,99999999 | 1 |
| TKT | 32 | 32 | 3,79205E-17 | 0,99999999 | 1 |
| RECQL5 | 1 | 1 | -0,215647931 | 0,133736667 | 1 |
| PGP | 15 | 15 | 0 | 1 | 1 |
| PDCD5 | 6 | 6 | -4,78979E-18 | 0,999999998 | 1 |
| PLTP | 6 | 4 | -0,023455288 | 0,584897155 | 1 |
| MAEA | 4 | 4 | 0,063558736 | 0,209983024 | 1 |

| | | | | | |
|---|---|---|---|---|---|
| ASAP2 | 2 | 2 | 0,040503475 | 0,4659248 | 1 |
| THNSL2 | 4 | 2 | 1,58656E-12 | 0,999999185 | 1 |
| P2RY12 | 5 | 5 | 0 | 1 | 1 |
| NRBP1 | 7 | 6 | 0 | 1 | 1 |
| NANP | 6 | 4 | -0,062028113 | 0,385360054 | 1 |
| CNDP2 | 27 | 24 | 2,53579E-16 | 0,99999995 | 1 |
| TUBA4A | 10 | 10 | -5,952E-16 | 0,999999934 | 1 |
| SCFD2 | 4 | 4 | 0 | 1 | 1 |
| UFL1 | 5 | 4 | 4,12558E-17 | 0,999999978 | 1 |
| CPNE8 | 2 | 2 | -0,016295145 | 0,689550875 | 1 |
| NTM | 9 | 9 | 2,40923E-16 | 0,999999956 | 1 |
| ZRANB2 | 4 | 1 | 0 | 1 | 1 |
| BPGM | 8 | 6 | 3,96658E-26 | 1 | 1 |
| EEPD1 | 3 | 3 | -0,085363185 | 0,17367446 | 1 |
| F8A1 | 5 | 5 | 0,042798954 | 0,327527321 | 1 |
| AVL9 | 12 | 12 | -3,35471E-16 | 0,999999952 | 1 |
| PTPRT | 4 | 4 | 1,40815E-16 | 0,999999981 | 1 |
| SNX1 | 20 | 20 | 1,29337E-16 | 0,999999956 | 1 |
| PCBD1 | 3 | 3 | 1,22662E-16 | 0,999999988 | 1 |
| TTPAL | 4 | 4 | 0 | 1 | 1 |
| PEX11B | 6 | 6 | 0 | 1 | 1 |
| TTC37 | 8 | 8 | -6,62479E-17 | 0,999999982 | 1 |
| CCDC71 | 1 | 1 | 0,275201343 | 0,166715344 | 1 |
| NDUFB11 | 12 | 12 | 0 | 1 | 1 |
| ATG4C | 2 | 2 | 0 | 1 | 1 |
| AK4 | 14 | 14 | 1,63568E-15 | 0,99999991 | 1 |
| GMPPA | 8 | 8 | 0,037821852 | 0,250957438 | 1 |
| AK3 | 15 | 15 | -2,01756E-15 | 0,999999912 | 1 |
| PRMT3 | 3 | 3 | -0,038342466 | 0,471667867 | 1 |
| LAGE3 | 1 | 1 | 0 | 1 | 1 |
| APP | 21 | 21 | 0 | 1 | 1 |
| ACSS1 | 18 | 18 | -0,185116007 | 0,130936704 | 1 |
| SEC24C | 14 | 14 | 0 | 1 | 1 |

| | | | | | |
|---|---|---|---|---|---|
| GZF1 | 1 | 1 | -0,081225155 | 0,504682112 | 1 |
| HDAC6 | 11 | 11 | 2,6079E-17 | 0,999999985 | 1 |
| RAC3;RAC1 | 1 | 1 | 0 | 1 | 1 |
| RAC1 | 5 | 5 | 7,15925E-18 | 0,999999993 | 1 |
| COQ8A | 7 | 7 | 2,35101E-16 | 0,999999981 | 1 |
| SERPINA3N | 4 | 2 | 0 | 1 | 1 |
| MAT2B | 14 | 12 | -3,15719E-17 | 0,999999991 | 1 |
| CFL2 | 8 | 8 | 9,04913E-19 | 0,999999999 | 1 |
| CFL1 | 16 | 16 | 1,90363E-17 | 0,999999988 | 1 |
| GYKL1 | 5 | 5 | -5,06811E-17 | 0,99999998 | 1 |
| LYSMD1 | 6 | 6 | 0 | 1 | 1 |
| GPI | 48 | 48 | 0 | 1 | 1 |
| METTL26 | 5 | 4 | -0,081057108 | 0,183226201 | 1 |
| IARS2 | 37 | 37 | -2,33721E-16 | 0,99999997 | 1 |
| TUBG2 | 5 | 5 | 0 | 1 | 1 |
| RHOG | 8 | 8 | -1,24446E-15 | 0,999999906 | 1 |
| SHTN1 | 2 | 2 | 0 | 1 | 1 |
| TWF2 | 16 | 16 | 3,33691E-21 | 1 | 1 |
| SIPA1L2 | 5 | 5 | -0,078950317 | 0,13213624 | 1 |
| TRIM32 | 6 | 6 | 3,26932E-17 | 0,999999986 | 1 |
| PTDSS1 | 1 | 1 | 0 | 1 | 1 |
| TUBB4B;TUBB4A | 6 | 6 | 6,79065E-17 | 0,999999982 | 1 |
| VMN1R87 | 1 | 1 | 0 | 1 | 1 |
| NIPSNAP3B | 9 | 9 | -1,05771E-17 | 0,999999998 | 1 |
| RYR2 | 5 | 5 | 0 | 1 | 1 |
| STX6 | 7 | 7 | 0 | 1 | 1 |
| UNC5A | 6 | 6 | 9,00686E-15 | 0,999999827 | 1 |
| NAMPT | 18 | 17 | 0,053909355 | 0,284428112 | 1 |
| GFRA1 | 2 | 2 | 0 | 1 | 1 |
| ARL15 | 4 | 4 | 0,022544709 | 0,496101507 | 1 |
| HNRNPA0 | 5 | 3 | 3,27862E-13 | 0,99999862 | 1 |
| SEPSECS | 3 | 3 | 0,208919929 | 0,1665237 | 1 |
| COMMD10 | 3 | 3 | -1,61073E-17 | 0,999999996 | 1 |

| | | | | | |
|---|---|---|---|---|---|
| VAPA | 11 | 11 | 7,68681E-18 | 0,999999995 | 1 |
| PHYHIP | 15 | 15 | 0,055376914 | 0,25770001 | 1 |
| MAP6D1 | 8 | 8 | 6,02057E-16 | 0,999999962 | 1 |
| TUSC2 | 2 | 2 | 1,18523E-16 | 0,999999987 | 1 |
| PDXDC1 | 14 | 10 | 8,11111E-18 | 0,999999996 | 1 |
| STX16 | 7 | 7 | 8,30113E-17 | 0,99999998 | 1 |
| PRELID1 | 1 | 1 | 0 | 1 | 1 |
| DST | 8 | 8 | -6,10127E-17 | 0,999999979 | 1 |
| PPP1R1B | 5 | 5 | 5,65626E-19 | 1 | 1 |
| NDUFA9 | 29 | 29 | 0 | 1 | 1 |
| FAM120B | 1 | 1 | 0,036665328 | 0,610503723 | 1 |
| CTDP1 | 3 | 1 | 0,183469334 | 0,314942022 | 1 |
| CA2 | 16 | 15 | -2,09662E-16 | 0,999999963 | 1 |
| AGO2 | 12 | 12 | 1,34223E-17 | 0,999999995 | 1 |
| ARMCX2 | 1 | 1 | 0 | 1 | 1 |
| PCID2 | 1 | 1 | 0 | 1 | 1 |
| ZFAND6 | 1 | 1 | 0,234243533 | 0,268341796 | 1 |
| STARD10 | 2 | 2 | 0 | 1 | 1 |
| PORCN | 1 | 1 | 0 | 1 | 1 |
| SPEG | 6 | 6 | 1,64937E-17 | 0,999999993 | 1 |
| DIP2B | 20 | 19 | -0,010558579 | 0,578436485 | 1 |
| SMG8 | 2 | 2 | 0 | 1 | 1 |
| TMX2 | 10 | 10 | 1,47996E-16 | 0,999999951 | 1 |
| ATOX1 | 4 | 4 | 3,28441E-17 | 0,999999995 | 1 |
| EPHA7 | 5 | 5 | 0 | 1 | 1 |
| WARS2 | 6 | 6 | -4,88809E-16 | 0,999999945 | 1 |
| EHBP1 | 9 | 9 | -0,026725284 | 0,370087888 | 1 |
| HNRNPLL | 18 | 7 | 0,145939766 | 0,242314576 | 1 |
| TSNAX | 15 | 13 | -2,23553E-17 | 0,999999993 | 1 |
| ATP6V1B2 | 26 | 26 | 0,018589389 | 0,454589458 | 1 |
| ATP6V1B1 | 1 | 1 | 0 | 1 | 1 |
| AAMP | 4 | 3 | 5,08915E-19 | 0,999999999 | 1 |
| PALM | 1 | 1 | 0 | 1 | 1 |

| | | | | | |
|---|---|---|---|---|---|
| CUL2 | 32 | 30 | 0 | 1 | 1 |
| EMC7 | 6 | 6 | -3,14707E-18 | 0,999999999 | 1 |
| GRM8 | 3 | 3 | -0,127715221 | 0,122357103 | 1 |
| S100A16 | 1 | 1 | 0 | 1 | 1 |
| ME3;ME2 | 1 | 1 | -0,048901906 | 0,410758594 | 1 |
| ACOT9 | 30 | 29 | -2,76094E-17 | 0,999999992 | 1 |
| WDR13 | 19 | 18 | 0,043083422 | 0,134468081 | 1 |
| SYNPO | 18 | 18 | -0,084883291 | 0,272205085 | 1 |
| ARL6IP5 | 4 | 4 | 0 | 1 | 1 |
| FGF14 | 2 | 2 | 0 | 1 | 1 |
| CYRIB | 20 | 19 | -0,021518804 | 0,455896568 | 1 |
| MRPS34 | 7 | 7 | -2,46061E-18 | 0,999999996 | 1 |
| TPMT | 6 | 5 | 0,068853196 | 0,363035903 | 1 |
| NSUN3 | 1 | 1 | 0 | 1 | 1 |
| CYRIA | 15 | 13 | 0 | 1 | 1 |
| CRMP1 | 7 | 7 | 0,060208625 | 0,160592718 | 1 |
| SEC63 | 4 | 4 | 8,99521E-15 | 0,999999643 | 1 |
| HSD17B12 | 9 | 9 | 3,89282E-14 | 0,999999326 | 1 |
| COASY | 9 | 9 | 3,75115E-17 | 0,999999989 | 1 |
| APOOL | 7 | 7 | -3,66862E-15 | 0,999999859 | 1 |
| PTPRJ | 11 | 11 | 0,021526799 | 0,480667944 | 1 |
| NYAP2 | 4 | 4 | 0 | 1 | 1 |
| CLPP | 13 | 13 | -3,94887E-16 | 0,999999953 | 1 |
| NDUFB10 | 10 | 10 | 0 | 1 | 1 |
| HCCS | 3 | 3 | 0 | 1 | 1 |
| SLC38A3 | 6 | 6 | 4,9402E-18 | 0,999999997 | 1 |
| GSPT2 | 18 | 16 | 5,03125E-16 | 0,999999928 | 1 |
| PLS1 | 6 | 5 | 0 | 1 | 1 |
| MTCO1 | 3 | 3 | 1,60014E-18 | 0,999999998 | 1 |
| TPH2 | 10 | 10 | 0,032829802 | 0,522773026 | 1 |
| EIF1A;EIF1AX | 3 | 2 | -3,00139E-17 | 0,999999996 | 1 |
| TMCC3 | 5 | 5 | -0,077788813 | 0,227673781 | 1 |
| SARDH | 8 | 7 | -0,072511504 | 0,179055545 | 1 |

| | | | | | |
|---|---|---|---|---|---|
| NRXN1 | 26 | 26 | -2,42305E-18 | 0,999999998 | 1 |
| TUBA3B;TUBA4A;TU | 2 | 2 | 0 | 1 | 1 |
| ERLEC1 | 5 | 5 | 2,88078E-17 | 0,99999999 | 1 |
| PTPRS;PTPRD | 7 | 7 | 0,0499422 | 0,244847222 | 1 |
| SF1 | 6 | 2 | -5,85585E-18 | 0,999999998 | 1 |
| IRF7 | 1 | 1 | 0 | 1 | 1 |
| RTN4 | 5 | 5 | 0 | 1 | 1 |
| NUBPL | 5 | 5 | 3,06313E-18 | 0,999999997 | 1 |
| ARPC3 | 11 | 11 | 2,97465E-15 | 0,999999784 | 1 |
| PDE10A | 6 | 6 | 2,27054E-17 | 0,99999999 | 1 |
| NAXE | 11 | 10 | -0,019945676 | 0,459222591 | 1 |
| RPS11 | 12 | 11 | 0 | 1 | 1 |
| MFF | 5 | 5 | 5,3659E-17 | 0,999999988 | 1 |
| KCNJ3 | 5 | 5 | 0 | 1 | 1 |
| MDGA1 | 5 | 5 | 0,050541552 | 0,292170797 | 1 |
| EIF4H | 11 | 11 | 0 | 1 | 1 |
| GLB1 | 8 | 8 | 0,025034128 | 0,479992099 | 1 |
| PGAP4 | 1 | 1 | 0 | 1 | 1 |
| VPS50 | 27 | 27 | 0,041690766 | 0,202996474 | 1 |
| DIRAS2;DIRAS1 | 2 | 2 | -2,92706E-17 | 0,999999995 | 1 |
| MEMO1 | 5 | 5 | 0 | 1 | 1 |
| HAPLN1 | 21 | 20 | -0,031567411 | 0,429879427 | 1 |
| RAP1B | 13 | 13 | 0,034039826 | 0,417980313 | 1 |
| RAB3D;RAP1B | 1 | 1 | 0 | 1 | 1 |
| RAB3A | 15 | 15 | 4,5074E-17 | 0,999999981 | 1 |
| RAB3C | 11 | 10 | 0,0321987 | 0,331540673 | 1 |
| ACTBL2 | 8 | 8 | 0 | 1 | 1 |
| SGIP1 | 21 | 21 | 2,5607E-16 | 0,99999994 | 1 |
| SIRT2 | 16 | 16 | -0,033217601 | 0,537580184 | 1 |
| SDHB | 18 | 18 | -0,020055728 | 0,549858978 | 1 |
| MAPT | 23 | 23 | -0,011276032 | 0,488544653 | 1 |
| THEM6 | 4 | 4 | 7,28886E-18 | 0,999999994 | 1 |
| PIP4K2C | 20 | 20 | 0,057463848 | 0,155242449 | 1 |

| | | | | | |
|---|---|---|---|---|---|
| OLFM1 | 13 | 13 | 6,2211E-17 | 0,999999977 | 1 |
| SLC1A4 | 12 | 12 | 0,029478097 | 0,203896616 | 1 |
| ZC3H3 | 1 | 1 | -0,053577159 | 0,500005868 | 1 |
| DSTN | 15 | 14 | 8,66562E-17 | 0,999999974 | 1 |
| GSTM5 | 25 | 25 | -5,79875E-16 | 0,999999947 | 1 |
| SLC1A2 | 16 | 16 | 0 | 1 | 1 |
| SLC1A3 | 14 | 14 | 2,40469E-17 | 0,999999995 | 1 |
| SYNE2 | 3 | 3 | -0,057785561 | 0,177283386 | 1 |
| TNRC18 | 2 | 2 | 0 | 1 | 1 |
| TXN | 4 | 4 | 0,011923375 | 0,597598289 | 1 |
| ACTB | 17 | 17 | 1,30276E-16 | 0,999999973 | 1 |
| TMEM200C | 1 | 1 | 0 | 1 | 1 |
| CSRP1 | 10 | 10 | 0 | 1 | 1 |
| PRUNE1 | 18 | 16 | 0,043724949 | 0,244169687 | 1 |
| PPP6R3 | 16 | 15 | -2,72309E-17 | 0,999999992 | 1 |
| KCTD12 | 13 | 13 | 2,20557E-17 | 0,999999984 | 1 |
| DNM1 | 4 | 4 | 0 | 1 | 1 |
| DNAJA1 | 17 | 17 | 0 | 1 | 1 |
| MTARC2 | 16 | 16 | -3,6786E-16 | 0,999999955 | 1 |
| KIAA0513 | 18 | 18 | 0 | 1 | 1 |
| PPP2R5B | 9 | 9 | -1,58893E-16 | 0,999999962 | 1 |
| CDC42;RHOG;RAC2 | 1 | 1 | 0 | 1 | 1 |
| PFN1 | 15 | 15 | 0,029113478 | 0,454689975 | 1 |
| NAE1 | 26 | 23 | 0 | 1 | 1 |
| RANGAP1 | 11 | 8 | 7,799E-19 | 0,999999999 | 1 |
| RMND1 | 6 | 6 | 0,021727788 | 0,582383117 | 1 |
| S100A1 | 4 | 4 | 3,33988E-16 | 0,999999981 | 1 |
| FLOT2 | 28 | 28 | 0,019277951 | 0,402836866 | 1 |
| GFUS | 8 | 8 | -3,4823E-17 | 0,999999985 | 1 |
| SRSF2 | 5 | 1 | 0 | 1 | 1 |
| APOE | 25 | 21 | 5,32496E-17 | 0,999999976 | 1 |
| CCSAP | 11 | 10 | 0,016821882 | 0,601063532 | 1 |
| MTIF2 | 6 | 6 | -0,136232446 | 0,120509641 | 1 |

| | | | | | |
|---|---|---|---|---|---|
| AARS2 | 8 | 8 | -2,50399E-17 | 0,999999987 | 1 |
| ARFGEF2;ARFGEF1 | 8 | 8 | 2,40459E-17 | 0,999999992 | 1 |
| TWF1 | 17 | 17 | 0,022428849 | 0,255036702 | 1 |
| AGPAT4 | 6 | 6 | 0 | 1 | 1 |
| TM9SF3 | 6 | 6 | -0,076033528 | 0,154077062 | 1 |
| WDR48 | 22 | 22 | 0 | 1 | 1 |
| HSPA5 | 41 | 41 | -1,24661E-18 | 0,999999998 | 1 |
| HSPA2 | 28 | 28 | 0 | 1 | 1 |
| AGAP1 | 7 | 7 | 4,75082E-17 | 0,99999998 | 1 |
| NIPSNAP2 | 15 | 15 | -2,07769E-16 | 0,999999977 | 1 |
| MCU | 20 | 20 | 2,99927E-17 | 0,999999993 | 1 |
| UROS | 7 | 7 | 0 | 1 | 1 |
| SOWAHC | 1 | 1 | 0 | 1 | 1 |
| DNAAF3 | 1 | 1 | 0 | 1 | 1 |
| PEX19 | 2 | 2 | -0,060389256 | 0,358196871 | 1 |
| ISLR2 | 7 | 7 | 0,008453741 | 0,624517088 | 1 |
| NPL | 5 | 5 | -2,48484E-19 | 1 | 1 |
| TTC38 | 9 | 6 | -6,82215E-19 | 0,999999999 | 1 |
| ATP5F1E | 3 | 3 | -1,995E-17 | 0,99999999 | 1 |
| TRIM23 | 1 | 1 | 5,98906E-18 | 0,999999995 | 1 |
| CPSF1 | 1 | 1 | 0 | 1 | 1 |
| sp\|Q8R3C1\|CB042_ | 1 | 1 | 0 | 1 | 1 |
| AGPAT3 | 8 | 8 | 0 | 1 | 1 |
| CCT6B | 3 | 3 | 1,61705E-16 | 0,999999959 | 1 |
| DAGLA | 15 | 15 | -9,65487E-17 | 0,999999959 | 1 |
| PURG | 7 | 6 | 0 | 1 | 1 |
| RPS4X | 21 | 19 | 0,058813764 | 0,397114671 | 1 |
| ERC1 | 17 | 17 | 2,91328E-17 | 0,99999998 | 1 |
| ELAVL3 | 13 | 11 | -4,7165E-18 | 0,999999998 | 1 |
| PRDX6 | 5 | 5 | 0,014893771 | 0,57692602 | 1 |
| TRAK1 | 1 | 1 | 0 | 1 | 1 |
| FAIM2 | 2 | 2 | 0,023487924 | 0,484791824 | 1 |
| FHDC1 | 1 | 1 | 0 | 1 | 1 |

| | | | | | |
|---|---|---|---|---|---|
| GPRIN3 | 9 | 9 | -8,26203E-18 | 0,999999995 | 1 |
| TCEAL5 | 8 | 3 | -0,032883933 | 0,474241902 | 1 |
| HS1BP3 | 5 | 5 | -0,100923752 | 0,145581148 | 1 |
| UBE4A | 7 | 7 | -0,01011433 | 0,634214021 | 1 |
| KPNA4 | 5 | 4 | 1,32672E-17 | 0,999999993 | 1 |
| ASAH1 | 15 | 15 | 1,12555E-17 | 0,999999999 | 1 |
| EIF2S2 | 9 | 8 | 3,80157E-17 | 0,999999993 | 1 |
| RAB3B | 11 | 11 | -2,61938E-18 | 0,999999999 | 1 |
| ISOC2A | 2 | 2 | -1,45388E-19 | 1 | 1 |
| CRACD | 6 | 5 | 2,4454E-16 | 0,999999991 | 1 |
| EMC8 | 4 | 4 | 2,41465E-18 | 0,999999996 | 1 |
| NMD3 | 1 | 1 | 0 | 1 | 1 |
| DTX3 | 3 | 3 | 1,14194E-12 | 0,999997175 | 1 |
| ELMOD1 | 3 | 3 | 3,73412E-16 | 0,999999963 | 1 |
| NT5C2 | 9 | 9 | 0 | 1 | 1 |
| DYNC1LI2 | 13 | 13 | -8,36418E-17 | 0,999999968 | 1 |
| ARF3;ARF1 | 6 | 6 | 1,30797E-18 | 0,999999999 | 1 |
| ARF5 | 15 | 15 | 0,094854871 | 0,204566752 | 1 |
| ARF2 | 4 | 4 | -9,97481E-19 | 0,999999999 | 1 |
| SLC25A51 | 11 | 11 | 0 | 1 | 1 |
| PSMB3 | 8 | 8 | 0,039320094 | 0,421000202 | 1 |
| EIF4E | 6 | 6 | 0,036508343 | 0,380647075 | 1 |
| H4F16 | 8 | 6 | 0,646867085 | 0,159682384 | 1 |
| SLC44A2 | 9 | 9 | 1,15805E-17 | 0,999999997 | 1 |
| API5 | 8 | 3 | -6,6017E-18 | 0,999999995 | 1 |
| HSFY2 | 1 | 1 | 0 | 1 | 1 |
| HPCAL1 | 8 | 8 | 0,040168106 | 0,264197927 | 1 |
| PTGR3 | 11 | 11 | 0 | 1 | 1 |
| MT3 | 2 | 2 | 0 | 1 | 1 |
| GAL3ST3 | 2 | 2 | -0,088253567 | 0,141785658 | 1 |
| MCF2L | 1 | 1 | 0 | 1 | 1 |
| CALR | 21 | 21 | 3,58208E-19 | 0,999999999 | 1 |
| LDHB | 26 | 25 | -1,54605E-18 | 1 | 1 |

| | | | | | |
|---|---|---|---|---|---|
| NEDD4 | 12 | 11 | 0 | 1 | 1 |
| CADM4 | 12 | 12 | -1,51096E-15 | 0,999999853 | 1 |
| TAGLN2 | 12 | 7 | 7,45857E-19 | 0,999999999 | 1 |
| SUPT5H | 14 | 5 | 0 | 1 | 1 |
| ATP6V1F | 9 | 9 | 3,94728E-17 | 0,999999989 | 1 |
| TMX3 | 8 | 8 | 2,77895E-17 | 0,999999988 | 1 |
| SAMM50 | 19 | 19 | 5,95489E-18 | 0,999999998 | 1 |
| KCNA1 | 10 | 10 | 9,93132E-17 | 0,99999996 | 1 |
| RABGAP1 | 17 | 17 | 7,5173E-17 | 0,999999972 | 1 |
| GRIPAP1 | 32 | 32 | -1,05321E-16 | 0,999999957 | 1 |
| SLMAP | 7 | 7 | 0 | 1 | 1 |
| STX17 | 6 | 6 | 0,069111546 | 0,135134247 | 1 |
| PRDX6 | 24 | 24 | -2,90996E-18 | 0,999999998 | 1 |
| SLC25A46 | 11 | 11 | 6,12669E-17 | 0,999999989 | 1 |
| PCMT1 | 15 | 14 | 0,040187103 | 0,141680942 | 1 |
| CDC42 | 7 | 7 | 0,10016734 | 0,124507902 | 1 |
| VSNL1 | 23 | 23 | 0,038512807 | 0,151017792 | 1 |
| JAM3 | 7 | 7 | -0,009110475 | 0,743930027 | 1 |
| ERP44 | 11 | 11 | 2,3777E-16 | 0,999999938 | 1 |
| PPFIA4 | 25 | 25 | -9,81908E-17 | 0,999999956 | 1 |
| ITM2B | 7 | 7 | 2,74813E-16 | 0,999999946 | 1 |
| GTPBP1 | 8 | 8 | 0,014556355 | 0,625127994 | 1 |
| CRTAC1 | 8 | 8 | 0 | 1 | 1 |
| GRIN2A | 29 | 29 | 0,054476994 | 0,116305928 | 1 |
| GGT7 | 13 | 13 | -1,01404E-17 | 0,999999994 | 1 |
| ARHGAP21 | 2 | 2 | 0,022164817 | 0,570485573 | 1 |
| KIF5B | 28 | 26 | 0 | 1 | 1 |
| MBOAT2 | 1 | 1 | 0 | 1 | 1 |
| ATG9A | 9 | 9 | -1,65874E-16 | 0,99999999 | 1 |
| NRXN1 | 13 | 13 | 0 | 1 | 1 |
| FKBP15 | 5 | 5 | -0,03207709 | 0,452327785 | 1 |
| PMPCA | 21 | 21 | -0,104756681 | 0,164729502 | 1 |
| SLC3A2 | 23 | 23 | 3,43786E-17 | 0,999999985 | 1 |

| | | | | | |
|---|---|---|---|---|---|
| PPP2R2A | 20 | 19 | 4,68346E-16 | 0,999999924 | 1 |
| FSCN1 | 30 | 30 | -5,97122E-17 | 0,999999978 | 1 |
| NPM1 | 10 | 6 | 0 | 1 | 1 |
| DDAH1 | 24 | 24 | 0 | 1 | 1 |
| AKR1D1 | 1 | 1 | -0,22813498 | 0,145336182 | 1 |
| UBE4B | 15 | 15 | 0 | 1 | 1 |
| GMPS | 30 | 29 | 0,018220724 | 0,478606201 | 1 |
| TTC19 | 7 | 7 | -4,27075E-17 | 0,999999986 | 1 |
| CD9 | 3 | 3 | -0,081373304 | 0,141260639 | 1 |
| HSDL2 | 6 | 6 | 0 | 1 | 1 |
| AKT3 | 14 | 14 | 0,038282282 | 0,291993215 | 1 |
| EPB41L3 | 17 | 17 | -1,4967E-16 | 0,999999961 | 1 |
| SHROOM2 | 13 | 12 | 0 | 1 | 1 |
| VAC14 | 16 | 16 | -0,037341405 | 0,11899695 | 1 |
| TTLL12 | 14 | 13 | 3,30759E-17 | 0,999999981 | 1 |
| KYAT1 | 10 | 7 | 0 | 1 | 1 |
| PDLIM1 | 3 | 3 | -0,113243146 | 0,131144652 | 1 |
| LGI2 | 8 | 8 | -3,45521E-17 | 0,999999987 | 1 |
| G3BP1 | 8 | 7 | -1,19032E-15 | 0,999999893 | 1 |
| KCND2 | 11 | 11 | 0 | 1 | 1 |
| ACAD9 | 29 | 29 | -0,03903255 | 0,40872986 | 1 |
| ATXN7L3B | 1 | 1 | 3,67504E-23 | 1 | 1 |
| VPS33A | 8 | 8 | -1,36185E-19 | 1 | 1 |
| EIF3M | 8 | 6 | 0 | 1 | 1 |
| EIF2B3 | 10 | 9 | -4,59454E-17 | 0,999999984 | 1 |
| TMX1 | 4 | 4 | -5,94807E-18 | 0,999999996 | 1 |
| TLR4 | 1 | 1 | 0 | 1 | 1 |
| TMEM65 | 6 | 6 | 1,23739E-17 | 0,999999996 | 1 |
| EXOC6B | 16 | 16 | 0 | 1 | 1 |
| PDHB | 26 | 26 | -4,90155E-18 | 0,999999997 | 1 |
| DDX39B | 8 | 4 | 0,16817698 | 0,188766516 | 1 |
| TIMM10B | 3 | 3 | 2,83398E-17 | 0,999999998 | 1 |
| KALRN;ARHGEF25 | 1 | 1 | 0 | 1 | 1 |

| | | | | | |
|---|---|---|---|---|---|
| TIAM1 | 4 | 3 | 0 | 1 | 1 |
| PPIB | 13 | 13 | 0 | 1 | 1 |
| PPIL1 | 5 | 5 | 0 | 1 | 1 |
| RPL9 | 8 | 8 | 0,108197236 | 0,119456955 | 1 |
| PDLIM5 | 3 | 3 | -2,05497E-16 | 0,999999959 | 1 |
| STX12 | 10 | 10 | 0 | 1 | 1 |
| NDUFB6 | 9 | 9 | -4,57457E-17 | 0,999999995 | 1 |
| PDIA4 | 34 | 34 | -3,77018E-16 | 0,999999936 | 1 |
| SEPTIN4 | 4 | 4 | -0,034495706 | 0,438577763 | 1 |
| NDUFS3 | 17 | 17 | 0 | 1 | 1 |
| ANK3;ANK2 | 6 | 5 | 7,7361E-17 | 0,999999969 | 1 |
| DAB1 | 4 | 4 | 0 | 1 | 1 |
| PDLIM4 | 1 | 1 | 0 | 1 | 1 |
| FEZ1 | 3 | 3 | 0 | 1 | 1 |
| PGRMC2 | 4 | 4 | -2,12177E-13 | 0,999998675 | 1 |
| USP46 | 4 | 4 | 0 | 1 | 1 |
| HIBADH | 11 | 11 | -1,89345E-16 | 0,999999972 | 1 |
| PGRMC1 | 11 | 11 | -1,03302E-17 | 0,999999993 | 1 |
| ATP5MF | 5 | 5 | -2,32395E-17 | 0,999999995 | 1 |
| EIF4A2 | 11 | 11 | 0,048413381 | 0,193822572 | 1 |
| STARD5 | 2 | 2 | -0,052374842 | 0,374266464 | 1 |
| AFG3L2 | 42 | 42 | -1,50094E-15 | 0,999999903 | 1 |
| PLAA | 24 | 24 | 0,025190038 | 0,286057439 | 1 |
| PCK2 | 23 | 23 | 0 | 1 | 1 |
| DCUN1D1 | 8 | 8 | 3,21391E-17 | 0,999999984 | 1 |
| MCM6 | 1 | 1 | 2,25077E-18 | 0,999999997 | 1 |
| CADM2 | 18 | 18 | -3,48437E-18 | 0,999999994 | 1 |
| POR | 22 | 22 | 0 | 1 | 1 |
| B630019K06RIK | 3 | 3 | -0,089753869 | 0,123425727 | 1 |
| FSCN2 | 1 | 1 | -0,138207404 | 0,324947371 | 1 |
| MTX1 | 8 | 8 | -2,42357E-16 | 0,999999968 | 1 |
| ERP29 | 9 | 9 | 0 | 1 | 1 |
| ARSB | 8 | 8 | 0 | 1 | 1 |

| | | | | | |
|---|---|---|---|---|---|
| CAPN2 | 22 | 19 | 2,27716E-18 | 0,999999997 | 1 |
| RASA3 | 14 | 14 | -2,17242E-17 | 0,999999986 | 1 |
| ADPRH | 13 | 13 | 5,70255E-14 | 0,999999622 | 1 |
| TRAPPC1 | 4 | 4 | 0,083671343 | 0,126583489 | 1 |
| SELENBP1 | 20 | 20 | -5,06379E-17 | 0,999999974 | 1 |
| IGHM | 9 | 2 | 4,426936769 | 0,99988134 | 1 |
| IGLON5 | 4 | 4 | -0,057597867 | 0,143936198 | 1 |
| AHCYL2 | 13 | 12 | 0 | 1 | 1 |
| GSTZ1 | 11 | 11 | -0,053806898 | 0,396915108 | 1 |
| UBAP2L | 18 | 15 | 0,002655117 | 0,811445954 | 1 |
| SLC12A7 | 1 | 1 | 0 | 1 | 1 |
| WDR37 | 17 | 17 | 0,039814758 | 0,2750622 | 1 |
| SYN2 | 31 | 31 | 0,063971576 | 0,174161368 | 1 |
| ARHGAP5 | 18 | 18 | -0,015376291 | 0,520946665 | 1 |
| ARNT | 1 | 1 | 0,07128977 | 0,391619234 | 1 |
| PCDHGA2 | 1 | 1 | -1,52594E-18 | 0,999999998 | 1 |
| FUCA1 | 1 | 1 | 0 | 1 | 1 |
| ACOT1 | 3 | 3 | -3,96637E-13 | 0,999999081 | 1 |
| G6PDX | 24 | 23 | 2,05095E-12 | 0,999995967 | 1 |
| DNAJB11 | 8 | 8 | 2,34254E-17 | 0,999999986 | 1 |
| MDP1 | 5 | 5 | 2,23995E-16 | 0,999999969 | 1 |
| DNAJA2 | 19 | 19 | 0,038293161 | 0,156849443 | 1 |
| CALM1;CALM2;CALN | 9 | 9 | 0 | 1 | 1 |
| NETO1 | 2 | 2 | 1,16113E-17 | 0,999999995 | 1 |
| SOS1 | 2 | 2 | 0,056950619 | 0,234338634 | 1 |
| EGLN3 | 1 | 1 | 0 | 1 | 1 |
| VIM | 13 | 12 | 2,21983E-16 | 0,999999976 | 1 |
| DNAJB1 | 7 | 7 | 0 | 1 | 1 |
| CPNE5 | 6 | 6 | -2,01326E-17 | 0,999999996 | 1 |
| MAP4K3 | 7 | 7 | 3,39495E-17 | 0,999999986 | 1 |
| DNAJB4 | 8 | 7 | 0 | 1 | 1 |
| MCCC1 | 29 | 29 | -0,019088298 | 0,622635738 | 1 |
| WNK1 | 10 | 10 | -2,20927E-16 | 0,999999958 | 1 |

| | | | | | |
|---|---|---|---|---|---|
| FHIP1B | 8 | 6 | -2,14798E-17 | 0,999999998 | 1 |
| FGFR3 | 2 | 2 | -0,096802332 | 0,138462989 | 1 |
| MPP3 | 11 | 11 | 0 | 1 | 1 |
| ZC3H13 | 1 | 1 | 0 | 1 | 1 |
| HMCN1 | 1 | 1 | 0 | 1 | 1 |
| ELP2 | 8 | 8 | 0 | 1 | 1 |
| UEVLD | 2 | 2 | 0 | 1 | 1 |
| MACROD2 | 9 | 9 | -2,17548E-18 | 0,999999999 | 1 |
| RHOA | 5 | 5 | 4,43172E-17 | 0,999999985 | 1 |
| AHI1 | 4 | 4 | -1,01654E-14 | 0,99999971 | 1 |
| CAPRIN1 | 14 | 14 | -0,099828074 | 0,164454264 | 1 |
| RPS21 | 6 | 4 | -4,59459E-16 | 0,999999947 | 1 |
| NDUFC2 | 8 | 8 | -1,87365E-17 | 0,999999993 | 1 |
| LMAN2 | 8 | 8 | 1,52009E-17 | 0,999999989 | 1 |
| SPRED1 | 6 | 6 | -0,054785171 | 0,210344197 | 1 |
| NUDT16;NUDT16L1 | 1 | 1 | 0 | 1 | 1 |
| ABAT | 38 | 38 | -0,091391115 | 0,234259072 | 1 |
| OAS1F | 1 | 1 | 0 | 1 | 1 |
| ITIH3 | 4 | 3 | -7,61305E-19 | 1 | 1 |
| UBE2I | 4 | 2 | 5,51832E-18 | 0,999999998 | 1 |
| SEMA4D | 4 | 4 | -0,014852834 | 0,620680413 | 1 |
| PPME1 | 18 | 16 | 2,85273E-18 | 0,999999996 | 1 |
| ATP6AP2 | 4 | 4 | 0 | 1 | 1 |
| DNAJA4 | 12 | 12 | 1,48402E-16 | 0,999999961 | 1 |
| DRG1 | 9 | 9 | 1,70318E-17 | 0,999999994 | 1 |
| GUK1 | 10 | 10 | 4,84034E-17 | 0,999999977 | 1 |
| DYNLL1 | 8 | 8 | 0,044151488 | 0,350806006 | 1 |
| SH3BGRL | 9 | 8 | 0 | 1 | 1 |
| DYNLL2 | 2 | 2 | 0 | 1 | 1 |
| TLE2 | 1 | 1 | 0 | 1 | 1 |
| UGT8 | 2 | 2 | 0,132357816 | 0,14683093 | 1 |
| YWHAZ | 27 | 26 | 0,053710703 | 0,265684042 | 1 |
| LSMEM1 | 1 | 1 | 0 | 1 | 1 |

| | | | | | |
|---|---|---|---|---|---|
| IDH3G | 13 | 13 | 8,7501E-17 | 0,999999992 | 1 |
| GM49601;SEPTIN5 | 2 | 2 | -2,96783E-16 | 0,999999971 | 1 |
| MRPS6 | 1 | 1 | -0,155605262 | 0,151933225 | 1 |
| NDUFV2 | 15 | 14 | 0 | 1 | 1 |
| USP51 | 1 | 1 | -0,219233595 | 0,196229869 | 1 |
| SUCLG2 | 18 | 18 | -0,099544252 | 0,249583169 | 1 |
| EXOC6 | 6 | 6 | 4,62321E-15 | 0,99999982 | 1 |
| SLC27A4 | 23 | 23 | -8,39026E-17 | 0,999999984 | 1 |
| SLC24A2 | 7 | 7 | 0,031935827 | 0,441406803 | 1 |
| SULT4A1 | 11 | 11 | 0 | 1 | 1 |
| RPS29 | 2 | 2 | -6,69235E-19 | 0,999999999 | 1 |
| SBDS | 7 | 7 | 1,20378E-16 | 0,999999969 | 1 |
| MYO1D | 27 | 27 | -0,056825202 | 0,423596178 | 1 |
| ELMO2;ELMO1 | 3 | 3 | 2,67514E-16 | 0,999999961 | 1 |
| CYB5R1 | 15 | 15 | 3,76596E-17 | 0,999999984 | 1 |
| CDH10 | 7 | 7 | 6,4049E-17 | 0,999999985 | 1 |
| VARS2 | 11 | 11 | -1,77531E-17 | 0,999999992 | 1 |
| PSMD7 | 14 | 13 | 0,042685004 | 0,245964844 | 1 |
| HINT3 | 2 | 2 | 0,004817364 | 0,782455773 | 1 |
| NEK7 | 2 | 2 | 4,10022E-19 | 1 | 1 |
| CSNK1A1 | 9 | 9 | 3,77837E-17 | 0,999999983 | 1 |
| CHEK1 | 1 | 1 | 0,121076021 | 0,115523682 | 1 |
| CAMKK1 | 17 | 13 | 9,32003E-17 | 0,999999983 | 1 |
| RPS6KA1 | 5 | 5 | -0,045830823 | 0,357704915 | 1 |
| PITPNC1 | 9 | 9 | -3,37551E-18 | 0,999999998 | 1 |
| CYB5R3 | 17 | 17 | 2,59704E-17 | 0,999999985 | 1 |
| GABBR1 | 15 | 15 | 1,161E-16 | 0,999999972 | 1 |
| ZYG11B | 8 | 7 | 0,034703872 | 0,282048945 | 1 |
| DHRS7 | 4 | 4 | 0,010029056 | 0,696263342 | 1 |
| TUBGCP3 | 6 | 6 | -2,43484E-18 | 0,999999998 | 1 |
| TSTD3 | 2 | 2 | -0,011962028 | 0,715292269 | 1 |
| PCSK2 | 5 | 5 | 0 | 1 | 1 |
| UBE2M | 9 | 9 | 0,006407312 | 0,72812544 | 1 |

| | | | | | |
|---|---|---|---|---|---|
| FMR1 | 6 | 5 | 0,057862766 | 0,298075292 | 1 |
| GCC2 | 5 | 5 | 0,075497995 | 0,193197526 | 1 |
| LPCAT1 | 1 | 1 | 0 | 1 | 1 |
| NRK | 1 | 1 | 0 | 1 | 1 |
| PLXNC1 | 11 | 11 | 0,034401082 | 0,444462633 | 1 |
| DLG1 | 6 | 5 | -0,036443644 | 0,291856181 | 1 |
| VCAM1 | 15 | 15 | -1,31418E-17 | 0,999999995 | 1 |
| PSMD8 | 10 | 10 | 0 | 1 | 1 |
| ENTPD1 | 1 | 1 | 0,002962086 | 0,854449577 | 1 |
| MESD | 5 | 5 | -0,013921191 | 0,616156013 | 1 |
| GM7356 | 1 | 1 | -1,28408E-18 | 0,999999998 | 1 |
| USP10 | 9 | 9 | 8,92601E-16 | 0,999999924 | 1 |
| RPS26 | 2 | 2 | 2,90707E-16 | 0,999999974 | 1 |
| ARIH1 | 5 | 5 | 0,041336175 | 0,235947023 | 1 |
| ANP32B | 4 | 2 | 0 | 1 | 1 |
| HSP90B1 | 45 | 45 | 5,91036E-16 | 0,999999954 | 1 |
| PITHD1 | 11 | 11 | 0 | 1 | 1 |
| CCNY | 9 | 9 | 0,050431631 | 0,287414782 | 1 |
| ACTR3B;ACTR3 | 6 | 5 | 0,093279943 | 0,182781087 | 1 |
| IGKV5-39 | 1 | 1 | 0 | 1 | 1 |
| CPNE7;CPNE4;CPNE3 | 1 | 1 | 0,091297129 | 0,176128053 | 1 |
| RASGEF1A | 3 | 3 | 6,61923E-13 | 0,999998247 | 1 |
| VWA8 | 35 | 35 | -0,042162899 | 0,420163056 | 1 |
| DDX3Y;DDX3X | 17 | 16 | 1,57653E-16 | 0,999999964 | 1 |
| ACTR2 | 22 | 22 | 0,049345742 | 0,184884973 | 1 |
| HEPACAM | 11 | 11 | -9,03442E-18 | 0,99999999 | 1 |
| TBC1D13 | 6 | 6 | 0 | 1 | 1 |
| ANO3 | 1 | 1 | 0 | 1 | 1 |
| ME3 | 28 | 28 | -4,29413E-17 | 0,999999992 | 1 |
| 2310061I04RIK | 8 | 8 | 3,4088E-17 | 0,999999995 | 1 |
| CUL4B | 17 | 14 | 0 | 1 | 1 |
| ITIH2 | 2 | 2 | -8,1926E-18 | 1 | 1 |
| LRRC59 | 7 | 7 | 0 | 1 | 1 |

| | | | | | |
|---|---|---|---|---|---|
| TMED9 | 4 | 4 | 0 | 1 | 1 |
| TMED4 | 4 | 4 | 0,046230258 | 0,312336146 | 1 |
| BRSK1;BRSK2 | 2 | 2 | 0,012831649 | 0,675349927 | 1 |
| PLCB1 | 73 | 72 | 0,007701369 | 0,633308568 | 1 |
| MAP2 | 16 | 16 | -7,88102E-18 | 0,999999995 | 1 |
| RTN4 | 3 | 2 | 1,61334E-17 | 0,999999994 | 1 |
| CCDC25 | 4 | 3 | 0,12633885 | 0,181099575 | 1 |
| RARS2 | 11 | 11 | -1,22927E-17 | 0,999999992 | 1 |
| LDHA | 26 | 25 | 5,29644E-16 | 0,999999924 | 1 |
| GOLM1 | 2 | 2 | 5,45151E-18 | 0,999999997 | 1 |
| MOGS | 12 | 12 | 2,25521E-18 | 0,999999996 | 1 |
| SLC12A6 | 7 | 7 | 0,026317276 | 0,297813396 | 1 |
| AARSD1 | 12 | 10 | 0 | 1 | 1 |
| KMT2C | 1 | 1 | 0 | 1 | 1 |
| RAB4A | 9 | 8 | 0,057814908 | 0,199619835 | 1 |
| CFAP418 | 3 | 3 | 0,02106303 | 0,547242298 | 1 |
| KCMF1 | 3 | 3 | 0 | 1 | 1 |
| NDRG4 | 9 | 9 | 0 | 1 | 1 |
| SMAP2 | 9 | 8 | 0 | 1 | 1 |
| MDH1 | 19 | 18 | 1,66805E-17 | 0,999999995 | 1 |
| DNAJB2 | 7 | 7 | -1,92379E-16 | 0,999999952 | 1 |
| ALDH18A1 | 23 | 23 | -8,11967E-18 | 0,999999995 | 1 |
| CCAR1 | 1 | 1 | 0,066130999 | 0,627364094 | 1 |
| CLTB | 10 | 10 | -3,77537E-17 | 0,999999986 | 1 |
| EPM2AIP1 | 22 | 21 | 2,11263E-17 | 0,999999986 | 1 |
| GRHPR | 15 | 15 | -6,77469E-17 | 0,999999985 | 1 |
| OGDHL | 49 | 49 | -1,31692E-15 | 0,999999905 | 1 |
| UCK1 | 5 | 4 | 0 | 1 | 1 |
| MAPRE2 | 14 | 14 | 0 | 1 | 1 |
| APPL2 | 15 | 15 | -2,43045E-17 | 0,999999985 | 1 |
| FIBP | 7 | 7 | 0,038340899 | 0,269950262 | 1 |
| GPC4 | 12 | 12 | -0,016194628 | 0,571176676 | 1 |
| TPI1 | 20 | 18 | -3,8503E-17 | 0,999999984 | 1 |

| | | | | | |
|---|---|---|---|---|---|
| DPYSL4 | 28 | 28 | -5,88191E-17 | 0,999999978 | 1 |
| PLEKHO2 | 2 | 2 | -1,49435E-17 | 0,99999999 | 1 |
| GDPD5 | 1 | 1 | 0 | 1 | 1 |
| DIP2C | 15 | 15 | 0,029576139 | 0,404518742 | 1 |
| CEP170;CEP170B | 2 | 2 | -8,1913E-17 | 0,999999992 | 1 |
| GDI2 | 38 | 35 | 0 | 1 | 1 |
| TNRC6B | 3 | 3 | 6,03879E-14 | 0,999999474 | 1 |
| F3 | 7 | 7 | -1,18577E-15 | 0,99999992 | 1 |
| RNF187 | 1 | 1 | 0 | 1 | 1 |
| BCAT1 | 11 | 10 | -0,047589518 | 0,362680062 | 1 |
| TPT1 | 7 | 6 | 1,49766E-17 | 0,999999991 | 1 |
| BLVRA | 17 | 17 | -0,048919874 | 0,236874732 | 1 |
| HNRNPD | 13 | 8 | 0,072269953 | 0,222380551 | 1 |
| APPL1 | 23 | 22 | 1,7397E-17 | 0,999999991 | 1 |
| PRKCZ | 2 | 2 | 0 | 1 | 1 |
| CARMIL1 | 8 | 8 | 1,90316E-17 | 0,999999988 | 1 |
| GRK2 | 23 | 23 | 4,80723E-17 | 0,999999984 | 1 |
| ADRBK2 | 3 | 3 | 0,000160073 | 0,959388431 | 1 |
| SMOK3A;SMOK3B | 1 | 1 | 0 | 1 | 1 |
| GRK6 | 4 | 4 | 0,027632494 | 0,425292195 | 1 |
| PHKG1 | 3 | 3 | -0,001796648 | 0,882372782 | 1 |
| CAMK1 | 8 | 8 | -0,011236799 | 0,643352747 | 1 |
| CAMK1D | 15 | 14 | 0,061634751 | 0,12275379 | 1 |
| PRKAA2 | 10 | 10 | 0,056181601 | 0,164187916 | 1 |
| SNRK | 5 | 5 | 0 | 1 | 1 |
| CELF4 | 4 | 4 | 8,36455E-15 | 0,999999765 | 1 |
| CDK18 | 4 | 4 | -1,03668E-16 | 0,999999984 | 1 |
| CDK5 | 13 | 13 | 6,52527E-17 | 0,999999978 | 1 |
| CDK14 | 6 | 6 | 0,039630099 | 0,280809741 | 1 |
| PDPR | 16 | 16 | -2,42769E-16 | 0,99999997 | 1 |
| TPBG | 3 | 3 | 3,77658E-18 | 0,999999998 | 1 |
| ZC3H7B | 2 | 2 | -8,92972E-19 | 0,999999999 | 1 |
| RPS27L | 1 | 1 | 0,122796198 | 0,324692502 | 1 |

| | | | | | |
|---|---|---|---|---|---|
| CCDC22 | 10 | 10 | -0,017021763 | 0,430205655 | 1 |
| FRAS1 | 1 | 1 | 0,140409644 | 0,190734949 | 1 |
| SEC24B | 9 | 8 | 0 | 1 | 1 |
| NDUFAF4 | 8 | 8 | 0 | 1 | 1 |
| TMED10 | 9 | 9 | 0 | 1 | 1 |
| KIF1A | 3 | 3 | -0,079602163 | 0,181803495 | 1 |
| AIDA | 6 | 6 | 0,040969258 | 0,336897002 | 1 |
| HPRT1 | 15 | 14 | -3,97213E-18 | 0,999999997 | 1 |
| PGLS | 10 | 9 | 2,92035E-16 | 0,999999963 | 1 |
| CAMSAP3 | 1 | 1 | -3,14325E-17 | 0,999999988 | 1 |
| SNAP29 | 9 | 9 | 0,013019564 | 0,520077001 | 1 |
| CYLD | 11 | 11 | -1,03241E-16 | 0,999999975 | 1 |
| SMARCA2 | 2 | 1 | 0 | 1 | 1 |
| CASTOR2 | 6 | 5 | 0,000919149 | 0,921042436 | 1 |
| CHN1 | 6 | 5 | -1,66889E-17 | 0,999999997 | 1 |
| PIP4K2B | 25 | 24 | 2,09974E-16 | 0,999999947 | 1 |
| PSMD13 | 20 | 17 | 0 | 1 | 1 |
| URGCP | 3 | 1 | 0,019018512 | 0,744389821 | 1 |
| FNTA | 9 | 7 | -2,42411E-17 | 0,999999989 | 1 |
| SLC2A1 | 7 | 7 | -5,33943E-18 | 0,999999997 | 1 |
| HNMT | 10 | 6 | 0 | 1 | 1 |
| COPB1 | 18 | 17 | -0,078591307 | 0,143825256 | 1 |
| SYT1 | 32 | 32 | 0,048006158 | 0,119630342 | 1 |
| ASRGL1 | 13 | 12 | -6,25075E-17 | 0,99999999 | 1 |
| CAMSAP1 | 9 | 9 | 0 | 1 | 1 |
| CKAP4 | 31 | 31 | -0,017137711 | 0,521119573 | 1 |
| LRBA | 2 | 2 | 1,39748E-17 | 0,999999994 | 1 |
| NAA15;NAA16 | 5 | 3 | -3,18179E-15 | 0,999999858 | 1 |
| ERMN | 10 | 10 | -0,023214265 | 0,618839985 | 1 |
| ANP32A | 16 | 11 | 0,087436241 | 0,15877868 | 1 |
| ANP32E | 3 | 2 | 1,06977E-13 | 0,99999975 | 1 |
| ARHGAP35 | 26 | 26 | 0 | 1 | 1 |
| ACTA1;ACTBL2 | 12 | 11 | -1,82772E-17 | 0,99999999 | 1 |

| | | | | | |
|---|---|---|---|---|---|
| PPP1R9A | 7 | 7 | -0,040601081 | 0,429444364 | 1 |
| WDR54 | 7 | 7 | 6,88689E-17 | 0,999999986 | 1 |
| MLKL | 1 | 1 | 0 | 1 | 1 |
| GALT | 1 | 1 | 0 | 1 | 1 |
| LRRC58 | 1 | 1 | 0 | 1 | 1 |
| SIRT3 | 3 | 3 | 0 | 1 | 1 |
| HSPB11 | 2 | 2 | 0 | 1 | 1 |
| ZMYND10 | 1 | 1 | 0 | 1 | 1 |
| ACTG2 | 2 | 2 | 0 | 1 | 1 |
| SMIM8 | 1 | 1 | 0,121973743 | 0,127318001 | 1 |
| HNRNPR | 3 | 2 | -4,3324E-17 | 0,999999996 | 1 |
| VPS16 | 1 | 1 | 0 | 1 | 1 |
| TAGLN;TAGLN3 | 1 | 1 | 0 | 1 | 1 |
| DCAF7 | 7 | 7 | 1,12679E-16 | 0,999999961 | 1 |
| RPS15 | 3 | 2 | 0 | 1 | 1 |
| DDI2 | 8 | 8 | 0 | 1 | 1 |
| NDUFB8 | 8 | 8 | -1,30204E-16 | 0,99999997 | 1 |
| MMAA | 5 | 5 | -1,74471E-15 | 0,999999934 | 1 |
| EIF3D | 14 | 11 | -0,025617549 | 0,431777712 | 1 |
| CLVS2 | 7 | 6 | 0,040034271 | 0,286193768 | 1 |
| PIP4K2A | 17 | 17 | 0,03642219 | 0,263929007 | 1 |
| UBE2Q1 | 7 | 6 | 0 | 1 | 1 |
| PLEKHA2 | 3 | 3 | -9,71019E-17 | 0,999999991 | 1 |
| PLEKHA1 | 2 | 2 | 4,14686E-16 | 0,999999959 | 1 |
| LYN | 4 | 4 | -7,04621E-18 | 0,999999997 | 1 |
| ESPN | 1 | 1 | 0 | 1 | 1 |
| PHYHIPL | 16 | 16 | 0 | 1 | 1 |
| THY1 | 8 | 8 | 0,022560173 | 0,328125078 | 1 |
| NT5C3A | 7 | 5 | -5,63121E-16 | 0,999999932 | 1 |
| CSF1 | 1 | 1 | -4,69455E-19 | 0,999999999 | 1 |
| SNX16 | 6 | 6 | 1,14111E-05 | 0,985780736 | 1 |
| RPS17 | 10 | 9 | 0 | 1 | 1 |
| CYTH1 | 3 | 3 | -3,10006E-19 | 1 | 1 |

| | | | | | |
|---|---|---|---|---|---|
| SHPRH | 1 | 1 | 0 | 1 | 1 |
| TEX2 | 11 | 11 | 1,85482E-17 | 0,999999991 | 1 |
| FHIP1A | 1 | 1 | -1,99881E-18 | 0,999999998 | 1 |
| GM8251 | 2 | 2 | 0,082173393 | 0,154952952 | 1 |
| PSMD12 | 25 | 25 | 5,20061E-17 | 0,999999975 | 1 |
| NSDHL | 3 | 3 | 1,06134E-17 | 0,999999997 | 1 |
| DDX19A | 8 | 6 | 0 | 1 | 1 |
| BAG4 | 4 | 4 | 0 | 1 | 1 |
| SLC30A3 | 5 | 5 | -6,37106E-19 | 0,999999999 | 1 |
| NEGR1 | 10 | 10 | 1,40189E-16 | 0,999999981 | 1 |
| VBP1 | 12 | 11 | 0,027957554 | 0,387653054 | 1 |
| BRAF | 3 | 3 | 0,076798867 | 0,316574543 | 1 |
| HARS1;HARS2 | 3 | 3 | 1,43727E-19 | 1 | 1 |
| MYL6 | 8 | 8 | 0 | 1 | 1 |
| VMP1 | 2 | 2 | 0 | 1 | 1 |
| MAP1LC3A | 4 | 4 | -0,03122534 | 0,468979992 | 1 |
| BRAF | 23 | 22 | 0,025326818 | 0,231315072 | 1 |
| ATP1B3 | 9 | 9 | -4,79511E-16 | 0,999999958 | 1 |
| AKAP13 | 1 | 1 | 0,251546082 | 0,134788849 | 1 |
| KNDC1 | 2 | 2 | -9,38721E-18 | 0,999999994 | 1 |
| ARHGEF18 | 5 | 5 | 4,22098E-19 | 0,999999999 | 1 |
| ELMO3 | 1 | 1 | -0,187013618 | 0,262807846 | 1 |
| SH2D5 | 2 | 2 | -5,88139E-16 | 0,999999979 | 1 |
| NLGN3;NLGN1 | 1 | 1 | 0,082810357 | 0,257124115 | 1 |
| JAKMIP2 | 4 | 4 | 0 | 1 | 1 |
| DPH7 | 2 | 2 | 0 | 1 | 1 |
| RAD23A | 5 | 4 | -3,11698E-19 | 1 | 1 |
| C1QA | 5 | 5 | 0,038032353 | 0,533967281 | 1 |
| HDAC4 | 10 | 9 | 0 | 1 | 1 |
| MARK2 | 14 | 14 | 4,90193E-17 | 0,999999979 | 1 |
| PHKA1 | 9 | 9 | -0,002496177 | 0,823288627 | 1 |
| MTR | 8 | 5 | 4,23782E-18 | 0,999999996 | 1 |
| FAM171B | 12 | 12 | 0 | 1 | 1 |

| | | | | | |
|---|---|---|---|---|---|
| ABCD3 | 9 | 9 | 0,056393935 | 0,202946192 | 1 |
| PUM2;PUM1 | 2 | 2 | -4,7304E-19 | 1 | 1 |
| TTYH2 | 1 | 1 | -1,20975E-16 | 0,999999981 | 1 |
| PFN2 | 10 | 10 | 0,000905591 | 0,897813387 | 1 |
| STX1A | 27 | 27 | 0,033326427 | 0,179497587 | 1 |
| MCTP1 | 3 | 3 | 1,47141E-17 | 0,999999992 | 1 |
| ELP6 | 1 | 1 | 0 | 1 | 1 |
| TIMM29 | 11 | 11 | -5,01446E-17 | 0,999999983 | 1 |
| DOCK4 | 35 | 34 | 1,89768E-16 | 0,999999944 | 1 |
| NME2 | 6 | 6 | 3,54207E-16 | 0,999999941 | 1 |
| NME1 | 18 | 18 | 0,056401063 | 0,169636583 | 1 |
| CARHSP1 | 4 | 3 | 0 | 1 | 1 |
| REEP1 | 2 | 2 | 0 | 1 | 1 |
| GLS | 11 | 11 | -8,121E-17 | 0,999999993 | 1 |
| RAB6A | 11 | 11 | -4,584E-16 | 0,999999919 | 1 |
| NRIP3 | 3 | 3 | 0,094022929 | 0,315033107 | 1 |
| NRIP2 | 3 | 3 | 3,99178E-17 | 0,999999988 | 1 |
| MRPL4 | 6 | 6 | -0,043899091 | 0,294722377 | 1 |
| KLC2 | 4 | 4 | -0,051401692 | 0,20236878 | 1 |
| HCN2 | 9 | 9 | 1,35978E-17 | 0,99999999 | 1 |
| GRID1 | 7 | 7 | 0,02105832 | 0,410893772 | 1 |
| CBX3 | 7 | 3 | -3,75193E-19 | 1 | 1 |
| HSPE1 | 12 | 11 | -0,140488587 | 0,167733934 | 1 |
| MLF2 | 6 | 6 | -0,057608971 | 0,164642118 | 1 |
| KCNQ2 | 11 | 11 | -0,013418792 | 0,534072753 | 1 |
| KRAS | 13 | 13 | 0,005204198 | 0,734599437 | 1 |
| NKAIN4 | 1 | 1 | 0,061234514 | 0,373519818 | 1 |
| NCK1 | 5 | 5 | 8,23946E-19 | 0,999999999 | 1 |
| PPCS | 3 | 3 | 0 | 1 | 1 |
| CACNA2D1 | 2 | 2 | 1,30035E-15 | 0,999999946 | 1 |
| NDUFB7 | 8 | 8 | 0 | 1 | 1 |
| ANK1 | 19 | 19 | -0,031198297 | 0,277430519 | 1 |
| USP24 | 26 | 26 | 0 | 1 | 1 |

| | | | | | |
|---|---|---|---|---|---|
| IMMT | 16 | 15 | -3,56515E-16 | 0,99999997 | 1 |
| CBARP | 8 | 8 | 0,029571356 | 0,259831692 | 1 |
| TBC1D8B | 6 | 6 | 1,24073E-16 | 0,999999979 | 1 |
| SCN8A | 5 | 5 | 0 | 1 | 1 |
| SCN3A;SCN2A | 1 | 1 | 4,15299E-19 | 0,999999999 | 1 |
| THOP1 | 5 | 2 | -3,49673E-17 | 0,999999989 | 1 |
| CACNB3 | 11 | 11 | 0,033345306 | 0,276806207 | 1 |
| VPS4A | 5 | 5 | 0 | 1 | 1 |
| PUM1 | 6 | 5 | 0 | 1 | 1 |
| RPL36A | 2 | 1 | 0 | 1 | 1 |
| RAB21 | 10 | 10 | 0,04844187 | 0,153416829 | 1 |
| LGALS1 | 8 | 7 | -1,32507E-15 | 0,999999941 | 1 |
| SLITRK4 | 1 | 1 | 0,005694477 | 0,823597911 | 1 |
| GSK3B | 9 | 8 | 5,0039E-17 | 0,999999982 | 1 |
| MAP4K4 | 5 | 5 | 6,09206E-16 | 0,999999927 | 1 |
| CNR1 | 9 | 9 | 8,19423E-16 | 0,999999886 | 1 |
| CATSPERB | 1 | 1 | 0 | 1 | 1 |
| SLC25A10 | 8 | 8 | 7,73986E-18 | 0,999999998 | 1 |
| ARHGAP26 | 11 | 11 | 0 | 1 | 1 |
| GBA2 | 11 | 11 | 0 | 1 | 1 |
| SDCBP | 5 | 5 | -6,44695E-18 | 0,999999995 | 1 |
| CAMK1G | 1 | 1 | 4,05569E-20 | 1 | 1 |
| BLVRB | 7 | 6 | -1,86031E-16 | 0,999999967 | 1 |
| CAMSAP2 | 17 | 16 | -0,026634066 | 0,361637349 | 1 |
| CRKL | 14 | 12 | 3,67614E-17 | 0,999999986 | 1 |
| MCAT | 7 | 7 | -0,04830432 | 0,320692811 | 1 |
| TENM1 | 10 | 10 | -1,42134E-17 | 0,999999988 | 1 |
| SGSM1 | 11 | 11 | 4,18844E-17 | 0,999999983 | 1 |
| SFN;YWHAG;YWHAE | 4 | 4 | 0,097362772 | 0,168232332 | 1 |
| BRAP | 3 | 1 | 0 | 1 | 1 |
| CTU1 | 1 | 1 | -0,002120024 | 0,926365275 | 1 |
| IGKC | 7 | 1 | 0 | 1 | 1 |
| ABCC8 | 7 | 7 | -0,031953673 | 0,330234172 | 1 |

| | | | | | |
|---|---|---|---|---|---|
| MAPK8IP3 | 24 | 24 | 0,011797634 | 0,400866324 | 1 |
| WDR24 | 3 | 3 | 0 | 1 | 1 |
| TMEM106B | 6 | 6 | 9,71598E-18 | 0,999999991 | 1 |
| MICOS13 | 4 | 4 | -3,01555E-15 | 0,999999886 | 1 |
| CIRBP | 3 | 1 | -0,176818666 | 0,277338411 | 1 |
| MAG | 3 | 3 | 0 | 1 | 1 |
| OXSR1 | 7 | 7 | 0,001996124 | 0,857652038 | 1 |
| PEF1 | 6 | 6 | 3,06376E-18 | 0,999999996 | 1 |
| TMEM38B | 1 | 1 | -0,011913362 | 0,744496924 | 1 |
| SEPTIN11;SEPTIN6 | 7 | 7 | 0 | 1 | 1 |
| CALML3 | 3 | 3 | 0,085112426 | 0,241961018 | 1 |
| TRIM9 | 3 | 3 | 0,050792495 | 0,217976924 | 1 |
| RAP1A | 4 | 4 | 7,12931E-18 | 0,999999992 | 1 |
| COG3 | 2 | 1 | 0,291264376 | 0,238111478 | 1 |
| PPM1B | 3 | 3 | 6,5788E-18 | 0,999999995 | 1 |
| MARK2 | 3 | 3 | 0 | 1 | 1 |
| NT5C | 6 | 6 | -1,22698E-18 | 0,999999998 | 1 |
| MBP | 8 | 8 | -0,122150788 | 0,301855977 | 1 |
| RPS20 | 5 | 5 | 0,021508608 | 0,513417484 | 1 |
| PAK1 | 25 | 24 | 0,01941617 | 0,213026443 | 1 |
| DDOST | 11 | 11 | 1,31753E-16 | 0,999999967 | 1 |
| TENM1;TENM3 | 1 | 1 | 0 | 1 | 1 |
| GNA11;GNAQ | 7 | 7 | 3,67952E-15 | 0,999999859 | 1 |
| PALS2 | 1 | 1 | 2,08099E-17 | 0,999999991 | 1 |
| CETN1;CETN2 | 2 | 2 | 0 | 1 | 1 |
| GNAI1;GNAI2 | 4 | 4 | 0 | 1 | 1 |
| ANKRD49 | 1 | 1 | -0,203316425 | 0,254604411 | 1 |
| OLFML2B | 1 | 1 | 0 | 1 | 1 |
| BIN2 | 2 | 2 | 9,9133E-19 | 1 | 1 |
| YTHDF2 | 3 | 3 | 0 | 1 | 1 |
| NDUFS4 | 8 | 8 | -6,53391E-18 | 0,999999997 | 1 |
| SRPRB | 5 | 5 | 3,18436E-14 | 0,999999481 | 1 |
| NAA25 | 11 | 9 | 1,18546E-17 | 0,999999993 | 1 |

| | | | | | |
|---|---|---|---|---|---|
| UQCC2 | 4 | 4 | 9,33901E-19 | 0,999999999 | 1 |
| MRPL40 | 3 | 3 | 0 | 1 | 1 |
| EIF2B5 | 4 | 3 | 0 | 1 | 1 |
| PAOX | 10 | 8 | -5,52021E-18 | 0,999999996 | 1 |
| NDUFV3 | 7 | 7 | -0,021933832 | 0,69537502 | 1 |
| PLIN4 | 3 | 2 | 0 | 1 | 1 |
| DIRAS1 | 8 | 8 | 2,73368E-17 | 0,999999993 | 1 |
| GGPS1 | 6 | 6 | 2,26195E-19 | 0,999999999 | 1 |
| ADCY5;ADCY6 | 2 | 2 | 1,40189E-15 | 0,999999915 | 1 |
| CAB39 | 16 | 16 | 0,016513687 | 0,439215333 | 1 |
| MYH11 | 4 | 4 | -0,021050876 | 0,5273657 | 1 |
| BFSP2 | 1 | 1 | 0 | 1 | 1 |
| CTBP1 | 12 | 12 | 3,88134E-17 | 0,999999979 | 1 |
| KRT6A;KRT72;KRT5 | 4 | 3 | 0,19013753 | 0,433701185 | 1 |
| KRT73 | 5 | 5 | 0 | 1 | 1 |
| SNX6 | 16 | 16 | -1,6358E-17 | 0,99999999 | 1 |
| ARG1 | 1 | 1 | 0 | 1 | 1 |
| CMTM4 | 2 | 2 | 3,39412E-18 | 0,999999998 | 1 |
| RADIL | 1 | 1 | 0 | 1 | 1 |
| MRPL49 | 3 | 3 | -5,91356E-15 | 0,99999982 | 1 |
| ZC3H15 | 4 | 4 | 0,192882625 | 0,12183248 | 1 |
| PDZD8 | 7 | 7 | -1,68621E-16 | 0,999999982 | 1 |
| RPL27 | 6 | 5 | 6,2348E-15 | 0,999999719 | 1 |
| CCDC85A | 6 | 6 | -5,45349E-16 | 0,999999977 | 1 |
| RPS12 | 3 | 2 | -0,029223205 | 0,492349064 | 1 |
| PLG | 6 | 3 | 0,13335542 | 0,243327262 | 1 |
| STK38L | 5 | 5 | 5,86771E-17 | 0,99999999 | 1 |
| ROCK1;ROCK2 | 8 | 8 | 0,003067146 | 0,808158926 | 1 |
| CSNK2A1 | 18 | 18 | 1,38383E-16 | 0,999999959 | 1 |
| MAP2K7 | 5 | 5 | 0 | 1 | 1 |
| NAA50 | 9 | 7 | 0 | 1 | 1 |
| MCEE | 5 | 5 | -1,72538E-15 | 0,999999901 | 1 |
| GMPR2;GMPR | 2 | 2 | 0 | 1 | 1 |

| | | | | | |
|---|---|---|---|---|---|
| CERT1 | 7 | 3 | -0,088618477 | 0,370854286 | 1 |
| PARK7 | 20 | 16 | 2,06974E-17 | 0,999999986 | 1 |
| CDH9 | 4 | 4 | 1,00613E-18 | 0,999999999 | 1 |
| MT-CYB | 1 | 1 | 0,064375052 | 0,348060353 | 1 |
| SLC25A22 | 13 | 13 | 9,28236E-17 | 0,999999972 | 1 |
| QTRT1 | 2 | 2 | 3,29516E-18 | 0,999999997 | 1 |
| EPB41 | 2 | 2 | 0 | 1 | 1 |
| TMED8 | 4 | 4 | -9,89778E-17 | 0,999999971 | 1 |
| CFAP44 | 1 | 1 | 0 | 1 | 1 |
| CAPZA1 | 8 | 8 | 0,058835496 | 0,278717429 | 1 |
| SRP54 | 5 | 5 | 0 | 1 | 1 |
| CLCC1 | 5 | 4 | -4,16611E-19 | 1 | 1 |
| GDAP1L1 | 16 | 16 | -4,01064E-18 | 0,999999997 | 1 |
| PPP2R2C | 2 | 2 | 2,14355E-17 | 0,999999994 | 1 |
| MTHFSD | 1 | 1 | 0 | 1 | 1 |
| TRAPPC9 | 17 | 17 | 0 | 1 | 1 |
| MRPS10 | 2 | 2 | 0 | 1 | 1 |
| MGAT4B | 1 | 1 | 0 | 1 | 1 |
| WASF2 | 2 | 2 | 5,24045E-17 | 0,99999999 | 1 |
| FDX2 | 5 | 5 | -0,050710693 | 0,275835836 | 1 |
| NDUFAF6 | 1 | 1 | 0 | 1 | 1 |
| NTRK2;NTRK3 | 3 | 3 | 1,97368E-17 | 0,99999999 | 1 |
| DOCK7 | 8 | 8 | -1,1579E-58 | 1 | 1 |
| SLC6A9 | 4 | 4 | 4,05722E-18 | 0,999999996 | 1 |
| PSMD6 | 20 | 20 | 3,73041E-15 | 0,999999741 | 1 |
| GDI1 | 49 | 48 | 1,34293E-16 | 0,999999985 | 1 |
| ACSL4 | 9 | 9 | -2,11992E-18 | 0,999999999 | 1 |
| CDH2 | 13 | 13 | 7,39982E-18 | 0,999999996 | 1 |
| GLRX5 | 3 | 3 | 1,05937E-18 | 1 | 1 |
| RAB22A | 5 | 5 | -4,59684E-17 | 0,999999985 | 1 |
| RALGAPB | 14 | 14 | 2,85276E-18 | 0,999999998 | 1 |
| TUBA1A | 2 | 2 | 0 | 1 | 1 |
| TRAPPC3 | 6 | 5 | 0 | 1 | 1 |

| | | | | | |
|---|---|---|---|---|---|
| DPP6 | 28 | 28 | 0,039268371 | 0,247205767 | 1 |
| PPTC7 | 5 | 5 | -0,006476642 | 0,772782491 | 1 |
| PRDX4 | 6 | 6 | 1,99257E-16 | 0,999999965 | 1 |
| MAPK8;MAPK9 | 1 | 1 | 0 | 1 | 1 |
| MTRES1 | 2 | 2 | 1,38444E-15 | 0,999999931 | 1 |
| B3GLCT | 2 | 2 | 0,032257497 | 0,464355926 | 1 |
| PURA | 16 | 16 | -3,7596E-17 | 0,99999998 | 1 |
| APMAP | 14 | 14 | 0 | 1 | 1 |
| EMD | 6 | 6 | 0 | 1 | 1 |
| MAPRE3 | 16 | 16 | 0 | 1 | 1 |
| TECR | 11 | 11 | 7,66081E-17 | 0,999999976 | 1 |
| NECAB1 | 10 | 8 | -1,14491E-14 | 0,99999969 | 1 |
| AGO3;AGO1 | 2 | 2 | -4,76365E-17 | 0,999999984 | 1 |
| GOLGA7B | 3 | 3 | 0,036702587 | 0,367636635 | 1 |
| TMCC1 | 7 | 7 | 0 | 1 | 1 |
| CTTN | 27 | 27 | 0 | 1 | 1 |
| MTPN | 10 | 9 | 0 | 1 | 1 |
| EML6 | 5 | 5 | -2,73623E-17 | 0,999999983 | 1 |
| SGSM1 | 1 | 1 | 0 | 1 | 1 |
| UBTD2 | 3 | 3 | -1,62799E-18 | 0,999999998 | 1 |
| PYGL | 4 | 3 | 1,74188E-17 | 0,999999994 | 1 |
| CAD | 18 | 18 | -3,72428E-18 | 0,999999999 | 1 |
| ATP6V1G1 | 3 | 3 | 1,02978E-15 | 0,999999928 | 1 |
| AGRN | 2 | 2 | 0 | 1 | 1 |
| RPL13 | 9 | 8 | 0,068551769 | 0,392568857 | 1 |
| PEX3 | 1 | 1 | -0,112906877 | 0,147051719 | 1 |
| FAM120A | 23 | 20 | 0 | 1 | 1 |
| KRIT1 | 2 | 2 | 0 | 1 | 1 |
| NYAP1 | 1 | 1 | 0 | 1 | 1 |
| VANGL2 | 1 | 1 | 0 | 1 | 1 |
| GSTP2 | 1 | 1 | 0 | 1 | 1 |
| NAP1L4 | 11 | 10 | -1,09297E-17 | 0,999999992 | 1 |
| KPNA1 | 8 | 8 | -2,08522E-17 | 0,999999989 | 1 |

| | | | | | |
|---|---|---|---|---|---|
| KPNA6 | 4 | 4 | -8,32881E-19 | 0,999999999 | 1 |
| TRAPPC5 | 8 | 8 | 1,22598E-17 | 0,999999993 | 1 |
| EXOC4 | 25 | 25 | 9,83738E-18 | 0,999999996 | 1 |
| HEBP1 | 7 | 7 | 2,02094E-16 | 0,999999975 | 1 |
| PKM | 4 | 4 | 1,48251E-16 | 0,999999974 | 1 |
| CPLX1 | 5 | 5 | 0 | 1 | 1 |
| AKT1S1 | 2 | 2 | 4,64278E-17 | 0,999999983 | 1 |
| CFAP36 | 11 | 10 | -0,053991184 | 0,325958399 | 1 |
| KCTD8 | 6 | 6 | -0,030893433 | 0,317418709 | 1 |
| SCN1A;SCN3A;SCN2/ | 7 | 7 | 2,72038E-17 | 0,999999993 | 1 |
| ANK2 | 3 | 3 | -9,47173E-18 | 0,999999992 | 1 |
| ATG5 | 4 | 4 | 8,16423E-18 | 0,999999994 | 1 |
| OGFR | 9 | 8 | -1,17291E-17 | 0,999999998 | 1 |
| KCTD16 | 12 | 12 | -0,014928436 | 0,44647351 | 1 |
| NUDT19 | 1 | 1 | 0 | 1 | 1 |
| ARVCF | 13 | 13 | 5,37669E-17 | 0,999999985 | 1 |
| BCL2L1 | 2 | 2 | 0 | 1 | 1 |
| PCDH10 | 6 | 6 | 1,39348E-17 | 0,999999994 | 1 |
| SERBP1 | 15 | 10 | 0,03929947 | 0,321753608 | 1 |
| SPTAN1 | 5 | 5 | -0,039317464 | 0,368640206 | 1 |
| EEF1AKMT1 | 2 | 2 | -4,49455E-23 | 1 | 1 |
| DYNLT1 | 1 | 1 | 0 | 1 | 1 |
| NEB | 1 | 1 | 0 | 1 | 1 |
| DNPEP | 16 | 15 | 0,008241516 | 0,575860264 | 1 |
| STXBP5L | 25 | 25 | 0,034406472 | 0,165014381 | 1 |
| EPB41;EPB41L3 | 4 | 4 | 0 | 1 | 1 |
| FAM228A | 1 | 1 | 0 | 1 | 1 |
| SLC10A4 | 2 | 2 | 4,25484E-15 | 0,999999842 | 1 |
| GSTO1 | 16 | 16 | 1,43682E-37 | 1 | 1 |
| IMPA1 | 15 | 14 | -7,36361E-18 | 0,999999996 | 1 |
| GULO | 1 | 1 | 0 | 1 | 1 |
| FRMPD3 | 7 | 7 | 0 | 1 | 1 |
| ALG5 | 1 | 1 | 0 | 1 | 1 |

| | | | | | |
|---|---|---|---|---|---|
| DLG5 | 1 | 1 | 0 | 1 | 1 |
| CIAPIN1 | 5 | 5 | 0 | 1 | 1 |
| PLSCR3 | 3 | 3 | 1,92083E-17 | 0,999999997 | 1 |
| IPO9 | 18 | 17 | 2,9207E-17 | 0,999999982 | 1 |
| KLHL26 | 4 | 4 | 6,99961E-18 | 0,999999993 | 1 |
| PPAT | 8 | 8 | -1,22504E-17 | 0,999999995 | 1 |
| DKK3 | 5 | 5 | 3,21325E-18 | 0,999999998 | 1 |
| ACHE | 6 | 6 | -5,09908E-16 | 0,99999997 | 1 |
| TUBGCP4 | 2 | 2 | -0,045593589 | 0,377721457 | 1 |
| LASP1 | 18 | 18 | -4,31233E-16 | 0,999999915 | 1 |
| CEP104 | 1 | 1 | 0 | 1 | 1 |
| EARS2 | 10 | 10 | 5,93363E-18 | 0,999999998 | 1 |
| STOM | 4 | 4 | 0,049271802 | 0,380946327 | 1 |
| SLC49A4 | 1 | 1 | 0 | 1 | 1 |
| GLDC | 15 | 15 | -0,119089905 | 0,127136391 | 1 |
| CLINT1 | 12 | 12 | -3,09275E-17 | 0,999999984 | 1 |
| VPS4B;VPS4A | 2 | 2 | 5,71696E-16 | 0,99999993 | 1 |
| SLC44A1 | 9 | 9 | -7,74133E-18 | 0,999999997 | 1 |
| ETHE1 | 5 | 5 | -3,89115E-19 | 1 | 1 |
| SFXN5 | 12 | 12 | -0,11022337 | 0,270899595 | 1 |
| TSN | 15 | 13 | -9,94381E-16 | 0,999999931 | 1 |
| CDC42BPA | 15 | 15 | -1,22769E-16 | 0,999999963 | 1 |
| PITRM1 | 32 | 32 | -2,20206E-16 | 0,999999968 | 1 |
| STT3B | 8 | 8 | 0,052079908 | 0,381975391 | 1 |
| PSD | 11 | 11 | -1,49301E-17 | 0,999999991 | 1 |
| STT3A | 11 | 11 | 2,51177E-15 | 0,999999795 | 1 |
| INPP1 | 17 | 15 | -4,27147E-18 | 0,999999995 | 1 |
| MOB4 | 8 | 7 | 0 | 1 | 1 |
| MBOAT7 | 3 | 3 | 0,073655532 | 0,266910442 | 1 |
| CHGB | 3 | 3 | 0 | 1 | 1 |
| GPM6B | 1 | 1 | 1,41509E-18 | 0,999999998 | 1 |
| UBR4 | 8 | 8 | 0 | 1 | 1 |
| ACP1 | 4 | 3 | -0,065545941 | 0,131824609 | 1 |

| | | | | | |
|---|---|---|---|---|---|
| CR1L | 5 | 5 | 0 | 1 | 1 |
| TTC39C | 7 | 7 | 0,05100315 | 0,27178091 | 1 |
| TMX4 | 6 | 6 | 0 | 1 | 1 |
| RABGGTB | 5 | 5 | -7,65043E-16 | 0,999999929 | 1 |
| NARS1 | 26 | 23 | 9,39883E-16 | 0,999999902 | 1 |
| DARS2 | 10 | 10 | 0 | 1 | 1 |
| RWDD2B | 1 | 1 | -0,051668958 | 0,47922233 | 1 |
| NUDCD1 | 7 | 6 | 0 | 1 | 1 |
| ILF3 | 8 | 4 | 0 | 1 | 1 |
| HAX1 | 5 | 5 | 9,16663E-18 | 0,999999999 | 1 |
| LARS1 | 39 | 28 | 6,1087E-17 | 0,999999987 | 1 |
| RALGPS1 | 1 | 1 | 0,037558284 | 0,587389047 | 1 |
| GPR37 | 2 | 2 | -0,076556577 | 0,130670392 | 1 |
| SLITRK5 | 6 | 6 | 5,48026E-17 | 0,999999984 | 1 |
| RAB23 | 10 | 10 | 0,046468774 | 0,264655109 | 1 |
| RGS12 | 8 | 8 | 0 | 1 | 1 |
| GNG4 | 1 | 1 | 0 | 1 | 1 |
| GNG2 | 2 | 2 | 0 | 1 | 1 |
| CNKSR2 | 2 | 2 | 0 | 1 | 1 |
| RPN2 | 16 | 16 | 0,013081278 | 0,456751428 | 1 |
| PPP1R14C | 1 | 1 | 0 | 1 | 1 |
| HNRNPA1 | 12 | 10 | 0,08851634 | 0,372224218 | 1 |
| HNRNPA3 | 10 | 9 | 0,120497277 | 0,293139116 | 1 |
| SERPINB9 | 7 | 7 | 0 | 1 | 1 |
| GPT2 | 6 | 6 | 0,065942099 | 0,259058217 | 1 |
| YBX1;YBX3 | 3 | 3 | 0,059864236 | 0,492950978 | 1 |
| C2CD3 | 1 | 1 | 0,132991903 | 0,1386529 | 1 |
| RNF126 | 1 | 1 | -0,004926009 | 0,86075783 | 1 |
| MTDH | 15 | 14 | -2,19161E-17 | 0,999999988 | 1 |
| PIN1 | 6 | 6 | 0,064237418 | 0,309955741 | 1 |
| TMUB1 | 2 | 2 | 0 | 1 | 1 |
| ATP6V1G2 | 12 | 12 | 3,03562E-17 | 0,999999983 | 1 |
| CD82 | 5 | 5 | -2,0778E-16 | 0,999999974 | 1 |

| | | | | | |
|---|---|---|---|---|---|
| WASHC1 | 4 | 4 | 0 | 1 | 1 |
| ZNF445 | 1 | 1 | 0 | 1 | 1 |
| UQCRH | 4 | 4 | 0,088844893 | 0,147558228 | 1 |
| UBN1 | 1 | 1 | 0 | 1 | 1 |
| FAM162A | 9 | 9 | -5,60023E-17 | 0,999999991 | 1 |
| UBQLN2 | 17 | 12 | -0,030380421 | 0,167366344 | 1 |
| TUBB4A | 14 | 14 | -1,37336E-16 | 0,999999986 | 1 |
| MTCH2 | 12 | 12 | 0,003277282 | 0,839758153 | 1 |
| PNPLA6 | 2 | 2 | 4,76704E-15 | 0,999999862 | 1 |
| ALDH1A2 | 5 | 1 | 0 | 1 | 1 |
| ALDH1B1 | 21 | 20 | -5,97395E-14 | 0,999999456 | 1 |
| NDRG1 | 9 | 8 | -0,039102642 | 0,39635998 | 1 |
| MARK4 | 7 | 7 | 5,99816E-17 | 0,99999998 | 1 |
| DVL2 | 2 | 2 | 1,35948E-15 | 0,99999996 | 1 |
| MED7 | 1 | 1 | 0 | 1 | 1 |
| RABGAP1L | 3 | 3 | 5,2056E-16 | 0,999999948 | 1 |
| MLC1 | 6 | 6 | 0,034608838 | 0,311217301 | 1 |
| PCDH9 | 14 | 14 | -3,74215E-18 | 0,999999996 | 1 |
| COMMD8 | 2 | 2 | 0 | 1 | 1 |
| FKBPL | 1 | 1 | 0 | 1 | 1 |
| IGBP1B | 1 | 1 | 0,016933087 | 0,721909238 | 1 |
| NDUFAF2 | 10 | 10 | -4,63239E-17 | 0,999999988 | 1 |
| GPM6A | 7 | 7 | 0,037511367 | 0,456120773 | 1 |
| TYW5 | 1 | 1 | 0 | 1 | 1 |
| FIS1 | 7 | 7 | 0 | 1 | 1 |
| TMEM151B | 1 | 1 | -0,049241219 | 0,355769604 | 1 |
| PYCARD | 1 | 1 | 0 | 1 | 1 |
| HNRNPA2B1 | 13 | 12 | 0,099966661 | 0,349728392 | 1 |
| NES | 1 | 1 | -0,080986078 | 0,393684805 | 1 |
| SPOPL | 1 | 1 | 0 | 1 | 1 |
| TSC22D1 | 4 | 4 | -3,44193E-15 | 0,999999842 | 1 |
| ATP6V1B2;ATP6V1B1 | 8 | 8 | 0,043292224 | 0,226100746 | 1 |
| TUBB4B;TUBB5;TUBI | 1 | 1 | 0 | 1 | 1 |

| | | | | | |
|---|---|---|---|---|---|
| MIEN1 | 2 | 2 | 0 | 1 | 1 |
| CFAP95 | 1 | 1 | 0,015498541 | 0,699949591 | 1 |
| RAB1B | 9 | 9 | 3,53495E-17 | 0,999999981 | 1 |
| SNX27 | 18 | 17 | 0,005657657 | 0,589135215 | 1 |
| KCND3 | 7 | 7 | 0,023554423 | 0,520591904 | 1 |
| TTL | 7 | 7 | -0,030095404 | 0,478354888 | 1 |
| NTNG2 | 3 | 3 | 0,023163487 | 0,515754849 | 1 |
| DDAH2;DDAH1 | 1 | 1 | 0 | 1 | 1 |
| PRXL2A | 4 | 4 | 0 | 1 | 1 |
| STX7 | 9 | 9 | -3,28033E-17 | 0,999999982 | 1 |
| CPEB4;CPEB3;CPEB2 | 3 | 3 | 0 | 1 | 1 |
| SET | 9 | 2 | 6,91285E-15 | 0,999999864 | 1 |
| BEST3 | 1 | 1 | 0 | 1 | 1 |
| GBE1 | 15 | 15 | -1,23808E-14 | 0,999999786 | 1 |
| UBE2K | 8 | 8 | 0,045532151 | 0,342424147 | 1 |
| UBL3 | 2 | 2 | 0 | 1 | 1 |
| RPL5 | 18 | 16 | 4,05383E-17 | 0,999999985 | 1 |
| MAGEE2 | 3 | 3 | 0 | 1 | 1 |
| GNA11 | 12 | 12 | 0,033092753 | 0,297954514 | 1 |
| UCHL5 | 5 | 4 | 0 | 1 | 1 |
| HSCB | 7 | 7 | -0,072381161 | 0,290247111 | 1 |
| UCHL1 | 17 | 17 | 0 | 1 | 1 |
| PTPDC1 | 2 | 1 | 0,137365588 | 0,334442109 | 1 |
| MAIP1 | 5 | 5 | -1,54911E-17 | 0,999999998 | 1 |
| ATP1B2 | 14 | 14 | 3,71179E-16 | 0,999999955 | 1 |
| MAG | 11 | 11 | -0,054526648 | 0,465824634 | 1 |
| PYCR2 | 15 | 15 | -1,35874E-16 | 0,999999979 | 1 |
| PPT1 | 7 | 7 | 0 | 1 | 1 |
| ATP8A1 | 38 | 38 | 0,043844185 | 0,170625426 | 1 |
| SEZ6L2 | 12 | 12 | -0,035144717 | 0,184886729 | 1 |
| TMEM132B | 16 | 16 | 0,01000134 | 0,573984713 | 1 |
| RPS28 | 6 | 6 | -1,81618E-15 | 0,999999865 | 1 |
| TSPYL4 | 4 | 3 | 0,104064698 | 0,151998538 | 1 |

| | | | | | |
|---|---|---|---|---|---|
| KALRN;TRIO | 8 | 8 | 5,47276E-17 | 0,999999979 | 1 |
| RANBP6;IPO5 | 2 | 2 | 0 | 1 | 1 |
| MEGF9 | 1 | 1 | 0 | 1 | 1 |
| SLC25A16 | 3 | 3 | -0,044294356 | 0,545058759 | 1 |
| PRDX2 | 10 | 10 | -3,40739E-17 | 0,999999988 | 1 |
| HERC4 | 8 | 8 | 0,040793927 | 0,408187688 | 1 |
| ASPM | 1 | 1 | 0,161304488 | 0,367262937 | 1 |
| ATG13 | 1 | 1 | 5,54104E-15 | 0,99999986 | 1 |
| SLC7A10 | 1 | 1 | 2,76986E-19 | 0,999999999 | 1 |
| SLC7A8 | 5 | 5 | 0 | 1 | 1 |
| FBXO21 | 7 | 6 | -8,55325E-16 | 0,99999995 | 1 |
| FAIM | 4 | 4 | 1,698E-17 | 0,999999992 | 1 |
| IREB2 | 2 | 2 | -2,40111E-15 | 0,999999925 | 1 |
| UBB;UBC;RPS27A;UE | 8 | 8 | 0 | 1 | 1 |
| NEDD8 | 4 | 4 | 2,03132E-16 | 0,99999998 | 1 |
| CLCN3 | 2 | 2 | 0 | 1 | 1 |
| CCDC51 | 8 | 8 | -3,35173E-17 | 0,999999989 | 1 |
| CTNNB1;JUP | 2 | 2 | -2,40365E-16 | 0,999999991 | 1 |
| ERI3 | 2 | 2 | 0 | 1 | 1 |
| CLASP1;CLASP2 | 4 | 4 | -1,09242E-16 | 0,999999968 | 1 |
| TSPAN9 | 1 | 1 | 0,081705783 | 0,245116619 | 1 |
| ADAM11 | 10 | 10 | 0,037255301 | 0,316619415 | 1 |
| PPIA | 19 | 19 | 0 | 1 | 1 |
| ARAF | 4 | 4 | 3,64465E-16 | 0,999999957 | 1 |
| MYL1;MYL3 | 2 | 2 | 9,79251E-19 | 0,999999998 | 1 |
| GRIP2 | 3 | 3 | -0,006662879 | 0,726937992 | 1 |
| STK39 | 7 | 7 | 0,022311091 | 0,525996653 | 1 |
| MCTS1 | 5 | 3 | 0,109145963 | 0,116635045 | 1 |
| SLC45A1 | 1 | 1 | 0 | 1 | 1 |
| EMC6 | 1 | 1 | 0 | 1 | 1 |
| OPCML | 2 | 2 | 0,027583457 | 0,499073583 | 1 |
| SMG9 | 1 | 1 | 0 | 1 | 1 |
| MSRA | 10 | 10 | 0 | 1 | 1 |

| | | | | | |
|---|---|---|---|---|---|
| LSM12 | 3 | 3 | -6,23196E-18 | 0,999999994 | 1 |
| CNOT2 | 2 | 2 | -2,77951E-17 | 0,999999989 | 1 |
| HACD3 | 8 | 8 | 0 | 1 | 1 |
| NGEF | 14 | 14 | 0 | 1 | 1 |
| DBN1 | 1 | 1 | -0,088487249 | 0,376192907 | 1 |
| MT-CO3 | 1 | 1 | 0 | 1 | 1 |
| SASH1 | 3 | 3 | -0,005694511 | 0,785059929 | 1 |
| SNCB | 6 | 6 | 3,30754E-16 | 0,999999959 | 1 |
| RHOB | 9 | 9 | 8,51958E-17 | 0,99999997 | 1 |
| ARFIP1 | 8 | 8 | 7,75908E-15 | 0,999999708 | 1 |
| RTCB | 21 | 19 | 0 | 1 | 1 |
| PYM1 | 3 | 3 | 3,77442E-17 | 0,999999987 | 1 |
| EEF1A1;EEF1A2 | 14 | 13 | 0,041981271 | 0,222350373 | 1 |
| CFAP300 | 1 | 1 | 0,174976126 | 0,346152958 | 1 |
| UBXN2B | 4 | 4 | 4,54331E-18 | 0,999999995 | 1 |
| GOPC | 4 | 4 | -0,046784925 | 0,286408094 | 1 |
| SPOPL | 1 | 1 | 0 | 1 | 1 |
| NMT2 | 11 | 11 | 1,64695E-16 | 0,999999973 | 1 |
| VDAC1 | 23 | 23 | -1,07404E-17 | 0,999999996 | 1 |
| UQCRC1 | 24 | 24 | -5,69656E-17 | 0,999999986 | 1 |
| MRPL48 | 1 | 1 | 4,98709E-17 | 0,999999986 | 1 |
| NDUFA10 | 19 | 19 | -3,00745E-16 | 0,999999972 | 1 |
| REEP2 | 4 | 4 | -0,031844844 | 0,383668124 | 1 |
| CRYL1 | 14 | 13 | 1,5324E-15 | 0,999999903 | 1 |
| DYNLRB1 | 4 | 4 | 1,5258E-16 | 0,999999971 | 1 |
| RNF213 | 1 | 1 | 0,015590892 | 0,721029012 | 1 |
| UVRAG | 4 | 4 | 0,07338007 | 0,14475041 | 1 |
| EML4 | 9 | 8 | -0,034280394 | 0,382999515 | 1 |
| STXBP6 | 4 | 4 | -4,10621E-19 | 0,999999999 | 1 |
| CIAO2B | 3 | 3 | 0 | 1 | 1 |
| GFPT1 | 5 | 5 | 0 | 1 | 1 |
| SAR1A | 5 | 5 | 0,025336802 | 0,688397864 | 1 |
| DENND10 | 7 | 7 | 1,6564E-17 | 0,999999989 | 1 |

| | | | | | |
|---|---|---|---|---|---|
| ABI2;ABI1 | 9 | 9 | 0 | 1 | 1 |
| PPFIA3;PPFIA2 | 1 | 1 | 0 | 1 | 1 |
| KIFBP | 12 | 10 | 1,32276E-17 | 0,999999996 | 1 |
| ABI3 | 3 | 3 | -0,067398197 | 0,248505009 | 1 |
| USP32 | 18 | 18 | 9,95273E-15 | 0,999999643 | 1 |
| ARFGEF1 | 14 | 13 | 0 | 1 | 1 |
| GALNT2 | 2 | 2 | 0 | 1 | 1 |
| MAP9 | 1 | 1 | 0 | 1 | 1 |
| MAOB | 20 | 20 | -0,019600426 | 0,60651259 | 1 |
| SETD3 | 3 | 3 | 0 | 1 | 1 |
| PLPP3 | 9 | 9 | 4,61216E-17 | 0,999999993 | 1 |
| MTO1 | 7 | 7 | -1,12163E-16 | 0,99999997 | 1 |
| RAP1GAP;RAP1GAP2 | 2 | 2 | 5,60216E-14 | 0,999999514 | 1 |
| CTSB | 12 | 12 | 9,37832E-16 | 0,999999928 | 1 |
| LRRC4C | 5 | 5 | 1,27771E-16 | 0,999999961 | 1 |
| PLXDC2 | 4 | 4 | 0 | 1 | 1 |
| SHMT2 | 15 | 14 | -3,71666E-18 | 0,999999996 | 1 |
| HSD17B10 | 12 | 12 | 1,55004E-16 | 0,999999978 | 1 |
| RDH11 | 6 | 6 | 2,19549E-17 | 0,999999986 | 1 |
| OLFR332 | 1 | 1 | 0 | 1 | 1 |
| PRKCA | 18 | 18 | 6,10132E-16 | 0,999999928 | 1 |
| EGFR | 7 | 7 | 2,49546E-17 | 0,999999985 | 1 |
| HSD11B1 | 3 | 3 | -1,74279E-16 | 0,99999997 | 1 |
| B4GALNT1 | 2 | 2 | -8,74274E-16 | 0,999999947 | 1 |
| VPS26A | 12 | 12 | 0,023437176 | 0,419692251 | 1 |
| HAP1 | 3 | 3 | -0,051840007 | 0,467435358 | 1 |
| EIF3E | 26 | 20 | 9,52205E-17 | 0,999999983 | 1 |
| HSPA13 | 5 | 5 | 5,26413E-19 | 1 | 1 |
| INPP5A | 12 | 12 | 0 | 1 | 1 |
| MAP7 | 8 | 8 | -5,37888E-18 | 0,999999995 | 1 |
| MRPS27 | 11 | 11 | -0,048594452 | 0,398343244 | 1 |
| PRKCSH | 11 | 10 | -3,63045E-16 | 0,999999927 | 1 |
| TIGAR | 5 | 5 | 0,016557821 | 0,635582181 | 1 |

| | | | | | |
|---|---|---|---|---|---|
| EIF3G | 6 | 5 | -9,71667E-18 | 0,999999991 | 1 |
| PXMP4 | 1 | 1 | -0,082317799 | 0,387816094 | 1 |
| TTC21B | 1 | 1 | -0,099768973 | 0,236793291 | 1 |
| NDUFA11 | 5 | 5 | 1,41309E-17 | 0,999999991 | 1 |
| ALDH3B1 | 6 | 6 | -1,18591E-14 | 0,999999727 | 1 |
| MAGI2 | 24 | 24 | 0 | 1 | 1 |
| CACYBP | 12 | 10 | 0 | 1 | 1 |
| MTX1 | 3 | 3 | 0 | 1 | 1 |
| IMPDH2 | 14 | 12 | 9,86118E-17 | 0,99999997 | 1 |
| CHRM1 | 1 | 1 | 0 | 1 | 1 |
| ATP5ME | 8 | 8 | -1,41257E-17 | 0,999999995 | 1 |
| UBE2E2 | 2 | 1 | 0 | 1 | 1 |
| KCNJ6 | 3 | 3 | -3,2128E-17 | 0,999999996 | 1 |
| COPS7A | 9 | 9 | 5,3649E-12 | 0,999992446 | 1 |
| NECTIN4 | 1 | 1 | -0,076254445 | 0,355340042 | 1 |
| LAMTOR3 | 2 | 2 | 0,08627578 | 0,19906813 | 1 |
| MOB1B;MOB1A | 2 | 2 | 0 | 1 | 1 |
| ABHD17C | 1 | 1 | 0 | 1 | 1 |
| RUFY3 | 4 | 3 | 5,28816E-18 | 0,999999996 | 1 |
| SERPINA1B | 5 | 3 | -0,111004983 | 0,280087903 | 1 |
| PPP2CB | 2 | 2 | 0,084086113 | 0,249790489 | 1 |
| NENF | 2 | 2 | -0,040646213 | 0,394823082 | 1 |
| MAP2K2 | 5 | 4 | 0 | 1 | 1 |
| RAB11FIP2;RAB11FII | 1 | 1 | 2,57574E-19 | 1 | 1 |
| ABHD6 | 17 | 17 | 0 | 1 | 1 |
| CLEC2L | 1 | 1 | 0 | 1 | 1 |
| GGA3 | 8 | 7 | -4,00178E-17 | 0,999999988 | 1 |
| LRRC57 | 12 | 12 | 0 | 1 | 1 |
| ARHGEF7 | 24 | 24 | 0,000120008 | 0,947589012 | 1 |
| DOC2A | 3 | 3 | -6,94874E-19 | 0,999999999 | 1 |
| ELAVL2;ELAVL4 | 3 | 2 | -0,061182305 | 0,381534673 | 1 |
| SYNRG | 17 | 17 | -0,011594978 | 0,4830707 | 1 |
| NIPBL | 1 | 1 | 0 | 1 | 1 |

| | | | | | |
|---|---|---|---|---|---|
| PMM1 | 9 | 9 | 2,89084E-16 | 0,999999951 | 1 |
| CA14 | 3 | 3 | -7,80815E-16 | 0,999999933 | 1 |
| METTL7A1 | 2 | 2 | 0 | 1 | 1 |
| PAFAH1B2 | 5 | 5 | -3,31683E-16 | 0,999999949 | 1 |
| FTH1 | 18 | 16 | -9,74338E-19 | 0,999999998 | 1 |
| FABP5 | 11 | 10 | -4,06656E-16 | 0,999999959 | 1 |
| UBLCP1 | 4 | 3 | 0 | 1 | 1 |
| UFSP2 | 7 | 7 | 0,03889423 | 0,268925059 | 1 |
| ACAP3 | 2 | 2 | 0 | 1 | 1 |
| KCTD17 | 3 | 3 | -4,65341E-19 | 0,999999999 | 1 |
| TFAM | 13 | 13 | -7,76898E-17 | 0,999999982 | 1 |
| SH3GLB1 | 11 | 11 | 0 | 1 | 1 |
| SLC25A20 | 3 | 3 | -2,8837E-17 | 0,999999991 | 1 |
| HSP90B1;HSP90AB1 | 2 | 2 | 0,02188462 | 0,617193032 | 1 |
| CEP131 | 1 | 1 | 0 | 1 | 1 |
| RIMOC1 | 4 | 4 | 0 | 1 | 1 |
| ARR3 | 1 | 1 | 0 | 1 | 1 |
| ARHGEF2 | 17 | 17 | -5,81656E-18 | 0,999999998 | 1 |
| WDR91 | 7 | 7 | -0,010743926 | 0,591418109 | 1 |
| RPS9 | 16 | 13 | 0,13563105 | 0,144134878 | 1 |
| PFDN5 | 7 | 6 | 0 | 1 | 1 |
| PEX5L | 3 | 3 | 0 | 1 | 1 |
| ATP6V1H | 37 | 37 | 0,005887853 | 0,575730467 | 1 |
| OXSR1;STK39 | 1 | 1 | 0 | 1 | 1 |
| UBE2D2 | 2 | 1 | 0 | 1 | 1 |
| PDE6D | 6 | 5 | 9,56902E-19 | 0,999999998 | 1 |
| SCCPDH | 13 | 13 | -1,69289E-16 | 0,999999965 | 1 |
| GNB4 | 5 | 5 | 8,09348E-17 | 0,999999981 | 1 |
| LEMD2 | 1 | 1 | 0 | 1 | 1 |
| RANBP9 | 8 | 8 | 0 | 1 | 1 |
| SKI | 1 | 1 | 0 | 1 | 1 |
| SCAMP2 | 1 | 1 | 0 | 1 | 1 |
| AFDN | 28 | 28 | -0,032090972 | 0,282391096 | 1 |

| | | | | | |
|---|---|---|---|---|---|
| ARL2 | 7 | 7 | 1,25575E-16 | 0,999999961 | 1 |
| DPM3 | 1 | 1 | -0,128698307 | 0,124523442 | 1 |
| KIF5C;KIF5A;KIF5B | 11 | 11 | 1,07927E-16 | 0,999999959 | 1 |
| PPP1R1A | 2 | 2 | 0 | 1 | 1 |
| SYT5 | 8 | 8 | 2,80013E-13 | 0,999998331 | 1 |
| MRPL43 | 2 | 2 | -0,095035975 | 0,184866651 | 1 |
| NDUFB1 | 2 | 2 | 4,33749E-17 | 0,999999994 | 1 |
| STAT1 | 8 | 8 | 0,177240869 | 0,177091375 | 1 |
| MVB12B | 5 | 5 | 0 | 1 | 1 |
| sp\|Q8CCC3\|CL056_I | 1 | 1 | 0 | 1 | 1 |
| APOD | 4 | 4 | 0 | 1 | 1 |
| RWDD4 | 2 | 2 | -1,23111E-16 | 0,999999968 | 1 |
| TEX264 | 2 | 2 | -8,18786E-18 | 0,999999998 | 1 |
| RAB6B | 17 | 16 | 0 | 1 | 1 |
| SLC4A10;SLC4A7;SLC | 1 | 1 | 7,43471E-18 | 0,999999995 | 1 |
| NT5C3B | 8 | 6 | 0 | 1 | 1 |
| GPCPD1 | 16 | 14 | 9,48825E-17 | 0,999999982 | 1 |
| EVI2A | 1 | 1 | -0,032265892 | 0,612846228 | 1 |
| PDE4A | 8 | 8 | 1,43163E-14 | 0,99999953 | 1 |
| MYH13 | 2 | 1 | 0 | 1 | 1 |
| LRRC73 | 2 | 2 | 1,30535E-16 | 0,999999983 | 1 |
| FBXO2 | 8 | 7 | 0,058809933 | 0,160568752 | 1 |
| YIF1B | 2 | 2 | 8,16593E-15 | 0,999999743 | 1 |
| RNF14 | 5 | 5 | -6,51722E-17 | 0,999999977 | 1 |
| MTIF3 | 1 | 1 | 0 | 1 | 1 |
| MYO7A | 2 | 1 | 0 | 1 | 1 |
| ZNF106 | 1 | 1 | -0,064065991 | 0,32530118 | 1 |
| BABAM2 | 6 | 5 | 0,01743253 | 0,558977077 | 1 |
| HDDC2 | 6 | 4 | 0 | 1 | 1 |
| TAMM41 | 4 | 4 | -8,75147E-18 | 0,999999993 | 1 |
| TRAPPC2L | 5 | 5 | 0,036536869 | 0,341599958 | 1 |
| CCZ1 | 7 | 7 | -1,59613E-17 | 0,999999993 | 1 |
| FAM98B | 6 | 3 | -1,46161E-20 | 1 | 1 |

| | | | | | |
|---|---|---|---|---|---|
| REXO1 | 1 | 1 | 0,159051173 | 0,122768806 | 1 |
| CMPK1 | 18 | 18 | -1,90943E-17 | 0,999999987 | 1 |
| ALPK2 | 1 | 1 | 0 | 1 | 1 |
| SYAP1 | 8 | 8 | -2,65749E-18 | 0,999999999 | 1 |
| SAR1B | 8 | 8 | 0,007567214 | 0,826316602 | 1 |
| RPL31 | 5 | 5 | 0,050072109 | 0,545492849 | 1 |
| COX15 | 4 | 4 | 0 | 1 | 1 |
| BPNT2 | 6 | 6 | -0,068726236 | 0,123979179 | 1 |
| H60C | 1 | 1 | -0,033608648 | 0,520128831 | 1 |
| DSG1A;DSG1B | 8 | 2 | 0,348359445 | 0,17362316 | 1 |
| GAS7 | 15 | 15 | -1,65491E-16 | 0,999999965 | 1 |
| RBBP7 | 3 | 1 | 0 | 1 | 1 |
| PHAF1 | 5 | 5 | 1,53209E-15 | 0,99999987 | 1 |
| EIF3K | 4 | 3 | -1,9145E-19 | 1 | 1 |
| CORO2A | 8 | 8 | 1,36713E-17 | 0,999999996 | 1 |
| RAB11A | 1 | 1 | 0 | 1 | 1 |
| PMFBP1 | 1 | 1 | -0,108921396 | 0,21007356 | 1 |
| ITGAM | 10 | 10 | 0,013324263 | 0,760716777 | 1 |
| CTSL | 1 | 1 | 0 | 1 | 1 |
| EPN2 | 9 | 8 | 0 | 1 | 1 |
| PRPS1 | 6 | 6 | 9,14448E-16 | 0,999999907 | 1 |
| NDUFA13 | 20 | 20 | -7,99024E-17 | 0,999999984 | 1 |
| UBE2V1 | 5 | 4 | 0 | 1 | 1 |
| KCTD4 | 6 | 6 | 0,033278247 | 0,462696039 | 1 |
| SKP1 | 14 | 14 | 0,029453994 | 0,268776773 | 1 |
| SLC4A10;SLC4A7 | 1 | 1 | 0 | 1 | 1 |
| CRLF3 | 3 | 3 | 4,29267E-19 | 1 | 1 |
| CLPB | 15 | 15 | -4,43914E-16 | 0,999999943 | 1 |
| ARCN1 | 17 | 17 | -3,51666E-15 | 0,999999832 | 1 |
| CRYZL1 | 7 | 7 | 0,03352219 | 0,227021516 | 1 |
| UBA6 | 24 | 22 | 2,72921E-17 | 0,999999984 | 1 |
| ZFP64 | 1 | 1 | 0,059261495 | 0,478794265 | 1 |
| NDUFS8 | 17 | 17 | -1,16508E-17 | 0,999999997 | 1 |

| | | | | | |
|---|---|---|---|---|---|
| SLITRK2 | 2 | 2 | -5,49321E-19 | 0,999999999 | 1 |
| ZAR1 | 1 | 1 | 0 | 1 | 1 |
| NDUFV3 | 5 | 5 | -1,58801E-16 | 0,999999971 | 1 |
| UFD1 | 7 | 6 | 5,43064E-16 | 0,999999928 | 1 |
| B4GAT1 | 3 | 3 | 3,14843E-18 | 0,999999999 | 1 |
| RBM39 | 2 | 1 | 0 | 1 | 1 |
| RNF25 | 4 | 4 | 0 | 1 | 1 |
| OCIAD1 | 1 | 1 | 0 | 1 | 1 |
| FAM53C | 1 | 1 | -1,63659E-18 | 0,999999998 | 1 |
| GSAP | 1 | 1 | 0 | 1 | 1 |
| DPP6 | 5 | 5 | 0 | 1 | 1 |
| CTU2 | 5 | 5 | 0,000362994 | 0,942015028 | 1 |
| PI4KB | 2 | 2 | 1,72725E-17 | 0,99999999 | 1 |
| MRPS22 | 8 | 8 | -0,001605114 | 0,885898734 | 1 |
| CYB5A | 4 | 4 | 2,1963E-17 | 0,999999982 | 1 |
| SLC7A14 | 13 | 13 | 1,52755E-16 | 0,99999995 | 1 |
| OTUD7B;OTUD7A | 2 | 2 | -3,98982E-16 | 0,999999939 | 1 |
| MTMR12 | 6 | 6 | 1,37239E-17 | 0,999999989 | 1 |
| GBP9 | 1 | 1 | 0 | 1 | 1 |
| HSPBP1 | 6 | 6 | 0,001448855 | 0,879141822 | 1 |
| STX5 | 2 | 2 | 7,25355E-18 | 0,999999997 | 1 |
| LARP1 | 14 | 12 | -2,1052E-17 | 0,999999992 | 1 |
| RNF170 | 3 | 3 | 0,011420153 | 0,657067939 | 1 |
| MEAK7 | 2 | 2 | 1,17114E-17 | 0,999999994 | 1 |
| SLC25A31 | 3 | 1 | 0,011481845 | 0,739518738 | 1 |
| MAP1LC3B | 3 | 3 | -4,54166E-17 | 0,999999992 | 1 |
| FCSK | 7 | 7 | -1,9851E-16 | 0,999999956 | 1 |
| RAB33A | 3 | 3 | 3,29058E-19 | 0,999999999 | 1 |
| RPL17 | 10 | 10 | 0,084526972 | 0,373434431 | 1 |
| NUDT16 | 4 | 4 | 6,38684E-17 | 0,999999981 | 1 |
| TBK1 | 9 | 9 | 0 | 1 | 1 |
| FAM50A | 1 | 1 | 0 | 1 | 1 |
| MTPAP | 6 | 6 | 2,42832E-17 | 0,999999997 | 1 |

| | | | | | |
|---|---|---|---|---|---|
| LIN7B;LIN7C;LIN7A | 3 | 3 | 1,1369E-19 | 1 | 1 |
| DENND1A | 4 | 4 | 3,816E-15 | 0,999999815 | 1 |
| BMPR2 | 6 | 6 | 0,007160651 | 0,693392929 | 1 |
| HERC2 | 2 | 2 | 9,11866E-18 | 0,999999993 | 1 |
| TFCP2 | 3 | 3 | 5,33497E-16 | 0,999999964 | 1 |
| P4HTM | 1 | 1 | 0 | 1 | 1 |
| LMTK2 | 4 | 4 | -1,56076E-15 | 0,999999906 | 1 |
| MAPRE1 | 11 | 11 | 0 | 1 | 1 |
| C2CD6 | 2 | 2 | -2,14872E-16 | 0,999999993 | 1 |
| CSTB | 4 | 4 | 0 | 1 | 1 |
| FNTB | 7 | 7 | 0 | 1 | 1 |
| PIK3R2 | 3 | 3 | -2,8461E-18 | 0,999999996 | 1 |
| CMTM5 | 2 | 2 | -2,5862E-18 | 0,999999997 | 1 |
| IMMT | 6 | 6 | -7,31957E-16 | 0,999999946 | 1 |
| CHRM4 | 1 | 1 | 0 | 1 | 1 |
| CYGB | 9 | 8 | -1,7316E-17 | 0,999999996 | 1 |
| PTPN11 | 32 | 30 | 0 | 1 | 1 |
| MPP1 | 12 | 12 | 6,59324E-16 | 0,999999923 | 1 |
| IL22RA1 | 1 | 1 | 0 | 1 | 1 |
| UNC13A;UNC13B | 9 | 9 | 1,92614E-16 | 0,999999942 | 1 |
| MYDGF | 6 | 6 | 1,05829E-17 | 0,999999992 | 1 |
| GPRASP2 | 4 | 4 | -5,46232E-16 | 0,999999952 | 1 |
| YPEL5 | 2 | 1 | 0,042139362 | 0,652353527 | 1 |
| ABL2 | 9 | 9 | -1,69144E-17 | 0,999999996 | 1 |
| SORBS2 | 1 | 1 | 0 | 1 | 1 |
| CYP2J9 | 4 | 4 | -8,11922E-16 | 0,99999994 | 1 |
| SEPHS1 | 4 | 4 | 4,83462E-16 | 0,999999961 | 1 |
| JMY | 1 | 1 | 0 | 1 | 1 |
| FAM177A1 | 4 | 4 | 1,20186E-15 | 0,999999945 | 1 |
| RNF141 | 2 | 2 | -0,007237581 | 0,772395035 | 1 |
| MAP3K5 | 7 | 7 | 0 | 1 | 1 |
| AQP4 | 6 | 6 | 0,09036999 | 0,143157305 | 1 |
| NR3C1 | 6 | 6 | -2,04081E-17 | 0,999999995 | 1 |

| | | | | | |
|---|---|---|---|---|---|
| CACNG3 | 4 | 4 | -1,01786E-17 | 0,999999994 | 1 |
| NHLRC2 | 13 | 13 | 0,016290099 | 0,551428135 | 1 |
| MYL4 | 2 | 2 | -6,91259E-16 | 0,999999945 | 1 |
| SLC1A2 | 11 | 11 | -8,19144E-18 | 0,999999994 | 1 |
| SH3BGRL2 | 8 | 8 | 1,27593E-20 | 1 | 1 |
| HARS2 | 10 | 10 | 0 | 1 | 1 |
| FNBP1L | 2 | 2 | 0,019034558 | 0,680246462 | 1 |
| ANAPC4 | 5 | 5 | 1,17855E-18 | 0,999999998 | 1 |
| PRNP | 10 | 10 | 7,96265E-17 | 0,999999978 | 1 |
| NECTIN3 | 1 | 1 | 0,057989698 | 0,402368935 | 1 |
| PEAK1 | 6 | 6 | 7,4271E-18 | 0,999999995 | 1 |
| TMEM100 | 1 | 1 | 0 | 1 | 1 |
| SCAF1 | 1 | 1 | 0 | 1 | 1 |
| TSPAN2 | 3 | 3 | -0,076910681 | 0,287581226 | 1 |
| CAMK2A | 16 | 16 | 0,07246957 | 0,132987009 | 1 |
| WWP2 | 2 | 2 | 0 | 1 | 1 |
| SIKE1 | 1 | 1 | 0 | 1 | 1 |
| FGF12 | 2 | 2 | 3,24848E-16 | 0,999999976 | 1 |
| MPI | 19 | 19 | 1,32022E-14 | 0,99999967 | 1 |
| GET3 | 12 | 12 | 0,040113271 | 0,189855219 | 1 |
| WDR4 | 1 | 1 | 0 | 1 | 1 |
| VPS29 | 10 | 9 | 0,022936863 | 0,424088525 | 1 |
| OCIAD1 | 4 | 4 | -0,123923467 | 0,22023508 | 1 |
| TPM1 | 7 | 7 | -8,77064E-18 | 0,999999996 | 1 |
| CLCN4 | 1 | 1 | 0,061664467 | 0,344583071 | 1 |
| KCNA6 | 6 | 6 | 0 | 1 | 1 |
| SDC4 | 3 | 3 | 0 | 1 | 1 |
| EIF3J1;EIF3J2 | 7 | 7 | 0 | 1 | 1 |
| AP3S2 | 5 | 5 | 5,89511E-18 | 0,999999993 | 1 |
| AAMDC | 7 | 5 | 0 | 1 | 1 |
| SEZ6L | 8 | 5 | -4,76837E-13 | 0,999998568 | 1 |
| GMFB | 8 | 8 | 2,88276E-17 | 0,999999987 | 1 |
| UHRF1 | 1 | 1 | 0,08838345 | 0,431176328 | 1 |

| | | | | | |
|---|---|---|---|---|---|
| CDH13 | 10 | 10 | -8,78779E-16 | 0,999999935 | 1 |
| SLC17A6 | 7 | 7 | -0,007019221 | 0,743868456 | 1 |
| PFDN6 | 2 | 2 | -4,87585E-19 | 0,999999999 | 1 |
| ANO10 | 2 | 2 | 0 | 1 | 1 |
| AW551984 | 1 | 1 | 0 | 1 | 1 |
| MMS19 | 2 | 2 | 1,07685E-13 | 0,999999163 | 1 |
| KRT18 | 1 | 1 | 0 | 1 | 1 |
| SLC7A6OS | 2 | 2 | -6,26716E-15 | 0,999999844 | 1 |
| HIGD1A | 1 | 1 | 0 | 1 | 1 |
| sp\|Q9D9H8\|CB069_ | 4 | 4 | 8,01357E-18 | 0,999999998 | 1 |
| LIG1 | 1 | 1 | 0,21919071 | 0,367880262 | 1 |
| sp\|Q9D1K7\|CT027_ | 4 | 4 | 1,65557E-16 | 0,999999965 | 1 |
| EIF4A3 | 3 | 3 | -5,51748E-17 | 0,999999993 | 1 |
| PEX16 | 1 | 1 | 0 | 1 | 1 |
| VEZT | 3 | 3 | -0,06395902 | 0,351383939 | 1 |
| PPFIA1 | 4 | 4 | 0 | 1 | 1 |
| TMEM109 | 2 | 2 | -1,09352E-17 | 0,999999997 | 1 |
| SYPL1 | 2 | 2 | 0 | 1 | 1 |
| KIDINS220 | 9 | 9 | 0 | 1 | 1 |
| SYT7 | 3 | 3 | 0 | 1 | 1 |
| DPM1 | 7 | 7 | -1,67075E-14 | 0,99999963 | 1 |
| EPN2 | 1 | 1 | 0 | 1 | 1 |
| NUDT3 | 9 | 9 | 0 | 1 | 1 |
| TUBB3 | 15 | 15 | -3,09109E-17 | 0,999999986 | 1 |
| TUBB4B;TUBB5;TUB | 6 | 6 | -1,45899E-16 | 0,999999978 | 1 |
| PSMF1 | 4 | 4 | 8,69393E-18 | 0,999999997 | 1 |
| DNM2;DNM3 | 8 | 8 | -6,57039E-18 | 0,999999998 | 1 |
| RAB18 | 7 | 7 | 0 | 1 | 1 |
| HECTD1 | 3 | 2 | -4,00088E-18 | 0,999999996 | 1 |
| CAND1;CAND2 | 3 | 3 | 0 | 1 | 1 |
| GNPAT | 2 | 2 | -1,1548E-17 | 0,999999998 | 1 |
| RHOC;RHOA | 5 | 5 | 0,058160835 | 0,298280372 | 1 |
| PDCD6IP | 1 | 1 | 0 | 1 | 1 |

| | | | | | |
|---|---|---|---|---|---|
| AKR1B1 | 17 | 16 | 2,93969E-17 | 0,999999987 | 1 |
| SDF2 | 2 | 2 | 1,2836E-16 | 0,999999993 | 1 |
| NXN | 2 | 2 | 0 | 1 | 1 |
| PFDN1 | 4 | 4 | 2,03095E-17 | 0,99999999 | 1 |
| TIPRL | 13 | 12 | -3,13825E-17 | 0,99999999 | 1 |
| NUP98 | 1 | 1 | 0 | 1 | 1 |
| DAZL | 1 | 1 | 2,95042E-19 | 0,999999999 | 1 |
| ENTPD3 | 2 | 2 | 0 | 1 | 1 |
| TRIM46 | 7 | 6 | 0 | 1 | 1 |
| MTAP | 7 | 7 | 0 | 1 | 1 |
| TIMM22 | 2 | 2 | -4,60278E-19 | 0,999999999 | 1 |
| MRPL22 | 3 | 3 | 0 | 1 | 1 |
| KARS1 | 26 | 24 | 1,30498E-16 | 0,999999969 | 1 |
| PFDN2 | 8 | 7 | 0 | 1 | 1 |
| STAT2 | 1 | 1 | 0 | 1 | 1 |
| DNAJC3 | 5 | 5 | -1,09542E-12 | 0,999996933 | 1 |
| ATP6AP1 | 7 | 7 | 0,0243261 | 0,493252858 | 1 |
| NDUFS6 | 7 | 7 | -2,10046E-17 | 0,999999995 | 1 |
| ABCF2 | 7 | 6 | 0 | 1 | 1 |
| NDUFA6 | 9 | 9 | 0 | 1 | 1 |
| IKZF3 | 1 | 1 | 0 | 1 | 1 |
| GM26992 | 1 | 1 | -0,016628649 | 0,655992548 | 1 |
| HCLS1 | 3 | 3 | 0 | 1 | 1 |
| SRBD1 | 1 | 1 | 0 | 1 | 1 |
| EEF1A2 | 14 | 14 | 0 | 1 | 1 |
| SH2D3C | 3 | 3 | -0,080964905 | 0,147241252 | 1 |
| SARM1 | 11 | 11 | 0,03373324 | 0,305661027 | 1 |
| AIF1 | 2 | 2 | 0 | 1 | 1 |
| EEF1A1 | 14 | 14 | 4,35961E-15 | 0,999999804 | 1 |
| HMGCL | 9 | 9 | -0,02544238 | 0,429606109 | 1 |
| RAB4A;RAB4B | 2 | 2 | 0,057898496 | 0,159470744 | 1 |
| URB1 | 1 | 1 | 0 | 1 | 1 |
| ETNK1 | 5 | 3 | -6,31661E-17 | 0,999999996 | 1 |

| | | | | | |
|---|---|---|---|---|---|
| MCAM | 4 | 4 | -1,37468E-16 | 0,999999977 | 1 |
| METAP2 | 9 | 8 | 0 | 1 | 1 |
| LACTB | 16 | 16 | -0,029646431 | 0,506641622 | 1 |
| KLHL22 | 6 | 6 | 1,54565E-17 | 0,999999993 | 1 |
| NUDT4 | 4 | 4 | 1,67637E-18 | 0,999999997 | 1 |
| LIN7C | 6 | 5 | -4,4913E-19 | 1 | 1 |
| HNRNPAB | 8 | 7 | 2,46172E-17 | 0,999999993 | 1 |
| GNAI3 | 7 | 7 | 1,1992E-16 | 0,999999973 | 1 |
| PEBP1 | 16 | 16 | -1,23473E-16 | 0,999999983 | 1 |
| RAB31 | 4 | 4 | 0 | 1 | 1 |
| CSNK1G2;CSNK1G3 | 3 | 3 | 1,46265E-19 | 1 | 1 |
| BRK1 | 3 | 3 | 0 | 1 | 1 |
| CNTFR | 4 | 4 | 0 | 1 | 1 |
| MRPS15 | 4 | 4 | -1,33508E-17 | 0,999999992 | 1 |
| SNX4 | 15 | 15 | 1,45883E-15 | 0,999999868 | 1 |
| NDUFB4 | 7 | 7 | 0 | 1 | 1 |
| FAF1 | 9 | 8 | -3,70891E-18 | 0,999999998 | 1 |
| MAGI1 | 7 | 7 | -0,000748103 | 0,912459101 | 1 |
| GRIA4 | 4 | 4 | -0,058215984 | 0,224120358 | 1 |
| NEFL;NEFH;VIM;INA | 1 | 1 | 0 | 1 | 1 |
| KRT5 | 37 | 15 | 0,277972888 | 0,444878497 | 1 |
| SLC25A42 | 8 | 8 | 1,46822E-16 | 0,999999974 | 1 |
| CELSR1 | 1 | 1 | 0 | 1 | 1 |
| ERGIC1 | 5 | 5 | 0 | 1 | 1 |
| DNAJC8 | 3 | 1 | 0 | 1 | 1 |
| UBASH3B | 4 | 4 | 0 | 1 | 1 |
| ATAD1 | 11 | 11 | -0,037196856 | 0,436935196 | 1 |
| CAMK2A;CAMK2B | 1 | 1 | 0 | 1 | 1 |
| TALDO1 | 25 | 18 | 0 | 1 | 1 |
| ELAVL1 | 7 | 5 | 4,3157E-16 | 0,99999996 | 1 |
| SHC2 | 2 | 2 | -1,01399E-15 | 0,999999973 | 1 |
| BPHL | 14 | 14 | -0,034961082 | 0,525409078 | 1 |
| SLC22A4 | 2 | 2 | 5,22931E-14 | 0,999999438 | 1 |

| | | | | | |
|---|---|---|---|---|---|
| THG1L | 2 | 2 | 0,120883368 | 0,142459442 | 1 |
| MLEC | 8 | 8 | 0,00624708 | 0,715296846 | 1 |
| MRPL38 | 3 | 3 | -0,020819077 | 0,618495207 | 1 |
| GPR155 | 2 | 2 | 0,104385894 | 0,230810118 | 1 |
| MYD88 | 1 | 1 | -0,192412953 | 0,132925123 | 1 |
| STON2 | 7 | 7 | 3,52605E-17 | 0,999999984 | 1 |
| WASHC5 | 8 | 7 | -1,44793E-15 | 0,999999872 | 1 |
| ZYX | 7 | 7 | -0,026658733 | 0,48511896 | 1 |
| SNF8 | 4 | 4 | -8,12734E-18 | 0,999999998 | 1 |
| ACADSB | 11 | 11 | -0,023414219 | 0,610147602 | 1 |
| VSNL1;HPCAL4 | 4 | 4 | 1,31098E-16 | 0,999999977 | 1 |
| HSPA14 | 1 | 1 | 0 | 1 | 1 |
| RANBP1 | 5 | 5 | -0,008987363 | 0,668375067 | 1 |
| KRT1;KRT6A;KRT2;KI | 1 | 1 | 0 | 1 | 1 |
| CIAO3 | 4 | 4 | -1,08976E-17 | 0,999999999 | 1 |
| LRRC8A;LRRC8C | 1 | 1 | 0,07866697 | 0,315915363 | 1 |
| GPR20 | 1 | 1 | 0 | 1 | 1 |
| LCMT1 | 12 | 12 | 7,72661E-17 | 0,999999976 | 1 |
| CYP51A1 | 4 | 4 | -0,061295298 | 0,186925536 | 1 |
| GLG1 | 9 | 9 | -0,018548836 | 0,56653478 | 1 |
| TMED7 | 3 | 3 | -3,73915E-18 | 0,999999998 | 1 |
| MSRB2 | 5 | 5 | -0,080069309 | 0,259456429 | 1 |
| RWDD1 | 4 | 4 | 2,29349E-18 | 0,999999997 | 1 |
| TIMMDC1 | 6 | 6 | -3,04411E-17 | 0,999999992 | 1 |
| USP9Y | 11 | 11 | 0,031275311 | 0,218020993 | 1 |
| SPAST | 6 | 6 | 1,63351E-17 | 0,999999986 | 1 |
| KATNAL1 | 6 | 6 | 0,037052119 | 0,272378452 | 1 |
| WNK2;WNK3 | 1 | 1 | 0 | 1 | 1 |
| AP3M1 | 7 | 7 | 0,014404918 | 0,591032623 | 1 |
| MRS2 | 8 | 8 | -4,02516E-19 | 1 | 1 |
| TUBA1C;TUBAL3;TUI | 2 | 2 | 0 | 1 | 1 |
| TENM3 | 10 | 10 | -1,02087E-16 | 0,999999967 | 1 |
| HSPA1L;HSPA2 | 2 | 2 | 0,031400138 | 0,345636695 | 1 |

| | | | | | |
|---|---|---|---|---|---|
| CPNE5;CPNE8 | 3 | 3 | -2,64821E-18 | 0,999999999 | 1 |
| ILK | 2 | 2 | 0,020396189 | 0,584785853 | 1 |
| SCN1A | 15 | 15 | -7,79007E-15 | 0,999999671 | 1 |
| NDUFA8 | 8 | 8 | -4,0932E-23 | 1 | 1 |
| DAP3 | 9 | 9 | -1,47457E-17 | 0,999999992 | 1 |
| PRKAG1;PRKAG2 | 1 | 1 | 0,076100383 | 0,197283943 | 1 |
| RPS15A | 9 | 7 | 0,02851519 | 0,532375962 | 1 |
| NIBAN2 | 3 | 3 | -7,11506E-20 | 1 | 1 |
| KCNC3 | 4 | 4 | 0 | 1 | 1 |
| SMCR8 | 14 | 14 | 0 | 1 | 1 |
| COA6 | 1 | 1 | 0 | 1 | 1 |
| SNX15 | 4 | 4 | 2,65762E-16 | 0,999999966 | 1 |
| MRPS11 | 1 | 1 | 0 | 1 | 1 |
| TIAL1 | 2 | 2 | 0 | 1 | 1 |
| PCBP3 | 6 | 5 | 1,03174E-14 | 0,999999775 | 1 |
| SLC4A10;SLC4A8 | 2 | 2 | 3,0074E-18 | 0,999999997 | 1 |
| OGDH | 8 | 8 | 4,99646E-15 | 0,999999936 | 1 |
| KDSR | 2 | 2 | 0,096114276 | 0,116265782 | 1 |
| ARL3 | 12 | 11 | 0 | 1 | 1 |
| PBDC1 | 4 | 4 | 0,008138134 | 0,706904144 | 1 |
| BOLA2 | 3 | 3 | 2,9522E-16 | 0,999999991 | 1 |
| B3GAT3 | 4 | 4 | 0 | 1 | 1 |
| VPS26B | 12 | 12 | 0 | 1 | 1 |
| RAB5C;RAB5B;RAB5/ | 4 | 4 | 0,079814987 | 0,183907598 | 1 |
| CBFB | 1 | 1 | 0 | 1 | 1 |
| POLDIP2 | 9 | 9 | -1,5339E-16 | 0,999999966 | 1 |
| AGAP1 | 3 | 3 | -0,076988021 | 0,122255821 | 1 |
| 1700109H08RIK | 1 | 1 | 0 | 1 | 1 |
| ATP5PF | 5 | 5 | -1,58192E-16 | 0,999999983 | 1 |
| PTPRF | 6 | 6 | -1,12389E-15 | 0,9999999 | 1 |
| TM9SF4 | 4 | 4 | -4,38565E-16 | 0,999999941 | 1 |
| TRPC4 | 3 | 3 | 0 | 1 | 1 |
| TH | 2 | 2 | 0 | 1 | 1 |

| | | | | | |
|---|---|---|---|---|---|
| AP4B1 | 1 | 1 | 0 | 1 | 1 |
| MAPRE1;MAPRE3 | 2 | 2 | 0 | 1 | 1 |
| SSR4 | 6 | 6 | 0,055105023 | 0,33167877 | 1 |
| PPP2R2B | 1 | 1 | 0 | 1 | 1 |
| PUM2 | 1 | 1 | 0 | 1 | 1 |
| PURB | 14 | 14 | -1,45519E-16 | 0,999999958 | 1 |
| GNAO1 | 6 | 6 | 0,126715423 | 0,135973314 | 1 |
| NCAM1 | 6 | 6 | 2,78901E-18 | 0,999999998 | 1 |
| TOMM20 | 4 | 4 | -0,064604246 | 0,363538304 | 1 |
| DST;MACF1 | 3 | 3 | -3,1332E-19 | 0,999999999 | 1 |
| TMEM43 | 6 | 6 | 9,84006E-17 | 0,999999982 | 1 |
| N6AMT1 | 3 | 2 | 0 | 1 | 1 |
| DDT | 13 | 13 | 6,45462E-17 | 0,999999986 | 1 |
| SLC25A1 | 10 | 10 | 2,3484E-16 | 0,999999955 | 1 |
| LPGAT1 | 4 | 4 | 1,09396E-14 | 0,999999703 | 1 |
| SLC25A40 | 3 | 3 | -1,35461E-15 | 0,999999915 | 1 |
| 1700012B07RIK | 1 | 1 | 4,99179E-21 | 1 | 1 |
| DOCK8 | 2 | 2 | -5,30353E-19 | 0,999999999 | 1 |
| SORBS1 | 5 | 5 | -0,035838917 | 0,375991162 | 1 |
| NDRG2 | 13 | 13 | 1,71696E-17 | 0,999999989 | 1 |
| INPP4A | 4 | 4 | 0,028363242 | 0,502495493 | 1 |
| MDGA2 | 4 | 4 | -0,007694053 | 0,690522658 | 1 |
| PDCL3 | 2 | 1 | -8,11108E-20 | 1 | 1 |
| PCSK1 | 1 | 1 | 0 | 1 | 1 |
| MAPKAP1 | 1 | 1 | -0,062247858 | 0,487279946 | 1 |
| MARS1 | 1 | 1 | 0,011239415 | 0,788203988 | 1 |
| SCN1A;SCN3A;SCN2A | 4 | 4 | -4,82759E-19 | 0,999999999 | 1 |
| RIT2 | 3 | 3 | 0 | 1 | 1 |
| CPNE3 | 5 | 5 | 0,007884051 | 0,741572627 | 1 |
| IST1 | 6 | 6 | -2,18995E-19 | 1 | 1 |
| POFUT1 | 1 | 1 | 0 | 1 | 1 |
| COQ10A | 2 | 2 | 0 | 1 | 1 |
| U2SURP | 1 | 1 | 0 | 1 | 1 |

| | | | | | |
|---|---|---|---|---|---|
| SCFD1 | 5 | 5 | 0 | 1 | 1 |
| PTPA | 14 | 14 | 7,99976E-18 | 0,999999997 | 1 |
| FIG4 | 2 | 2 | 0,019691835 | 0,659849288 | 1 |
| PDE3A | 1 | 1 | 0 | 1 | 1 |
| LIMA1 | 2 | 2 | 0 | 1 | 1 |
| NARS2 | 5 | 5 | 9,91575E-17 | 0,999999986 | 1 |
| ABRACL | 1 | 1 | 2,56156E-15 | 0,999999879 | 1 |
| PDE4B | 9 | 9 | -0,012999093 | 0,5442272 | 1 |
| IGSF9B | 2 | 2 | -1,35065E-16 | 0,999999989 | 1 |
| COMMD7 | 4 | 4 | 0 | 1 | 1 |
| HIF1AN | 1 | 1 | 0,019594195 | 0,657575032 | 1 |
| NDUFAF7 | 9 | 9 | -0,018122144 | 0,65785292 | 1 |
| RPL22L1 | 3 | 3 | 0,04015824 | 0,427993824 | 1 |
| DYNC1I1;DYNC1I2 | 2 | 2 | 1,46949E-19 | 1 | 1 |
| DOCK1 | 7 | 7 | -1,63031E-16 | 0,999999968 | 1 |
| PGK2;PGK1 | 8 | 8 | 7,82213E-17 | 0,999999972 | 1 |
| PTPRA;PTPRE | 1 | 1 | 0 | 1 | 1 |
| WWP1 | 3 | 3 | 0,053176051 | 0,292771498 | 1 |
| TIMM8A2;TIMM8A1 | 1 | 1 | 0 | 1 | 1 |
| GLRX2 | 1 | 1 | -4,33392E-20 | 1 | 1 |
| WDR81 | 4 | 4 | -0,07425695 | 0,221804061 | 1 |
| WDR61 | 6 | 4 | -1,38519E-16 | 0,99999997 | 1 |
| PACS2 | 6 | 6 | -3,56988E-17 | 0,999999986 | 1 |
| NFU1 | 6 | 6 | -4,70968E-18 | 0,999999995 | 1 |
| ARHGEF1 | 3 | 3 | -1,77428E-14 | 0,99999963 | 1 |
| SNAP91;PICALM | 2 | 2 | 1,80275E-18 | 0,999999997 | 1 |
| TBC1D7 | 1 | 1 | 0,04140563 | 0,501541965 | 1 |
| ABHD17A | 2 | 2 | 0 | 1 | 1 |
| RAB27B;RAB27A | 2 | 2 | -4,9792E-18 | 0,999999999 | 1 |
| NDUFA12 | 15 | 15 | -3,58764E-16 | 0,999999949 | 1 |
| MCRIP1 | 1 | 1 | 0 | 1 | 1 |
| DYRK1A | 4 | 4 | 0 | 1 | 1 |
| HMGB1 | 9 | 6 | 0,045460388 | 0,608646567 | 1 |

| | | | | | |
|---|---|---|---|---|---|
| WNK1;WNK2;WNK3 | 2 | 2 | -7,57356E-19 | 0,999999998 | 1 |
| NLRP9C | 1 | 1 | 0 | 1 | 1 |
| GSTM1;GSTM2;GSTN | 3 | 3 | -0,027177409 | 0,446847057 | 1 |
| GID4 | 1 | 1 | 0 | 1 | 1 |
| LRRC40 | 9 | 9 | 8,98743E-16 | 0,999999918 | 1 |
| SH3BP1 | 5 | 5 | 1,89771E-18 | 0,999999998 | 1 |
| CNNM2 | 6 | 6 | 0 | 1 | 1 |
| BAG2 | 2 | 2 | 0 | 1 | 1 |
| COA3 | 2 | 2 | -1,3576E-17 | 0,999999995 | 1 |
| MAGEE1 | 1 | 1 | 0 | 1 | 1 |
| PTPN3 | 3 | 3 | -4,20409E-20 | 1 | 1 |
| EPHB3 | 7 | 7 | -1,18197E-18 | 1 | 1 |
| UCHL3 | 13 | 12 | 1,10397E-15 | 0,999999865 | 1 |
| AMZ2 | 2 | 2 | 3,26289E-18 | 0,999999999 | 1 |
| VIM;KRT6A;KRT76;KI | 1 | 1 | 0 | 1 | 1 |
| KRT72 | 1 | 1 | 0 | 1 | 1 |
| KRT1;KRT77;KRT73 | 1 | 1 | 0 | 1 | 1 |
| RGS7;RGS6 | 4 | 4 | 1,44951E-18 | 0,999999997 | 1 |
| UPF3B | 2 | 2 | 9,15195E-19 | 0,999999999 | 1 |
| RAB8B | 6 | 6 | 0,0222078 | 0,503765229 | 1 |
| PROX1 | 1 | 1 | 0 | 1 | 1 |
| SUMO1 | 2 | 1 | 0 | 1 | 1 |
| SLC4A8 | 5 | 5 | 3,11919E-15 | 0,999999888 | 1 |
| SLC4A7 | 8 | 7 | 0 | 1 | 1 |
| AIMP2 | 6 | 3 | 4,25248E-14 | 0,999999405 | 1 |
| ANK2 | 6 | 6 | -2,15283E-17 | 0,999999989 | 1 |
| TXNRD1 | 20 | 20 | 0 | 1 | 1 |
| SNRPD3 | 3 | 3 | 0,058301833 | 0,461547029 | 1 |
| PTP4A2;PTP4A1 | 3 | 3 | 3,65964E-17 | 0,999999994 | 1 |
| NDST2 | 1 | 1 | 0 | 1 | 1 |
| FADS2 | 3 | 3 | 0 | 1 | 1 |
| NUP133 | 1 | 1 | 0 | 1 | 1 |
| WDR26 | 9 | 9 | 4,85397E-17 | 0,99999997 | 1 |

| | | | | | |
|---|---|---|---|---|---|
| AP1G2;AP1G1 | 2 | 2 | -8,88472E-19 | 0,999999999 | 1 |
| SKIV2L | 1 | 1 | 0 | 1 | 1 |
| CMIP | 3 | 3 | -8,16253E-18 | 0,999999996 | 1 |
| ASCC3 | 1 | 1 | -0,029313377 | 0,687452926 | 1 |
| GGA1 | 11 | 10 | 0 | 1 | 1 |
| GM10717 | 1 | 1 | 0 | 1 | 1 |
| LIN52 | 1 | 1 | 0 | 1 | 1 |
| PPP1CB | 6 | 6 | 3,30974E-17 | 0,999999985 | 1 |
| RGS17 | 1 | 1 | 0 | 1 | 1 |
| VPS35L | 7 | 7 | -0,039512808 | 0,317098123 | 1 |
| EFNB2 | 2 | 2 | 0,055539305 | 0,378883853 | 1 |
| NPC1 | 3 | 3 | 2,40227E-20 | 1 | 1 |
| GSTM7 | 14 | 14 | 2,46048E-18 | 0,999999999 | 1 |
| KCNN2 | 3 | 3 | 0 | 1 | 1 |
| MAN2B1 | 3 | 3 | 0,018073782 | 0,657520048 | 1 |
| HID1 | 10 | 10 | 0,059136581 | 0,270592811 | 1 |
| ABCC9 | 1 | 1 | 0 | 1 | 1 |
| ZFP735 | 1 | 1 | 0 | 1 | 1 |
| ITPR1;ITPR2 | 2 | 2 | 4,85737E-18 | 0,999999997 | 1 |
| NOL7 | 1 | 1 | 0 | 1 | 1 |
| MCMBP | 2 | 1 | 0 | 1 | 1 |
| MRPL19 | 7 | 7 | -9,81369E-16 | 0,999999919 | 1 |
| MRPL16 | 3 | 3 | -2,8322E-17 | 0,999999991 | 1 |
| HKDC1 | 8 | 8 | -2,44322E-18 | 0,999999999 | 1 |
| ACSF3 | 13 | 13 | 0 | 1 | 1 |
| COQ5 | 5 | 5 | -1,07682E-17 | 0,999999994 | 1 |
| PREPL | 18 | 14 | 2,49683E-18 | 0,999999998 | 1 |
| RGS19 | 1 | 1 | 0 | 1 | 1 |
| INPP5F | 10 | 9 | -0,005008308 | 0,752773965 | 1 |
| SSR1 | 3 | 3 | -2,78329E-17 | 0,999999989 | 1 |
| RPL32 | 1 | 1 | 0 | 1 | 1 |
| NDST3;NDST4 | 1 | 1 | 0 | 1 | 1 |
| MARK3 | 12 | 12 | -8,88426E-17 | 0,999999977 | 1 |

| | | | | | |
|---|---|---|---|---|---|
| PLCH2;PLCH1 | 2 | 2 | 0,098611522 | 0,271367324 | 1 |
| CABLES2 | 1 | 1 | 0 | 1 | 1 |
| PPOX | 5 | 5 | 0 | 1 | 1 |
| PRKCE;PRKCH | 1 | 1 | 0,014943866 | 0,694776195 | 1 |
| SLC25A23 | 22 | 22 | -6,99897E-18 | 0,999999996 | 1 |
| MRPS2 | 5 | 5 | 0 | 1 | 1 |
| MADD | 3 | 2 | -5,79441E-15 | 0,999999906 | 1 |
| TATDN1 | 3 | 2 | 0 | 1 | 1 |
| SCN9A | 1 | 1 | 0 | 1 | 1 |
| LRRC20 | 2 | 1 | 0 | 1 | 1 |
| ACYP1 | 5 | 5 | 0 | 1 | 1 |
| SNX7 | 4 | 4 | 1,32834E-18 | 0,999999999 | 1 |
| NRXN3;NRXN1 | 3 | 3 | 0 | 1 | 1 |
| SLC13A5 | 2 | 2 | -0,118424156 | 0,165291609 | 1 |
| PARL | 2 | 2 | 0 | 1 | 1 |
| FGF1 | 2 | 2 | 5,77927E-15 | 0,999999763 | 1 |
| CSNK2B | 5 | 5 | 5,03745E-17 | 0,999999979 | 1 |
| UBR2 | 1 | 1 | 0 | 1 | 1 |
| UBE2Z | 9 | 8 | 0 | 1 | 1 |
| BMPR1A | 2 | 2 | 0 | 1 | 1 |
| D3ERTD751E | 2 | 1 | 0 | 1 | 1 |
| CASK | 1 | 1 | 0 | 1 | 1 |
| C1QC | 4 | 4 | 0,145432064 | 0,259271148 | 1 |
| RAN | 12 | 11 | 1,05792E-17 | 0,999999992 | 1 |
| PPIF | 6 | 6 | -0,026007961 | 0,421730542 | 1 |
| TTN | 7 | 6 | 0,025265439 | 0,427221301 | 1 |
| DNM1;DNM2 | 8 | 8 | 0,026911743 | 0,393419003 | 1 |
| TUBB1;TUBB4B;TUBI | 4 | 4 | 1,08221E-16 | 0,999999988 | 1 |
| NT5E | 2 | 2 | -1,00244E-11 | 0,99999424 | 1 |
| HAPLN2 | 8 | 8 | 7,40004E-18 | 0,999999996 | 1 |
| ANKS1B | 4 | 4 | 0 | 1 | 1 |
| COX6B1 | 7 | 7 | 0,030513974 | 0,374118313 | 1 |
| DHRS4 | 8 | 8 | -0,055280396 | 0,15700514 | 1 |

| | | | | | |
|---|---|---|---|---|---|
| CAMK2G | 3 | 3 | 5,21569E-14 | 0,999999343 | 1 |
| SLC6A15;SLC6A17 | 1 | 1 | 0,054794031 | 0,509198756 | 1 |
| SLC6A6;SLC6A2;SLC6 | 1 | 1 | 0,049155103 | 0,412310082 | 1 |
| PON2 | 5 | 5 | 1,68195E-17 | 0,99999999 | 1 |
| GADD45GIP1 | 2 | 2 | 0 | 1 | 1 |
| ITPK1 | 5 | 5 | -0,065670103 | 0,288233357 | 1 |
| SCN1A;SCN3A | 1 | 1 | 0 | 1 | 1 |
| TXNL1 | 18 | 17 | 4,67271E-17 | 0,999999994 | 1 |
| BLES03 | 9 | 7 | 0 | 1 | 1 |
| AP3B1 | 14 | 14 | 1,75297E-15 | 0,999999829 | 1 |
| SRP14 | 3 | 3 | 0,145173695 | 0,136008175 | 1 |
| HAGHL | 5 | 5 | -3,67048E-19 | 0,999999999 | 1 |
| SBNO1 | 3 | 1 | 0 | 1 | 1 |
| SKT | 3 | 3 | -0,03878589 | 0,551696568 | 1 |
| RPAP1 | 1 | 1 | 0 | 1 | 1 |
| ATP2B4 | 5 | 5 | -2,17784E-17 | 0,99999999 | 1 |
| CELF3 | 1 | 1 | 0 | 1 | 1 |
| GNAS;GNAL | 5 | 5 | 4,57542E-16 | 0,999999946 | 1 |
| MRPL1 | 10 | 10 | -1,37909E-17 | 0,999999996 | 1 |
| COMMD4 | 1 | 1 | 0,057980688 | 0,47737565 | 1 |
| DYNC2I2 | 1 | 1 | 0 | 1 | 1 |
| DGKA | 1 | 1 | 0 | 1 | 1 |
| FHIT | 3 | 3 | -1,71631E-17 | 0,999999991 | 1 |
| SEC14L3 | 1 | 1 | -0,260021416 | 0,186862138 | 1 |
| GRIP1 | 9 | 9 | 0,006149257 | 0,724959581 | 1 |
| CORO1B | 17 | 15 | 0 | 1 | 1 |
| CLCN6 | 8 | 8 | 0,039481468 | 0,178519778 | 1 |
| TPP1 | 4 | 4 | 5,05073E-19 | 0,999999999 | 1 |
| LGALSL | 7 | 7 | 0 | 1 | 1 |
| ZBTB8OS | 1 | 1 | 0 | 1 | 1 |
| NLRX1 | 8 | 8 | 0 | 1 | 1 |
| SRPK1 | 4 | 3 | 0 | 1 | 1 |
| CREBZF | 1 | 1 | 0 | 1 | 1 |

| | | | | | |
|---|---|---|---|---|---|
| PHACTR3 | 2 | 2 | 2,175E-18 | 0,999999999 | 1 |
| PIN4 | 1 | 1 | 0,101634041 | 0,255258925 | 1 |
| GFER | 4 | 4 | -1,19334E-16 | 0,999999973 | 1 |
| ISCA2 | 6 | 6 | -3,4093E-16 | 0,999999956 | 1 |
| GMPR2 | 5 | 5 | 0,029757219 | 0,338734221 | 1 |
| PRPS2 | 13 | 12 | 0,013977057 | 0,65659545 | 1 |
| RPS6KB1 | 3 | 3 | 0 | 1 | 1 |
| ATP5PO | 18 | 18 | -1,32001E-17 | 0,999999993 | 1 |
| PDK1;PDK2 | 1 | 1 | 0 | 1 | 1 |
| RRM2B | 3 | 3 | -9,16133E-17 | 0,999999983 | 1 |
| KPTN | 1 | 1 | 0 | 1 | 1 |
| MARS2 | 8 | 8 | -1,27694E-17 | 0,999999996 | 1 |
| NACC1 | 1 | 1 | -0,130415934 | 0,191530925 | 1 |
| NUDT7 | 1 | 1 | 0 | 1 | 1 |
| SLC39A10 | 9 | 9 | -1,91614E-16 | 0,999999956 | 1 |
| MBP | 3 | 3 | -0,121816593 | 0,238041378 | 1 |
| SEC23A;SEC23B | 3 | 3 | 3,32528E-16 | 0,999999977 | 1 |
| GRIN2C | 1 | 1 | 0 | 1 | 1 |
| PTPN1 | 4 | 4 | -0,013464469 | 0,572481957 | 1 |
| BCCIP | 2 | 2 | 9,0715E-18 | 0,999999998 | 1 |
| CAMK2G | 4 | 4 | -0,025190283 | 0,409059897 | 1 |
| KCNJ13 | 4 | 2 | 0,192895789 | 0,266573887 | 1 |
| PTP4A2 | 3 | 3 | 6,65957E-17 | 0,999999982 | 1 |
| SLC24A4 | 1 | 1 | 0 | 1 | 1 |
| TAS2R41 | 1 | 1 | 0 | 1 | 1 |
| PMPCB;UQCRC1 | 1 | 1 | -4,48599E-18 | 0,999999995 | 1 |
| TTN | 1 | 1 | 0,158749914 | 0,293470171 | 1 |
| NAAA | 4 | 4 | -2,56045E-15 | 0,999999931 | 1 |
| SLC6A2 | 2 | 2 | 0,003821105 | 0,796615483 | 1 |
| ELL | 1 | 1 | 0 | 1 | 1 |
| sp\|Q8BN57\|CC033_ | 2 | 2 | 0 | 1 | 1 |
| SNX3 | 6 | 6 | 0 | 1 | 1 |
| SNX12 | 6 | 6 | 9,69054E-17 | 0,999999975 | 1 |

| | | | | | |
|---|---|---|---|---|---|
| ITCH | 7 | 7 | -3,1017E-14 | 0,999999437 | 1 |
| CDK14 | 5 | 5 | -1,72765E-17 | 0,999999994 | 1 |
| PMM2 | 13 | 13 | 0 | 1 | 1 |
| SLC25A44 | 7 | 7 | -4,07936E-19 | 0,999999999 | 1 |
| CD101 | 2 | 2 | 0 | 1 | 1 |
| COQ4 | 4 | 4 | 0 | 1 | 1 |
| PLXNA1;PLXNA4 | 3 | 3 | 4,04424E-19 | 1 | 1 |
| SESTD1 | 4 | 4 | 0,034878709 | 0,391439605 | 1 |
| TTR | 6 | 2 | -0,756456442 | 0,999991878 | 1 |
| DOCK11 | 6 | 6 | -2,40576E-14 | 0,999999436 | 1 |
| MRPS25 | 4 | 4 | -0,035084792 | 0,383417439 | 1 |
| TXN2 | 2 | 2 | 1,03184E-18 | 0,999999999 | 1 |
| MTMR9 | 5 | 5 | 0,030225467 | 0,351985714 | 1 |
| DDX5 | 6 | 6 | 1,38236E-17 | 0,999999995 | 1 |
| GLS2 | 2 | 2 | -3,00339E-19 | 0,999999999 | 1 |
| UCKL1 | 6 | 6 | 5,78235E-16 | 0,999999946 | 1 |
| HERC1 | 4 | 4 | 0 | 1 | 1 |
| FRS2 | 1 | 1 | -0,103516257 | 0,216971839 | 1 |
| TCF25 | 4 | 4 | 1,60847E-16 | 0,999999974 | 1 |
| ZC3HAV1 | 1 | 1 | 0,065528375 | 0,466097156 | 1 |
| SDHC | 4 | 4 | 4,39403E-19 | 0,999999999 | 1 |
| FAU | 1 | 1 | -0,061192115 | 0,336967068 | 1 |
| KCNJ11 | 2 | 2 | 2,69997E-19 | 0,999999999 | 1 |
| CPEB2 | 3 | 3 | 0,042942581 | 0,438538136 | 1 |
| CPEB3 | 2 | 2 | -1,3991E-15 | 0,999999935 | 1 |
| ECHDC1 | 8 | 8 | 0 | 1 | 1 |
| SLC37A4 | 1 | 1 | 7,50079E-17 | 0,999999983 | 1 |
| ACP2 | 3 | 3 | -1,79383E-15 | 0,999999907 | 1 |
| MAN2A2 | 2 | 2 | -7,04419E-14 | 0,99999931 | 1 |
| GABARAP | 1 | 1 | 0 | 1 | 1 |
| RGS6 | 5 | 5 | 2,89329E-17 | 0,99999999 | 1 |
| PLXNB2;PLXND1;PLX | 1 | 1 | 0 | 1 | 1 |
| PLXNA1;PLXNA2 | 2 | 2 | 5,64766E-20 | 1 | 1 |

| | | | | | |
|---|---|---|---|---|---|
| NT5C1B | 1 | 1 | 0 | 1 | 1 |
| DET1 | 1 | 1 | -0,047266646 | 0,647648938 | 1 |
| RUNDC3A | 7 | 7 | 0,016211551 | 0,495064949 | 1 |
| NAP1L1;NAP1L4 | 2 | 2 | -0,006746703 | 0,724387434 | 1 |
| PELO | 2 | 2 | 0 | 1 | 1 |
| CAMK2A | 9 | 9 | 4,72945E-17 | 0,999999982 | 1 |
| ALDH3B2;ALDH3B1 | 2 | 2 | -1,16605E-16 | 0,999999981 | 1 |
| MMP14 | 1 | 1 | -1,06837E-19 | 1 | 1 |
| UBE2H | 5 | 4 | 0,046601241 | 0,401439672 | 1 |
| SLC6A6 | 1 | 1 | 0,043622658 | 0,397888094 | 1 |
| SLC6A11 | 13 | 13 | 6,02721E-17 | 0,999999978 | 1 |
| MYCBP2 | 13 | 11 | 0 | 1 | 1 |
| CNRIP1 | 8 | 8 | 5,00796E-16 | 0,999999929 | 1 |
| GMDS | 9 | 9 | -1,33419E-17 | 0,999999996 | 1 |
| KCNA3 | 2 | 2 | 0 | 1 | 1 |
| NDUFA4 | 6 | 6 | -1,1939E-16 | 0,999999982 | 1 |
| SEC61B | 3 | 3 | 0 | 1 | 1 |
| WASF2;WASF1 | 1 | 1 | 0 | 1 | 1 |
| NSG2 | 1 | 1 | 0 | 1 | 1 |
| CLDN11 | 3 | 3 | -0,150897744 | 0,191781089 | 1 |
| ANKLE2 | 1 | 1 | 0 | 1 | 1 |
| HK2;HK1 | 3 | 3 | 4,62375E-16 | 0,999999975 | 1 |
| LMCD1 | 3 | 2 | -6,42463E-15 | 0,999999866 | 1 |
| GM3839 | 30 | 29 | 0 | 1 | 1 |
| EGR4 | 1 | 1 | 0,050385978 | 0,576054131 | 1 |
| SLC25A22;SLC25A18 | 3 | 3 | 0 | 1 | 1 |
| MSTO1 | 3 | 3 | -2,23803E-19 | 1 | 1 |
| HTRA1 | 5 | 5 | -0,035533739 | 0,5650605 | 1 |
| CADM1 | 3 | 3 | 8,94255E-17 | 0,999999987 | 1 |
| EIF2B4 | 5 | 5 | -3,44616E-13 | 0,999999081 | 1 |
| PIKFYVE | 8 | 8 | 1,66813E-17 | 0,999999988 | 1 |
| DDX3Y | 5 | 4 | -1,3166E-17 | 0,999999991 | 1 |
| SH2D6 | 1 | 1 | 0 | 1 | 1 |

| | | | | | |
|---|---|---|---|---|---|
| ATP8A1;ATP8B1 | 1 | 1 | 0,072728066 | 0,261208461 | 1 |
| GYS1 | 8 | 8 | -5,28861E-17 | 0,999999983 | 1 |
| ANKRD17 | 2 | 2 | -6,38376E-18 | 0,999999998 | 1 |
| TMEM151A | 1 | 1 | -0,03104534 | 0,635429236 | 1 |
| MBP | 1 | 1 | -0,080025482 | 0,474289812 | 1 |
| SLC12A5;SLC12A7 | 1 | 1 | 0 | 1 | 1 |
| GNL2 | 1 | 1 | 0,154123835 | 0,441673131 | 1 |
| TBC1D5 | 5 | 5 | 1,68309E-16 | 0,999999961 | 1 |
| SHF | 2 | 2 | 0 | 1 | 1 |
| SDSL | 3 | 3 | 0 | 1 | 1 |
| DTYMK | 6 | 6 | 4,26899E-17 | 0,999999987 | 1 |
| PTPRD | 2 | 2 | -0,003255993 | 0,871740255 | 1 |
| GFPT2 | 4 | 4 | 0 | 1 | 1 |
| GYG1 | 4 | 4 | 0 | 1 | 1 |
| RPS6KA3 | 3 | 3 | -5,90006E-16 | 0,999999935 | 1 |
| MAP4K5 | 3 | 3 | -0,053624167 | 0,424128585 | 1 |
| CSNK1E | 4 | 4 | -3,05596E-19 | 0,999999999 | 1 |
| SIRT5 | 11 | 11 | -8,36465E-17 | 0,999999971 | 1 |
| ACOT13 | 7 | 7 | -4,99684E-18 | 0,999999999 | 1 |
| SHC3 | 3 | 3 | -6,20962E-17 | 0,999999992 | 1 |
| LUZP1 | 8 | 8 | 0 | 1 | 1 |
| SIDT2 | 1 | 1 | 0 | 1 | 1 |
| TDRP | 2 | 2 | 0 | 1 | 1 |
| MYO1C | 4 | 4 | 0,125960181 | 0,124805723 | 1 |
| CSNK1D | 2 | 2 | -0,07281965 | 0,195869878 | 1 |
| RAB5B | 6 | 6 | 0,027753793 | 0,42685125 | 1 |
| CRYZL2 | 7 | 7 | -2,90108E-16 | 0,999999958 | 1 |
| UCHL5 | 6 | 6 | 0 | 1 | 1 |
| SEPHS2 | 3 | 3 | 4,87236E-18 | 0,999999996 | 1 |
| TAGLN3;TAGLN2 | 4 | 4 | 6,22822E-16 | 0,999999942 | 1 |
| STUM | 1 | 1 | 0,002268783 | 0,875848783 | 1 |
| GLYCTK | 1 | 1 | 0 | 1 | 1 |
| PTPRS | 1 | 1 | 0 | 1 | 1 |

| | | | | | |
|---|---|---|---|---|---|
| SLC25A4;SLC25A5 | 6 | 6 | -5,16329E-15 | 0,999999861 | 1 |
| MBP | 2 | 2 | -0,056161662 | 0,546648739 | 1 |
| CELF1;CELF2 | 2 | 2 | 0 | 1 | 1 |
| UHRF1BP1 | 1 | 1 | 0 | 1 | 1 |
| EIF1;EIF1B | 3 | 3 | 0,07079766 | 0,308897272 | 1 |
| GABRB2 | 10 | 10 | 0,005874672 | 0,665810955 | 1 |
| TSR2 | 4 | 3 | -7,46324E-18 | 0,999999993 | 1 |
| KCNQ5 | 2 | 2 | -0,054165439 | 0,39226957 | 1 |
| LSM3 | 2 | 2 | 2,01907E-15 | 0,999999909 | 1 |
| PRRT1 | 4 | 4 | 0 | 1 | 1 |
| CAMK4 | 2 | 2 | -9,66331E-19 | 1 | 1 |
| SLC25A25 | 15 | 15 | -3,43338E-16 | 0,999999953 | 1 |
| ARPIN | 4 | 4 | 0 | 1 | 1 |
| FBXO3 | 10 | 10 | 2,2797E-14 | 0,999999531 | 1 |
| GMPR | 11 | 11 | 0 | 1 | 1 |
| MTX3 | 9 | 9 | -5,13663E-17 | 0,999999984 | 1 |
| IL33 | 1 | 1 | 0 | 1 | 1 |
| TBCC | 5 | 5 | -0,011556375 | 0,661294013 | 1 |
| RUFY3 | 19 | 19 | 4,49538E-18 | 0,999999998 | 1 |
| RTN4IP1 | 9 | 9 | -1,51939E-16 | 0,999999977 | 1 |
| CSK | 6 | 6 | 3,94E-16 | 0,999999953 | 1 |
| ADCY3 | 5 | 5 | 0,022231727 | 0,55074583 | 1 |
| ZDHHC15 | 1 | 1 | 0 | 1 | 1 |
| PIGT | 3 | 3 | 0,039797937 | 0,329773806 | 1 |
| AK1 | 17 | 16 | 0 | 1 | 1 |
| ADCY1 | 10 | 10 | 0 | 1 | 1 |
| CLEC16A | 2 | 2 | 0,02637622 | 0,535939235 | 1 |
| SLC9A3R2 | 6 | 6 | -6,37064E-18 | 0,999999998 | 1 |
| SLC22A23 | 6 | 6 | -9,15321E-18 | 0,999999995 | 1 |
| PTGES3 | 6 | 6 | 0,118203612 | 0,200449022 | 1 |
| ANAPC1 | 4 | 4 | 0,026675202 | 0,534309533 | 1 |
| TIMM10 | 4 | 4 | -1,64181E-19 | 1 | 1 |
| GNG3 | 2 | 2 | 0 | 1 | 1 |

| | | | | | |
|---|---|---|---|---|---|
| PARS2 | 8 | 8 | 0 | 1 | 1 |
| COMTD1 | 2 | 2 | -2,53498E-17 | 0,999999993 | 1 |
| RNF150 | 1 | 1 | 0 | 1 | 1 |
| HNRNPC | 6 | 5 | 3,95527E-17 | 0,999999992 | 1 |
| ACOT1;ACOT2 | 10 | 10 | 0 | 1 | 1 |
| SEPTIN4;SEPTIN5 | 4 | 4 | 1,49269E-18 | 0,999999999 | 1 |
| FKBP1B | 3 | 3 | 0 | 1 | 1 |
| EPPK1;PLEC | 3 | 3 | 7,80438E-18 | 0,999999994 | 1 |
| VDAC2 | 14 | 14 | 0 | 1 | 1 |
| HNRNPD;HNRNPDL | 1 | 1 | 0,072366824 | 0,390789314 | 1 |
| MSI2 | 4 | 3 | -1,2412E-18 | 1 | 1 |
| RBFOX1 | 2 | 2 | 3,65322E-16 | 0,999999979 | 1 |
| RBFOX3 | 2 | 2 | -4,10106E-19 | 1 | 1 |
| GBA | 4 | 4 | -5,80377E-17 | 0,999999978 | 1 |
| PREX2 | 6 | 6 | 0 | 1 | 1 |
| PTBP1 | 4 | 2 | 0 | 1 | 1 |
| PPIL3 | 2 | 1 | -0,170123152 | 0,366962926 | 1 |
| INPP5J | 6 | 6 | 1,31611E-17 | 0,999999995 | 1 |
| MMAB | 6 | 6 | -5,68378E-15 | 0,999999804 | 1 |
| CLIC1 | 6 | 5 | 1,51021E-17 | 0,999999994 | 1 |
| ANK3;ANK1;ANK2 | 2 | 2 | -2,57195E-18 | 0,999999998 | 1 |
| NTRK2 | 5 | 5 | 0 | 1 | 1 |
| CCT6A;CCT6B | 5 | 4 | 0 | 1 | 1 |
| GATD3 | 13 | 13 | -1,2969E-16 | 0,999999974 | 1 |
| COX7A2 | 3 | 3 | -8,28837E-17 | 0,999999989 | 1 |
| CKM | 2 | 2 | 8,23594E-18 | 0,999999996 | 1 |
| CUTC | 5 | 2 | -4,66216E-16 | 1 | 1 |
| PHKB | 7 | 7 | -2,81213E-16 | 0,999999949 | 1 |
| ATP4A | 2 | 2 | 0 | 1 | 1 |
| H2-D1 | 4 | 3 | 0,197793418 | 0,290809235 | 1 |
| ACBD5 | 5 | 5 | -1,27517E-17 | 0,99999999 | 1 |
| ADAM23 | 9 | 9 | 0 | 1 | 1 |
| PTER | 3 | 3 | -0,178697768 | 0,173442463 | 1 |

| | | | | | |
|---|---|---|---|---|---|
| UNC119 | 1 | 1 | -0,117861274 | 0,427319155 | 1 |
| MTMR2 | 12 | 12 | 0,027417776 | 0,337195632 | 1 |
| BNIP3L | 2 | 2 | -5,02046E-16 | 0,999999942 | 1 |
| MERTK | 2 | 2 | -9,68089E-18 | 0,999999998 | 1 |
| HSD17B10 | 5 | 5 | -2,67439E-18 | 0,999999999 | 1 |
| PSMD10 | 1 | 1 | 0,198535134 | 0,171406418 | 1 |
| CNOT7 | 3 | 2 | 2,78177E-17 | 0,999999995 | 1 |
| DAG1 | 6 | 6 | -4,35309E-17 | 0,99999998 | 1 |
| CADPS | 1 | 1 | 0,110798912 | 0,119034068 | 1 |
| CNOT11 | 3 | 3 | 0 | 1 | 1 |
| NOVA1 | 4 | 3 | 2,75074E-16 | 0,999999975 | 1 |
| MFF | 11 | 11 | -5,90051E-17 | 0,999999992 | 1 |
| EFNB1 | 2 | 2 | 3,41848E-17 | 0,99999999 | 1 |
| PLPBP | 13 | 12 | 0 | 1 | 1 |
| SYBU | 3 | 3 | -1,11256E-16 | 0,999999985 | 1 |
| PRKAR1B | 9 | 9 | -1,04802E-10 | 0,999960522 | 1 |
| TMEM263 | 2 | 2 | 6,66081E-09 | 0,999794564 | 1 |
| HBEGF | 1 | 1 | -0,064254901 | 0,362662875 | 1 |
| TMEM163 | 5 | 5 | 1,9564E-16 | 0,999999982 | 1 |
| LMAN1 | 2 | 2 | -1,97679E-15 | 0,999999894 | 1 |
| SCRN2 | 5 | 4 | 0,07805663 | 0,317626232 | 1 |
| TUBB4B;TUBB5;TUBI | 2 | 2 | 6,00825E-18 | 0,999999997 | 1 |
| FLRT2 | 4 | 4 | 0,063677023 | 0,167582974 | 1 |
| DGUOK | 3 | 3 | 0 | 1 | 1 |
| CBR1;CBR3 | 1 | 1 | 0 | 1 | 1 |
| NIPA1 | 3 | 3 | -7,51423E-20 | 1 | 1 |
| HECW1 | 4 | 4 | -0,028811708 | 0,588504524 | 1 |
| ETFB | 8 | 8 | -0,060366239 | 0,423859459 | 1 |
| SERAC1 | 5 | 5 | 0 | 1 | 1 |
| PIK3C3 | 10 | 9 | -3,34928E-17 | 0,999999983 | 1 |
| CYTH3;CYTH2;CYTH1 | 1 | 1 | 0 | 1 | 1 |
| SLC25A35 | 4 | 4 | 0,008375072 | 0,731649514 | 1 |
| SVIP | 2 | 2 | -0,224781197 | 0,121202024 | 1 |

| | | | | | |
|---|---|---|---|---|---|
| YME1L1 | 10 | 10 | -0,03035832 | 0,499803177 | 1 |
| CDS1 | 2 | 2 | -0,070632909 | 0,239934981 | 1 |
| CYTH2 | 4 | 4 | 0,004258232 | 0,765925249 | 1 |
| ABCA5 | 3 | 3 | 0 | 1 | 1 |
| NAP1L1 | 10 | 9 | -1,58263E-17 | 0,999999989 | 1 |
| SCYL1 | 2 | 2 | -6,64143E-15 | 0,999999797 | 1 |
| TKTL2 | 1 | 1 | 0 | 1 | 1 |
| CDHR4 | 1 | 1 | 0 | 1 | 1 |
| CRACDL | 2 | 2 | -0,009204577 | 0,80766462 | 1 |
| SFPQ | 2 | 2 | -1,32046E-18 | 0,999999999 | 1 |
| NPEPL1 | 3 | 3 | 7,03019E-20 | 1 | 1 |
| TACO1 | 5 | 5 | -8,03379E-17 | 0,999999986 | 1 |
| AKAP1 | 2 | 2 | 5,95751E-18 | 0,999999996 | 1 |
| IMPDH1 | 3 | 3 | -2,46658E-16 | 0,999999978 | 1 |
| UBFD1 | 8 | 6 | 0 | 1 | 1 |
| GALNT17 | 7 | 7 | -0,068616605 | 0,227480658 | 1 |
| PGRMC1;PGRMC2 | 1 | 1 | 1,46742E-19 | 1 | 1 |
| ARHGAP33 | 3 | 3 | 0 | 1 | 1 |
| EGFLAM | 1 | 1 | 0 | 1 | 1 |
| PLBD2 | 5 | 5 | 0 | 1 | 1 |
| MRPL47 | 2 | 2 | -1,28166E-17 | 0,999999996 | 1 |
| RMND5A | 3 | 3 | 2,73612E-15 | 0,999999868 | 1 |
| GLMN | 3 | 3 | 1,5267E-17 | 0,999999991 | 1 |
| SLC25A33 | 2 | 2 | -0,07502885 | 0,356722479 | 1 |
| SMYD5 | 2 | 2 | 8,15476E-18 | 0,999999997 | 1 |
| DAPK1 | 2 | 2 | 0 | 1 | 1 |
| SNX18 | 5 | 5 | 1,80908E-18 | 0,999999999 | 1 |
| POLR2M | 2 | 2 | -0,028393261 | 0,635720703 | 1 |
| ZDHHC5 | 7 | 7 | -0,006156744 | 0,758100845 | 1 |
| ENOX1 | 1 | 1 | 0,153381217 | 0,355983475 | 1 |
| CORO2B | 19 | 19 | 0,016142254 | 0,437520746 | 1 |
| PLGRKT | 3 | 3 | -0,073138474 | 0,203178582 | 1 |
| RAPGEF6 | 3 | 3 | 0 | 1 | 1 |

| | | | | | |
|---|---|---|---|---|---|
| RAE1 | 3 | 3 | 3,74667E-16 | 0,999999966 | 1 |
| EMC10 | 1 | 1 | 0 | 1 | 1 |
| SPATA5;KATNAL2 | 1 | 1 | 0 | 1 | 1 |
| BORCS5 | 4 | 4 | -3,66812E-16 | 0,99999993 | 1 |
| MPDU1 | 2 | 2 | 6,06956E-20 | 1 | 1 |
| TRUB2 | 1 | 1 | 0 | 1 | 1 |
| ELAVL2 | 7 | 7 | 4,83507E-16 | 0,999999942 | 1 |
| PTPRK | 4 | 4 | 5,72193E-16 | 0,999999951 | 1 |
| CTNND2;PKP4 | 4 | 4 | -9,00271E-18 | 0,999999993 | 1 |
| MMP17 | 2 | 2 | 0 | 1 | 1 |
| MRTFB | 8 | 8 | -0,050250471 | 0,202096942 | 1 |
| CDC34 | 2 | 2 | 1,27917E-14 | 0,999999746 | 1 |
| WNK1;WNK3 | 1 | 1 | -0,10883764 | 0,160546362 | 1 |
| WNK1;WNK2;WNK4; | 1 | 1 | 0 | 1 | 1 |
| KCNA4 | 5 | 5 | 0 | 1 | 1 |
| KCNA10;KCNA1;KCN | 1 | 1 | 0 | 1 | 1 |
| CNNM3 | 8 | 8 | 7,1123E-17 | 0,999999979 | 1 |
| EVI5L | 9 | 9 | 5,60146E-19 | 1 | 1 |
| MVD | 4 | 3 | -8,40812E-16 | 0,999999945 | 1 |
| NDUFB3 | 6 | 6 | -4,45577E-17 | 0,999999991 | 1 |
| PLP1 | 8 | 8 | -0,090023475 | 0,349765617 | 1 |
| ALDH3A1 | 3 | 1 | 0 | 1 | 1 |
| LRRC4 | 2 | 2 | 0 | 1 | 1 |
| SLIT1 | 3 | 3 | 0,042191258 | 0,412981392 | 1 |
| SHANK1 | 1 | 1 | 0 | 1 | 1 |
| PPP3CC | 2 | 2 | 0,081376194 | 0,299938538 | 1 |
| SLC25A36 | 1 | 1 | 0 | 1 | 1 |
| KIF2A;KIF2B | 1 | 1 | 8,02766E-19 | 0,999999999 | 1 |
| RPAP3 | 2 | 2 | 0,054613319 | 0,371072499 | 1 |
| VPS37B | 3 | 3 | -6,49826E-16 | 0,99999994 | 1 |
| SCARB2 | 4 | 4 | 4,06866E-18 | 0,999999998 | 1 |
| BRSK2 | 4 | 4 | 0 | 1 | 1 |
| STK11 | 5 | 5 | 6,07589E-15 | 0,999999767 | 1 |

| | | | | | |
|---|---|---|---|---|---|
| COX5A | 7 | 7 | -5,87254E-17 | 0,999999988 | 1 |
| TEX26 | 1 | 1 | 0 | 1 | 1 |
| FMR1 | 2 | 2 | -0,063595361 | 0,245627321 | 1 |
| SLC36A1 | 1 | 1 | 0,093257089 | 0,345132479 | 1 |
| IL1RAP | 3 | 3 | -0,064226486 | 0,228437372 | 1 |
| SLC25A4;SLC25A5;SL | 4 | 4 | 0 | 1 | 1 |
| MATK | 4 | 4 | 0 | 1 | 1 |
| LSM6 | 4 | 3 | 0 | 1 | 1 |
| ENO1;ENO2 | 1 | 1 | 0 | 1 | 1 |
| EDIL3 | 4 | 4 | -1,96444E-18 | 0,999999999 | 1 |
| FBXO22 | 10 | 7 | 0 | 1 | 1 |
| NLGN4L;NLGN2 | 2 | 2 | 2,73717E-18 | 0,999999997 | 1 |
| GIPC1 | 9 | 9 | 0 | 1 | 1 |
| SYNJ1 | 1 | 1 | 0 | 1 | 1 |
| TNFRSF21 | 6 | 6 | 0 | 1 | 1 |
| DLG2 | 3 | 3 | 1,17981E-19 | 1 | 1 |
| PPIH | 3 | 2 | 0 | 1 | 1 |
| PDLIM5 | 4 | 4 | 3,10338E-17 | 0,999999993 | 1 |
| CPPED1 | 5 | 4 | -0,045749235 | 0,353994871 | 1 |
| KRT77 | 4 | 1 | 0 | 1 | 1 |
| NRGN | 2 | 2 | 0,204110639 | 0,999995776 | 1 |
| PEX6 | 1 | 1 | 0,154439694 | 0,136633941 | 1 |
| BCKDHA | 3 | 3 | 0,103055214 | 0,119721522 | 1 |
| MAGED2 | 1 | 1 | 0 | 1 | 1 |
| LPIN2 | 4 | 3 | 0 | 1 | 1 |
| HCFC1 | 6 | 5 | 1,67605E-18 | 0,999999998 | 1 |
| MTND1 | 7 | 7 | -1,34832E-16 | 0,999999987 | 1 |
| UBE2J1 | 3 | 3 | 0 | 1 | 1 |
| RNF24 | 1 | 1 | 0 | 1 | 1 |
| SYNJ2BP | 6 | 6 | -1,25239E-17 | 0,999999998 | 1 |
| RPRD2 | 1 | 1 | -0,051676141 | 0,523567305 | 1 |
| GRIA1 | 1 | 1 | 0 | 1 | 1 |
| LRFN4 | 4 | 4 | 5,032E-16 | 0,999999951 | 1 |

| | | | | | |
|---|---|---|---|---|---|
| TMED2 | 4 | 4 | 0 | 1 | 1 |
| P2RX7 | 2 | 2 | 1,2218E-11 | 0,99999007 | 1 |
| CFAP298 | 1 | 1 | 0 | 1 | 1 |
| OPCML | 2 | 2 | 3,56249E-19 | 0,999999999 | 1 |
| SEC13 | 3 | 3 | 3,11805E-18 | 0,999999997 | 1 |
| CSTF2 | 1 | 1 | 0,134444613 | 0,334378449 | 1 |
| COMT | 5 | 5 | -3,00298E-18 | 0,999999997 | 1 |
| CNTN6 | 2 | 2 | 1,13258E-19 | 1 | 1 |
| NUDT17 | 2 | 2 | 0 | 1 | 1 |
| GSTT3 | 1 | 1 | 0 | 1 | 1 |
| FKBP1A | 8 | 7 | 8,77401E-18 | 0,999999995 | 1 |
| SDK2 | 9 | 9 | 1,57079E-17 | 0,99999999 | 1 |
| ITPA | 8 | 8 | 5,85944E-16 | 0,999999932 | 1 |
| KRT14;KRT42 | 6 | 4 | 0,256949286 | 0,49249786 | 1 |
| RUNDC3B | 4 | 4 | 0 | 1 | 1 |
| GOLGA5 | 5 | 5 | -1,85638E-15 | 0,99999993 | 1 |
| KRT6A;KRT5 | 3 | 2 | 0,18011895 | 0,482404099 | 1 |
| PDXDC1 | 5 | 3 | 0,016938019 | 0,662659847 | 1 |
| BOLA1 | 3 | 3 | -4,90179E-09 | 0,999787088 | 1 |
| CNTN4 | 5 | 5 | -9,32411E-17 | 0,999999978 | 1 |
| PES1 | 1 | 1 | 0 | 1 | 1 |
| NANS | 8 | 8 | 0 | 1 | 1 |
| FGL1 | 1 | 1 | 0 | 1 | 1 |
| SEMA6C | 1 | 1 | -0,038096003 | 0,468891312 | 1 |
| RRAS2 | 3 | 3 | 0 | 1 | 1 |
| COPS5 | 21 | 20 | 0,010911844 | 0,582833316 | 1 |
| PLPP6 | 2 | 2 | 0,048189919 | 0,46654314 | 1 |
| DCAF11 | 5 | 3 | 0,14656844 | 0,200231355 | 1 |
| NAT14 | 1 | 1 | 0 | 1 | 1 |
| EMC4 | 2 | 2 | 1,20659E-18 | 0,999999998 | 1 |
| PPP1R3D | 2 | 2 | 1,64126E-17 | 0,999999996 | 1 |
| SNTB1 | 2 | 2 | 0,079848124 | 0,168090173 | 1 |
| DCHS2 | 1 | 1 | 0 | 1 | 1 |

| | | | | | |
|---|---|---|---|---|---|
| EFR3A | 15 | 15 | -0,000172339 | 0,93592964 | 1 |
| FERMT2 | 1 | 1 | -0,038660082 | 0,492392882 | 1 |
| IFT22 | 4 | 4 | 0,044153043 | 0,353158657 | 1 |
| PTAR1 | 3 | 2 | 0 | 1 | 1 |
| SRC;FYN | 3 | 3 | 4,30112E-18 | 0,999999997 | 1 |
| S100A13 | 2 | 2 | -3,81699E-17 | 0,999999993 | 1 |
| VPS36 | 9 | 8 | 2,57646E-16 | 0,999999962 | 1 |
| UBE3C | 11 | 11 | 0 | 1 | 1 |
| OPALIN | 2 | 2 | -4,54873E-18 | 0,999999998 | 1 |
| RFT1 | 1 | 1 | 0 | 1 | 1 |
| AHCYL2 | 1 | 1 | 0 | 1 | 1 |
| CBLN4 | 1 | 1 | -0,069222568 | 0,287366589 | 1 |
| TECPR2 | 7 | 7 | 0 | 1 | 1 |
| SGPL1 | 5 | 5 | 0 | 1 | 1 |
| CAMK2G | 6 | 6 | 0 | 1 | 1 |
| MYCBP2 | 4 | 4 | -3,8332E-18 | 0,999999995 | 1 |
| MRPS36 | 4 | 4 | -2,07233E-18 | 0,999999997 | 1 |
| PLA2G7 | 5 | 5 | -0,038415245 | 0,326336679 | 1 |
| SCP2 | 6 | 6 | 3,97847E-17 | 0,999999992 | 1 |
| TPCN1 | 1 | 1 | 0 | 1 | 1 |
| AGFG2 | 5 | 5 | 0,035577299 | 0,522114974 | 1 |
| MAP4K4;MINK1 | 1 | 1 | 0 | 1 | 1 |
| F11R | 2 | 2 | -3,03429E-18 | 0,999999996 | 1 |
| MAP3K15 | 1 | 1 | -0,017590651 | 0,661990569 | 1 |
| HACD2 | 2 | 2 | 2,99476E-17 | 0,999999991 | 1 |
| PISD | 2 | 2 | -0,048741165 | 0,338152051 | 1 |
| GAS8 | 2 | 2 | -0,064955024 | 0,19177468 | 1 |
| TMEM254 | 1 | 1 | 0 | 1 | 1 |
| FKBP2 | 1 | 1 | 0 | 1 | 1 |
| DDX17 | 4 | 3 | 2,60851E-16 | 0,999999987 | 1 |
| SCN1B | 6 | 6 | 0,060486269 | 0,145117919 | 1 |
| GPRASP1 | 1 | 1 | 0 | 1 | 1 |
| EMB | 4 | 3 | 0,05209917 | 0,344081896 | 1 |

| | | | | | |
|---|---|---|---|---|---|
| SFT2D3 | 1 | 1 | -0,118322748 | 0,201961386 | 1 |
| ELP3 | 5 | 5 | 1,24067E-15 | 0,999999903 | 1 |
| DNAAF10 | 1 | 1 | 0 | 1 | 1 |
| SLC30A6 | 1 | 1 | -1,32627E-17 | 0,999999992 | 1 |
| MARK3;MARK2 | 1 | 1 | 0 | 1 | 1 |
| ABI2 | 1 | 1 | 0 | 1 | 1 |
| SHISA7 | 13 | 13 | 0 | 1 | 1 |
| MELK | 2 | 2 | 0 | 1 | 1 |
| FUBP1 | 3 | 1 | 0 | 1 | 1 |
| KIF13A | 1 | 1 | 0 | 1 | 1 |
| ACVR2A | 1 | 1 | 0 | 1 | 1 |
| GAD2 | 11 | 10 | 0,004793591 | 0,794521712 | 1 |
| NCOA7 | 8 | 8 | -1,92688E-19 | 1 | 1 |
| PHF24 | 4 | 4 | 0 | 1 | 1 |
| EEF1AKMT2 | 3 | 3 | 2,65836E-18 | 0,999999998 | 1 |
| TMEM178B | 4 | 4 | 1,19385E-16 | 0,99999997 | 1 |
| KIF3C | 1 | 1 | -0,009285878 | 0,78273732 | 1 |
| PGS1 | 8 | 8 | -4,32566E-06 | 0,992668239 | 1 |
| PDAP1 | 9 | 9 | 0,01550756 | 0,543762744 | 1 |
| MADD | 1 | 1 | 0 | 1 | 1 |
| UBE2V1;UBE2V2 | 8 | 8 | -1,94978E-16 | 0,999999963 | 1 |
| FAHD1 | 9 | 9 | -0,081691079 | 0,180566778 | 1 |
| PLLP | 2 | 2 | -2,70963E-15 | 0,99999989 | 1 |
| TNFRSF14 | 1 | 1 | 0,004890185 | 0,836369362 | 1 |
| PLEKHA5 | 4 | 4 | -0,051154202 | 0,276457779 | 1 |
| PTPMT1 | 7 | 7 | 0 | 1 | 1 |
| FER1L4 | 1 | 1 | 0 | 1 | 1 |
| ANAPC2 | 2 | 2 | 0 | 1 | 1 |
| TPD52L2 | 9 | 9 | 1,67236E-17 | 0,999999996 | 1 |
| NUDT8 | 1 | 1 | 0 | 1 | 1 |
| SBF1 | 3 | 2 | 2,17751E-17 | 0,999999994 | 1 |
| FABP1 | 1 | 1 | 0 | 1 | 1 |
| PPP1CA | 7 | 7 | 2,33911E-17 | 0,999999982 | 1 |

| | | | | | |
|---|---|---|---|---|---|
| MGRN1 | 3 | 3 | 0,03001378 | 0,532201359 | 1 |
| PCBP2 | 5 | 5 | 0,020764945 | 0,567588157 | 1 |
| MRPL15 | 4 | 4 | 0 | 1 | 1 |
| SERPINH1 | 2 | 2 | 0 | 1 | 1 |
| CCP110 | 2 | 2 | 0 | 1 | 1 |
| AP1G2 | 1 | 1 | 0 | 1 | 1 |
| CAB39L | 5 | 5 | 0,074117422 | 0,166357347 | 1 |
| TOR1A | 2 | 2 | 2,01174E-11 | 0,9999933 | 1 |
| PUF60 | 8 | 8 | 0 | 1 | 1 |
| TIA1 | 1 | 1 | -0,044553687 | 0,657427558 | 1 |
| CALM3 | 1 | 1 | 0 | 1 | 1 |
| APOO | 7 | 7 | -1,10988E-16 | 0,999999988 | 1 |
| SEMA4B | 4 | 4 | -0,017551056 | 0,57993045 | 1 |
| SLC5A7 | 5 | 5 | 6,01425E-17 | 0,999999976 | 1 |
| AHSG | 5 | 1 | -0,162787095 | 0,260273845 | 1 |
| NUMB | 5 | 5 | 5,13421E-16 | 0,99999994 | 1 |
| CNTNAP5A | 3 | 3 | -0,135034234 | 0,155879455 | 1 |
| CNTNAP5C | 1 | 1 | 0 | 1 | 1 |
| PRXL2B | 6 | 6 | 0,022990849 | 0,363540052 | 1 |
| NEK9 | 5 | 5 | 4,0078E-13 | 0,999998043 | 1 |
| DLG3;DLG1 | 2 | 2 | 0,09774682 | 0,117108138 | 1 |
| SYT2 | 7 | 7 | -0,023370926 | 0,530839325 | 1 |
| ASPHD2 | 4 | 4 | 3,34634E-17 | 0,999999993 | 1 |
| TSPAN7 | 3 | 3 | 9,50572E-16 | 0,999999918 | 1 |
| MRPS17 | 2 | 2 | -0,074056876 | 0,311629997 | 1 |
| MAPK14 | 2 | 2 | 8,32562E-17 | 0,999999988 | 1 |
| CAMKMT | 1 | 1 | 0 | 1 | 1 |
| WASF1 | 13 | 13 | 1,54351E-16 | 0,999999958 | 1 |
| CUSTOS | 2 | 2 | 0,201660605 | 0,177616309 | 1 |
| EIF5B | 3 | 1 | 5,74402E-20 | 1 | 1 |
| CUZD1 | 1 | 1 | -0,230668758 | 0,16328179 | 1 |
| NDUFS5 | 8 | 8 | -8,18355E-19 | 1 | 1 |
| MKNK2 | 1 | 1 | 0 | 1 | 1 |

| | | | | | |
|---|---|---|---|---|---|
| SELENOI | 1 | 1 | -8,31947E-19 | 0,999999998 | 1 |
| CD38 | 3 | 3 | 0,062221778 | 0,188344639 | 1 |
| UBL4A | 7 | 7 | 0 | 1 | 1 |
| PRL8A1 | 1 | 1 | 0 | 1 | 1 |
| BCKDHA | 10 | 10 | -0,046962641 | 0,380686442 | 1 |
| CYB5B | 8 | 8 | 0 | 1 | 1 |
| PABIR1 | 3 | 2 | 0,179991802 | 0,131363463 | 1 |
| EIF6 | 3 | 3 | 0 | 1 | 1 |
| MCUB | 3 | 3 | 0 | 1 | 1 |
| NRXN2 | 1 | 1 | 0 | 1 | 1 |
| RAC2 | 5 | 5 | 1,77188E-16 | 0,999999973 | 1 |
| AKR1B10 | 4 | 4 | -0,076590066 | 0,179252891 | 1 |
| PRMT9 | 1 | 1 | 0,09892522 | 0,496120147 | 1 |
| EXOC5 | 13 | 13 | 1,23611E-16 | 0,999999951 | 1 |
| SYNJ1 | 3 | 3 | 6,18704E-20 | 1 | 1 |
| BID | 1 | 1 | 0 | 1 | 1 |
| CACNA1C;CACNA1S | 1 | 1 | 0,015983749 | 0,657308618 | 1 |
| SCLT1 | 1 | 1 | 0 | 1 | 1 |
| FUT8 | 1 | 1 | 0 | 1 | 1 |
| CWF19L2 | 1 | 1 | 4,06293E-18 | 0,999999996 | 1 |
| CCDC68 | 1 | 1 | 0 | 1 | 1 |
| VAMP4 | 2 | 2 | -1,51123E-15 | 0,999999888 | 1 |
| SLC7A9 | 1 | 1 | -5,66987E-19 | 0,999999999 | 1 |
| SV2C | 2 | 1 | -4,50617E-20 | 1 | 1 |
| VIM;INA | 1 | 1 | 0 | 1 | 1 |
| OSBPL10 | 6 | 6 | 0,036893135 | 0,356184652 | 1 |
| sp\|Q8K1L6\|CP074_I | 1 | 1 | 0 | 1 | 1 |
| VCPKMT | 1 | 1 | 0 | 1 | 1 |
| NUDT5 | 4 | 4 | 6,43072E-17 | 0,999999975 | 1 |
| CLCN7 | 2 | 2 | 0 | 1 | 1 |
| DLG2 | 8 | 8 | -9,76135E-19 | 1 | 1 |
| CISD2 | 3 | 3 | -9,6523E-16 | 0,999999927 | 1 |
| CAB39;CAB39L | 4 | 4 | 0 | 1 | 1 |

| | | | | | |
|---|---|---|---|---|---|
| GSTM2 | 6 | 3 | 0,092060079 | 0,967947867 | 1 |
| OLFR1047 | 1 | 1 | 0 | 1 | 1 |
| SETD2 | 1 | 1 | 0 | 1 | 1 |
| PTK7 | 3 | 3 | 0 | 1 | 1 |
| PDS5B | 1 | 1 | 0 | 1 | 1 |
| ZCCHC2 | 1 | 1 | 0 | 1 | 1 |
| AKT1;AKT2 | 1 | 1 | -0,04729236 | 0,465595072 | 1 |
| KCNC4 | 2 | 2 | 1,82448E-09 | 0,999882402 | 1 |
| TBCE | 8 | 7 | 2,13639E-15 | 0,999999914 | 1 |
| CAMK2G;CAMK2B | 5 | 5 | 1,35831E-17 | 0,999999989 | 1 |
| ATG16L1 | 6 | 6 | 0 | 1 | 1 |
| SYVN1 | 2 | 2 | 0 | 1 | 1 |
| MOBP | 2 | 2 | -0,042170024 | 0,659645073 | 1 |
| KCNIP4 | 2 | 2 | 3,99588E-17 | 0,999999992 | 1 |
| PLCB3 | 6 | 6 | -2,69174E-16 | 0,999999949 | 1 |
| STAM2;STAM | 1 | 1 | 0 | 1 | 1 |
| HNRNPF | 4 | 3 | 9,65019E-16 | 0,999999944 | 1 |
| AGPAT1 | 4 | 3 | 3,14597E-16 | 0,999999946 | 1 |
| MRPS28 | 2 | 2 | 1,70585E-16 | 0,999999983 | 1 |
| TRP53I11 | 4 | 4 | 0 | 1 | 1 |
| ULK2 | 2 | 2 | -5,70277E-16 | 0,999999953 | 1 |
| TNFRSF11B | 1 | 1 | 0,112363421 | 0,278725009 | 1 |
| PCDHAC2 | 3 | 3 | 0 | 1 | 1 |
| ARHGAP12 | 1 | 1 | 0 | 1 | 1 |
| VMN2R98;VMN2R1( | 1 | 1 | 0 | 1 | 1 |
| PLP2 | 1 | 1 | 0 | 1 | 1 |
| ARHGEF26 | 3 | 3 | 7,07116E-16 | 0,999999923 | 1 |
| XIRP2 | 1 | 1 | 0 | 1 | 1 |
| M6PR | 4 | 4 | 5,40853E-18 | 0,999999995 | 1 |
| TRPM1 | 1 | 1 | 0 | 1 | 1 |
| HELZ2 | 2 | 2 | 0 | 1 | 1 |
| DCAF6 | 1 | 1 | 0 | 1 | 1 |
| TMEM63C | 4 | 4 | 0,073952172 | 0,191817108 | 1 |

| | | | | | |
|---|---|---|---|---|---|
| AMPD3 | 6 | 6 | 0 | 1 | 1 |
| LRFN3 | 4 | 4 | 0 | 1 | 1 |
| PIGK | 4 | 4 | -7,23821E-18 | 0,999999994 | 1 |
| SCAPER | 2 | 1 | -0,151150042 | 0,345333313 | 1 |
| CTSO | 1 | 1 | 0 | 1 | 1 |
| FOXRED2 | 1 | 1 | 0 | 1 | 1 |
| KLHL14 | 1 | 1 | 0 | 1 | 1 |
| LAMTOR1 | 5 | 5 | 0,056005317 | 0,199257349 | 1 |
| KIF14 | 1 | 1 | 0 | 1 | 1 |
| DNAH1 | 1 | 1 | 0 | 1 | 1 |
| ELMO1 | 10 | 10 | -3,83588E-18 | 0,999999996 | 1 |
| PTDSS2 | 2 | 2 | 1,3867E-17 | 0,999999991 | 1 |
| DCTN3 | 7 | 7 | -3,18148E-43 | 1 | 1 |
| FAM53B | 1 | 1 | 0 | 1 | 1 |
| SAMD9L | 1 | 1 | 0,113911137 | 0,2501802 | 1 |
| SLC25A24 | 5 | 5 | -1,51769E-13 | 0,999998915 | 1 |
| AGTPBP1 | 8 | 8 | 0 | 1 | 1 |
| NF2 | 2 | 2 | -7,99509E-20 | 1 | 1 |
| IMPACT | 16 | 16 | 5,11534E-18 | 0,999999998 | 1 |
| MFN1 | 3 | 3 | 1,21073E-16 | 0,999999972 | 1 |
| COMMD2 | 4 | 4 | 0 | 1 | 1 |
| TOP3B | 1 | 1 | 0 | 1 | 1 |
| KBTBD2 | 3 | 3 | 0 | 1 | 1 |
| CD81 | 5 | 5 | 5,94269E-18 | 0,999999999 | 1 |
| EIF3L | 23 | 21 | 0 | 1 | 1 |
| MYO1E | 1 | 1 | 0,043927997 | 0,491919571 | 1 |
| PCYT1A | 5 | 5 | 0,055647906 | 0,275433408 | 1 |
| KAZN | 2 | 2 | -2,64219E-17 | 0,999999991 | 1 |
| PBSN | 1 | 1 | 0 | 1 | 1 |
| RBSN | 1 | 1 | -0,227226313 | 0,202446695 | 1 |
| FBXL20 | 1 | 1 | 0 | 1 | 1 |
| CACNA2D1;CACNA2I | 1 | 1 | 0 | 1 | 1 |
| CLCN2 | 2 | 2 | -7,29627E-15 | 0,99999982 | 1 |

| | | | | | |
|---|---|---|---|---|---|
| TRIP12 | 6 | 5 | 3,58666E-15 | 0,999999865 | 1 |
| ACP1 | 1 | 1 | 0 | 1 | 1 |
| NRXN3 | 12 | 12 | 0,011214717 | 0,590016422 | 1 |
| GNAL | 4 | 4 | -4,23735E-18 | 0,999999999 | 1 |
| CCDC90B | 3 | 3 | 0 | 1 | 1 |
| FAM185A | 2 | 2 | -8,19877E-17 | 0,999999978 | 1 |
| AKTIP | 2 | 2 | -7,4902E-18 | 0,999999995 | 1 |
| SIDT1 | 4 | 4 | 1,37584E-16 | 0,999999964 | 1 |
| FAM98A | 2 | 2 | -0,056992274 | 0,233067354 | 1 |
| sp\|Q8BHB7\|CP046_ | 1 | 1 | 0 | 1 | 1 |
| ZKSCAN8 | 1 | 1 | -5,35712E-15 | 0,999999854 | 1 |
| LIFR | 1 | 1 | 0 | 1 | 1 |
| BECN1 | 3 | 3 | 2,89305E-11 | 0,999987939 | 1 |
| TAOK3 | 3 | 3 | 0 | 1 | 1 |
| NUP85 | 1 | 1 | 0 | 1 | 1 |
| SNAP25 | 8 | 8 | -1,15704E-17 | 0,999999992 | 1 |
| RPL38 | 4 | 3 | 8,46236E-16 | 0,999999937 | 1 |
| EPHB1 | 8 | 8 | 0 | 1 | 1 |
| SELENOT | 3 | 3 | 1,32901E-16 | 0,999999965 | 1 |
| TMEM63B | 4 | 4 | 1,23288E-18 | 0,999999998 | 1 |
| INO80 | 1 | 1 | 0,101612805 | 0,299754903 | 1 |
| ANKRD13D | 4 | 4 | 3,43364E-18 | 0,999999998 | 1 |
| GAREM1 | 3 | 3 | -0,022734109 | 0,527127788 | 1 |
| RPL37A | 3 | 3 | 0,104592873 | 0,207978473 | 1 |
| NDUFAF3 | 5 | 5 | -6,30375E-17 | 0,999999984 | 1 |
| GABRB1;GABRB2 | 5 | 5 | 0,033669443 | 0,214356495 | 1 |
| TGFB2 | 1 | 1 | 0 | 1 | 1 |
| CRYBG3 | 2 | 2 | 2,32062E-17 | 0,999999994 | 1 |
| TSC22D1;TSC22D4 | 1 | 1 | -0,09814932 | 0,397099154 | 1 |
| ULK1 | 2 | 2 | 0 | 1 | 1 |
| COG6 | 2 | 1 | 0 | 1 | 1 |
| POMT2 | 1 | 1 | 0 | 1 | 1 |
| NLRP4A | 1 | 1 | 0 | 1 | 1 |

| | | | | | |
|---|---|---|---|---|---|
| FOCAD | 1 | 1 | 0 | 1 | 1 |
| WDR45B | 4 | 4 | 4,5651E-18 | 0,999999997 | 1 |
| CSTF3 | 1 | 1 | 0 | 1 | 1 |
| PCK2;PCK1 | 1 | 1 | 0,06120552 | 0,425555758 | 1 |
| MAP3K15;MAP3K5 | 2 | 2 | -2,17033E-17 | 0,999999991 | 1 |
| FGFR1OP2 | 3 | 3 | 1,35351E-05 | 0,987150046 | 1 |
| NRBP2 | 9 | 9 | 0,016711728 | 0,436744783 | 1 |
| NAT8L | 3 | 3 | -0,01850211 | 0,643425626 | 1 |
| SLC14A1 | 3 | 3 | 3,55442E-19 | 0,999999999 | 1 |
| PPP2R3D | 1 | 1 | 0 | 1 | 1 |
| NOS1AP | 2 | 2 | 0 | 1 | 1 |
| BAX | 8 | 7 | 3,05946E-16 | 0,999999969 | 1 |
| NTRK2;NTRK1;NTRK3 | 2 | 2 | 0 | 1 | 1 |
| INSR;INSRR | 1 | 1 | -0,070581496 | 0,294120565 | 1 |
| KCNJ16 | 1 | 1 | -0,18152877 | 0,160077332 | 1 |
| WNK1;WNK2;WNK4 | 2 | 2 | -6,98531E-17 | 0,999999984 | 1 |
| TUSC3 | 3 | 3 | 1,58507E-16 | 0,999999979 | 1 |
| ITGB2 | 15 | 15 | 0,056979866 | 0,492351017 | 1 |
| STN1 | 2 | 2 | 0 | 1 | 1 |
| ADCY8 | 5 | 5 | -2,39676E-16 | 0,999999963 | 1 |
| SLC25A14 | 2 | 2 | -5,78735E-18 | 0,999999999 | 1 |
| ABCC1 | 1 | 1 | 0 | 1 | 1 |
| RNF11 | 2 | 2 | 0,073476107 | 0,245982004 | 1 |
| UTRN | 1 | 1 | 0 | 1 | 1 |
| DNAJB14 | 2 | 2 | 0 | 1 | 1 |
| NGLY1 | 4 | 4 | 2,55136E-17 | 0,999999994 | 1 |
| FGFR1 | 2 | 2 | 0 | 1 | 1 |
| GRM4 | 2 | 2 | 0 | 1 | 1 |
| CTBP2 | 3 | 3 | 0 | 1 | 1 |
| SPRYD4 | 7 | 7 | -3,98885E-17 | 0,999999989 | 1 |
| SLC18A3 | 2 | 2 | 0 | 1 | 1 |
| KIF2B | 1 | 1 | 0 | 1 | 1 |
| CHD4 | 1 | 1 | 0 | 1 | 1 |

| | | | | | |
|---|---|---|---|---|---|
| PNKD | 1 | 1 | 0,007887893 | 0,764603991 | 1 |
| PPP2R3C | 1 | 1 | 0 | 1 | 1 |
| CCDC92 | 6 | 6 | 4,75747E-16 | 0,999999933 | 1 |
| SLC35G2 | 4 | 4 | 0 | 1 | 1 |
| PSMD4 | 8 | 7 | 0 | 1 | 1 |
| MAGOH;MAGOHB | 1 | 1 | 0 | 1 | 1 |
| MYLK3;MYLK4;MYLK | 1 | 1 | 0 | 1 | 1 |
| UQCC1 | 6 | 6 | -5,03787E-16 | 0,999999946 | 1 |
| HCK | 1 | 1 | 0 | 1 | 1 |
| APBA2 | 7 | 7 | 4,16525E-17 | 0,999999981 | 1 |
| XPNPEP1 | 1 | 1 | -0,255752508 | 0,165845569 | 1 |
| MUC19 | 1 | 1 | 0 | 1 | 1 |
| SEC14L1 | 2 | 2 | 1,2638E-17 | 0,999999997 | 1 |
| CAVIN1 | 1 | 1 | 0 | 1 | 1 |
| GNA14 | 1 | 1 | 4,50852E-19 | 0,999999999 | 1 |
| LOXHD1 | 1 | 1 | 0 | 1 | 1 |
| RASGRP1 | 4 | 4 | 0 | 1 | 1 |
| EHD4;EHD3;EHD1 | 4 | 4 | 0,024907071 | 0,512075141 | 1 |
| MYO9A | 1 | 1 | 2,62602E-17 | 0,999999987 | 1 |
| CNOT10 | 2 | 2 | -5,9282E-19 | 0,999999999 | 1 |
| SH3BP4 | 1 | 1 | 0 | 1 | 1 |
| RPA3 | 1 | 1 | 0,195642952 | 0,41831301 | 1 |
| HSPA1L;HSPA2;HSPA | 2 | 2 | 0,013040403 | 0,569277366 | 1 |
| IPO8 | 2 | 2 | 0 | 1 | 1 |
| PIP4P1 | 2 | 2 | 0 | 1 | 1 |
| PIP4P2 | 2 | 2 | 0 | 1 | 1 |
| IGSF10 | 2 | 2 | 0 | 1 | 1 |
| LIN7A | 7 | 7 | 0,025174269 | 0,506993929 | 1 |
| RBM12B1 | 1 | 1 | 0 | 1 | 1 |
| STX8 | 3 | 3 | -2,66053E-17 | 0,999999986 | 1 |
| APBA3 | 1 | 1 | 0 | 1 | 1 |
| COQ7 | 6 | 6 | -7,50952E-19 | 1 | 1 |
| PLCL1;PLCL2 | 1 | 1 | 0 | 1 | 1 |

| | | | | | |
|---|---|---|---|---|---|
| RANBP6 | 2 | 2 | -1,68467E-16 | 0,999999966 | 1 |
| DYM | 2 | 2 | 0,061388994 | 0,485353387 | 1 |
| SPG21 | 2 | 2 | -1,15449E-18 | 0,999999998 | 1 |
| RAB33B;RAB33A | 1 | 1 | 0 | 1 | 1 |
| MRPL21 | 4 | 4 | -2,08826E-18 | 0,999999997 | 1 |
| CPQ | 1 | 1 | 0 | 1 | 1 |
| TAF6 | 1 | 1 | 0 | 1 | 1 |
| CWC25 | 1 | 1 | 0 | 1 | 1 |
| NAB1 | 1 | 1 | 0,166176711 | 0,314251908 | 1 |
| MRPL14 | 4 | 4 | 0 | 1 | 1 |
| RAB8A;RAB10;RAB8I | 1 | 1 | -0,03040001 | 0,667564861 | 1 |
| sp\|Q8K207\|CA021_I | 1 | 1 | 0 | 1 | 1 |
| TOP1 | 1 | 1 | 0 | 1 | 1 |
| ERAP1 | 1 | 1 | 0 | 1 | 1 |
| RICTOR | 1 | 1 | 0 | 1 | 1 |
| CHMP7 | 7 | 7 | 0 | 1 | 1 |
| GDAP1 | 17 | 17 | -3,53914E-16 | 0,999999966 | 1 |
| CHMP6 | 2 | 2 | 0 | 1 | 1 |
| B3GALT9 | 2 | 2 | 3,95897E-18 | 0,999999996 | 1 |
| RPLP2 | 4 | 4 | 0,035513805 | 0,32965491 | 1 |
| NAA15 | 12 | 9 | 0 | 1 | 1 |
| NSL1 | 1 | 1 | 0 | 1 | 1 |
| GSDME | 8 | 7 | -7,57666E-18 | 0,999999998 | 1 |
| CHMP3 | 3 | 3 | 0 | 1 | 1 |
| MAPK8 | 5 | 5 | 0 | 1 | 1 |
| UBE3B | 3 | 3 | 0 | 1 | 1 |
| CAPNS1 | 9 | 9 | 0,017658525 | 0,571056645 | 1 |
| DNAJB5 | 2 | 2 | 0 | 1 | 1 |
| DNAJC24 | 1 | 1 | 0,024012981 | 0,609693904 | 1 |
| GNAS | 1 | 1 | 0 | 1 | 1 |
| SCN4B | 1 | 1 | -0,107305893 | 0,576914341 | 1 |
| UPRT | 2 | 2 | 0 | 1 | 1 |
| ILF2 | 4 | 2 | 1,49995E-17 | 0,999999995 | 1 |

| | | | | | |
|---|---|---|---|---|---|
| ISOC2B | 2 | 2 | 0 | 1 | 1 |
| STRIP1;STRIP2 | 2 | 2 | -0,033356547 | 0,52712909 | 1 |
| POLR1C | 2 | 2 | -4,19991E-17 | 0,999999996 | 1 |
| NDRG2 | 1 | 1 | -4,1925E-17 | 0,999999986 | 1 |
| GM21698;GM21663 | 1 | 1 | 0 | 1 | 1 |
| GNA12;GNA13 | 1 | 1 | -0,036906392 | 0,455900532 | 1 |
| GET1 | 1 | 1 | 0 | 1 | 1 |
| EFCAB5 | 1 | 1 | 0,155027971 | 0,156689248 | 1 |
| PHLPP2 | 1 | 1 | -0,019979328 | 0,698355475 | 1 |
| ERCC6L | 1 | 1 | 0 | 1 | 1 |
| SGO2 | 1 | 1 | 0 | 1 | 1 |
| KCNA10 | 2 | 2 | 0,080689661 | 0,355240192 | 1 |
| ELP4 | 5 | 5 | -9,55174E-19 | 0,999999998 | 1 |
| TMEM177 | 1 | 1 | 0 | 1 | 1 |
| GALNS | 1 | 1 | 2,32271E-19 | 1 | 1 |
| NBAS | 4 | 4 | 1,11332E-17 | 0,999999998 | 1 |
| TPM3 | 6 | 6 | 0 | 1 | 1 |
| LIRE1 | 1 | 1 | 0 | 1 | 1 |
| CEP97 | 2 | 2 | -1,00322E-16 | 0,999999985 | 1 |
| MRPS21 | 2 | 2 | -1,04615E-19 | 1 | 1 |
| IGSF21 | 8 | 8 | 3,1126E-16 | 0,999999967 | 1 |
| ASB6 | 1 | 1 | 0 | 1 | 1 |
| DENND4A | 2 | 2 | 0 | 1 | 1 |
| UBAC1 | 3 | 3 | 0,12495222 | 0,359707284 | 1 |
| SEC31B | 1 | 1 | 0 | 1 | 1 |
| COX20 | 2 | 2 | 5,21029E-16 | 0,999999953 | 1 |
| MTCO2 | 12 | 12 | -6,69827E-17 | 0,999999994 | 1 |
| DNAL1 | 4 | 3 | 0 | 1 | 1 |
| ABCA1 | 2 | 2 | 1,19452E-17 | 0,999999993 | 1 |
| LENG8 | 1 | 1 | 0 | 1 | 1 |
| PRPF31 | 2 | 1 | 0,197149517 | 0,373773025 | 1 |
| TUBB2A;TUBB6;TUB | 3 | 3 | 0,042599987 | 0,652042291 | 1 |
| TUBB4B;TUBB5;TUBI | 2 | 2 | 2,04702E-19 | 1 | 1 |

| | | | | | |
|---|---|---|---|---|---|
| CARNMT1 | 1 | 1 | 0,046841368 | 0,536686788 | 1 |
| PKP2 | 9 | 9 | 6,52524E-16 | 0,999999921 | 1 |
| VAV1 | 1 | 1 | 0 | 1 | 1 |
| GCA | 2 | 2 | -5,54925E-19 | 0,999999999 | 1 |
| DCAKD | 6 | 6 | 3,49023E-17 | 0,999999985 | 1 |
| COQ9 | 12 | 12 | -9,92273E-17 | 0,999999983 | 1 |
| SMAP | 4 | 1 | 0 | 1 | 1 |
| CEP152 | 1 | 1 | 0 | 1 | 1 |
| LRRTM2 | 4 | 4 | 0 | 1 | 1 |
| RCAN1 | 6 | 6 | 0 | 1 | 1 |
| UBE2D3 | 1 | 1 | -0,037542858 | 0,585210291 | 1 |
| PHYHIPL | 4 | 4 | 1,02075E-18 | 0,999999998 | 1 |
| MAP2K3;MAP2K6 | 1 | 1 | 0 | 1 | 1 |
| NAGA | 3 | 3 | -0,085595952 | 0,121351008 | 1 |
| EFNA3 | 1 | 1 | 0 | 1 | 1 |
| GUCY1A1 | 2 | 2 | -0,036056245 | 0,637348959 | 1 |
| TUBB2A | 2 | 2 | 0 | 1 | 1 |
| TUBB4B | 1 | 1 | 0 | 1 | 1 |
| TUBB2B | 2 | 2 | 0 | 1 | 1 |
| MARCHF5 | 6 | 6 | -1,96884E-21 | 1 | 1 |
| CDK17 | 5 | 5 | 2,15784E-18 | 0,999999997 | 1 |
| 1700014D04RIK | 1 | 1 | 0 | 1 | 1 |
| ACTL6B | 4 | 4 | 0 | 1 | 1 |
| TANC2 | 3 | 3 | -2,70214E-15 | 0,999999868 | 1 |
| SHOC2 | 4 | 4 | 3,79076E-16 | 0,999999945 | 1 |
| SPAG17 | 1 | 1 | 0 | 1 | 1 |
| SERINC1 | 3 | 3 | 0,049314553 | 0,401876583 | 1 |
| DNAH14 | 2 | 2 | 0 | 1 | 1 |
| NEU1 | 1 | 1 | -0,079516929 | 0,400920225 | 1 |
| ZFP407 | 1 | 1 | 0,108707783 | 0,530576685 | 1 |
| ELAC1 | 1 | 1 | 0 | 1 | 1 |
| TSHR | 2 | 2 | 5,01459E-14 | 0,999999578 | 1 |
| MAP2 | 1 | 1 | 0 | 1 | 1 |

| | | | | | |
|---|---|---|---|---|---|
| KMT2D | 1 | 1 | 0 | 1 | 1 |
| CX3CL1 | 2 | 2 | -2,56581E-16 | 0,999999961 | 1 |
| NTRK2 | 3 | 3 | 4,55926E-16 | 0,999999968 | 1 |
| MAP1LC3A;MAP1LC: | 2 | 2 | -1,09098E-17 | 0,999999995 | 1 |
| CBWD1 | 2 | 2 | 0,0498354 | 0,999195496 | 1 |
| P2YR13 | 1 | 1 | 0 | 1 | 1 |
| GSK3B | 5 | 5 | 1,61326E-17 | 0,999999989 | 1 |
| ADHFE1 | 4 | 4 | -1,85144E-17 | 0,999999993 | 1 |
| GABRA2 | 5 | 5 | -1,42421E-18 | 0,999999998 | 1 |
| R3HDM1 | 1 | 1 | 0 | 1 | 1 |
| CCDC91 | 3 | 3 | -2,35251E-18 | 0,999999998 | 1 |
| ARF4 | 6 | 6 | 1,1244E-16 | 0,999999975 | 1 |
| EID2 | 1 | 1 | -0,084943133 | 0,665346328 | 1 |
| UBA5 | 15 | 14 | 0,013136017 | 0,545041839 | 1 |
| DDX19B | 1 | 1 | 1,09463E-17 | 0,999999995 | 1 |
| HMG20B | 1 | 1 | -1,37087E-19 | 1 | 1 |
| SEC62 | 2 | 2 | 0,046556362 | 0,442252783 | 1 |
| COQ8B | 2 | 2 | 0 | 1 | 1 |
| ZNFX1 | 1 | 1 | 0 | 1 | 1 |
| XRN1 | 1 | 1 | -2,06333E-17 | 0,999999994 | 1 |
| TPM3;TPM1;TPM2 | 4 | 3 | -2,47729E-16 | 0,999999985 | 1 |
| PPM1B;PPM1A | 5 | 3 | 0 | 1 | 1 |
| FBXW5 | 1 | 1 | 0 | 1 | 1 |
| SNX19 | 2 | 2 | 0 | 1 | 1 |
| WASHC3 | 1 | 1 | 0 | 1 | 1 |
| CSPG4B | 1 | 1 | 0 | 1 | 1 |
| VMN2R16 | 1 | 1 | 0,15961897 | 0,281211731 | 1 |
| SPRYD7 | 3 | 3 | 1,29466E-17 | 0,999999996 | 1 |
| ATP2B2 | 4 | 4 | 0 | 1 | 1 |
| SH3BGRL3 | 5 | 5 | 0,059505955 | 0,153843318 | 1 |
| ATCAY | 7 | 7 | 0,061280391 | 0,134687242 | 1 |
| KIF5A | 5 | 5 | 0,033914221 | 0,322741377 | 1 |
| KRT14;KRT16 | 8 | 5 | 0,647794227 | 0,128784227 | 1 |

| | | | | | |
|---|---|---|---|---|---|
| TUBB1;TUBB4B;TUBI | 1 | 1 | 0 | 1 | 1 |
| CLASP1;CLASP2 | 1 | 1 | 0 | 1 | 1 |
| PLEKHH1;PLEKHH2 | 1 | 1 | 0 | 1 | 1 |
| DNM1 | 6 | 6 | 4,02801E-16 | 0,999999949 | 1 |
| TBCK | 4 | 4 | 0,031334714 | 0,401205896 | 1 |
| CEP120 | 1 | 1 | -0,065967131 | 0,494295798 | 1 |
| TNS2 | 2 | 2 | 0,006663637 | 0,772134542 | 1 |
| SHMT1;SHMT2 | 1 | 1 | 0 | 1 | 1 |
| OXNAD1 | 3 | 3 | -1,80198E-16 | 0,999999974 | 1 |
| CRHBP | 2 | 2 | 0 | 1 | 1 |
| ZFP804B | 1 | 1 | 0,092148671 | 0,225992014 | 1 |
| ERLIN2;ERLIN1 | 3 | 3 | 0 | 1 | 1 |
| PSME1 | 6 | 6 | 0,017553606 | 0,596343739 | 1 |
| TUBB4B;TUBB5;TUBI | 6 | 6 | 6,42546E-16 | 0,999999941 | 1 |
| LAMA5 | 1 | 1 | 0 | 1 | 1 |
| SCRG1 | 1 | 1 | 0 | 1 | 1 |
| MAG | 1 | 1 | -0,128950871 | 0,25057911 | 1 |
| CDK1 | 1 | 1 | 0 | 1 | 1 |
| CACNA1C;CACNA1F | 1 | 1 | -1,32448E-14 | 0,999999751 | 1 |
| FAM210B | 2 | 2 | 0,126846575 | 0,150254513 | 1 |
| OSGEP | 6 | 3 | 0 | 1 | 1 |
| PSD3 | 2 | 2 | 0,055928227 | 0,278742432 | 1 |
| MESP2 | 1 | 1 | 0 | 1 | 1 |
| TGOLN1 | 2 | 2 | 0 | 1 | 1 |
| DCN | 4 | 3 | 5,04727E-18 | 0,999999998 | 1 |
| SZRD1 | 1 | 1 | 0 | 1 | 1 |
| TAF5 | 1 | 1 | 0 | 1 | 1 |
| ANKH | 3 | 3 | 0 | 1 | 1 |
| SLC2A4 | 1 | 1 | 0 | 1 | 1 |
| KRTCAP2 | 1 | 1 | 0 | 1 | 1 |
| USP33 | 3 | 3 | 0,023309548 | 0,581920608 | 1 |
| RPH3A;DOC2A | 1 | 1 | 0 | 1 | 1 |
| MAPK9 | 6 | 6 | 0 | 1 | 1 |

| | | | | | |
|---|---|---|---|---|---|
| SSBP1 | 10 | 10 | -2,69357E-16 | 0,999999964 | 1 |
| MRPL20 | 1 | 1 | 0 | 1 | 1 |
| OSBPL9 | 3 | 3 | -0,072956221 | 0,211760097 | 1 |
| TXNDC17 | 3 | 3 | -2,89996E-17 | 0,999999994 | 1 |
| AKT1;AKT2;AKT3 | 3 | 3 | 2,67749E-17 | 0,999999991 | 1 |
| DSTYK | 1 | 1 | 0,125452781 | 0,25839962 | 1 |
| MON1A | 3 | 3 | 7,21487E-17 | 0,999999986 | 1 |
| VPS72 | 1 | 1 | 0 | 1 | 1 |
| CRYGE;CRYGF | 1 | 1 | 0,214896378 | 0,152983201 | 1 |
| SURF1 | 3 | 3 | -1,59905E-18 | 0,999999999 | 1 |
| HNRNPF | 2 | 2 | 4,09629E-18 | 0,999999997 | 1 |
| 4930512M02RIK | 1 | 1 | 0,126448401 | 0,136251342 | 1 |
| HSPA1L;HSPA2;HSPA | 3 | 3 | 0 | 1 | 1 |
| CDH8 | 2 | 2 | 2,35916E-16 | 0,999999965 | 1 |
| ABCA9 | 4 | 4 | 0 | 1 | 1 |
| UFM1 | 2 | 2 | 0 | 1 | 1 |
| PLXNA3 | 1 | 1 | 0 | 1 | 1 |
| SLC27A2 | 1 | 1 | 0 | 1 | 1 |
| AFAP1 | 1 | 1 | 0 | 1 | 1 |
| ITCH;NEDD4L | 1 | 1 | 0 | 1 | 1 |
| PSME3 | 6 | 2 | 1,22853E-18 | 1 | 1 |
| ADGB | 2 | 2 | 2,42008E-16 | 0,999999981 | 1 |
| FAM107A | 1 | 1 | 0 | 1 | 1 |
| CDC27 | 2 | 2 | 1,72817E-18 | 0,999999998 | 1 |
| PCDHA3 | 1 | 1 | 0,262134215 | 0,24681775 | 1 |
| SCN7A | 1 | 1 | 2,17127E-19 | 1 | 1 |
| PPP6R2 | 1 | 1 | 0 | 1 | 1 |
| POLR2B | 4 | 2 | 0 | 1 | 1 |
| DNAJB6 | 1 | 1 | 0 | 1 | 1 |
| PABPC1L | 1 | 1 | 0 | 1 | 1 |
| NAA10;NAA12 | 3 | 3 | -1,90937E-17 | 0,999999993 | 1 |
| MIX23 | 3 | 3 | 7,14576E-19 | 0,999999999 | 1 |
| EPB41L3 | 2 | 2 | 0 | 1 | 1 |

| | | | | | |
|---|---|---|---|---|---|
| TMPPE | 2 | 2 | 1,60733E-16 | 0,999999979 | 1 |
| PRSS33 | 1 | 1 | 0 | 1 | 1 |
| ARHGEF11 | 13 | 13 | -1,82236E-17 | 0,999999992 | 1 |
| LMF1 | 1 | 1 | 0 | 1 | 1 |
| CLDN1 | 1 | 1 | 0 | 1 | 1 |
| IER5 | 1 | 1 | 0 | 1 | 1 |
| GAPDH | 2 | 2 | 0 | 1 | 1 |
| STAMBP | 3 | 3 | -3,27447E-15 | 0,999999862 | 1 |
| BZW2 | 3 | 3 | 0 | 1 | 1 |
| ANO6 | 1 | 1 | 0 | 1 | 1 |
| GRIA2;GRIA4 | 1 | 1 | 0 | 1 | 1 |
| GOLGA7 | 2 | 2 | 0,011327951 | 0,696361365 | 1 |
| TMEM192 | 1 | 1 | 0 | 1 | 1 |
| UBE2D1 | 2 | 2 | 0,060964048 | 0,37872558 | 1 |
| UBE2D2B | 2 | 2 | 0,03659781 | 0,503199824 | 1 |
| RNF14 | 1 | 1 | 0 | 1 | 1 |
| MTATP8 | 3 | 3 | -1,29202E-15 | 0,999999932 | 1 |
| FLRT3 | 4 | 4 | 1,3157E-16 | 0,999999975 | 1 |
| ABLIM3 | 1 | 1 | 0 | 1 | 1 |
| WDR45 | 3 | 3 | 0 | 1 | 1 |
| DNAJB7 | 2 | 2 | -1,60351E-17 | 0,999999993 | 1 |
| PHF14 | 1 | 1 | 0 | 1 | 1 |
| MYC | 1 | 1 | 0 | 1 | 1 |
| SARG | 1 | 1 | 0 | 1 | 1 |
| SNX10 | 2 | 2 | -2,25026E-19 | 1 | 1 |
| KHDRBS1 | 1 | 1 | 0,130550102 | 0,258588007 | 1 |
| ADRM1 | 8 | 6 | 0 | 1 | 1 |
| ODAD2 | 1 | 1 | 0 | 1 | 1 |
| GNAI1;GNAI2;GNAI3 | 4 | 4 | 2,4187E-13 | 0,999998589 | 1 |
| HNRNPDL | 2 | 1 | 0 | 1 | 1 |
| GUCY2C | 1 | 1 | 0 | 1 | 1 |
| CEACAM13 | 1 | 1 | -0,27187448 | 0,507268842 | 1 |
| CYP3A16 | 1 | 1 | 0 | 1 | 1 |

| | | | | | |
|---|---|---|---|---|---|
| PRKCH | 1 | 1 | 0 | 1 | 1 |
| PLPP1 | 1 | 1 | 0 | 1 | 1 |
| SMAP1 | 4 | 4 | 5,03611E-20 | 1 | 1 |
| SLC35F3 | 1 | 1 | 0 | 1 | 1 |
| FAM210A | 5 | 5 | 4,14605E-17 | 0,999999992 | 1 |
| AGAP3;AGAP1 | 2 | 2 | -1,4855E-17 | 0,999999996 | 1 |
| KCNQ2 | 1 | 1 | 0 | 1 | 1 |
| CHRM2 | 1 | 1 | 0 | 1 | 1 |
| ZC3HAV1 | 1 | 1 | -2,06012E-18 | 0,999999997 | 1 |
| TMCO1 | 3 | 3 | -9,86194E-17 | 0,999999983 | 1 |
| SARS1 | 1 | 1 | 0,05945527 | 0,427221035 | 1 |
| PTMA | 4 | 2 | 0 | 1 | 1 |
| SPAG9;MAPK8IP3 | 6 | 6 | 0,024885424 | 0,41524904 | 1 |
| LIG3 | 1 | 1 | 0 | 1 | 1 |
| DOCK4;DOCK3 | 3 | 3 | -1,04962E-15 | 0,999999931 | 1 |
| ATP6V0A4;ATP6V0A | 1 | 1 | 0,030960323 | 0,513727655 | 1 |
| ANO1 | 1 | 1 | 0 | 1 | 1 |
| FKBP2 | 4 | 4 | 0 | 1 | 1 |
| GSTT1 | 2 | 2 | 0 | 1 | 1 |
| BSDC1 | 4 | 4 | 0,078154817 | 0,148841814 | 1 |
| RP1 | 2 | 2 | 0 | 1 | 1 |
| CCDC106 | 1 | 1 | -0,091063406 | 0,343755073 | 1 |
| A630010A05RIK | 1 | 1 | 0 | 1 | 1 |
| NDUFA5 | 8 | 8 | -5,09714E-18 | 0,999999997 | 1 |
| LIMCH1 | 1 | 1 | 0 | 1 | 1 |
| GRM1 | 8 | 8 | 0 | 1 | 1 |
| NEFH;KRT6A;KRT2;K | 2 | 2 | 1,366003442 | 0,58117378 | 1 |
| NEFL;INA | 2 | 1 | 0 | 1 | 1 |
| sp\|Q9CPS8\|SMAKA_ | 1 | 1 | -0,218058264 | 0,423659079 | 1 |
| GPAA1 | 3 | 3 | -3,87472E-20 | 1 | 1 |
| GJC3 | 2 | 2 | -0,078654396 | 0,256423928 | 1 |
| SNX17 | 5 | 5 | 0 | 1 | 1 |
| GNG7 | 3 | 3 | 3,13675E-13 | 0,999998553 | 1 |

| | | | | | |
|---|---|---|---|---|---|
| MRPL12 | 5 | 5 | -2,08852E-15 | 0,999999901 | 1 |
| CEP164 | 1 | 1 | 0 | 1 | 1 |
| SYT1;SYT5 | 5 | 5 | 0,031925194 | 0,289129022 | 1 |
| GTF3C4 | 1 | 1 | 2,95084E-17 | 0,999999991 | 1 |
| CHD2;CHD1 | 1 | 1 | 0 | 1 | 1 |
| BCAT2 | 3 | 3 | -2,46496E-18 | 0,999999997 | 1 |
| RPL28 | 3 | 3 | 0,086722319 | 0,438734134 | 1 |
| BSG | 9 | 9 | 0,027299129 | 0,463262208 | 1 |
| UGT2B1 | 1 | 1 | 0 | 1 | 1 |
| GPT2;GPT | 1 | 1 | 0 | 1 | 1 |
| FHL2 | 1 | 1 | 0 | 1 | 1 |
| WNT7A | 1 | 1 | 0 | 1 | 1 |
| MACF1 | 2 | 2 | -0,088957605 | 0,276568412 | 1 |
| RGS2 | 1 | 1 | 0 | 1 | 1 |
| CAST | 1 | 1 | 0,062715165 | 0,476518102 | 1 |
| GTF3C1 | 2 | 2 | 6,13836E-20 | 1 | 1 |
| TACC2 | 1 | 1 | 0 | 1 | 1 |
| LASP1;NEBL | 1 | 1 | 0 | 1 | 1 |
| OSBPL8;OSBPL5 | 1 | 1 | 0 | 1 | 1 |
| HOOK1 | 2 | 1 | 0 | 1 | 1 |
| PPP1R13B | 9 | 9 | -6,58325E-17 | 0,999999979 | 1 |
| RFK | 3 | 3 | 0 | 1 | 1 |
| SYCP1 | 1 | 1 | 0 | 1 | 1 |
| RPS24 | 3 | 2 | 0,181398072 | 0,391233092 | 1 |
| GNL3 | 1 | 1 | -0,032324632 | 0,616512993 | 1 |
| RPL30 | 8 | 8 | 0,097627998 | 0,160709284 | 1 |
| SYN2;SYN3 | 2 | 2 | 2,88327E-16 | 0,999999967 | 1 |
| PLAG1 | 1 | 1 | 1,14994E-18 | 0,999999998 | 1 |
| ZGRF1 | 1 | 1 | 0 | 1 | 1 |
| MPC1 | 4 | 4 | -0,099654702 | 0,157857743 | 1 |
| ANKS1B | 2 | 2 | 0 | 1 | 1 |
| MMS22L | 1 | 1 | 0 | 1 | 1 |
| CDC14A | 1 | 1 | 0 | 1 | 1 |

| | | | | | |
|---|---|---|---|---|---|
| GLRX | 3 | 3 | 0 | 1 | 1 |
| DNM1 | 3 | 3 | 0 | 1 | 1 |
| LIN9 | 1 | 1 | 0 | 1 | 1 |
| TICAM1 | 1 | 1 | 0,047430287 | 0,497899875 | 1 |
| ARXES1;ARXES2 | 3 | 3 | -6,96138E-18 | 0,999999992 | 1 |
| LZIC | 7 | 6 | 9,44423E-17 | 0,99999998 | 1 |
| DAO | 1 | 1 | -0,163702897 | 0,229398301 | 1 |
| CALR3 | 1 | 1 | 0 | 1 | 1 |
| SERPINA1B;SERPINA | 6 | 1 | 0 | 1 | 1 |
| COL28A1 | 1 | 1 | 0 | 1 | 1 |
| KIF21A | 1 | 1 | -7,78702E-19 | 0,999999998 | 1 |
| ARRB2 | 3 | 3 | -3,03795E-17 | 0,999999987 | 1 |
| PDE12 | 7 | 6 | 3,40171E-19 | 1 | 1 |
| ISCU | 5 | 5 | 2,57312E-18 | 0,999999999 | 1 |
| RALA;RALB | 6 | 6 | 3,88925E-17 | 0,999999982 | 1 |
| EP400 | 1 | 1 | 1,35683E-17 | 0,999999992 | 1 |
| ABL1;ABL2 | 3 | 3 | 2,80971E-17 | 0,999999991 | 1 |
| AAK1;BMP2K | 2 | 2 | 0,01606537 | 0,614109365 | 1 |
| PURB;PURG | 2 | 2 | 0 | 1 | 1 |
| PPP1R2 | 5 | 4 | -5,3302E-17 | 0,999999979 | 1 |
| KRT24;KRT36;KRT18; | 1 | 1 | 0 | 1 | 1 |
| PSMB6 | 4 | 4 | 0 | 1 | 1 |
| MINPP1 | 1 | 1 | 0 | 1 | 1 |
| ZFYVE19 | 3 | 3 | 0 | 1 | 1 |
| PDIA6 | 2 | 2 | -8,30085E-18 | 0,999999995 | 1 |
| RAB3D | 3 | 3 | -1,98411E-17 | 0,999999994 | 1 |
| MRTFB;MRTFA;MYO | 1 | 1 | 0 | 1 | 1 |
| CDK14;CDK1;CDK5;C | 1 | 1 | 0 | 1 | 1 |
| STK24;STK25 | 3 | 3 | 0,030348051 | 0,512975395 | 1 |
| KCNH1 | 1 | 1 | 0,113551122 | 0,234146686 | 1 |
| WWOX | 4 | 4 | 0 | 1 | 1 |
| OXA1L | 2 | 2 | -0,04183166 | 0,612852723 | 1 |
| NUBP2 | 3 | 3 | 0,02777232 | 0,63453182 | 1 |

| | | | | | |
|---|---|---|---|---|---|
| RPL13A | 3 | 3 | 0 | 1 | 1 |
| ALG11 | 1 | 1 | 0 | 1 | 1 |
| CARD14 | 1 | 1 | -0,006619781 | 0,845582035 | 1 |
| SZT2 | 3 | 3 | -0,01002325 | 0,700152272 | 1 |
| KRT6A;KRT76;KRT5 | 2 | 2 | 0,824985839 | 0,249856838 | 1 |
| VPS37C | 2 | 2 | 0,164159394 | 0,132826222 | 1 |
| PCNX3 | 2 | 2 | 0 | 1 | 1 |
| CROCC | 2 | 2 | 0 | 1 | 1 |
| VPS8 | 3 | 3 | -2,35181E-17 | 0,999999995 | 1 |
| DIAPH2 | 3 | 3 | -1,70633E-16 | 0,999999967 | 1 |
| GATC | 1 | 1 | 0 | 1 | 1 |
| TMEM63B | 1 | 1 | 1,34597E-20 | 1 | 1 |
| RDH12 | 1 | 1 | -3,48655E-19 | 0,999999999 | 1 |
| USP25 | 3 | 1 | 0,013971722 | 0,806515288 | 1 |
| LANCL3 | 1 | 1 | 0,178171849 | 0,157532138 | 1 |
| sp\|Q922C1\|CS044_I | 1 | 1 | -0,035789978 | 0,475842766 | 1 |
| DNAH2 | 1 | 1 | 0 | 1 | 1 |
| TRPM3 | 1 | 1 | 0 | 1 | 1 |
| HIKESHI | 3 | 3 | 0,007334043 | 0,763708095 | 1 |
| TBL1XR1 | 1 | 1 | -0,048684157 | 0,584752774 | 1 |
| TCAIM | 1 | 1 | 0 | 1 | 1 |
| PTK2 | 7 | 6 | -4,13632E-17 | 0,999999986 | 1 |
| ENO1;ENO3 | 2 | 2 | 4,7818E-16 | 0,999999958 | 1 |
| TBPL2 | 1 | 1 | 0 | 1 | 1 |
| CCDC32 | 1 | 1 | 0 | 1 | 1 |
| UBP1 | 4 | 4 | 1,19806E-12 | 0,999996642 | 1 |
| KRT18;KRT17 | 1 | 1 | 0 | 1 | 1 |
| MINDY1 | 3 | 3 | -0,067997583 | 0,221637387 | 1 |
| PSD2 | 2 | 2 | 0 | 1 | 1 |
| HAS1 | 1 | 1 | 0 | 1 | 1 |
| ZSWIM8 | 1 | 1 | -0,089912254 | 0,223638301 | 1 |
| COMMD5 | 1 | 1 | 0,172518268 | 0,229822868 | 1 |
| NCOR2 | 1 | 1 | 0 | 1 | 1 |

| | | | | | |
|---|---|---|---|---|---|
| MAPK15 | 1 | 1 | 0,029115747 | 0,581900371 | 1 |
| DENR | 5 | 5 | -1,37552E-17 | 0,999999994 | 1 |
| NLGN1 | 2 | 2 | 1,99312E-16 | 0,999999964 | 1 |
| DCUN1D2 | 2 | 2 | 1,55562E-20 | 1 | 1 |
| PRUNE2 | 4 | 4 | -1,02432E-16 | 0,999999977 | 1 |
| ULK3 | 1 | 1 | 0 | 1 | 1 |
| APOH | 1 | 1 | 0 | 1 | 1 |
| NR2C2 | 1 | 1 | 0 | 1 | 1 |
| RRAS;RRAS2 | 3 | 3 | 0,060854308 | 0,424957415 | 1 |
| OGDH | 2 | 2 | -0,031391808 | 0,63218142 | 1 |
| SMC1A | 1 | 1 | 0,004071237 | 0,866639974 | 1 |
| CDK5RAP3 | 1 | 1 | 0 | 1 | 1 |
| DNM1;DNM2;DNM3 | 7 | 7 | 2,17544E-17 | 0,999999986 | 1 |
| LRTM2 | 2 | 2 | 5,03261E-19 | 0,999999999 | 1 |
| FXN | 2 | 2 | -0,127072029 | 0,214240325 | 1 |
| LRFN2;LRFN4 | 1 | 1 | 0 | 1 | 1 |
| MAP3K10 | 1 | 1 | 0 | 1 | 1 |
| TIMM13 | 3 | 3 | -1,24158E-16 | 0,999999987 | 1 |
| TXNDC9 | 4 | 4 | 0 | 1 | 1 |
| TBCEL | 7 | 6 | -0,010896549 | 0,661342853 | 1 |
| IRGM1 | 5 | 5 | 3,36222E-16 | 0,999999969 | 1 |
| DCLK2;DCLK1 | 1 | 1 | 0 | 1 | 1 |
| SPO11 | 1 | 1 | -0,057368484 | 0,321869868 | 1 |
| NCEH1 | 15 | 14 | 4,09829E-16 | 0,999999927 | 1 |
| STMN3;STMN1;STM | 1 | 1 | -4,94649E-20 | 1 | 1 |
| HOOK1;HOOK3 | 1 | 1 | 0,262422158 | 0,128253329 | 1 |
| ODF2 | 1 | 1 | 4,61976E-17 | 0,999999988 | 1 |
| ZNF608 | 1 | 1 | 0 | 1 | 1 |
| GRM2;GRM4 | 2 | 2 | -4,48264E-16 | 0,999999955 | 1 |
| GNG12 | 5 | 5 | -8,21632E-16 | 0,999999908 | 1 |
| OBSCN | 1 | 1 | 0 | 1 | 1 |
| ADD1 | 1 | 1 | -0,106293561 | 0,182908395 | 1 |
| ADD1 | 1 | 1 | 0 | 1 | 1 |

| | | | | | |
|---|---|---|---|---|---|
| ARHGAP22 | 1 | 1 | -0,039813176 | 0,427080361 | 1 |
| ATPAF2 | 2 | 2 | 0 | 1 | 1 |
| SLC12A5;SLC12A6 | 3 | 3 | 6,12497E-19 | 0,999999999 | 1 |
| BBS9 | 1 | 1 | 0 | 1 | 1 |
| DNAH10 | 1 | 1 | 0 | 1 | 1 |
| SPECC1 | 1 | 1 | 0 | 1 | 1 |
| ARHGAP36 | 1 | 1 | 0,013585951 | 0,746173202 | 1 |
| GPC6 | 1 | 1 | 0 | 1 | 1 |
| GSE1 | 1 | 1 | 0 | 1 | 1 |
| LAMC3 | 1 | 1 | 0,031755319 | 0,623825547 | 1 |
| KRT6A | 19 | 6 | 1,83955E-16 | 0,999999985 | 1 |
| PEX1 | 2 | 2 | 2,54633E-10 | 0,999965771 | 1 |
| TAX1BP1 | 4 | 4 | 9,77977E-21 | 1 | 1 |
| APOL7B;APOL7E | 1 | 1 | 0 | 1 | 1 |
| GOLGA4 | 1 | 1 | 4,39136E-20 | 1 | 1 |
| BBS7 | 1 | 1 | 0 | 1 | 1 |
| MCUR1 | 4 | 4 | -0,032053632 | 0,524350286 | 1 |
| KIF1B | 3 | 3 | 0 | 1 | 1 |
| 4931408C20RIK | 2 | 2 | -1,10124E-18 | 0,999999999 | 1 |
| ARC | 1 | 1 | 0 | 1 | 1 |
| LMBRD1 | 1 | 1 | 0 | 1 | 1 |
| DMAC2 | 1 | 1 | -4,67281E-20 | 1 | 1 |
| TRIM58 | 1 | 1 | -0,077178391 | 0,346481883 | 1 |
| KCNT1 | 1 | 1 | 0 | 1 | 1 |
| PFKFB2;PFKFB4 | 1 | 1 | 0 | 1 | 1 |
| C1QB | 4 | 4 | 0,055779058 | 0,542684869 | 1 |
| KRT14;KRT42;KRT17 | 3 | 2 | 6,41548E-18 | 0,999999998 | 1 |
| TOR1AIP1 | 1 | 1 | 0 | 1 | 1 |
| SCYL2 | 13 | 12 | 0 | 1 | 1 |
| HDAC5 | 4 | 4 | -4,87072E-16 | 0,999999945 | 1 |
| PPP1R12B | 2 | 2 | -0,049059026 | 0,49245893 | 1 |
| MIGA2 | 7 | 7 | -9,50956E-17 | 0,99999997 | 1 |
| TRAPPC6B | 5 | 5 | 0 | 1 | 1 |

| | | | | | |
|---|---|---|---|---|---|
| FASTKD1 | 1 | 1 | 0 | 1 | 1 |
| ANKS1B | 1 | 1 | 0 | 1 | 1 |
| PRC1 | 1 | 1 | 0 | 1 | 1 |
| TRIM6 | 1 | 1 | 0,002831594 | 0,942237483 | 1 |
| PCDH1 | 6 | 6 | 0,046494833 | 0,292774739 | 1 |
| HMGCS2;HMGCS1 | 1 | 1 | 0,092312482 | 0,404226275 | 1 |
| PDILT | 1 | 1 | 0 | 1 | 1 |
| TLR13 | 1 | 1 | 0 | 1 | 1 |
| SEZ6L2 | 2 | 2 | -0,029506354 | 0,48236522 | 1 |
| PLXNA2;PLXNA4 | 5 | 5 | -7,77199E-17 | 0,999999977 | 1 |
| MYO1F | 1 | 1 | 0 | 1 | 1 |
| ARMC8 | 3 | 3 | -0,012511276 | 0,703387732 | 1 |
| COG5 | 2 | 2 | -0,130260536 | 0,253158396 | 1 |
| GNAI1;GNAI2;GNAO | 5 | 5 | 0,023218146 | 0,501814124 | 1 |
| PPP2R2A;PPP2R2C | 2 | 2 | 0 | 1 | 1 |
| NCF2 | 1 | 1 | 0 | 1 | 1 |
| MARK4;MARK1 | 1 | 1 | 0,026884671 | 0,537392659 | 1 |
| TMEM9B | 1 | 1 | 0 | 1 | 1 |
| CHKA | 1 | 1 | -0,09869618 | 0,261901999 | 1 |
| LARP6 | 3 | 3 | -3,258E-17 | 0,999999984 | 1 |
| IL1RAPL1 | 2 | 2 | -2,07217E-18 | 0,999999999 | 1 |
| PTPRC | 1 | 1 | 0 | 1 | 1 |
| SNX24 | 1 | 1 | 0,070318735 | 0,518087525 | 1 |
| WDR6 | 2 | 2 | 0 | 1 | 1 |
| RHBDD2 | 1 | 1 | 0 | 1 | 1 |
| CYB5D2 | 2 | 2 | 1,16448E-17 | 0,999999997 | 1 |
| RIOX1 | 1 | 1 | 0 | 1 | 1 |
| COQ10B | 1 | 1 | 0 | 1 | 1 |
| PDK1;PDK3 | 1 | 1 | 0 | 1 | 1 |
| MAPK10 | 4 | 4 | 5,32049E-18 | 0,999999997 | 1 |
| AKR1C14 | 1 | 1 | 0 | 1 | 1 |
| AMIGO2 | 1 | 1 | 0,102112791 | 0,266903455 | 1 |
| RIMS3 | 2 | 2 | 0 | 1 | 1 |

| | | | | | |
|---|---|---|---|---|---|
| COX7A1 | 1 | 1 | 0 | 1 | 1 |
| CEPT1 | 1 | 1 | 0 | 1 | 1 |
| GSTM4 | 4 | 3 | -0,149902294 | 0,139100663 | 1 |
| RRAS | 3 | 3 | 0 | 1 | 1 |
| RBP3 | 1 | 1 | 0 | 1 | 1 |
| DAZAP1 | 2 | 1 | 0,393037735 | 0,138805171 | 1 |
| TMEM126B | 1 | 1 | 0 | 1 | 1 |
| VLDLR | 2 | 2 | 4,97612E-19 | 0,999999999 | 1 |
| ATP7A | 1 | 1 | 0 | 1 | 1 |
| ATP4A;ATP1A4 | 1 | 1 | 0 | 1 | 1 |
| TMEM256 | 2 | 2 | -1,21553E-16 | 0,999999986 | 1 |
| ACOX3 | 5 | 5 | -8,28981E-14 | 0,999999235 | 1 |
| RRAGA | 1 | 1 | 0 | 1 | 1 |
| COA7 | 3 | 3 | -6,8017E-18 | 0,999999994 | 1 |
| GRSF1 | 2 | 2 | 0 | 1 | 1 |
| PTK2;PTK2B | 1 | 1 | 0 | 1 | 1 |
| SLC20A1 | 2 | 2 | -0,107988797 | 0,16234578 | 1 |
| NRN1 | 4 | 4 | 0,0260717 | 0,635620052 | 1 |
| CDK16;CDK17 | 2 | 2 | 6,93445E-19 | 0,999999999 | 1 |
| PRSS1 | 1 | 1 | 0 | 1 | 1 |
| SYT7 | 3 | 3 | -0,083844923 | 0,229039135 | 1 |
| ITGB5 | 1 | 1 | 0 | 1 | 1 |
| ITGB6 | 1 | 1 | 1,57588E-20 | 1 | 1 |
| PPM1L | 3 | 3 | -0,073893399 | 0,146257846 | 1 |
| KCNC2 | 3 | 3 | -6,30495E-41 | 1 | 1 |
| CD34 | 2 | 2 | 0 | 1 | 1 |
| CD2AP | 1 | 1 | 0 | 1 | 1 |
| MRPL17 | 1 | 1 | -5,49819E-17 | 0,999999986 | 1 |
| CDC5L | 1 | 1 | 0,172699773 | 0,117247871 | 1 |
| SLC25A27 | 1 | 1 | 8,99027E-18 | 0,999999994 | 1 |
| MOCS3 | 2 | 2 | 0 | 1 | 1 |
| ICOSLG | 1 | 1 | 0 | 1 | 1 |
| FAM171A1 | 7 | 7 | 0,020212108 | 0,518461982 | 1 |

| | | | | | |
|---|---|---|---|---|---|
| TCAF1 | 2 | 2 | 0 | 1 | 1 |
| PCDH1 | 5 | 5 | 0,021270847 | 0,511424328 | 1 |
| RER1 | 4 | 4 | 1,58171E-17 | 0,99999999 | 1 |
| LYRM7 | 2 | 2 | 1,67498E-15 | 0,999999935 | 1 |
| WWC1 | 2 | 2 | 0,079935779 | 0,341923272 | 1 |
| YRDC | 2 | 1 | 0 | 1 | 1 |
| SLC30A5 | 2 | 2 | 0 | 1 | 1 |
| SOGA1 | 2 | 2 | 0 | 1 | 1 |
| EPHA5 | 1 | 1 | 0 | 1 | 1 |
| LSAMP | 1 | 1 | 0 | 1 | 1 |
| KIF5C;KIF5A | 6 | 6 | 1,77375E-18 | 0,999999997 | 1 |
| CCAR1 | 1 | 1 | 0 | 1 | 1 |
| NUCB2 | 2 | 2 | -0,097499235 | 0,210283361 | 1 |
| TFCP2L1 | 1 | 1 | 0 | 1 | 1 |
| TUBB6;TUBB3 | 4 | 4 | -8,99326E-18 | 0,999999998 | 1 |
| TUBB4B;TUBB5;TUBI | 7 | 7 | 0 | 1 | 1 |
| CLSTN3 | 2 | 2 | 0 | 1 | 1 |
| KIF3A;KIF4 | 1 | 1 | -2,74615E-18 | 0,999999997 | 1 |
| ZFP101 | 1 | 1 | -0,147904962 | 0,212923683 | 1 |
| MTATP6 | 1 | 1 | 0 | 1 | 1 |
| PTS | 5 | 4 | 0 | 1 | 1 |
| KCTD10 | 1 | 1 | 0 | 1 | 1 |
| SCFD1 | 7 | 7 | 0 | 1 | 1 |
| TKTL1 | 2 | 2 | 0,033017391 | 0,514734585 | 1 |
| ANK2 | 3 | 3 | -7,89318E-18 | 0,999999995 | 1 |
| ATP1A2;ATP1A1;ATF | 6 | 6 | 1,67529E-15 | 0,999999899 | 1 |
| IQGAP2;IQGAP1 | 2 | 2 | 0,086981726 | 0,142173243 | 1 |
| TUBA3B;TUBA1A | 1 | 1 | 0 | 1 | 1 |
| LRRTM4 | 4 | 4 | 0 | 1 | 1 |
| GNB4;GNB1 | 2 | 2 | 1,28085E-19 | 1 | 1 |
| UQCR11 | 1 | 1 | 0 | 1 | 1 |
| OLFR141 | 1 | 1 | 0,292736021 | 0,136635812 | 1 |
| STING1 | 1 | 1 | -2,12702E-21 | 1 | 1 |

| | | | | | |
|---|---|---|---|---|---|
| FABP3 | 10 | 7 | -2,30991E-17 | 0,999999995 | 1 |
| GID8 | 3 | 3 | 0,041961858 | 0,457740791 | 1 |
| PIK3CA | 1 | 1 | 0 | 1 | 1 |
| NTF3 | 1 | 1 | 0 | 1 | 1 |
| CTSS | 4 | 3 | 0,000991786 | 0,931790607 | 1 |
| SBF2 | 2 | 2 | -2,88348E-16 | 0,999999967 | 1 |
| NEDD4 | 14 | 14 | 1,20698E-16 | 0,999999964 | 1 |
| CYBC1 | 1 | 1 | 0 | 1 | 1 |
| SRSF5;SRSF6;SRSF4 | 1 | 1 | 0 | 1 | 1 |
| MT-ND2 | 2 | 2 | 1,26328E-13 | 0,999999301 | 1 |
| ST6GALNAC1 | 1 | 1 | 0 | 1 | 1 |
| CNTRL | 1 | 1 | -0,024256548 | 0,705474612 | 1 |
| PPP1R12B;PPP1R12/ | 2 | 2 | -1,30315E-18 | 0,999999999 | 1 |
| SUPT6H | 4 | 1 | 0 | 1 | 1 |
| SLC35A4 | 1 | 1 | 0 | 1 | 1 |
| STK16 | 1 | 1 | 0 | 1 | 1 |
| NAA16 | 2 | 2 | -4,7874E-17 | 0,999999987 | 1 |
| LRRTM4;LRRTM3 | 1 | 1 | 0 | 1 | 1 |
| DNM2 | 1 | 1 | 0 | 1 | 1 |
| THSD7A | 5 | 5 | 0 | 1 | 1 |
| CAPG | 4 | 4 | 0,090203033 | 0,468304768 | 1 |
| 0610012G03RIK | 1 | 1 | 0 | 1 | 1 |
| RPL19 | 3 | 3 | 4,82806E-16 | 0,999999958 | 1 |
| PRKCD | 2 | 2 | 4,55911E-18 | 0,999999999 | 1 |
| PODXL2 | 1 | 1 | 0 | 1 | 1 |
| sp\|Q8WUR0\|CS012_ | 1 | 1 | 0 | 1 | 1 |
| SPRY3 | 1 | 1 | 0 | 1 | 1 |
| SYNGAP1 | 1 | 1 | 0 | 1 | 1 |
| SELENOF | 1 | 1 | 0 | 1 | 1 |
| TBC1D22B | 4 | 3 | 0 | 1 | 1 |
| TBC1D22A | 7 | 7 | -2,81043E-17 | 0,99999999 | 1 |
| GABRA2;GABRA1 | 2 | 2 | -7,72562E-15 | 0,99999982 | 1 |
| TTC27 | 1 | 1 | 0 | 1 | 1 |

| | | | | | |
|---|---|---|---|---|---|
| UNC13A;UNC13C;UN | 5 | 5 | 0 | 1 | 1 |
| ARL10 | 1 | 1 | 0 | 1 | 1 |
| STXBP1 | 2 | 2 | 0,087966317 | 0,134895532 | 1 |
| PIK3R1 | 3 | 3 | 8,14921E-13 | 0,999998428 | 1 |
| LCT | 2 | 2 | 0 | 1 | 1 |
| EPPK1 | 13 | 1 | 0,131078358 | 0,162040787 | 1 |
| MRPL28 | 4 | 4 | -7,24432E-19 | 0,999999999 | 1 |
| ANKRD46 | 1 | 1 | 0 | 1 | 1 |
| GLT1D1 | 1 | 1 | 0 | 1 | 1 |
| KPNA2 | 1 | 1 | 0 | 1 | 1 |
| PET117 | 1 | 1 | 0 | 1 | 1 |
| DPP6 | 1 | 1 | 0 | 1 | 1 |
| GIPC2 | 2 | 1 | 0 | 1 | 1 |
| SCO2 | 3 | 3 | -0,019350631 | 0,657171048 | 1 |
| SLC4A9 | 1 | 1 | 0 | 1 | 1 |
| WDR35 | 1 | 1 | -0,077995144 | 0,333221097 | 1 |
| TELO2 | 2 | 2 | 0 | 1 | 1 |
| SPRY2 | 2 | 2 | 0,117338671 | 0,122434836 | 1 |
| ARHGEF25 | 2 | 2 | 2,68513E-17 | 0,999999992 | 1 |
| HDAC4 | 1 | 1 | 0 | 1 | 1 |
| NUDT10;NUDT11 | 7 | 6 | 0 | 1 | 1 |
| RGS7 | 1 | 1 | 0 | 1 | 1 |
| NRCAM | 1 | 1 | 0,002386733 | 0,88105806 | 1 |
| PLXNA2 | 7 | 7 | 0 | 1 | 1 |
| TMEM143 | 3 | 3 | 3,77482E-19 | 0,999999999 | 1 |
| PON3 | 1 | 1 | 0 | 1 | 1 |
| ELK3 | 1 | 1 | -0,231086723 | 0,179120285 | 1 |
| BCAP29 | 3 | 3 | -7,31779E-18 | 0,999999995 | 1 |
| CAMK2A;CAMK2D;C. | 1 | 1 | 0 | 1 | 1 |
| ITPR3 | 3 | 3 | 0 | 1 | 1 |
| MRM1 | 1 | 1 | -0,123831135 | 0,143254714 | 1 |
| KIF21B | 1 | 1 | -0,076657673 | 0,399408048 | 1 |
| MTMR3 | 4 | 4 | 0 | 1 | 1 |

| | | | | | |
|---|---|---|---|---|---|
| CASP1 | 1 | 1 | 0 | 1 | 1 |
| UIMC1 | 1 | 1 | 0 | 1 | 1 |
| DZANK1 | 2 | 2 | 0 | 1 | 1 |
| RAB8A;RAB10;RAB8I | 1 | 1 | 0 | 1 | 1 |
| RILP | 1 | 1 | 0 | 1 | 1 |
| CHADL | 1 | 1 | -0,006127835 | 0,8085707 | 1 |
| YES1 | 1 | 1 | 0 | 1 | 1 |
| RAPGEFL1 | 5 | 5 | 0,037010207 | 0,412693171 | 1 |
| H2BC15;H2BC14;H2I | 5 | 4 | 0,116388297 | 0,53880154 | 1 |
| NTMT1 | 2 | 2 | 0 | 1 | 1 |
| MPP4 | 1 | 1 | 0 | 1 | 1 |
| DEF8 | 1 | 1 | -0,085060052 | 0,347736825 | 1 |
| DPH1 | 1 | 1 | 0 | 1 | 1 |
| STARD3NL | 2 | 2 | 1,57868E-14 | 0,999999692 | 1 |
| ENPP4 | 1 | 1 | 0 | 1 | 1 |
| CAMLG | 2 | 2 | 0 | 1 | 1 |
| AKR1B7;AKR1B8 | 1 | 1 | 0 | 1 | 1 |
| SLC13A1 | 1 | 1 | 0,055922089 | 0,298539025 | 1 |
| MROH1 | 1 | 1 | 2,09407E-15 | 0,999999922 | 1 |
| sp\|Q9CWU4\|CA052_ | 1 | 1 | 0,124175785 | 0,469339069 | 1 |
| TP53BP2 | 1 | 1 | -1,84758E-19 | 0,999999999 | 1 |
| GFOD2 | 1 | 1 | -2,19686E-20 | 1 | 1 |
| JAK3 | 1 | 1 | -0,088818805 | 0,237433011 | 1 |
| HGH1 | 1 | 1 | 0 | 1 | 1 |
| GDAP2 | 3 | 3 | 1,50661E-17 | 0,999999991 | 1 |
| AMFR | 3 | 3 | -1,51273E-15 | 0,999999888 | 1 |
| CALB1;CALB2 | 1 | 1 | -1,78779E-19 | 1 | 1 |
| DNAH9 | 1 | 1 | 0 | 1 | 1 |
| SORCS3 | 4 | 4 | -2,87283E-18 | 0,999999998 | 1 |
| SURF6 | 1 | 1 | -0,084273805 | 0,564034126 | 1 |
| SMURF1 | 1 | 1 | 0,066613603 | 0,327680581 | 1 |
| SCG2 | 4 | 4 | -5,06618E-16 | 0,999999922 | 1 |
| HSD11B2 | 1 | 1 | 0 | 1 | 1 |

| | | | | | |
|---|---|---|---|---|---|
| SAMD10 | 1 | 1 | 0 | 1 | 1 |
| IGHV4-1 | 1 | 1 | 0 | 1 | 1 |
| LCMT2;SEPTIN5 | 1 | 1 | 0 | 1 | 1 |
| EXOC1 | 3 | 3 | 0 | 1 | 1 |
| NUP93 | 2 | 2 | 0,057125738 | 0,31979931 | 1 |
| METAP1 | 2 | 1 | -0,121052501 | 0,391623667 | 1 |
| 2410002F23RIK | 3 | 3 | -0,059976921 | 0,33727584 | 1 |
| OLFR1180 | 1 | 1 | 0,018188297 | 0,664562012 | 1 |
| RNF112 | 1 | 1 | 0 | 1 | 1 |
| ARHGAP33;ARHGAP: | 1 | 1 | 0 | 1 | 1 |
| IYD | 1 | 1 | 0 | 1 | 1 |
| SLC33A1 | 2 | 2 | 0,131435108 | 0,228838597 | 1 |
| OPHN1 | 2 | 2 | 0 | 1 | 1 |
| NDUFAB1 | 5 | 4 | 1,97082E-16 | 0,999999969 | 1 |
| CHPF | 1 | 1 | 0 | 1 | 1 |
| VTA1 | 6 | 6 | 1,86585E-17 | 0,999999991 | 1 |
| KCNB1 | 2 | 2 | -0,002720568 | 0,857938339 | 1 |
| LRFN2 | 4 | 4 | 0,022574517 | 0,439131204 | 1 |
| AMIGO1 | 3 | 3 | -0,036788349 | 0,397416504 | 1 |
| SLC25A32 | 3 | 3 | 0 | 1 | 1 |
| SNN | 1 | 1 | 0 | 1 | 1 |
| EFCAB8 | 1 | 1 | 0 | 1 | 1 |
| DOK6 | 1 | 1 | 0 | 1 | 1 |
| NRXN2;NRXN3 | 1 | 1 | 0 | 1 | 1 |
| ALG10B | 1 | 1 | 0 | 1 | 1 |
| LRRTM3 | 1 | 1 | 0 | 1 | 1 |
| REPS1;REPS2 | 1 | 1 | -0,01459223 | 0,67134517 | 1 |
| PARVA | 2 | 2 | 8,18876E-13 | 0,999997445 | 1 |
| KCTD2 | 1 | 1 | 0,033593724 | 0,543946463 | 1 |
| DUSP23 | 1 | 1 | -1,61772E-20 | 1 | 1 |
| CDK16 | 4 | 4 | 0,0036915 | 0,792577639 | 1 |
| PRKDC | 1 | 1 | 0 | 1 | 1 |
| AKT1;AKT3 | 1 | 1 | -0,012532933 | 0,716697125 | 1 |

| | | | | | |
|---|---|---|---|---|---|
| MTM1 | 2 | 1 | -0,021213945 | 0,689577945 | 1 |
| GPATCH2L | 1 | 1 | 0 | 1 | 1 |
| UGCG | 1 | 1 | 0 | 1 | 1 |
| RHOT2 | 1 | 1 | 0,056393743 | 0,40534481 | 1 |
| EPG5 | 4 | 4 | 1,10573E-19 | 1 | 1 |
| TMEM11 | 3 | 3 | 8,34532E-17 | 0,999999977 | 1 |
| PCMTD1 | 2 | 2 | -0,010359724 | 0,767548084 | 1 |
| ANKMY1 | 1 | 1 | 0,138910487 | 0,234965852 | 1 |
| EPHB6 | 2 | 2 | -7,37294E-10 | 0,99992326 | 1 |
| PANX2 | 2 | 2 | 0 | 1 | 1 |
| ITSN2 | 1 | 1 | 0 | 1 | 1 |
| NCF1 | 1 | 1 | 0 | 1 | 1 |
| SLC35F6 | 1 | 1 | 0 | 1 | 1 |
| CHAC2 | 3 | 1 | 0 | 1 | 1 |
| SLC25A29 | 3 | 3 | -1,76558E-16 | 0,999999969 | 1 |
| KTN1 | 1 | 1 | 0 | 1 | 1 |
| S100A6 | 1 | 1 | 0 | 1 | 1 |
| ACO2 | 1 | 1 | -3,44881E-17 | 0,999999989 | 1 |
| P4HA1 | 2 | 2 | -2,61682E-16 | 0,999999989 | 1 |
| RBL2 | 1 | 1 | -0,036192961 | 0,543275963 | 1 |
| DDX10 | 1 | 1 | 0 | 1 | 1 |
| PARP14 | 1 | 1 | 0 | 1 | 1 |
| HDAC9;HDAC5 | 1 | 1 | -0,131787834 | 0,254325024 | 1 |
| ARHGAP10 | 1 | 1 | 0 | 1 | 1 |
| OS9 | 1 | 1 | 0 | 1 | 1 |
| CNTN3;CNTN4 | 1 | 1 | 0 | 1 | 1 |
| HLCS | 2 | 1 | 0 | 1 | 1 |
| RDH13 | 3 | 3 | -1,10164E-12 | 0,999998198 | 1 |
| TTC3 | 2 | 2 | 1,16962E-18 | 0,999999999 | 1 |
| SPRY4 | 2 | 2 | -0,071073566 | 0,238167254 | 1 |
| ERBIN | 3 | 3 | 0 | 1 | 1 |
| RAB8A;RAB4A;RAB1( | 1 | 1 | 0 | 1 | 1 |
| LGALS8 | 1 | 1 | 0 | 1 | 1 |

| | | | | | |
|---|---|---|---|---|---|
| RAPH1 | 7 | 7 | -1,47559E-17 | 0,999999992 | 1 |
| GABRD | 1 | 1 | -0,061734231 | 0,45124669 | 1 |
| MMP25 | 1 | 1 | 0 | 1 | 1 |
| CCDC190 | 1 | 1 | 0 | 1 | 1 |
| ANGPTL6 | 1 | 1 | -7,06906E-18 | 0,999999997 | 1 |
| DEF6 | 1 | 1 | 0 | 1 | 1 |
| SALL2 | 1 | 1 | 0 | 1 | 1 |
| VAMP2 | 4 | 4 | 1,86369E-14 | 0,999999579 | 1 |
| LSP1 | 1 | 1 | 0 | 1 | 1 |
| STAU1 | 1 | 1 | -1,91149E-15 | 0,99999993 | 1 |
| SPTAN1 | 2 | 2 | -0,064717856 | 0,164112341 | 1 |
| TMEM175 | 2 | 2 | -1,56987E-14 | 0,999999776 | 1 |
| DNAL4 | 1 | 1 | 0 | 1 | 1 |
| SLC25A12;SLC25A13 | 6 | 6 | 0 | 1 | 1 |
| FBXL2 | 1 | 1 | 0 | 1 | 1 |
| MKRN2 | 1 | 1 | -0,025282373 | 0,632695991 | 1 |
| TMEM9;TMEM9B | 1 | 1 | 0,041514458 | 0,414731335 | 1 |
| XKR4 | 2 | 2 | 0 | 1 | 1 |
| SAMD14 | 3 | 3 | -0,038517556 | 0,379677945 | 1 |
| TBX15 | 1 | 1 | 0 | 1 | 1 |
| RGS7BP | 4 | 4 | 0,029761984 | 0,418379474 | 1 |
| ITGB1 | 4 | 4 | 0 | 1 | 1 |
| GABRA1;GABRA5 | 2 | 2 | 1,28875E-17 | 0,999999995 | 1 |
| MPC2 | 7 | 7 | -6,03692E-17 | 0,999999985 | 1 |
| SELENOS | 1 | 1 | 0 | 1 | 1 |
| ARHGEF4 | 2 | 1 | 0 | 1 | 1 |
| CLDN12 | 1 | 1 | 0,058490383 | 0,379264627 | 1 |
| SLC23A2 | 3 | 3 | -3,26736E-16 | 0,999999964 | 1 |
| IGHV1-62-1 | 1 | 1 | 0 | 1 | 1 |
| PRELP | 2 | 2 | 8,73221E-17 | 0,999999983 | 1 |
| USP38 | 2 | 2 | 4,1812E-17 | 0,999999987 | 1 |
| JAK1 | 3 | 3 | 0 | 1 | 1 |
| MRPS36 | 1 | 1 | 0 | 1 | 1 |

| | | | | | |
|---|---|---|---|---|---|
| NSG1 | 1 | 1 | 0,003945713 | 0,855731445 | 1 |
| GRIA1;GRIA2 | 2 | 2 | 0 | 1 | 1 |
| GM11639 | 2 | 2 | 0 | 1 | 1 |
| COMMD1 | 2 | 2 | -6,04561E-17 | 0,999999989 | 1 |
| B4GALNT3 | 1 | 1 | 0 | 1 | 1 |
| IFT122 | 1 | 1 | 0 | 1 | 1 |
| PMVK | 2 | 1 | 0 | 1 | 1 |
| PDSS2 | 1 | 1 | 0 | 1 | 1 |
| MLIP | 2 | 2 | 0 | 1 | 1 |
| DNAAF2 | 1 | 1 | 0 | 1 | 1 |
| KIF13B | 1 | 1 | 0 | 1 | 1 |
| UNC5D | 1 | 1 | 0 | 1 | 1 |
| PLS1;LCP1 | 1 | 1 | 0,067715148 | 0,321138706 | 1 |
| HK2 | 2 | 2 | 1,95334E-14 | 0,999999688 | 1 |
| SLC29A2 | 1 | 1 | 0 | 1 | 1 |
| PHACTR1;PHACTR4 | 1 | 1 | 0 | 1 | 1 |
| WASF3 | 8 | 7 | -2,82689E-16 | 0,999999956 | 1 |
| DVL1 | 2 | 2 | 0 | 1 | 1 |
| CNST | 2 | 2 | -0,115193174 | 0,204305329 | 1 |
| ZFP189 | 1 | 1 | 0 | 1 | 1 |
| CPEB2 | 1 | 1 | -0,040425498 | 0,548692137 | 1 |
| LRRFIP1 | 1 | 1 | 0,069285474 | 0,445550387 | 1 |
| MAP4K4 | 1 | 1 | 0 | 1 | 1 |
| RFC2 | 1 | 1 | 0 | 1 | 1 |
| AGO4;AGO3;AGO1 | 1 | 1 | -0,128206639 | 0,204108681 | 1 |
| CTTNBP2NL | 1 | 1 | 0 | 1 | 1 |
| SMPDL3B | 3 | 3 | 0 | 1 | 1 |
| MAP3K20 | 1 | 1 | 0 | 1 | 1 |
| RAB3GAP1 | 1 | 1 | 0 | 1 | 1 |
| ATP6V0A2;ATP6V0A | 1 | 1 | 0 | 1 | 1 |
| GAB1 | 2 | 2 | 3,33274E-18 | 0,999999998 | 1 |
| CSMD1 | 2 | 2 | 0,106646932 | 0,205271409 | 1 |
| OLFM2 | 2 | 2 | -0,008889381 | 0,768111655 | 1 |

| | | | | | |
|---|---|---|---|---|---|
| HM13 | 4 | 4 | -0,012171057 | 0,655053032 | 1 |
| MORC3 | 1 | 1 | 0 | 1 | 1 |
| MYO5A | 4 | 4 | 0 | 1 | 1 |
| ARFGAP3 | 3 | 3 | 0 | 1 | 1 |
| S1PR1 | 4 | 4 | 1,69795E-17 | 0,99999999 | 1 |
| DCP1A | 1 | 1 | 0 | 1 | 1 |
| VPS26C | 3 | 3 | 0 | 1 | 1 |
| LGI4 | 2 | 2 | 0 | 1 | 1 |
| MRPS18B | 2 | 2 | 0 | 1 | 1 |
| KLC1 | 1 | 1 | 0 | 1 | 1 |
| EMC3 | 5 | 5 | 0 | 1 | 1 |
| CAMK2G;CAMK2B | 2 | 2 | 2,94324E-17 | 0,999999987 | 1 |
| CNOT9 | 3 | 3 | 0 | 1 | 1 |
| ANKRD13A | 2 | 2 | 0 | 1 | 1 |
| KIF21A | 1 | 1 | 0 | 1 | 1 |
| MRPS16 | 2 | 2 | -6,76521E-17 | 0,999999982 | 1 |
| LRIT1 | 1 | 1 | -0,134361707 | 0,276115874 | 1 |
| PRKG1;PRKG2 | 1 | 1 | 0 | 1 | 1 |
| GNPDA2 | 6 | 6 | 0 | 1 | 1 |
| CD2BP2 | 1 | 1 | 0,096813877 | 0,661341745 | 1 |
| TMOD3 | 2 | 1 | 0,33811959 | 0,177471078 | 1 |
| INPP5B | 3 | 3 | 0 | 1 | 1 |
| ZDHHC20 | 1 | 1 | 0 | 1 | 1 |
| PLAAT3 | 2 | 2 | -5,5152E-18 | 0,999999998 | 1 |
| C4BPA | 1 | 1 | 0 | 1 | 1 |
| UBE2B | 1 | 1 | 0 | 1 | 1 |
| LDHC;LDHA | 1 | 1 | 0 | 1 | 1 |
| TPM2 | 1 | 1 | 0 | 1 | 1 |
| MOXD1 | 1 | 1 | 0 | 1 | 1 |
| SERPINB9C;SERPINB9 | 1 | 1 | 0 | 1 | 1 |
| SERPINB8 | 1 | 1 | 0 | 1 | 1 |
| SLC5A3 | 2 | 2 | 0 | 1 | 1 |
| HDHD5 | 5 | 5 | -0,030740027 | 0,517800258 | 1 |

| | | | | | |
|---|---|---|---|---|---|
| CARS2 | 2 | 2 | -8,96512E-18 | 0,999999994 | 1 |
| NCOA1 | 2 | 2 | 1,47382E-12 | 0,999999977 | 1 |
| CACNA1B;CACNA1A | 1 | 1 | 0,070768234 | 0,381709333 | 1 |
| SNX14 | 1 | 1 | 0 | 1 | 1 |
| CD47 | 3 | 3 | 2,86109E-16 | 0,999999965 | 1 |
| HS2ST1 | 2 | 2 | -0,075393354 | 0,32233273 | 1 |
| MOB2 | 1 | 1 | 0 | 1 | 1 |
| NUCB1 | 3 | 3 | 0 | 1 | 1 |
| PCBP3;PCBP2 | 1 | 1 | 0 | 1 | 1 |
| FEZF1 | 1 | 1 | -0,047756857 | 0,571464079 | 1 |
| FCGR1 | 3 | 3 | 0,090386497 | 0,313787709 | 1 |
| MFHAS1 | 1 | 1 | -1,81525E-15 | 0,999999925 | 1 |
| KLHL3 | 2 | 2 | 0,086626316 | 0,17236636 | 1 |
| GDF10 | 1 | 1 | 0 | 1 | 1 |
| NAA30 | 3 | 3 | 0 | 1 | 1 |
| TAF1C | 1 | 1 | 0 | 1 | 1 |
| MRPL44 | 1 | 1 | 0 | 1 | 1 |
| IVNS1ABP | 1 | 1 | 0 | 1 | 1 |
| FAM91A1 | 4 | 4 | 0,073202364 | 0,158970309 | 1 |
| CAMSAP3;CAMSAP2 | 1 | 1 | 0 | 1 | 1 |
| PCNT | 1 | 1 | 0,052865744 | 0,561670605 | 1 |
| CFAP20 | 2 | 2 | 2,94989E-19 | 1 | 1 |
| SESN1 | 1 | 1 | 0 | 1 | 1 |
| MRPL55 | 3 | 3 | -6,69107E-14 | 0,99999936 | 1 |
| WDR73 | 1 | 1 | 0 | 1 | 1 |
| CAMK2G | 2 | 2 | 0,022774159 | 0,609287534 | 1 |
| ZFP386 | 1 | 1 | 0 | 1 | 1 |
| GSTM1;GSTM2 | 1 | 1 | 0 | 1 | 1 |
| UFSP1 | 1 | 1 | 0 | 1 | 1 |
| USP35 | 3 | 3 | -2,48915E-16 | 0,99999998 | 1 |
| DESI1 | 1 | 1 | 0 | 1 | 1 |
| ABLIM2 | 1 | 1 | -4,86063E-19 | 0,999999999 | 1 |
| NPR2 | 1 | 1 | -0,039890134 | 0,539448395 | 1 |

| | | | | | |
|---|---|---|---|---|---|
| NEFL;NEFH;INA | 1 | 1 | 0 | 1 | 1 |
| ATP6V1C2 | 1 | 1 | 0 | 1 | 1 |
| FAM3C | 2 | 2 | -1,44042E-19 | 1 | 1 |
| JPT1 | 3 | 2 | 3,57994E-16 | 0,999999966 | 1 |
| CDC42EP1 | 1 | 1 | 0 | 1 | 1 |
| RTN1 | 3 | 3 | 0,012713033 | 0,682771061 | 1 |
| TENM4;TENM1 | 2 | 2 | 0,002974017 | 0,84047449 | 1 |
| NPTXR;NPCD | 2 | 2 | 0,056581722 | 0,429494331 | 1 |
| DPYSL2;CRMP1 | 3 | 3 | 0 | 1 | 1 |
| FAT3 | 2 | 2 | 0 | 1 | 1 |
| MYZAP | 1 | 1 | -0,071714769 | 0,451888529 | 1 |
| GOLGA1 | 1 | 1 | -0,047440586 | 0,430623953 | 1 |
| GRIK3;GRIK2 | 1 | 1 | 0 | 1 | 1 |
| ATP5MJ | 2 | 2 | -9,26506E-16 | 0,999999926 | 1 |
| TEKT3 | 1 | 1 | 0 | 1 | 1 |
| CDIPT | 6 | 6 | 0,046757615 | 0,298363709 | 1 |
| PHACTR1;RAI14 | 1 | 1 | 0 | 1 | 1 |
| OLFR1234 | 1 | 1 | 0 | 1 | 1 |
| MS4A6D | 1 | 1 | 0 | 1 | 1 |
| HK2;HK1;HK3 | 3 | 3 | 0,130604495 | 0,341942836 | 1 |
| MAP3K7 | 3 | 3 | 0 | 1 | 1 |
| VASP | 1 | 1 | -0,227482513 | 0,146736725 | 1 |
| FAM136A | 2 | 2 | -5,76869E-15 | 0,999999842 | 1 |
| ARF3;ARF1;ARF2 | 7 | 7 | 1,40608E-16 | 0,999999973 | 1 |
| TK2 | 3 | 3 | -0,091785254 | 0,247450193 | 1 |
| STRN;STRN3 | 1 | 1 | -0,094894848 | 0,34235802 | 1 |
| DTL | 1 | 1 | 0 | 1 | 1 |
| STXBP6 | 1 | 1 | 0 | 1 | 1 |
| HIPK3 | 1 | 1 | 0 | 1 | 1 |
| MAPK10 | 4 | 2 | 0 | 1 | 1 |
| DPH5 | 2 | 1 | 0 | 1 | 1 |
| SCG3 | 3 | 3 | -5,90812E-18 | 0,999999995 | 1 |
| RBM8A | 2 | 2 | 0,068128544 | 0,46642924 | 1 |

| | | | | | |
|---|---|---|---|---|---|
| INSR | 3 | 3 | 0 | 1 | 1 |
| TUBB4B;TUBB2A;TU | 4 | 3 | -5,61818E-14 | 0,999999497 | 1 |
| ADCYAP1R1 | 3 | 3 | -0,074601955 | 0,280494631 | 1 |
| ACO2 | 2 | 2 | -0,02038011 | 0,648082303 | 1 |
| CYRIA;CYRIB | 5 | 4 | 0 | 1 | 1 |
| MYLK4 | 1 | 1 | 0 | 1 | 1 |
| KRT14;KRT42;KRT17 | 3 | 2 | 2,23263E-16 | 0,999999987 | 1 |
| H1F10 | 2 | 2 | 7,17367E-18 | 0,999999998 | 1 |
| SEC24A | 1 | 1 | 0 | 1 | 1 |
| LAMP5 | 2 | 2 | 0 | 1 | 1 |
| STX18 | 2 | 2 | 1,17891E-16 | 0,999999994 | 1 |
| GCSH | 3 | 3 | -2,67329E-18 | 0,999999998 | 1 |
| POU3F2 | 1 | 1 | -0,023871629 | 0,674937728 | 1 |
| CCDC113 | 1 | 1 | 0 | 1 | 1 |
| PIR | 4 | 3 | -0,046286984 | 0,399569468 | 1 |
| GNAZ;GNA11;GNAQ | 2 | 2 | 0,032123407 | 0,432837111 | 1 |
| BBOF1 | 1 | 1 | 0 | 1 | 1 |
| MLST8 | 5 | 5 | -7,37343E-18 | 0,999999996 | 1 |
| MLH1 | 1 | 1 | -0,043182418 | 0,51852901 | 1 |
| ROBO1 | 1 | 1 | 0 | 1 | 1 |
| CETN4 | 1 | 1 | -0,12472057 | 0,315659199 | 1 |
| PATL1 | 1 | 1 | 0 | 1 | 1 |
| PLA2R1 | 1 | 1 | 0 | 1 | 1 |
| CFL1;CFL2;DSTN | 1 | 1 | 0 | 1 | 1 |
| YWHAQ;YWHAB | 1 | 1 | 3,5872E-21 | 1 | 1 |
| YWHAG;YWHAZ;YWI | 2 | 1 | 0 | 1 | 1 |
| OLFR1188 | 1 | 1 | 0,226839748 | 0,351129703 | 1 |
| MUC16 | 2 | 1 | 0,210689737 | 0,13572196 | 1 |
| BPIFB9A | 1 | 1 | 0,042162138 | 0,46304311 | 1 |
| UMODL1 | 1 | 1 | 0 | 1 | 1 |
| ZP3R | 1 | 1 | 0 | 1 | 1 |
| ATP12A | 1 | 1 | 0 | 1 | 1 |
| SNX11 | 2 | 2 | -0,080920882 | 0,321897357 | 1 |

| | | | | | |
|---|---|---|---|---|---|
| VPS9D1 | 1 | 1 | 0,05790246 | 0,486340313 | 1 |
| TARS3;TARS1 | 2 | 2 | -1,87996E-16 | 1 | 1 |
| AGO1 | 4 | 4 | 0 | 1 | 1 |
| GRIA3;GRIA4;GRIA2 | 1 | 1 | 0 | 1 | 1 |
| PKIG | 1 | 1 | 0 | 1 | 1 |
| EIF4G2 | 1 | 1 | -0,013210926 | 0,73316466 | 1 |
| ITFG1 | 7 | 7 | 2,11504E-17 | 0,999999984 | 1 |
| ARHGEF10 | 1 | 1 | 0 | 1 | 1 |
| CAPZB | 3 | 3 | 0 | 1 | 1 |
| STAC3 | 2 | 2 | -0,066456768 | 0,340662144 | 1 |
| KCNIP2 | 1 | 1 | 0,168081816 | 0,116973321 | 1 |
| ZMPSTE24 | 4 | 4 | -1,42652E-19 | 1 | 1 |
| PLEKHA5 | 4 | 4 | -1,14597E-16 | 0,999999976 | 1 |
| LPCAT2 | 1 | 1 | 0 | 1 | 1 |
| LPAR1 | 1 | 1 | 0,122846981 | 0,501378113 | 1 |
| ANKS1B | 2 | 2 | 8,48547E-16 | 0,999999967 | 1 |
| CPLX3 | 1 | 1 | 0,075829697 | 0,385391749 | 1 |
| KCNA10;KCNA1;KCN | 1 | 1 | 0 | 1 | 1 |
| U2AF2 | 1 | 1 | 0 | 1 | 1 |
| DERL1 | 1 | 1 | 0,333020646 | 0,132485134 | 1 |
| NEUROD2 | 1 | 1 | 0 | 1 | 1 |
| NEO1 | 2 | 2 | 7,03414E-20 | 1 | 1 |
| TRIM2;TRIM3 | 1 | 1 | 0 | 1 | 1 |
| SGPP1 | 1 | 1 | 0 | 1 | 1 |
| KRT42;KRT17 | 3 | 2 | -8,26157E-17 | 0,999999985 | 1 |
| MMGT1 | 2 | 2 | 0 | 1 | 1 |
| GPM6B | 1 | 1 | 0 | 1 | 1 |
| NPY2R | 2 | 2 | 0 | 1 | 1 |
| RAPH1 | 1 | 1 | -0,112328821 | 0,458663302 | 1 |
| DNLZ | 1 | 1 | 0 | 1 | 1 |
| ZCCHC7 | 1 | 1 | 0 | 1 | 1 |
| PGM5 | 2 | 2 | 0 | 1 | 1 |
| TBC1D23 | 3 | 3 | -0,034439875 | 0,503513276 | 1 |

| | | | | | |
|---|---|---|---|---|---|
| IDH1;IDH2 | 2 | 2 | -0,065322358 | 0,283987281 | 1 |
| TRIM9 | 3 | 3 | 7,16598E-18 | 0,999999994 | 1 |
| IGSF3 | 1 | 1 | -0,078581608 | 0,524063781 | 1 |
| MTMR10 | 1 | 1 | 0 | 1 | 1 |
| RHOT1;RHOT2 | 1 | 1 | 0 | 1 | 1 |
| GPD1 | 2 | 1 | 0 | 1 | 1 |
| CHMP5 | 1 | 1 | 0 | 1 | 1 |
| KRT77;KRT79;KRT5 | 1 | 1 | 0 | 1 | 1 |
| FMNL2;FMNL3;FMN | 1 | 1 | 0 | 1 | 1 |
| ARHGAP30 | 2 | 2 | 2,23177E-18 | 0,999999998 | 1 |
| TNFAIP8L3 | 3 | 3 | 0 | 1 | 1 |
| ARHGAP31 | 2 | 2 | 0 | 1 | 1 |
| ATP7B | 1 | 1 | 0 | 1 | 1 |
| LCLAT1 | 5 | 5 | 0 | 1 | 1 |
| KRT79 | 1 | 1 | 1,4304E-17 | 0,999999996 | 1 |
| PPFIA2 | 1 | 1 | -0,004654979 | 0,842469412 | 1 |
| JCAD | 1 | 1 | -0,081595135 | 0,373890603 | 1 |
| MRPL18 | 1 | 1 | -0,082892573 | 0,375164304 | 1 |
| SMAP2;SMAP1 | 1 | 1 | 0 | 1 | 1 |
| ANKRD28 | 4 | 4 | -4,21949E-15 | 0,999999808 | 1 |
| ZW10 | 2 | 1 | 0 | 1 | 1 |
| LRRC7;ERBIN | 2 | 2 | -1,79087E-16 | 0,999999977 | 1 |
| COX7C | 2 | 2 | -1,55944E-14 | 0,99999981 | 1 |
| INPP4A | 1 | 1 | -2,28685E-16 | 0,999999977 | 1 |
| AMBRA1 | 1 | 1 | 0 | 1 | 1 |
| SLC7A2 | 4 | 4 | -2,75893E-18 | 0,999999997 | 1 |
| LARP4B | 1 | 1 | 0,266869495 | 0,122657128 | 1 |
| FBL | 1 | 1 | 0,116115303 | 0,212850434 | 1 |
| ATL2 | 4 | 4 | -1,36228E-16 | 0,999999966 | 1 |
| ENSA | 2 | 2 | -2,47436E-17 | 0,999999995 | 1 |
| PALS1 | 2 | 2 | 0,067629709 | 0,249971778 | 1 |
| DTX4 | 1 | 1 | 0 | 1 | 1 |
| CNTD1 | 1 | 1 | -0,027864823 | 0,588787002 | 1 |

| | | | | | |
|---|---|---|---|---|---|
| TUBB4B;TUBB5;TUBI | 4 | 4 | 8,10278E-18 | 0,999999995 | 1 |
| PSME4 | 2 | 2 | -7,61272E-19 | 1 | 1 |
| FMO3 | 1 | 1 | 0 | 1 | 1 |
| FAN1 | 2 | 2 | -1,69406E-19 | 1 | 1 |
| KHDC1A;KHDC1C | 1 | 1 | 0,010051983 | 0,751459644 | 1 |
| STK32B | 1 | 1 | 0 | 1 | 1 |
| STK26 | 1 | 1 | 0 | 1 | 1 |
| CDYL | 1 | 1 | 0,085653546 | 0,301099781 | 1 |
| CHMP1B2 | 1 | 1 | 0 | 1 | 1 |
| CERS6 | 2 | 2 | -8,11829E-18 | 0,999999996 | 1 |
| CHMP1B1 | 1 | 1 | 0 | 1 | 1 |
| PTPN6 | 2 | 2 | 4,55367E-18 | 0,999999998 | 1 |
| PEX5L | 3 | 3 | -9,53514E-20 | 1 | 1 |
| AQP1 | 2 | 1 | 0,145349609 | 0,407157764 | 1 |
| PECR | 1 | 1 | 0 | 1 | 1 |
| MOCS2 | 2 | 1 | 0,061741862 | 0,411007022 | 1 |
| KCNA1;KCNA3;KCNA | 1 | 1 | 0 | 1 | 1 |
| LRRFIP2 | 1 | 1 | 0 | 1 | 1 |
| RABIF | 3 | 3 | -3,13691E-18 | 0,999999997 | 1 |
| SLC24A2 | 4 | 4 | 0 | 1 | 1 |
| MOV10L1 | 1 | 1 | 0 | 1 | 1 |
| ATAD5 | 1 | 1 | 0,145126056 | 0,162425327 | 1 |
| SCPEP1 | 1 | 1 | 0,027525546 | 0,721074375 | 1 |
| UBXN4 | 1 | 1 | 0 | 1 | 1 |
| TMSB15B1 | 1 | 1 | 0 | 1 | 1 |
| CC2D1A | 1 | 1 | 0 | 1 | 1 |
| DDX20 | 1 | 1 | 4,40456E-20 | 1 | 1 |
| PHOSPHO2 | 1 | 1 | 0,188326532 | 0,16233203 | 1 |
| AMT | 2 | 2 | -0,042530719 | 0,568877715 | 1 |
| CDC42 | 2 | 2 | 0 | 1 | 1 |
| CDC42 | 2 | 2 | 0,030444569 | 0,50519461 | 1 |
| EVA1A | 1 | 1 | -0,087098271 | 0,146046494 | 1 |
| EFCAB9 | 1 | 1 | 0 | 1 | 1 |

| | | | | | |
|---|---|---|---|---|---|
| MRPL23 | 4 | 4 | 0 | 1 | 1 |
| CTSZ | 2 | 1 | -0,067161798 | 0,29189703 | 1 |
| DCX;DCLK1 | 1 | 1 | 0 | 1 | 1 |
| DNM1L | 3 | 3 | 0 | 1 | 1 |
| CTSC | 1 | 1 | 0 | 1 | 1 |
| PPP1CC | 2 | 2 | -0,007294388 | 0,694636529 | 1 |
| TRIM9 | 1 | 1 | 0 | 1 | 1 |
| SAT2 | 1 | 1 | 0 | 1 | 1 |
| EPB41L2;EPB41L3 | 1 | 1 | -2,15726E-17 | 0,999999991 | 1 |
| SINHCAF | 1 | 1 | 0 | 1 | 1 |
| GRB10;GRB14 | 1 | 1 | 0 | 1 | 1 |
| RNF31 | 3 | 3 | -1,1583E-18 | 1 | 1 |
| GRIN1 | 1 | 1 | 0 | 1 | 1 |
| sp\|P0C913\|OCC1_M | 1 | 1 | 0,179793367 | 0,163598793 | 1 |
| NRXN2;NRXN1 | 3 | 3 | -1,19977E-15 | 0,999999912 | 1 |
| D6WSU163E | 2 | 2 | -2,66046E-18 | 0,999999999 | 1 |
| RUFY1;RUFY3 | 1 | 1 | 0 | 1 | 1 |
| MSANTD4 | 2 | 2 | 0 | 1 | 1 |
| SEC23B | 1 | 1 | 0,137354138 | 0,134761317 | 1 |
| IKZF2 | 1 | 1 | 0 | 1 | 1 |
| CRTC1 | 9 | 8 | 3,40752E-17 | 0,999999985 | 1 |
| ZC3H12B | 1 | 1 | 0 | 1 | 1 |
| SYN1;SYN2 | 7 | 7 | 0,031720286 | 0,513573624 | 1 |
| OAF | 1 | 1 | 0 | 1 | 1 |
| GZMK | 1 | 1 | 0 | 1 | 1 |
| FOXL2 | 1 | 1 | 0 | 1 | 1 |
| ZHX1 | 1 | 1 | -0,190346632 | 0,128159977 | 1 |
| ADAMTS4 | 1 | 1 | 0 | 1 | 1 |
| TMPO | 3 | 3 | -1,01428E-17 | 0,999999997 | 1 |
| PINX1 | 1 | 1 | 0 | 1 | 1 |
| CMYA5 | 1 | 1 | 0 | 1 | 1 |
| POU4F3 | 1 | 1 | 0 | 1 | 1 |
| ZNF609 | 1 | 1 | 0 | 1 | 1 |

| | | | | | |
|---|---|---|---|---|---|
| FER1L6 | 1 | 1 | 0 | 1 | 1 |
| SOX15;SOX16 | 1 | 1 | 0 | 1 | 1 |
| EGF | 1 | 1 | 0 | 1 | 1 |
| SLC25A44 | 1 | 1 | -0,014102643 | 0,758407367 | 1 |
| COX17 | 1 | 1 | -0,073529025 | 0,364517773 | 1 |
| RELCH | 1 | 1 | 0 | 1 | 1 |
| ECI3 | 1 | 1 | 0 | 1 | 1 |
| KDM4B | 1 | 1 | 0 | 1 | 1 |
| LATS2 | 1 | 1 | 0 | 1 | 1 |
| ARHGAP6 | 1 | 1 | -0,07295039 | 0,391070082 | 1 |
| RANBP17;XPO7 | 1 | 1 | 0 | 1 | 1 |
| DGKI;DGKZ | 2 | 2 | 0 | 1 | 1 |
| BAIAP2 | 3 | 3 | -4,79538E-17 | 0,999999983 | 1 |
| GAL3ST4 | 1 | 1 | 0 | 1 | 1 |
| FMNL3 | 1 | 1 | 0,022404147 | 0,599272391 | 1 |
| SH2D7 | 1 | 1 | 0 | 1 | 1 |
| ACVRL1 | 1 | 1 | 0 | 1 | 1 |
| TMEM134 | 1 | 1 | 0 | 1 | 1 |
| LPP | 1 | 1 | 0 | 1 | 1 |
| CHRNB1 | 1 | 1 | -0,051622074 | 0,418764534 | 1 |
| MPLKIP | 1 | 1 | 0,083606984 | 0,388593308 | 1 |
| SYN1 | 3 | 3 | 1,15216E-18 | 0,999999999 | 1 |
| PARP1 | 1 | 1 | 0 | 1 | 1 |
| OOSP2 | 1 | 1 | -0,092693975 | 0,213258592 | 1 |
| MORC2A | 1 | 1 | 0 | 1 | 1 |
| CHD1 | 1 | 1 | -0,037932211 | 0,476646336 | 1 |
| PCBP2 | 1 | 1 | 0 | 1 | 1 |
| PCBP2 | 1 | 1 | 1,39624E-17 | 0,999999992 | 1 |
| COL6A3 | 1 | 1 | 0 | 1 | 1 |
| NCKAP5 | 1 | 1 | -0,052710925 | 0,519251457 | 1 |
| NECAP2 | 1 | 1 | 0,140555658 | 0,507492621 | 1 |
| TONSL | 2 | 2 | 0 | 1 | 1 |
| MYO1B | 1 | 1 | -0,061175513 | 0,622843092 | 1 |

| | | | | | |
|---|---|---|---|---|---|
| BSN;PCLO | 3 | 3 | 6,40631E-20 | 1 | 1 |
| ADNP | 1 | 1 | 0 | 1 | 1 |
| KLC1;KLC2 | 4 | 4 | 0 | 1 | 1 |
| CPLX1;CPLX2 | 5 | 5 | 3,89681E-05 | 0,981154591 | 1 |
| RECQL4 | 1 | 1 | 0 | 1 | 1 |
| SNX25 | 2 | 2 | -0,131266217 | 0,139069523 | 1 |
| DRGX | 1 | 1 | 0 | 1 | 1 |
| EPB41L2 | 3 | 3 | -0,044913829 | 0,353129715 | 1 |
| DBI | 4 | 4 | -3,48764E-20 | 1 | 1 |
| DGKB | 1 | 1 | 0 | 1 | 1 |
| TMCC1;TMCC2 | 1 | 1 | 0 | 1 | 1 |
| 4930402K13RIK | 1 | 1 | 0 | 1 | 1 |
| CAMK2A;CAMK2D | 2 | 2 | 0,154635893 | 0,32585194 | 1 |
| MACF1 | 2 | 2 | 0 | 1 | 1 |
| STXBP5L | 1 | 1 | 1,12468E-19 | 1 | 1 |
| MATN4 | 5 | 5 | 0 | 1 | 1 |
| NUDT14 | 3 | 2 | -1,16621E-17 | 0,999999995 | 1 |
| SNX3;SNX12 | 2 | 2 | 1,853E-18 | 0,999999998 | 1 |
| COL24A1 | 1 | 1 | 0 | 1 | 1 |
| TRUB1 | 2 | 2 | 3,18171E-18 | 0,999999998 | 1 |
| MAPRE3 | 1 | 1 | 0,058884268 | 0,349159218 | 1 |
| SNCB;SNCG | 2 | 2 | 9,52257E-19 | 0,999999999 | 1 |
| RNF157 | 1 | 1 | 0 | 1 | 1 |
| MAP3K7CL | 1 | 1 | 0 | 1 | 1 |
| TMEM129 | 1 | 1 | 1,43476E-18 | 0,999999998 | 1 |
| CDR2 | 1 | 1 | 0 | 1 | 1 |
| MANF | 2 | 2 | 0 | 1 | 1 |
| TRIM33 | 1 | 1 | 0 | 1 | 1 |
| MYO5C | 1 | 1 | 0 | 1 | 1 |
| SGSM2;SGSM1 | 1 | 1 | -0,145972036 | 0,123746513 | 1 |
| WWC2 | 2 | 2 | 0,078892013 | 0,337122091 | 1 |
| ASIC1 | 1 | 1 | 0 | 1 | 1 |
| STX1A;STX1B | 1 | 1 | 0 | 1 | 1 |

| | | | | | |
|---|---|---|---|---|---|
| ACTN4 | 2 | 2 | 0 | 1 | 1 |
| MAK | 2 | 2 | 0,051231597 | 0,325415992 | 1 |
| BCKDK | 4 | 4 | -1,22699E-17 | 0,999999993 | 1 |
| CYTH4 | 2 | 2 | -3,92419E-18 | 0,999999999 | 1 |
| CLN6 | 1 | 1 | 0 | 1 | 1 |
| CDH2;CDH4 | 2 | 2 | 0,060879603 | 0,222915249 | 1 |
| NRBF2 | 2 | 2 | 0 | 1 | 1 |
| RBM15B | 1 | 1 | 0 | 1 | 1 |
| SNX3 | 1 | 1 | 0 | 1 | 1 |
| SNX12 | 1 | 1 | 0,063307664 | 0,366993574 | 1 |
| RABEPK | 3 | 1 | 1,72501E-17 | 0,999999991 | 1 |
| RIPOR3 | 1 | 1 | 0 | 1 | 1 |
| KRT34 | 1 | 1 | 0,105246211 | 0,222827867 | 1 |
| ACTG1 | 1 | 1 | 0 | 1 | 1 |
| TRPC6 | 1 | 1 | 0 | 1 | 1 |
| TMEFF1 | 2 | 2 | 0 | 1 | 1 |
| RPL39 | 1 | 1 | -0,168333331 | 0,242786877 | 1 |
| DLGAP1 | 1 | 1 | 0 | 1 | 1 |
| PCDHGC5;PCDHGA9; | 2 | 2 | 0 | 1 | 1 |
| QRICH2 | 1 | 1 | 0 | 1 | 1 |
| ARAF | 1 | 1 | 0 | 1 | 1 |
| KSR2 | 2 | 2 | -3,36412E-16 | 0,999999972 | 1 |
| SNU13 | 1 | 1 | 0,299676153 | 0,156075649 | 1 |
| GCNT3 | 1 | 1 | 0 | 1 | 1 |
| CUBN | 1 | 1 | -2,41806E-20 | 1 | 1 |
| SSR3 | 1 | 1 | 0 | 1 | 1 |
| ARFGAP1 | 1 | 1 | 0 | 1 | 1 |
| MMTAG2 | 2 | 2 | 0 | 1 | 1 |
| MYH10;MYH14 | 2 | 2 | -1,82239E-17 | 0,99999999 | 1 |
| TGM4 | 1 | 1 | 0 | 1 | 1 |
| SCAF4 | 1 | 1 | 0 | 1 | 1 |
| EEF1D | 1 | 1 | 0 | 1 | 1 |
| CLASP1 | 2 | 2 | 3,75725E-16 | 0,999999975 | 1 |

| | | | | | |
|---|---|---|---|---|---|
| KRT10;KRT13 | 2 | 2 | 0,293085603 | 0,487454995 | 1 |
| TIA1;TIAL1 | 3 | 2 | 0,128717783 | 0,194406801 | 1 |
| TMEM186 | 1 | 1 | 0 | 1 | 1 |
| PPP3CA | 1 | 1 | 0 | 1 | 1 |
| PPP3CA | 2 | 2 | 1,12883E-17 | 0,999999991 | 1 |
| NSD2 | 1 | 1 | 0 | 1 | 1 |
| KRT12 | 2 | 1 | 0 | 1 | 1 |
| MYO10 | 1 | 1 | 0 | 1 | 1 |
| YARS1 | 2 | 1 | 0,225271179 | 0,335972687 | 1 |
| RHOB;RHOC;RHOA | 1 | 1 | 0,101097984 | 0,222894475 | 1 |
| HCN2;HCN1;HCN3 | 1 | 1 | 0 | 1 | 1 |
| TEX28 | 1 | 1 | 0,050803371 | 0,424052893 | 1 |
| TLE5 | 1 | 1 | 0 | 1 | 1 |
| KLC3 | 1 | 1 | 0,288648536 | 0,246160619 | 1 |
| TRIML1 | 1 | 1 | 0 | 1 | 1 |
| KCNA1;KCNA3;KCNA | 1 | 1 | 0 | 1 | 1 |
| SURF4 | 2 | 1 | 2,11911E-15 | 0,999999904 | 1 |
| CHCHD2 | 1 | 1 | -0,130918643 | 0,134859586 | 1 |
| MYH3 | 1 | 1 | 0 | 1 | 1 |
| NGLY1 | 1 | 1 | 0 | 1 | 1 |
| CCDC150 | 1 | 1 | 0,244967959 | 0,18621698 | 1 |
| CHD7;CHD9 | 1 | 1 | 0 | 1 | 1 |
| LRRC75A | 1 | 1 | 0,104397027 | 0,166728585 | 1 |
| HIPK2 | 1 | 1 | 0 | 1 | 1 |
| CSRNP1 | 1 | 1 | 0 | 1 | 1 |
| DENND11 | 3 | 3 | 0 | 1 | 1 |
| EBAG9 | 1 | 1 | 0 | 1 | 1 |
| CC2D1B | 1 | 1 | -0,118250399 | 0,227322361 | 1 |
| PURB;PURA;PURG | 1 | 1 | 0 | 1 | 1 |
| SELENOW | 1 | 1 | 0,065973764 | 0,500108897 | 1 |
| ABCD2 | 2 | 2 | 2,00019E-18 | 0,999999999 | 1 |
| L1TD1 | 1 | 1 | 0 | 1 | 1 |
| GALNT16 | 3 | 3 | 3,61788E-16 | 0,999999985 | 1 |

| | | | | | |
|---|---|---|---|---|---|
| ENPP2 | 1 | 1 | 0 | 1 | 1 |
| TMPRSS11B | 1 | 1 | 0,048096862 | 0,598236015 | 1 |
| MFAP3;MFAP3L | 1 | 1 | 0 | 1 | 1 |
| TAF1 | 1 | 1 | 0 | 1 | 1 |
| PHKA2 | 2 | 2 | -1,62891E-14 | 0,999999711 | 1 |
| LRRC4B;LRRC4C | 1 | 1 | 0,050450663 | 0,382599806 | 1 |
| SARAF | 1 | 1 | 0 | 1 | 1 |
| TEKT2 | 1 | 1 | 0 | 1 | 1 |
| RBFOX2 | 1 | 1 | 0 | 1 | 1 |
| ADSS1;ADSS2 | 5 | 4 | -1,73401E-14 | 0,999999777 | 1 |
| SYNGAP1 | 1 | 1 | 0 | 1 | 1 |
| HLTF | 2 | 2 | -6,08206E-17 | 0,999999982 | 1 |
| SERPINA16 | 1 | 1 | 0,027603849 | 0,615744691 | 1 |
| CUX1 | 1 | 1 | -2,29442E-17 | 0,99999999 | 1 |
| AGAP3 | 1 | 1 | 3,25291E-18 | 0,999999996 | 1 |
| CCM2 | 2 | 2 | -4,84292E-17 | 0,999999986 | 1 |
| UBE2J1 | 1 | 1 | 0 | 1 | 1 |
| CACNB1 | 1 | 1 | 0,133299893 | 0,223813778 | 1 |
| QTRT2 | 2 | 1 | 0,19128623 | 0,125298355 | 1 |
| RECQL | 1 | 1 | 0 | 1 | 1 |
| DCAF8 | 2 | 1 | 0 | 1 | 1 |
| NIN | 1 | 1 | 0,196368751 | 0,187181771 | 1 |
| MEIS3 | 1 | 1 | -0,034228787 | 0,568787854 | 1 |
| HSPA12A;HSPA12B | 2 | 2 | -3,07053E-19 | 1 | 1 |
| RALGAPA2 | 1 | 1 | 0 | 1 | 1 |
| CACNB2;CACNB4;CA | 4 | 4 | 6,82895E-24 | 1 | 1 |
| PTMS | 4 | 3 | 7,20694E-19 | 0,999999999 | 1 |
| ADAMTS17 | 1 | 1 | 0 | 1 | 1 |
| KRT1 | 34 | 5 | 4,13686E-15 | 0,999999928 | 1 |
| TMC7 | 1 | 1 | -0,060522005 | 0,430552462 | 1 |
| SMTNL2 | 1 | 1 | 0 | 1 | 1 |
| SMAD1;SMAD5 | 2 | 2 | -4,07581E-19 | 0,999999999 | 1 |
| LDAH | 1 | 1 | -0,10550375 | 0,375594508 | 1 |

| | | | | | |
|---|---|---|---|---|---|
| GOLT1B | 2 | 2 | -2,90461E-16 | 0,999999991 | 1 |
| ALDH1A1;ALDH1A2;/ | 1 | 1 | 0 | 1 | 1 |
| CHCHD3 | 2 | 2 | 0 | 1 | 1 |
| SNX32 | 3 | 3 | 0 | 1 | 1 |
| PJA1 | 1 | 1 | 0,122110634 | 0,226606862 | 1 |
| DNAJC19 | 1 | 1 | 0 | 1 | 1 |
| SNX7 | 1 | 1 | -2,41294E-17 | 0,999999988 | 1 |
| ZC3H14 | 1 | 1 | 0 | 1 | 1 |
| MAL2 | 2 | 2 | 0 | 1 | 1 |
| HCN3 | 1 | 1 | 0,07519899 | 0,401505264 | 1 |
| KEAP1 | 1 | 1 | 0,016376704 | 0,796871924 | 1 |
| RAP2A;RAP2C | 1 | 1 | 0,095606315 | 0,265111886 | 1 |
| SST | 1 | 1 | 0,13028155 | 0,214034799 | 1 |
| SPATA2 | 3 | 3 | 0 | 1 | 1 |
| ENTPD6 | 1 | 1 | 0 | 1 | 1 |
| LPP | 1 | 1 | 0 | 1 | 1 |
| VPS37A | 1 | 1 | 0 | 1 | 1 |
| FOXO4 | 1 | 1 | 0 | 1 | 1 |
| ATP2B3 | 2 | 2 | 0 | 1 | 1 |
| DMAC1 | 1 | 1 | 0 | 1 | 1 |
| SFN | 14 | 3 | 3,41867E-14 | 0,999999637 | 1 |
| RTN3 | 4 | 4 | 4,9709E-16 | 0,999999936 | 1 |
| GOLPH3 | 1 | 1 | 0 | 1 | 1 |
| NTAQ1 | 1 | 1 | 0 | 1 | 1 |
| XKR7 | 1 | 1 | 0 | 1 | 1 |
| DAAM2;DAAM1 | 3 | 3 | 0,039770955 | 0,36219729 | 1 |
| FER | 2 | 2 | 1,56694E-12 | 0,999998829 | 1 |
| KCNF1 | 1 | 1 | 0 | 1 | 1 |
| FAM107B | 1 | 1 | 0 | 1 | 1 |
| LEXM | 1 | 1 | 0 | 1 | 1 |
| FAM184B | 1 | 1 | 2,47105E-17 | 0,999999989 | 1 |
| CTNND1 | 1 | 1 | 0 | 1 | 1 |
| SLC39A11 | 1 | 1 | 0 | 1 | 1 |

| | | | | | |
|---|---|---|---|---|---|
| PPCDC | 2 | 2 | 9,41482E-19 | 0,999999999 | 1 |
| NACAD | 1 | 1 | 0 | 1 | 1 |
| PIK3IP1 | 1 | 1 | 0,037682603 | 0,589319803 | 1 |
| PTK6 | 1 | 1 | 0 | 1 | 1 |
| RB1 | 2 | 2 | -1,30111E-15 | 0,99999992 | 1 |
| GRAMD4 | 1 | 1 | 0 | 1 | 1 |
| ADAMTS16 | 1 | 1 | 0 | 1 | 1 |
| ATG14 | 1 | 1 | 0 | 1 | 1 |
| RASSF2 | 2 | 2 | 6,87436E-19 | 0,999999999 | 1 |
| TMEM132E | 2 | 2 | 0,04009373 | 0,381969125 | 1 |
| PDLIM5 | 1 | 1 | 0 | 1 | 1 |
| AP3S1 | 1 | 1 | 0 | 1 | 1 |
| DCTN5 | 1 | 1 | 0 | 1 | 1 |
| NRAS | 1 | 1 | 0,019885482 | 0,616617597 | 1 |
| HRAS | 3 | 3 | 1,21633E-19 | 1 | 1 |
| LSM14B | 1 | 1 | 0 | 1 | 1 |
| RNPEPL1 | 1 | 1 | 0,04776679 | 0,575289824 | 1 |
| TAB1 | 3 | 3 | -5,06461E-17 | 0,999999986 | 1 |
| INPP5K | 2 | 2 | 0,057594924 | 0,356954786 | 1 |
| GM21876 | 1 | 1 | 0,067922437 | 0,515342629 | 1 |
| CD63 | 1 | 1 | 0 | 1 | 1 |
| OSBPL11 | 2 | 2 | 1,29258E-15 | 0,999999943 | 1 |
| INPP5E | 1 | 1 | 0 | 1 | 1 |
| PHACTR4 | 1 | 1 | 0 | 1 | 1 |
| SMDT1 | 1 | 1 | 0 | 1 | 1 |
| PTPRD | 1 | 1 | 0,139577489 | 0,150651728 | 1 |
| CDIP1 | 1 | 1 | 0 | 1 | 1 |
| TMOD4 | 1 | 1 | 0 | 1 | 1 |
| SLCO3A1 | 1 | 1 | 0 | 1 | 1 |
| SRP19 | 1 | 1 | 0 | 1 | 1 |
| PRRC1 | 1 | 1 | 0 | 1 | 1 |
| 6330409D20RIK | 1 | 1 | 0 | 1 | 1 |
| NR1H4 | 1 | 1 | 0 | 1 | 1 |

| | | | | | |
|---|---|---|---|---|---|
| CMTM6 | 1 | 1 | 0 | 1 | 1 |
| CROT | 1 | 1 | 0 | 1 | 1 |
| NASP | 2 | 1 | 0 | 1 | 1 |
| HEPH | 1 | 1 | -0,109527935 | 0,241655199 | 1 |
| EGLN1 | 2 | 2 | -1,16766E-16 | 0,999999982 | 1 |
| CSMD3 | 1 | 1 | 2,53917E-22 | 1 | 1 |
| CASP3 | 1 | 1 | 0 | 1 | 1 |
| ATP6V0C | 4 | 4 | 0,084884591 | 0,547427984 | 1 |
| PABPC4L;PABPC4;PA | 1 | 1 | -0,142574918 | 0,376094214 | 1 |
| SLC12A3 | 1 | 1 | 0,058233087 | 0,331817378 | 1 |
| ITSN1 | 1 | 1 | 0 | 1 | 1 |
| SLC5A6 | 1 | 1 | 0,143957276 | 0,184321145 | 1 |
| SLC38A2 | 1 | 1 | 0 | 1 | 1 |
| OLFM3 | 1 | 1 | -0,010985404 | 0,770589311 | 1 |
| AGBL1 | 1 | 1 | 0 | 1 | 1 |
| DNER | 1 | 1 | 0,078931694 | 0,290428041 | 1 |
| LEKR1 | 1 | 1 | 0 | 1 | 1 |
| CEBPZOS | 1 | 1 | 0 | 1 | 1 |
| SYNGAP1 | 1 | 1 | -0,019749927 | 0,676012546 | 1 |
| RYR3;RYR1;RYR2 | 3 | 3 | -0,020823243 | 0,684141019 | 1 |
| GARNL3 | 1 | 1 | -0,088796502 | 0,210707046 | 1 |
| CHCHD1 | 1 | 1 | -0,146237818 | 0,25643305 | 1 |
| AASDHPPT | 1 | 1 | 0 | 1 | 1 |
| SLC6A4 | 1 | 1 | 3,11748E-19 | 0,999999999 | 1 |
| RGSL1 | 1 | 1 | 0 | 1 | 1 |
| SYTL5 | 1 | 1 | 0 | 1 | 1 |
| CCNYL1 | 2 | 2 | -0,030719657 | 0,509340544 | 1 |
| BET1L | 1 | 1 | 0 | 1 | 1 |
| PKN1 | 1 | 1 | 0 | 1 | 1 |
| ARPP21 | 3 | 3 | 0 | 1 | 1 |
| CTNND1 | 1 | 1 | 0 | 1 | 1 |
| SNX13 | 1 | 1 | 0 | 1 | 1 |
| TPM3;TPM1 | 1 | 1 | 0 | 1 | 1 |

| | | | | | |
|---|---|---|---|---|---|
| OSBPL7 | 1 | 1 | 2,71448E-19 | 1 | 1 |
| SCMH1 | 1 | 1 | 0 | 1 | 1 |
| IL19 | 1 | 1 | -4,28189E-19 | 0,999999999 | 1 |
| FHIP2B | 1 | 1 | 0 | 1 | 1 |
| DACT3 | 1 | 1 | 0 | 1 | 1 |
| ATP11C | 1 | 1 | 0 | 1 | 1 |
| CYP2D10 | 1 | 1 | 0 | 1 | 1 |
| PHKG2 | 1 | 1 | 0 | 1 | 1 |
| SPOCK1 | 1 | 1 | -0,016779706 | 0,655114495 | 1 |
| OVOS | 1 | 1 | 0 | 1 | 1 |
| TTC23L | 1 | 1 | 0 | 1 | 1 |
| FRMPD4 | 3 | 3 | 0 | 1 | 1 |
| KRT78 | 5 | 4 | 0,172228767 | 0,477416102 | 1 |
| PTPRR | 3 | 3 | 0 | 1 | 1 |
| MVK | 5 | 5 | 0,047595061 | 0,371850903 | 1 |
| UBE2R2 | 2 | 1 | 0 | 1 | 1 |
| ARL14EP | 1 | 1 | 0 | 1 | 1 |
| ARIH2 | 4 | 4 | 0 | 1 | 1 |
| PLEKHA6 | 2 | 2 | -0,026884055 | 0,50669001 | 1 |
| BET1 | 1 | 1 | 0 | 1 | 1 |
| SRA1 | 1 | 1 | 0 | 1 | 1 |
| SFXN4 | 1 | 1 | 0 | 1 | 1 |
| GM2A | 2 | 2 | 0 | 1 | 1 |
| CCS | 2 | 2 | 3,40398E-18 | 0,999999997 | 1 |
| FBXL12 | 1 | 1 | -0,111128537 | 0,368748473 | 1 |
| GLI3 | 1 | 1 | 0 | 1 | 1 |
| ADD1 | 1 | 1 | 0 | 1 | 1 |
| SMPDL3A | 1 | 1 | 0 | 1 | 1 |
| TUBGCP5 | 1 | 1 | 0 | 1 | 1 |
| TGS1 | 1 | 1 | -0,061494885 | 0,517039614 | 1 |
| MYH11;MYH14 | 1 | 1 | 0 | 1 | 1 |
| POGLUT2 | 1 | 1 | 0 | 1 | 1 |
| BAG1 | 1 | 1 | -0,105407598 | 0,211461075 | 1 |

| | | | | | |
|---|---|---|---|---|---|
| ANKRD50 | 1 | 1 | 0 | 1 | 1 |
| 1700009N14RIK | 1 | 1 | 0 | 1 | 1 |
| DHX29 | 2 | 2 | -0,10379614 | 0,177540878 | 1 |
| MBP | 2 | 2 | -5,62621E-16 | 0,999999964 | 1 |
| SMARCD1 | 1 | 1 | 0 | 1 | 1 |
| PAQR9 | 1 | 1 | 0 | 1 | 1 |
| RETREG3 | 2 | 2 | -0,104568326 | 0,176616194 | 1 |
| SYNDIG1 | 1 | 1 | 0 | 1 | 1 |
| LRBA | 1 | 1 | -0,022790977 | 0,696552509 | 1 |
| CCDC141 | 1 | 1 | 0,056705381 | 0,44199326 | 1 |
| TNFRSF13B | 1 | 1 | 0 | 1 | 1 |
| FAM234A | 1 | 1 | -6,67834E-19 | 0,999999999 | 1 |
| AAGAB | 2 | 2 | 1,89982E-16 | 0,99999997 | 1 |
| CFAP99 | 1 | 1 | 0 | 1 | 1 |
| PTBP3 | 2 | 2 | 9,67E-17 | 0,999999991 | 1 |
| AFTPH | 3 | 3 | -3,44095E-17 | 0,999999991 | 1 |
| IFT27 | 1 | 1 | 0 | 1 | 1 |
| HYDIN | 1 | 1 | 0 | 1 | 1 |
| TRIM15 | 1 | 1 | 0 | 1 | 1 |
| SH3D21 | 1 | 1 | -0,060381324 | 0,286047945 | 1 |
| CX3CR1 | 1 | 1 | 0 | 1 | 1 |
| ZFP646 | 1 | 1 | 0 | 1 | 1 |
| AMOT | 1 | 1 | 0 | 1 | 1 |
| FRS3 | 1 | 1 | 0 | 1 | 1 |
| STRN3 | 2 | 2 | 0 | 1 | 1 |
| LAMA3 | 2 | 2 | -3,63187E-17 | 1 | 1 |
| CSNK1G2 | 1 | 1 | 0 | 1 | 1 |
| INF2 | 2 | 2 | 3,46041E-19 | 1 | 1 |
| TRMT10C | 3 | 3 | 0 | 1 | 1 |
| CYFIP1 | 2 | 1 | 0,07735197 | 0,187421302 | 1 |
| GAS7 | 1 | 1 | 0,386149981 | 0,152390935 | 1 |
| ZFYVE27 | 1 | 1 | 0 | 1 | 1 |
| SYNJ1 | 1 | 1 | 0 | 1 | 1 |

| | | | | | |
|---|---|---|---|---|---|
| SYNJ1 | 1 | 1 | 0 | 1 | 1 |
| TYRO3 | 1 | 1 | -0,051665577 | 0,508846875 | 1 |
| TLR5 | 1 | 1 | -0,138675591 | 0,228393627 | 1 |
| CAMK2A | 2 | 2 | 2,19915E-16 | 0,999999962 | 1 |
| NAV3 | 2 | 2 | 0 | 1 | 1 |
| PKIA | 1 | 1 | 0 | 1 | 1 |
| TMEM200A | 1 | 1 | 0 | 1 | 1 |
| FHIP2A | 1 | 1 | 0 | 1 | 1 |
| PCDHGA6 | 1 | 1 | 0 | 1 | 1 |
| ARHGEF6;ARHGEF7 | 2 | 2 | 3,25422E-16 | 0,999999947 | 1 |
| YTHDC2 | 1 | 1 | 0 | 1 | 1 |
| LYZ1 | 1 | 1 | 0 | 1 | 1 |
| CCDC28A | 1 | 1 | 0 | 1 | 1 |
| HOXA13 | 1 | 1 | 0,104678791 | 0,327548836 | 1 |
| POLR2M | 1 | 1 | 0 | 1 | 1 |
| CLPTM1L | 3 | 3 | -0,002498809 | 0,849688017 | 1 |
| FEM1B | 1 | 1 | 0 | 1 | 1 |
| RNF146 | 1 | 1 | 0 | 1 | 1 |
| ANKS1B | 1 | 1 | 0 | 1 | 1 |
| DMAP1 | 1 | 1 | 0 | 1 | 1 |
| FASTKD2 | 1 | 1 | 0 | 1 | 1 |
| CDC123 | 1 | 1 | 0 | 1 | 1 |
| GNAT2 | 1 | 1 | 0 | 1 | 1 |
| SEPTIN7 | 3 | 2 | 3,40549E-11 | 0,999989693 | 1 |
| TRIM45 | 1 | 1 | 0 | 1 | 1 |
| AKAP2 | 3 | 3 | -4,76742E-18 | 0,999999999 | 1 |
| SUSD5 | 1 | 1 | -0,154998151 | 0,369627187 | 1 |
| PPP1R16A | 1 | 1 | 0 | 1 | 1 |
| TRAK2 | 1 | 1 | 0 | 1 | 1 |
| B3GNT7 | 1 | 1 | 0 | 1 | 1 |
| GPR17 | 1 | 1 | 0 | 1 | 1 |
| PTK2 | 1 | 1 | -0,023402204 | 0,604612316 | 1 |
| SNX9 | 1 | 1 | 0,178095986 | 0,137639012 | 1 |

| | | | | | |
|---|---|---|---|---|---|
| MDM1 | 1 | 1 | -0,148012537 | 0,328818655 | 1 |
| TSPAN14 | 1 | 1 | 0 | 1 | 1 |
| TENM4;TENM2;TENI | 1 | 1 | 0 | 1 | 1 |
| MSI1 | 1 | 1 | 0 | 1 | 1 |
| DPH2 | 1 | 1 | 0 | 1 | 1 |
| PLEKHB1 | 3 | 3 | -2,71499E-15 | 0,999999892 | 1 |
| SDF4 | 2 | 2 | 4,59312E-19 | 0,999999999 | 1 |
| MAPK10;MAPK8 | 5 | 4 | 1,38049E-16 | 0,999999965 | 1 |
| DCX;DCLK2 | 1 | 1 | 0 | 1 | 1 |
| PIK3R2;PIK3R1 | 1 | 1 | 0 | 1 | 1 |
| CDH24 | 1 | 1 | 0 | 1 | 1 |
| TMEM63A | 1 | 1 | 0 | 1 | 1 |
| DNAJC30 | 2 | 2 | 0 | 1 | 1 |
| MEIOC | 1 | 1 | 0 | 1 | 1 |
| D17H6S53E | 1 | 1 | 0 | 1 | 1 |
| RHOB;RHOA | 1 | 1 | 0 | 1 | 1 |
| TMEM151A | 1 | 1 | 0 | 1 | 1 |
| CLUH | 1 | 1 | -0,051131433 | 0,500134257 | 1 |
| SDHAF2 | 1 | 1 | 0 | 1 | 1 |
| ADAL | 1 | 1 | 0 | 1 | 1 |
| MAP4 | 1 | 1 | 0 | 1 | 1 |
| TERF2 | 1 | 1 | 0,012168525 | 0,820638556 | 1 |
| POGLUT1 | 3 | 3 | 0 | 1 | 1 |
| ANKRD27 | 1 | 1 | 0 | 1 | 1 |
| CCDC134 | 2 | 2 | -0,042465747 | 0,491636609 | 1 |
| ANGEL2 | 1 | 1 | 0 | 1 | 1 |
| B3GALNT1 | 1 | 1 | 0,126118501 | 0,144394796 | 1 |
| RFXAP | 1 | 1 | 0 | 1 | 1 |
| CHGA | 1 | 1 | 6,69606E-11 | 0,999981546 | 1 |
| VAMP8 | 1 | 1 | 0 | 1 | 1 |
| CDKN1B | 2 | 2 | -2,91739E-18 | 0,999999999 | 1 |
| SUMO2 | 1 | 1 | 0 | 1 | 1 |
| NPTX2 | 1 | 1 | -2,1477E-15 | 0,999999885 | 1 |

| | | | | | |
|---|---|---|---|---|---|
| ARHGAP17 | 1 | 1 | 0 | 1 | 1 |
| FCHO2 | 1 | 1 | 0 | 1 | 1 |
| PRKCG;PRKCB | 1 | 1 | 4,71906E-18 | 0,999999997 | 1 |
| SCN1A;SCN2A;SCN9/ | 1 | 1 | 0 | 1 | 1 |
| RAPGEF5 | 1 | 1 | -0,005361122 | 0,872963109 | 1 |
| TMEM68 | 2 | 2 | 0 | 1 | 1 |
| EI24 | 1 | 1 | -0,064974328 | 0,300498833 | 1 |
| MRPL53 | 2 | 2 | 0 | 1 | 1 |
| UBAP1 | 1 | 1 | 0,030396502 | 0,675061388 | 1 |
| EPN2 | 1 | 1 | 0 | 1 | 1 |
| POLR3G | 1 | 1 | 0 | 1 | 1 |
| FXYD1 | 1 | 1 | 0 | 1 | 1 |
| PALMD | 2 | 2 | -0,069529644 | 0,198962196 | 1 |
| SEPTIN12 | 2 | 2 | 3,76117E-17 | 0,999999995 | 1 |
| 2010109A12RIK | 1 | 1 | 0 | 1 | 1 |
| CD300LF | 1 | 1 | 0 | 1 | 1 |
| TMEM19 | 1 | 1 | 0 | 1 | 1 |
| ACTN1 | 2 | 2 | 4,68237E-17 | 0,999999991 | 1 |
| CAMK2G | 1 | 1 | 0 | 1 | 1 |
| CAPN7 | 1 | 1 | 0 | 1 | 1 |
| FAM189A1 | 1 | 1 | 0 | 1 | 1 |
| TNIK;MINK1 | 2 | 2 | 3,66249E-19 | 0,999999999 | 1 |
| RELL2 | 1 | 1 | 0 | 1 | 1 |
| ATP5F1A | 3 | 3 | -0,06039702 | 0,452307905 | 1 |
| KMT2A | 1 | 1 | 0 | 1 | 1 |
| CTC1 | 2 | 1 | 0,081486381 | 0,528391257 | 1 |
| NDRG2 | 1 | 1 | 5,5257E-20 | 1 | 1 |
| ANKRD52 | 1 | 1 | 0,067270467 | 0,379526804 | 1 |
| ARAF | 1 | 1 | -7,3154E-18 | 0,999999994 | 1 |
| PLCG2 | 1 | 1 | 0,015732933 | 0,730431262 | 1 |
| PRKD1;PRKD3 | 1 | 1 | 0 | 1 | 1 |
| SCN1A;SCN3A;SCN2/ | 1 | 1 | 0 | 1 | 1 |
| RAB3D;RAB8A;RAB8 | 1 | 1 | 0 | 1 | 1 |

| | | | | | |
|---|---|---|---|---|---|
| SCN1A;SCN3A;SCN2/ | 2 | 2 | 0,083788978 | 0,248787661 | 1 |
| ACVR1B | 1 | 1 | 0,068459641 | 0,330694184 | 1 |
| KCNT2 | 1 | 1 | 0 | 1 | 1 |
| CLIP1 | 1 | 1 | 0 | 1 | 1 |
| AFF4 | 1 | 1 | -0,011892444 | 0,804186135 | 1 |
| CACNB2;CACNB1 | 1 | 1 | 0 | 1 | 1 |
| ATRX | 1 | 1 | -0,197513654 | 0,327073635 | 1 |
| RTN4RL1 | 1 | 1 | 0,108499358 | 0,216306192 | 1 |
| MGAM | 1 | 1 | 0,09592969 | 0,180730004 | 1 |
| MBTPS1 | 1 | 1 | 0 | 1 | 1 |
| ABCA8B | 1 | 1 | 0 | 1 | 1 |
| FGD5 | 1 | 1 | 0 | 1 | 1 |
| RPS6KL1 | 1 | 1 | 0,026013623 | 0,622738822 | 1 |
| MTURN | 1 | 1 | 0,004817467 | 0,882223015 | 1 |
| FCGR2 | 1 | 1 | 0 | 1 | 1 |
| ABHD4 | 3 | 3 | 0 | 1 | 1 |
| NKIRAS1 | 1 | 1 | 0 | 1 | 1 |
| MOSPD2 | 1 | 1 | 0 | 1 | 1 |
| SCML2 | 1 | 1 | -0,09682963 | 0,229025006 | 1 |
| BNIP2 | 1 | 1 | 0 | 1 | 1 |
| CUL9 | 1 | 1 | 0 | 1 | 1 |
| DDHD1 | 1 | 1 | 0 | 1 | 1 |
| NDUFS2 | 1 | 1 | 0 | 1 | 1 |
| FBXO6 | 2 | 2 | 1,82167E-15 | 0,999999914 | 1 |
| PTPRS | 1 | 1 | 0,032311434 | 0,522227085 | 1 |
| BLOC1S1 | 1 | 1 | 0 | 1 | 1 |
| FZR1 | 1 | 1 | 0 | 1 | 1 |
| YIPF5 | 1 | 1 | 0 | 1 | 1 |
| ST8SIA3 | 2 | 2 | -5,54991E-20 | 1 | 1 |
| SH3RF1 | 2 | 2 | 1,51087E-17 | 0,999999994 | 1 |
| SNAP25 | 2 | 2 | 1,10066E-18 | 0,999999998 | 1 |
| PSAP | 1 | 1 | 0 | 1 | 1 |
| CACNA1A;CACNA1E | 1 | 1 | -0,045245061 | 0,38591978 | 1 |

| | | | | | |
|---|---|---|---|---|---|
| TMEM9 | 1 | 1 | 0 | 1 | 1 |
| SLC1A1 | 1 | 1 | 0 | 1 | 1 |
| CA10 | 1 | 1 | 0 | 1 | 1 |
| KIF5A;KIF5B | 2 | 2 | -8,73802E-16 | 0,999999938 | 1 |
| ERGIC3 | 1 | 1 | -0,077510163 | 0,294984864 | 1 |
| EIF4E2 | 1 | 1 | 0,096390037 | 0,416102713 | 1 |
| ARPC3 | 1 | 1 | 0 | 1 | 1 |
| KCNQ3 | 1 | 1 | 0 | 1 | 1 |
| B2M | 1 | 1 | 0,031706594 | 0,614257434 | 1 |
| CSPG5 | 1 | 1 | 0 | 1 | 1 |
| RBBP4;RBBP7 | 3 | 1 | 0 | 1 | 1 |
| FOXRED1 | 2 | 2 | 0 | 1 | 1 |
| DENND6A | 1 | 1 | 0 | 1 | 1 |
| LRP11 | 1 | 1 | 0 | 1 | 1 |
| PARK7 | 1 | 1 | 0 | 1 | 1 |
| ZFP442 | 1 | 1 | -8,86594E-18 | 0,999999994 | 1 |
| COL6A5 | 1 | 1 | 0 | 1 | 1 |
| CUTA | 1 | 1 | 0 | 1 | 1 |
| H1-4 | 1 | 1 | 0,295126592 | 0,311293861 | 1 |
| SCN3A | 1 | 1 | 0 | 1 | 1 |
| PABPN1 | 1 | 1 | 0 | 1 | 1 |
| SLC12A6;GM21985 | 2 | 2 | 6,60945E-18 | 0,999999995 | 1 |
| FBXL4 | 2 | 2 | 1,34549E-16 | 0,999999977 | 1 |
| WDSUB1 | 1 | 1 | 0 | 1 | 1 |
| ADAMTS10 | 1 | 1 | 0 | 1 | 1 |
| SFI1 | 1 | 1 | 0 | 1 | 1 |
| TMEM145 | 1 | 1 | 0 | 1 | 1 |
| KALRN | 1 | 1 | -0,017946568 | 0,682672705 | 1 |
| DBNDD2 | 1 | 1 | 0,074263874 | 0,307586086 | 1 |
| TRANK1 | 1 | 1 | 0 | 1 | 1 |
| RASSF2;RASSF4 | 1 | 1 | 0 | 1 | 1 |
| ABCA3 | 1 | 1 | 0 | 1 | 1 |
| TPGS1 | 1 | 1 | 0 | 1 | 1 |

| | | | | | |
|---|---|---|---|---|---|
| CHRM3 | 1 | 1 | 0 | 1 | 1 |
| ERMAP | 1 | 1 | 0 | 1 | 1 |
| ACSM3 | 1 | 1 | 0 | 1 | 1 |
| SLC7A1 | 1 | 1 | 0 | 1 | 1 |
| FRMPD4 | 1 | 1 | 0 | 1 | 1 |
| DNM1L | 1 | 1 | 0 | 1 | 1 |
| CAPG | 2 | 2 | 1,67316E-18 | 0,999999999 | 1 |
| CYSTM1 | 1 | 1 | 0 | 1 | 1 |
| IRF3 | 2 | 2 | -4,02564E-18 | 0,999999996 | 1 |
| NEURL1 | 1 | 1 | 0 | 1 | 1 |
| SNCG | 1 | 1 | 0 | 1 | 1 |
| ARID2 | 1 | 1 | 0 | 1 | 1 |
| IMPA1 | 1 | 1 | 0 | 1 | 1 |
| SMG6 | 2 | 2 | 0,16472305 | 0,132124018 | 1 |
| IGF1R | 2 | 2 | -9,44323E-19 | 0,999999999 | 1 |
| ACKR1 | 1 | 1 | -2,5322E-19 | 1 | 1 |
| PRELID3B | 1 | 1 | -5,72887E-19 | 0,999999999 | 1 |
| NUP214 | 1 | 1 | -0,105452969 | 0,405182255 | 1 |
| CDA | 1 | 1 | 0 | 1 | 1 |
| SLC4A4 | 1 | 1 | 0 | 1 | 1 |
| IQSEC2 | 1 | 1 | 0,028327833 | 0,666998902 | 1 |
| CD47 | 1 | 1 | 0 | 1 | 1 |
| PTN | 1 | 1 | 0,144190752 | 0,264715591 | 1 |
| CPLANE1 | 1 | 1 | 0 | 1 | 1 |
| PIP5K1C;PIP5K1A | 1 | 1 | 0 | 1 | 1 |
| LYNX1 | 1 | 1 | 0 | 1 | 1 |
| FBXO42 | 1 | 1 | 0 | 1 | 1 |
| KIAA0319 | 1 | 1 | 0 | 1 | 1 |
| KIFC2 | 1 | 1 | 0 | 1 | 1 |
| RAB3D;RAB3B | 1 | 1 | 0 | 1 | 1 |
| DPH6 | 2 | 2 | 0 | 1 | 1 |
| CXADR | 2 | 2 | 0 | 1 | 1 |
| RELCH | 1 | 1 | 0 | 1 | 1 |

| | | | | | |
|---|---|---|---|---|---|
| ATP1A2;ATP12A | 1 | 1 | -1,3237E-17 | 0,999999991 | 1 |
| YIPF3 | 1 | 1 | 0 | 1 | 1 |
| SUMO2;SUMO3 | 1 | 1 | 0 | 1 | 1 |
| BCKDHB | 1 | 1 | -0,065630431 | 0,498348164 | 1 |
| CYP4F3;CYP4F14 | 1 | 1 | 0 | 1 | 1 |
| RHOF | 2 | 2 | -1,75694E-15 | 0,999999903 | 1 |
| MIB1 | 1 | 1 | 2,02057E-19 | 0,999999999 | 1 |
| ATXN7L2 | 1 | 1 | 0 | 1 | 1 |
| GM1043 | 1 | 1 | 0,008426712 | 0,795837326 | 1 |
| CSNK1D | 1 | 1 | 0 | 1 | 1 |
| PDE4DIP | 1 | 1 | 0,231242829 | 0,306250025 | 1 |
| RGN | 1 | 1 | 0 | 1 | 1 |
| ATP8A1 | 1 | 1 | 0 | 1 | 1 |
| PTPRF;PTPRS;PTPRD | 2 | 2 | 2,14978E-15 | 0,999999916 | 1 |
| DCLK1 | 2 | 2 | -0,208328306 | 0,128491308 | 1 |
| KLK10 | 1 | 1 | 0 | 1 | 1 |
| TG | 1 | 1 | -0,192924912 | 0,149876953 | 1 |
| DPY30 | 1 | 1 | 0 | 1 | 1 |
| DIP2C | 1 | 1 | 1,63967E-17 | 0,999999991 | 1 |
| MAST2 | 1 | 1 | -0,111823234 | 0,120463376 | 1 |
| BCL11A | 1 | 1 | 0,073160949 | 0,235611548 | 1 |
| MECP2 | 1 | 1 | 0 | 1 | 1 |
| SMAD2 | 1 | 1 | 0 | 1 | 1 |
| GSC2;GSC | 1 | 1 | 0 | 1 | 1 |
| MGST1 | 1 | 1 | 9,70085E-19 | 0,999999999 | 1 |
| ZFP958 | 1 | 1 | 0 | 1 | 1 |
| DNAH8 | 1 | 1 | -0,062801328 | 0,437222216 | 1 |
| NFKB1 | 1 | 1 | -0,038021425 | 0,69861746 | 1 |
| APPBP2 | 1 | 1 | 0 | 1 | 1 |
| HPDL | 1 | 1 | 0 | 1 | 1 |
| FUNDC1 | 1 | 1 | 0 | 1 | 1 |
| GABBR1 | 1 | 1 | 0 | 1 | 1 |
| CFL1;CFL2 | 3 | 3 | 5,08907E-17 | 0,999999982 | 1 |

| | | | | | |
|---|---|---|---|---|---|
| DEPDC5 | 1 | 1 | 0 | 1 | 1 |
| NWD1 | 1 | 1 | 0 | 1 | 1 |
| MRPL10 | 2 | 2 | 7,98258E-13 | 0,999997634 | 1 |
| MRPL30 | 1 | 1 | 0 | 1 | 1 |
| GRB10 | 1 | 1 | 0 | 1 | 1 |
| FBXW15 | 1 | 1 | 0 | 1 | 1 |
| MAP4K2 | 1 | 1 | -0,103033933 | 0,3301793 | 1 |
| RYR1;RYR2 | 1 | 1 | 0 | 1 | 1 |
| GAS2L1 | 1 | 1 | 0 | 1 | 1 |
| NBEAL2 | 2 | 2 | -0,02205185 | 0,691398578 | 1 |
| ANO8 | 1 | 1 | 0,127129299 | 0,325367149 | 1 |
| TMEM14C | 1 | 1 | 0 | 1 | 1 |
| ANKS1A | 2 | 1 | 0 | 1 | 1 |
| SLC47A2 | 1 | 1 | 0 | 1 | 1 |
| STK11IP | 1 | 1 | -0,147966654 | 0,190959821 | 1 |
| GM14692 | 1 | 1 | 0 | 1 | 1 |
| BTBD11 | 1 | 1 | -9,89989E-18 | 0,999999993 | 1 |
| QRICH1 | 1 | 1 | 0 | 1 | 1 |
| FEM1AA | 1 | 1 | 0 | 1 | 1 |
| DOCK9 | 2 | 2 | -0,000641008 | 0,950051291 | 1 |
| LIMS1 | 1 | 1 | 0,128079023 | 0,148507689 | 1 |
| TDRD3 | 1 | 1 | 0 | 1 | 1 |
| AGBL4 | 1 | 1 | 0 | 1 | 1 |
| TBC1D9 | 1 | 1 | -0,214864769 | 0,305982146 | 1 |
| LDHC;LDHA;LDHB | 1 | 1 | 0 | 1 | 1 |
| MATR3 | 1 | 1 | 0 | 1 | 1 |
| PCDHGA8 | 1 | 1 | 0 | 1 | 1 |
| RAB19 | 1 | 1 | 0 | 1 | 1 |
| FER1L5 | 1 | 1 | 0 | 1 | 1 |
| SIPA1L3;SIPA1L1 | 1 | 1 | 0 | 1 | 1 |
| OVCA2 | 1 | 1 | -0,001671014 | 0,910154059 | 1 |
| ABCB9 | 2 | 2 | 0 | 1 | 1 |
| CRIPT | 1 | 1 | 0 | 1 | 1 |

| | | | | | |
|---|---|---|---|---|---|
| FBXW11 | 1 | 1 | 0 | 1 | 1 |
| CFAP46 | 1 | 1 | 0 | 1 | 1 |
| AGFG2;AGFG1 | 1 | 1 | 0 | 1 | 1 |
| PLA2G6 | 2 | 2 | 1,3835E-16 | 0,999999974 | 1 |
| SH2D1A | 1 | 1 | 0 | 1 | 1 |
| MYH11;MYH10;MYH | 1 | 1 | 0 | 1 | 1 |
| IGDCC4 | 1 | 1 | 0,082058587 | 0,31351547 | 1 |
| CRAT | 1 | 1 | -0,172107849 | 0,13981069 | 1 |
| LDAH | 1 | 1 | 0 | 1 | 1 |
| MPV17 | 1 | 1 | 0 | 1 | 1 |
| MTHFD1L;MTHFD1 | 1 | 1 | 0 | 1 | 1 |
| CCDC85A | 1 | 1 | 0 | 1 | 1 |
| EPHA3;EPHA4;EPHA! | 1 | 1 | 0,050314772 | 0,336015579 | 1 |
| LIN7B;LIN7A | 2 | 2 | -3,98478E-19 | 1 | 1 |
| PCDHGA9;PCDHGA8 | 1 | 1 | 0 | 1 | 1 |
| TMEM94 | 1 | 1 | 0,02732002 | 0,570284976 | 1 |
| GLT6D1 | 1 | 1 | 0 | 1 | 1 |
| NUDT16L1 | 1 | 1 | 0 | 1 | 1 |
| GSTT2 | 1 | 1 | 0 | 1 | 1 |
| GIMAP8 | 1 | 1 | 0 | 1 | 1 |
| VMN2R3 | 1 | 1 | 0 | 1 | 1 |
| KCNA5 | 1 | 1 | 0,093687227 | 0,182077519 | 1 |
| CPVL | 1 | 1 | 0 | 1 | 1 |
| ACO1;IREB2 | 1 | 1 | 0 | 1 | 1 |
| SKOR2 | 1 | 1 | 0 | 1 | 1 |
| IL23R | 1 | 1 | 0 | 1 | 1 |
| ADCY10 | 1 | 1 | -0,023544987 | 0,647751753 | 1 |
| ZBTB48 | 1 | 1 | 0 | 1 | 1 |
| SFT2D2 | 1 | 1 | 0 | 1 | 1 |
| VPS13B | 2 | 2 | -0,102037997 | 0,459223692 | 1 |
| MTG1 | 1 | 1 | -0,255380535 | 0,117889136 | 1 |
| ODR4 | 1 | 1 | -0,041469026 | 0,463290534 | 1 |
| RPL35 | 1 | 1 | 0,046070866 | 0,573688424 | 1 |

| | | | | | |
|---|---|---|---|---|---|
| GRAMD1A | 1 | 1 | 0 | 1 | 1 |
| RABL3 | 1 | 1 | 0 | 1 | 1 |
| ENPP5 | 1 | 1 | 0 | 1 | 1 |
| DEGS1 | 2 | 2 | -4,63634E-18 | 0,999999997 | 1 |
| OLFR1458 | 1 | 1 | -0,079597975 | 0,266441449 | 1 |
| PPM1J;PPM1H | 2 | 2 | 0 | 1 | 1 |
| RTL8C | 1 | 1 | 0 | 1 | 1 |
| GATM | 4 | 4 | -1,45855E-17 | 0,99999999 | 1 |
| SNRPE | 1 | 1 | 0 | 1 | 1 |
| SHC1 | 1 | 1 | -0,145658109 | 0,311251541 | 1 |
| CYP1A2 | 1 | 1 | 0 | 1 | 1 |
| TRPC3 | 1 | 1 | 0 | 1 | 1 |
| RTP3 | 1 | 1 | 0 | 1 | 1 |
| ENO3 | 3 | 3 | 0,02608146 | 0,615693325 | 1 |
| TMEM223 | 1 | 1 | 0 | 1 | 1 |
| PLEKHA6 | 1 | 1 | 0 | 1 | 1 |
| KCNAB3 | 1 | 1 | -0,057111218 | 0,537334849 | 1 |
| PRX | 1 | 1 | 0 | 1 | 1 |
| OSTC | 1 | 1 | 0 | 1 | 1 |
| RPP25L | 1 | 1 | 0 | 1 | 1 |
| ANKZF1 | 1 | 1 | 0 | 1 | 1 |
| ABCA17 | 1 | 1 | 0 | 1 | 1 |
| IDS | 2 | 1 | 0 | 1 | 1 |
| GABBR1 | 1 | 1 | 0,009459655 | 0,791038046 | 1 |
| TYMP | 1 | 1 | 0 | 1 | 1 |
| ICMT | 1 | 1 | 0 | 1 | 1 |
| WDFY2 | 1 | 1 | 0,099017053 | 0,306018156 | 1 |
| TICAM2 | 1 | 1 | -0,193081869 | 0,243327928 | 1 |
| ALDH16A1 | 1 | 1 | 0,215991248 | 0,288706258 | 1 |
| CCHCR1 | 1 | 1 | 0 | 1 | 1 |
| FMNL2;FMNL1 | 1 | 1 | -0,076553908 | 0,219826334 | 1 |
| ABHD14B | 1 | 1 | 0 | 1 | 1 |
| ORAI2 | 1 | 1 | 0 | 1 | 1 |

| | | | | | |
|---|---|---|---|---|---|
| SLC25A53 | 1 | 1 | 0 | 1 | 1 |
| SGSH | 1 | 1 | 0 | 1 | 1 |
| MRPS26 | 2 | 2 | 0 | 1 | 1 |
| NDUFA1 | 2 | 2 | 2,15429E-19 | 0,999999999 | 1 |
| MRPL50 | 1 | 1 | 0,145333768 | 0,182252393 | 1 |
| ADAMTS20 | 1 | 1 | 0 | 1 | 1 |
| TNS3;TNS2 | 1 | 1 | -0,090884312 | 0,208277346 | 1 |
| ADORA1 | 1 | 1 | 0 | 1 | 1 |
| ACP1 | 1 | 1 | 0 | 1 | 1 |
| VSTM2B | 1 | 1 | -0,170421187 | 0,275298549 | 1 |
| TAX1BP3 | 1 | 1 | 0 | 1 | 1 |
| STXBP1 | 3 | 3 | 3,00945E-17 | 0,999999995 | 1 |
| TGOLN2 | 1 | 1 | 0 | 1 | 1 |
| ROBO4 | 1 | 1 | 0 | 1 | 1 |
| SEZ6 | 1 | 1 | 0 | 1 | 1 |
| DNAJB4;DNAJB1 | 1 | 1 | 0,092865556 | 0,23678505 | 1 |
| ARMC8 | 1 | 1 | 1,00826E-18 | 0,999999999 | 1 |
| KLHDC7A | 1 | 1 | 0 | 1 | 1 |
| UBAP1 | 1 | 1 | 0 | 1 | 1 |
| ABCG2;ABCG3 | 1 | 1 | 0,066786964 | 0,404610794 | 1 |
| ALDH1L2 | 1 | 1 | -1,11265E-20 | 1 | 1 |
| SCN3A;SCN2A;SCN9/ | 2 | 2 | -9,21452E-17 | 0,999999975 | 1 |
| sp\|Q8VE95\|CH082_ | 1 | 1 | -0,1003998 | 0,223755148 | 1 |
| ATG2A | 1 | 1 | 0 | 1 | 1 |
| CYP26B1 | 1 | 1 | 0 | 1 | 1 |
| HNRNPH1 | 1 | 1 | 8,02792E-17 | 0,999999987 | 1 |
| CUL9;CUL7 | 1 | 1 | -0,051768928 | 0,476996439 | 1 |
| UBXN4 | 1 | 1 | 0 | 1 | 1 |
| SAXO2 | 1 | 1 | 0 | 1 | 1 |
| COPS7A | 1 | 1 | 0 | 1 | 1 |
| MAVS | 1 | 1 | 0 | 1 | 1 |
| SBF1 | 1 | 1 | 0 | 1 | 1 |
| JAM2 | 1 | 1 | 6,826E-19 | 0,999999999 | 1 |

| | | | | | |
|---|---|---|---|---|---|
| PDF | 1 | 1 | 0 | 1 | 1 |
| FGF13 | 1 | 1 | 0 | 1 | 1 |
| MEP1B | 1 | 1 | -2,11826E-21 | 1 | 1 |
| ATP13A3 | 1 | 1 | 0 | 1 | 1 |
| PANK1 | 1 | 1 | -0,10297713 | 0,254393765 | 1 |
| KCNA1;KCNA3;KCNA | 2 | 2 | 3,49969E-18 | 0,999999997 | 1 |
| PTRHD1 | 1 | 1 | -0,091061903 | 0,354494983 | 1 |
| NDUFS1 | 1 | 1 | 0 | 1 | 1 |
| SNAPIN | 1 | 1 | 0 | 1 | 1 |
| ACOT8 | 1 | 1 | 0,054040452 | 0,359834045 | 1 |
| PLAT | 1 | 1 | 0 | 1 | 1 |
| PKP3 | 1 | 1 | 0 | 1 | 1 |
| ARFRP1 | 1 | 1 | 0 | 1 | 1 |
| SLC27A6 | 1 | 1 | 0 | 1 | 1 |
| SSU72 | 1 | 1 | 0 | 1 | 1 |
| UNC13B | 1 | 1 | 0 | 1 | 1 |
| EPHB3;EPHB4 | 1 | 1 | 0 | 1 | 1 |
| ACAD10 | 2 | 2 | -9,20758E-17 | 0,999999983 | 1 |
| RGMA | 1 | 1 | 0 | 1 | 1 |
| TOR1B | 1 | 1 | 0 | 1 | 1 |
| PPIP5K1;PPIP5K2 | 1 | 1 | 0 | 1 | 1 |
| LRP2 | 1 | 1 | 0 | 1 | 1 |
| MSL3 | 1 | 1 | 0 | 1 | 1 |
| SLC17A8;SLC17A6 | 1 | 1 | 0 | 1 | 1 |
| STXBP5L;STXBP5 | 1 | 1 | 0 | 1 | 1 |
| KATNAL2 | 1 | 1 | 0 | 1 | 1 |
| CSF2RB2;CSF2RB | 1 | 1 | 0 | 1 | 1 |
| EPHB2;EPHA7 | 1 | 1 | 0 | 1 | 1 |
| ARL6IP1 | 1 | 1 | 0 | 1 | 1 |
| HSPA12B | 1 | 1 | 0 | 1 | 1 |
| CRK | 1 | 1 | 0 | 1 | 1 |
| RAB3D;RAB3A | 1 | 1 | 0 | 1 | 1 |
| IGHV9-4 | 1 | 1 | 0 | 1 | 1 |

| | | | | | |
|---|---|---|---|---|---|
| CD44 | 3 | 3 | -5,11151E-16 | 0,999999951 | 1 |
| PXN | 1 | 1 | 0 | 1 | 1 |
| VMN2R70 | 1 | 1 | 0 | 1 | 1 |
| HSD17B7 | 1 | 1 | 0 | 1 | 1 |
| UGT1A6 | 1 | 1 | 0 | 1 | 1 |
| METTL5 | 1 | 1 | 0 | 1 | 1 |
| KRT1;KRT6A;KRT2;KI | 2 | 2 | 0,117778748 | 0,482736135 | 1 |
| KRT86;KRT81 | 1 | 1 | 0 | 1 | 1 |
| OCIAD1 | 1 | 1 | 0 | 1 | 1 |
| ABHD17B | 2 | 2 | 0,051539073 | 0,331772687 | 1 |
| CCDC158 | 1 | 1 | 0,016185135 | 0,811940831 | 1 |
| MYO5B | 1 | 1 | -0,026192997 | 0,563284756 | 1 |
| B4GALT4 | 1 | 1 | 0 | 1 | 1 |
| MBNL2 | 1 | 1 | 0 | 1 | 1 |
| GNA12 | 1 | 1 | 0 | 1 | 1 |
| SCN1A;SCN3A;SCN2/ | 1 | 1 | 0,078064376 | 0,255140996 | 1 |
| PDCD6IP | 1 | 1 | 0 | 1 | 1 |
| CHST14 | 1 | 1 | -0,109118902 | 0,431676619 | 1 |
| DOP1B | 1 | 1 | 0 | 1 | 1 |
| HDAC1;HDAC2 | 1 | 1 | 0 | 1 | 1 |
| OTUD7B | 1 | 1 | 0 | 1 | 1 |
| KCTD21 | 1 | 1 | 0 | 1 | 1 |
| GOLGA7;GOLGA7B | 1 | 1 | 0 | 1 | 1 |
| UTP20 | 1 | 1 | -0,152642577 | 0,234615761 | 1 |
| CEP85L | 1 | 1 | 0 | 1 | 1 |
| SPRED2 | 1 | 1 | 0 | 1 | 1 |
| MYO1A;MYO1B | 1 | 1 | 0 | 1 | 1 |
| EHD4;EHD3 | 2 | 2 | 0,064676782 | 0,232308526 | 1 |
| TBC1D25 | 1 | 1 | 0 | 1 | 1 |
| RLN1 | 1 | 1 | 2,09185E-15 | 0,999999899 | 1 |
| TUBB4B;TUBB5;TUBI | 1 | 1 | 0 | 1 | 1 |
| ATAD2 | 1 | 1 | 0 | 1 | 1 |
| TIMM23 | 2 | 2 | -5,8571E-17 | 0,999999991 | 1 |

| | | | | | |
|---|---|---|---|---|---|
| TAT | 1 | 1 | 0 | 1 | 1 |
| CACNB4;CACNB1 | 1 | 1 | 0 | 1 | 1 |
| TMCO5A | 1 | 1 | -0,136789309 | 0,278694178 | 1 |
| UNC119B;UNC119 | 1 | 1 | 0 | 1 | 1 |
| CLDN10 | 1 | 1 | 0 | 1 | 1 |
| NT5C1A | 1 | 1 | 0,060697185 | 0,454153949 | 1 |
| CRK | 1 | 1 | 0 | 1 | 1 |
| H3-5;H3C11;H3C15;I | 2 | 1 | 0,5756003 | 0,162808786 | 1 |
| RIMS1;RIMS2 | 1 | 1 | 0 | 1 | 1 |
| ERBB4 | 1 | 1 | 0 | 1 | 1 |
| ADGRB3;ADGRB2 | 1 | 1 | 0 | 1 | 1 |
| SMIM26 | 1 | 1 | 0 | 1 | 1 |
| MSTN | 1 | 1 | 0 | 1 | 1 |
| MDGA2 | 1 | 1 | 0,113578099 | 0,160153711 | 1 |
| SERPINA3I | 1 | 1 | 0 | 1 | 1 |
| WDR82 | 1 | 1 | 0 | 1 | 1 |
| HDAC9 | 1 | 1 | 0 | 1 | 1 |
| RAPGEF3;RAPGEF4 | 1 | 1 | 0,068119475 | 0,26520902 | 1 |
| sp\|E0CYV9\|CD054_I | 1 | 1 | 0 | 1 | 1 |
| PTPRA | 1 | 1 | 0 | 1 | 1 |
| SNRNP25 | 1 | 1 | 3,20611E-18 | 0,999999997 | 1 |
| GPX3 | 1 | 1 | 0,261075162 | 0,170956851 | 1 |
| ETFBKMT | 1 | 1 | 0,028986976 | 0,536888508 | 1 |
| MMP15 | 1 | 1 | 0 | 1 | 1 |
| TAF6L | 1 | 1 | 0 | 1 | 1 |
| MMP23 | 1 | 1 | 0 | 1 | 1 |
| BCKDHB | 1 | 1 | -8,27471E-23 | 1 | 1 |
| TMBIM1 | 1 | 1 | 0 | 1 | 1 |
| MRPS5 | 3 | 3 | 0 | 1 | 1 |
| CEMIP2 | 1 | 1 | -0,085767325 | 0,69670338 | 1 |
| LRRC74B | 1 | 1 | 0 | 1 | 1 |
| CPEB1 | 1 | 1 | -0,053703218 | 0,727128776 | 1 |
| H1-3 | 3 | 2 | 0 | 1 | 1 |

| | | | | | |
|---|---|---|---|---|---|
| SHANK3 | 1 | 1 | -0,039883981 | 0,536677348 | 1 |
| PSIP1 | 1 | 1 | -3,01999E-17 | 0,999999989 | 1 |
| 1700029H14RIK | 1 | 1 | -0,095740022 | 0,620024293 | 1 |
| KIAA0895L | 1 | 1 | 0 | 1 | 1 |
| MYBPC3 | 1 | 1 | -0,030340696 | 0,639970935 | 1 |
| SIX6 | 1 | 1 | 1,65379E-18 | 0,999999998 | 1 |
| TRO | 1 | 1 | 0 | 1 | 1 |
| MMRN1 | 1 | 1 | 0 | 1 | 1 |
| KRT15 | 19 | 2 | 1,01833E-19 | 1 | 1 |
| CLU | 1 | 1 | 0 | 1 | 1 |
| ZCCHC18 | 1 | 1 | -0,051587238 | 0,539131061 | 1 |
| TMEM237 | 1 | 1 | 0 | 1 | 1 |
| TIMM21 | 2 | 2 | -3,56735E-17 | 0,999999989 | 1 |
| PAN3 | 1 | 1 | 0 | 1 | 1 |
| ADCY6 | 1 | 1 | 0 | 1 | 1 |
| COL1A2 | 7 | 2 | 1,33863E-15 | 1 | 1 |
| NRXN1 | 1 | 1 | -0,073227429 | 0,42333231 | 1 |
| SLC8A3 | 1 | 1 | 0 | 1 | 1 |
| KCTD12 | 1 | 1 | 0 | 1 | 1 |
| YIPF2 | 1 | 1 | 0 | 1 | 1 |
| MME | 1 | 1 | 0 | 1 | 1 |
| PTCD2 | 1 | 1 | 0 | 1 | 1 |
| TUBB1 | 1 | 1 | 0,113940583 | 0,349162802 | 1 |
| KCNJ15 | 1 | 1 | 0 | 1 | 1 |
| HERC3 | 1 | 1 | 0 | 1 | 1 |
| CACFD1 | 1 | 1 | 1,60028E-14 | 0,999999746 | 1 |
| RBM26 | 1 | 1 | -0,103236019 | 0,548494506 | 1 |
| PRSS58 | 1 | 1 | -0,057633185 | 0,611204081 | 1 |
| MTFMT | 1 | 1 | -0,109325148 | 0,261310839 | 1 |
| GTPBP8 | 1 | 1 | -0,122662163 | 0,229598213 | 1 |
| TENM4;TENM2 | 1 | 1 | 0 | 1 | 1 |
| TULP4 | 1 | 1 | 0 | 1 | 1 |
| YIPF4 | 1 | 1 | 0 | 1 | 1 |

| | | | | | |
|---|---|---|---|---|---|
| UCP3 | 1 | 1 | -5,79919E-15 | 0,999999867 | 1 |
| PGK2 | 1 | 1 | -0,096578868 | 0,488209889 | 1 |
| DNAJC28 | 1 | 1 | 0 | 1 | 1 |
| LIPT2 | 1 | 1 | -9,26202E-19 | 0,999999999 | 1 |
| OLFR1141 | 1 | 1 | 0 | 1 | 1 |
| SLC17A8 | 1 | 1 | -3,01438E-19 | 0,999999999 | 1 |
| SMC1B | 1 | 1 | 0 | 1 | 1 |
| TOMM6 | 1 | 1 | 0 | 1 | 1 |
| SLIT3 | 1 | 1 | 0 | 1 | 1 |
| OAS2 | 1 | 1 | 0,171579484 | 0,337627729 | 1 |
| PRPH | 1 | 1 | 0 | 1 | 1 |
| RPL36 | 1 | 1 | 0 | 1 | 1 |
| SMIM4 | 1 | 1 | 0 | 1 | 1 |
| S100A14 | 1 | 1 | 0,476920367 | 0,185414216 | 1 |
| SPECC1L | 1 | 1 | 5,29243E-19 | 0,999999999 | 1 |
| TM9SF1 | 1 | 1 | 0 | 1 | 1 |
| CHD6 | 1 | 1 | 0 | 1 | 1 |
| PTCD1 | 1 | 1 | 0 | 1 | 1 |
| STOX2 | 1 | 1 | 0 | 1 | 1 |
| ARAP1 | 1 | 1 | 0 | 1 | 1 |
| ARG2 | 1 | 1 | 0 | 1 | 1 |
| CDH4 | 1 | 1 | 0 | 1 | 1 |
| HBS1L | 1 | 1 | 0 | 1 | 1 |
| CLCN3;CLCN4;CLCN5 | 1 | 1 | 0 | 1 | 1 |
| NCOR1 | 1 | 1 | 0,070733242 | 0,752003848 | 1 |
| UQCC3 | 1 | 1 | 0 | 1 | 1 |
| TEX9 | 1 | 1 | -0,267842625 | 0,226232908 | 1 |
| ZBTB4 | 1 | 1 | -0,022348719 | 0,688158203 | 1 |
| MACF1;DST | 1 | 1 | -0,261109115 | 0,154355383 | 1 |
| PLA2G4C | 1 | 1 | 0 | 1 | 1 |
| MUL1 | 2 | 1 | 0,179844988 | 0,197243417 | 1 |
| APC2 | 1 | 1 | 0 | 1 | 1 |
| ATP8A1 | 1 | 1 | 0 | 1 | 1 |

| | | | | | |
|---|---|---|---|---|---|
| ACAD11 | 1 | 1 | -0,105005532 | 0,286725394 | 1 |
| FDX1 | 1 | 1 | -0,046772821 | 0,543130127 | 1 |
| SGSM2 | 1 | 1 | 0 | 1 | 1 |
| ITGAE | 1 | 1 | 0 | 1 | 1 |
| FAXC | 1 | 1 | -0,23149138 | 0,156126575 | 1 |
| IFIH1 | 1 | 1 | 0 | 1 | 1 |
| FLNC | 1 | 1 | 0 | 1 | 1 |
| TIFAB | 1 | 1 | 0 | 1 | 1 |
| OLFR576 | 1 | 1 | -3,37212E-20 | 1 | 1 |
| HMGN5 | 5 | 0 | | | |
| AGT | 4 | 0 | | | |
| CCDC50 | 1 | 0 | | | |
| CCAR2 | 30 | 0 | | | |
| FTO | 6 | 1 | | | |
| CMTR1 | 2 | 0 | | | |
| SCLY | 3 | 1 | | | |
| GBP2 | 3 | 0 | | | |
| APOL9B;APOL9A | 1 | 0 | | | |
| HEXIM1 | 3 | 0 | | | |
| LUC7L | 1 | 0 | | | |
| PPP4R3A | 8 | 0 | | | |
| THUMPD1 | 13 | 0 | | | |
| SRSF7 | 3 | 0 | | | |
| SERPINA6 | 3 | 0 | | | |
| FBLN1 | 1 | 0 | | | |
| FGB | 12 | 0 | | | |
| TRMT5 | 3 | 0 | | | |
| PPM1G | 15 | 0 | | | |
| GSTA2;GSTA1 | 1 | 0 | | | |
| GTF2A2 | 1 | 0 | | | |
| FCHSD2 | 1 | 0 | | | |
| HTATSF1 | 5 | 0 | | | |
| ZNF428 | 1 | 0 | | | |

| | | |
|---|---|---|
| POLR2E | 1 | 0 |
| NCBP1 | 3 | 0 |
| GTF2B | 2 | 0 |
| CES1C;CES1D | 1 | 0 |
| TTC33 | 1 | 0 |
| FGA | 8 | 1 |
| C4B | 3 | 0 |
| RBM12 | 3 | 0 |
| SPTBN5 | 1 | 0 |
| RNF220 | 2 | 0 |
| TBPL1 | 2 | 0 |
| WRNIP1 | 4 | 0 |
| ADIPOQ | 2 | 0 |
| RPS6KA5 | 5 | 0 |
| FAM169A | 12 | 0 |
| DMTF1 | 1 | 0 |
| RAMAC | 1 | 0 |
| SUB1 | 11 | 1 |
| YAP1 | 1 | 0 |
| CES1D | 1 | 0 |
| S100A11 | 3 | 0 |
| HAL | 3 | 0 |
| HEATR3 | 1 | 0 |
| HP | 8 | 0 |
| sp\|P01654\|KV3A1_I | 1 | 0 |
| HDGFL3 | 6 | 1 |
| UBE2E1;UBE2E2 | 2 | 0 |
| GTF2I | 1 | 1 |
| SERPINA1B;SERPINA | 1 | 0 |
| SERPINA1D | 5 | 0 |
| CFAP74 | 1 | 0 |
| RPS6KC1 | 1 | 0 |
| CBX5 | 2 | 0 |

| | | |
|---|---|---|
| DUS3L | 4 | 0 |
| HPX | 18 | 0 |
| H2-Q10;H2-D1 | 1 | 0 |
| APEX1 | 5 | 0 |
| DOHH | 1 | 0 |
| FREM1 | 1 | 0 |
| RHOC | 1 | 0 |
| ECEL1 | 1 | 0 |
| APOL10A | 1 | 0 |
| MYOZ2 | 1 | 1 |
| INMT | 3 | 0 |
| GAN | 1 | 0 |
| ETNPPL | 3 | 1 |
| CFDP1 | 4 | 0 |
| ZC3HC1 | 2 | 0 |
| UBE2W | 1 | 0 |
| OGFOD1 | 5 | 0 |
| MUP2 | 2 | 0 |
| STAT5A;STAT5B | 1 | 0 |
| NFYB | 2 | 0 |
| ZKSCAN16 | 1 | 1 |
| CRP | 2 | 0 |
| KNG1 | 4 | 1 |
| UBAP2 | 2 | 1 |
| PPP4R2 | 2 | 0 |
| PRPF39 | 1 | 0 |
| DHX15 | 8 | 1 |
| URI1 | 1 | 0 |
| RBBP8 | 1 | 0 |
| sp\|P06330\|HVM51_ | 3 | 0 |
| SNRPA | 1 | 0 |
| FGGY | 1 | 0 |
| MTREX | 1 | 0 |

| | | |
|---|---|---|
| ILKAP | 1 | 0 |
| PSMB10 | 1 | 0 |
| USP13 | 2 | 0 |
| SERPINA1B;SERPINA | 4 | 1 |
| SERPINA1C | 6 | 0 |
| CWF19L1 | 2 | 0 |
| TIMELESS | 1 | 0 |
| KCTD8;KCTD16 | 1 | 1 |
| JMJD6 | 1 | 0 |
| HNRNPAB;HNRNPDL | 1 | 0 |
| SARNP | 2 | 1 |
| GAREM2 | 1 | 0 |
| KIAA0100 | 1 | 0 |
| H2-EB2 | 1 | 0 |
| ANXA1 | 17 | 1 |
| XRCC5 | 2 | 0 |
| sp\|P01631\|KV2A7_I | 1 | 0 |
| IGKV12-41 | 1 | 1 |
| GC | 5 | 0 |
| PSME3IP1 | 2 | 0 |
| TOPBP1 | 1 | 0 |
| CSTF2 | 1 | 0 |
| LSM8 | 2 | 1 |
| ADAMTSL2 | 1 | 0 |
| PBX1 | 1 | 0 |
| AKAP8 | 2 | 0 |
| DCHS1 | 1 | 0 |
| NEDD4;NEDD4L | 1 | 0 |
| TOGARAM1 | 1 | 0 |
| METTL3 | 1 | 0 |
| SSB | 10 | 0 |
| OSGEP | 1 | 0 |
| DRG1;DRG2 | 1 | 0 |

| | | |
|---|---|---|
| CDK9 | 1 | 0 |
| GTF2F1 | 1 | 0 |
| SERPINB6B | 1 | 0 |
| DHPS | 4 | 0 |
| HNRNPM | 1 | 0 |
| CLSTN2 | 1 | 0 |
| ACAT2 | 1 | 1 |
| FOXK1 | 2 | 0 |
| GM45623 | 1 | 1 |
| TRMT61A | 2 | 0 |
| FETUB | 1 | 0 |
| PDCD2 | 1 | 0 |
| PPIH | 1 | 0 |
| PON1 | 2 | 0 |
| SERPINA1E | 6 | 0 |
| PAPOLA;PAPOLB | 1 | 0 |
| OLFR552 | 1 | 0 |
| ZSCAN22 | 1 | 0 |
| CBX1 | 6 | 0 |
| JADE3 | 1 | 0 |
| SERPINB12 | 1 | 0 |
| HMBS | 4 | 0 |
| CFD | 1 | 0 |
| TYW3 | 1 | 0 |
| DRP2 | 1 | 1 |
| PKNOX2 | 1 | 0 |
| DRAP1 | 1 | 0 |
| ADH1 | 1 | 0 |
| GANC | 7 | 1 |
| NR3C2 | 1 | 0 |
| DCAF5 | 3 | 0 |
| SDAD1 | 1 | 0 |
| TCEAL1 | 1 | 0 |

| | | |
|---|---|---|
| POLR2I | 1 | 1 |
| COG4 | 2 | 0 |
| ODAD4 | 1 | 1 |
| MAP7D3 | 1 | 0 |
| ELOA | 1 | 0 |
| UBL7 | 2 | 0 |
| TSC22D1 | 1 | 0 |
| KBTBD4 | 1 | 0 |
| LRG1 | 2 | 0 |
| MYH11 | 1 | 0 |
| HSPB1 | 1 | 1 |
| PAPOLA | 1 | 0 |
| TP53RKB | 1 | 0 |
| NUDT21 | 2 | 0 |
| ITIH4 | 5 | 1 |
| NEK2;IRAK1 | 1 | 0 |
| DHX9 | 1 | 0 |
| MOCS2 | 1 | 0 |
| PIBF1 | 1 | 0 |
| KIF20B | 1 | 0 |
| D130040H23RIK | 1 | 0 |
| KIF23 | 1 | 0 |
| PIP5KL1 | 1 | 0 |
| sp\|P04945\|KV6AB_I | 1 | 0 |
| IGKV6-17 | 2 | 0 |
| sp\|P01644\|KV5AB_I | 4 | 0 |
| ANKRD44 | 1 | 0 |
| UBR5 | 1 | 1 |
| HAT1 | 1 | 0 |
| NAA20 | 2 | 0 |
| QSOX1 | 1 | 0 |
| GABPA | 1 | 0 |
| YEATS2 | 1 | 0 |

| | | |
|---|---|---|
| LONRF2 | 1 | 0 |
| LHFPL4 | 1 | 0 |
| TOMM34 | 1 | 0 |
| SLC22A6 | 1 | 0 |
| 1810024B03RIK | 1 | 0 |
| UBE2A | 1 | 0 |
| ZCCHC8 | 1 | 0 |
| CDK3 | 1 | 0 |
| WDR12 | 1 | 0 |
| TXNL4A | 1 | 0 |
| TRMT6 | 3 | 0 |
| CCDC159 | 1 | 0 |
| NFYC | 2 | 0 |
| ASB3 | 1 | 0 |
| RRAGD | 1 | 0 |
| APBA1;APBA2 | 2 | 0 |
| TERB1 | 1 | 0 |
| PRSS44 | 1 | 1 |
| SNRPA1 | 1 | 0 |
| RIPK2 | 1 | 0 |
| ATF2 | 1 | 0 |
| SFR1 | 3 | 0 |
| UBR7 | 2 | 0 |
| IL18 | 2 | 0 |
| RTF1 | 3 | 0 |
| PUS7 | 1 | 0 |
| PSMG3 | 2 | 0 |
| NAA15 | 1 | 0 |
| IGHA | 1 | 0 |
| HEBP2 | 1 | 0 |
| SLC6A13 | 1 | 0 |
| SNRPD1 | 1 | 1 |
| ARL4A | 1 | 1 |

| | | |
|---|---|---|
| PPP1R37 | 2 | 1 |
| IGHG1 | 1 | 0 |
| OLFR1257 | 1 | 0 |
| OLFR1228;OLFR1222 | 1 | 1 |
| IRX2 | 1 | 0 |
| PLCH1 | 1 | 1 |
| EPHA1 | 1 | 0 |
| SETD6 | 1 | 0 |
| HROB | 1 | 0 |
| CRY1 | 1 | 0 |
| TCP11L2 | 1 | 0 |
| ATG4B | 1 | 0 |
| CDCA2 | 1 | 1 |
| CNBD1 | 1 | 0 |
| ANGPTL8 | 1 | 0 |
| ORM2;ORM1 | 1 | 0 |
| SSBP2;SSBP3 | 1 | 0 |
| ERICH6B | 1 | 0 |
| NCBP2 | 1 | 0 |
| ANXA8 | 5 | 0 |
| INSRR | 1 | 1 |
| WDCP | 1 | 0 |
| TC2N | 1 | 0 |
| HTR3A | 1 | 0 |
| TDP2 | 1 | 0 |
| ARL2BP | 2 | 1 |
| UBXN2A | 1 | 0 |
| ICA | 1 | 0 |
| GTF2A1 | 1 | 0 |
| RPRD1A | 1 | 0 |
| TCEA3 | 1 | 1 |
| SLC25A37 | 1 | 0 |
| SUMO3 | 1 | 0 |

| | | |
|---|---|---|
| DAB2 | 1 | 0 |
| APOA2 | 1 | 0 |
| RPS5 | 1 | 1 |
| FAT1 | 1 | 0 |
| HEATR1 | 1 | 0 |
| GLI1 | 1 | 0 |
| PHAX | 1 | 0 |
| FAM151B | 1 | 0 |
| sp\|P01680\|KV4A1_I | 1 | 0 |
| SATB2 | 1 | 1 |
| APOC3 | 1 | 0 |
| TOR1AIP2 | 1 | 0 |
| LGALS3 | 5 | 0 |
| SUPT16H | 1 | 0 |
| SAAL1 | 1 | 0 |
| APCS | 1 | 0 |
| SCYL3 | 1 | 0 |
| TGM3 | 7 | 1 |
| sp\|Q8BHN7\|CL029_ | 1 | 0 |
| TRMT1 | 1 | 0 |
| FLNB | 1 | 0 |
| FBXO33 | 1 | 0 |
| IGHG1 | 3 | 0 |
| IGHG2C | 3 | 0 |
| METTL1 | 2 | 0 |
| PSMB9 | 1 | 0 |
| LARP7 | 1 | 0 |
| SUPT4H1A;SUPT4H1 | 1 | 0 |
| NOLC1 | 1 | 0 |
| H2-Q10;H2-D1 | 1 | 0 |
| HSPA12A | 1 | 1 |
| RBFOX1 | 1 | 0 |
| ORM1 | 2 | 0 |

| | | |
|---|---|---|
| HMGB2 | 3 | 0 |
| HDAC2 | 1 | 0 |
| HDGFL2;HDGFL3 | 1 | 0 |
| POLR2A | 1 | 1 |
| RALGPS2 | 1 | 0 |
| SLC6A21 | 1 | 0 |
| UPP1 | 1 | 0 |
| SLC41A3 | 1 | 1 |
| CALM4 | 2 | 1 |
| HEG1 | 1 | 1 |
| ULK4 | 1 | 1 |
| IFI202 | 5 | 1 |
| KRT6A;KRT76 | 1 | 1 |
| HRNR | 5 | 1 |
| CD177 | 2 | 1 |
| PPP1R15A | 1 | 1 |
| DSC1 | 1 | 1 |
| ALDH3B2 | 1 | 1 |
| S100A9 | 5 | 1 |
| GRP | 1 | 1 |
| LTF | 14 | 1 |
| NRDE2 | 1 | 1 |
| MEP1A | 1 | 1 |
| NGP | 6 | |
| SERPINB5 | 11 | |
| IGKV4-63 | 1 | |
| MSH2 | 1 | |
| DSG1B | 1 | |
| ITPR1 | 1 | |
| CHIL3 | 7 | |
| WTAP | 1 | |
| NEU2 | 1 | |
| INTS7 | 1 | |

| | |
|---|---|
| URAH | 2 |
| GM32742 | 1 |
| POLQ | 1 |
| LCN6 | 1 |
| DOLK | 1 |
| MPO | 12 |
| FLG2 | 3 |
| NCCRP1 | 1 |
| ABCA6 | 1 |
| KRT13 | 2 |
| CFAP47 | 1 |
| TRIM66 | 1 |
| NKTR | 1 |
| FBXL5 | 1 |
| CCHCR1 | 1 |
| TMEM119 | 1 |
| RUFY2 | 1 |
| TACR3 | 1 |
| S100A8 | 1 |
| KRT84 | 3 |
| A1BG | 2 |
| HELQ | 1 |
| SAA1 | 1 |
| ENDOU | 7 |
| KRT33B | 1 |
| sp\|P01654\|KV3A1_M | 1 |
| H2-D1;H2-T23 | 1 |
| IGKV4-57 | 1 |
| LYZ2 | 1 |
| MPZ | 1 |
| POF1B | 6 |
| ORM2 | 1 |
| CFAP57 | 1 |

| | |
|---|---|
| KRT6B | 5 |
| AHNAK2 | 1 |
| KRT36 | 1 |
| KRT35;KRT34 | 1 |
| PKHD1L1 | 1 |
| FLG | 3 |
| LCN2 | 3 |
| KRT6A;KRT79 | 1 |
| IGHV1-18;IGHV1-22; | 1 |
| GM5478;GM5414 | 1 |
| SERPINB3B | 1 |
| LMNB1 | 3 |
| ZFP60 | 1 |
| H1-5 | 2 |
| KRT24 | 6 |
| TREX2 | 3 |
| IGHV9-3 | 3 |
| OBP1B | 1 |
| CAMP | 2 |
| LGALS7 | 6 |
| SERPINB2 | 4 |
| TRIM29 | 4 |
| DSG1A | 1 |
| SBSN | 2 |
| IGKV1-135;IGKV1-13 | 1 |
| COL1A1 | 6 |
| MMP9 | 2 |
| FLG | 2 |
| CDH22 | 1 |
| SERPINB6D | 1 |
| IGHV1-20;IGHV1-37 | 1 |
| GM1553 | 1 |
| IGHG2B | 2 |

| | |
|---|---|
| SUN2 | 1 |
| ITGB2L | 1 |
| PRTN3 | 1 |
| CYBB | 1 |
| IGKV17-127;IGKV17- | 1 |
| IGHV5-9-1 | 1 |
| IGHV5-12 | 2 |
| KRT1;KRT2;KRT77 | 1 |
| ELANE | 1 |
| DSC3 | 1 |
| PPL | 2 |
| KRT10 | 1 |
| KRT19;KRT15 | 1 |
| OLFM4 | 1 |
| CSTDC5;CSTDC6 | 1 |
| OBP1A | 1 |
| PIP | 2 |
| LY6D | 1 |
| DSG3 | 1 |
| KRT13;KRT15 | 1 |
| CDSN | 1 |
