## Supplementary material for "SFRP1 upregulation causes hippocampal synaptic dysfunction and memory impairment": Suppl Table 3a

| gene_symbols_or_id | unique_peptides | peptides_used_for_dea_contrast: Control_Hom vs SFRP1-TG_Hom | foldchange.log2_msqrob_contrast: Control_Hom vs SFRP1-TG_Hom | pvalue_msqrob_contrast: Control_Hom vs SFRP1-TG_Hom | qvalue_msqrob_contrast: Control_Hom vs SFRP1-TG_Hom |
|---|---|---|---|---|---|
| WDR7 | 66 | 54 | 0,120255866 | 7,62048E-08 | 0,00045601 |
| EIF3B | 22 | 19 | -0,125243027 | 2,23629E-07 | 0,000669097 |
| THBS4 | 6 | 6 | -0,467788995 | 4,61938E-07 | 0,000691059 |
| EIF4G3 | 24 | 20 | -0,196263651 | 3,59799E-07 | 0,000691059 |
| SLC9A3R1 | 19 | 17 | -0,142462989 | 1,17959E-06 | 0,001176448 |
| ACSBG1 | 34 | 30 | -0,12418982 | 1,1318E-06 | 0,001176448 |
| SPARCL1 | 15 | 14 | -0,238959877 | 1,51544E-06 | 0,001295482 |
| DLGAP3 | 21 | 13 | 0,146762452 | 4,24867E-06 | 0,003178003 |
| FAM169A | 12 | 10 | -0,187363708 | 6,18064E-06 | 0,004109438 |
| CYGB | 9 | 8 | -0,20695318 | 6,92825E-06 | 0,004145862 |
| DBNL | 23 | 21 | -0,113629418 | 7,64383E-06 | 0,004158242 |
| DMXL2 | 123 | 90 | 0,127453901 | 9,19276E-06 | 0,004584121 |
| SF1 | 6 | 5 | -0,246671886 | 1,25576E-05 | 0,005780381 |
| EWSR1 | 5 | 3 | -0,315469415 | 1,42004E-05 | 0,006069665 |
| MTREX | 1 | 1 | -0,624173862 | 1,79252E-05 | 0,007150944 |
| EPB41L2 | 37 | 33 | -0,114756452 | 2,33812E-05 | 0,008744565 |
| EIF3E | 26 | 20 | -0,102318604 | 3,31247E-05 | 0,011659897 |
| CNN3 | 10 | 6 | -0,171896623 | 4,30375E-05 | 0,014307585 |
| CSNK1A1 | 1 | 1 | -0,493397456 | 7,6421E-05 | 0,02304827 |
| ZC3HC1 | 2 | 2 | -0,452927809 | 7,7033E-05 | 0,02304827 |
| LRRC40 | 9 | 9 | -0,187859864 | 9,04188E-05 | 0,02576504 |
| SEPHS1 | 4 | 4 | -0,245909761 | 0,000114878 | 0,02988838 |
| NSF | 72 | 62 | 0,069607053 | 0,000110188 | 0,02988838 |
| PLXNB1 | 28 | 19 | -0,097516199 | 0,000135618 | 0,033813983 |
| LZIC | 7 | 6 | -0,192442936 | 0,000150287 | 0,034589171 |
| OMG | 10 | 9 | -0,116609773 | 0,000144862 | 0,034589171 |
| OSCP1 | 9 | 8 | -0,188856938 | 0,000172897 | 0,038319131 |
| STRN4 | 16 | 13 | -0,098717079 | 0,000207828 | 0,044415717 |

| | | | | | |
|---|---|---|---|---|---|
| EIF3L | 23 | 21 | -0,092949931 | 0,000233516 | 0,04818485 |
| UFL1 | 5 | 3 | -0,192468626 | 0,000253472 | 0,050559171 |
| KCMF1 | 3 | 3 | -0,221091389 | 0,000291212 | 0,05428284 |
| sp\|Q8C3W1\|CA198_M | 9 | 7 | -0,169926328 | 0,000291064 | 0,05428284 |
| HOOK3 | 21 | 17 | -0,165313537 | 0,000299354 | 0,05428284 |
| AP2M1 | 35 | 29 | 0,068042544 | 0,000321299 | 0,054932979 |
| IQSEC3 | 16 | 13 | 0,129763155 | 0,000317439 | 0,054932979 |
| NRBP1 | 7 | 7 | -0,144710707 | 0,000372671 | 0,060271975 |
| KIF3A | 17 | 16 | -0,099114718 | 0,000365486 | 0,060271975 |
| HNRNPC | 6 | 6 | -0,274203818 | 0,000385113 | 0,060645217 |
| USP14 | 27 | 25 | -0,087974043 | 0,000411581 | 0,060987046 |
| HK1 | 61 | 46 | 0,098246142 | 0,000399559 | 0,060987046 |
| BRINP1 | 15 | 6 | 0,128990682 | 0,000417859 | 0,060987046 |
| NGEF | 14 | 12 | -0,188624519 | 0,000500434 | 0,066546547 |
| GJA1 | 19 | 17 | -0,177280265 | 0,000488943 | 0,066546547 |
| TCEA1 | 13 | 11 | -0,114528393 | 0,000494088 | 0,066546547 |
| NDUFS2 | 27 | 24 | 0,114105553 | 0,000499673 | 0,066546547 |
| PITPNC1 | 9 | 6 | -0,212813443 | 0,000514026 | 0,066868141 |
| EIF3F | 15 | 11 | -0,127944511 | 0,000549926 | 0,068142077 |
| SF3B3 | 11 | 9 | -0,111850298 | 0,000555032 | 0,068142077 |
| ACAP3 | 2 | 1 | 0,342315183 | 0,000557982 | 0,068142077 |
| CLU | 16 | 16 | -0,144973642 | 0,00059579 | 0,071304103 |
| PTPRZ1 | 34 | 30 | -0,071470667 | 0,00070435 | 0,08264371 |
| GNL1 | 13 | 11 | 0,223949645 | 0,000742104 | 0,08539906 |
| MTSS2 | 6 | 5 | -0,232943441 | 0,000770711 | 0,085406206 |
| PEF1 | 6 | 5 | -0,156170573 | 0,000763347 | 0,085406206 |
| MARCKSL1 | 2 | 2 | -0,414277281 | 0,000850045 | 0,092484945 |
| RASGRF1 | 18 | 16 | -0,092438432 | 0,000869404 | 0,092902063 |
| CAMK2A | 16 | 15 | 0,096532612 | 0,000890442 | 0,093480783 |
| DTNA | 9 | 6 | -0,171089121 | 0,000907215 | 0,093599512 |
| TBC1D15 | 8 | 6 | -0,174298656 | 0,000949422 | 0,094689054 |
| SEC31A | 30 | 24 | -0,132107361 | 0,000933943 | 0,094689054 |
| FUBP1 | 3 | 2 | -0,273484606 | 0,001099262 | 0,107835784 |

| | | | | | |
|---|---|---|---|---|---|
| SNRPA1 | 1 | 1 | -0,332308631 | 0,001222782 | 0,110865603 |
| STXBP6 | 4 | 3 | -0,262895615 | 0,001169347 | 0,110865603 |
| CSE1L | 36 | 31 | -0,105772452 | 0,001191088 | 0,110865603 |
| EIF3H | 12 | 9 | -0,105546575 | 0,001220076 | 0,110865603 |
| MCUB | 3 | 1 | 0,45025026 | 0,001168067 | 0,110865603 |
| HDGFL2 | 8 | 5 | -0,154161898 | 0,001285625 | 0,111495328 |
| NF1 | 13 | 12 | -0,107818765 | 0,001282073 | 0,111495328 |
| EPB41L1 | 62 | 56 | -0,050031292 | 0,001253869 | 0,111495328 |
| CACNG3 | 4 | 1 | -0,338788189 | 0,001311761 | 0,112136803 |
| GABBR1 | 15 | 9 | 0,119143 | 0,001357329 | 0,114397988 |
| FERMT2 | 15 | 12 | -0,142721024 | 0,001404764 | 0,116751509 |
| FUS | 9 | 7 | -0,164082363 | 0,001598634 | 0,131044191 |
| RPLP0 | 13 | 12 | -0,107781442 | 0,001706325 | 0,137981734 |
| BABAM2 | 6 | 5 | -0,142221899 | 0,001792937 | 0,140877894 |
| CNKSR2 | 35 | 23 | 0,076741907 | 0,001812767 | 0,140877894 |
| NT5DC3 | 30 | 24 | 0,122668466 | 0,001799308 | 0,140877894 |
| GORASP2 | 9 | 9 | -0,219453077 | 0,001962215 | 0,147343087 |
| NCK1 | 5 | 3 | -0,146236 | 0,001969827 | 0,147343087 |
| MYO18A | 70 | 49 | 0,050071286 | 0,001969459 | 0,147343087 |
| PPP3CA | 1 | 1 | -0,295900764 | 0,002019036 | 0,149159437 |
| SFXN3 | 15 | 12 | 0,079087955 | 0,002048324 | 0,149477707 |
| HNRNPK | 22 | 20 | -0,175091417 | 0,002115632 | 0,151734357 |
| CPNE4 | 18 | 14 | 0,133219389 | 0,002129961 | 0,151734357 |
| AP1S2 | 2 | 2 | -0,197191943 | 0,002235141 | 0,157353943 |
| USP47 | 26 | 20 | -0,071776405 | 0,002322261 | 0,161586183 |
| EIF2S2 | 9 | 8 | -0,092917808 | 0,002371412 | 0,163109535 |
| IQSEC1 | 43 | 31 | 0,05752237 | 0,00243589 | 0,165296307 |
| TOMM70 | 28 | 23 | 0,104370296 | 0,002458451 | 0,165296307 |
| DOCK10 | 11 | 6 | -0,159206086 | 0,002520076 | 0,167215583 |
| UBL4A | 7 | 7 | -0,132470158 | 0,002598772 | 0,167215583 |
| CCDC177 | 19 | 12 | -0,101828321 | 0,002555089 | 0,167215583 |
| PCK2 | 23 | 16 | 0,174894928 | 0,002584378 | 0,167215583 |
| NASP | 2 | 2 | -0,184058311 | 0,002734567 | 0,174081378 |

| | | | | | |
|---|---|---|---|---|---|
| FAM120A | 23 | 21 | -0,080912811 | 0,002829147 | 0,178206502 |
| CLTC | 130 | 110 | 0,084760156 | 0,003013218 | 0,185155388 |
| NRP1 | 16 | 11 | 0,095060804 | 0,003032291 | 0,185155388 |
| VDAC1 | 23 | 21 | 0,114371118 | 0,002994744 | 0,185155388 |
| VPS37A | 1 | 1 | -0,424615144 | 0,003183297 | 0,192412629 |
| IMMT | 55 | 47 | 0,093430113 | 0,003225333 | 0,193003938 |
| SRSF1 | 3 | 1 | -0,349717553 | 0,003529538 | 0,205055872 |
| EFHD2 | 20 | 19 | -0,127474613 | 0,003523874 | 0,205055872 |
| COX6A1 | 3 | 2 | 0,194998629 | 0,003516093 | 0,205055872 |
| BAG6 | 22 | 17 | -0,125198748 | 0,003840417 | 0,214729388 |
| SRPK2 | 11 | 9 | -0,105738838 | 0,003905855 | 0,214729388 |
| ASS1 | 15 | 12 | -0,092807302 | 0,00374116 | 0,214729388 |
| ALDH1B1 | 21 | 14 | 0,070791887 | 0,003801219 | 0,214729388 |
| MFN2 | 31 | 16 | 0,087827152 | 0,003876152 | 0,214729388 |
| SLC35A4 | 1 | 1 | 0,318583968 | 0,003911347 | 0,214729388 |
| GLS | 11 | 9 | 0,123527614 | 0,004248362 | 0,231110907 |
| SHROOM2 | 13 | 10 | -0,174953772 | 0,00442949 | 0,231527011 |
| EIF3G | 6 | 6 | -0,155472683 | 0,004466287 | 0,231527011 |
| HNRNPUL2 | 29 | 23 | -0,128676104 | 0,004456665 | 0,231527011 |
| SEC23IP | 18 | 13 | -0,09683008 | 0,004396033 | 0,231527011 |
| SHANK1 | 52 | 37 | 0,109300139 | 0,004321803 | 0,231527011 |
| GLIPR2 | 2 | 2 | 0,182702984 | 0,004488157 | 0,231527011 |
| SUPT6H | 4 | 2 | -0,166077929 | 0,004917588 | 0,236312853 |
| DNAJB4 | 8 | 5 | -0,148790916 | 0,004831643 | 0,236312853 |
| SNX3 | 6 | 6 | -0,119621379 | 0,004936348 | 0,236312853 |
| APLP1 | 12 | 8 | -0,117229043 | 0,00489154 | 0,236312853 |
| NAPB | 26 | 25 | 0,049467813 | 0,004652464 | 0,236312853 |
| TECPR1 | 20 | 15 | 0,073253851 | 0,004879774 | 0,236312853 |
| ADAM11 | 10 | 8 | 0,105853221 | 0,004782292 | 0,236312853 |
| ME1 | 29 | 26 | 0,171981466 | 0,004891577 | 0,236312853 |
| FAM177A1 | 4 | 2 | 0,323912012 | 0,004809678 | 0,236312853 |
| ADCYAP1R1 | 3 | 2 | -0,22861038 | 0,005105456 | 0,238680051 |
| SNX27 | 18 | 14 | -0,076986913 | 0,005055283 | 0,238680051 |

| | | | | | |
|---|---|---|---|---|---|
| ATP6V0A1 | 43 | 37 | 0,065453182 | 0,005092297 | 0,238680051 |
| POLR2H | 7 | 6 | -0,140848619 | 0,005265573 | 0,242223657 |
| WDR91 | 7 | 7 | -0,123965873 | 0,005232385 | 0,242223657 |
| ARIH1 | 5 | 5 | -0,109842293 | 0,00530269 | 0,242223657 |
| ZCCHC8 | 1 | 1 | -0,318134032 | 0,005408179 | 0,245170767 |
| RPS6KA1 | 5 | 5 | 0,170999878 | 0,005620903 | 0,252898361 |
| EIF4G1 | 1 | 1 | -0,286821415 | 0,0057515 | 0,25293484 |
| HNRNPF | 2 | 2 | -0,199680685 | 0,005746061 | 0,25293484 |
| ENTPD2 | 9 | 7 | -0,180084723 | 0,005790788 | 0,25293484 |
| CIAO2B | 3 | 2 | -0,179050639 | 0,005768824 | 0,25293484 |
| ATXN2L | 18 | 16 | -0,143463204 | 0,006028025 | 0,259508663 |
| CAMK2D | 14 | 10 | -0,138364024 | 0,00600397 | 0,259508663 |
| ERC2 | 45 | 29 | 0,071865474 | 0,006278547 | 0,268363038 |
| SYT12 | 13 | 11 | 0,100810418 | 0,006417942 | 0,272375632 |
| MOB1B;MOB1A | 2 | 2 | -0,18466344 | 0,00658096 | 0,277327202 |
| PCBP1 | 12 | 12 | -0,112081233 | 0,006720704 | 0,277889513 |
| GK | 19 | 15 | 0,08123898 | 0,00673362 | 0,277889513 |
| OGDH | 50 | 44 | 0,098325814 | 0,00670651 | 0,277889513 |
| R3HDM2 | 5 | 4 | -0,142435407 | 0,006887075 | 0,282275749 |
| GNG4 | 1 | 1 | -0,341163624 | 0,007231567 | 0,284695355 |
| HAPLN1 | 21 | 17 | -0,212080096 | 0,007111806 | 0,284695355 |
| YTHDF2 | 3 | 2 | -0,172214073 | 0,007144611 | 0,284695355 |
| MPP2 | 22 | 19 | -0,076691436 | 0,007069904 | 0,284695355 |
| MYH10;MYH14 | 2 | 2 | 0,18189149 | 0,007127265 | 0,284695355 |
| PPIP5K1;PPIP5K2 | 1 | 1 | 0,34567529 | 0,007204584 | 0,284695355 |
| PCCB | 28 | 16 | 0,145838337 | 0,007398168 | 0,289350551 |
| SERPINA3K | 23 | 16 | -0,628368383 | 0,007674831 | 0,292328254 |
| MAPT | 6 | 5 | -0,263536663 | 0,007955348 | 0,292328254 |
| URGCP | 3 | 1 | -0,258110125 | 0,007807214 | 0,292328254 |
| SEC13 | 3 | 3 | -0,12983937 | 0,00758019 | 0,292328254 |
| ATXN10 | 23 | 16 | -0,09171985 | 0,008060522 | 0,292328254 |
| VARS1 | 31 | 29 | -0,051826132 | 0,007857902 | 0,292328254 |
| NDUFV1 | 26 | 21 | 0,124745616 | 0,007726999 | 0,292328254 |

| | | | | | |
|---|---|---|---|---|---|
| PRUNE2 | 4 | 2 | 0,13162616 | 0,007886268 | 0,292328254 |
| ALDH1A1 | 19 | 18 | 0,182348799 | 0,008005411 | 0,292328254 |
| ZCCHC7 | 1 | 1 | 0,267962677 | 0,008050031 | 0,292328254 |
| PLCH2;PLCH1 | 2 | 1 | 0,347648246 | 0,007755223 | 0,292328254 |
| KRT6A | 19 | 1 | 0,655317266 | 0,008045386 | 0,292328254 |
| NDUFS7 | 9 | 7 | 0,102204857 | 0,00812861 | 0,293021699 |
| OLFR1234 | 1 | 1 | 0,427819051 | 0,00824313 | 0,295370592 |
| AGO2 | 12 | 10 | -0,115063712 | 0,008338998 | 0,297027161 |
| MADD | 47 | 37 | 0,04890688 | 0,008538234 | 0,302324228 |
| NT5C1A | 1 | 1 | -0,342908422 | 0,008845398 | 0,306576332 |
| DYNLT1 | 1 | 1 | -0,282369181 | 0,008761122 | 0,306576332 |
| GPRC5B | 3 | 1 | -0,232784529 | 0,008989143 | 0,306576332 |
| UCHL5 | 5 | 3 | -0,142595427 | 0,008992288 | 0,306576332 |
| USO1 | 36 | 32 | -0,103816885 | 0,009016951 | 0,306576332 |
| EIF4G1 | 22 | 21 | -0,094746221 | 0,008754855 | 0,306576332 |
| PA2G4 | 20 | 17 | -0,09084668 | 0,008892125 | 0,306576332 |
| SCFD1 | 5 | 3 | -0,154997831 | 0,009217801 | 0,309883812 |
| GUCY1B1 | 18 | 14 | -0,085829143 | 0,009215046 | 0,309883812 |
| PROM1 | 4 | 1 | -0,368782272 | 0,009694727 | 0,313066118 |
| DPY19L4 | 1 | 1 | -0,284314245 | 0,009783316 | 0,313066118 |
| RBFOX3;RBFOX1 | 5 | 2 | -0,144892988 | 0,009546391 | 0,313066118 |
| MSTO1 | 3 | 2 | -0,142216368 | 0,009770564 | 0,313066118 |
| ACBD3 | 6 | 5 | -0,112968444 | 0,00973596 | 0,313066118 |
| UBE3A | 26 | 21 | -0,063756431 | 0,009554215 | 0,313066118 |
| SYNGAP1 | 74 | 51 | 0,104998258 | 0,009419002 | 0,313066118 |
| RIMS1 | 12 | 9 | 0,116917657 | 0,009666443 | 0,313066118 |
| CISD1 | 9 | 7 | 0,117424665 | 0,009781454 | 0,313066118 |
| TRIM32 | 6 | 5 | -0,138268259 | 0,009899624 | 0,315102929 |
| MRPL11 | 6 | 2 | 0,285712116 | 0,009987265 | 0,316210536 |
| USP10 | 9 | 9 | -0,099642491 | 0,010069862 | 0,317147657 |
| PKIA | 1 | 1 | -0,487347285 | 0,010192982 | 0,319026885 |
| NPM1 | 10 | 8 | -0,270651356 | 0,010236157 | 0,319026885 |
| NCDN | 35 | 32 | -0,171903295 | 0,01030652 | 0,319555523 |

| | | | | | |
|---|---|---|---|---|---|
| PCCA | 41 | 31 | 0,095270763 | 0,01050079 | 0,323900667 |
| LAMP5 | 2 | 2 | -0,154518211 | 0,01063849 | 0,326465243 |
| DLD | 22 | 19 | 0,096863665 | 0,010915958 | 0,333270886 |
| PSMD4 | 8 | 8 | -0,08422051 | 0,011006367 | 0,334325376 |
| MRPS21 | 2 | 1 | 0,388190201 | 0,011113988 | 0,335889403 |
| RGS10 | 4 | 4 | -0,144131545 | 0,01128583 | 0,339368889 |
| MTRES1 | 2 | 1 | -0,258604695 | 0,011562588 | 0,345952639 |
| EIF4E2 | 1 | 1 | -0,313988684 | 0,011644463 | 0,346668993 |
| PAK3 | 11 | 7 | -0,112228241 | 0,011819435 | 0,350136144 |
| G3BP2 | 16 | 12 | -0,159426367 | 0,011972676 | 0,352928527 |
| MADD | 3 | 1 | -0,231227026 | 0,012043068 | 0,353263314 |
| ISCA2 | 6 | 5 | 0,125933192 | 0,012290467 | 0,357020166 |
| PDCD2 | 1 | 1 | 0,249830796 | 0,012242411 | 0,357020166 |
| KIF21B | 1 | 1 | -0,36849329 | 0,012426295 | 0,359221982 |
| KBTBD11 | 18 | 15 | -0,087527797 | 0,012614396 | 0,362906466 |
| ATP5F1C | 20 | 15 | 0,083078814 | 0,012703463 | 0,363720217 |
| CDC42EP4 | 10 | 9 | -0,142714855 | 0,012879087 | 0,364553674 |
| UPF2 | 7 | 3 | -0,126360763 | 0,012818606 | 0,364553674 |
| HSPD1 | 52 | 39 | 0,095910337 | 0,012915337 | 0,364553674 |
| CRACDL | 20 | 14 | -0,128987723 | 0,013003767 | 0,365326491 |
| RANGAP1 | 11 | 11 | -0,123603334 | 0,013119062 | 0,366843294 |
| TBC1D17 | 11 | 10 | -0,072916662 | 0,013314596 | 0,367163788 |
| ABHD12 | 17 | 13 | -0,066484192 | 0,013267685 | 0,367163788 |
| ATP1A3 | 49 | 44 | 0,056563736 | 0,013273547 | 0,367163788 |
| ARHGEF12 | 19 | 15 | -0,080062574 | 0,013769275 | 0,377960295 |
| SMPDL3B | 3 | 1 | -0,310429211 | 0,013984582 | 0,380231348 |
| BZW1 | 12 | 10 | -0,191273554 | 0,014042635 | 0,380231348 |
| CYB5R3 | 17 | 9 | 0,096647941 | 0,013974837 | 0,380231348 |
| CES1C;CES1D | 1 | 1 | 0,318214249 | 0,014210437 | 0,383041694 |
| GPC4 | 12 | 7 | -0,095305896 | 0,014346018 | 0,384962199 |
| PACSIN1 | 43 | 30 | 0,044194614 | 0,014455401 | 0,386165716 |
| DYNC1I2 | 9 | 9 | -0,10278803 | 0,014665034 | 0,390024726 |
| CDK17 | 5 | 3 | -0,150092245 | 0,015148938 | 0,399344693 |

| | | | | | |
|---|---|---|---|---|---|
| NDUFC2 | 8 | 7 | 0,08562264 | 0,015127953 | 0,399344693 |
| PLCH2 | 27 | 21 | 0,133473728 | 0,015296648 | 0,401469931 |
| PPP4R3A | 8 | 5 | -0,107393551 | 0,01548171 | 0,404552644 |
| SRI | 6 | 4 | -0,122831446 | 0,016118497 | 0,40473234 |
| PITPNM2 | 21 | 15 | -0,097693383 | 0,016246468 | 0,40473234 |
| RPL11 | 6 | 6 | -0,091728612 | 0,015921201 | 0,40473234 |
| CRKL | 14 | 10 | -0,079803278 | 0,016018808 | 0,40473234 |
| ASAP1 | 20 | 17 | -0,071409812 | 0,015643941 | 0,40473234 |
| ATP1B1 | 19 | 16 | 0,045035946 | 0,015670722 | 0,40473234 |
| RYR2 | 19 | 13 | 0,079679263 | 0,015958324 | 0,40473234 |
| SLC25A12;SLC25A13 | 6 | 5 | 0,109985053 | 0,016300216 | 0,40473234 |
| NDUFA4 | 6 | 6 | 0,125190411 | 0,015798543 | 0,40473234 |
| AGK | 26 | 16 | 0,127298784 | 0,016190336 | 0,40473234 |
| CA4 | 7 | 6 | 0,138441381 | 0,01573846 | 0,40473234 |
| ANKS1B | 3 | 2 | 0,138714229 | 0,016049059 | 0,40473234 |
| DGKE | 9 | 5 | -0,120802142 | 0,016520032 | 0,408495339 |
| DDX19A | 8 | 5 | -0,100271645 | 0,016642397 | 0,409827577 |
| ATAD1 | 11 | 5 | 0,126540941 | 0,016783131 | 0,409919423 |
| UQCR10 | 3 | 3 | 0,127069798 | 0,016728391 | 0,409919423 |
| IREB2 | 2 | 2 | -0,173890826 | 0,016866581 | 0,410283012 |
| SNX10 | 2 | 1 | -0,2431039 | 0,017047448 | 0,41300377 |
| EIF3M | 8 | 4 | -0,144881531 | 0,017593079 | 0,420251732 |
| MTMR12 | 6 | 4 | -0,135123394 | 0,017433572 | 0,420251732 |
| GRM2 | 15 | 9 | -0,099976611 | 0,017561421 | 0,420251732 |
| AP2A2 | 54 | 50 | 0,072964953 | 0,017627538 | 0,420251732 |
| CDK18 | 4 | 4 | -0,119714193 | 0,017817301 | 0,421417912 |
| ADCY5;ADCY6 | 2 | 2 | 0,153484529 | 0,017748469 | 0,421417912 |
| MYO9A | 1 | 1 | -0,211750549 | 0,017996447 | 0,422796545 |
| SPAG9 | 29 | 27 | -0,096951639 | 0,018016898 | 0,422796545 |
| VAMP2 | 4 | 4 | 0,084039175 | 0,018231539 | 0,424503999 |
| ACO2 | 53 | 42 | 0,089629661 | 0,018229688 | 0,424503999 |
| COX20 | 2 | 1 | 0,233736739 | 0,01909478 | 0,442880487 |
| AMFR | 3 | 1 | -0,27155031 | 0,019530581 | 0,44607251 |

| | | | | | |
|---|---|---|---|---|---|
| PHACTR1;RAI14 | 1 | 1 | -0,221940139 | 0,019432238 | 0,44607251 |
| BCAT1 | 11 | 10 | 0,100420944 | 0,019519974 | 0,44607251 |
| ANKRD27 | 1 | 1 | 0,262240344 | 0,019353485 | 0,44607251 |
| VDAC3 | 12 | 10 | 0,100053017 | 0,019705263 | 0,448350928 |
| SLC8A1 | 30 | 18 | 0,068542085 | 0,019804789 | 0,448908542 |
| CBLN4 | 1 | 1 | -0,219499364 | 0,019980089 | 0,451173022 |
| BAX | 8 | 8 | -0,076469163 | 0,020094025 | 0,452040012 |
| SEPTIN8 | 30 | 27 | 0,044573727 | 0,020170602 | 0,452063219 |
| PIP5K1A | 10 | 8 | -0,140140652 | 0,020389064 | 0,453561939 |
| APPL2 | 15 | 14 | -0,087179091 | 0,02035203 | 0,453561939 |
| MRTFB;MRTFA;MYOCD | 1 | 1 | -0,185554675 | 0,020482809 | 0,453959737 |
| PREPL | 18 | 16 | -0,08360013 | 0,020574003 | 0,454298281 |
| PRKAR1A | 17 | 14 | -0,069120845 | 0,020824436 | 0,45813759 |
| EXTL2 | 4 | 3 | -0,164183954 | 0,020940751 | 0,459008994 |
| S1PR1 | 4 | 2 | -0,176558199 | 0,021261537 | 0,459364744 |
| DDX6 | 14 | 12 | -0,059764618 | 0,02109682 | 0,459364744 |
| PHB2 | 20 | 17 | 0,072750532 | 0,021159566 | 0,459364744 |
| ATP5A1 | 50 | 41 | 0,0821861 | 0,021264043 | 0,459364744 |
| HSD11B1 | 3 | 1 | -0,303578005 | 0,021856963 | 0,469907149 |
| LAGE3 | 1 | 1 | -0,244632383 | 0,022144689 | 0,469907149 |
| PCBP3 | 6 | 5 | -0,147997331 | 0,022008467 | 0,469907149 |
| DPYSL4 | 28 | 24 | -0,09607865 | 0,022120897 | 0,469907149 |
| IPO5 | 35 | 28 | -0,075880815 | 0,021999238 | 0,469907149 |
| DCXR | 3 | 3 | -0,178758592 | 0,022464776 | 0,470898429 |
| FN3K | 5 | 4 | -0,122096246 | 0,022504165 | 0,470898429 |
| CHAT | 6 | 6 | -0,088109408 | 0,022506175 | 0,470898429 |
| ZZEF1 | 18 | 13 | -0,079370857 | 0,022448566 | 0,470898429 |
| OGFR | 9 | 7 | -0,144159619 | 0,022699203 | 0,473282341 |
| AACS | 11 | 10 | -0,10450576 | 0,022955126 | 0,475306131 |
| RAB23 | 10 | 7 | -0,080709332 | 0,022884488 | 0,475306131 |
| TXNDC17 | 3 | 3 | 0,167768111 | 0,023222954 | 0,479193637 |
| NOVA2 | 4 | 3 | -0,142802628 | 0,023802368 | 0,48392969 |
| SEC16A | 11 | 7 | -0,113991796 | 0,023787295 | 0,48392969 |

| | | | | | |
|---|---|---|---|---|---|
| HNRNPH2 | 6 | 6 | -0,09640557 | 0,023647364 | 0,48392969 |
| FH | 29 | 22 | 0,083068546 | 0,023625273 | 0,48392969 |
| NDUFA12 | 15 | 11 | 0,083911367 | 0,023856828 | 0,48392969 |
| SUMO2 | 1 | 1 | -0,237154245 | 0,024153673 | 0,488295867 |
| ARL5A | 2 | 1 | -0,20713892 | 0,024399306 | 0,488312531 |
| CDC37L1 | 3 | 2 | -0,123354561 | 0,024240099 | 0,488312531 |
| SYNGAP1;DAB2IP | 2 | 2 | 0,177739566 | 0,024342641 | 0,488312531 |
| ANKRD34A | 6 | 6 | -0,138686555 | 0,024729986 | 0,493280786 |
| NDUFA5 | 8 | 6 | 0,113092263 | 0,024854822 | 0,494123775 |
| CDC42BPA | 15 | 13 | 0,076947694 | 0,025168397 | 0,498700955 |
| RAPH1 | 1 | 1 | -0,454585793 | 0,025594212 | 0,49984448 |
| GABPA | 1 | 1 | -0,383453241 | 0,025708604 | 0,49984448 |
| CHMP7 | 7 | 5 | -0,204355348 | 0,025998039 | 0,49984448 |
| TSC22D1 | 4 | 3 | -0,19350804 | 0,025825519 | 0,49984448 |
| PCBP2 | 5 | 5 | -0,120273917 | 0,025676934 | 0,49984448 |
| CYP46A1 | 21 | 14 | -0,07100823 | 0,02606141 | 0,49984448 |
| MPP3 | 8 | 6 | 0,071280042 | 0,026015556 | 0,49984448 |
| GRIN2B | 39 | 23 | 0,072327756 | 0,025371839 | 0,49984448 |
| SYT1 | 32 | 28 | 0,076157724 | 0,025471383 | 0,49984448 |
| ITIH2 | 2 | 1 | 0,278217959 | 0,025816901 | 0,49984448 |
| YBX1;YBX3 | 3 | 2 | -0,188349786 | 0,026921116 | 0,505003008 |
| SERBP1 | 15 | 14 | -0,10220851 | 0,02678391 | 0,505003008 |
| USP8 | 9 | 6 | -0,101413098 | 0,026690245 | 0,505003008 |
| NSFL1C | 24 | 22 | -0,043369991 | 0,026835954 | 0,505003008 |
| SCO1 | 2 | 2 | 0,171583054 | 0,026582908 | 0,505003008 |
| WDR41 | 6 | 4 | 0,178834991 | 0,026916257 | 0,505003008 |
| MAPK10;MAPK8;MAPK | 1 | 1 | 0,219515336 | 0,026804531 | 0,505003008 |
| ERC1 | 17 | 8 | 0,08997224 | 0,027213995 | 0,506315131 |
| SAT2 | 1 | 1 | 0,216739851 | 0,027076655 | 0,506315131 |
| FAM184B | 1 | 1 | 0,273206493 | 0,027244898 | 0,506315131 |
| NACA | 8 | 6 | -0,079666487 | 0,027623391 | 0,51175967 |
| SORBS1 | 2 | 1 | -0,300667552 | 0,027823437 | 0,513874843 |
| UPF1 | 26 | 23 | -0,047295804 | 0,028036506 | 0,514884582 |

| | | | | | |
|---|---|---|---|---|---|
| ADCY9 | 31 | 20 | 0,097642923 | 0,028050196 | 0,514884582 |
| UIMC1 | 1 | 1 | 0,240177631 | 0,028300602 | 0,517892369 |
| PDE1B | 18 | 16 | -0,133488331 | 0,028642846 | 0,51956937 |
| KALRN;TRIO | 8 | 7 | 0,072499874 | 0,028652723 | 0,51956937 |
| OPA1 | 62 | 38 | 0,075200211 | 0,028509855 | 0,51956937 |
| ALDH3A2 | 1 | 1 | 0,219444242 | 0,028802689 | 0,520710844 |
| GPD2 | 56 | 47 | 0,077932015 | 0,028927322 | 0,521388831 |
| AGT | 4 | 3 | -0,171965352 | 0,029014705 | 0,521393382 |
| MICU3 | 21 | 15 | 0,08781774 | 0,029121406 | 0,521744 |
| ZYG11B | 8 | 5 | -0,098788706 | 0,029230508 | 0,522135402 |
| MFSD6 | 3 | 2 | 0,246244937 | 0,029405205 | 0,523692693 |
| MICAL1 | 4 | 4 | -0,118012035 | 0,029501736 | 0,52385278 |
| HLCS | 2 | 1 | -0,318846519 | 0,029678289 | 0,523878708 |
| TIMM10 | 4 | 3 | 0,126676407 | 0,029635296 | 0,523878708 |
| ATP6V1B2 | 26 | 24 | 0,049633421 | 0,029780005 | 0,524128085 |
| RHOF | 2 | 2 | 0,196086043 | 0,030078504 | 0,526611292 |
| SIDT2 | 1 | 1 | 0,254596549 | 0,030097103 | 0,526611292 |
| MOB2 | 1 | 1 | -0,188877534 | 0,030716612 | 0,535883991 |
| PUF60 | 8 | 6 | -0,133982686 | 0,030883273 | 0,537225311 |
| PHYHIPL | 16 | 15 | -0,085498232 | 0,031235523 | 0,541777888 |
| CSRP1 | 10 | 7 | -0,118862158 | 0,031763802 | 0,541949268 |
| CIAPIN1 | 5 | 4 | -0,105329769 | 0,031788802 | 0,541949268 |
| PSMC2 | 27 | 24 | -0,069158394 | 0,031503502 | 0,541949268 |
| DLG4 | 33 | 28 | 0,036291951 | 0,031706354 | 0,541949268 |
| NDUFA10 | 19 | 14 | 0,107932672 | 0,031351774 | 0,541949268 |
| ATP5F1E | 3 | 3 | 0,117570076 | 0,031740558 | 0,541949268 |
| RPL18 | 6 | 5 | -0,161603909 | 0,031984501 | 0,543736519 |
| SERPINA1C | 6 | 5 | -0,403186386 | 0,032212301 | 0,545873742 |
| PDHB | 26 | 20 | 0,079802 | 0,032292665 | 0,545873742 |
| SLC25A5 | 12 | 9 | 0,065766309 | 0,032431631 | 0,546678536 |
| RAPGEF4 | 24 | 14 | 0,056406813 | 0,032563229 | 0,547354958 |
| DGKI | 7 | 6 | -0,09584517 | 0,03318992 | 0,556326282 |
| CCDC9 | 2 | 2 | -0,20143221 | 0,033492275 | 0,559826182 |

| | | | | | |
|---|---|---|---|---|---|
| TBCA | 6 | 6 | -0,107944455 | 0,03370752 | 0,56185459 |
| ARF4 | 6 | 3 | -0,134118667 | 0,033861359 | 0,562617087 |
| KHSRP | 9 | 7 | -0,132359053 | 0,033941305 | 0,562617087 |
| GRM3 | 22 | 18 | -0,089823594 | 0,034553922 | 0,568051293 |
| SYT3 | 16 | 9 | 0,081134533 | 0,03454086 | 0,568051293 |
| IGSF11 | 3 | 1 | 0,333047542 | 0,034453434 | 0,568051293 |
| RYR1;RYR2 | 1 | 1 | 0,186208945 | 0,034664271 | 0,5683041 |
| EPB41L3 | 17 | 15 | -0,096301826 | 0,034850624 | 0,569798186 |
| EIF4G2 | 22 | 19 | -0,105867502 | 0,03521769 | 0,574230676 |
| HNRNPD | 13 | 11 | -0,158182928 | 0,036186837 | 0,578058677 |
| DGKH | 18 | 15 | -0,150755377 | 0,036225268 | 0,578058677 |
| EDC4 | 9 | 8 | -0,132538719 | 0,035688749 | 0,578058677 |
| KATNB1 | 10 | 8 | -0,105678697 | 0,036128641 | 0,578058677 |
| ACTN4 | 23 | 21 | -0,094223733 | 0,03590695 | 0,578058677 |
| SNX5 | 14 | 11 | 0,061589045 | 0,035608635 | 0,578058677 |
| GNAI1 | 9 | 7 | 0,071096693 | 0,035754304 | 0,578058677 |
| UQCRC1 | 24 | 22 | 0,086173975 | 0,036189302 | 0,578058677 |
| ARHGEF2 | 17 | 14 | 0,055494456 | 0,036617178 | 0,582258796 |
| EPHA4 | 26 | 20 | 0,062466 | 0,036683083 | 0,582258796 |
| EIF4G2 | 1 | 1 | -0,21497018 | 0,037850133 | 0,596423323 |
| SART3 | 17 | 14 | -0,110484883 | 0,037802603 | 0,596423323 |
| PUM1 | 6 | 6 | -0,101369206 | 0,037874476 | 0,596423323 |
| CYRIA | 15 | 13 | -0,070496294 | 0,038084975 | 0,596598136 |
| SCCPDH | 13 | 6 | 0,081000594 | 0,038033702 | 0,596598136 |
| WIPF2 | 7 | 7 | -0,13325196 | 0,038458941 | 0,599318494 |
| PCDH7 | 14 | 7 | -0,123316229 | 0,038371629 | 0,599318494 |
| CYCS | 14 | 14 | 0,090543033 | 0,038580984 | 0,599658726 |
| AFG1L | 8 | 1 | -0,241596964 | 0,038731955 | 0,600445651 |
| PLIN3 | 8 | 6 | -0,199684295 | 0,039152898 | 0,605402943 |
| G3BP1 | 8 | 5 | -0,088241183 | 0,039427268 | 0,608074148 |
| TAGLN2 | 12 | 8 | -0,121357819 | 0,039862751 | 0,613210034 |
| SLC7A11 | 4 | 3 | -0,276395333 | 0,04091385 | 0,623207463 |
| CFAP36 | 11 | 7 | -0,097884426 | 0,04075097 | 0,623207463 |

| | | | | | |
|---|---|---|---|---|---|
| CYC1 | 17 | 14 | 0,071109529 | 0,040929233 | 0,623207463 |
| HK2;HK1 | 3 | 3 | 0,111544284 | 0,040803722 | 0,623207463 |
| HIBADH | 11 | 8 | 0,088774289 | 0,041255908 | 0,626587184 |
| SERPINA1B;SERPINA1D | 6 | 6 | -0,374236346 | 0,041738837 | 0,626796449 |
| HNRNPAB | 8 | 7 | -0,140162882 | 0,041499626 | 0,626796449 |
| SGTA | 15 | 13 | -0,118459526 | 0,041668444 | 0,626796449 |
| DMTN | 24 | 19 | -0,100636074 | 0,041793413 | 0,626796449 |
| PCYOX1 | 14 | 10 | 0,063431262 | 0,041664008 | 0,626796449 |
| CDKN1B | 2 | 1 | -0,396636844 | 0,042500453 | 0,627150857 |
| ERLIN1 | 2 | 2 | -0,193353034 | 0,042779576 | 0,627150857 |
| NCBP1 | 3 | 2 | -0,141161551 | 0,042266847 | 0,627150857 |
| PALMD | 2 | 2 | -0,110763223 | 0,0428165 | 0,627150857 |
| EIF4H | 11 | 9 | -0,075166125 | 0,042098266 | 0,627150857 |
| SV2B | 20 | 16 | 0,066109383 | 0,042766325 | 0,627150857 |
| SLC17A7 | 14 | 12 | 0,08138747 | 0,042770539 | 0,627150857 |
| TOMM40L | 6 | 3 | 0,106010738 | 0,042820841 | 0,627150857 |
| PLA2G15 | 4 | 2 | 0,122256614 | 0,04286509 | 0,627150857 |
| MFF | 11 | 7 | 0,122713707 | 0,042647483 | 0,627150857 |
| EPM2AIP1 | 22 | 20 | -0,089850426 | 0,043096141 | 0,627463034 |
| LRRC47 | 17 | 16 | -0,058021452 | 0,043066663 | 0,627463034 |
| OOSP2 | 1 | 1 | -0,155830346 | 0,043633619 | 0,632914042 |
| GLMN | 3 | 3 | -0,145250541 | 0,043682069 | 0,632914042 |
| SEC23A;SEC23B | 3 | 3 | -0,145719013 | 0,044027146 | 0,636373046 |
| CUSTOS | 2 | 1 | -0,189255198 | 0,044367737 | 0,638212825 |
| RELCH | 1 | 1 | 0,150034893 | 0,044326125 | 0,638212825 |
| RELN | 4 | 3 | -0,11513711 | 0,044704495 | 0,638604339 |
| HNRNPA3 | 10 | 8 | -0,098770427 | 0,044821829 | 0,638604339 |
| GAMT | 5 | 5 | 0,103575036 | 0,044732293 | 0,638604339 |
| USP32 | 18 | 9 | 0,108284703 | 0,044646403 | 0,638604339 |
| ROBO1 | 14 | 8 | 0,113106983 | 0,045073157 | 0,639141634 |
| ATP2B1 | 6 | 3 | 0,138825824 | 0,044969274 | 0,639141634 |
| HTATSF1 | 5 | 4 | -0,107282491 | 0,045250433 | 0,639985169 |
| CKB | 30 | 27 | -0,055967117 | 0,045346543 | 0,639985169 |

| | | | | | |
|---|---|---|---|---|---|
| PALS1 | 2 | 1 | -0,270809053 | 0,045663281 | 0,642938998 |
| PHKA1 | 9 | 7 | -0,089319437 | 0,045896643 | 0,644707781 |
| P2RY12 | 5 | 3 | -0,134548846 | 0,046346522 | 0,649502544 |
| PHKB | 7 | 7 | -0,081900549 | 0,046507219 | 0,650231776 |
| RAD23B | 18 | 15 | -0,073256073 | 0,046674661 | 0,651051685 |
| GABRA2 | 5 | 2 | -0,186134145 | 0,047343198 | 0,653124507 |
| EPN2 | 9 | 8 | -0,113633748 | 0,047478135 | 0,653124507 |
| PAIP1 | 6 | 6 | -0,084071522 | 0,047422787 | 0,653124507 |
| WDR26 | 9 | 6 | -0,078306602 | 0,047408138 | 0,653124507 |
| VBP1 | 12 | 11 | -0,07224375 | 0,04746878 | 0,653124507 |
| PDK2 | 12 | 7 | 0,079318235 | 0,047184424 | 0,653124507 |
| AP3D1 | 31 | 17 | 0,207318721 | 0,047607725 | 0,653405109 |
| GGA3 | 8 | 7 | -0,121839059 | 0,047915252 | 0,656120975 |
| LIN7B;LIN7C;LIN7A | 3 | 3 | -0,191401033 | 0,049462296 | 0,674334804 |
| STX16 | 7 | 4 | -0,097794477 | 0,049470752 | 0,674334804 |
| SETD3 | 10 | 9 | -0,065097734 | 0,049844809 | 0,676352236 |
| DLGAP1 | 9 | 3 | 0,110780966 | 0,049759223 | 0,676352236 |
