## Supplementary material for "SFRP1 upregulation causes hippocampal synaptic dysfunction and memory impairment": Suppl Table 3b

| gene_symbols_or_id | unique_peptides | peptides_used_for_dea_contrast: Control_Syn vs SFRP1-TG_Syn | foldchange.log2_msqrob_contrast: Control_Syn vs SFRP1-TG_Syn | pvalue_msqrob_contrast: Control_Syn vs SFRP1-TG_Syn | qvalue_msqrob_contrast: Control_Syn vs SFRP1-TG_Syn |
|---|---|---|---|---|---|
| GPC1 | 11 | 11 | 0,148551357 | 6,6978E-07 | 0,004031403 |
| TJP2 | 18 | 17 | -0,175466514 | 2,04293E-06 | 0,004185319 |
| WDR7 | 66 | 62 | 0,099547848 | 2,08605E-06 | 0,004185319 |
| WIPF2 | 7 | 7 | -0,177535697 | 3,01369E-06 | 0,004534857 |
| ANKS1B | 15 | 14 | 0,141902832 | 7,79528E-06 | 0,009383962 |
| ZER1 | 10 | 7 | 0,210179954 | 1,04331E-05 | 0,010466098 |
| TIAM2 | 2 | 1 | -0,451637706 | 4,08371E-05 | 0,018126734 |
| ATP6V0A1 | 43 | 40 | 0,066896086 | 3,53926E-05 | 0,018126734 |
| NRXN3 | 26 | 23 | 0,087528165 | 3,43383E-05 | 0,018126734 |
| EPHA4 | 26 | 23 | 0,088206354 | 4,09726E-05 | 0,018126734 |
| SYT12 | 13 | 13 | 0,131213028 | 3,29517E-05 | 0,018126734 |
| RAB39B | 11 | 11 | 0,13983937 | 4,21622E-05 | 0,018126734 |
| HOMER3 | 17 | 17 | 0,19229051 | 3,23827E-05 | 0,018126734 |
| PPP4C | 2 | 1 | 0,396977275 | 3,14982E-05 | 0,018126734 |
| EFR3B | 24 | 24 | 0,081160144 | 5,43192E-05 | 0,020922245 |
| CACNA1A | 13 | 12 | 0,154500863 | 5,56165E-05 | 0,020922245 |
| PBXIP1 | 3 | 3 | -0,240637356 | 6,5247E-05 | 0,022604052 |
| TJP1 | 30 | 25 | -0,113082946 | 7,67601E-05 | 0,024316795 |
| CA4 | 7 | 7 | 0,176444462 | 9,74142E-05 | 0,027798204 |
| SLC16A1 | 5 | 4 | 0,212579828 | 9,42485E-05 | 0,027798204 |
| TUBAL3 | 5 | 1 | 0,388599445 | 0,000101605 | 0,027798204 |
| AP2A2 | 54 | 51 | 0,087764579 | 0,000115547 | 0,029907213 |
| NRP1 | 16 | 14 | 0,164135095 | 0,000119251 | 0,029907213 |
| SLC44A2 | 3 | 3 | 0,322216941 | 0,000126631 | 0,030487662 |
| SDC3 | 2 | 2 | -0,352999852 | 0,000132637 | 0,030705537 |
| NUCB1 | 11 | 11 | -0,223516622 | 0,000162684 | 0,036266375 |

| | | | | | |
|---|---|---|---|---|---|
| ROGDI | 17 | 16 | 0,101345749 | 0,000169379 | 0,036410538 |
| PIP5K1C | 25 | 25 | 0,100756627 | 0,000200147 | 0,041540876 |
| MACF1 | 72 | 61 | -0,090762029 | 0,000213133 | 0,042761526 |
| IRS2 | 2 | 1 | -0,544376929 | 0,000263767 | 0,051213246 |
| PPP2R5C | 9 | 7 | 0,131003029 | 0,000313078 | 0,057132078 |
| CADM3 | 18 | 17 | 0,151867118 | 0,000313235 | 0,057132078 |
| CSNK1D | 1 | 1 | -0,45041982 | 0,000352519 | 0,058910426 |
| NRP2 | 10 | 9 | 0,13187915 | 0,000362134 | 0,058910426 |
| CPNE4 | 18 | 14 | 0,189571172 | 0,000347773 | 0,058910426 |
| NUP205 | 2 | 2 | 0,33192573 | 0,000356363 | 0,058910426 |
| FAM81A | 9 | 8 | -0,147256199 | 0,000399603 | 0,061038985 |
| CPE | 22 | 19 | 0,079069254 | 0,000400065 | 0,061038985 |
| ARHGAP39 | 27 | 26 | 0,082808084 | 0,000405642 | 0,061038985 |
| NPTX1 | 14 | 12 | 0,136459354 | 0,000432171 | 0,063444844 |
| SLC17A7 | 14 | 12 | 0,155121857 | 0,000483063 | 0,069227525 |
| EPHA6 | 5 | 4 | 0,175618878 | 0,000498651 | 0,06979948 |
| RAB2A;RAB2B | 9 | 7 | 0,17723437 | 0,000543514 | 0,074350228 |
| RAB1A | 14 | 10 | 0,12776863 | 0,000556564 | 0,074443579 |
| TPRG1L | 10 | 9 | 0,122940778 | 0,000578267 | 0,074769406 |
| PFKL | 37 | 32 | 0,172992589 | 0,000594187 | 0,074769406 |
| GALE | 8 | 1 | 0,466605358 | 0,000596267 | 0,074769406 |
| FAAH | 14 | 12 | 0,120982155 | 0,000612016 | 0,075178095 |
| CPD | 11 | 10 | -0,118794755 | 0,000652442 | 0,078541026 |
| CACNA2D3 | 31 | 31 | 0,084492887 | 0,000685026 | 0,079396127 |
| SCAI | 22 | 20 | 0,086258831 | 0,000685928 | 0,079396127 |
| ARL8B | 7 | 7 | 0,202610955 | 0,00070107 | 0,079617744 |
| CYP2D22 | 6 | 5 | -0,253645001 | 0,000727087 | 0,081043263 |
| MYO16 | 1 | 1 | -0,485453611 | 0,000791913 | 0,084416567 |
| THA1 | 3 | 2 | -0,250185725 | 0,000780487 | 0,084416567 |
| NLN | 13 | 11 | -0,184893518 | 0,000799426 | 0,084416567 |
| RAB15 | 9 | 8 | 0,117757802 | 0,000820012 | 0,085097499 |
| CNKSR2 | 35 | 32 | 0,059896901 | 0,00087005 | 0,087848037 |
| L1CAM | 4 | 4 | 0,198483357 | 0,000875707 | 0,087848037 |

| | | | | | |
|---|---|---|---|---|---|
| SH3GL3 | 12 | 12 | 0,091880178 | 0,000926777 | 0,091447045 |
| EHD3;EHD1 | 11 | 10 | 0,112912649 | 0,000948642 | 0,092094779 |
| DOCK10 | 11 | 8 | -0,134147783 | 0,001005026 | 0,096019819 |
| AKAP7 | 3 | 3 | -0,321012854 | 0,001042766 | 0,0965601 |
| RAB8A | 7 | 6 | 0,1551853 | 0,00103774 | 0,0965601 |
| CLVS1 | 12 | 8 | 0,10506215 | 0,001063904 | 0,097024843 |
| MRPL24 | 2 | 2 | -0,269145505 | 0,001207511 | 0,105333497 |
| ACTR10 | 15 | 14 | 0,089224898 | 0,001185634 | 0,105333497 |
| DDX3X | 4 | 3 | 0,167638039 | 0,001205127 | 0,105333497 |
| TMEM181A | 1 | 1 | -0,478021602 | 0,001587819 | 0,131804812 |
| KCNMA1 | 26 | 23 | 0,085067447 | 0,001557417 | 0,131804812 |
| LRRTM1 | 9 | 9 | 0,126404765 | 0,001597493 | 0,131804812 |
| BRCC3 | 6 | 4 | 0,223262136 | 0,001598563 | 0,131804812 |
| EPB41L2 | 37 | 37 | -0,096481478 | 0,001693813 | 0,135934143 |
| PRICKLE2 | 16 | 16 | 0,078622104 | 0,001689241 | 0,135934143 |
| FIBCD1 | 4 | 4 | 0,158110684 | 0,00182241 | 0,144167982 |
| SYNPR | 8 | 8 | 0,189621446 | 0,001844315 | 0,144167982 |
| USP12 | 1 | 1 | 0,389096438 | 0,001918348 | 0,148032532 |
| ARHGEF2 | 12 | 10 | -0,125544399 | 0,002254113 | 0,171740617 |
| LIPT1 | 1 | 1 | -0,334986213 | 0,002332552 | 0,175174148 |
| CAMK2D | 14 | 12 | -0,114946637 | 0,002357386 | 0,175174148 |
| RPL10A | 14 | 7 | 0,197313958 | 0,002474831 | 0,181658628 |
| CCDC136 | 5 | 5 | -0,140568468 | 0,0026242 | 0,190301912 |
| ACSL3 | 14 | 12 | -0,091148433 | 0,002672161 | 0,191415856 |
| ARL4C | 1 | 1 | 0,333682855 | 0,002734967 | 0,191415856 |
| RIN1 | 1 | 1 | 0,492856226 | 0,002718317 | 0,191415856 |
| DGKB | 16 | 12 | 0,166668335 | 0,002774529 | 0,19195276 |
| FGFR1;FGFR3 | 1 | 1 | -0,269109754 | 0,002957634 | 0,200357663 |
| SYT3 | 16 | 15 | 0,089754706 | 0,003029166 | 0,200357663 |
| RPL21 | 6 | 4 | 0,281243316 | 0,002979441 | 0,200357663 |
| VKORC1L1 | 1 | 1 | 0,483648071 | 0,003001584 | 0,200357663 |
| RAPGEF4 | 24 | 19 | 0,072668954 | 0,003154252 | 0,206363527 |
| CCDC88A | 12 | 8 | -0,160377575 | 0,003338592 | 0,212994213 |

| | | | | | |
|---|---|---|---|---|---|
| HBB-B1 | 13 | 8 | 0,398107228 | 0,003356116 | 0,212994213 |
| PHYHD1 | 4 | 1 | 0,46968236 | 0,003361763 | 0,212994213 |
| ROBO1;ROBO2 | 2 | 1 | 0,296916556 | 0,003469859 | 0,215310107 |
| ITGAM;GM49368 | 1 | 1 | 0,464823259 | 0,003450173 | 0,215310107 |
| PSEN1 | 1 | 1 | 0,366079339 | 0,003524532 | 0,216471017 |
| RBX1 | 1 | 1 | 0,308528611 | 0,00364443 | 0,221573966 |
| SMPD3 | 11 | 11 | 0,09246862 | 0,0037327 | 0,22244677 |
| AP4S1 | 1 | 1 | 0,471933378 | 0,003732594 | 0,22244677 |
| SENP6 | 1 | 1 | -0,248210053 | 0,003985487 | 0,235182792 |
| SLC6A7 | 13 | 13 | 0,104603972 | 0,004049046 | 0,236613675 |
| TMEM30A | 11 | 9 | 0,120007998 | 0,004359508 | 0,252306526 |
| FAM117B | 2 | 1 | -0,429116831 | 0,004668689 | 0,257345818 |
| PITPNM2 | 21 | 20 | -0,093950486 | 0,004703113 | 0,257345818 |
| FMN2 | 19 | 17 | -0,07431638 | 0,004585585 | 0,257345818 |
| ARHGEF12 | 19 | 18 | -0,072706637 | 0,004669902 | 0,257345818 |
| PPP2R5A | 14 | 13 | 0,065087196 | 0,004593926 | 0,257345818 |
| AP1S1 | 7 | 4 | 0,156637505 | 0,004656507 | 0,257345818 |
| CCT3 | 40 | 34 | 0,064529663 | 0,004911859 | 0,266346671 |
| RALGAPA1 | 14 | 12 | -0,121320363 | 0,005075763 | 0,271785947 |
| KIT | 17 | 14 | 0,127687025 | 0,005102477 | 0,271785947 |
| DPY19L4 | 1 | 1 | -0,298561263 | 0,005319626 | 0,280866935 |
| GALK1 | 9 | 3 | -0,135144611 | 0,005557691 | 0,290884733 |
| SERINC5 | 1 | 1 | -0,283691903 | 0,005762543 | 0,297285262 |
| CSNK1D | 2 | 1 | -0,215027247 | 0,005811779 | 0,297285262 |
| SCN3A;SCN2A;SCN9A | 4 | 4 | 0,147757801 | 0,005828154 | 0,297285262 |
| ATP6V0D1 | 19 | 19 | 0,065102564 | 0,005989134 | 0,300404973 |
| LAMTOR2 | 3 | 3 | 0,174480098 | 0,005987642 | 0,300404973 |
| SRCIN1 | 10 | 9 | -0,108267417 | 0,006211028 | 0,305660639 |
| ADD3 | 25 | 22 | -0,055721797 | 0,006246263 | 0,305660639 |
| SBF1 | 58 | 50 | 0,043100116 | 0,006163736 | 0,305660639 |
| DNM1 | 55 | 54 | 0,084412001 | 0,006378379 | 0,309608568 |
| HPCA | 7 | 7 | 0,216278646 | 0,006650317 | 0,31605311 |
| SRP68 | 7 | 4 | 0,22707341 | 0,006691637 | 0,31605311 |

| | | | | | |
|---|---|---|---|---|---|
| ARHGEF9;SPATA13;ARHGE | 1 | 1 | 0,408595422 | 0,006649569 | 0,31605311 |
| GPR155 | 2 | 1 | 0,62566653 | 0,006721183 | 0,31605311 |
| FAM120C | 10 | 6 | -0,116854016 | 0,006817548 | 0,318099411 |
| HIP1 | 7 | 7 | -0,144378698 | 0,006931624 | 0,319989853 |
| RAB10 | 11 | 11 | 0,09160893 | 0,006964391 | 0,319989853 |
| CNNM4 | 3 | 2 | 0,221613689 | 0,007117737 | 0,324558027 |
| TRMU | 4 | 4 | -0,18497737 | 0,007337457 | 0,328477942 |
| EIF5A | 13 | 9 | 0,180218967 | 0,007367424 | 0,328477942 |
| CP | 9 | 1 | 0,576458801 | 0,007340803 | 0,328477942 |
| GIT1 | 35 | 32 | -0,048463494 | 0,007482739 | 0,331166215 |
| CDV3 | 6 | 4 | -0,145619909 | 0,007595025 | 0,333682148 |
| KCNJ10 | 5 | 4 | 0,125687011 | 0,007753293 | 0,338167181 |
| MRPS14 | 1 | 1 | -0,254276626 | 0,008141681 | 0,352552379 |
| NQO1 | 8 | 3 | -0,148015011 | 0,008389456 | 0,353119832 |
| RAB9A | 6 | 5 | 0,127080037 | 0,008295723 | 0,353119832 |
| VPS25 | 5 | 5 | 0,135778788 | 0,008379405 | 0,353119832 |
| RAB27A | 2 | 2 | 0,13695741 | 0,008334123 | 0,353119832 |
| SPTAN1 | 2 | 2 | -0,233736228 | 0,008592957 | 0,354943756 |
| EEF1G | 20 | 16 | 0,077101222 | 0,008609701 | 0,354943756 |
| SV2B | 20 | 19 | 0,104775162 | 0,008538899 | 0,354943756 |
| CCDC9 | 2 | 1 | 0,433864844 | 0,008706465 | 0,356491231 |
| PLXNB1 | 28 | 25 | -0,049706031 | 0,008830153 | 0,357027008 |
| PACSIN1 | 43 | 33 | 0,075704864 | 0,008838183 | 0,357027008 |
| KLC2 | 15 | 14 | -0,060805337 | 0,009133775 | 0,364519664 |
| GPD1 | 17 | 14 | 0,07782166 | 0,009144786 | 0,364519664 |
| SLC25A18 | 11 | 11 | -0,23923433 | 0,009359477 | 0,370622988 |
| RAB14 | 20 | 20 | 0,081036233 | 0,009588077 | 0,377193703 |
| LRRC8D | 12 | 8 | 0,10867557 | 0,009746544 | 0,380937987 |
| NEDD4L | 18 | 17 | 0,058861373 | 0,010004983 | 0,382910881 |
| OLA1 | 25 | 14 | 0,062870554 | 0,009883719 | 0,382910881 |
| AGAP3 | 11 | 11 | 0,084667528 | 0,01005149 | 0,382910881 |
| GABRA5 | 5 | 5 | 0,152296778 | 0,009972571 | 0,382910881 |
| SMN1 | 1 | 1 | -0,526556955 | 0,010179615 | 0,382944381 |

| | | | | | |
|---|---|---|---|---|---|
| HPCAL1;NCALD | 6 | 6 | 0,085711417 | 0,010168224 | 0,382944381 |
| CUL9 | 2 | 1 | -0,267553585 | 0,010256676 | 0,3834468 |
| KTN1 | 23 | 17 | -0,098301898 | 0,010396554 | 0,386276893 |
| CPOX | 9 | 9 | -0,134380583 | 0,010619397 | 0,389744836 |
| RAB4B | 9 | 8 | 0,087601375 | 0,010579877 | 0,389744836 |
| NPTN | 2 | 2 | 0,229020652 | 0,010698636 | 0,390273259 |
| AK9 | 1 | 1 | -0,183081646 | 0,010939073 | 0,394265167 |
| SHISA4 | 1 | 1 | 0,344943363 | 0,010892722 | 0,394265167 |
| PCDHGC5 | 8 | 5 | 0,101204498 | 0,011008825 | 0,394417352 |
| SQOR | 8 | 6 | -0,202910442 | 0,01124351 | 0,400441918 |
| ANK3;ANK1 | 1 | 1 | -0,43115677 | 0,012374471 | 0,417705515 |
| WFS1 | 19 | 16 | -0,144529144 | 0,012166773 | 0,417705515 |
| MYO6 | 29 | 26 | -0,060222929 | 0,012127355 | 0,417705515 |
| CSNK2A2 | 18 | 17 | 0,064508186 | 0,01195335 | 0,417705515 |
| SEPTIN8 | 30 | 29 | 0,075415224 | 0,012394989 | 0,417705515 |
| RGS14 | 20 | 17 | 0,113294058 | 0,012491609 | 0,417705515 |
| SLC39A12 | 6 | 6 | 0,114315258 | 0,012140058 | 0,417705515 |
| DBH | 1 | 1 | 0,268878954 | 0,012239747 | 0,417705515 |
| KCTD6 | 3 | 2 | 0,308007349 | 0,011804145 | 0,417705515 |
| LSM14A | 3 | 1 | 0,327303175 | 0,012445935 | 0,417705515 |
| H1-4;H1-3 | 3 | 1 | 0,981025677 | 0,01200912 | 0,417705515 |
| ABHD3 | 4 | 3 | -0,22132543 | 0,012763844 | 0,420151678 |
| VCAN | 1 | 1 | -0,208155703 | 0,012766275 | 0,420151678 |
| NTNG1 | 5 | 5 | -0,099703354 | 0,012774175 | 0,420151678 |
| DNAJC13 | 29 | 25 | 0,057836899 | 0,012933676 | 0,420798883 |
| HBB-BS | 4 | 2 | 0,350741742 | 0,012896075 | 0,420798883 |
| MPST | 15 | 14 | -0,183135292 | 0,013092255 | 0,423668176 |
| NLGN4L | 1 | 1 | -0,311669588 | 0,013168968 | 0,423871761 |
| PSD3 | 34 | 30 | 0,069029626 | 0,013522863 | 0,430656694 |
| VWA5A | 18 | 12 | 0,139362493 | 0,013456182 | 0,430656694 |
| WDR20 | 4 | 2 | -0,171536031 | 0,013786178 | 0,432182326 |
| CYFIP2 | 70 | 59 | 0,072830448 | 0,013724978 | 0,432182326 |
| RPL37A | 3 | 1 | 0,31661003 | 0,013758304 | 0,432182326 |

| | | | | | |
|---|---|---|---|---|---|
| AKR1C13 | 1 | 1 | 0,188649616 | 0,013979823 | 0,435982152 |
| RHOT1 | 1 | 1 | -0,20234925 | 0,014199023 | 0,440535665 |
| NANP | 6 | 1 | -0,414721095 | 0,015208139 | 0,441363712 |
| PPM1K | 1 | 1 | -0,288604024 | 0,015216078 | 0,441363712 |
| UNC5CL | 1 | 1 | -0,271970568 | 0,014907659 | 0,441363712 |
| TXNDC12 | 2 | 2 | -0,135381277 | 0,015325638 | 0,441363712 |
| DMD | 6 | 4 | -0,13019078 | 0,01452115 | 0,441363712 |
| EPB41L1 | 62 | 57 | -0,038201308 | 0,015117552 | 0,441363712 |
| NAPG | 24 | 21 | 0,0560102 | 0,014785996 | 0,441363712 |
| RIMS1 | 12 | 11 | 0,058807606 | 0,015256713 | 0,441363712 |
| VPS26B | 7 | 7 | 0,094302721 | 0,014493433 | 0,441363712 |
| HAPLN4 | 10 | 8 | 0,09448473 | 0,015170136 | 0,441363712 |
| AP3S1 | 7 | 7 | 0,141161787 | 0,015143943 | 0,441363712 |
| TRAPPC2 | 2 | 2 | 0,157090156 | 0,015035607 | 0,441363712 |
| RPL22 | 4 | 3 | 0,172939883 | 0,014490066 | 0,441363712 |
| C9ORF72 | 3 | 3 | 0,200650546 | 0,014843733 | 0,441363712 |
| RPS13 | 10 | 8 | 0,226549553 | 0,014712268 | 0,441363712 |
| QKI | 4 | 4 | -0,114833598 | 0,015411171 | 0,44171351 |
| TOMM34 | 1 | 1 | 0,258920604 | 0,015528645 | 0,442971151 |
| MTERF2 | 3 | 2 | -0,175412777 | 0,015768799 | 0,447700005 |
| USO1 | 36 | 19 | 0,052210321 | 0,016254125 | 0,457166248 |
| GRIA2 | 36 | 35 | 0,093047169 | 0,016199356 | 0,457166248 |
| AKAP5 | 20 | 16 | -0,11687739 | 0,016435367 | 0,460113836 |
| AASS | 3 | 2 | -0,356201681 | 0,016594483 | 0,462417564 |
| USP46;USP12 | 3 | 3 | 0,13201852 | 0,016795141 | 0,465852325 |
| MAPK8IP1 | 2 | 2 | -0,193554146 | 0,01688137 | 0,466096188 |
| SPATA45 | 1 | 1 | -0,181820833 | 0,01697653 | 0,466583255 |
| TSHZ1 | 1 | 1 | -0,320696208 | 0,018128056 | 0,476474976 |
| AMIGO1 | 3 | 1 | -0,250199804 | 0,017734891 | 0,476474976 |
| SHISA9 | 2 | 2 | -0,151088129 | 0,017792071 | 0,476474976 |
| MAP1B | 109 | 94 | -0,091288929 | 0,018033519 | 0,476474976 |
| NHSL2 | 10 | 7 | -0,089765332 | 0,018064196 | 0,476474976 |
| FLOT1 | 29 | 26 | 0,055234319 | 0,018024616 | 0,476474976 |

| | | | | | |
|---|---|---|---|---|---|
| RRAGC | 16 | 8 | 0,096479054 | 0,017702887 | 0,476474976 |
| DIRAS2 | 12 | 8 | 0,147834366 | 0,018005471 | 0,476474976 |
| MAPK10;MAPK9 | 2 | 1 | 0,17727296 | 0,017715164 | 0,476474976 |
| ITGA6 | 5 | 3 | 0,22626851 | 0,017492098 | 0,476474976 |
| CALCOCO1 | 14 | 10 | -0,069426153 | 0,018220001 | 0,476809517 |
| SLC8A1 | 30 | 21 | 0,076729835 | 0,018370736 | 0,47867299 |
| EIF3F | 15 | 8 | 0,092646986 | 0,018603395 | 0,482645838 |
| LIAS | 4 | 4 | -0,131805923 | 0,018871832 | 0,483359825 |
| MAPK1 | 35 | 29 | 0,114086296 | 0,018865179 | 0,483359825 |
| HNRNPK | 22 | 9 | 0,172445803 | 0,018845957 | 0,483359825 |
| UBAP2L | 18 | 6 | 0,128093542 | 0,019183566 | 0,489262215 |
| ATP2B3 | 5 | 5 | 0,094774052 | 0,019380513 | 0,492199614 |
| TLCD4 | 2 | 2 | 0,145307943 | 0,019484966 | 0,492773155 |
| KYAT3 | 12 | 10 | -0,232690524 | 0,019690388 | 0,494414838 |
| PALS1 | 2 | 1 | 0,201067857 | 0,019714165 | 0,494414838 |
| ADCY9 | 31 | 29 | 0,076296074 | 0,019907451 | 0,496049319 |
| RPL18A | 6 | 3 | 0,448959613 | 0,019944166 | 0,496049319 |
| SPTBN4 | 44 | 40 | -0,11184401 | 0,021121028 | 0,519637719 |
| SPECC1 | 15 | 12 | -0,079770816 | 0,021000204 | 0,519637719 |
| PCDH7 | 14 | 13 | -0,074508528 | 0,02121065 | 0,519637719 |
| TNIK;MAP4K4;MINK1 | 9 | 9 | 0,078324758 | 0,02130407 | 0,519637719 |
| FARSB | 21 | 9 | 0,171412948 | 0,021324226 | 0,519637719 |
| CALU | 10 | 7 | -0,088361559 | 0,021729491 | 0,526947462 |
| SPIRE1 | 5 | 5 | 0,101636175 | 0,021799289 | 0,526947462 |
| DST | 41 | 36 | -0,086666701 | 0,021923366 | 0,527826969 |
| PCSK1N | 6 | 5 | -0,153858294 | 0,022322581 | 0,52963667 |
| CCT4 | 38 | 28 | 0,070356767 | 0,022202791 | 0,52963667 |
| YWHAB | 10 | 8 | 0,071408532 | 0,022114704 | 0,52963667 |
| PLXNA1 | 40 | 39 | 0,081074817 | 0,022350509 | 0,52963667 |
| MYH14 | 42 | 35 | -0,085681291 | 0,022787756 | 0,53162599 |
| STMN1;STMN2 | 4 | 4 | -0,077457102 | 0,022525629 | 0,53162599 |
| HMOX2 | 16 | 15 | -0,048958163 | 0,02274573 | 0,53162599 |
| CA11 | 1 | 1 | 0,192957749 | 0,022639674 | 0,53162599 |

| | | | | | |
|---|---|---|---|---|---|
| GPX1 | 13 | 12 | 0,106099976 | 0,022907935 | 0,532366253 |
| TPM1 | 1 | 1 | -0,203837939 | 0,023214512 | 0,537415947 |
| PRKAB2 | 7 | 7 | 0,115746863 | 0,023417002 | 0,540026564 |
| YWHAQ | 18 | 14 | 0,08338572 | 0,0235826 | 0,541769735 |
| TMEM70 | 2 | 2 | -0,192953003 | 0,023996214 | 0,547095492 |
| 2900026A02RIK | 11 | 8 | 0,110816133 | 0,023963215 | 0,547095492 |
| POMT1 | 1 | 1 | -0,251931896 | 0,024296924 | 0,547892101 |
| MIGA1 | 2 | 2 | -0,203428673 | 0,024304235 | 0,547892101 |
| FAM241B | 2 | 2 | -0,111189754 | 0,024216681 | 0,547892101 |
| TRPV2 | 6 | 5 | 0,077046659 | 0,024527824 | 0,548821456 |
| RPS23 | 5 | 3 | 0,29415059 | 0,024527201 | 0,548821456 |
| CSDC2 | 2 | 1 | -0,204091327 | 0,024712127 | 0,550897386 |
| FRYL | 3 | 2 | -0,20327304 | 0,024955566 | 0,552680235 |
| SPARCL1 | 15 | 11 | -0,083963553 | 0,025113615 | 0,552680235 |
| OSBP2 | 4 | 3 | 0,12233909 | 0,025251215 | 0,552680235 |
| PTP4A1 | 3 | 3 | 0,143184246 | 0,025213578 | 0,552680235 |
| CSNK1G1;CSNK1G3 | 1 | 1 | 0,18061595 | 0,025078379 | 0,552680235 |
| TGOLN1;TGOLN2 | 1 | 1 | -0,346350518 | 0,025753002 | 0,555475167 |
| ADI1 | 3 | 1 | -0,214615622 | 0,025818291 | 0,555475167 |
| AP2B1 | 35 | 32 | 0,058081263 | 0,025683043 | 0,555475167 |
| ATP8A2 | 8 | 5 | 0,09342174 | 0,025840347 | 0,555475167 |
| RPL10A | 1 | 1 | 0,224356021 | 0,025492736 | 0,555475167 |
| GPT | 7 | 2 | -0,170063869 | 0,026498719 | 0,565587917 |
| CNBP | 1 | 1 | 0,331122114 | 0,026424791 | 0,565587917 |
| SH2D3C | 3 | 1 | -0,221929904 | 0,026901791 | 0,567717211 |
| PLCD1 | 15 | 10 | 0,075981123 | 0,026948365 | 0,567717211 |
| GMPPB | 8 | 7 | 0,11192674 | 0,026830239 | 0,567717211 |
| ARF6 | 8 | 7 | 0,200903637 | 0,026975764 | 0,567717211 |
| PRPSAP1 | 11 | 7 | 0,079212447 | 0,027418783 | 0,575030155 |
| GFOD1 | 9 | 7 | -0,054800212 | 0,027553719 | 0,575853594 |
| PARD3B | 1 | 1 | -0,195456134 | 0,027757677 | 0,578108845 |
| CELSR2 | 3 | 2 | -0,21586216 | 0,028064304 | 0,580477831 |
| MIPEP | 6 | 6 | -0,141219225 | 0,028019455 | 0,580477831 |

| | | | | | |
|---|---|---|---|---|---|
| MAP7 | 1 | 1 | -0,23334759 | 0,029109311 | 0,581129871 |
| COPA | 24 | 21 | -0,127502683 | 0,029067598 | 0,581129871 |
| EML4 | 9 | 3 | -0,120833093 | 0,029173094 | 0,581129871 |
| MYO18A | 70 | 66 | -0,046659319 | 0,028419732 | 0,581129871 |
| DNAJC6 | 24 | 23 | 0,038618267 | 0,028434199 | 0,581129871 |
| PDPK1 | 17 | 16 | 0,067218916 | 0,028596362 | 0,581129871 |
| INPP4A | 9 | 9 | 0,072687302 | 0,028683018 | 0,581129871 |
| RAB7A | 21 | 19 | 0,07542358 | 0,029201051 | 0,581129871 |
| ACOX1 | 15 | 12 | 0,081587609 | 0,029091799 | 0,581129871 |
| NDRG3 | 14 | 11 | 0,085485767 | 0,029615657 | 0,581129871 |
| CAPZA2 | 14 | 14 | 0,087922735 | 0,029737131 | 0,581129871 |
| LANCL2 | 18 | 18 | 0,090642182 | 0,028623713 | 0,581129871 |
| PPM1E | 21 | 11 | 0,100624049 | 0,028264761 | 0,581129871 |
| DCTN6 | 6 | 5 | 0,106479956 | 0,029737166 | 0,581129871 |
| PRKAG1 | 6 | 5 | 0,114350195 | 0,029283404 | 0,581129871 |
| SRR | 12 | 8 | 0,130755879 | 0,029628534 | 0,581129871 |
| RAB2B | 3 | 3 | 0,138082491 | 0,02952781 | 0,581129871 |
| PRR36 | 14 | 12 | -0,14537763 | 0,030171979 | 0,582318015 |
| PALM | 4 | 4 | -0,123629549 | 0,030087236 | 0,582318015 |
| GSTM1;GSTM2;GSTM4 | 3 | 2 | -0,114536235 | 0,029995663 | 0,582318015 |
| RAB35 | 14 | 14 | 0,056299655 | 0,030184951 | 0,582318015 |
| SLIT1;SLIT2 | 1 | 1 | -0,214332088 | 0,030373632 | 0,583184168 |
| FLAD1 | 11 | 9 | -0,069851216 | 0,030676156 | 0,583184168 |
| PRKAR1A | 17 | 15 | -0,049079717 | 0,030648449 | 0,583184168 |
| SUGT1 | 17 | 12 | 0,066748684 | 0,030916431 | 0,583184168 |
| GSK3A | 9 | 9 | 0,074863034 | 0,030930416 | 0,583184168 |
| ALDH1A1 | 19 | 18 | 0,081974507 | 0,031004973 | 0,583184168 |
| ITM2C | 5 | 5 | 0,105910347 | 0,030862511 | 0,583184168 |
| HBA-A1 | 9 | 6 | 0,344475571 | 0,030972738 | 0,583184168 |
| CAMK2B | 19 | 17 | 0,056655193 | 0,031416186 | 0,589077958 |
| TTC9 | 1 | 1 | -0,212565839 | 0,032360746 | 0,602435449 |
| IMMT | 2 | 2 | -0,162641789 | 0,032356089 | 0,602435449 |
| SPOCK2 | 2 | 2 | 0,153099116 | 0,032428823 | 0,602435449 |

| | | | | | |
|---|---|---|---|---|---|
| MRTFA | 2 | 2 | -0,311664313 | 0,032850496 | 0,60612302 |
| MADD | 47 | 43 | 0,04322689 | 0,032826742 | 0,60612302 |
| ACAP2 | 13 | 13 | 0,085175138 | 0,032929428 | 0,60612302 |
| EPHB2 | 10 | 6 | 0,086815312 | 0,033068656 | 0,606830002 |
| C1QL3 | 2 | 2 | -0,16216255 | 0,033269865 | 0,608053456 |
| PCDH1 | 23 | 22 | 0,068243163 | 0,033337372 | 0,608053456 |
| GJA1 | 19 | 18 | -0,135705557 | 0,033810146 | 0,612961647 |
| RAB1A;RAB1B | 5 | 5 | 0,076139813 | 0,033763152 | 0,612961647 |
| GABRB3 | 5 | 5 | 0,086223987 | 0,034108396 | 0,614900736 |
| TUBA8 | 10 | 8 | 0,141767839 | 0,034121423 | 0,614900736 |
| ANP32A | 16 | 4 | 0,139692878 | 0,034462865 | 0,619199954 |
| CORO1A | 22 | 20 | 0,057429364 | 0,034815387 | 0,621821405 |
| CD47 | 5 | 5 | 0,122182039 | 0,034795966 | 0,621821405 |
| ATP6V0A2;ATP6V0A4;ATP | 1 | 1 | 0,177256351 | 0,035039923 | 0,623980166 |
| RAF1 | 2 | 1 | 0,256798101 | 0,035225821 | 0,625440173 |
| AIFM3 | 10 | 7 | -0,090338711 | 0,035539396 | 0,627306822 |
| UBXN6 | 19 | 16 | 0,054467689 | 0,035441275 | 0,627306822 |
| MRPS31 | 3 | 2 | -0,205113579 | 0,035986638 | 0,62783644 |
| PNPO | 6 | 4 | -0,105982941 | 0,03593407 | 0,62783644 |
| AP2S1 | 11 | 10 | 0,101915394 | 0,035826142 | 0,62783644 |
| CETN2 | 2 | 1 | 0,248427475 | 0,035863896 | 0,62783644 |
| EEF1B | 10 | 7 | 0,113800986 | 0,036215599 | 0,630004886 |
| ACTR3B | 15 | 12 | 0,063278485 | 0,036752761 | 0,633343209 |
| SLC6A17 | 20 | 20 | 0,076070557 | 0,0366711 | 0,633343209 |
| ARL8A;ARL8B | 6 | 6 | 0,192637409 | 0,036701594 | 0,633343209 |
| PURG | 7 | 1 | 0,265571527 | 0,036828397 | 0,633343209 |
| DDX1 | 15 | 13 | 0,063221862 | 0,036953818 | 0,633689539 |
| TRAPPC8 | 12 | 10 | 0,067466845 | 0,037243569 | 0,635096912 |
| GATD1 | 5 | 5 | 0,128351117 | 0,03724692 | 0,635096912 |
| ARFGAP2 | 8 | 6 | 0,10223448 | 0,037362576 | 0,635269333 |
| TSPAN15 | 1 | 1 | -0,267860315 | 0,037606573 | 0,63645572 |
| CMPK2 | 14 | 13 | -0,077180326 | 0,037995 | 0,63645572 |
| SRGAP3 | 43 | 39 | 0,03642608 | 0,037946441 | 0,63645572 |

| | | | | | |
|---|---|---|---|---|---|
| ATP9A | 11 | 10 | 0,075457665 | 0,038033898 | 0,63645572 |
| SYP | 9 | 8 | 0,110823455 | 0,037645116 | 0,63645572 |
| CYTH3 | 3 | 3 | 0,122165732 | 0,038066798 | 0,63645572 |
| HPCAL4 | 19 | 12 | 0,1428039 | 0,038226137 | 0,637349353 |
| ALB | 40 | 11 | 0,322022778 | 0,038413654 | 0,638706579 |
| LIMCH1 | 8 | 7 | -0,117035942 | 0,038828299 | 0,643822409 |
| CNIH2 | 2 | 2 | 0,125536828 | 0,038951091 | 0,644084112 |
| ACADS | 7 | 7 | -0,178345326 | 0,039807649 | 0,650171394 |
| MRPL9 | 3 | 3 | -0,164509804 | 0,039803133 | 0,650171394 |
| CASKIN1 | 52 | 51 | -0,077581746 | 0,03985932 | 0,650171394 |
| CYFIP1 | 28 | 26 | 0,065983808 | 0,039696568 | 0,650171394 |
| SLC25A17 | 1 | 1 | 0,152702008 | 0,039484334 | 0,650171394 |
| YWHAH | 18 | 17 | 0,063844855 | 0,039978868 | 0,650358942 |
| TSC1 | 17 | 14 | -0,076863289 | 0,04039611 | 0,652706354 |
| EIPR1 | 7 | 5 | 0,102567784 | 0,040448491 | 0,652706354 |
| BACE1 | 1 | 1 | 0,187654691 | 0,040432784 | 0,652706354 |
| GM14151 | 1 | 1 | -0,166926254 | 0,040818535 | 0,653865129 |
| CDS2 | 7 | 7 | -0,076535935 | 0,040828484 | 0,653865129 |
| HSPA8 | 39 | 36 | 0,057545919 | 0,040888029 | 0,653865129 |
| DLGAP1 | 9 | 8 | 0,070978127 | 0,040954835 | 0,653865129 |
| MACO1 | 2 | 1 | -0,235230787 | 0,041070093 | 0,653970608 |
| PLPPR2 | 8 | 8 | 0,095810604 | 0,041236159 | 0,654882429 |
| MANF | 3 | 3 | 0,212249893 | 0,042102502 | 0,666881475 |
| CLMN | 14 | 12 | -0,089561988 | 0,043029511 | 0,676864779 |
| VAC14 | 16 | 13 | -0,045642596 | 0,043082861 | 0,676864779 |
| CLDND1 | 4 | 4 | 0,098573256 | 0,043182601 | 0,676864779 |
| ANAPC1 | 4 | 1 | 0,278095718 | 0,0429263 | 0,676864779 |
| CDC42EP4 | 10 | 10 | -0,087402514 | 0,043627314 | 0,678534376 |
| STXBP5L | 6 | 5 | 0,073189199 | 0,043547006 | 0,678534376 |
| PRKAA1 | 4 | 4 | 0,110849291 | 0,043589911 | 0,678534376 |
| ZFPL1 | 1 | 1 | -0,249878425 | 0,043859874 | 0,67967944 |
| RPLP0 | 13 | 9 | 0,099540551 | 0,044091057 | 0,67967944 |
| MEAK7 | 2 | 1 | 0,181403682 | 0,044152627 | 0,67967944 |

| | | | | | |
|---|---|---|---|---|---|
| RPS8 | 5 | 3 | 0,393697088 | 0,043968002 | 0,67967944 |
| SLC2A8 | 1 | 1 | -0,219382933 | 0,044356583 | 0,681077223 |
| ERCC5 | 1 | 1 | -0,33873921 | 0,045070312 | 0,683558101 |
| ARHGAP32 | 12 | 12 | -0,050873906 | 0,044856791 | 0,683558101 |
| ACTR1A | 10 | 8 | 0,070764569 | 0,045085989 | 0,683558101 |
| RAC3 | 2 | 2 | 0,107018292 | 0,044792651 | 0,683558101 |
| GPR37L1 | 3 | 3 | 0,16608837 | 0,04502276 | 0,683558101 |
| RPL10 | 9 | 6 | 0,257154067 | 0,045623542 | 0,689970094 |
| PRKAG2 | 16 | 14 | 0,06686497 | 0,046022486 | 0,692612708 |
| PAN2 | 1 | 1 | 0,161870112 | 0,046028424 | 0,692612708 |
| MIA3 | 9 | 8 | -0,086444886 | 0,046360252 | 0,69413521 |
| KIAA1549 | 17 | 15 | 0,06157095 | 0,046304592 | 0,69413521 |
| ZHX2 | 1 | 1 | -0,194487342 | 0,046999366 | 0,694294662 |
| MARCKSL1 | 2 | 2 | -0,152113412 | 0,047080399 | 0,694294662 |
| TIMM17B | 2 | 2 | -0,126702 | 0,046737434 | 0,694294662 |
| LRPAP1 | 17 | 16 | -0,074801953 | 0,046997773 | 0,694294662 |
| RRBP1 | 20 | 20 | -0,045933993 | 0,047098098 | 0,694294662 |
| PDCD6IP | 37 | 31 | -0,028535902 | 0,047026179 | 0,694294662 |
| SEC11A | 3 | 3 | 0,124455523 | 0,047178355 | 0,694294662 |
| NHSL1 | 1 | 1 | -0,194329634 | 0,047629288 | 0,694535974 |
| USP30 | 2 | 1 | -0,17048003 | 0,047488001 | 0,694535974 |
| STRN4 | 16 | 14 | -0,055014128 | 0,047373389 | 0,694535974 |
| VPS53 | 17 | 15 | 0,048925828 | 0,047656315 | 0,694535974 |
| RASAL1 | 31 | 28 | 0,081963038 | 0,048598844 | 0,706561456 |
| RPLP1 | 2 | 1 | 0,197413144 | 0,049090976 | 0,711996593 |
| DDRGK1 | 3 | 2 | -0,177887003 | 0,049332093 | 0,713242347 |
| BRINP1 | 15 | 14 | 0,057334764 | 0,049413866 | 0,713242347 |
| SUSD2 | 2 | 2 | 0,116722828 | 0,04969293 | 0,715554412 |
| SKP1 | 14 | 11 | 0,063001023 | 0,04997466 | 0,717893739 |
